# Integrating NMR and Simulations Reveals Motions in the UUCG Tetraloop

**DOI:** 10.1101/690412

**Authors:** Sandro Bottaro, Parker J. Nichols, Beat Vögeli, Michele Parrinello, Kresten Lindorff-Larsen

## Abstract

We provide an atomic-level description of the structure and dynamics of the UUCG RNA stem-loop by combining molecular dynamics simulations with experimental data. The integration of simulations with exact nuclear Overhauser enhancements data allowed us to characterize two distinct states of this molecule. The most stable conformation corresponds to the consensus three-dimensional structure. The second state is characterized by the absence of the peculiar non-Watson-Crick interactions in the loop region. By using machine learning techniques we identify a set of experimental measurements that are most sensitive to the presence of non-native states. We find that although our MD ensemble, as well as the consensus UUCG tetraloop structures, are in good agreement with experiments, there are remaining discrepancies. Together, our results show that i) the structural interpretation of experimental data for dynamic RNAs is highly complex, even for a simple model system such as the UUCG tetraloop ii) the MD simulation overstabilize a non-native loop conformation, and iii) eNOE data support its presence with a population of ≈10%.

## INTRODUCTION

RNA loops are structural elements that cap A-form double helices, and as such are fundamental structural units in RNA molecules. The great majority of known RNA loops contain four nucleotides *Wolters* (*1992*), and these so-called tetraloops are one of the most common and well-studied three-dimensional RNA motifs. The great majority of known RNA tetraloops have the sequence GNRA or UNCG, where N is any nucleotide and R is guanine or adenine *Bottaro and Lindorff-Larsen* (*2017*). Their small size, together with their biological relevance, has made these systems primary targets for nuclear magnetic resonance (NMR) spectroscopy, X-ray-crystallography, and atomistic molecular dynamics (MD) simulation studies *Cheong et al.* (*1990*); *Woese et al.* (*1990*); *Ferner et al.* (*2008*).

The UUCG tetraloop has been long known to be highly stable, and both crystallographic and NMR studies suggest that this tetraloop adopts a well-defined three dimensional structure including a characteristic trans-Sugar-Watson (tSW) interaction between U6 and G9 *Ennifar et al.* (*2000*); *Nozinovic et al.* (*2010*) (Fig. 1). Experimentally, the UUCG tetraloop is used to stabilize the secondary structure of larger RNA molecules without interacting with other RNAs or proteins *Hall* (*2015*).

**Figure 1.**
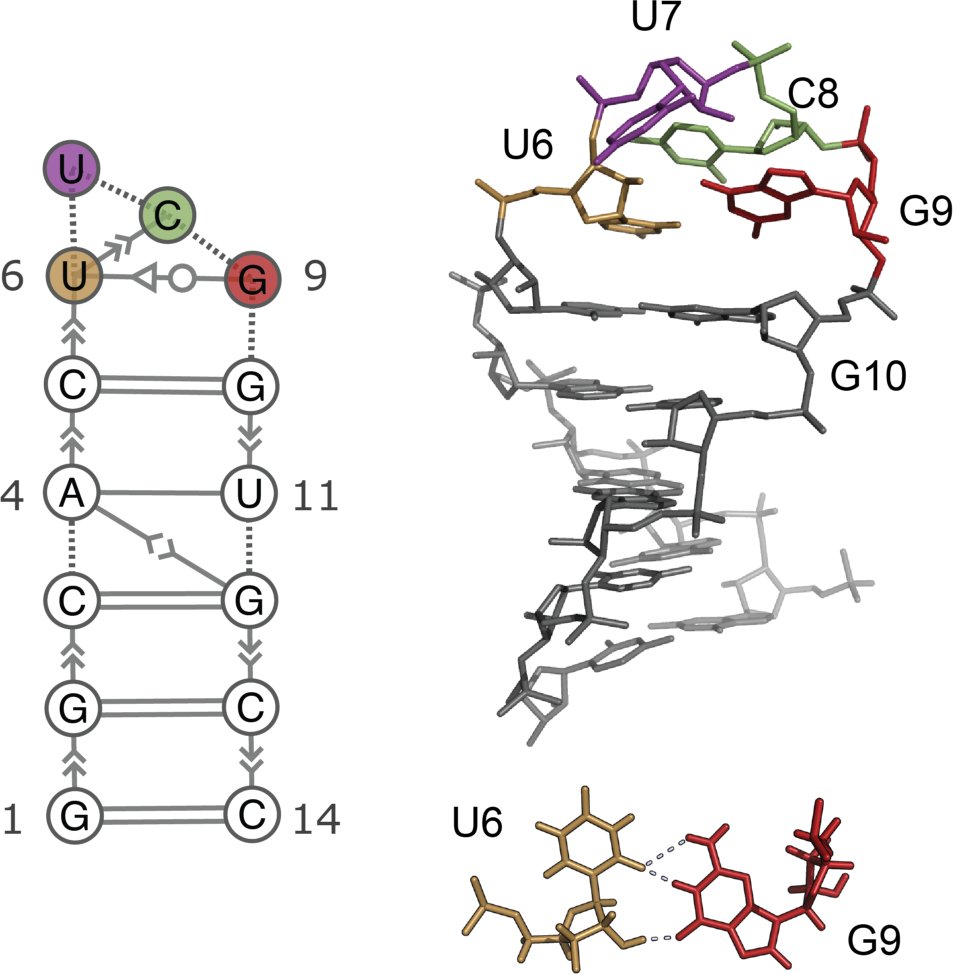
Consensus secondary structure (left) and three dimensional structure (right) of the UUCG tetraloop *Nozinovic et al.* (*2010*). The stem is formed by 5 consecutive Watson-Crick base-pairs capped by the loop U6-U7-C8-G9. One of the most distinctive feature of this structure is the trans-Sugar-Watson interaction between U6 and G9 (bottom). Extended secondary structure annotation follows the Leontis-Westhof nomenclature *Leontis and Westhof* (*2001*)

Despite its stability, the UUCG tetraloop is not rigid. In particular, three recent studies by independent groups indicate the presence of alternative loop conformations *Borkar et al.* (*2017*); *Hartlmüller et al.* (*2017*); *Nichols et al.* (*2018b*). Earlier NMR studies *Nozinovic et al.* (*2010*); *Duchardt and Schwalbe* (*2005*) also suggested the presence of loop dynamics, without providing a detailed structural interpretation of the data. More generally, the atomic-detailed characterization of RNA structure and dynamics requires specialized techniques and substantial experimental effort, including NMR measurements of nuclear Overhauser effects (NOE), scalar couplings, chemical shifts, residual dipolar couplings, cross-correlated relaxation rates as well as a wide range of relaxation-dispersion type NMR experiments *Salmon et al.* (*2014*); *Marušič et al.* (*2019*).

While NOEs are typically used to determine RNA and protein structures, they also contain dynamic information. Because ensemble-averaged NOEs are highly sensitive to the underlying distance 2uctuations, they may contain contributions even from minor populations. Normally, such information is diZcult to extract because standard NOE measurements are relatively inaccurate. It has, however, been demonstrated that a substantial part of the information content inherent to these probes can be obtained from exact NOE measurements (eNOEs) *Nichols et al.* (*2018b*). As opposed to conventional NOEs, eNOEs can be converted into tight upper and lower distance limit restraints *Vögeli* (*2014*); *Nichols et al.* (*2017*, 2018a).

Previous computational studies of the UUCG tetraloops focused either on the dynamics around the near-native state *Giambaşu et al.* (*2015*) or on the diZculty in separating force-field inaccuracies from insuZcient sampling *Banás et al.* (*2010*); *Bergonzo et al.* (*2015*). In a previous study we reported converged free-energy landscape for RNA 8-mer and 6-mer loops, and we have shown that native-like states are not the global free-energy minimum using the current AMBER RNA force-field *Bottaro et al.* (*2016*). This problem has been addressed in a new parameterization of the AMBER force-field, that improves the description of the UUCG 14-mer and other RNA systems *Tan et al.* (*2018*). Nevertheless, it remains diZcult to assess the accuracy of these simulations, because experiments alone do not provide an atomic-detailed description of structure and dynamics that serve as a benchmark.

Here, we use extensive atomistic MD simulations to map the conformational landscape of the UUCG tetraloop using enhanced sampling techniques and a recent force-field parameterization. To improve the description of this system further, we perform an a posteriori refinement of the MD simulation using experimental data via a Bayesian/maximum entropy procedure *Hummer and Köfinger* (*2015*); *Bottaro et al.* (*2018a*). We validate the eNOE-refined ensemble against independent NMR measurements and find an agreement that is on average comparable with NMR structures of the UUCG tetraloop deposited in the Protein Data Bank (PDB).

Our experimentally-refined ensemble reveals the presence of two conformational states. The dominant, major state (here called state A) is the consensus UUCG structure shown in Fig. 1. The second, previously unreported lowly-populated state (state B) is characterized by the absence of the signature U6-G9 non-Watson-Crick base pair, with the C8 and G9 bases exposed into solution. We employ a random forest classifier to identify the structural properties that discriminate state A from state B. Furthermore, we use the same method in the space of experimental data to identify specific measurements that are most sensitive to the presence of state B. By construction our refined ensemble better agrees with eNOE compared to the original MD simulation and to the consensus PBD structures.

The paper is organized as follows: we first compare the predictions obtained from MD simulation against different experimental datasets. We then discuss the effect of the refinement procedure, showing how it improves the agreement with experiments and how it affects the population of different conformations. We proceed by identifying the relevant degrees of freedom and contacts that characterize the two states. Finally, we identify experiments sensitive to the presence of state B. We accompany this paper with the commented code, in form of Jupyter notebooks, to reproduce step-by-step the complete analysis, including all figures and supplementary results presented in the manuscript.

## Results

### MD simulations and comparison with experimental data

We simulate the RNA 14-mer with sequence GGCACUUCGGUGCC starting from a completely extended conformation. Studying the folding free-energy landscape of this system is computationally expensive: for this reason previous attempts required *µs*-long simulations in combination with tempering protocols *Tan et al.* (*2018*); *Kuhrova et al.* (*2013*); *Chen and García* (*2013*).

Here, we combine two enhanced sampling techniques: solute tempering in the REST2 formulation *Wang et al.* (*2011*) and well-tempered metadynamics *Barducci et al.* (*2008*). We used a nucleic-acid specific metric, called eRMSD, *Bottaro et al.* (*2014*) as a collective variable for enhanced sampling. The MD simulation setup and convergence analysis are presented in supporting information 1 (SI1).

Before describing the conformational ensemble provided by MD, we compare the computational prediction with available NMR spectroscopy data. More precisely, we consider the following experimental datasets:

- Dataset A. Exact eNOEs *Nichols et al.* (*2018b*), consisting in 62 bidirectional exact NOE, 177 unidirectional eNOE and 77 generic normalized eNOE (gneNOE). This dataset alone was used to determine the structure of the UUCG tetraloop with PDB accession codes 6BY4 and 6BY5. In addition to the original dataset, we added 1 new eNOE and 6 new gn-eNOEs, as described in SI2.
- Dataset B. 97 ^3^J scalar couplings, 31 RDCs and 250 NOE distances. This data, among other NMR measurements, was used to calculate the consensus UUCG tetraloop structure (PDB 2KOC *Nozinovic et al.* (*2010*)).
- Dataset C. 38 (RDC1) plus 13 (RDC2) residual dipolar couplings. These RDCs have been used in conjunction with MD simulations to obtain a dynamic ensemble of the UUCG tetraloop. *Borkar et al.* (*2017*).
- Dataset D. 91 solvent paramagnetic resonance enhancement (sPRE) measurements *Hartlmüller et al.* (*2017*).

The orange bars in the four panels of Fig. 2 show the agreement between simulation and the different experimental datasets. The agreement between experiment and simulations is expressed using the reduced *χ*^2^ statistics, defined as the average square difference between the experimental measurement (*F*^*exp*^) and the back-calculated ensemble average (< *F* (x) >) normalized by the experimental error *σ*:

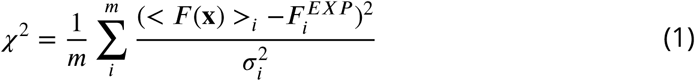

**Figure 2.**
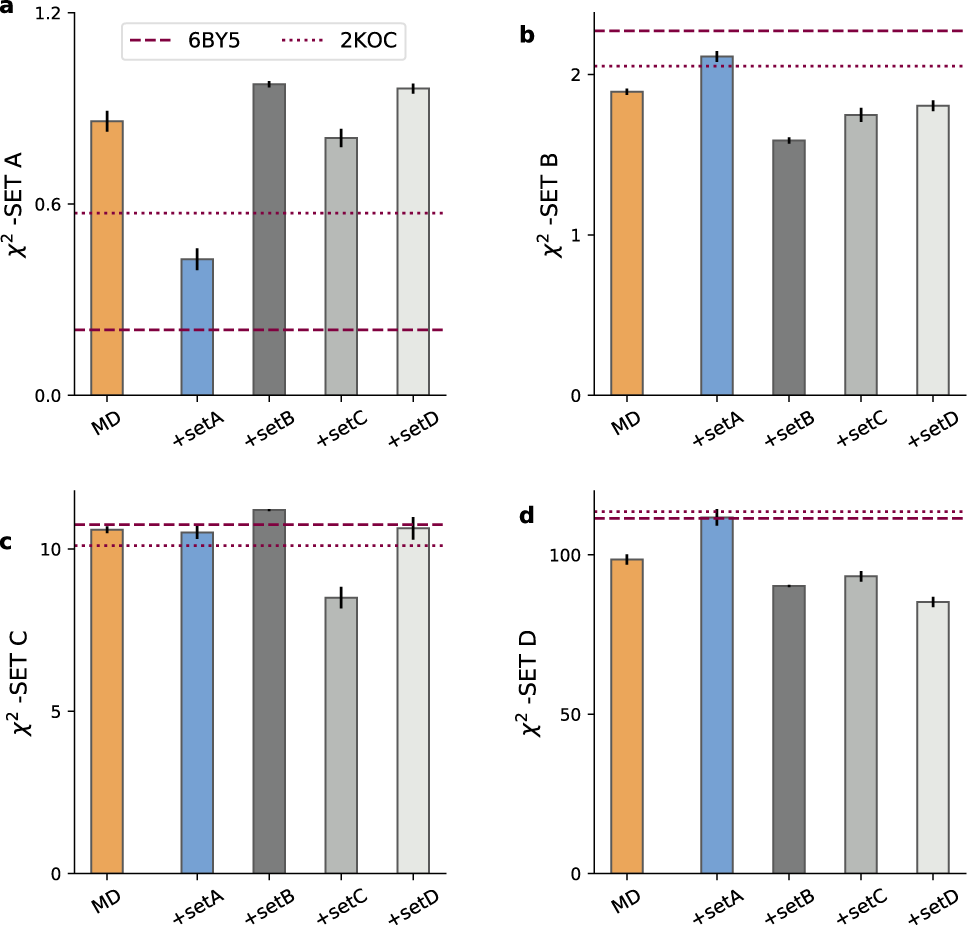
Comparison between experiment and simulations. **a)** *χ*^2^ between experimental dataset A against the MD ensemble (MD) and against the refined ensembles (MD+set A, MD+set B, MD+set C, MD+set D). As a reference, values calculated from all NMR models from PDB structures 2KOC and 6BY5 are shown as dashed lines. The agreement between the same ensembles and datasets B, C, D, are shown in panels **b,c**, and **d**, respectively. Error bars show the standard error estimated using four blocks.

Hence, the lower the *χ*^2^, the better the agreement. As a rule of thumb, *χ*^2^ < 1 can be considered small, as the difference between experiment and prediction is within experimental error. See SI2 and SI3 for additional details on this comparison. As a reference, we report in Fig. 2 the agreement calculated on the PDB ensembles 6BY5 *Nichols et al.* (*2018b*) and 2KOC *Nozinovic et al.* (*2010*). For set A, the agreement of the MD with experiment is considerably poorer than the one calculated on 6BY5. We recall that this latter ensemble was determined by fitting dataset A, we thus expect *χ*^2^ to be small in this case. On datasets B, D the MD better agrees with experiments compared to 2KOC, 6BY5, while differences on set C are smaller. When considering other statistics (e.g. root mean square error, Pearson correlation, number of violations), the same conclusions apply. Note that *χ*^2^ for sPRE and for scalar couplings (SI 3) is very large. This discrepancy may arise both from the imperfect ensembles, from the underestimation of the experimental error, as well as from the limitation of the function used to calculate the experimental quantity from the atomic positions (i.e. the forward model). As an example, the parameters in the Karplus equation for HCOP couplings critically depend on a single experimental data point measured in 1969 *Bentrude and Hargis* (*1969*).

### Bayesian/Maximum entropy refinement of the MD ensemble

As described above, our MD simulation provides a conformational ensemble consisting of a rich and diverse set of conformations, that, however, as a whole does not match all experimental data perfectly.

In order to improve the description provided by the MD simulation, we calculate a refined conformational ensemble by a posteriori including experimental information into simulations. In brief, the refinement is obtained by assigning a new weight to each MD snapshot, in such a way that the averages calculated with these new weights match a set of input (or “training”) experimental data within a given error. Among all the possible solutions to this underdetermined problem, we use the one that maximize the Shannon cross-entropy *Pitera and Chodera* (*2012*); *Boomsma et al.* (*2014*); *Orioli et al.* (*2019*).

Since we have four independent experimental datasets, we can refine the simulation by using one of them, using the other three as test set. We choose the free hyper-parameter of the algorithm (*θ*) by performing a k-fold cross-validation procedure (see SI 4). By construction, the refinement procedure improves the agreement on the training data, but it does not guarantee improved agreement on the test sets. Figure 2 shows the effect of the refinement for different combinations of training/test set. For example, the blue bar in panel **a** shows the agreement between experiment and the MD simulations refined against set A (MD+set A) evaluated on set A itself (training). The blue bars in panels **b,c,d** show the *χ*^2^ of MD+set A on datasets B,C,D (test). The MD+set B is trained on set B and tested on datasets A,C,D, and so on.

We observe that including into simulations dataset A has a small, yet detrimental effect on the agreement with sets B and D. The MD+set B ensemble shows improved agreement with dataset D, but performs worse than the original MD on set A and C. We observe a similar behavior for MD+set D. Among the four refined ensembles the MD+set C is the one that behaves better, as it shows a smaller or equal *χ*^2^ (relative to MD) on the three tests sets A,B,D.

Taken together, our results show that the MD and the refined ensembles fit available experimental data to a degree that is comparable to the one calculated from PDB structures 2KOC and 6BY5 (Fig. 2). However, there exists substantial discrepancies with experimental data, and none of the considered ensembles clearly outperforms the others.

### Free energy landscape

In this section we analyse in detail the refined MD ensembles, and discuss the differences with respect to the original simulation and previously determined structures. We consider the histogram of the distance (structural dissimilarity) from the consensus structure (PDB 2KOC). Distances are measured using the eRMSD, a nucleic-acid specific metric that takes into account both position and orientations between nucleobases *Bottaro et al.* (*2014*), although a similar picture is obtained using the standard RMSD metric.

The distribution obtained from the original MD ensemble is shown in orange in Fig. 3**a**, and repeated in the four panels. The two peaks correspond to structures where the stem is fully formed, but with different loop conformations. Structures in the left peak (eRMSD<0.7, state A) display the signature interactions present in the consensus structure, while in the second peak (state B) the loop is disordered. The population of state A in our MD is 60% ± 4, larger than the one calculated from the 180*µ*s simulated tempering simulation by Tan et al *Tan et al.* (*2018*) (≈ 40%).

**Figure 3.**
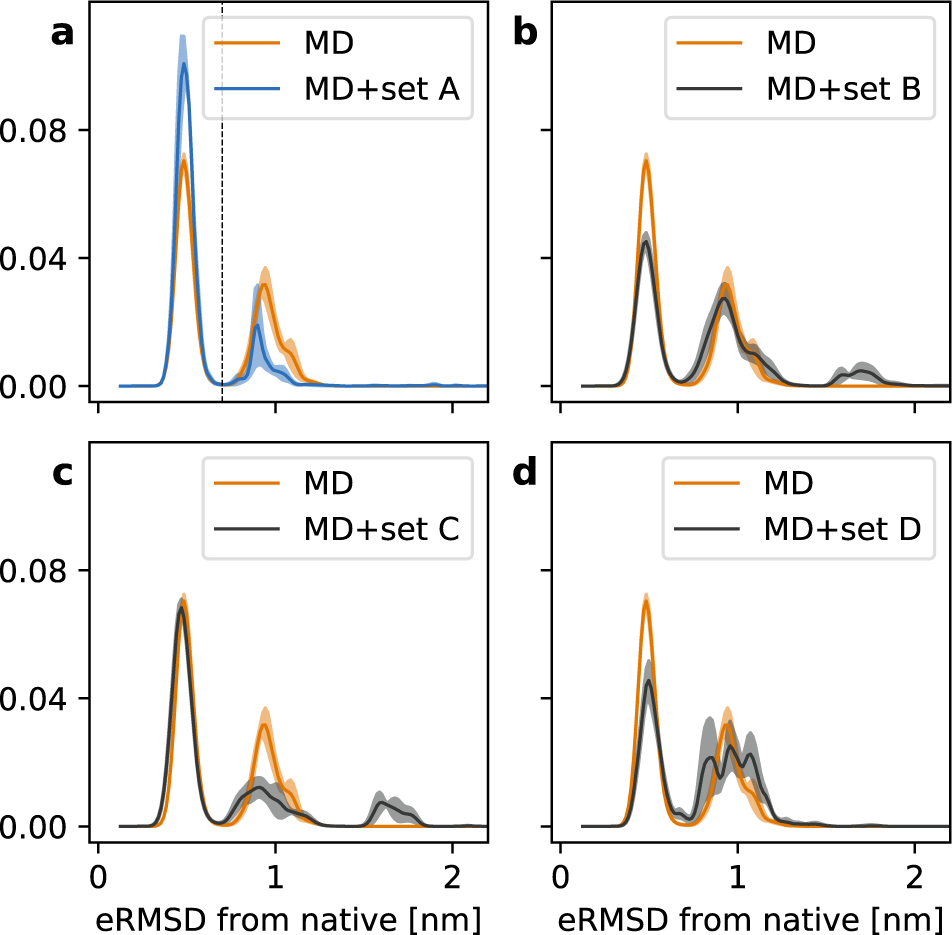
Histograms of the eRMSD from native. The original MD simulation (orange) is compared with the four refined ensembles: MD+set A in panel **a**, MD+set B in panel **b**, MD+set C in panel **c**, and MD+set D in **d.** Shades show the standard error estimated using four blocks. The vertical dashed line in panel **a** shows the separation between state A (eRMSD *<* 0.7) and state B (eRMSD ≥ 0.7).

The inclusion of experimental data in simulations affects the histogram in different ways. When including eNOE data from dataset A (Fig 3a, blue), we observe an increase in the population of state A to 83% ± 16. The refinement obtained with datasets B and D (Fig. 3 panels **b,d)** reduces the population of state A down to 44% ± 6 and 44% ± 10, respectively. In the MD+set C ensemble, the population of state A is unaffacted (65 ± 5). Interestingly, we observe one new peak appearing at eRMSD *>* 1.5, also present in the MD+set B ensemble. This peak corresponds to fully extended conformations, and it is present when including RDC measurements. If we trust our simulation, the forward model including the complex issue of estimating the alignment tensor, and the refinement procedure, we can conclude that RDC are sensitive to a small population of either dimers or unfolded structures. Unfortunately, we do not have the possibility to test this hypothesis, which is left for future investigations. Here, instead, we focus on the results obtained from MD+set A. First, because this ensemble has the highest population of state A, and it is thus less “surprising”. Second, because we better understand and control the experimental data, and lastly because we can take advantage of our previous experience in using NOE for RNA ensemble refinement *Bottaro et al.* (*2018a*).

### Structural differences between state A and B

Having discovered this new B-state, we proceed to analyse its structural features. While state A is known and structurally well-defined (Fig. 4a), it is not trivial from a simple visual inspection to identify which are the main features of state B (Fig. 4b). Here, we address this question by using a random forest classifier. In practice, we first extract samples from the MD+set A ensemble using a bootstrapping procedure. Second, we assign a label to each sample depending on the distance from native (state A if eRMSD*<*0.7, and state B otherwise). Last, we calculate structural properties (e.g. torsion angles, distances) for each of the the labelled sample, that are used to train a random forest classifier. In this context we are not interested in the decision tree per se, but rather in its ability to rank the importance of the input features in the classification problem. Thus, we can find the most relevant degrees of freedom that discriminate the two states. The result of such analysis on all dihedral angles *α, β, γ, δ, ϵ, ζ, χ* in the 14-mer reveals that the two highest-ranked angles are *ζ* in C8 and G9. Fig. 4d,e show the histograms for these two angles: the angle in C8 is a good classifier, as all samples from state A (empty curve) are in *gauche*+ conformation, while all samples from state B (filled curve) are in *gauche*−. Information on state A/B differences are also contained in G9 *ζ*, although the separation of the two states is not as striking as in C8. The importance of C8 and G9 is further confirmed when using the distance between the center of the six-membered rings in the nucleobases as input features. In this case, the distances between U6-C8 and U6-G9 are the two most important degrees of freedom that distinguish state A from state B. The distribution of these two distances is shown in Fig. 4c. In the consensus structure U6 and G9 interact through a trans sugar-Watson base-pair and U6 and C8 are stacked. State B is characterized instead by the absence of the stacking interaction (large U6-C8 distance), and of the non-Watson-Crick base-pair (large U6-G9 distance).

**Figure 4.**
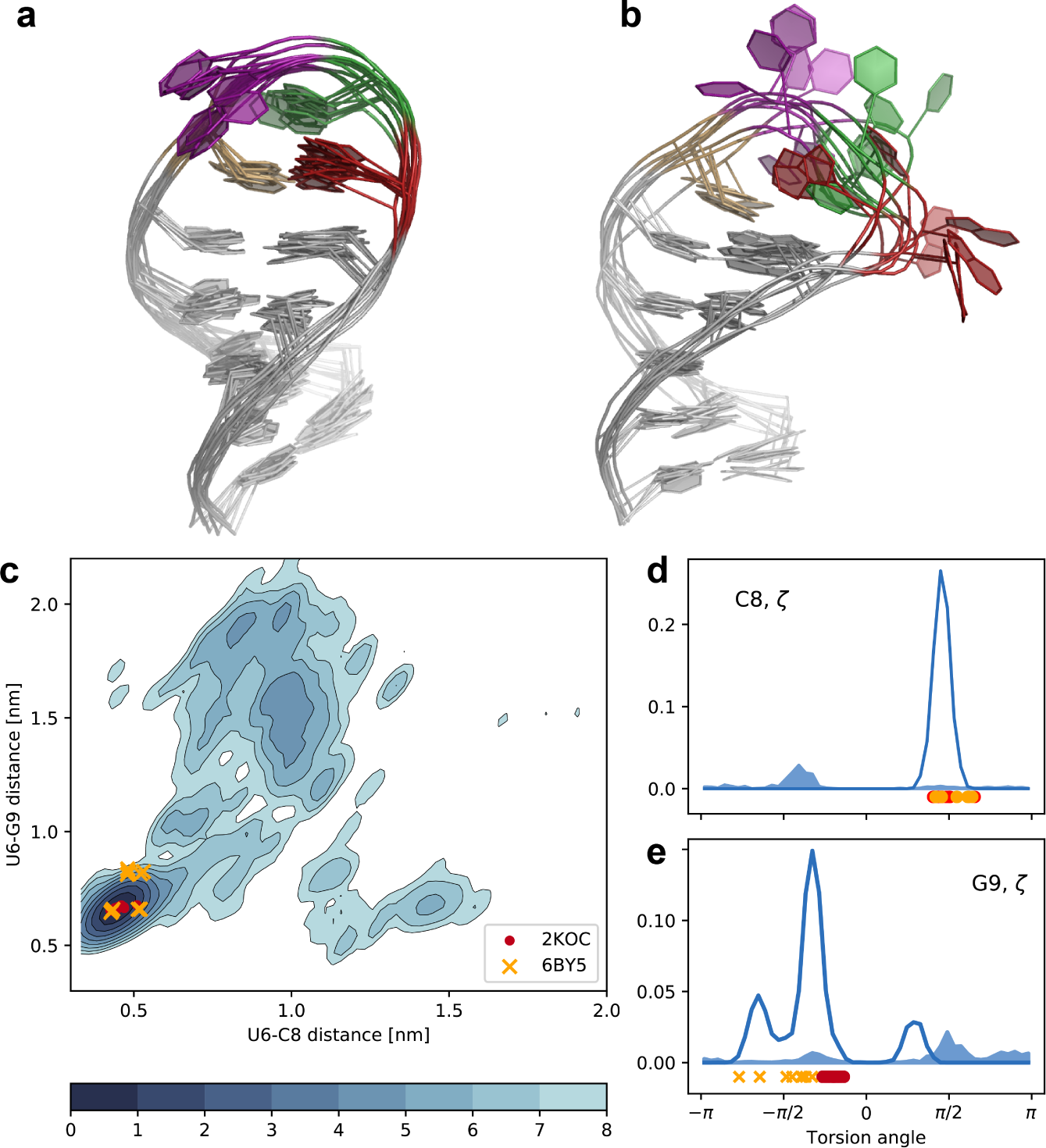
**a)** Representative (random) conformations sampled from state A. **b)** Representative (random) conformations sampled from state B. The color code is identical to Fig. 1: U6 in ochre, U7 in purple, C8 in green and G9 in red. **c)** Free energy surface projected onto the the U6-C8/U6-G9 distance between ring centres. The units of the colorbar are in *k*_B_T. **d)** Histogram of *ζ* dihedral angle in C8. The open filled area indicates conformations belonging to state A, and the filled area indicates conformations belonging to state B. **e)** Histogram of *ζ* dihedral angle in G9.

### Experimental measurements sensitive to the presence of state B

In the previous section we have described the structural differences between state A and state B, and we now seek for additional experimental validation. In particular, we would like to answer the following question: does the MD+set A ensemble provide a better loop description compared to the consensus NMR structure 2KOC?

The four experimental sets contain hundreds of datapoints in total, and it is therefore not trivial to identify specific measurements that probe directly the presence of state B. In machine learning this is called a feature importance problem, that we solve using a random forest classifier, as we did in the previous section. Here, however, the features are not structural properties, but back-calculated experimental data. We construct a random forest classifier for each experimental dataset, and we obtain four sub-datasets with the features (measurements) that are most important for classifying (i.e. separating) state A and B. Consistently with Fig. 4, we find that these measurements involve nucleotides in the loop region.

For each measurement in the sub-datasets, we calculate the difference between the ensemble average and experiments divided by the error *σ* (i.e. the Z-score).

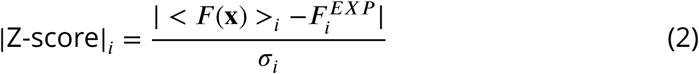

Figure 5 shows the Z-score calculated on the MD+set A ensemble (x-axis) against the same quantity calculated on the 2KOC structures. Points above the diagonal (such as 1 in panel a, 6 in panel d) indicate measurements for which the MD+set A ensemble better agrees with experimental data (|Z-score|_MD+set A_ < |Z-score|_2KOC_), while points below the diagonal (e.g. 3,4,7,8) indicate the opposite.

**Figure 5.**
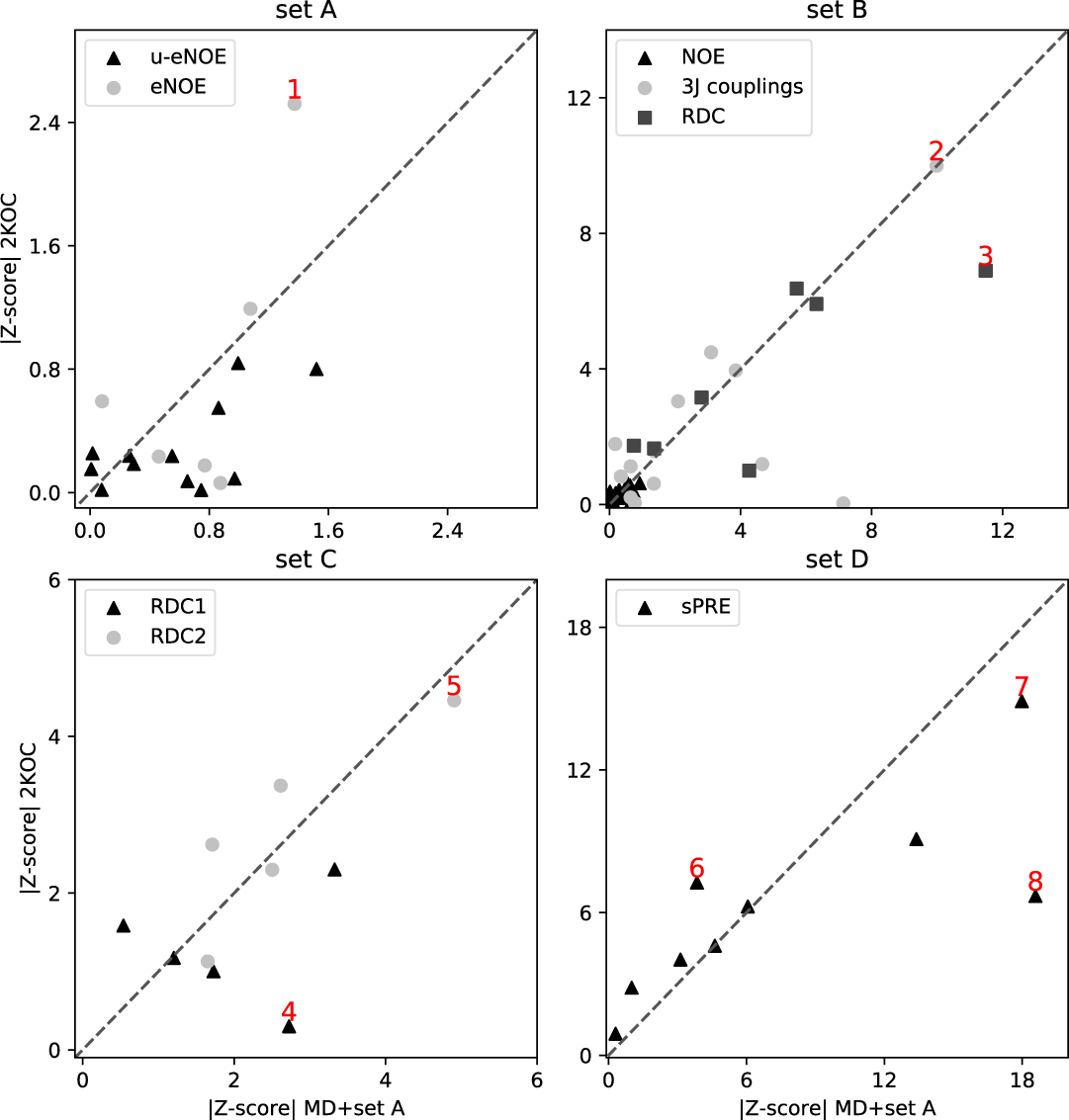
Comparison between 2KOC and MD+set A ensemble on experimental data that are most sensitive to the presence of state B. Scatter plots show the Z-score calculated on the MD+set A ensemble (x-axis) versus the same quantity calculated on the PDB ensemble 2KOC (y-axis). The four panels show data belonging to the four datasets: set A in panel **a**, set B in panel **b**, set C in panel **c**, and set D in panel **d**. Points discussed in the main text and shown in Fig. 6 are labeled in red.

Overall, there are more points below the diagonal (42) than above (36), suggesting that the original 2KOC ensemble provide a better description of the loop region. For example, the RDC in set C 8C3’-8H3 is within experimental error for 2KOC, but not for our MD+set A ensemble (point 4, Fig. 6). Conversely, the eNOE C8 H5’ to G9 H8 in dataset A is significantly closer to the experiment in the MD+set A ensemble (point 1, Fig. 6). Note that major discrepancies are present in both ensembles, such as point 2 (^3^J in G9 H5”-H4), point 3(RDC C8 C1’-H1’), point 5 (RDC G10 C8-H8), and point 7 (sPRE G10 1H5’). Again, we stress that these discrepancies can be ascribed to errors in the ensembles, but also to inaccuracies in the empirical model employed to calculate experimental data from structures, or to errors in the data.

**Figure 6.**
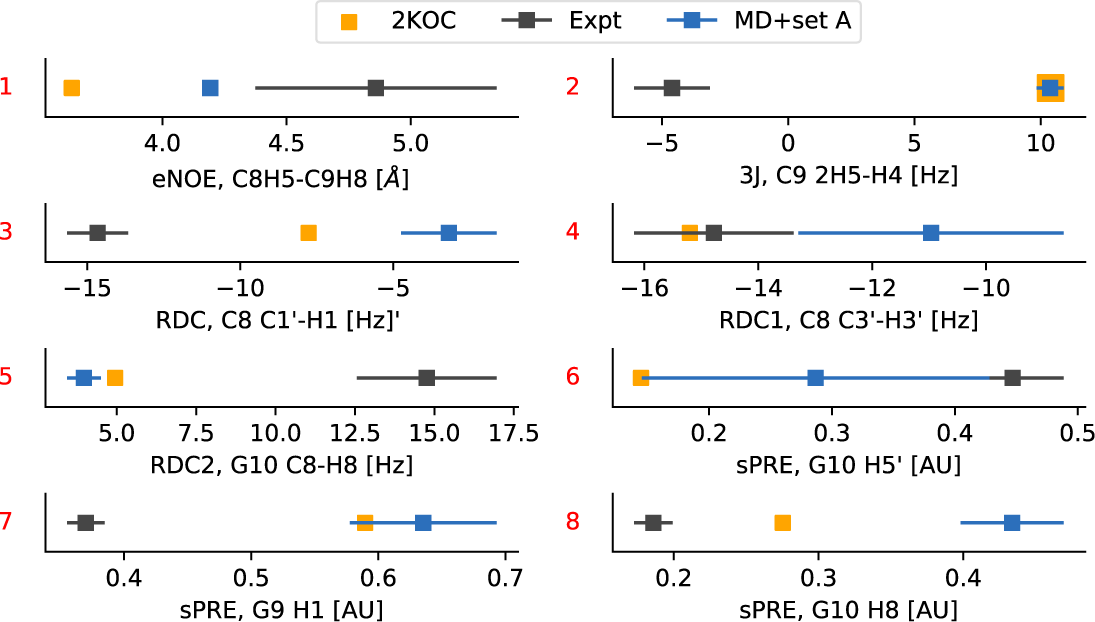
Comparison between calculated and experimental data for selected measurements discussed in the text and labelled as in Fig.5.

## Conclusions

Based on our extensive MD simulations and integrating them with exact NOE data, we report the free energy landscape of a prototype stem-loop RNA 14-mer known as the UUCG tetraloop. By combining a recently refined force field for RNA with enhanced sampling MD we were able to fold the tetraloop to its native conformation(s) as judged by very good agreement with several sets of experiments. The main finding of the present study is the presence of a low populated, non-native conformation (state B). The low-populated state differs from the consensus structure (state A) only in the loop region, and it is characterized by the absence of the tSW base-pair between U6 and G9, with C8 and G9 partially exposed into solution (Fig. 4). This result has been obtained by using atomistic MD simulations and eNOE, without the need of additional data.

The free-energy surfaces and estimated population provided here are based on the available experimental data, on the employed model, and the extent of our sampling. Therefore, they are subject to inaccuracies. However, both simulations and eNOE data are consistent with the presence of the B state as described in this paper. This interpretation is qualitatively consistent with several NMR studies, that also suggested the presence of dynamics in G9 *Duchardt and Schwalbe* (*2005*); *Nozinovic et al.* (*2010*); *Hartlmüller et al.* (*2017*). Conversely, on-resonance 13C R1*ρ* relaxation dispersion experiments on a UUCG tetraloop with a different stem sequence showed no significant exchange contributions, indicating the absence of motions with substantial chemical shift variation in the *µ*-ms timescale *Salmon et al.* (*2015*). Note also that G9-exposed structures were reported in previous MD simulations *Kuhrova et al.* (*2013*); *Bergonzo et al.* (*2015*); *Cesari et al.* (*2019*), suggesting our finding to be robust with respect to the choice of the force-field and water model.

In this work we have used eNOEs to reweight a posteriori the ensemble generated via enhanced sampling MD simulations. This refinement procedure is a computationally cheap post-processing *Graf et al.* (*2007*); *Hummer and Köfinger* (*2015*); *Bottaro et al.* (*2018a*) that allows one to try different combinations of training and test sets, as we did in this work *Orioli et al.* (*2019*). Note that refinement is in principle less powerful compared to on-the-2y methods that samples directly from the target probability distribution *Bonomi et al.* (*2017*); *Reißer et al.* (*2019*).

In our study we refine the simulation by matching RDC data (set B, C) or solvent PRE (set D) as well. Only when we use set C for training we obtain an improved or equal agreement on the test sets relative to the original MD simulation (Fig. 2). Additionally, different data affect the MD conformational ensemble in different ways (Fig. 3). Several reasons can contribute to this behaviour. First, we do not expect all experimental data to be perfectly compatible one with the other, because measurements were conducted in similar, but not identical conditions. Second, the forward models might not be accurate for arbitrary molecular conformations. For example, if the forward model accurately predicts the RDC given the native structure, but fails on unfolded/misfolded conformations, we obtain artefacts that cannot be easily accounted for in our refinement procedure. Note that this problem is typically less relevant when using experimental RDC, sPRE or chemical shift data for scoring structures *Sripakdeevong et al.* (*2014*); *Salmon et al.* (*2015*); *Hartlmüller et al.* (*2017*).

Based on the above observations, and considering our previous experience with eNOE data, we here analyse in detail the results obtained using MD refined using set A (Fig. 3a). The structural features that are most important to discriminate between state A and state B are identified using a random forest classifier. The problem of concisely interpret differences in biomolecular conformations has been recently pursued using a variety of machine learning methods, including linear discriminant analysis *Piccini et al.* (*2018*), decision trees *Brandt et al.* (*2018*), and others *Fleetwood et al.* (*2019*). In this work we extend this idea, and use back-calculated experimental data as input features for the random forest classifier. In this way we identify individual (available) measurements that are most sensitive to the presence of state B. Note that this approach can also be used in a generative fashion to design experiments that probe the existence of specific conformational states.

We closely inspect the selected set of measurements that are sensitive to state B (Fig. 5). In the majority of the cases we find the presence of the additional state to provide a worse agreement with experiment compared to the consensus NMR structure (PDB code 2KOC) (see e.g. Fig. 6, points 3,8). In other cases (Fig. 6, points 1,6), instead, the MD+set A performs better than 2KOC. Several other data significantly deviate from experiments in both ensembles (Fig. 6, points 2,5,7). This suggest the possibility that conformations that are different from state A are indeed present, but do not correspond to the state B as described in Fig.4b.

Finally, we note that the approach taken here is general and it is applicable to other RNA or protein systems *Escobedo et al.* (*2019*); *Crehuet et al.* (*2019*). Previous characterization of slow, larger motions in RNA molecules have mostly relied on relaxation-dispersion, chemical exchange saturation transfer or related NMR experiments that probe chemical shift differences between different conformational states. We hope that the integration of MD simulations and eNOE measurements provides further opportunities for characterizing the free energy landscapes of RNA molecules.

## Acknowledgements

The research is funded by a grant from The Velux Foundations (S.B. and K.L.-L.), the Lundbeck Foundation BRAINSTRUC initiative (K.L.-L.), a start-up package from the University of Colorado (B.V.), and by NSF grant 1917254 for Infrastructure Innovation for Biological Research (B.V.). The authors thank Giovanni Bussi for helpful comments. We also thank Tobias Madl, Harald Schwalbe, and coworkers for useful discussions and for sharing their data.

## Data Availability

Jupyter notebooks to reproduce the analysis and all figures are included as supporting information. The MD trajectory, together with the experimental data are hosted on github (https://github.com/KULL-Centre/papers/tree/master/2020/UUCG-dynamics-Bottaro-et-al). The Plumed input file to reproduce the simulation is hosted on the Plumed nest *Bonomi et al.* (*2019*) under the accession code 19.070.

## Methods

### Integrating MD simulation and experimental data

We combine the MD simulation with experimental data using a maximum entropy/Bayesian procedure *Boomsma et al.* (*2014*); *Beauchamp et al.* (*2014*); *Hummer and Köfinger* (*2015*). In our previous work, we have described this reweighting procedure as Bayesian/MaxEnt (BME) *Bottaro et al.* (*2018b*,a). In BME we use the experimental data to modify a posteriori the simulation so that the new conformational ensemble has the following properties: (i) the calculated averages are close to the experimental values taking uncertainty into account and (ii) it maximizes the relative Shannon entropy with respect to the original simulation ensemble. The modification comes in the form of a new set of weights 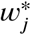, one for each simulation frame.

It can be shown that this problem can be cast as a minimization problem, in which one seeks the minimum of the function Γ with respect to the set of Lagrange multipliers 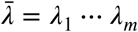, with *m* being the number of experimental constraints.

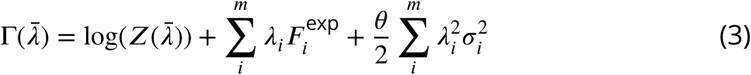

Here, *σ*_*i*_ are the uncertainties on the experimental measurements 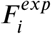 and include experimental errors and inaccuracies introduced by the calculation of the experimental quantity from the atomic positions (*F* (x)). *θ* is a free parameter, while the partition function *Z* is defined as

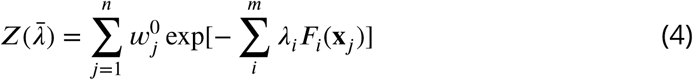

The sum over the index *j* runs over the *n* frames in the simulation, and 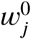 are the original weights. *w*^0^ = 1/*n* when using plain MD simulations or enhanced sampling techniques that sample directly from the target distribution (e.g. parallel tempering). In this paper we use WT-METAD, and the original weights *w*^0^ are estimated using the final bias potential *Branduardi et al.* (*2012*). The minimization of Eq. 3 yields a set of Lagrange multipliers 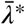 that are used to calculate the optimal weights

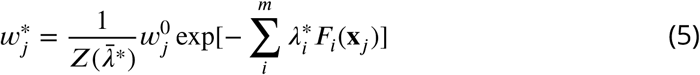

In the context of the UUCG tetraloop, we use the dataset A described in the previous section to refine the simulation ensemble, and cross-validate the results against datasets B, C, and D. Details on the comparison between simulations and experiments, on the BME procedure and on the choice of the regularization parameter *0* can be found in SI 4.

## Random Forest Classifier

The random forest analysis is set up according to the following procedure:

- Bootstrap *n* = 50000 samples from the MD simulation trajectories. The weight of each sample, 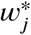, is calculated as described above.
- Samples are assigned to state A if the eRMSD from the first model of the NMR structure 2KOC is less than 0.7, and state B otherwise.
- Calculate a set of *m* features (torsion angles, distances between ring centres, experimental data) for each sample *Bottaro et al.* (*2019*).
- Construct a random forest classifier using *n* samples, *m* features and 2 classes (state A and state B). In our work, we use the implementation of the random forest algorithm *Breiman* (*2001*) available in sklearn 0.22 using a maximum tree depth of 3. 80% of the samples are used for training, while the remaining 20% are used to evaluate the accuracy of the classifier (>97% in all cases).
- Rank the *m* features by their importance. In the analysis of the experimental data in Fig. 5, only features with importance greater than 0.2 are shown.

