## Supplementary material for "Integrating NMR and Simulations Reveals Motions in the UUCG Tetraloop": Figure 5C (from another version of this preprint: version 2, 2019-09-11)

figure5C


### Figure 5 - HLDA on NOE¶

In this notebook we perform the HLDA analysis on potential NOE. To this end we perform the following steps:

1. Take all possible proton-proton pairs in the 14-mer between non-consecutive nucleotides.
2. Calculate all the distances, and remove the pairs for which the average NOE is larger than 0.6nm
3. This results in approximately 800 H-H pairs, that we use as descriptors in HLDA
4. We sort the distances according to the eigenvector coefficient.

In the last part of the notebook we use the calculated distances to produce figure 5 in the manuscript.
Note that the three new eNOE are NOT used in the BME refinement procedure.

In [20]:

```
import mdtraj as md
import numpy as np
import pickle
import os

top = "data/PDB/2koc_gmx.pdb"
t = md.load(top)

# find proton pairs in non-consecutive residues
pairs = []
labels = []
residues = [res for res in t.topology.residues]
for i in range(len(residues)):
    for k in range(i+1,len(residues)):
        for at1 in residues[i].atoms:
            n1 = str(at1)
            if("H" in str(at1)):
                for at2 in residues[k].atoms:
                    n2 = str(at2)
                    if("H" in str(at2)):                      
                        labels.append(n1+","+ n2)
                        pairs.append([at1.index,at2.index])

print("# Found %d proton-proton pairs" % len(pairs))

# read weights 
theta=30
weights_1 = np.array([float(line.split()[1]) for line in open("data/weights_set_A1_t_%d.dat" % (theta))])

# calculate the proton-proton pairs whose ensemble average is smaller than 0.6nm
# we use pickles because calculating 10k distances takes time
fname = "data/close_hh_indeces.p"
threshold =0.6 
traj_md = md.load("data/traj_temp_f_0.xtc",top=top)

if os.path.isfile(fname):
    low_idx = pickle.load(open(fname,'rb'), encoding='bytes')
else:
    dd_all = md.compute_distances(traj_md, pairs)
    # calculate 1/6 average
    averages_all = np.power(np.average(np.power(dd_all,-6.),weights=weights_1,axis=0),-1./6.)
    low_idx = np.where(averages_all<threshold)[0]
    pickle.dump(low_idx ,open(fname, 'wb'))

print("# The number of pairs below the threshold is %d" % len(low_idx))
```

```
# Found 10866 proton-proton pairs
# The number of pairs below the threshold is 789
```

In [21]:

```
#%rm data/close_hh_indeces.p
```

In [22]:

```
# define new pairs and labels
new_pairs = [pairs[j] for j in low_idx]
new_labels = [labels[j] for j in low_idx]

# calculate these distances
trajA = md.load("data/stateA.xtc",top=top)
trajB = md.load("data/stateB.xtc",top=top)

ddA = md.compute_distances(trajA, new_pairs)
ddB = md.compute_distances(trajB, new_pairs)
ddA6 =np.power(ddA,-6.)
ddB6 =np.power(ddB,-6.)

# read ermsd from pickle to get the weights corresponding to state A and B
fname = "data/observables/ermsd_full.p"
ermsd_full = pickle.load(open(fname,'rb'), encoding='bytes')
stateA = np.where(ermsd_full<0.7)[0]
stateB = np.where((ermsd_full>=0.7) & (ermsd_full<1.4))[0]  

weights_stateA = weights_1[stateA]
weights_stateB = weights_1[stateB]
```

In [23]:

```
# Do HLDA using my script

import sys
sys.path.append("/Users/sandrobottaro/Software/HLDA")
import HLDA as hlda

# initialize class
hlda_dd = hlda.HLDA()

# add data
n = hlda_dd.load_data(ddA6,oweights=weights_stateA)
n = hlda_dd.load_data(ddB6,oweights=weights_stateB)

# run the algorithm
hlda_dd.run_lda(harmonic=True)

# get the coefficients of the evigenvector corresponding to the largest eigenvalue
eigenvalues = hlda_dd.get_evals()
coefficients = hlda_dd.get_evecs()[:,-1]
```

```
# initializing HLDA class.
# Processing data with 25124 samples and 789 features 
# Processing data with 43169 samples and 789 features
```

In [24]:

```
# plot eigenvalues. Only the last one is non-zero
import matplotlib.pyplot as plt
import seaborn as sns

fig,ax = plt.subplots(1,1,figsize=(18,5))
plt.scatter(range(len(eigenvalues)),eigenvalues)
plt.xlabel("eigenvalue index")
plt.ylabel("eigenvalue")
plt.show()
```

In [25]:

```
# calculate the NOE for state A, state B and on the whole ensemble
averagesA = np.power(np.average(ddA6,weights=weights_stateA,axis=0),-1./6.)
averagesB = np.power(np.average(ddB6,weights=weights_stateB,axis=0),-1./6.)

dd_MD6 = np.power(md.compute_distances(traj_md, new_pairs),-6.)
averages_MD = np.power(np.average(dd_MD6,weights=weights_1,axis=0),-1./6.)
```

In [26]:

```
# plot H-H contacts with the highest coefficients 

colors = ["pale red","amber","cool blue","battleship grey"]
cp = sns.xkcd_palette(colors)

cmap1 = sns.diverging_palette(220, 20, n=7)

#fig = plt.figure(figsize=(7.5,3.5))
fig = plt.figure(figsize=(7.5,7.5))

plt.subplots_adjust(left=0.2, bottom=0.1, right=0.96, top=0.92, wspace=0.1, hspace=0.25)
ax = plt.subplot2grid((1, 3), (0, 0),rowspan=2)
ax1 = plt.subplot2grid((1, 3), (0, 1),rowspan=2,colspan=2)

ii = list(np.argsort(np.abs(coefficients))[-10:][::-1])

for k in range(len(ii)):
    if(coefficients[ii[k]]>0):
        ax.barh([len(ii)-k-1],[np.abs(coefficients[ii[k]])],color=cmap1[0],alpha=0.5)
    else:
        ax.barh([len(ii)-k-1],[np.abs(coefficients[ii[k]])],color=cmap1[-1],alpha=0.5)
uu= ax.set_yticks(range(len(ii)))
uu = ax.set_yticklabels([new_labels[j] for j in ii],rotation=0,fontsize=9.)
ax.set_xlabel("coefficient ")

#ax.set_yticks([0,10,20])

myd1 = []
myl1 = []
kk = []
for q in ii:
    print("dist %s %8.4f %8.4f %8.4f  "% (new_labels[q],averages_MD[q],averagesA[q],averagesB[q] ))
    #if(np.abs(averagesA[q]-averagesB[q])>0.0):
    myd1.append([averagesA[q],averagesB[q], averages_MD[q]])
    myl1.append(new_labels[q])
    kk.append(q)
myx1 = range(len(myl1))
myd1 = np.array(myd1)


ax1.scatter(myd1[:,0],myx1,color='k',label="state A",marker='o',s=40)
ax1.scatter(myd1[:,1],myx1,color='k',label="state B",marker='*',s=50)
ax1.scatter(myd1[:,2],myx1,color='0.5',label="MD+eNOE",marker='+',s=120)
#ax1.scatter(averages_2koc[kk],myx1,color=cp[0],label="2koc",marker='s',s=25)
#ax1.scatter(averages_6by5[kk],myx1,color=cp[1],label="6by5",marker='s',s=25)

#ax1.set_xticks(range(len(myl1)))
#ax1.set_ylim(0.3,1.1)
#ax1.set_yticks([0.4,0.6,0.8,1.0])
#ax1.set_xticklabels([myl1[h] for h in range(len(myl1))],rotation=0,fontsize=8)
ax1.legend(ncol=4,fontsize=9,loc="upper center",bbox_to_anchor=(0.46,1.07))

ax1.set_yticks([])
#ax1.yaxis.set_label_position("right")
ax1.set_xlabel("calculated NOE  (nm)")
plt.savefig("figures/distance_hlda.pdf")
plt.savefig("figures/distance_hlda.png",dpi=600)

plt.show()
```

```
dist C8-2HO',G10-1H5'   0.3855   1.0434   0.2800  
dist C8-2HO',G10-H4'   0.5359   0.9745   0.3906  
dist C8-H41,G10-2HO'   0.5131   1.1995   0.3728  
dist C8-H42,G10-2HO'   0.5702   1.1965   0.4146  
dist C8-1H2',G10-H8   0.5725   0.9312   0.4190  
dist C8-2HO',G10-H1'   0.5623   0.8421   0.4137  
dist C8-2HO',G10-H8   0.4369   0.8220   0.3182  
dist U7-H1',G9-H22   0.5645   0.9829   0.4120  
dist C8-1H2',G9-H4'   0.5557   0.7806   0.4118  
dist C8-1H2',G10-H1'   0.5551   0.8508   0.4076
```

Note that all these contacts are between C8 and G10!! We then asked Beat and Nichols to check in their spectra for the presence of such peaks. Some of them were identified, which we plot below.

In [27]:

```
#C8_H5;G10_H1'  5.8 0.58 LOWER
#C8_H5;G10_H8	5.87	0.587  LOWER
#C8_H2';G10_H8	7.06	0.706 LOWER
#C8_H3';G9_H8	5.78	0.578 LOWER
#C8_H2';G9_H8	5.92	0.592 LOWER
#C8_H1';G10_H4'	5.28	0.528 LOWER
#C8_H2';G9_H2' 4.62	0.462 LOWER
# 8    RCYT H1'  10   RGUA H8
#8    RCYT H1'  10   RGUA H4'     


import mdtraj as md
import numpy as np

def find(pdb,at1,at2):
    t = md.load(pdb).topology.atoms

    p1 = None
    p2 = None
    for i,at in enumerate(t):
        if(str(at)==at1): p1 = i
        if(str(at)==at2): p2 = i
    if(p1==None or p2==None):
        print("# Atoms not found %s %s %s %s" % (at1,str(p1),at2,str(p2)))
    return p1,p2


pdb = "data/PDB/2koc_gmx.pdb"
a1, a2 = find(pdb,"C8-H5","G10-H1'")
b1, b2 = find(pdb,"C8-H5","G10-H8")
c1, c2 = find(pdb,"C8-1H2'","G10-H8")
d1, d2 = find(pdb,"C8-H3'","G9-H8")
e1, e2 = find(pdb,"C8-1H2'","G9-H8")
f1, f2 = find(pdb,"C8-H1'","G10-H4'")
g1, g2 = find(pdb,"C8-1H2'","G9-1H2'")
h1, h2 = find(pdb,"C8-H1'","G10-H8")
j1, j2 = find(pdb,"G9-H1'","G10-H8")

k1, k2 = find(pdb,"U6-2HO'","G9-H1")
l1, l2 = find(pdb,"U6-1H2'","G9-H1")
m1, m2 = find(pdb,"C8-H4'","G10-H8")
#f1, f2 = find(pdb,"U6-H6","G9-H1")
#g1, g2 = find(pdb,"U6-H1'","G9-H1")

#exp = [[6.3,0.63],[5.77,0.577],[5.8,0.58]]
exp = [[5.8],[5.87],[7.06],[5.78],[5.92],[5.28],[4.62],[5],[5],[5.77],[3.13],[3.55],[5.26]]


pairs = [[a1,a2],[b1,b2],[c1,c2],[d1,d2],[e1,e2],[f1,f2],[g1,g2],[h1,h2],[j1,j2],[m1,m2],[k1,k2],[l1,l2]]

traj_md = md.load("data/traj_temp_f_0.xtc",top=pdb)
dd = 10.*md.compute_distances(traj_md,pairs)
dd6 =np.power(dd,-6.)
theta=30
blocks=4
bsize = int(dd.shape[0]/blocks)
avgs = []
i1 = 7
i2= 8
for j in range(blocks):
    
    st = j*bsize
    end = st + bsize
    ww = np.array([float(line.split()[1]) for line in open("data/weights_set_A1_t_%d_rep_%d.dat" % (theta,j))])
    avg = np.power(np.average(dd6[st:end],weights=ww,axis=0),-1./6.)
    spec = (dd6[st:end,i1].reshape(end-st,1)+dd6[st:end,i2].reshape(end-st,1))
    spec = np.power(np.average(spec,axis=0,weights=ww),-1./6.)
    avg = np.concatenate((avg,spec))
    avgs.append(avg)

final_avg = np.average(avgs,axis=0)
final_std = np.std(avgs,axis=0,ddof=1)/np.sqrt(blocks)
```

In [28]:

```
pdb1 = "data/PDB/2koc.pdb"

a1, a2 = find(pdb1,"C8-H5","G10-H1'")
b1, b2 = find(pdb1,"C8-H5","G10-H8")
c1, c2 = find(pdb1,"C8-H2'","G10-H8")

d1, d2 = find(pdb1,"C8-H3'","G9-H8")
e1, e2 = find(pdb1,"C8-H2'","G9-H8")

f1, f2 = find(pdb1,"C8-H1'","G10-H4'")
g1, g2 = find(pdb1,"C8-H2'","G9-H2'")
h1, h2 = find(pdb1,"C8-H1'","G10-H8")
j1, j2 = find(pdb1,"G9-H1'","G10-H8")

k1, k2 = find(pdb1,"U6-HO2'","G9-H1")
l1, l2 = find(pdb1,"U6-H2'","G9-H1")
m1, m2 = find(pdb1,"C8-H4'","G10-H8")

#a1, a2 = find(pdb1,"C8-H2'","G10-H8")
#b1, b2 = find(pdb1,"C8-H4'","G10-H8")
#c1, c2 = find(pdb1,"C8-H5","G10-H1'")


#f1, f2 = find(pdb1,"U6-H6","G9-H1")
#g1, g2 = find(pdb1,"U6-H1'","G9-H1")

#pairs1 = [[a1,a2],[b1,b2],[c1,c2]]
pairs1 = [[a1,a2],[b1,b2],[c1,c2],[d1,d2],[e1,e2],[f1,f2],[g1,g2],[h1,h2],[j1,j2],[m1,m2],[k1,k2],[l1,l2]]

traj1 = md.load(pdb1,top=pdb1)
dd1 = 10.*md.compute_distances(traj1,pairs1)
dd16 =np.power(dd1,-6.)
avg1 = np.power(np.average(dd16,axis=0),-1./6.)
spec = (dd16[:,i1].reshape(dd16.shape[0],1)+dd16[:,i2].reshape(dd16.shape[0],1))
spec = np.power(np.average(spec,axis=0),-1./6.)
avg1 = np.concatenate((avg1,spec))

pdb2 = "data/PDB/6by5.pdb"
a1, a2 = find(pdb2,"C8-H5","G10-H1'")
b1, b2 = find(pdb2,"C8-H5","G10-H8")
c1, c2 = find(pdb2,"C8-H2'","G10-H8")
d1, d2 = find(pdb2,"C8-H3'","G9-H8")
e1, e2 = find(pdb2,"C8-H2'","G9-H8")
f1, f2 = find(pdb2,"C8-H1'","G10-H4'")
g1, g2 = find(pdb2,"C8-H2'","G9-H2'")
h1, h2 = find(pdb2,"C8-H1'","G10-H8")
j1, j2 = find(pdb2,"G9-H1'","G10-H8")
k1, k2 = find(pdb2,"U6-HO2'","G9-H1")
l1, l2 = find(pdb2,"U6-H2'","G9-H1")
m1, m2 = find(pdb2,"C8-H4'","G10-H8")

#pairs2 = [[a1,a2],[b1,b2],[c1,c2]]
pairs2 = [[a1,a2],[b1,b2],[c1,c2],[d1,d2],[e1,e2],[f1,f2],[g1,g2],[h1,h2],[j1,j2],[m1,m2],[k1,k2],[l1,l2]]
traj2 = md.load(pdb2,top=pdb2)
dd2 = 10.*md.compute_distances(traj2,pairs2)
dd26 =np.power(dd2,-6.)
avg2 = np.power(np.average(dd26,axis=0),-1./6.)
spec = (dd26[:,i1].reshape(dd26.shape[0],1)+dd26[:,i2].reshape(dd26.shape[0],1))
spec = np.power(np.average(spec,axis=0),-1./6.)
avg2 = np.concatenate((avg2,spec))
```

```
/anaconda3/lib/python3.7/site-packages/mdtraj/formats/pdb/pdbfile.py:198: UserWarning: Unlikely unit cell vectors detected in PDB file likely resulting from a dummy CRYST1 record. Discarding unit cell vectors.
  'cell vectors.', category=UserWarning)
/anaconda3/lib/python3.7/site-packages/mdtraj/formats/pdb/pdbfile.py:198: UserWarning: Unlikely unit cell vectors detected in PDB file likely resulting from a dummy CRYST1 record. Discarding unit cell vectors.
  'cell vectors.', category=UserWarning)
/anaconda3/lib/python3.7/site-packages/mdtraj/formats/pdb/pdbfile.py:198: UserWarning: Unlikely unit cell vectors detected in PDB file likely resulting from a dummy CRYST1 record. Discarding unit cell vectors.
  'cell vectors.', category=UserWarning)
/anaconda3/lib/python3.7/site-packages/mdtraj/formats/pdb/pdbfile.py:198: UserWarning: Unlikely unit cell vectors detected in PDB file likely resulting from a dummy CRYST1 record. Discarding unit cell vectors.
  'cell vectors.', category=UserWarning)
/anaconda3/lib/python3.7/site-packages/mdtraj/formats/pdb/pdbfile.py:198: UserWarning: Unlikely unit cell vectors detected in PDB file likely resulting from a dummy CRYST1 record. Discarding unit cell vectors.
  'cell vectors.', category=UserWarning)
/anaconda3/lib/python3.7/site-packages/mdtraj/formats/pdb/pdbfile.py:198: UserWarning: Unlikely unit cell vectors detected in PDB file likely resulting from a dummy CRYST1 record. Discarding unit cell vectors.
  'cell vectors.', category=UserWarning)
/anaconda3/lib/python3.7/site-packages/mdtraj/formats/pdb/pdbfile.py:198: UserWarning: Unlikely unit cell vectors detected in PDB file likely resulting from a dummy CRYST1 record. Discarding unit cell vectors.
  'cell vectors.', category=UserWarning)
/anaconda3/lib/python3.7/site-packages/mdtraj/formats/pdb/pdbfile.py:198: UserWarning: Unlikely unit cell vectors detected in PDB file likely resulting from a dummy CRYST1 record. Discarding unit cell vectors.
  'cell vectors.', category=UserWarning)
/anaconda3/lib/python3.7/site-packages/mdtraj/formats/pdb/pdbfile.py:198: UserWarning: Unlikely unit cell vectors detected in PDB file likely resulting from a dummy CRYST1 record. Discarding unit cell vectors.
  'cell vectors.', category=UserWarning)
/anaconda3/lib/python3.7/site-packages/mdtraj/formats/pdb/pdbfile.py:198: UserWarning: Unlikely unit cell vectors detected in PDB file likely resulting from a dummy CRYST1 record. Discarding unit cell vectors.
  'cell vectors.', category=UserWarning)
/anaconda3/lib/python3.7/site-packages/mdtraj/formats/pdb/pdbfile.py:198: UserWarning: Unlikely unit cell vectors detected in PDB file likely resulting from a dummy CRYST1 record. Discarding unit cell vectors.
  'cell vectors.', category=UserWarning)
/anaconda3/lib/python3.7/site-packages/mdtraj/formats/pdb/pdbfile.py:198: UserWarning: Unlikely unit cell vectors detected in PDB file likely resulting from a dummy CRYST1 record. Discarding unit cell vectors.
  'cell vectors.', category=UserWarning)
/anaconda3/lib/python3.7/site-packages/mdtraj/formats/pdb/pdbfile.py:198: UserWarning: Unlikely unit cell vectors detected in PDB file likely resulting from a dummy CRYST1 record. Discarding unit cell vectors.
  'cell vectors.', category=UserWarning)
/anaconda3/lib/python3.7/site-packages/mdtraj/core/trajectory.py:419: UserWarning: top= kwarg ignored since file contains topology information
  warnings.warn('top= kwarg ignored since file contains topology information')
/anaconda3/lib/python3.7/site-packages/mdtraj/formats/pdb/pdbfile.py:198: UserWarning: Unlikely unit cell vectors detected in PDB file likely resulting from a dummy CRYST1 record. Discarding unit cell vectors.
  'cell vectors.', category=UserWarning)
/anaconda3/lib/python3.7/site-packages/mdtraj/formats/pdb/pdbfile.py:198: UserWarning: Unlikely unit cell vectors detected in PDB file likely resulting from a dummy CRYST1 record. Discarding unit cell vectors.
  'cell vectors.', category=UserWarning)
/anaconda3/lib/python3.7/site-packages/mdtraj/formats/pdb/pdbfile.py:198: UserWarning: Unlikely unit cell vectors detected in PDB file likely resulting from a dummy CRYST1 record. Discarding unit cell vectors.
  'cell vectors.', category=UserWarning)
/anaconda3/lib/python3.7/site-packages/mdtraj/formats/pdb/pdbfile.py:198: UserWarning: Unlikely unit cell vectors detected in PDB file likely resulting from a dummy CRYST1 record. Discarding unit cell vectors.
  'cell vectors.', category=UserWarning)
/anaconda3/lib/python3.7/site-packages/mdtraj/formats/pdb/pdbfile.py:198: UserWarning: Unlikely unit cell vectors detected in PDB file likely resulting from a dummy CRYST1 record. Discarding unit cell vectors.
  'cell vectors.', category=UserWarning)
/anaconda3/lib/python3.7/site-packages/mdtraj/formats/pdb/pdbfile.py:198: UserWarning: Unlikely unit cell vectors detected in PDB file likely resulting from a dummy CRYST1 record. Discarding unit cell vectors.
  'cell vectors.', category=UserWarning)
/anaconda3/lib/python3.7/site-packages/mdtraj/formats/pdb/pdbfile.py:198: UserWarning: Unlikely unit cell vectors detected in PDB file likely resulting from a dummy CRYST1 record. Discarding unit cell vectors.
  'cell vectors.', category=UserWarning)
/anaconda3/lib/python3.7/site-packages/mdtraj/formats/pdb/pdbfile.py:198: UserWarning: Unlikely unit cell vectors detected in PDB file likely resulting from a dummy CRYST1 record. Discarding unit cell vectors.
  'cell vectors.', category=UserWarning)
/anaconda3/lib/python3.7/site-packages/mdtraj/formats/pdb/pdbfile.py:198: UserWarning: Unlikely unit cell vectors detected in PDB file likely resulting from a dummy CRYST1 record. Discarding unit cell vectors.
  'cell vectors.', category=UserWarning)
/anaconda3/lib/python3.7/site-packages/mdtraj/formats/pdb/pdbfile.py:198: UserWarning: Unlikely unit cell vectors detected in PDB file likely resulting from a dummy CRYST1 record. Discarding unit cell vectors.
  'cell vectors.', category=UserWarning)
/anaconda3/lib/python3.7/site-packages/mdtraj/formats/pdb/pdbfile.py:198: UserWarning: Unlikely unit cell vectors detected in PDB file likely resulting from a dummy CRYST1 record. Discarding unit cell vectors.
  'cell vectors.', category=UserWarning)
/anaconda3/lib/python3.7/site-packages/mdtraj/formats/pdb/pdbfile.py:198: UserWarning: Unlikely unit cell vectors detected in PDB file likely resulting from a dummy CRYST1 record. Discarding unit cell vectors.
  'cell vectors.', category=UserWarning)
/anaconda3/lib/python3.7/site-packages/mdtraj/formats/pdb/pdbfile.py:198: UserWarning: Unlikely unit cell vectors detected in PDB file likely resulting from a dummy CRYST1 record. Discarding unit cell vectors.
  'cell vectors.', category=UserWarning)
/anaconda3/lib/python3.7/site-packages/mdtraj/formats/pdb/pdbfile.py:198: UserWarning: Unlikely unit cell vectors detected in PDB file likely resulting from a dummy CRYST1 record. Discarding unit cell vectors.
  'cell vectors.', category=UserWarning)
/anaconda3/lib/python3.7/site-packages/mdtraj/core/trajectory.py:419: UserWarning: top= kwarg ignored since file contains topology information
  warnings.warn('top= kwarg ignored since file contains topology information')
/anaconda3/lib/python3.7/site-packages/mdtraj/formats/pdb/pdbfile.py:198: UserWarning: Unlikely unit cell vectors detected in PDB file likely resulting from a dummy CRYST1 record. Discarding unit cell vectors.
  'cell vectors.', category=UserWarning)
```

In [29]:

```
import matplotlib.pyplot as plt
import seaborn as sns

cols = ["pumpkin","medium blue","cement","leaf","black","white"]
cols = ["black","medium blue","pumpkin","brick red","black","cement"]

cp = sns.xkcd_palette(cols)

fig,ax =plt.subplots(1,1,figsize=(3.5,3.5))
plt.subplots_adjust(left=0.25, bottom=0.12, right=0.9, top=0.85, wspace=0.1, hspace=0.25)


ax.errorbar(final_avg[0],1,xerr=final_std[0],color=cp[1],marker='o',markersize=8,label="MD + eNOE")
ax.errorbar(final_avg[1],0.5,xerr=final_std[1],color=cp[1],marker='o',markersize=8)
ax.errorbar(final_avg[2],0,xerr=final_std[2],color=cp[1],marker='o',markersize=8)

#ax.barh(1,exp[0][1]-exp[0][0],left=exp[0][0],height=0.15,edgecolor=cp[-2],zorder=0,color=cp[-1],alpha=0.5,label="exp")
#ax.barh(0.5,exp[1][1]-exp[1][0],left=exp[1][0],height=0.15,edgecolor=cp[-2],zorder=0,color=cp[-1],alpha=0.5)
#ax.barh(0,exp[2][0],height=0.15,edgecolor=cp[-2],zorder=0,color=cp[-1],alpha=0.75,hatch="//")
#ax.barh(0,exp[2][1]-exp[2][0],left=exp[2][0],height=0.15,edgecolor=cp[-2],zorder=0,color=cp[-1],alpha=0.5)


ax.scatter(avg1[0],1,color=cp[2],marker='o',s=40,label="2KOC")
ax.scatter(avg1[1],0.5,color=cp[2],marker='o',s=40)
ax.scatter(avg1[2],0,color=cp[2],marker='o',s=40)

ax.scatter(avg2[0],1,color=cp[3],marker='o',s=40,label="6BY5")
ax.scatter(avg2[1],0.5,color=cp[3],marker='o',s=40,zorder=10)
ax.scatter(avg2[2],0,color=cp[3],marker='o',s=40)

for q in range(3,5):
    dx = -0.5*q+1
    ax.errorbar(final_avg[q],dx,xerr=final_std[q],color=cp[1],marker='o',markersize=8)
    #ax.barh(dx,exp[q][1]-exp[q][0],left=exp[q][0],height=0.15,edgecolor=cp[-2],zorder=0,color=cp[-1],alpha=0.5)
    ax.scatter(avg1[q],dx,color=cp[2],marker='o',s=40)
    ax.scatter(avg2[q],dx,color=cp[3],marker='o',s=40)


ax.legend(ncol=2,fontsize=9,loc="upper center",bbox_to_anchor=(0.44,1.2))


ax.set_yticks([-1,-0.5,0,0.5,1])
ax.set_yticklabels(["U6 H2'\nG9-H1", "U6 HO2'\nG9-H1","C8 H5\n G10 H1'","C8 H4'\nG10 H8","C8 H2'\nG10 H8",])
ax.set_xlim(2.,10)
#ax1.yaxis.set_label_position("right")
ax.set_xlabel("NOE  ($\AA$)")
ax.spines['right'].set_visible(False)
ax.spines['top'].set_visible(False)
plt.savefig("figures/distance_hlda_noe1.pdf")
plt.savefig("figures/distance_hlda_noe1.png",dpi=600)

plt.show()
```

In [30]:

```
import matplotlib.pyplot as plt
import seaborn as sns

cols = ["pumpkin","medium blue","cement","leaf","black","white"]
cols = ["black","medium blue","pumpkin","brick red","black","cement"]

cp = sns.xkcd_palette(cols)

fig,ax =plt.subplots(1,1,figsize=(4,4))
plt.subplots_adjust(left=0.25, bottom=0.12, right=0.9, top=0.85, wspace=0.1, hspace=0.25)
pp = ["8 H5\n10 H1'","8 H5\n10 H8","8 H2'\n10 H8",\
    "8 H3'\n9 H8","8 H2'\n9 H8","8 H1'\n10 H4'",\
    "8 H2'\n9 H2'","8 H1'\n10 H8","9 H1'\n10 H8","8 H4'\n10 H8","6 HO2'\n9 H1","6 H2'\n9-H1","8/9 H1'\n10 H8"]

k = 8
oks = [0,1,2,9,10,12]
lab = []
ax.set_yticks(np.arange(k)[::-1]+1)
for i,el in enumerate(pp):
    
    print(el,final_avg[i],exp[i][0],avg1[i],avg2[i])
    if(i in oks):
        if(i==0):        
            ax.errorbar(final_avg[i],k,xerr=final_std[i],color=cp[1],marker='o',markersize=8,label="MD + eNOE")
            ax.scatter(avg1[i],k,color=cp[2],marker='o',s=40,label="2KOC")
            ax.scatter(avg2[i],k,color=cp[3],marker='o',s=40,label="6BY5")
        else:
            ax.errorbar(final_avg[i],k,xerr=final_std[i],color=cp[1],marker='o',markersize=8)
            ax.scatter(avg1[i],k,color=cp[2],marker='o',s=40)
            ax.scatter(avg2[i],k,color=cp[3],marker='o',s=40)

        if(i>=6):
            ax.barh(k,0.2*exp[i][0],left=0.9*exp[i][0],height=0.4,edgecolor=cp[-2],zorder=0,color=cp[-1],alpha=0.5)
        else:
            ax.barh(k,12,left=0.9*exp[i][0],height=0.4,edgecolor=cp[-2],zorder=0,color=cp[-1],alpha=0.75)
        k -= 1
        lab.append(pp[i])
ax.set_xlim(2.5,11)
ax.set_yticklabels(lab)
ax.set_xlabel("NOE  ($\AA$)")
ax.spines['right'].set_visible(False)
ax.spines['top'].set_visible(False)
plt.savefig("figures/distance_hlda_noe2.pdf")
plt.savefig("figures/distance_hlda_noe2.png",dpi=600)
plt.show()
plt.close()
```

```
8 H5
10 H1' 5.891326235383453 5.8 8.962134 9.08019
8 H5
10 H8 9.001857898080367 5.87 10.58514 9.555871
8 H2'
10 H8 6.461807584427399 7.06 9.757728 8.258541
8 H3'
9 H8 6.088436238643737 5.78 6.9300303 7.5827227
8 H2'
9 H8 6.183709391505947 5.92 7.792676 8.616932
8 H1'
10 H4' 6.806701444459584 5.28 9.289813 8.394944
8 H2'
9 H2' 6.8858702879154166 4.62 8.561864 8.288545
8 H1'
10 H8 5.745333593353949 5 7.0618997 5.183244
9 H1'
10 H8 6.564226622975962 5 6.6583 7.0731764
8 H4'
10 H8 6.173093999603631 5.77 6.892979 6.3549776
6 HO2'
9 H1 3.377016540132443 3.13 3.0758977 3.1230862
6 H2'
9-H1 3.8617091470587637 3.55 3.5278594 3.548772
8/9 H1'
10 H8 5.238084208252596 5.26 6.093243 5.060349
```

In [ ]:
