## Supplementary material for "Integrating NMR and Simulations Reveals Motions in the UUCG Tetraloop": Figure 4 (from another version of this preprint: version 2, 2019-09-11)

figure4


### Figure 4 - HLDA on torsion angles¶

In this notebook we perform the HLDA analysis on the torsion angles. We proceed in the following way:

1. We create two sub-trajectories stateA.xtc and stateB.xtc. Frames in stateA have eRMSD from native <0.7, while 0.7<eRMSD <1.4 in stateB.
2. We read the weights of the frames from the BME optimization
3. We calculate all torsion angles on the entire trajectory using barnaba.
4. We calculate for each torsion the quantity alphabeta=0.5\*(1. + cos(theta-theta\_2koc)). This is useful as it takes care of periodicity.
5. We perform HLDA using two classes and alphabeta as descriptors. The script to run HLDA can be found here
6. Finally, we plot the eigenvector coefficients.

In [5]:

```
import pickle
import numpy as np
import mdtraj as md

# read ermsd from pickle
fname = "data/observables/ermsd_full.p"
ermsd_full = pickle.load(open(fname,'rb'), encoding='bytes')

# create two trajectories corresponding to the two states
top = "data/PDB/2koc_gmx.pdb"
trj = "data/traj_temp_f_0.xtc"
trajmd = md.load(trj,top=top)
stateA = np.where(ermsd_full<0.7)[0]
stateB = np.where((ermsd_full>=0.7) & (ermsd_full<1.4))[0]  
trajmd[stateA].save("data/stateA.xtc")
trajmd[stateB].save("data/stateB.xtc")
```

In [6]:

```
# read weights 
theta=30
dd = np.array([[float(x) for x in line.split()] for line in open("data/weights_set_A1_t_%d.dat" % (theta))])
weights_0 = dd[:,0]
weights_1 = dd[:,1]

weights_stateA = weights_1[stateA]
weights_stateB = weights_1[stateB]
```

In [7]:

```
import barnaba as bb
from barnaba import definitions

# calculate all backbone angles using barnaba
anglesA,res = bb.backbone_angles("data/stateA.xtc",topology=top)
anglesB,res = bb.backbone_angles("data/stateB.xtc",topology=top)
fname = "data/observables/angles_2koc.p"
[angles_2koc,res_2koc] = pickle.load(open(fname,'rb'), encoding='bytes')
```

```
# Loading data/stateA.xtc 
# Loading data/stateB.xtc
```

In [8]:

```
# now we remove nans (at the termini) at get labels that will be useful later for plotting 
la = definitions.bb_angles
data_angles_A = []
data_angles_B = []
data_angles_2koc = []
labels = []
for i in range(anglesA.shape[1]):
    for j in range(anglesA.shape[2]):
        if(np.isfinite(anglesA[0,i,j])):
            #ddiff1 = adiff(anglesA[:,i,j],angles_2koc[0,i,j])
            #ddiff2 = adiff(anglesB[:,i,j],angles_2koc[0,i,j])
            data_angles_A.append(anglesA[:,i,j])
            data_angles_B.append(anglesB[:,i,j])
            data_angles_2koc.append(angles_2koc[0,i,j])
            labels.append("%d_%s" % (i+1,la[j]))

data_angles_A = np.array(data_angles_A)
data_angles_B = np.array(data_angles_B)
data_angles_2koc = np.array(data_angles_2koc)

# calculate features as the alphabeta in PLUMED
ddiffA = data_angles_A-data_angles_2koc[:,np.newaxis]
ddiffB = data_angles_B-data_angles_2koc[:,np.newaxis]
data_A =0.5*(1. + np.cos(np.arctan2(np.sin(ddiffA), np.cos(ddiffA)))).T
data_B =0.5*(1. + np.cos(np.arctan2(np.sin(ddiffB), np.cos(ddiffB)))).T
```

In [9]:

```
# Do HLDA using my script

import sys
sys.path.append("/Users/sandrobottaro/Software/HLDA")
import HLDA as hlda

# initialize class
hlda_angles = hlda.HLDA()

# add data
n = hlda_angles.load_data(data_A,oweights=weights_stateA)
n = hlda_angles.load_data(data_B,oweights=weights_stateB)

# run the algorithm
hlda_angles.run_lda(harmonic=True)

# get the coefficients of the evigenvector corresponding to the largest eigenvalue
eigenvalues = hlda_angles.get_evals()
coefficients = hlda_angles.get_evecs()[:,-1]
```

```
# initializing HLDA class.
# Processing data with 25124 samples and 94 features 
# Processing data with 43169 samples and 94 features
```

In [10]:

```
import matplotlib.pyplot as plt
import seaborn as sns

fig,ax = plt.subplots(1,1,figsize=(18,5))
plt.scatter(range(len(eigenvalues)),eigenvalues)
plt.xlabel("eigenvalue index")
plt.ylabel("eigenvalue")
plt.show()


fig,ax = plt.subplots(1,1,figsize=(18,5))
plt.scatter(range(len(eigenvalues)),coefficients)
plt.xlabel("eigenvalue index")
plt.ylabel("eigenvalue")
plt.show()
```

```
<Figure size 1800x500 with 1 Axes>
```

```
<Figure size 1800x500 with 1 Axes>
```

In [11]:

```
# here we plot the coefficients of the non-zero eigenvector

# We need to make some adjustmenst to the array to make the plot nice,otherwise it would be simple.
# reshape the array to be n_residues (14) by n_angles (7)

reshaped = np.zeros((14,7))
for j in range(len(coefficients)):
    i = int((j+2)%7)
    k = int((j+2)/7)
    reshaped[k,i]= coefficients[j]
# set manually alpha, beta in G1 and epsilon, zeta in C14 to nan
reshaped[0,0] = np.nan
reshaped[0,1] = np.nan
reshaped[-1,-2] = np.nan
reshaped[-1,-3] = np.nan

# define the figure
fig = plt.figure(figsize=(7.5,3.5))
ax = plt.subplot2grid((1, 3), (0, 0), colspan=2)
plt.subplots_adjust(left=0.05, bottom=0.12, right=0.93, top=0.98, wspace=0.07, hspace=0.2)
# define labels
angs= [r"$\alpha$",r"$\beta$",r"$\gamma$",r"$\delta$",r"$\epsilon$",r"$\zeta$",r"$\chi$"]
ress = ['G1', 'G2', 'C3', 'A4', 'C5', 'U6', 'U7', 'C8', 'G9', 'G10', 'U11', 'G12', 'C13','C14']
# set colormap
cmap = sns.diverging_palette(275, 150, s=80, l=55, n=7,as_cmap=True)
# draw heatmap
ax = sns.heatmap(reshaped.T/-1.,center=0,ax=ax,linewidths=.35,\
                yticklabels=angs,xticklabels=ress,cmap=cmap,cbar_kws={"pad":0.03},snap=True)
# set ylabels after so that we can rotate them
ax.set_yticklabels(ax.get_yticklabels(), rotation = 0, fontsize = 12)
ax.set_ylabel("dihedral angle")
ax.set_xlabel("Residue")

plt.show()
```

We plot on the side the distribution of the degree of freedom with the largest coefficient. According to HLDA, this is the degree of freedom that contributes the most in discriminating state A from state B.

In [12]:

```
# find index with largest eigenvalue
imax = np.argsort(np.abs(coefficients))[-1]
print("The angle with the largest coefficent is %s (%8.4f)" % (labels[imax],coefficients[imax]))
kmax = int(labels[imax].split("_")[0])-1
jmax = la.index(labels[imax].split("_")[1])

# get angles from pickles
fname = "data/observables/angles.p"
[angles,res] = pickle.load(open(fname,'rb'), encoding='bytes')


fname = "data/observables/angles_6by5.p"
[angles_6by5,res_6by5] = pickle.load(open(fname,'rb'), encoding='bytes')

angles_2koc[np.where(angles_2koc<0.0)] += 2.*np.pi
angles_2koc *= (180./np.pi)

angles_6by5[np.where(angles_6by5<0.0)] += 2.*np.pi
angles_6by5 *= (180./np.pi)
```

```
The angle with the largest coefficent is 8_zeta (-61.6249)
```

```
/anaconda3/lib/python3.7/site-packages/ipykernel_launcher.py:15: RuntimeWarning: invalid value encountered in less
  from ipykernel import kernelapp as app
/anaconda3/lib/python3.7/site-packages/ipykernel_launcher.py:18: RuntimeWarning: invalid value encountered in less
```

In [14]:

```
from scipy import stats
cols = ["black","medium blue","pumpkin","brick red","black","white"]
cp = sns.xkcd_palette(cols)

# plot again the heatmap
fig = plt.figure(figsize=(7.5,3.5))
ax = plt.subplot2grid((2, 3), (0, 0), colspan=2,rowspan=2)
plt.subplots_adjust(left=0.07, bottom=0.125, right=0.91, top=0.98, wspace=0.01, hspace=0.01)
# draw heatmap
ax = sns.heatmap(-1.*reshaped.T,center=0,ax=ax,linewidths=.35,\
                yticklabels=angs,xticklabels=ress,cmap=cmap,cbar_kws={"pad":0.03},snap=True)
# set ylabels after so that we can rotate them
ax.set_yticklabels(ax.get_yticklabels(), rotation = 0, fontsize = 12)
ax.set_ylabel("dihedral angle")
ax.set_xlabel("Residue")

# now plot the histogram
ax1 = plt.subplot2grid((2, 3), (0, 2),rowspan=2)
def histo(data,_weights,bins):
    
    data[np.where(data<0.0)] += 2.*np.pi
    data *= (180/np.pi)
    data = data[np.isfinite(data)]
    f = 2./data.std()
    kernel = stats.gaussian_kde(data,weights=_weights,bw_method=f)
    hh = kernel(bins)
    return hh

bins = np.linspace(0,360,200)
yC = histo(np.copy(angles[:,kmax,jmax]),weights_1,bins)
ax1.fill(bins,yC,c=cp[1],ls="-",label="MD+eNOE",alpha=0.5)


ax1.set_yticks([0,0.02,0.04])
ax1.set_ylim(-0.006,0.05)
ax1.set_xticks([0,120,240,360])
#ax1.set_xticklabels([])
ax1.yaxis.tick_right()
ax1.scatter(angles_2koc[:,kmax,jmax],[-0.004]*len(angles_2koc),color=cp[2],label="2KOC",marker="o",s=8)
ax1.scatter(angles_6by5[:,kmax,jmax],[-0.002]*len(angles_6by5),color=cp[3],label="6BY5",marker="o",s=8)
ax1.set_xlabel("C8($\zeta$) [deg]")
ax1.yaxis.set_label_position("right")
ax1.set_ylabel("Probability density")
ax1.legend(ncol=2,fontsize=8)


ax.text(-1.8,0.6,"a",fontsize=13,weight = 'bold')
ax1.text(-53,0.047,"b",fontsize=13,weight = 'bold')
ax1.text(85,0.001,"g+",fontsize=9,ha='center')
ax1.text(290,0.001,"g-",fontsize=9,ha='center')

plt.savefig("figures/angles_hlda.pdf")
plt.savefig("figures/angles_hlda.png",dpi=600)
plt.show()
```

In [ ]:
