## Supplementary material for "Integrating NMR and Simulations Reveals Motions in the UUCG Tetraloop": Figure 3C (from another version of this preprint: version 2, 2019-09-11)

figure3C


### Plot free energy surfaces¶

In this notebook we plot the free energy surfaces onto the eRMSD from native versus C6-G9 distance, using all eNOE, INCLUDING the new data.

1. We calculate different quantities from trajectories. This is a rather dull excercise, that we did in a separate notebook. Here, we just read the data from pickle files
2. We do a 2-dimensional histogram using the distance between C6 and G9 and the loop RMSD.
3. Identify on the FES the regions corresponding to state A and state B.
4. Draw 100 samples from the trajectory with the optmimized weights. These samples will be used to make the figure with the 3D structures
5. Calculate population and free energy difference between state A and state B.
6. Overlay on the FES all NUUCGN structures from the PDB where NN form a watson-crick base-pair and add them to the projection.

In [9]:

```
import pickle
import numpy as np

# ERMSD - stem loop (residues 5-10)
fname = "data/observables/ermsd_loop_2koc.p"
ermsd_loop_2koc = pickle.load(open(fname,'rb'), encoding='bytes')

fname = "data/observables/ermsd_loop_6by5.p"
ermsd_loop_6by5 = pickle.load(open(fname,'rb'), encoding='bytes')

fname = "data/observables/ermsd_loop.p"
ermsd_loop = pickle.load(open(fname,'rb'), encoding='bytes')

# vectors connecting center of nucleobases
fname = "data/observables/rvecs_2koc.p"
[rvecs_2koc,res_2koc] = pickle.load(open(fname,'rb'), encoding='bytes')

fname = "data/observables/rvecs_6by5.p"
[rvecs_6by5,res_6by5] = pickle.load(open(fname,'rb'), encoding='bytes')

fname = "data/observables/rvecs.p"
[rvecs,res] = pickle.load(open(fname,'rb'), encoding='bytes')

d_69 = np.sqrt(np.sum(rvecs[:,5,8,:]**2,axis=1))
d_69_2koc = np.sqrt(np.sum(rvecs_2koc[:,5,8,:]**2,axis=1))
d_69_6by5 = np.sqrt(np.sum(rvecs_6by5[:,5,8,:]**2,axis=1))
```

1. Make the histogram (KDE takes time). Note that the weighted KDE is only available with a new scipy version.

In [10]:

```
from scipy import stats

theta = 30
kbt=0.008314462*280
# AAAA USING DATA with NEW ENOE ####
data_tmp = np.array([[float(x) for x in line.split()] for line in open("data/weights_set_A1_t_%d.dat" % (theta),"r")])
weights0f = data_tmp[:,0]
weights1f = data_tmp[:,1]

datax = ermsd_loop
datay = d_69
xa=np.arange(1.05*np.min(datax),1.05*np.max(datax),0.025)
ya=np.arange(1.05*np.min(datay),1.05*np.max(datay),0.025)

kernel = stats.gaussian_kde(np.array([datax,datay]),weights=weights1f,bw_method=0.15)

X, Y = np.meshgrid(xa,ya)
grid_coords = np.vstack([X.ravel(), Y.ravel()])
H = np.reshape(kernel(grid_coords).T, X.shape)

Z= -(kbt*np.log(H))
Z -= np.min(Z)
```

```
/anaconda3/lib/python3.7/site-packages/ipykernel_launcher.py:21: RuntimeWarning: divide by zero encountered in log
```

Plot the 2D fes. Note that we define by hand two rectangular regions that correspond to state A and state B.

In [11]:

```
import matplotlib.pyplot as plt
import matplotlib.patches as patches

import seaborn as sns 

levels = [0,2.5,5,7.5,10,12.5,15,17.5]
sns.set_palette(sns.color_palette("cubehelix", len(levels)-1))
cmap = sns.cubehelix_palette(len(levels)-1, as_cmap=True,reverse=True,rot=-0.2)

fig,ax = plt.subplots(1,1,figsize=(7.5, 6))


# define region A and region B (by hand)
region_A = [[0.05,0.3],[0.85,1]]
region_B = [[1.0,1.0],[1.8,2.15]]


x1 = np.where((xa<region_A[1][0]) & (xa > region_A[0][0]) )[0]
y1 = np.where((ya<region_A[1][1]) & (ya > region_A[0][1]) )[0]

x2 = np.where((xa<region_B[1][0]) & (xa > region_B[0][0]) )[0]
y2 = np.where((ya<region_B[1][1]) & (ya > region_B[0][1]) )[0]
rect = patches.Rectangle((region_A[0][0],region_A[0][1]),region_A[1][0]-region_A[0][0],region_A[1][1]-region_A[0][1],\
                         linewidth=0.5,edgecolor='k',facecolor='none')
ax.add_patch(rect)
rect = patches.Rectangle((region_B[0][0],region_B[0][1]),region_B[1][0]-region_B[0][0],region_B[1][1]-region_B[0][1],\
                         linewidth=0.5,edgecolor='k',facecolor='none')
ax.add_patch(rect)


plt.subplots_adjust(left=0.1, bottom=0.1, right=0.95, top=0.98, wspace=0.07, hspace=0.1)

plt.xlim(0,1.9)
plt.ylim(0,2.2)
CS = ax.contourf(X, Y, Z,levels=levels,cmap=cmap)
CB = fig.colorbar(CS)

#plt.scatter(ermsd_pdbs,dists1,s=1,c='0.2',label='PDB structures')
#plt.legend(ncol=1,loc="lower right")
plt.xlabel("eRMSD from native (residues 4-11)")
plt.ylabel("C6-G9 distance (nm)")
#plt.savefig("figures/2d_projection.png",dpi=600)
#plt.savefig("figures/2d_projection.pdf")
plt.show()
plt.close()
```

now we draw 100 samples from the trajectory with the desired weights. These samples will be used to make the figure with the 3D structures

In [12]:

```
import mdtraj as md
native = "data/PDB/2koc_gmx.pdb"

top = "data/PDB/2koc_gmx.pdb"
trj = "data/traj_temp_f_0.xtc"
trajmd = md.load(trj,top=top)
# choose 100 samples
ff = np.random.choice(len(weights1f),size=100,p=weights1f/np.sum(weights1f),replace=False)
stateAi = []
stateBi = []
for f in ff:
    ermsd = ermsd_loop[f]
    dista = d_69[f]
    if((ermsd<region_A[1][0]) and (ermsd > region_A[0][0])):
        if((dista<region_A[1][1]) and (dista > region_A[0][1])):
            stateAi.append(f)
    if((ermsd<region_B[1][0]) and (ermsd > region_B[0][0])):
        if((dista<region_B[1][1]) and (dista > region_B[0][1])):
            stateBi.append(f)

trajmd[stateAi].superpose(md.load(native)).save("data/samples_stateA.pdb")  
trajmd[stateBi].superpose(md.load(native)).save("data/samples_stateB.pdb")
trajmd[ff].superpose(md.load(native)).save("data/samples_final.pdb")
```

In [14]:

```
weights_A = []
for j in range(4):
    dd = np.array([[float(x) for x in line.split()] for line in open("data/weights_set_A1_t_%d_rep_%d.dat" % (theta,j))])
    weights_A.append(dd[:,1])
weights_A = np.array(weights_A)

pops = []
bsize = int(ermsd_loop.shape[0]/4)
for j in range(4):
    start = j*bsize
    end = start + bsize
    datax = ermsd_loop[start:end]
    datay = d_69[start:end]
    kernel = stats.gaussian_kde(np.array([datax,datay]),weights=weights_A[j],bw_method=0.1)
    X, Y = np.meshgrid(xa,ya)
    grid_coords = np.vstack([X.ravel(), Y.ravel()])
    H = np.reshape(kernel(grid_coords).T, X.shape)
    apop = (np.sum(H[x1,:][:,y1])/np.sum(H))
    bpop = (np.sum(H[x2,:][:,y2])/np.sum(H))
    pops.append([apop,bpop])
pops = np.array(pops)
```

In [15]:

```
print("STATE A population %5.1f +/- %5.1f" % \
      (100*np.average(pops,axis=0)[0],100*np.std(pops,axis=0,ddof=1)[0]/2.))
print("STATE B population %5.1f +/- %5.1f" % \
      (100*np.average(pops,axis=0)[1],100*np.std(pops,axis=0,ddof=1)[1]/2.))
dfs = [-kbt*(np.log(pops[i,0]) - np.log(pops[i,1])) for i in range(pops.shape[0])]
print("Delta F: %5.2f+/-%5.2f Kj/mol" % (np.average(dfs,axis=0),np.std(dfs,axis=0,ddof=1)))
```

```
STATE A population  84.0 +/-   7.1
STATE B population  11.1 +/-   6.1
Delta F: -5.71+/- 2.87 Kj/mol
```

To add some more info to the plot, we take all the CUUCGG structures from the PDB where CG form a watson-crick base-pair and add them to the projection.

In [16]:

```
import glob as glob
import barnaba as bb
ref = "data/PDB/CUUCGG/2KOC_5_C_A_0_00001.pdb"
ermsd_pdbs = []
dists = []
ll = glob.glob("data/PDB/CUUCGG/*.pdb")
for f in ll:
    #ermsd_pdbs.append(bb.ermsd(ref,f,residues_ref=[1,2,3,4,5,6],residues_target=[1,2,3,4,5,6]))
    ermsd_pdbs.append(bb.ermsd(ref,f))
    #Note that distances between C8 and G10 are smaller in state B compared to state A.
    rvec,res = bb.dump_rvec(f,cutoff=100)
    dists.append(rvec[0])
```

```
# Loaded reference data/PDB/CUUCGG/2KOC_5_C_A_0_00001.pdb 
# Loaded target data/PDB/CUUCGG/1EKZ_13_C_B_0_00006.pdb 
# Loading data/PDB/CUUCGG/1EKZ_13_C_B_0_00006.pdb 
# Loaded reference data/PDB/CUUCGG/2KOC_5_C_A_0_00001.pdb 
# Loaded target data/PDB/CUUCGG/2N6T_19_C_A_0_00009.pdb 
# Loading data/PDB/CUUCGG/2N6T_19_C_A_0_00009.pdb 
# Loaded reference data/PDB/CUUCGG/2KOC_5_C_A_0_00001.pdb 
# Loaded target data/PDB/CUUCGG/1M5L_17_C_A_0_00011.pdb 
# Loading data/PDB/CUUCGG/1M5L_17_C_A_0_00011.pdb 
# Loaded reference data/PDB/CUUCGG/2KOC_5_C_A_0_00001.pdb 
# Loaded target data/PDB/CUUCGG/1M5L_17_C_A_0_00005.pdb 
# Loading data/PDB/CUUCGG/1M5L_17_C_A_0_00005.pdb 
# Loaded reference data/PDB/CUUCGG/2KOC_5_C_A_0_00001.pdb 
# Loaded target data/PDB/CUUCGG/1EKZ_13_C_B_0_00012.pdb 
# Loading data/PDB/CUUCGG/1EKZ_13_C_B_0_00012.pdb 
# Loaded reference data/PDB/CUUCGG/2KOC_5_C_A_0_00001.pdb 
# Loaded target data/PDB/CUUCGG/4v9i-pdb-bundle1_419_C_A_0_00000.pdb 
# Loading data/PDB/CUUCGG/4v9i-pdb-bundle1_419_C_A_0_00000.pdb 
# Loaded reference data/PDB/CUUCGG/2KOC_5_C_A_0_00001.pdb 
# Loaded target data/PDB/CUUCGG/4v5d-pdb-bundle3_419_C_A_0_00000.pdb 
# Loading data/PDB/CUUCGG/4v5d-pdb-bundle3_419_C_A_0_00000.pdb 
# Loaded reference data/PDB/CUUCGG/2KOC_5_C_A_0_00001.pdb 
# Loaded target data/PDB/CUUCGG/1F7F_12_C_A_0_00001.pdb 
# Loading data/PDB/CUUCGG/1F7F_12_C_A_0_00001.pdb 
# Loaded reference data/PDB/CUUCGG/2KOC_5_C_A_0_00001.pdb 
# Loaded target data/PDB/CUUCGG/6c5l-pdb-bundle3_419_C_A_0_00000.pdb 
# Loading data/PDB/CUUCGG/6c5l-pdb-bundle3_419_C_A_0_00000.pdb 
# Loaded reference data/PDB/CUUCGG/2KOC_5_C_A_0_00001.pdb 
# Loaded target data/PDB/CUUCGG/6enf-pdb-bundle1_1449_C_a_0_00003.pdb 
# Loading data/PDB/CUUCGG/6enf-pdb-bundle1_1449_C_a_0_00003.pdb 
# Loaded reference data/PDB/CUUCGG/2KOC_5_C_A_0_00001.pdb 
# Loaded target data/PDB/CUUCGG/1EBQ_12_C_A_0_00002.pdb 
# Loading data/PDB/CUUCGG/1EBQ_12_C_A_0_00002.pdb 
# Loaded reference data/PDB/CUUCGG/2KOC_5_C_A_0_00001.pdb 
# Loaded target data/PDB/CUUCGG/1VQ9_1769_C_0_0_00000.pdb 
# Loading data/PDB/CUUCGG/1VQ9_1769_C_0_0_00000.pdb 
# Loaded reference data/PDB/CUUCGG/2KOC_5_C_A_0_00001.pdb 
# Loaded target data/PDB/CUUCGG/5mdw-pdb-bundle1_419_C_2_0_00002.pdb 
# Loading data/PDB/CUUCGG/5mdw-pdb-bundle1_419_C_2_0_00002.pdb 
# Loaded reference data/PDB/CUUCGG/2KOC_5_C_A_0_00001.pdb 
# Loaded target data/PDB/CUUCGG/1AUD_28_C_B_0_00027.pdb 
# Loading data/PDB/CUUCGG/1AUD_28_C_B_0_00027.pdb 
# Loaded reference data/PDB/CUUCGG/2KOC_5_C_A_0_00001.pdb 
# Loaded target data/PDB/CUUCGG/4v9l-pdb-bundle3_419_C_A_0_00000.pdb 
# Loading data/PDB/CUUCGG/4v9l-pdb-bundle3_419_C_A_0_00000.pdb 
# Loaded reference data/PDB/CUUCGG/2KOC_5_C_A_0_00001.pdb 
# Loaded target data/PDB/CUUCGG/4v5a-pdb-bundle1_419_C_A_0_00000.pdb 
# Loading data/PDB/CUUCGG/4v5a-pdb-bundle1_419_C_A_0_00000.pdb 
# Loaded reference data/PDB/CUUCGG/2KOC_5_C_A_0_00001.pdb 
# Loaded target data/PDB/CUUCGG/4wf1-pdb-bundle2_1691_C_A_0_00000.pdb 
# Loading data/PDB/CUUCGG/4wf1-pdb-bundle2_1691_C_A_0_00000.pdb 
# Loaded reference data/PDB/CUUCGG/2KOC_5_C_A_0_00001.pdb 
# Loaded target data/PDB/CUUCGG/4v63-pdb-bundle1_419_C_A_0_00000.pdb 
# Loading data/PDB/CUUCGG/4v63-pdb-bundle1_419_C_A_0_00000.pdb 
# Loaded reference data/PDB/CUUCGG/2KOC_5_C_A_0_00001.pdb 
# Loaded target data/PDB/CUUCGG/4v7u-pdb-bundle3_207_C_A_0_00000.pdb 
# Loading data/PDB/CUUCGG/4v7u-pdb-bundle3_207_C_A_0_00000.pdb 
# Loaded reference data/PDB/CUUCGG/2KOC_5_C_A_0_00001.pdb 
# Loaded target data/PDB/CUUCGG/1FYP_11_C_A_0_00027.pdb 
# Loading data/PDB/CUUCGG/1FYP_11_C_A_0_00027.pdb 
# Loaded reference data/PDB/CUUCGG/2KOC_5_C_A_0_00001.pdb 
# Loaded target data/PDB/CUUCGG/5hd1-pdb-bundle2_415_C_A_0_00000.pdb 
# Loading data/PDB/CUUCGG/5hd1-pdb-bundle2_415_C_A_0_00000.pdb 
# Loaded reference data/PDB/CUUCGG/2KOC_5_C_A_0_00001.pdb 
# Loaded target data/PDB/CUUCGG/4X64_419_C_A_0_00000.pdb 
# Loading data/PDB/CUUCGG/4X64_419_C_A_0_00000.pdb 
# Loaded reference data/PDB/CUUCGG/2KOC_5_C_A_0_00001.pdb 
# Loaded target data/PDB/CUUCGG/1M82_10_C_A_0_00023.pdb 
# Loading data/PDB/CUUCGG/1M82_10_C_A_0_00023.pdb 
# Loaded reference data/PDB/CUUCGG/2KOC_5_C_A_0_00001.pdb 
# Loaded target data/PDB/CUUCGG/5j8a-pdb-bundle3_1691_C_d_0_00000.pdb 
# Loading data/PDB/CUUCGG/5j8a-pdb-bundle3_1691_C_d_0_00000.pdb 
# Loaded reference data/PDB/CUUCGG/2KOC_5_C_A_0_00001.pdb 
# Loaded target data/PDB/CUUCGG/3CF5_1708_C_X_0_00000.pdb 
# Loading data/PDB/CUUCGG/3CF5_1708_C_X_0_00000.pdb 
# Loaded reference data/PDB/CUUCGG/2KOC_5_C_A_0_00001.pdb 
# Loaded target data/PDB/CUUCGG/4WF9_1735_C_X_0_00000.pdb 
# Loading data/PDB/CUUCGG/4WF9_1735_C_X_0_00000.pdb 
# Loaded reference data/PDB/CUUCGG/2KOC_5_C_A_0_00001.pdb 
# Loaded target data/PDB/CUUCGG/5tgm-pdb-bundle3_230_C_d_0_00000.pdb 
# Loading data/PDB/CUUCGG/5tgm-pdb-bundle3_230_C_d_0_00000.pdb 
# Loaded reference data/PDB/CUUCGG/2KOC_5_C_A_0_00001.pdb 
# Loaded target data/PDB/CUUCGG/4v9r-pdb-bundle4_1691_C_A_0_00000.pdb 
# Loading data/PDB/CUUCGG/4v9r-pdb-bundle4_1691_C_A_0_00000.pdb 
# Loaded reference data/PDB/CUUCGG/2KOC_5_C_A_0_00001.pdb 
# Loaded target data/PDB/CUUCGG/5IEM_25_C_A_0_00000.pdb 
# Loading data/PDB/CUUCGG/5IEM_25_C_A_0_00000.pdb 
# Loaded reference data/PDB/CUUCGG/2KOC_5_C_A_0_00001.pdb 
# Loaded target data/PDB/CUUCGG/1K01_1708_C_A_0_00000.pdb 
# Loading data/PDB/CUUCGG/1K01_1708_C_A_0_00000.pdb 
# Loaded reference data/PDB/CUUCGG/2KOC_5_C_A_0_00001.pdb 
# Loaded target data/PDB/CUUCGG/4v50-pdb-bundle1_1028_C_A_0_00002.pdb 
# Loading data/PDB/CUUCGG/4v50-pdb-bundle1_1028_C_A_0_00002.pdb 
# Loaded reference data/PDB/CUUCGG/2KOC_5_C_A_0_00001.pdb 
# Loaded target data/PDB/CUUCGG/5ib8-pdb-bundle2_1691_C_A_0_00000.pdb 
# Loading data/PDB/CUUCGG/5ib8-pdb-bundle2_1691_C_A_0_00000.pdb 
# Loaded reference data/PDB/CUUCGG/2KOC_5_C_A_0_00001.pdb 
# Loaded target data/PDB/CUUCGG/2KOC_5_C_A_0_00008.pdb 
# Loading data/PDB/CUUCGG/2KOC_5_C_A_0_00008.pdb 
# Loaded reference data/PDB/CUUCGG/2KOC_5_C_A_0_00001.pdb 
# Loaded target data/PDB/CUUCGG/6boh-pdb-bundle2_419_C_1_0_00000.pdb 
# Loading data/PDB/CUUCGG/6boh-pdb-bundle2_419_C_1_0_00000.pdb 
# Loaded reference data/PDB/CUUCGG/2KOC_5_C_A_0_00001.pdb 
# Loaded target data/PDB/CUUCGG/1F7Y_8_C_B_0_00000.pdb 
# Loading data/PDB/CUUCGG/1F7Y_8_C_B_0_00000.pdb 
# Loaded reference data/PDB/CUUCGG/2KOC_5_C_A_0_00001.pdb 
# Loaded target data/PDB/CUUCGG/2KMJ_29_C_A_0_00005.pdb 
# Loading data/PDB/CUUCGG/2KMJ_29_C_A_0_00005.pdb 
# Loaded reference data/PDB/CUUCGG/2KOC_5_C_A_0_00001.pdb 
# Loaded target data/PDB/CUUCGG/2KTZ_19_C_A_0_00007.pdb 
# Loading data/PDB/CUUCGG/2KTZ_19_C_A_0_00007.pdb 
# Loaded reference data/PDB/CUUCGG/2KOC_5_C_A_0_00001.pdb 
# Loaded target data/PDB/CUUCGG/4v6g-pdb-bundle3_419_C_A_0_00000.pdb 
# Loading data/PDB/CUUCGG/4v6g-pdb-bundle3_419_C_A_0_00000.pdb 
# Loaded reference data/PDB/CUUCGG/2KOC_5_C_A_0_00001.pdb 
# Loaded target data/PDB/CUUCGG/1VQL_1769_C_0_0_00000.pdb 
# Loading data/PDB/CUUCGG/1VQL_1769_C_0_0_00000.pdb 
# Loaded reference data/PDB/CUUCGG/2KOC_5_C_A_0_00001.pdb 
# Loaded target data/PDB/CUUCGG/1Z31_270_C_A_0_00008.pdb 
# Loading data/PDB/CUUCGG/1Z31_270_C_A_0_00008.pdb 
# Loaded reference data/PDB/CUUCGG/2KOC_5_C_A_0_00001.pdb 
# Loaded target data/PDB/CUUCGG/6gsk-pdb-bundle1_419_C_A_0_00000.pdb 
# Loading data/PDB/CUUCGG/6gsk-pdb-bundle1_419_C_A_0_00000.pdb 
# Loaded reference data/PDB/CUUCGG/2KOC_5_C_A_0_00001.pdb 
# Loaded target data/PDB/CUUCGG/5ndj-pdb-bundle2_1741_C_A_0_00000.pdb 
# Loading data/PDB/CUUCGG/5ndj-pdb-bundle2_1741_C_A_0_00000.pdb 
# Loaded reference data/PDB/CUUCGG/2KOC_5_C_A_0_00001.pdb 
# Loaded target data/PDB/CUUCGG/1IKD_7_C_A_0_00002.pdb 
# Loading data/PDB/CUUCGG/1IKD_7_C_A_0_00002.pdb 
# Loaded reference data/PDB/CUUCGG/2KOC_5_C_A_0_00001.pdb 
# Loaded target data/PDB/CUUCGG/2L2K_19_C_A_0_00001.pdb 
# Loading data/PDB/CUUCGG/2L2K_19_C_A_0_00001.pdb 
# Loaded reference data/PDB/CUUCGG/2KOC_5_C_A_0_00001.pdb 
# Loaded target data/PDB/CUUCGG/4www-pdb-bundle3_1691_C_G_0_00000.pdb 
# Loading data/PDB/CUUCGG/4www-pdb-bundle3_1691_C_G_0_00000.pdb 
# Loaded reference data/PDB/CUUCGG/2KOC_5_C_A_0_00001.pdb 
# Loaded target data/PDB/CUUCGG/1IKD_7_C_A_0_00016.pdb 
# Loading data/PDB/CUUCGG/1IKD_7_C_A_0_00016.pdb 
# Loaded reference data/PDB/CUUCGG/2KOC_5_C_A_0_00001.pdb 
# Loaded target data/PDB/CUUCGG/2L2K_19_C_A_0_00015.pdb 
# Loading data/PDB/CUUCGG/2L2K_19_C_A_0_00015.pdb 
# Loaded reference data/PDB/CUUCGG/2KOC_5_C_A_0_00001.pdb 
# Loaded target data/PDB/CUUCGG/5wis-pdb-bundle2_419_C_A_0_00000.pdb 
# Loading data/PDB/CUUCGG/5wis-pdb-bundle2_419_C_A_0_00000.pdb 
# Loaded reference data/PDB/CUUCGG/2KOC_5_C_A_0_00001.pdb 
# Loaded target data/PDB/CUUCGG/5wis-pdb-bundle1_1738_C_A_0_00000.pdb 
# Loading data/PDB/CUUCGG/5wis-pdb-bundle1_1738_C_A_0_00000.pdb 
# Loaded reference data/PDB/CUUCGG/2KOC_5_C_A_0_00001.pdb 
# Loaded target data/PDB/CUUCGG/1C0O_5_C_A_0_00008.pdb 
# Loading data/PDB/CUUCGG/1C0O_5_C_A_0_00008.pdb 
# Loaded reference data/PDB/CUUCGG/2KOC_5_C_A_0_00001.pdb 
# Loaded target data/PDB/CUUCGG/4w2f-pdb-bundle2_1738_C_A_0_00000.pdb 
# Loading data/PDB/CUUCGG/4w2f-pdb-bundle2_1738_C_A_0_00000.pdb 
# Loaded reference data/PDB/CUUCGG/2KOC_5_C_A_0_00001.pdb 
# Loaded target data/PDB/CUUCGG/1C0O_5_C_A_0_00009.pdb 
# Loading data/PDB/CUUCGG/1C0O_5_C_A_0_00009.pdb 
# Loaded reference data/PDB/CUUCGG/2KOC_5_C_A_0_00001.pdb 
# Loaded target data/PDB/CUUCGG/4v6g-pdb-bundle2_1691_C_A_0_00000.pdb 
# Loading data/PDB/CUUCGG/4v6g-pdb-bundle2_1691_C_A_0_00000.pdb 
# Loaded reference data/PDB/CUUCGG/2KOC_5_C_A_0_00001.pdb 
# Loaded target data/PDB/CUUCGG/6cfk-pdb-bundle2_419_C_A_0_00000.pdb 
# Loading data/PDB/CUUCGG/6cfk-pdb-bundle2_419_C_A_0_00000.pdb 
# Loaded reference data/PDB/CUUCGG/2KOC_5_C_A_0_00001.pdb 
# Loaded target data/PDB/CUUCGG/4X65_419_C_A_0_00000.pdb 
# Loading data/PDB/CUUCGG/4X65_419_C_A_0_00000.pdb 
# Loaded reference data/PDB/CUUCGG/2KOC_5_C_A_0_00001.pdb 
# Loaded target data/PDB/CUUCGG/4u20-pdb-bundle1_419_C_A_0_00001.pdb 
# Loading data/PDB/CUUCGG/4u20-pdb-bundle1_419_C_A_0_00001.pdb 
# Loaded reference data/PDB/CUUCGG/2KOC_5_C_A_0_00001.pdb 
# Loaded target data/PDB/CUUCGG/4v9k-pdb-bundle2_1691_C_A_0_00000.pdb 
# Loading data/PDB/CUUCGG/4v9k-pdb-bundle2_1691_C_A_0_00000.pdb 
# Loaded reference data/PDB/CUUCGG/2KOC_5_C_A_0_00001.pdb 
# Loaded target data/PDB/CUUCGG/5jc9-pdb-bundle1_1449_C_A_0_00003.pdb 
# Loading data/PDB/CUUCGG/5jc9-pdb-bundle1_1449_C_A_0_00003.pdb 
# Loaded reference data/PDB/CUUCGG/2KOC_5_C_A_0_00001.pdb 
# Loaded target data/PDB/CUUCGG/2L2K_19_C_A_0_00014.pdb 
# Loading data/PDB/CUUCGG/2L2K_19_C_A_0_00014.pdb 
# Loaded reference data/PDB/CUUCGG/2KOC_5_C_A_0_00001.pdb 
# Loaded target data/PDB/CUUCGG/1IKD_7_C_A_0_00017.pdb 
# Loading data/PDB/CUUCGG/1IKD_7_C_A_0_00017.pdb 
# Loaded reference data/PDB/CUUCGG/2KOC_5_C_A_0_00001.pdb 
# Loaded target data/PDB/CUUCGG/4v6c-pdb-bundle1_419_C_U_0_00001.pdb 
# Loading data/PDB/CUUCGG/4v6c-pdb-bundle1_419_C_U_0_00001.pdb 
# Loaded reference data/PDB/CUUCGG/2KOC_5_C_A_0_00001.pdb 
# Loaded target data/PDB/CUUCGG/6nd6-pdb-bundle3_1691_C_A_0_00000.pdb 
# Loading data/PDB/CUUCGG/6nd6-pdb-bundle3_1691_C_A_0_00000.pdb 
# Loaded reference data/PDB/CUUCGG/2KOC_5_C_A_0_00001.pdb 
# Loaded target data/PDB/CUUCGG/2L2K_19_C_A_0_00000.pdb 
# Loading data/PDB/CUUCGG/2L2K_19_C_A_0_00000.pdb 
# Loaded reference data/PDB/CUUCGG/2KOC_5_C_A_0_00001.pdb 
# Loaded target data/PDB/CUUCGG/1IKD_7_C_A_0_00003.pdb 
# Loading data/PDB/CUUCGG/1IKD_7_C_A_0_00003.pdb 
# Loaded reference data/PDB/CUUCGG/2KOC_5_C_A_0_00001.pdb 
# Loaded target data/PDB/CUUCGG/4v4h-pdb-bundle1_207_C_A_0_00000.pdb 
# Loading data/PDB/CUUCGG/4v4h-pdb-bundle1_207_C_A_0_00000.pdb 
# Loaded reference data/PDB/CUUCGG/2KOC_5_C_A_0_00001.pdb 
# Loaded target data/PDB/CUUCGG/1Z31_270_C_A_0_00009.pdb 
# Loading data/PDB/CUUCGG/1Z31_270_C_A_0_00009.pdb 
# Loaded reference data/PDB/CUUCGG/2KOC_5_C_A_0_00001.pdb 
# Loaded target data/PDB/CUUCGG/1F7H_12_C_A_0_00010.pdb 
# Loading data/PDB/CUUCGG/1F7H_12_C_A_0_00010.pdb 
# Loaded reference data/PDB/CUUCGG/2KOC_5_C_A_0_00001.pdb 
# Loaded target data/PDB/CUUCGG/2O44_1708_C_A_0_00000.pdb 
# Loading data/PDB/CUUCGG/2O44_1708_C_A_0_00000.pdb 
# Loaded reference data/PDB/CUUCGG/2KOC_5_C_A_0_00001.pdb 
# Loaded target data/PDB/CUUCGG/3CC4_1769_C_0_0_00000.pdb 
# Loading data/PDB/CUUCGG/3CC4_1769_C_0_0_00000.pdb 
# Loaded reference data/PDB/CUUCGG/2KOC_5_C_A_0_00001.pdb 
# Loaded target data/PDB/CUUCGG/2KMJ_29_C_A_0_00004.pdb 
# Loading data/PDB/CUUCGG/2KMJ_29_C_A_0_00004.pdb 
# Loaded reference data/PDB/CUUCGG/2KOC_5_C_A_0_00001.pdb 
# Loaded target data/PDB/CUUCGG/5j30-pdb-bundle1_1691_C_A_0_00000.pdb 
# Loading data/PDB/CUUCGG/5j30-pdb-bundle1_1691_C_A_0_00000.pdb 
# Loaded reference data/PDB/CUUCGG/2KOC_5_C_A_0_00001.pdb 
# Loaded target data/PDB/CUUCGG/2KTZ_19_C_A_0_00006.pdb 
# Loading data/PDB/CUUCGG/2KTZ_19_C_A_0_00006.pdb 
# Loaded reference data/PDB/CUUCGG/2KOC_5_C_A_0_00001.pdb 
# Loaded target data/PDB/CUUCGG/5aj0-pdb-bundle2_2851_C_A_0_00000.pdb 
# Loading data/PDB/CUUCGG/5aj0-pdb-bundle2_2851_C_A_0_00000.pdb 
# Loaded reference data/PDB/CUUCGG/2KOC_5_C_A_0_00001.pdb 
# Loaded target data/PDB/CUUCGG/4u25-pdb-bundle3_419_C_A_0_00001.pdb 
# Loading data/PDB/CUUCGG/4u25-pdb-bundle3_419_C_A_0_00001.pdb 
# Loaded reference data/PDB/CUUCGG/2KOC_5_C_A_0_00001.pdb 
# Loaded target data/PDB/CUUCGG/2KOC_5_C_A_0_00009.pdb 
# Loading data/PDB/CUUCGG/2KOC_5_C_A_0_00009.pdb 
# Loaded reference data/PDB/CUUCGG/2KOC_5_C_A_0_00001.pdb 
# Loaded target data/PDB/CUUCGG/4u27-pdb-bundle1_1028_C_A_0_00002.pdb 
# Loading data/PDB/CUUCGG/4u27-pdb-bundle1_1028_C_A_0_00002.pdb 
# Loaded reference data/PDB/CUUCGG/2KOC_5_C_A_0_00001.pdb 
# Loaded target data/PDB/CUUCGG/4u4q-pdb-bundle2_1923_C_A_0_00002.pdb 
# Loading data/PDB/CUUCGG/4u4q-pdb-bundle2_1923_C_A_0_00002.pdb 
# Loaded reference data/PDB/CUUCGG/2KOC_5_C_A_0_00001.pdb 
# Loaded target data/PDB/CUUCGG/5IEM_25_C_A_0_00001.pdb 
# Loading data/PDB/CUUCGG/5IEM_25_C_A_0_00001.pdb 
# Loaded reference data/PDB/CUUCGG/2KOC_5_C_A_0_00001.pdb 
# Loaded target data/PDB/CUUCGG/4v50-pdb-bundle1_419_C_A_0_00001.pdb 
# Loading data/PDB/CUUCGG/4v50-pdb-bundle1_419_C_A_0_00001.pdb 
# Loaded reference data/PDB/CUUCGG/2KOC_5_C_A_0_00001.pdb 
# Loaded target data/PDB/CUUCGG/5nwy-pdb-bundle2_1449_C_0_0_00003.pdb 
# Loading data/PDB/CUUCGG/5nwy-pdb-bundle2_1449_C_0_0_00003.pdb 
# Loaded reference data/PDB/CUUCGG/2KOC_5_C_A_0_00001.pdb 
# Loaded target data/PDB/CUUCGG/5j4b-pdb-bundle3_1691_C_A_0_00000.pdb 
# Loading data/PDB/CUUCGG/5j4b-pdb-bundle3_1691_C_A_0_00000.pdb 
# Loaded reference data/PDB/CUUCGG/2KOC_5_C_A_0_00001.pdb 
# Loaded target data/PDB/CUUCGG/1M82_10_C_A_0_00022.pdb 
# Loading data/PDB/CUUCGG/1M82_10_C_A_0_00022.pdb 
# Loaded reference data/PDB/CUUCGG/2KOC_5_C_A_0_00001.pdb 
# Loaded target data/PDB/CUUCGG/5j7l-pdb-bundle2_1691_C_U_0_00000.pdb 
# Loading data/PDB/CUUCGG/5j7l-pdb-bundle2_1691_C_U_0_00000.pdb 
# Loaded reference data/PDB/CUUCGG/2KOC_5_C_A_0_00001.pdb 
# Loaded target data/PDB/CUUCGG/1N8R_1769_C_A_0_00000.pdb 
# Loading data/PDB/CUUCGG/1N8R_1769_C_A_0_00000.pdb 
# Loaded reference data/PDB/CUUCGG/2KOC_5_C_A_0_00001.pdb 
# Loaded target data/PDB/CUUCGG/5dge-pdb-bundle3_230_C_d_0_00000.pdb 
# Loading data/PDB/CUUCGG/5dge-pdb-bundle3_230_C_d_0_00000.pdb 
# Loaded reference data/PDB/CUUCGG/2KOC_5_C_A_0_00001.pdb 
# Loaded target data/PDB/CUUCGG/1Y69_1708_C_0_0_00000.pdb 
# Loading data/PDB/CUUCGG/1Y69_1708_C_0_0_00000.pdb 
# Loaded reference data/PDB/CUUCGG/2KOC_5_C_A_0_00001.pdb 
# Loaded target data/PDB/CUUCGG/1FYP_11_C_A_0_00026.pdb 
# Loading data/PDB/CUUCGG/1FYP_11_C_A_0_00026.pdb 
# Loaded reference data/PDB/CUUCGG/2KOC_5_C_A_0_00001.pdb 
# Loaded target data/PDB/CUUCGG/4u4q-pdb-bundle3_230_C_d_0_00000.pdb 
# Loading data/PDB/CUUCGG/4u4q-pdb-bundle3_230_C_d_0_00000.pdb 
# Loaded reference data/PDB/CUUCGG/2KOC_5_C_A_0_00001.pdb 
# Loaded target data/PDB/CUUCGG/4v9i-pdb-bundle4_1737_C_A_0_00000.pdb 
# Loading data/PDB/CUUCGG/4v9i-pdb-bundle4_1737_C_A_0_00000.pdb 
# Loaded reference data/PDB/CUUCGG/2KOC_5_C_A_0_00001.pdb 
# Loaded target data/PDB/CUUCGG/1AUD_28_C_B_0_00026.pdb 
# Loading data/PDB/CUUCGG/1AUD_28_C_B_0_00026.pdb 
# Loaded reference data/PDB/CUUCGG/2KOC_5_C_A_0_00001.pdb 
# Loaded target data/PDB/CUUCGG/4v8d-pdb-bundle2_1691_C_A_0_00000.pdb 
# Loading data/PDB/CUUCGG/4v8d-pdb-bundle2_1691_C_A_0_00000.pdb 
# Loaded reference data/PDB/CUUCGG/2KOC_5_C_A_0_00001.pdb 
# Loaded target data/PDB/CUUCGG/1EBQ_12_C_A_0_00003.pdb 
# Loading data/PDB/CUUCGG/1EBQ_12_C_A_0_00003.pdb 
# Loaded reference data/PDB/CUUCGG/2KOC_5_C_A_0_00001.pdb 
# Loaded target data/PDB/CUUCGG/4w2i-pdb-bundle3_419_C_A_0_00000.pdb 
# Loading data/PDB/CUUCGG/4w2i-pdb-bundle3_419_C_A_0_00000.pdb 
# Loaded reference data/PDB/CUUCGG/2KOC_5_C_A_0_00001.pdb 
# Loaded target data/PDB/CUUCGG/3jbv-pdb-bundle1_1028_C_A_0_00002.pdb 
# Loading data/PDB/CUUCGG/3jbv-pdb-bundle1_1028_C_A_0_00002.pdb 
# Loaded reference data/PDB/CUUCGG/2KOC_5_C_A_0_00001.pdb 
# Loaded target data/PDB/CUUCGG/4u4z-pdb-bundle4_1923_C_I_0_00002.pdb 
# Loading data/PDB/CUUCGG/4u4z-pdb-bundle4_1923_C_I_0_00002.pdb 
# Loaded reference data/PDB/CUUCGG/2KOC_5_C_A_0_00001.pdb 
# Loaded target data/PDB/CUUCGG/1M5L_17_C_A_0_00004.pdb 
# Loading data/PDB/CUUCGG/1M5L_17_C_A_0_00004.pdb 
# Loaded reference data/PDB/CUUCGG/2KOC_5_C_A_0_00001.pdb 
# Loaded target data/PDB/CUUCGG/1EKZ_13_C_B_0_00013.pdb 
# Loading data/PDB/CUUCGG/1EKZ_13_C_B_0_00013.pdb 
# Loaded reference data/PDB/CUUCGG/2KOC_5_C_A_0_00001.pdb 
# Loaded target data/PDB/CUUCGG/4v9r-pdb-bundle1_419_C_A_0_00000.pdb 
# Loading data/PDB/CUUCGG/4v9r-pdb-bundle1_419_C_A_0_00000.pdb 
# Loaded reference data/PDB/CUUCGG/2KOC_5_C_A_0_00001.pdb 
# Loaded target data/PDB/CUUCGG/5j3c-pdb-bundle3_1691_C_A_0_00000.pdb 
# Loading data/PDB/CUUCGG/5j3c-pdb-bundle3_1691_C_A_0_00000.pdb 
# Loaded reference data/PDB/CUUCGG/2KOC_5_C_A_0_00001.pdb 
# Loaded target data/PDB/CUUCGG/2N6T_19_C_A_0_00008.pdb 
# Loading data/PDB/CUUCGG/2N6T_19_C_A_0_00008.pdb 
# Loaded reference data/PDB/CUUCGG/2KOC_5_C_A_0_00001.pdb 
# Loaded target data/PDB/CUUCGG/1EKZ_13_C_B_0_00007.pdb 
# Loading data/PDB/CUUCGG/1EKZ_13_C_B_0_00007.pdb 
# Loaded reference data/PDB/CUUCGG/2KOC_5_C_A_0_00001.pdb 
# Loaded target data/PDB/CUUCGG/1M5L_17_C_A_0_00010.pdb 
# Loading data/PDB/CUUCGG/1M5L_17_C_A_0_00010.pdb 
# Loaded reference data/PDB/CUUCGG/2KOC_5_C_A_0_00001.pdb 
# Loaded target data/PDB/CUUCGG/1F7I_12_C_A_0_00008.pdb 
# Loading data/PDB/CUUCGG/1F7I_12_C_A_0_00008.pdb 
# Loaded reference data/PDB/CUUCGG/2KOC_5_C_A_0_00001.pdb 
# Loaded target data/PDB/CUUCGG/1EKZ_13_C_B_0_00011.pdb 
# Loading data/PDB/CUUCGG/1EKZ_13_C_B_0_00011.pdb 
# Loaded reference data/PDB/CUUCGG/2KOC_5_C_A_0_00001.pdb 
# Loaded target data/PDB/CUUCGG/1M5L_17_C_A_0_00006.pdb 
# Loading data/PDB/CUUCGG/1M5L_17_C_A_0_00006.pdb 
# Loaded reference data/PDB/CUUCGG/2KOC_5_C_A_0_00001.pdb 
# Loaded target data/PDB/CUUCGG/1M5L_17_C_A_0_00012.pdb 
# Loading data/PDB/CUUCGG/1M5L_17_C_A_0_00012.pdb 
# Loaded reference data/PDB/CUUCGG/2KOC_5_C_A_0_00001.pdb 
# Loaded target data/PDB/CUUCGG/5e81-pdb-bundle2_1691_C_A_0_00000.pdb 
# Loading data/PDB/CUUCGG/5e81-pdb-bundle2_1691_C_A_0_00000.pdb 
# Loaded reference data/PDB/CUUCGG/2KOC_5_C_A_0_00001.pdb 
# Loaded target data/PDB/CUUCGG/4v6g-pdb-bundle1_419_C_A_0_00000.pdb 
# Loading data/PDB/CUUCGG/4v6g-pdb-bundle1_419_C_A_0_00000.pdb 
# Loaded reference data/PDB/CUUCGG/2KOC_5_C_A_0_00001.pdb 
# Loaded target data/PDB/CUUCGG/4v7l-pdb-bundle2_1691_C_A_0_00000.pdb 
# Loading data/PDB/CUUCGG/4v7l-pdb-bundle2_1691_C_A_0_00000.pdb 
# Loaded reference data/PDB/CUUCGG/2KOC_5_C_A_0_00001.pdb 
# Loaded target data/PDB/CUUCGG/2L2J_19_C_A_0_00018.pdb 
# Loading data/PDB/CUUCGG/2L2J_19_C_A_0_00018.pdb 
# Loaded reference data/PDB/CUUCGG/2KOC_5_C_A_0_00001.pdb 
# Loaded target data/PDB/CUUCGG/1RAW_21_C_A_0_00008.pdb 
# Loading data/PDB/CUUCGG/1RAW_21_C_A_0_00008.pdb 
# Loaded reference data/PDB/CUUCGG/2KOC_5_C_A_0_00001.pdb 
# Loaded target data/PDB/CUUCGG/1EBQ_12_C_A_0_00001.pdb 
# Loading data/PDB/CUUCGG/1EBQ_12_C_A_0_00001.pdb 
# Loaded reference data/PDB/CUUCGG/2KOC_5_C_A_0_00001.pdb 
# Loaded target data/PDB/CUUCGG/1AUD_28_C_B_0_00030.pdb 
# Loading data/PDB/CUUCGG/1AUD_28_C_B_0_00030.pdb 
# Loaded reference data/PDB/CUUCGG/2KOC_5_C_A_0_00001.pdb 
# Loaded target data/PDB/CUUCGG/4v8n-pdb-bundle4_1691_C_A_0_00000.pdb 
# Loading data/PDB/CUUCGG/4v8n-pdb-bundle4_1691_C_A_0_00000.pdb 
# Loaded reference data/PDB/CUUCGG/2KOC_5_C_A_0_00001.pdb 
# Loaded target data/PDB/CUUCGG/1I6U_16_C_D_0_00001.pdb 
# Loading data/PDB/CUUCGG/1I6U_16_C_D_0_00001.pdb 
# Loaded reference data/PDB/CUUCGG/2KOC_5_C_A_0_00001.pdb 
# Loaded target data/PDB/CUUCGG/1FYP_11_C_A_0_00018.pdb 
# Loading data/PDB/CUUCGG/1FYP_11_C_A_0_00018.pdb 
# Loaded reference data/PDB/CUUCGG/2KOC_5_C_A_0_00001.pdb 
# Loaded target data/PDB/CUUCGG/4woi-pdb-bundle4_1449_C_A_0_00003.pdb 
# Loading data/PDB/CUUCGG/4woi-pdb-bundle4_1449_C_A_0_00003.pdb 
# Loaded reference data/PDB/CUUCGG/2KOC_5_C_A_0_00001.pdb 
# Loaded target data/PDB/CUUCGG/6enu-pdb-bundle1_1449_C_a_0_00003.pdb 
# Loading data/PDB/CUUCGG/6enu-pdb-bundle1_1449_C_a_0_00003.pdb 
# Loaded reference data/PDB/CUUCGG/2KOC_5_C_A_0_00001.pdb 
# Loaded target data/PDB/CUUCGG/1AUD_28_C_B_0_00024.pdb 
# Loading data/PDB/CUUCGG/1AUD_28_C_B_0_00024.pdb 
# Loaded reference data/PDB/CUUCGG/2KOC_5_C_A_0_00001.pdb 
# Loaded target data/PDB/CUUCGG/1FYP_11_C_A_0_00024.pdb 
# Loading data/PDB/CUUCGG/1FYP_11_C_A_0_00024.pdb 
# Loaded reference data/PDB/CUUCGG/2KOC_5_C_A_0_00001.pdb 
# Loaded target data/PDB/CUUCGG/4v9d-pdb-bundle2_1691_C_N_0_00000.pdb 
# Loading data/PDB/CUUCGG/4v9d-pdb-bundle2_1691_C_N_0_00000.pdb 
# Loaded reference data/PDB/CUUCGG/2KOC_5_C_A_0_00001.pdb 
# Loaded target data/PDB/CUUCGG/1M82_10_C_A_0_00008.pdb 
# Loading data/PDB/CUUCGG/1M82_10_C_A_0_00008.pdb 
# Loaded reference data/PDB/CUUCGG/2KOC_5_C_A_0_00001.pdb 
# Loaded target data/PDB/CUUCGG/4wzd-pdb-bundle2_1691_C_A_0_00000.pdb 
# Loading data/PDB/CUUCGG/4wzd-pdb-bundle2_1691_C_A_0_00000.pdb 
# Loaded reference data/PDB/CUUCGG/2KOC_5_C_A_0_00001.pdb 
# Loaded target data/PDB/CUUCGG/4wt1-pdb-bundle2_1691_C_A_0_00000.pdb 
# Loading data/PDB/CUUCGG/4wt1-pdb-bundle2_1691_C_A_0_00000.pdb 
# Loaded reference data/PDB/CUUCGG/2KOC_5_C_A_0_00001.pdb 
# Loaded target data/PDB/CUUCGG/4v52-pdb-bundle2_1691_C_A_0_00000.pdb 
# Loading data/PDB/CUUCGG/4v52-pdb-bundle2_1691_C_A_0_00000.pdb 
# Loaded reference data/PDB/CUUCGG/2KOC_5_C_A_0_00001.pdb 
# Loaded target data/PDB/CUUCGG/1M82_10_C_A_0_00020.pdb 
# Loading data/PDB/CUUCGG/1M82_10_C_A_0_00020.pdb 
# Loaded reference data/PDB/CUUCGG/2KOC_5_C_A_0_00001.pdb 
# Loaded target data/PDB/CUUCGG/4v63-pdb-bundle3_419_C_A_0_00000.pdb 
# Loading data/PDB/CUUCGG/4v63-pdb-bundle3_419_C_A_0_00000.pdb 
# Loaded reference data/PDB/CUUCGG/2KOC_5_C_A_0_00001.pdb 
# Loaded target data/PDB/CUUCGG/4v5a-pdb-bundle3_419_C_A_0_00000.pdb 
# Loading data/PDB/CUUCGG/4v5a-pdb-bundle3_419_C_A_0_00000.pdb 
# Loaded reference data/PDB/CUUCGG/2KOC_5_C_A_0_00001.pdb 
# Loaded target data/PDB/CUUCGG/4v5j-pdb-bundle4_1691_C_A_0_00000.pdb 
# Loading data/PDB/CUUCGG/4v5j-pdb-bundle4_1691_C_A_0_00000.pdb 
# Loaded reference data/PDB/CUUCGG/2KOC_5_C_A_0_00001.pdb 
# Loaded target data/PDB/CUUCGG/5IEM_25_C_A_0_00003.pdb 
# Loading data/PDB/CUUCGG/5IEM_25_C_A_0_00003.pdb 
# Loaded reference data/PDB/CUUCGG/2KOC_5_C_A_0_00001.pdb 
# Loaded target data/PDB/CUUCGG/4v9p-pdb-bundle4_1449_C_A_0_00003.pdb 
# Loading data/PDB/CUUCGG/4v9p-pdb-bundle4_1449_C_A_0_00003.pdb 
# Loaded reference data/PDB/CUUCGG/2KOC_5_C_A_0_00001.pdb 
# Loaded target data/PDB/CUUCGG/5e7k-pdb-bundle2_1691_C_A_0_00000.pdb 
# Loading data/PDB/CUUCGG/5e7k-pdb-bundle2_1691_C_A_0_00000.pdb 
# Loaded reference data/PDB/CUUCGG/2KOC_5_C_A_0_00001.pdb 
# Loaded target data/PDB/CUUCGG/1vy7-pdb-bundle4_1691_C_A_0_00000.pdb 
# Loading data/PDB/CUUCGG/1vy7-pdb-bundle4_1691_C_A_0_00000.pdb 
# Loaded reference data/PDB/CUUCGG/2KOC_5_C_A_0_00001.pdb 
# Loaded target data/PDB/CUUCGG/2KTZ_19_C_A_0_00004.pdb 
# Loading data/PDB/CUUCGG/2KTZ_19_C_A_0_00004.pdb 
# Loaded reference data/PDB/CUUCGG/2KOC_5_C_A_0_00001.pdb 
# Loaded target data/PDB/CUUCGG/4v6c-pdb-bundle2_1691_C_A_0_00000.pdb 
# Loading data/PDB/CUUCGG/4v6c-pdb-bundle2_1691_C_A_0_00000.pdb 
# Loaded reference data/PDB/CUUCGG/2KOC_5_C_A_0_00001.pdb 
# Loaded target data/PDB/CUUCGG/2KMJ_29_C_A_0_00006.pdb 
# Loading data/PDB/CUUCGG/2KMJ_29_C_A_0_00006.pdb 
# Loaded reference data/PDB/CUUCGG/2KOC_5_C_A_0_00001.pdb 
# Loaded target data/PDB/CUUCGG/4X66_419_C_A_0_00000.pdb 
# Loading data/PDB/CUUCGG/4X66_419_C_A_0_00000.pdb 
# Loaded reference data/PDB/CUUCGG/2KOC_5_C_A_0_00001.pdb 
# Loaded target data/PDB/CUUCGG/5umd-pdb-bundle1_2191_C_A_0_00000.pdb 
# Loading data/PDB/CUUCGG/5umd-pdb-bundle1_2191_C_A_0_00000.pdb 
# Loaded reference data/PDB/CUUCGG/2KOC_5_C_A_0_00001.pdb 
# Loaded target data/PDB/CUUCGG/1G70_54_C_A_0_00008.pdb 
# Loading data/PDB/CUUCGG/1G70_54_C_A_0_00008.pdb 
# Loaded reference data/PDB/CUUCGG/2KOC_5_C_A_0_00001.pdb 
# Loaded target data/PDB/CUUCGG/1IKD_7_C_A_0_00029.pdb 
# Loading data/PDB/CUUCGG/1IKD_7_C_A_0_00029.pdb 
# Loaded reference data/PDB/CUUCGG/2KOC_5_C_A_0_00001.pdb 
# Loaded target data/PDB/CUUCGG/6cae-pdb-bundle3_1691_C_A_0_00000.pdb 
# Loading data/PDB/CUUCGG/6cae-pdb-bundle3_1691_C_A_0_00000.pdb 
# Loaded reference data/PDB/CUUCGG/2KOC_5_C_A_0_00001.pdb 
# Loaded target data/PDB/CUUCGG/4v5d-pdb-bundle1_419_C_A_0_00000.pdb 
# Loading data/PDB/CUUCGG/4v5d-pdb-bundle1_419_C_A_0_00000.pdb 
# Loaded reference data/PDB/CUUCGG/2KOC_5_C_A_0_00001.pdb 
# Loaded target data/PDB/CUUCGG/1F7H_12_C_A_0_00006.pdb 
# Loading data/PDB/CUUCGG/1F7H_12_C_A_0_00006.pdb 
# Loaded reference data/PDB/CUUCGG/2KOC_5_C_A_0_00001.pdb 
# Loaded target data/PDB/CUUCGG/2L2K_19_C_A_0_00016.pdb 
# Loading data/PDB/CUUCGG/2L2K_19_C_A_0_00016.pdb 
# Loaded reference data/PDB/CUUCGG/2KOC_5_C_A_0_00001.pdb 
# Loaded target data/PDB/CUUCGG/1IKD_7_C_A_0_00015.pdb 
# Loading data/PDB/CUUCGG/1IKD_7_C_A_0_00015.pdb 
# Loaded reference data/PDB/CUUCGG/2KOC_5_C_A_0_00001.pdb 
# Loaded target data/PDB/CUUCGG/6nd6-pdb-bundle4_419_C_A_0_00000.pdb 
# Loading data/PDB/CUUCGG/6nd6-pdb-bundle4_419_C_A_0_00000.pdb 
# Loaded reference data/PDB/CUUCGG/2KOC_5_C_A_0_00001.pdb 
# Loaded target data/PDB/CUUCGG/5we4-pdb-bundle2_419_C_a_0_00001.pdb 
# Loading data/PDB/CUUCGG/5we4-pdb-bundle2_419_C_a_0_00001.pdb 
# Loaded reference data/PDB/CUUCGG/2KOC_5_C_A_0_00001.pdb 
# Loaded target data/PDB/CUUCGG/4v5d-pdb-bundle2_1691_C_A_0_00000.pdb 
# Loading data/PDB/CUUCGG/4v5d-pdb-bundle2_1691_C_A_0_00000.pdb 
# Loaded reference data/PDB/CUUCGG/2KOC_5_C_A_0_00001.pdb 
# Loaded target data/PDB/CUUCGG/2L2K_19_C_A_0_00002.pdb 
# Loading data/PDB/CUUCGG/2L2K_19_C_A_0_00002.pdb 
# Loaded reference data/PDB/CUUCGG/2KOC_5_C_A_0_00001.pdb 
# Loaded target data/PDB/CUUCGG/4v9a-pdb-bundle4_1691_C_A_0_00000.pdb 
# Loading data/PDB/CUUCGG/4v9a-pdb-bundle4_1691_C_A_0_00000.pdb 
# Loaded reference data/PDB/CUUCGG/2KOC_5_C_A_0_00001.pdb 
# Loaded target data/PDB/CUUCGG/2HGH_32_C_B_0_00019.pdb 
# Loading data/PDB/CUUCGG/2HGH_32_C_B_0_00019.pdb 
# Loaded reference data/PDB/CUUCGG/2KOC_5_C_A_0_00001.pdb 
# Loaded target data/PDB/CUUCGG/4u4u-pdb-bundle2_1923_C_A_0_00002.pdb 
# Loading data/PDB/CUUCGG/4u4u-pdb-bundle2_1923_C_A_0_00002.pdb 
# Loaded reference data/PDB/CUUCGG/2KOC_5_C_A_0_00001.pdb 
# Loaded target data/PDB/CUUCGG/2KZL_27_C_A_0_00003.pdb 
# Loading data/PDB/CUUCGG/2KZL_27_C_A_0_00003.pdb 
# Loaded reference data/PDB/CUUCGG/2KOC_5_C_A_0_00001.pdb 
# Loaded target data/PDB/CUUCGG/5mdy-pdb-bundle1_1691_C_1_0_00000.pdb 
# Loading data/PDB/CUUCGG/5mdy-pdb-bundle1_1691_C_1_0_00000.pdb 
# Loaded reference data/PDB/CUUCGG/2KOC_5_C_A_0_00001.pdb 
# Loaded target data/PDB/CUUCGG/4v4q-pdb-bundle3_1449_C_A_0_00003.pdb 
# Loading data/PDB/CUUCGG/4v4q-pdb-bundle3_1449_C_A_0_00003.pdb 
# Loaded reference data/PDB/CUUCGG/2KOC_5_C_A_0_00001.pdb 
# Loaded target data/PDB/CUUCGG/4v5s-pdb-bundle2_1691_C_A_0_00000.pdb 
# Loading data/PDB/CUUCGG/4v5s-pdb-bundle2_1691_C_A_0_00000.pdb 
# Loaded reference data/PDB/CUUCGG/2KOC_5_C_A_0_00001.pdb 
# Loaded target data/PDB/CUUCGG/4w2i-pdb-bundle1_419_C_A_0_00000.pdb 
# Loading data/PDB/CUUCGG/4w2i-pdb-bundle1_419_C_A_0_00000.pdb 
# Loaded reference data/PDB/CUUCGG/2KOC_5_C_A_0_00001.pdb 
# Loaded target data/PDB/CUUCGG/5ngm-pdb-bundle1_1038_C_A_0_00002.pdb 
# Loading data/PDB/CUUCGG/5ngm-pdb-bundle1_1038_C_A_0_00002.pdb 
# Loaded reference data/PDB/CUUCGG/2KOC_5_C_A_0_00001.pdb 
# Loaded target data/PDB/CUUCGG/2HGH_32_C_B_0_00018.pdb 
# Loading data/PDB/CUUCGG/2HGH_32_C_B_0_00018.pdb 
# Loaded reference data/PDB/CUUCGG/2KOC_5_C_A_0_00001.pdb 
# Loaded target data/PDB/CUUCGG/2OTL_1769_C_0_0_00000.pdb 
# Loading data/PDB/CUUCGG/2OTL_1769_C_0_0_00000.pdb 
# Loaded reference data/PDB/CUUCGG/2KOC_5_C_A_0_00001.pdb 
# Loaded target data/PDB/CUUCGG/2L2K_19_C_A_0_00003.pdb 
# Loading data/PDB/CUUCGG/2L2K_19_C_A_0_00003.pdb 
# Loaded reference data/PDB/CUUCGG/2KOC_5_C_A_0_00001.pdb 
# Loaded target data/PDB/CUUCGG/4v54-pdb-bundle1_1028_C_A_0_00002.pdb 
# Loading data/PDB/CUUCGG/4v54-pdb-bundle1_1028_C_A_0_00002.pdb 
# Loaded reference data/PDB/CUUCGG/2KOC_5_C_A_0_00001.pdb 
# Loaded target data/PDB/CUUCGG/5j4c-pdb-bundle4_419_C_A_0_00000.pdb 
# Loading data/PDB/CUUCGG/5j4c-pdb-bundle4_419_C_A_0_00000.pdb 
# Loaded reference data/PDB/CUUCGG/2KOC_5_C_A_0_00001.pdb 
# Loaded target data/PDB/CUUCGG/1YIT_1769_C_0_0_00000.pdb 
# Loading data/PDB/CUUCGG/1YIT_1769_C_0_0_00000.pdb 
# Loaded reference data/PDB/CUUCGG/2KOC_5_C_A_0_00001.pdb 
# Loaded target data/PDB/CUUCGG/2L2K_19_C_A_0_00017.pdb 
# Loading data/PDB/CUUCGG/2L2K_19_C_A_0_00017.pdb 
# Loaded reference data/PDB/CUUCGG/2KOC_5_C_A_0_00001.pdb 
# Loaded target data/PDB/CUUCGG/4woi-pdb-bundle1_1449_C_A_0_00003.pdb 
# Loading data/PDB/CUUCGG/4woi-pdb-bundle1_1449_C_A_0_00003.pdb 
# Loaded reference data/PDB/CUUCGG/2KOC_5_C_A_0_00001.pdb 
# Loaded target data/PDB/CUUCGG/6gsk-pdb-bundle2_1691_C_A_0_00000.pdb 
# Loading data/PDB/CUUCGG/6gsk-pdb-bundle2_1691_C_A_0_00000.pdb 
# Loaded reference data/PDB/CUUCGG/2KOC_5_C_A_0_00001.pdb 
# Loaded target data/PDB/CUUCGG/4v4h-pdb-bundle3_1028_C_A_0_00002.pdb 
# Loading data/PDB/CUUCGG/4v4h-pdb-bundle3_1028_C_A_0_00002.pdb 
# Loaded reference data/PDB/CUUCGG/2KOC_5_C_A_0_00001.pdb 
# Loaded target data/PDB/CUUCGG/1IKD_7_C_A_0_00028.pdb 
# Loading data/PDB/CUUCGG/1IKD_7_C_A_0_00028.pdb 
# Loaded reference data/PDB/CUUCGG/2KOC_5_C_A_0_00001.pdb 
# Loaded target data/PDB/CUUCGG/1G70_54_C_A_0_00009.pdb 
# Loading data/PDB/CUUCGG/1G70_54_C_A_0_00009.pdb 
# Loaded reference data/PDB/CUUCGG/2KOC_5_C_A_0_00001.pdb 
# Loaded target data/PDB/CUUCGG/2KTZ_19_C_A_0_00005.pdb 
# Loading data/PDB/CUUCGG/2KTZ_19_C_A_0_00005.pdb 
# Loaded reference data/PDB/CUUCGG/2KOC_5_C_A_0_00001.pdb 
# Loaded target data/PDB/CUUCGG/2KMJ_29_C_A_0_00007.pdb 
# Loading data/PDB/CUUCGG/2KMJ_29_C_A_0_00007.pdb 
# Loaded reference data/PDB/CUUCGG/2KOC_5_C_A_0_00001.pdb 
# Loaded target data/PDB/CUUCGG/2VQE_419_C_A_0_00000.pdb 
# Loading data/PDB/CUUCGG/2VQE_419_C_A_0_00000.pdb 
# Loaded reference data/PDB/CUUCGG/2KOC_5_C_A_0_00001.pdb 
# Loaded target data/PDB/CUUCGG/4v8f-pdb-bundle3_1691_C_S_0_00000.pdb 
# Loading data/PDB/CUUCGG/4v8f-pdb-bundle3_1691_C_S_0_00000.pdb 
# Loaded reference data/PDB/CUUCGG/2KOC_5_C_A_0_00001.pdb 
# Loaded target data/PDB/CUUCGG/5IEM_25_C_A_0_00002.pdb 
# Loading data/PDB/CUUCGG/5IEM_25_C_A_0_00002.pdb 
# Loaded reference data/PDB/CUUCGG/2KOC_5_C_A_0_00001.pdb 
# Loaded target data/PDB/CUUCGG/4v7s-pdb-bundle1_1449_C_A_0_00003.pdb 
# Loading data/PDB/CUUCGG/4v7s-pdb-bundle1_1449_C_A_0_00003.pdb 
# Loaded reference data/PDB/CUUCGG/2KOC_5_C_A_0_00001.pdb 
# Loaded target data/PDB/CUUCGG/5j88-pdb-bundle1_1449_C_A_0_00003.pdb 
# Loading data/PDB/CUUCGG/5j88-pdb-bundle1_1449_C_A_0_00003.pdb 
# Loaded reference data/PDB/CUUCGG/2KOC_5_C_A_0_00001.pdb 
# Loaded target data/PDB/CUUCGG/1EBR_12_C_A_0_00004.pdb 
# Loading data/PDB/CUUCGG/1EBR_12_C_A_0_00004.pdb 
# Loaded reference data/PDB/CUUCGG/2KOC_5_C_A_0_00001.pdb 
# Loaded target data/PDB/CUUCGG/5j4d-pdb-bundle1_1691_C_B_0_00001.pdb 
# Loading data/PDB/CUUCGG/5j4d-pdb-bundle1_1691_C_B_0_00001.pdb 
# Loaded reference data/PDB/CUUCGG/2KOC_5_C_A_0_00001.pdb 
# Loaded target data/PDB/CUUCGG/5iqr-pdb-bundle2_419_C_2_0_00002.pdb 
# Loading data/PDB/CUUCGG/5iqr-pdb-bundle2_419_C_2_0_00002.pdb 
# Loaded reference data/PDB/CUUCGG/2KOC_5_C_A_0_00001.pdb 
# Loaded target data/PDB/CUUCGG/5wfs-pdb-bundle1_1691_C_A_0_00000.pdb 
# Loading data/PDB/CUUCGG/5wfs-pdb-bundle1_1691_C_A_0_00000.pdb 
# Loaded reference data/PDB/CUUCGG/2KOC_5_C_A_0_00001.pdb 
# Loaded target data/PDB/CUUCGG/1M82_10_C_A_0_00021.pdb 
# Loading data/PDB/CUUCGG/1M82_10_C_A_0_00021.pdb 
# Loaded reference data/PDB/CUUCGG/2KOC_5_C_A_0_00001.pdb 
# Loaded target data/PDB/CUUCGG/5o61-pdb-bundle2_419_C_A_0_00000.pdb 
# Loading data/PDB/CUUCGG/5o61-pdb-bundle2_419_C_A_0_00000.pdb 
# Loaded reference data/PDB/CUUCGG/2KOC_5_C_A_0_00001.pdb 
# Loaded target data/PDB/CUUCGG/5w4k-pdb-bundle1_1738_C_A_0_00000.pdb 
# Loading data/PDB/CUUCGG/5w4k-pdb-bundle1_1738_C_A_0_00000.pdb 
# Loaded reference data/PDB/CUUCGG/2KOC_5_C_A_0_00001.pdb 
# Loaded target data/PDB/CUUCGG/4u25-pdb-bundle1_419_C_A_0_00001.pdb 
# Loading data/PDB/CUUCGG/4u25-pdb-bundle1_419_C_A_0_00001.pdb 
# Loaded reference data/PDB/CUUCGG/2KOC_5_C_A_0_00001.pdb 
# Loaded target data/PDB/CUUCGG/5el6-pdb-bundle4_1691_C_A_0_00000.pdb 
# Loading data/PDB/CUUCGG/5el6-pdb-bundle4_1691_C_A_0_00000.pdb 
# Loaded reference data/PDB/CUUCGG/2KOC_5_C_A_0_00001.pdb 
# Loaded target data/PDB/CUUCGG/1M82_10_C_A_0_00009.pdb 
# Loading data/PDB/CUUCGG/1M82_10_C_A_0_00009.pdb 
# Loaded reference data/PDB/CUUCGG/2KOC_5_C_A_0_00001.pdb 
# Loaded target data/PDB/CUUCGG/4v50-pdb-bundle3_419_C_A_0_00001.pdb 
# Loading data/PDB/CUUCGG/4v50-pdb-bundle3_419_C_A_0_00001.pdb 
# Loaded reference data/PDB/CUUCGG/2KOC_5_C_A_0_00001.pdb 
# Loaded target data/PDB/CUUCGG/1FYP_11_C_A_0_00025.pdb 
# Loading data/PDB/CUUCGG/1FYP_11_C_A_0_00025.pdb 
# Loaded reference data/PDB/CUUCGG/2KOC_5_C_A_0_00001.pdb 
# Loaded target data/PDB/CUUCGG/4u25-pdb-bundle2_1691_C_A_0_00000.pdb 
# Loading data/PDB/CUUCGG/4u25-pdb-bundle2_1691_C_A_0_00000.pdb 
# Loaded reference data/PDB/CUUCGG/2KOC_5_C_A_0_00001.pdb 
# Loaded target data/PDB/CUUCGG/4u50-pdb-bundle4_1923_C_I_0_00002.pdb 
# Loading data/PDB/CUUCGG/4u50-pdb-bundle4_1923_C_I_0_00002.pdb 
# Loaded reference data/PDB/CUUCGG/2KOC_5_C_A_0_00001.pdb 
# Loaded target data/PDB/CUUCGG/5mgp-pdb-bundle2_419_C_a_0_00001.pdb 
# Loading data/PDB/CUUCGG/5mgp-pdb-bundle2_419_C_a_0_00001.pdb 
# Loaded reference data/PDB/CUUCGG/2KOC_5_C_A_0_00001.pdb 
# Loaded target data/PDB/CUUCGG/1VQ6_1769_C_0_0_00000.pdb 
# Loading data/PDB/CUUCGG/1VQ6_1769_C_0_0_00000.pdb 
# Loaded reference data/PDB/CUUCGG/2KOC_5_C_A_0_00001.pdb 
# Loaded target data/PDB/CUUCGG/1FYP_11_C_A_0_00019.pdb 
# Loading data/PDB/CUUCGG/1FYP_11_C_A_0_00019.pdb 
# Loaded reference data/PDB/CUUCGG/2KOC_5_C_A_0_00001.pdb 
# Loaded target data/PDB/CUUCGG/2ZM6_419_C_A_0_00000.pdb 
# Loading data/PDB/CUUCGG/2ZM6_419_C_A_0_00000.pdb 
# Loaded reference data/PDB/CUUCGG/2KOC_5_C_A_0_00001.pdb 
# Loaded target data/PDB/CUUCGG/1S72_1769_C_0_0_00000.pdb 
# Loading data/PDB/CUUCGG/1S72_1769_C_0_0_00000.pdb 
# Loaded reference data/PDB/CUUCGG/2KOC_5_C_A_0_00001.pdb 
# Loaded target data/PDB/CUUCGG/6ek0-pdb-bundle1_2872_C_A_0_00000.pdb 
# Loading data/PDB/CUUCGG/6ek0-pdb-bundle1_2872_C_A_0_00000.pdb 
# Loaded reference data/PDB/CUUCGG/2KOC_5_C_A_0_00001.pdb 
# Loaded target data/PDB/CUUCGG/1F7F_12_C_A_0_00003.pdb 
# Loading data/PDB/CUUCGG/1F7F_12_C_A_0_00003.pdb 
# Loaded reference data/PDB/CUUCGG/2KOC_5_C_A_0_00001.pdb 
# Loaded target data/PDB/CUUCGG/1EBQ_12_C_A_0_00000.pdb 
# Loading data/PDB/CUUCGG/1EBQ_12_C_A_0_00000.pdb 
# Loaded reference data/PDB/CUUCGG/2KOC_5_C_A_0_00001.pdb 
# Loaded target data/PDB/CUUCGG/1RAW_21_C_A_0_00009.pdb 
# Loading data/PDB/CUUCGG/1RAW_21_C_A_0_00009.pdb 
# Loaded reference data/PDB/CUUCGG/2KOC_5_C_A_0_00001.pdb 
# Loaded target data/PDB/CUUCGG/4v9o-pdb-bundle7_1691_C_A_0_00000.pdb 
# Loading data/PDB/CUUCGG/4v9o-pdb-bundle7_1691_C_A_0_00000.pdb 
# Loaded reference data/PDB/CUUCGG/2KOC_5_C_A_0_00001.pdb 
# Loaded target data/PDB/CUUCGG/5dfe-pdb-bundle1_1691_C_G_0_00000.pdb 
# Loading data/PDB/CUUCGG/5dfe-pdb-bundle1_1691_C_G_0_00000.pdb 
# Loaded reference data/PDB/CUUCGG/2KOC_5_C_A_0_00001.pdb 
# Loaded target data/PDB/CUUCGG/2L2J_19_C_A_0_00019.pdb 
# Loading data/PDB/CUUCGG/2L2J_19_C_A_0_00019.pdb 
# Loaded reference data/PDB/CUUCGG/2KOC_5_C_A_0_00001.pdb 
# Loaded target data/PDB/CUUCGG/5lze-pdb-bundle2_1691_C_A_0_00000.pdb 
# Loading data/PDB/CUUCGG/5lze-pdb-bundle2_1691_C_A_0_00000.pdb 
# Loaded reference data/PDB/CUUCGG/2KOC_5_C_A_0_00001.pdb 
# Loaded target data/PDB/CUUCGG/4u1u-pdb-bundle1_1028_C_A_0_00002.pdb 
# Loading data/PDB/CUUCGG/4u1u-pdb-bundle1_1028_C_A_0_00002.pdb 
# Loaded reference data/PDB/CUUCGG/2KOC_5_C_A_0_00001.pdb 
# Loaded target data/PDB/CUUCGG/4v4h-pdb-bundle3_207_C_A_0_00000.pdb 
# Loading data/PDB/CUUCGG/4v4h-pdb-bundle3_207_C_A_0_00000.pdb 
# Loaded reference data/PDB/CUUCGG/2KOC_5_C_A_0_00001.pdb 
# Loaded target data/PDB/CUUCGG/5j8a-pdb-bundle1_1449_C_b_0_00007.pdb 
# Loading data/PDB/CUUCGG/5j8a-pdb-bundle1_1449_C_b_0_00007.pdb 
# Loaded reference data/PDB/CUUCGG/2KOC_5_C_A_0_00001.pdb 
# Loaded target data/PDB/CUUCGG/1M5L_17_C_A_0_00013.pdb 
# Loading data/PDB/CUUCGG/1M5L_17_C_A_0_00013.pdb 
# Loaded reference data/PDB/CUUCGG/2KOC_5_C_A_0_00001.pdb 
# Loaded target data/PDB/CUUCGG/5j5b-pdb-bundle1_419_C_b_0_00005.pdb 
# Loading data/PDB/CUUCGG/5j5b-pdb-bundle1_419_C_b_0_00005.pdb 
# Loaded reference data/PDB/CUUCGG/2KOC_5_C_A_0_00001.pdb 
# Loaded target data/PDB/CUUCGG/1EKZ_13_C_B_0_00004.pdb 
# Loading data/PDB/CUUCGG/1EKZ_13_C_B_0_00004.pdb 
# Loaded reference data/PDB/CUUCGG/2KOC_5_C_A_0_00001.pdb 
# Loaded target data/PDB/CUUCGG/1EKZ_13_C_B_0_00010.pdb 
# Loading data/PDB/CUUCGG/1EKZ_13_C_B_0_00010.pdb 
# Loaded reference data/PDB/CUUCGG/2KOC_5_C_A_0_00001.pdb 
# Loaded target data/PDB/CUUCGG/1F7I_12_C_A_0_00009.pdb 
# Loading data/PDB/CUUCGG/1F7I_12_C_A_0_00009.pdb 
# Loaded reference data/PDB/CUUCGG/2KOC_5_C_A_0_00001.pdb 
# Loaded target data/PDB/CUUCGG/1M5L_17_C_A_0_00007.pdb 
# Loading data/PDB/CUUCGG/1M5L_17_C_A_0_00007.pdb 
# Loaded reference data/PDB/CUUCGG/2KOC_5_C_A_0_00001.pdb 
# Loaded target data/PDB/CUUCGG/2L2J_19_C_A_0_00009.pdb 
# Loading data/PDB/CUUCGG/2L2J_19_C_A_0_00009.pdb 
# Loaded reference data/PDB/CUUCGG/2KOC_5_C_A_0_00001.pdb 
# Loaded target data/PDB/CUUCGG/1EKZ_13_C_B_0_00028.pdb 
# Loading data/PDB/CUUCGG/1EKZ_13_C_B_0_00028.pdb 
# Loaded reference data/PDB/CUUCGG/2KOC_5_C_A_0_00001.pdb 
# Loaded target data/PDB/CUUCGG/5x8p-pdb-bundle2_977_C_a_0_00000.pdb 
# Loading data/PDB/CUUCGG/5x8p-pdb-bundle2_977_C_a_0_00000.pdb 
# Loaded reference data/PDB/CUUCGG/2KOC_5_C_A_0_00001.pdb 
# Loaded target data/PDB/CUUCGG/5Y87_28_C_D_0_00001.pdb 
# Loading data/PDB/CUUCGG/5Y87_28_C_D_0_00001.pdb 
# Loaded reference data/PDB/CUUCGG/2KOC_5_C_A_0_00001.pdb 
# Loaded target data/PDB/CUUCGG/6CAR_419_C_A_0_00000.pdb 
# Loading data/PDB/CUUCGG/6CAR_419_C_A_0_00000.pdb 
# Loaded reference data/PDB/CUUCGG/2KOC_5_C_A_0_00001.pdb 
# Loaded target data/PDB/CUUCGG/1M5L_17_C_A_0_00003.pdb 
# Loading data/PDB/CUUCGG/1M5L_17_C_A_0_00003.pdb 
# Loaded reference data/PDB/CUUCGG/2KOC_5_C_A_0_00001.pdb 
# Loaded target data/PDB/CUUCGG/1EKZ_13_C_B_0_00014.pdb 
# Loading data/PDB/CUUCGG/1EKZ_13_C_B_0_00014.pdb 
# Loaded reference data/PDB/CUUCGG/2KOC_5_C_A_0_00001.pdb 
# Loaded target data/PDB/CUUCGG/4v9o-pdb-bundle4_1449_C_A_0_00003.pdb 
# Loading data/PDB/CUUCGG/4v9o-pdb-bundle4_1449_C_A_0_00003.pdb 
# Loaded reference data/PDB/CUUCGG/2KOC_5_C_A_0_00001.pdb 
# Loaded target data/PDB/CUUCGG/1EKZ_13_C_B_0_00000.pdb 
# Loading data/PDB/CUUCGG/1EKZ_13_C_B_0_00000.pdb 
# Loaded reference data/PDB/CUUCGG/2KOC_5_C_A_0_00001.pdb 
# Loaded target data/PDB/CUUCGG/5lze-pdb-bundle1_1449_C_a_0_00003.pdb 
# Loading data/PDB/CUUCGG/5lze-pdb-bundle1_1449_C_a_0_00003.pdb 
# Loaded reference data/PDB/CUUCGG/2KOC_5_C_A_0_00001.pdb 
# Loaded target data/PDB/CUUCGG/3CCU_1769_C_0_0_00000.pdb 
# Loading data/PDB/CUUCGG/3CCU_1769_C_0_0_00000.pdb 
# Loaded reference data/PDB/CUUCGG/2KOC_5_C_A_0_00001.pdb 
# Loaded target data/PDB/CUUCGG/1YIJ_1769_C_0_0_00000.pdb 
# Loading data/PDB/CUUCGG/1YIJ_1769_C_0_0_00000.pdb 
# Loaded reference data/PDB/CUUCGG/2KOC_5_C_A_0_00001.pdb 
# Loaded target data/PDB/CUUCGG/1EBQ_12_C_A_0_00004.pdb 
# Loading data/PDB/CUUCGG/1EBQ_12_C_A_0_00004.pdb 
# Loaded reference data/PDB/CUUCGG/2KOC_5_C_A_0_00001.pdb 
# Loaded target data/PDB/CUUCGG/2N6W_32_C_A_0_00008.pdb 
# Loading data/PDB/CUUCGG/2N6W_32_C_A_0_00008.pdb 
# Loaded reference data/PDB/CUUCGG/2KOC_5_C_A_0_00001.pdb 
# Loaded target data/PDB/CUUCGG/5j4c-pdb-bundle2_419_C_A_0_00000.pdb 
# Loading data/PDB/CUUCGG/5j4c-pdb-bundle2_419_C_A_0_00000.pdb 
# Loaded reference data/PDB/CUUCGG/2KOC_5_C_A_0_00001.pdb 
# Loaded target data/PDB/CUUCGG/4u53-pdb-bundle2_1923_C_A_0_00002.pdb 
# Loading data/PDB/CUUCGG/4u53-pdb-bundle2_1923_C_A_0_00002.pdb 
# Loaded reference data/PDB/CUUCGG/2KOC_5_C_A_0_00001.pdb 
# Loaded target data/PDB/CUUCGG/1FYP_11_C_A_0_00021.pdb 
# Loading data/PDB/CUUCGG/1FYP_11_C_A_0_00021.pdb 
# Loaded reference data/PDB/CUUCGG/2KOC_5_C_A_0_00001.pdb 
# Loaded target data/PDB/CUUCGG/5hcr-pdb-bundle3_1737_C_A_0_00000.pdb 
# Loading data/PDB/CUUCGG/5hcr-pdb-bundle3_1737_C_A_0_00000.pdb 
# Loaded reference data/PDB/CUUCGG/2KOC_5_C_A_0_00001.pdb 
# Loaded target data/PDB/CUUCGG/1AUD_28_C_B_0_00009.pdb 
# Loading data/PDB/CUUCGG/1AUD_28_C_B_0_00009.pdb 
# Loaded reference data/PDB/CUUCGG/2KOC_5_C_A_0_00001.pdb 
# Loaded target data/PDB/CUUCGG/4v8f-pdb-bundle1_1691_C_A_0_00000.pdb 
# Loading data/PDB/CUUCGG/4v8f-pdb-bundle1_1691_C_A_0_00000.pdb 
# Loaded reference data/PDB/CUUCGG/2KOC_5_C_A_0_00001.pdb 
# Loaded target data/PDB/CUUCGG/1FYP_11_C_A_0_00009.pdb 
# Loading data/PDB/CUUCGG/1FYP_11_C_A_0_00009.pdb 
# Loaded reference data/PDB/CUUCGG/2KOC_5_C_A_0_00001.pdb 
# Loaded target data/PDB/CUUCGG/4v90-pdb-bundle1_419_C_A_0_00000.pdb 
# Loading data/PDB/CUUCGG/4v90-pdb-bundle1_419_C_A_0_00000.pdb 
# Loaded reference data/PDB/CUUCGG/2KOC_5_C_A_0_00001.pdb 
# Loaded target data/PDB/CUUCGG/5dfe-pdb-bundle2_419_C_a_0_00001.pdb 
# Loading data/PDB/CUUCGG/5dfe-pdb-bundle2_419_C_a_0_00001.pdb 
# Loaded reference data/PDB/CUUCGG/2KOC_5_C_A_0_00001.pdb 
# Loaded target data/PDB/CUUCGG/1AUD_28_C_B_0_00021.pdb 
# Loading data/PDB/CUUCGG/1AUD_28_C_B_0_00021.pdb 
# Loaded reference data/PDB/CUUCGG/2KOC_5_C_A_0_00001.pdb 
# Loaded target data/PDB/CUUCGG/4v8f-pdb-bundle2_419_C_A_0_00000.pdb 
# Loading data/PDB/CUUCGG/4v8f-pdb-bundle2_419_C_A_0_00000.pdb 
# Loaded reference data/PDB/CUUCGG/2KOC_5_C_A_0_00001.pdb 
# Loaded target data/PDB/CUUCGG/1M82_10_C_A_0_00025.pdb 
# Loading data/PDB/CUUCGG/1M82_10_C_A_0_00025.pdb 
# Loaded reference data/PDB/CUUCGG/2KOC_5_C_A_0_00001.pdb 
# Loaded target data/PDB/CUUCGG/4v6c-pdb-bundle3_1028_C_N_0_00002.pdb 
# Loading data/PDB/CUUCGG/4v6c-pdb-bundle3_1028_C_N_0_00002.pdb 
# Loaded reference data/PDB/CUUCGG/2KOC_5_C_A_0_00001.pdb 
# Loaded target data/PDB/CUUCGG/6cae-pdb-bundle2_419_C_A_0_00000.pdb 
# Loading data/PDB/CUUCGG/6cae-pdb-bundle2_419_C_A_0_00000.pdb 
# Loaded reference data/PDB/CUUCGG/2KOC_5_C_A_0_00001.pdb 
# Loaded target data/PDB/CUUCGG/5j5b-pdb-bundle1_1449_C_A_0_00003.pdb 
# Loading data/PDB/CUUCGG/5j5b-pdb-bundle1_1449_C_A_0_00003.pdb 
# Loaded reference data/PDB/CUUCGG/2KOC_5_C_A_0_00001.pdb 
# Loaded target data/PDB/CUUCGG/1P6V_26_C_B_0_00000.pdb 
# Loading data/PDB/CUUCGG/1P6V_26_C_B_0_00000.pdb 
# Loaded reference data/PDB/CUUCGG/2KOC_5_C_A_0_00001.pdb 
# Loaded target data/PDB/CUUCGG/1M82_10_C_A_0_00019.pdb 
# Loading data/PDB/CUUCGG/1M82_10_C_A_0_00019.pdb 
# Loaded reference data/PDB/CUUCGG/2KOC_5_C_A_0_00001.pdb 
# Loaded target data/PDB/CUUCGG/1EBR_12_C_A_0_00000.pdb 
# Loading data/PDB/CUUCGG/1EBR_12_C_A_0_00000.pdb 
# Loaded reference data/PDB/CUUCGG/2KOC_5_C_A_0_00001.pdb 
# Loaded target data/PDB/CUUCGG/4wu1-pdb-bundle1_419_C_A_0_00000.pdb 
# Loading data/PDB/CUUCGG/4wu1-pdb-bundle1_419_C_A_0_00000.pdb 
# Loaded reference data/PDB/CUUCGG/2KOC_5_C_A_0_00001.pdb 
# Loaded target data/PDB/CUUCGG/5IEM_25_C_A_0_00006.pdb 
# Loading data/PDB/CUUCGG/5IEM_25_C_A_0_00006.pdb 
# Loaded reference data/PDB/CUUCGG/2KOC_5_C_A_0_00001.pdb 
# Loaded target data/PDB/CUUCGG/1M1K_1769_C_A_0_00000.pdb 
# Loading data/PDB/CUUCGG/1M1K_1769_C_A_0_00000.pdb 
# Loaded reference data/PDB/CUUCGG/2KOC_5_C_A_0_00001.pdb 
# Loaded target data/PDB/CUUCGG/4wpo-pdb-bundle1_1738_C_A_0_00000.pdb 
# Loading data/PDB/CUUCGG/4wpo-pdb-bundle1_1738_C_A_0_00000.pdb 
# Loaded reference data/PDB/CUUCGG/2KOC_5_C_A_0_00001.pdb 
# Loaded target data/PDB/CUUCGG/4v7s-pdb-bundle2_1691_C_A_0_00000.pdb 
# Loading data/PDB/CUUCGG/4v7s-pdb-bundle2_1691_C_A_0_00000.pdb 
# Loaded reference data/PDB/CUUCGG/2KOC_5_C_A_0_00001.pdb 
# Loaded target data/PDB/CUUCGG/2KMJ_29_C_A_0_00003.pdb 
# Loading data/PDB/CUUCGG/2KMJ_29_C_A_0_00003.pdb 
# Loaded reference data/PDB/CUUCGG/2KOC_5_C_A_0_00001.pdb 
# Loaded target data/PDB/CUUCGG/6ha1-pdb-bundle1_1736_C_A_0_00001.pdb 
# Loading data/PDB/CUUCGG/6ha1-pdb-bundle1_1736_C_A_0_00001.pdb 
# Loaded reference data/PDB/CUUCGG/2KOC_5_C_A_0_00001.pdb 
# Loaded target data/PDB/CUUCGG/2KTZ_19_C_A_0_00001.pdb 
# Loading data/PDB/CUUCGG/2KTZ_19_C_A_0_00001.pdb 
# Loaded reference data/PDB/CUUCGG/2KOC_5_C_A_0_00001.pdb 
# Loaded target data/PDB/CUUCGG/5dgf-pdb-bundle3_230_C_b_0_00000.pdb 
# Loading data/PDB/CUUCGG/5dgf-pdb-bundle3_230_C_b_0_00000.pdb 
# Loaded reference data/PDB/CUUCGG/2KOC_5_C_A_0_00001.pdb 
# Loaded target data/PDB/CUUCGG/6fyy-pdb-bundle1_229_C_2_0_00000.pdb 
# Loading data/PDB/CUUCGG/6fyy-pdb-bundle1_229_C_2_0_00000.pdb 
# Loaded reference data/PDB/CUUCGG/2KOC_5_C_A_0_00001.pdb 
# Loaded target data/PDB/CUUCGG/4v87-pdb-bundle2_419_C_A_0_00000.pdb 
# Loading data/PDB/CUUCGG/4v87-pdb-bundle2_419_C_A_0_00000.pdb 
# Loaded reference data/PDB/CUUCGG/2KOC_5_C_A_0_00001.pdb 
# Loaded target data/PDB/CUUCGG/5wfs-pdb-bundle2_1449_C_a_0_00003.pdb 
# Loading data/PDB/CUUCGG/5wfs-pdb-bundle2_1449_C_a_0_00003.pdb 
# Loaded reference data/PDB/CUUCGG/2KOC_5_C_A_0_00001.pdb 
# Loaded target data/PDB/CUUCGG/4v8x-pdb-bundle1_419_C_A_0_00000.pdb 
# Loading data/PDB/CUUCGG/4v8x-pdb-bundle1_419_C_A_0_00000.pdb 
# Loaded reference data/PDB/CUUCGG/2KOC_5_C_A_0_00001.pdb 
# Loaded target data/PDB/CUUCGG/2L2K_19_C_A_0_00013.pdb 
# Loading data/PDB/CUUCGG/2L2K_19_C_A_0_00013.pdb 
# Loaded reference data/PDB/CUUCGG/2KOC_5_C_A_0_00001.pdb 
# Loaded target data/PDB/CUUCGG/1IKD_7_C_A_0_00010.pdb 
# Loading data/PDB/CUUCGG/1IKD_7_C_A_0_00010.pdb 
# Loaded reference data/PDB/CUUCGG/2KOC_5_C_A_0_00001.pdb 
# Loaded target data/PDB/CUUCGG/5ibb-pdb-bundle2_1691_C_A_0_00000.pdb 
# Loading data/PDB/CUUCGG/5ibb-pdb-bundle2_1691_C_A_0_00000.pdb 
# Loaded reference data/PDB/CUUCGG/2KOC_5_C_A_0_00001.pdb 
# Loaded target data/PDB/CUUCGG/5wfk-pdb-bundle2_419_C_a_0_00001.pdb 
# Loading data/PDB/CUUCGG/5wfk-pdb-bundle2_419_C_a_0_00001.pdb 
# Loaded reference data/PDB/CUUCGG/2KOC_5_C_A_0_00001.pdb 
# Loaded target data/PDB/CUUCGG/2L2K_19_C_A_0_00007.pdb 
# Loading data/PDB/CUUCGG/2L2K_19_C_A_0_00007.pdb 
# Loaded reference data/PDB/CUUCGG/2KOC_5_C_A_0_00001.pdb 
# Loaded target data/PDB/CUUCGG/5vp2-pdb-bundle3_1691_C_A_0_00000.pdb 
# Loading data/PDB/CUUCGG/5vp2-pdb-bundle3_1691_C_A_0_00000.pdb 
# Loaded reference data/PDB/CUUCGG/2KOC_5_C_A_0_00001.pdb 
# Loaded target data/PDB/CUUCGG/4v54-pdb-bundle3_1449_C_K_0_00003.pdb 
# Loading data/PDB/CUUCGG/4v54-pdb-bundle3_1449_C_K_0_00003.pdb 
# Loaded reference data/PDB/CUUCGG/2KOC_5_C_A_0_00001.pdb 
# Loaded target data/PDB/CUUCGG/4woi-pdb-bundle4_207_C_A_0_00000.pdb 
# Loading data/PDB/CUUCGG/4woi-pdb-bundle4_207_C_A_0_00000.pdb 
# Loaded reference data/PDB/CUUCGG/2KOC_5_C_A_0_00001.pdb 
# Loaded target data/PDB/CUUCGG/1F7G_12_C_A_0_00009.pdb 
# Loading data/PDB/CUUCGG/1F7G_12_C_A_0_00009.pdb 
# Loaded reference data/PDB/CUUCGG/2KOC_5_C_A_0_00001.pdb 
# Loaded target data/PDB/CUUCGG/2HGH_32_C_B_0_00008.pdb 
# Loading data/PDB/CUUCGG/2HGH_32_C_B_0_00008.pdb 
# Loaded reference data/PDB/CUUCGG/2KOC_5_C_A_0_00001.pdb 
# Loaded target data/PDB/CUUCGG/4woi-pdb-bundle2_1691_C_A_0_00000.pdb 
# Loading data/PDB/CUUCGG/4woi-pdb-bundle2_1691_C_A_0_00000.pdb 
# Loaded reference data/PDB/CUUCGG/2KOC_5_C_A_0_00001.pdb 
# Loaded target data/PDB/CUUCGG/5lzd-pdb-bundle1_419_C_a_0_00001.pdb 
# Loading data/PDB/CUUCGG/5lzd-pdb-bundle1_419_C_a_0_00001.pdb 
# Loaded reference data/PDB/CUUCGG/2KOC_5_C_A_0_00001.pdb 
# Loaded target data/PDB/CUUCGG/1F7G_12_C_A_0_00008.pdb 
# Loading data/PDB/CUUCGG/1F7G_12_C_A_0_00008.pdb 
# Loaded reference data/PDB/CUUCGG/2KOC_5_C_A_0_00001.pdb 
# Loaded target data/PDB/CUUCGG/2HGH_32_C_B_0_00009.pdb 
# Loading data/PDB/CUUCGG/2HGH_32_C_B_0_00009.pdb 
# Loaded reference data/PDB/CUUCGG/2KOC_5_C_A_0_00001.pdb 
# Loaded target data/PDB/CUUCGG/4yzv-pdb-bundle3_419_C_A_0_00000.pdb 
# Loading data/PDB/CUUCGG/4yzv-pdb-bundle3_419_C_A_0_00000.pdb 
# Loaded reference data/PDB/CUUCGG/2KOC_5_C_A_0_00001.pdb 
# Loaded target data/PDB/CUUCGG/5D5L_60_C_B_0_00001.pdb 
# Loading data/PDB/CUUCGG/5D5L_60_C_B_0_00001.pdb 
# Loaded reference data/PDB/CUUCGG/2KOC_5_C_A_0_00001.pdb 
# Loaded target data/PDB/CUUCGG/5i4l-pdb-bundle2_1923_C_A_0_00002.pdb 
# Loading data/PDB/CUUCGG/5i4l-pdb-bundle2_1923_C_A_0_00002.pdb 
# Loaded reference data/PDB/CUUCGG/2KOC_5_C_A_0_00001.pdb 
# Loaded target data/PDB/CUUCGG/4u4y-pdb-bundle3_230_C_d_0_00000.pdb 
# Loading data/PDB/CUUCGG/4u4y-pdb-bundle3_230_C_d_0_00000.pdb 
# Loaded reference data/PDB/CUUCGG/2KOC_5_C_A_0_00001.pdb 
# Loaded target data/PDB/CUUCGG/4v63-pdb-bundle4_1691_C_A_0_00000.pdb 
# Loading data/PDB/CUUCGG/4v63-pdb-bundle4_1691_C_A_0_00000.pdb 
# Loaded reference data/PDB/CUUCGG/2KOC_5_C_A_0_00001.pdb 
# Loaded target data/PDB/CUUCGG/2L2K_19_C_A_0_00006.pdb 
# Loading data/PDB/CUUCGG/2L2K_19_C_A_0_00006.pdb 
# Loaded reference data/PDB/CUUCGG/2KOC_5_C_A_0_00001.pdb 
# Loaded target data/PDB/CUUCGG/4v7j-pdb-bundle4_1691_C_A_0_00000.pdb 
# Loading data/PDB/CUUCGG/4v7j-pdb-bundle4_1691_C_A_0_00000.pdb 
# Loaded reference data/PDB/CUUCGG/2KOC_5_C_A_0_00001.pdb 
# Loaded target data/PDB/CUUCGG/2L2K_19_C_A_0_00012.pdb 
# Loading data/PDB/CUUCGG/2L2K_19_C_A_0_00012.pdb 
# Loaded reference data/PDB/CUUCGG/2KOC_5_C_A_0_00001.pdb 
# Loaded target data/PDB/CUUCGG/6mtd-pdb-bundle1_2872_C_5_0_00000.pdb 
# Loading data/PDB/CUUCGG/6mtd-pdb-bundle1_2872_C_5_0_00000.pdb 
# Loaded reference data/PDB/CUUCGG/2KOC_5_C_A_0_00001.pdb 
# Loaded target data/PDB/CUUCGG/5vpo-pdb-bundle2_1691_C_A_0_00000.pdb 
# Loading data/PDB/CUUCGG/5vpo-pdb-bundle2_1691_C_A_0_00000.pdb 
# Loaded reference data/PDB/CUUCGG/2KOC_5_C_A_0_00001.pdb 
# Loaded target data/PDB/CUUCGG/4v9o-pdb-bundle1_1691_C_C_0_00000.pdb 
# Loading data/PDB/CUUCGG/4v9o-pdb-bundle1_1691_C_C_0_00000.pdb 
# Loaded reference data/PDB/CUUCGG/2KOC_5_C_A_0_00001.pdb 
# Loaded target data/PDB/CUUCGG/4v54-pdb-bundle4_1691_C_A_0_00000.pdb 
# Loading data/PDB/CUUCGG/4v54-pdb-bundle4_1691_C_A_0_00000.pdb 
# Loaded reference data/PDB/CUUCGG/2KOC_5_C_A_0_00001.pdb 
# Loaded target data/PDB/CUUCGG/2KMJ_29_C_A_0_00002.pdb 
# Loading data/PDB/CUUCGG/2KMJ_29_C_A_0_00002.pdb 
# Loaded reference data/PDB/CUUCGG/2KOC_5_C_A_0_00001.pdb 
# Loaded target data/PDB/CUUCGG/2KTZ_19_C_A_0_00000.pdb 
# Loading data/PDB/CUUCGG/2KTZ_19_C_A_0_00000.pdb 
# Loaded reference data/PDB/CUUCGG/2KOC_5_C_A_0_00001.pdb 
# Loaded target data/PDB/CUUCGG/2M4Q_11_C_1_0_00008.pdb 
# Loading data/PDB/CUUCGG/2M4Q_11_C_1_0_00008.pdb 
# Loaded reference data/PDB/CUUCGG/2KOC_5_C_A_0_00001.pdb 
# Loaded target data/PDB/CUUCGG/4v9p-pdb-bundle7_1691_C_A_0_00000.pdb 
# Loading data/PDB/CUUCGG/4v9p-pdb-bundle7_1691_C_A_0_00000.pdb 
# Loaded reference data/PDB/CUUCGG/2KOC_5_C_A_0_00001.pdb 
# Loaded target data/PDB/CUUCGG/4wra-pdb-bundle2_1691_C_A_0_00000.pdb 
# Loading data/PDB/CUUCGG/4wra-pdb-bundle2_1691_C_A_0_00000.pdb 
# Loaded reference data/PDB/CUUCGG/2KOC_5_C_A_0_00001.pdb 
# Loaded target data/PDB/CUUCGG/5j4c-pdb-bundle1_1738_C_A_0_00000.pdb 
# Loading data/PDB/CUUCGG/5j4c-pdb-bundle1_1738_C_A_0_00000.pdb 
# Loaded reference data/PDB/CUUCGG/2KOC_5_C_A_0_00001.pdb 
# Loaded target data/PDB/CUUCGG/5IEM_25_C_A_0_00007.pdb 
# Loading data/PDB/CUUCGG/5IEM_25_C_A_0_00007.pdb 
# Loaded reference data/PDB/CUUCGG/2KOC_5_C_A_0_00001.pdb 
# Loaded target data/PDB/CUUCGG/1EBR_12_C_A_0_00001.pdb 
# Loading data/PDB/CUUCGG/1EBR_12_C_A_0_00001.pdb 
# Loaded reference data/PDB/CUUCGG/2KOC_5_C_A_0_00001.pdb 
# Loaded target data/PDB/CUUCGG/1M82_10_C_A_0_00018.pdb 
# Loading data/PDB/CUUCGG/1M82_10_C_A_0_00018.pdb 
# Loaded reference data/PDB/CUUCGG/2KOC_5_C_A_0_00001.pdb 
# Loaded target data/PDB/CUUCGG/5uq7-pdb-bundle2_419_C_a_0_00000.pdb 
# Loading data/PDB/CUUCGG/5uq7-pdb-bundle2_419_C_a_0_00000.pdb 
# Loaded reference data/PDB/CUUCGG/2KOC_5_C_A_0_00001.pdb 
# Loaded target data/PDB/CUUCGG/4u1u-pdb-bundle4_1691_C_A_0_00000.pdb 
# Loading data/PDB/CUUCGG/4u1u-pdb-bundle4_1691_C_A_0_00000.pdb 
# Loaded reference data/PDB/CUUCGG/2KOC_5_C_A_0_00001.pdb 
# Loaded target data/PDB/CUUCGG/6enu-pdb-bundle2_1691_C_A_0_00000.pdb 
# Loading data/PDB/CUUCGG/6enu-pdb-bundle2_1691_C_A_0_00000.pdb 
# Loaded reference data/PDB/CUUCGG/2KOC_5_C_A_0_00001.pdb 
# Loaded target data/PDB/CUUCGG/1M82_10_C_A_0_00024.pdb 
# Loading data/PDB/CUUCGG/1M82_10_C_A_0_00024.pdb 
# Loaded reference data/PDB/CUUCGG/2KOC_5_C_A_0_00001.pdb 
# Loaded target data/PDB/CUUCGG/4v7v-pdb-bundle3_419_C_A_0_00001.pdb 
# Loading data/PDB/CUUCGG/4v7v-pdb-bundle3_419_C_A_0_00001.pdb 
# Loaded reference data/PDB/CUUCGG/2KOC_5_C_A_0_00001.pdb 
# Loaded target data/PDB/CUUCGG/1M82_10_C_A_0_00030.pdb 
# Loading data/PDB/CUUCGG/1M82_10_C_A_0_00030.pdb 
# Loaded reference data/PDB/CUUCGG/2KOC_5_C_A_0_00001.pdb 
# Loaded target data/PDB/CUUCGG/1AUD_28_C_B_0_00020.pdb 
# Loading data/PDB/CUUCGG/1AUD_28_C_B_0_00020.pdb 
# Loaded reference data/PDB/CUUCGG/2KOC_5_C_A_0_00001.pdb 
# Loaded target data/PDB/CUUCGG/4v52-pdb-bundle1_1449_C_A_0_00003.pdb 
# Loading data/PDB/CUUCGG/4v52-pdb-bundle1_1449_C_A_0_00003.pdb 
# Loaded reference data/PDB/CUUCGG/2KOC_5_C_A_0_00001.pdb 
# Loaded target data/PDB/CUUCGG/1FYP_11_C_A_0_00008.pdb 
# Loading data/PDB/CUUCGG/1FYP_11_C_A_0_00008.pdb 
# Loaded reference data/PDB/CUUCGG/2KOC_5_C_A_0_00001.pdb 
# Loaded target data/PDB/CUUCGG/4v57-pdb-bundle3_1449_C_A_0_00003.pdb 
# Loading data/PDB/CUUCGG/4v57-pdb-bundle3_1449_C_A_0_00003.pdb 
# Loaded reference data/PDB/CUUCGG/2KOC_5_C_A_0_00001.pdb 
# Loaded target data/PDB/CUUCGG/4v5l-pdb-bundle2_1691_C_A_0_00000.pdb 
# Loading data/PDB/CUUCGG/4v5l-pdb-bundle2_1691_C_A_0_00000.pdb 
# Loaded reference data/PDB/CUUCGG/2KOC_5_C_A_0_00001.pdb 
# Loaded target data/PDB/CUUCGG/6eri-pdb-bundle2_978_C_A_0_00000.pdb 
# Loading data/PDB/CUUCGG/6eri-pdb-bundle2_978_C_A_0_00000.pdb 
# Loaded reference data/PDB/CUUCGG/2KOC_5_C_A_0_00001.pdb 
# Loaded target data/PDB/CUUCGG/6nd6-pdb-bundle2_419_C_A_0_00000.pdb 
# Loading data/PDB/CUUCGG/6nd6-pdb-bundle2_419_C_A_0_00000.pdb 
# Loaded reference data/PDB/CUUCGG/2KOC_5_C_A_0_00001.pdb 
# Loaded target data/PDB/CUUCGG/1FYP_11_C_A_0_00020.pdb 
# Loading data/PDB/CUUCGG/1FYP_11_C_A_0_00020.pdb 
# Loaded reference data/PDB/CUUCGG/2KOC_5_C_A_0_00001.pdb 
# Loaded target data/PDB/CUUCGG/1F7F_12_C_A_0_00006.pdb 
# Loading data/PDB/CUUCGG/1F7F_12_C_A_0_00006.pdb 
# Loaded reference data/PDB/CUUCGG/2KOC_5_C_A_0_00001.pdb 
# Loaded target data/PDB/CUUCGG/4wf1-pdb-bundle3_419_C_A_0_00001.pdb 
# Loading data/PDB/CUUCGG/4wf1-pdb-bundle3_419_C_A_0_00001.pdb 
# Loaded reference data/PDB/CUUCGG/2KOC_5_C_A_0_00001.pdb 
# Loaded target data/PDB/CUUCGG/4v7s-pdb-bundle1_419_C_A_0_00001.pdb 
# Loading data/PDB/CUUCGG/4v7s-pdb-bundle1_419_C_A_0_00001.pdb 
# Loaded reference data/PDB/CUUCGG/2KOC_5_C_A_0_00001.pdb 
# Loaded target data/PDB/CUUCGG/4u51-pdb-bundle3_230_C_d_0_00000.pdb 
# Loading data/PDB/CUUCGG/4u51-pdb-bundle3_230_C_d_0_00000.pdb 
# Loaded reference data/PDB/CUUCGG/2KOC_5_C_A_0_00001.pdb 
# Loaded target data/PDB/CUUCGG/2N6W_32_C_A_0_00009.pdb 
# Loading data/PDB/CUUCGG/2N6W_32_C_A_0_00009.pdb 
# Loaded reference data/PDB/CUUCGG/2KOC_5_C_A_0_00001.pdb 
# Loaded target data/PDB/CUUCGG/6cfl-pdb-bundle3_1691_C_A_0_00000.pdb 
# Loading data/PDB/CUUCGG/6cfl-pdb-bundle3_1691_C_A_0_00000.pdb 
# Loaded reference data/PDB/CUUCGG/2KOC_5_C_A_0_00001.pdb 
# Loaded target data/PDB/CUUCGG/5it8-pdb-bundle1_419_C_b_0_00005.pdb 
# Loading data/PDB/CUUCGG/5it8-pdb-bundle1_419_C_b_0_00005.pdb 
# Loaded reference data/PDB/CUUCGG/2KOC_5_C_A_0_00001.pdb 
# Loaded target data/PDB/CUUCGG/1EKZ_13_C_B_0_00001.pdb 
# Loading data/PDB/CUUCGG/1EKZ_13_C_B_0_00001.pdb 
# Loaded reference data/PDB/CUUCGG/2KOC_5_C_A_0_00001.pdb 
# Loaded target data/PDB/CUUCGG/1M5L_17_C_A_0_00002.pdb 
# Loading data/PDB/CUUCGG/1M5L_17_C_A_0_00002.pdb 
# Loaded reference data/PDB/CUUCGG/2KOC_5_C_A_0_00001.pdb 
# Loaded target data/PDB/CUUCGG/4wq1-pdb-bundle3_419_C_A_0_00000.pdb 
# Loading data/PDB/CUUCGG/4wq1-pdb-bundle3_419_C_A_0_00000.pdb 
# Loaded reference data/PDB/CUUCGG/2KOC_5_C_A_0_00001.pdb 
# Loaded target data/PDB/CUUCGG/1EKZ_13_C_B_0_00015.pdb 
# Loading data/PDB/CUUCGG/1EKZ_13_C_B_0_00015.pdb 
# Loaded reference data/PDB/CUUCGG/2KOC_5_C_A_0_00001.pdb 
# Loaded target data/PDB/CUUCGG/4v7u-pdb-bundle1_1028_C_A_0_00002.pdb 
# Loading data/PDB/CUUCGG/4v7u-pdb-bundle1_1028_C_A_0_00002.pdb 
# Loaded reference data/PDB/CUUCGG/2KOC_5_C_A_0_00001.pdb 
# Loaded target data/PDB/CUUCGG/1EKZ_13_C_B_0_00029.pdb 
# Loading data/PDB/CUUCGG/1EKZ_13_C_B_0_00029.pdb 
# Loaded reference data/PDB/CUUCGG/2KOC_5_C_A_0_00001.pdb 
# Loaded target data/PDB/CUUCGG/2L2J_19_C_A_0_00008.pdb 
# Loading data/PDB/CUUCGG/2L2J_19_C_A_0_00008.pdb 
# Loaded reference data/PDB/CUUCGG/2KOC_5_C_A_0_00001.pdb 
# Loaded target data/PDB/CUUCGG/4LFB_419_C_A_0_00000.pdb 
# Loading data/PDB/CUUCGG/4LFB_419_C_A_0_00000.pdb 
# Loaded reference data/PDB/CUUCGG/2KOC_5_C_A_0_00001.pdb 
# Loaded target data/PDB/CUUCGG/1M5L_17_C_A_0_00014.pdb 
# Loading data/PDB/CUUCGG/1M5L_17_C_A_0_00014.pdb 
# Loaded reference data/PDB/CUUCGG/2KOC_5_C_A_0_00001.pdb 
# Loaded target data/PDB/CUUCGG/4w2h-pdb-bundle4_1691_C_A_0_00000.pdb 
# Loading data/PDB/CUUCGG/4w2h-pdb-bundle4_1691_C_A_0_00000.pdb 
# Loaded reference data/PDB/CUUCGG/2KOC_5_C_A_0_00001.pdb 
# Loaded target data/PDB/CUUCGG/1EKZ_13_C_B_0_00003.pdb 
# Loading data/PDB/CUUCGG/1EKZ_13_C_B_0_00003.pdb 
# Loaded reference data/PDB/CUUCGG/2KOC_5_C_A_0_00001.pdb 
# Loaded target data/PDB/CUUCGG/1M5L_17_C_A_0_00000.pdb 
# Loading data/PDB/CUUCGG/1M5L_17_C_A_0_00000.pdb 
# Loaded reference data/PDB/CUUCGG/2KOC_5_C_A_0_00001.pdb 
# Loaded target data/PDB/CUUCGG/6DLS_27_C_A_0_00000.pdb 
# Loading data/PDB/CUUCGG/6DLS_27_C_A_0_00000.pdb 
# Loaded reference data/PDB/CUUCGG/2KOC_5_C_A_0_00001.pdb 
# Loaded target data/PDB/CUUCGG/4v84-pdb-bundle4_1691_C_A_0_00000.pdb 
# Loading data/PDB/CUUCGG/4v84-pdb-bundle4_1691_C_A_0_00000.pdb 
# Loaded reference data/PDB/CUUCGG/2KOC_5_C_A_0_00001.pdb 
# Loaded target data/PDB/CUUCGG/4v6a-pdb-bundle4_1691_C_A_0_00000.pdb 
# Loading data/PDB/CUUCGG/4v6a-pdb-bundle4_1691_C_A_0_00000.pdb 
# Loaded reference data/PDB/CUUCGG/2KOC_5_C_A_0_00001.pdb 
# Loaded target data/PDB/CUUCGG/4B3M_419_C_A_0_00000.pdb 
# Loading data/PDB/CUUCGG/4B3M_419_C_A_0_00000.pdb 
# Loaded reference data/PDB/CUUCGG/2KOC_5_C_A_0_00001.pdb 
# Loaded target data/PDB/CUUCGG/4u4y-pdb-bundle2_1923_C_A_0_00002.pdb 
# Loading data/PDB/CUUCGG/4u4y-pdb-bundle2_1923_C_A_0_00002.pdb 
# Loaded reference data/PDB/CUUCGG/2KOC_5_C_A_0_00001.pdb 
# Loaded target data/PDB/CUUCGG/4wsd-pdb-bundle4_1691_C_A_0_00000.pdb 
# Loading data/PDB/CUUCGG/4wsd-pdb-bundle4_1691_C_A_0_00000.pdb 
# Loaded reference data/PDB/CUUCGG/2KOC_5_C_A_0_00001.pdb 
# Loaded target data/PDB/CUUCGG/1vy5-pdb-bundle2_1691_C_A_0_00000.pdb 
# Loading data/PDB/CUUCGG/1vy5-pdb-bundle2_1691_C_A_0_00000.pdb 
# Loaded reference data/PDB/CUUCGG/2KOC_5_C_A_0_00001.pdb 
# Loaded target data/PDB/CUUCGG/5w4k-pdb-bundle2_419_C_A_0_00000.pdb 
# Loading data/PDB/CUUCGG/5w4k-pdb-bundle2_419_C_A_0_00000.pdb 
# Loaded reference data/PDB/CUUCGG/2KOC_5_C_A_0_00001.pdb 
# Loaded target data/PDB/CUUCGG/5mdz-pdb-bundle1_419_C_2_0_00002.pdb 
# Loading data/PDB/CUUCGG/5mdz-pdb-bundle1_419_C_2_0_00002.pdb 
# Loaded reference data/PDB/CUUCGG/2KOC_5_C_A_0_00001.pdb 
# Loaded target data/PDB/CUUCGG/4v7j-pdb-bundle1_419_C_V_0_00000.pdb 
# Loading data/PDB/CUUCGG/4v7j-pdb-bundle1_419_C_V_0_00000.pdb 
# Loaded reference data/PDB/CUUCGG/2KOC_5_C_A_0_00001.pdb 
# Loaded target data/PDB/CUUCGG/1FYP_11_C_A_0_00022.pdb 
# Loading data/PDB/CUUCGG/1FYP_11_C_A_0_00022.pdb 
# Loaded reference data/PDB/CUUCGG/2KOC_5_C_A_0_00001.pdb 
# Loaded target data/PDB/CUUCGG/5iqr-pdb-bundle2_1449_C_2_0_00004.pdb 
# Loading data/PDB/CUUCGG/5iqr-pdb-bundle2_1449_C_2_0_00004.pdb 
# Loaded reference data/PDB/CUUCGG/2KOC_5_C_A_0_00001.pdb 
# Loaded target data/PDB/CUUCGG/4v5l-pdb-bundle1_419_C_A_0_00000.pdb 
# Loading data/PDB/CUUCGG/4v5l-pdb-bundle1_419_C_A_0_00000.pdb 
# Loaded reference data/PDB/CUUCGG/2KOC_5_C_A_0_00001.pdb 
# Loaded target data/PDB/CUUCGG/4KHP_415_C_A_0_00000.pdb 
# Loading data/PDB/CUUCGG/4KHP_415_C_A_0_00000.pdb 
# Loaded reference data/PDB/CUUCGG/2KOC_5_C_A_0_00001.pdb 
# Loaded target data/PDB/CUUCGG/1AUD_28_C_B_0_00022.pdb 
# Loading data/PDB/CUUCGG/1AUD_28_C_B_0_00022.pdb 
# Loaded reference data/PDB/CUUCGG/2KOC_5_C_A_0_00001.pdb 
# Loaded target data/PDB/CUUCGG/4v7t-pdb-bundle3_1028_C_A_0_00002.pdb 
# Loading data/PDB/CUUCGG/4v7t-pdb-bundle3_1028_C_A_0_00002.pdb 
# Loaded reference data/PDB/CUUCGG/2KOC_5_C_A_0_00001.pdb 
# Loaded target data/PDB/CUUCGG/1M82_10_C_A_0_00026.pdb 
# Loading data/PDB/CUUCGG/1M82_10_C_A_0_00026.pdb 
# Loaded reference data/PDB/CUUCGG/2KOC_5_C_A_0_00001.pdb 
# Loaded target data/PDB/CUUCGG/4v8x-pdb-bundle3_419_C_A_0_00000.pdb 
# Loading data/PDB/CUUCGG/4v8x-pdb-bundle3_419_C_A_0_00000.pdb 
# Loaded reference data/PDB/CUUCGG/2KOC_5_C_A_0_00001.pdb 
# Loaded target data/PDB/CUUCGG/6b4v-pdb-bundle1_419_C_A_0_00000.pdb 
# Loading data/PDB/CUUCGG/6b4v-pdb-bundle1_419_C_A_0_00000.pdb 
# Loaded reference data/PDB/CUUCGG/2KOC_5_C_A_0_00001.pdb 
# Loaded target data/PDB/CUUCGG/4v64-pdb-bundle3_1449_C_A_0_00003.pdb 
# Loading data/PDB/CUUCGG/4v64-pdb-bundle3_1449_C_A_0_00003.pdb 
# Loaded reference data/PDB/CUUCGG/2KOC_5_C_A_0_00001.pdb 
# Loaded target data/PDB/CUUCGG/4v9c-pdb-bundle2_1691_C_A_0_00000.pdb 
# Loading data/PDB/CUUCGG/4v9c-pdb-bundle2_1691_C_A_0_00000.pdb 
# Loaded reference data/PDB/CUUCGG/2KOC_5_C_A_0_00001.pdb 
# Loaded target data/PDB/CUUCGG/5IEM_25_C_A_0_00005.pdb 
# Loading data/PDB/CUUCGG/5IEM_25_C_A_0_00005.pdb 
# Loaded reference data/PDB/CUUCGG/2KOC_5_C_A_0_00001.pdb 
# Loaded target data/PDB/CUUCGG/5fdu-pdb-bundle3_1691_C_A_0_00000.pdb 
# Loading data/PDB/CUUCGG/5fdu-pdb-bundle3_1691_C_A_0_00000.pdb 
# Loaded reference data/PDB/CUUCGG/2KOC_5_C_A_0_00001.pdb 
# Loaded target data/PDB/CUUCGG/1EBR_12_C_A_0_00003.pdb 
# Loading data/PDB/CUUCGG/1EBR_12_C_A_0_00003.pdb 
# Loaded reference data/PDB/CUUCGG/2KOC_5_C_A_0_00001.pdb 
# Loaded target data/PDB/CUUCGG/6DCC_11_C_B_0_00000.pdb 
# Loading data/PDB/CUUCGG/6DCC_11_C_B_0_00000.pdb 
# Loaded reference data/PDB/CUUCGG/2KOC_5_C_A_0_00001.pdb 
# Loaded target data/PDB/CUUCGG/4v67-pdb-bundle4_1691_C_A_0_00000.pdb 
# Loading data/PDB/CUUCGG/4v67-pdb-bundle4_1691_C_A_0_00000.pdb 
# Loaded reference data/PDB/CUUCGG/2KOC_5_C_A_0_00001.pdb 
# Loaded target data/PDB/CUUCGG/4v8b-pdb-bundle4_1691_C_A_0_00000.pdb 
# Loading data/PDB/CUUCGG/4v8b-pdb-bundle4_1691_C_A_0_00000.pdb 
# Loaded reference data/PDB/CUUCGG/2KOC_5_C_A_0_00001.pdb 
# Loaded target data/PDB/CUUCGG/2KTZ_19_C_A_0_00002.pdb 
# Loading data/PDB/CUUCGG/2KTZ_19_C_A_0_00002.pdb 
# Loaded reference data/PDB/CUUCGG/2KOC_5_C_A_0_00001.pdb 
# Loaded target data/PDB/CUUCGG/6b4v-pdb-bundle2_419_C_0_0_00000.pdb 
# Loading data/PDB/CUUCGG/6b4v-pdb-bundle2_419_C_0_0_00000.pdb 
# Loaded reference data/PDB/CUUCGG/2KOC_5_C_A_0_00001.pdb 
# Loaded target data/PDB/CUUCGG/2KMJ_29_C_A_0_00000.pdb 
# Loading data/PDB/CUUCGG/2KMJ_29_C_A_0_00000.pdb 
# Loaded reference data/PDB/CUUCGG/2KOC_5_C_A_0_00001.pdb 
# Loaded target data/PDB/CUUCGG/4ybb-pdb-bundle2_1691_C_I_0_00000.pdb 
# Loading data/PDB/CUUCGG/4ybb-pdb-bundle2_1691_C_I_0_00000.pdb 
# Loaded reference data/PDB/CUUCGG/2KOC_5_C_A_0_00001.pdb 
# Loaded target data/PDB/CUUCGG/2KOC_5_C_A_0_00019.pdb 
# Loading data/PDB/CUUCGG/2KOC_5_C_A_0_00019.pdb 
# Loaded reference data/PDB/CUUCGG/2KOC_5_C_A_0_00001.pdb 
# Loaded target data/PDB/CUUCGG/5wdt-pdb-bundle2_419_C_a_0_00001.pdb 
# Loading data/PDB/CUUCGG/5wdt-pdb-bundle2_419_C_a_0_00001.pdb 
# Loaded reference data/PDB/CUUCGG/2KOC_5_C_A_0_00001.pdb 
# Loaded target data/PDB/CUUCGG/4u4r-pdb-bundle4_1923_C_I_0_00002.pdb 
# Loading data/PDB/CUUCGG/4u4r-pdb-bundle4_1923_C_I_0_00002.pdb 
# Loaded reference data/PDB/CUUCGG/2KOC_5_C_A_0_00001.pdb 
# Loaded target data/PDB/CUUCGG/4wsd-pdb-bundle3_419_C_A_0_00000.pdb 
# Loading data/PDB/CUUCGG/4wsd-pdb-bundle3_419_C_A_0_00000.pdb 
# Loaded reference data/PDB/CUUCGG/2KOC_5_C_A_0_00001.pdb 
# Loaded target data/PDB/CUUCGG/4v95-pdb-bundle2_1691_C_A_0_00000.pdb 
# Loading data/PDB/CUUCGG/4v95-pdb-bundle2_1691_C_A_0_00000.pdb 
# Loaded reference data/PDB/CUUCGG/2KOC_5_C_A_0_00001.pdb 
# Loaded target data/PDB/CUUCGG/6hhq-pdb-bundle2_1923_C_e_0_00002.pdb 
# Loading data/PDB/CUUCGG/6hhq-pdb-bundle2_1923_C_e_0_00002.pdb 
# Loaded reference data/PDB/CUUCGG/2KOC_5_C_A_0_00001.pdb 
# Loaded target data/PDB/CUUCGG/1IKD_7_C_A_0_00007.pdb 
# Loading data/PDB/CUUCGG/1IKD_7_C_A_0_00007.pdb 
# Loaded reference data/PDB/CUUCGG/2KOC_5_C_A_0_00001.pdb 
# Loaded target data/PDB/CUUCGG/2L2K_19_C_A_0_00004.pdb 
# Loading data/PDB/CUUCGG/2L2K_19_C_A_0_00004.pdb 
# Loaded reference data/PDB/CUUCGG/2KOC_5_C_A_0_00001.pdb 
# Loaded target data/PDB/CUUCGG/1IKD_7_C_A_0_00013.pdb 
# Loading data/PDB/CUUCGG/1IKD_7_C_A_0_00013.pdb 
# Loaded reference data/PDB/CUUCGG/2KOC_5_C_A_0_00001.pdb 
# Loaded target data/PDB/CUUCGG/2L2K_19_C_A_0_00010.pdb 
# Loading data/PDB/CUUCGG/2L2K_19_C_A_0_00010.pdb 
# Loaded reference data/PDB/CUUCGG/2KOC_5_C_A_0_00001.pdb 
# Loaded target data/PDB/CUUCGG/5el6-pdb-bundle3_419_C_A_0_00000.pdb 
# Loading data/PDB/CUUCGG/5el6-pdb-bundle3_419_C_A_0_00000.pdb 
# Loaded reference data/PDB/CUUCGG/2KOC_5_C_A_0_00001.pdb 
# Loaded target data/PDB/CUUCGG/5obm-pdb-bundle3_1923_C_B_0_00002.pdb 
# Loading data/PDB/CUUCGG/5obm-pdb-bundle3_1923_C_B_0_00002.pdb 
# Loaded reference data/PDB/CUUCGG/2KOC_5_C_A_0_00001.pdb 
# Loaded target data/PDB/CUUCGG/5f8k-pdb-bundle1_1738_C_A_0_00000.pdb 
# Loading data/PDB/CUUCGG/5f8k-pdb-bundle1_1738_C_A_0_00000.pdb 
# Loaded reference data/PDB/CUUCGG/2KOC_5_C_A_0_00001.pdb 
# Loaded target data/PDB/CUUCGG/1YKV_220_C_B_0_00000.pdb 
# Loading data/PDB/CUUCGG/1YKV_220_C_B_0_00000.pdb 
# Loaded reference data/PDB/CUUCGG/2KOC_5_C_A_0_00001.pdb 
# Loaded target data/PDB/CUUCGG/5j91-pdb-bundle2_1691_C_U_0_00000.pdb 
# Loading data/PDB/CUUCGG/5j91-pdb-bundle2_1691_C_U_0_00000.pdb 
# Loaded reference data/PDB/CUUCGG/2KOC_5_C_A_0_00001.pdb 
# Loaded target data/PDB/CUUCGG/4v50-pdb-bundle4_1691_C_A_0_00000.pdb 
# Loading data/PDB/CUUCGG/4v50-pdb-bundle4_1691_C_A_0_00000.pdb 
# Loaded reference data/PDB/CUUCGG/2KOC_5_C_A_0_00001.pdb 
# Loaded target data/PDB/CUUCGG/5el4-pdb-bundle2_1691_C_A_0_00000.pdb 
# Loading data/PDB/CUUCGG/5el4-pdb-bundle2_1691_C_A_0_00000.pdb 
# Loaded reference data/PDB/CUUCGG/2KOC_5_C_A_0_00001.pdb 
# Loaded target data/PDB/CUUCGG/4X62_419_C_A_0_00000.pdb 
# Loading data/PDB/CUUCGG/4X62_419_C_A_0_00000.pdb 
# Loaded reference data/PDB/CUUCGG/2KOC_5_C_A_0_00001.pdb 
# Loaded target data/PDB/CUUCGG/6I0Y_1691_C_A_0_00000.pdb 
# Loading data/PDB/CUUCGG/6I0Y_1691_C_A_0_00000.pdb 
# Loaded reference data/PDB/CUUCGG/2KOC_5_C_A_0_00001.pdb 
# Loaded target data/PDB/CUUCGG/4v7s-pdb-bundle3_419_C_A_0_00001.pdb 
# Loading data/PDB/CUUCGG/4v7s-pdb-bundle3_419_C_A_0_00001.pdb 
# Loaded reference data/PDB/CUUCGG/2KOC_5_C_A_0_00001.pdb 
# Loaded target data/PDB/CUUCGG/4v7w-pdb-bundle2_1691_C_A_0_00000.pdb 
# Loading data/PDB/CUUCGG/4v7w-pdb-bundle2_1691_C_A_0_00000.pdb 
# Loaded reference data/PDB/CUUCGG/2KOC_5_C_A_0_00001.pdb 
# Loaded target data/PDB/CUUCGG/1C0O_5_C_A_0_00018.pdb 
# Loading data/PDB/CUUCGG/1C0O_5_C_A_0_00018.pdb 
# Loaded reference data/PDB/CUUCGG/2KOC_5_C_A_0_00001.pdb 
# Loaded target data/PDB/CUUCGG/4wf1-pdb-bundle1_419_C_A_0_00001.pdb 
# Loading data/PDB/CUUCGG/4wf1-pdb-bundle1_419_C_A_0_00001.pdb 
# Loaded reference data/PDB/CUUCGG/2KOC_5_C_A_0_00001.pdb 
# Loaded target data/PDB/CUUCGG/2KZL_27_C_A_0_00005.pdb 
# Loading data/PDB/CUUCGG/2KZL_27_C_A_0_00005.pdb 
# Loaded reference data/PDB/CUUCGG/2KOC_5_C_A_0_00001.pdb 
# Loaded target data/PDB/CUUCGG/4u6f-pdb-bundle2_1923_C_A_0_00002.pdb 
# Loading data/PDB/CUUCGG/4u6f-pdb-bundle2_1923_C_A_0_00002.pdb 
# Loaded reference data/PDB/CUUCGG/2KOC_5_C_A_0_00001.pdb 
# Loaded target data/PDB/CUUCGG/3I56_1769_C_0_0_00000.pdb 
# Loading data/PDB/CUUCGG/3I56_1769_C_0_0_00000.pdb 
# Loaded reference data/PDB/CUUCGG/2KOC_5_C_A_0_00001.pdb 
# Loaded target data/PDB/CUUCGG/4wq1-pdb-bundle1_419_C_A_0_00000.pdb 
# Loading data/PDB/CUUCGG/4wq1-pdb-bundle1_419_C_A_0_00000.pdb 
# Loaded reference data/PDB/CUUCGG/2KOC_5_C_A_0_00001.pdb 
# Loaded target data/PDB/CUUCGG/2L2K_19_C_A_0_00011.pdb 
# Loading data/PDB/CUUCGG/2L2K_19_C_A_0_00011.pdb 
# Loaded reference data/PDB/CUUCGG/2KOC_5_C_A_0_00001.pdb 
# Loaded target data/PDB/CUUCGG/1IKD_7_C_A_0_00012.pdb 
# Loading data/PDB/CUUCGG/1IKD_7_C_A_0_00012.pdb 
# Loaded reference data/PDB/CUUCGG/2KOC_5_C_A_0_00001.pdb 
# Loaded target data/PDB/CUUCGG/2L2K_19_C_A_0_00005.pdb 
# Loading data/PDB/CUUCGG/2L2K_19_C_A_0_00005.pdb 
# Loaded reference data/PDB/CUUCGG/2KOC_5_C_A_0_00001.pdb 
# Loaded target data/PDB/CUUCGG/1IKD_7_C_A_0_00006.pdb 
# Loading data/PDB/CUUCGG/1IKD_7_C_A_0_00006.pdb 
# Loaded reference data/PDB/CUUCGG/2KOC_5_C_A_0_00001.pdb 
# Loaded target data/PDB/CUUCGG/5tbw-pdb-bundle2_1923_C_e_0_00002.pdb 
# Loading data/PDB/CUUCGG/5tbw-pdb-bundle2_1923_C_e_0_00002.pdb 
# Loaded reference data/PDB/CUUCGG/2KOC_5_C_A_0_00001.pdb 
# Loaded target data/PDB/CUUCGG/2KOC_5_C_A_0_00018.pdb 
# Loading data/PDB/CUUCGG/2KOC_5_C_A_0_00018.pdb 
# Loaded reference data/PDB/CUUCGG/2KOC_5_C_A_0_00001.pdb 
# Loaded target data/PDB/CUUCGG/4v7v-pdb-bundle1_419_C_A_0_00001.pdb 
# Loading data/PDB/CUUCGG/4v7v-pdb-bundle1_419_C_A_0_00001.pdb 
# Loaded reference data/PDB/CUUCGG/2KOC_5_C_A_0_00001.pdb 
# Loaded target data/PDB/CUUCGG/4u6f-pdb-bundle3_230_C_d_0_00000.pdb 
# Loading data/PDB/CUUCGG/4u6f-pdb-bundle3_230_C_d_0_00000.pdb 
# Loaded reference data/PDB/CUUCGG/2KOC_5_C_A_0_00001.pdb 
# Loaded target data/PDB/CUUCGG/2KTZ_19_C_A_0_00003.pdb 
# Loading data/PDB/CUUCGG/2KTZ_19_C_A_0_00003.pdb 
# Loaded reference data/PDB/CUUCGG/2KOC_5_C_A_0_00001.pdb 
# Loaded target data/PDB/CUUCGG/2KMJ_29_C_A_0_00001.pdb 
# Loading data/PDB/CUUCGG/2KMJ_29_C_A_0_00001.pdb 
# Loaded reference data/PDB/CUUCGG/2KOC_5_C_A_0_00001.pdb 
# Loaded target data/PDB/CUUCGG/4v88-pdb-bundle3_230_C_d_0_00000.pdb 
# Loading data/PDB/CUUCGG/4v88-pdb-bundle3_230_C_d_0_00000.pdb 
# Loaded reference data/PDB/CUUCGG/2KOC_5_C_A_0_00001.pdb 
# Loaded target data/PDB/CUUCGG/4v7y-pdb-bundle4_1691_C_A_0_00000.pdb 
# Loading data/PDB/CUUCGG/4v7y-pdb-bundle4_1691_C_A_0_00000.pdb 
# Loaded reference data/PDB/CUUCGG/2KOC_5_C_A_0_00001.pdb 
# Loaded target data/PDB/CUUCGG/4IO9_1708_C_X_0_00000.pdb 
# Loading data/PDB/CUUCGG/4IO9_1708_C_X_0_00000.pdb 
# Loaded reference data/PDB/CUUCGG/2KOC_5_C_A_0_00001.pdb 
# Loaded target data/PDB/CUUCGG/1EBR_12_C_A_0_00002.pdb 
# Loading data/PDB/CUUCGG/1EBR_12_C_A_0_00002.pdb 
# Loaded reference data/PDB/CUUCGG/2KOC_5_C_A_0_00001.pdb 
# Loaded target data/PDB/CUUCGG/6DMC_57_C_B_0_00001.pdb 
# Loading data/PDB/CUUCGG/6DMC_57_C_B_0_00001.pdb 
# Loaded reference data/PDB/CUUCGG/2KOC_5_C_A_0_00001.pdb 
# Loaded target data/PDB/CUUCGG/5IEM_25_C_A_0_00004.pdb 
# Loading data/PDB/CUUCGG/5IEM_25_C_A_0_00004.pdb 
# Loaded reference data/PDB/CUUCGG/2KOC_5_C_A_0_00001.pdb 
# Loaded target data/PDB/CUUCGG/4v8c-pdb-bundle3_419_C_A_0_00000.pdb 
# Loading data/PDB/CUUCGG/4v8c-pdb-bundle3_419_C_A_0_00000.pdb 
# Loaded reference data/PDB/CUUCGG/2KOC_5_C_A_0_00001.pdb 
# Loaded target data/PDB/CUUCGG/6DME_57_C_A_0_00000.pdb 
# Loading data/PDB/CUUCGG/6DME_57_C_A_0_00000.pdb 
# Loaded reference data/PDB/CUUCGG/2KOC_5_C_A_0_00001.pdb 
# Loaded target data/PDB/CUUCGG/4wf1-pdb-bundle1_1449_C_A_0_00003.pdb 
# Loading data/PDB/CUUCGG/4wf1-pdb-bundle1_1449_C_A_0_00003.pdb 
# Loaded reference data/PDB/CUUCGG/2KOC_5_C_A_0_00001.pdb 
# Loaded target data/PDB/CUUCGG/4v5q-pdb-bundle4_1691_C_A_0_00000.pdb 
# Loading data/PDB/CUUCGG/4v5q-pdb-bundle4_1691_C_A_0_00000.pdb 
# Loaded reference data/PDB/CUUCGG/2KOC_5_C_A_0_00001.pdb 
# Loaded target data/PDB/CUUCGG/1M82_10_C_A_0_00027.pdb 
# Loading data/PDB/CUUCGG/1M82_10_C_A_0_00027.pdb 
# Loaded reference data/PDB/CUUCGG/2KOC_5_C_A_0_00001.pdb 
# Loaded target data/PDB/CUUCGG/1XSG_12_C_A_0_00000.pdb 
# Loading data/PDB/CUUCGG/1XSG_12_C_A_0_00000.pdb 
# Loaded reference data/PDB/CUUCGG/2KOC_5_C_A_0_00001.pdb 
# Loaded target data/PDB/CUUCGG/4v88-pdb-bundle4_1923_C_K_0_00002.pdb 
# Loading data/PDB/CUUCGG/4v88-pdb-bundle4_1923_C_K_0_00002.pdb 
# Loaded reference data/PDB/CUUCGG/2KOC_5_C_A_0_00001.pdb 
# Loaded target data/PDB/CUUCGG/1AUD_28_C_B_0_00023.pdb 
# Loading data/PDB/CUUCGG/1AUD_28_C_B_0_00023.pdb 
# Loaded reference data/PDB/CUUCGG/2KOC_5_C_A_0_00001.pdb 
# Loaded target data/PDB/CUUCGG/1FYP_11_C_A_0_00023.pdb 
# Loading data/PDB/CUUCGG/1FYP_11_C_A_0_00023.pdb 
# Loaded reference data/PDB/CUUCGG/2KOC_5_C_A_0_00001.pdb 
# Loaded target data/PDB/CUUCGG/5wis-pdb-bundle4_419_C_A_0_00000.pdb 
# Loading data/PDB/CUUCGG/5wis-pdb-bundle4_419_C_A_0_00000.pdb 
# Loaded reference data/PDB/CUUCGG/2KOC_5_C_A_0_00001.pdb 
# Loaded target data/PDB/CUUCGG/4ybb-pdb-bundle1_1449_C_V_0_00007.pdb 
# Loading data/PDB/CUUCGG/4ybb-pdb-bundle1_1449_C_V_0_00007.pdb 
# Loaded reference data/PDB/CUUCGG/2KOC_5_C_A_0_00001.pdb 
# Loaded target data/PDB/CUUCGG/4yzv-pdb-bundle1_419_C_A_0_00000.pdb 
# Loading data/PDB/CUUCGG/4yzv-pdb-bundle1_419_C_A_0_00000.pdb 
# Loaded reference data/PDB/CUUCGG/2KOC_5_C_A_0_00001.pdb 
# Loaded target data/PDB/CUUCGG/1F7F_12_C_A_0_00005.pdb 
# Loading data/PDB/CUUCGG/1F7F_12_C_A_0_00005.pdb 
# Loaded reference data/PDB/CUUCGG/2KOC_5_C_A_0_00001.pdb 
# Loaded target data/PDB/CUUCGG/1EKZ_13_C_B_0_00016.pdb 
# Loading data/PDB/CUUCGG/1EKZ_13_C_B_0_00016.pdb 
# Loaded reference data/PDB/CUUCGG/2KOC_5_C_A_0_00001.pdb 
# Loaded target data/PDB/CUUCGG/4u4n-pdb-bundle2_1923_C_A_0_00002.pdb 
# Loading data/PDB/CUUCGG/4u4n-pdb-bundle2_1923_C_A_0_00002.pdb 
# Loaded reference data/PDB/CUUCGG/2KOC_5_C_A_0_00001.pdb 
# Loaded target data/PDB/CUUCGG/1M5L_17_C_A_0_00001.pdb 
# Loading data/PDB/CUUCGG/1M5L_17_C_A_0_00001.pdb 
# Loaded reference data/PDB/CUUCGG/2KOC_5_C_A_0_00001.pdb 
# Loaded target data/PDB/CUUCGG/4v9d-pdb-bundle1_419_C_A_0_00001.pdb 
# Loading data/PDB/CUUCGG/4v9d-pdb-bundle1_419_C_A_0_00001.pdb 
# Loaded reference data/PDB/CUUCGG/2KOC_5_C_A_0_00001.pdb 
# Loaded target data/PDB/CUUCGG/1EKZ_13_C_B_0_00002.pdb 
# Loading data/PDB/CUUCGG/1EKZ_13_C_B_0_00002.pdb 
# Loaded reference data/PDB/CUUCGG/2KOC_5_C_A_0_00001.pdb 
# Loaded target data/PDB/CUUCGG/3CME_1769_C_0_0_00000.pdb 
# Loading data/PDB/CUUCGG/3CME_1769_C_0_0_00000.pdb 
# Loaded reference data/PDB/CUUCGG/2KOC_5_C_A_0_00001.pdb 
# Loaded target data/PDB/CUUCGG/6enf-pdb-bundle2_1691_C_A_0_00000.pdb 
# Loading data/PDB/CUUCGG/6enf-pdb-bundle2_1691_C_A_0_00000.pdb 
# Loaded reference data/PDB/CUUCGG/2KOC_5_C_A_0_00001.pdb 
# Loaded target data/PDB/CUUCGG/5czp-pdb-bundle2_419_C_V_0_00001.pdb 
# Loading data/PDB/CUUCGG/5czp-pdb-bundle2_419_C_V_0_00001.pdb 
# Loaded reference data/PDB/CUUCGG/2KOC_5_C_A_0_00001.pdb 
# Loaded target data/PDB/CUUCGG/4v8e-pdb-bundle2_419_C_A_0_00000.pdb 
# Loading data/PDB/CUUCGG/4v8e-pdb-bundle2_419_C_A_0_00000.pdb 
# Loaded reference data/PDB/CUUCGG/2KOC_5_C_A_0_00001.pdb 
# Loaded target data/PDB/CUUCGG/4tue-pdb-bundle2_1691_C_A_0_00000.pdb 
# Loading data/PDB/CUUCGG/4tue-pdb-bundle2_1691_C_A_0_00000.pdb 
# Loaded reference data/PDB/CUUCGG/2KOC_5_C_A_0_00001.pdb 
# Loaded target data/PDB/CUUCGG/2N3R_37_C_A_0_00011.pdb 
# Loading data/PDB/CUUCGG/2N3R_37_C_A_0_00011.pdb 
# Loaded reference data/PDB/CUUCGG/2KOC_5_C_A_0_00001.pdb 
# Loaded target data/PDB/CUUCGG/2L3E_18_C_A_0_00016.pdb 
# Loading data/PDB/CUUCGG/2L3E_18_C_A_0_00016.pdb 
# Loaded reference data/PDB/CUUCGG/2KOC_5_C_A_0_00001.pdb 
# Loaded target data/PDB/CUUCGG/5v8i-pdb-bundle3_1691_C_A_0_00000.pdb 
# Loading data/PDB/CUUCGG/5v8i-pdb-bundle3_1691_C_A_0_00000.pdb 
# Loaded reference data/PDB/CUUCGG/2KOC_5_C_A_0_00001.pdb 
# Loaded target data/PDB/CUUCGG/2L3E_18_C_A_0_00002.pdb 
# Loading data/PDB/CUUCGG/2L3E_18_C_A_0_00002.pdb 
# Loaded reference data/PDB/CUUCGG/2KOC_5_C_A_0_00001.pdb 
# Loaded target data/PDB/CUUCGG/2N3R_37_C_A_0_00005.pdb 
# Loading data/PDB/CUUCGG/2N3R_37_C_A_0_00005.pdb 
# Loaded reference data/PDB/CUUCGG/2KOC_5_C_A_0_00001.pdb 
# Loaded target data/PDB/CUUCGG/4u20-pdb-bundle4_1691_C_A_0_00000.pdb 
# Loading data/PDB/CUUCGG/4u20-pdb-bundle4_1691_C_A_0_00000.pdb 
# Loaded reference data/PDB/CUUCGG/2KOC_5_C_A_0_00001.pdb 
# Loaded target data/PDB/CUUCGG/3j7p-pdb-bundle1_2872_C_5_0_00000.pdb 
# Loading data/PDB/CUUCGG/3j7p-pdb-bundle1_2872_C_5_0_00000.pdb 
# Loaded reference data/PDB/CUUCGG/2KOC_5_C_A_0_00001.pdb 
# Loaded target data/PDB/CUUCGG/4lnt-pdb-bundle4_1691_C_A_0_00000.pdb 
# Loading data/PDB/CUUCGG/4lnt-pdb-bundle4_1691_C_A_0_00000.pdb 
# Loaded reference data/PDB/CUUCGG/2KOC_5_C_A_0_00001.pdb 
# Loaded target data/PDB/CUUCGG/5j88-pdb-bundle2_1691_C_T_0_00000.pdb 
# Loading data/PDB/CUUCGG/5j88-pdb-bundle2_1691_C_T_0_00000.pdb 
# Loaded reference data/PDB/CUUCGG/2KOC_5_C_A_0_00001.pdb 
# Loaded target data/PDB/CUUCGG/4www-pdb-bundle2_1449_C_V_0_00007.pdb 
# Loading data/PDB/CUUCGG/4www-pdb-bundle2_1449_C_V_0_00007.pdb 
# Loaded reference data/PDB/CUUCGG/2KOC_5_C_A_0_00001.pdb 
# Loaded target data/PDB/CUUCGG/4v9o-pdb-bundle3_1691_C_A_0_00000.pdb 
# Loading data/PDB/CUUCGG/4v9o-pdb-bundle3_1691_C_A_0_00000.pdb 
# Loaded reference data/PDB/CUUCGG/2KOC_5_C_A_0_00001.pdb 
# Loaded target data/PDB/CUUCGG/2UXD_419_C_A_0_00000.pdb 
# Loading data/PDB/CUUCGG/2UXD_419_C_A_0_00000.pdb 
# Loaded reference data/PDB/CUUCGG/2KOC_5_C_A_0_00001.pdb 
# Loaded target data/PDB/CUUCGG/4v8q-pdb-bundle1_1691_C_K_0_00000.pdb 
# Loading data/PDB/CUUCGG/4v8q-pdb-bundle1_1691_C_K_0_00000.pdb 
# Loaded reference data/PDB/CUUCGG/2KOC_5_C_A_0_00001.pdb 
# Loaded target data/PDB/CUUCGG/4v6f-pdb-bundle1_1691_C_A_0_00000.pdb 
# Loading data/PDB/CUUCGG/4v6f-pdb-bundle1_1691_C_A_0_00000.pdb 
# Loaded reference data/PDB/CUUCGG/2KOC_5_C_A_0_00001.pdb 
# Loaded target data/PDB/CUUCGG/5el5-pdb-bundle1_419_C_A_0_00000.pdb 
# Loading data/PDB/CUUCGG/5el5-pdb-bundle1_419_C_A_0_00000.pdb 
# Loaded reference data/PDB/CUUCGG/2KOC_5_C_A_0_00001.pdb 
# Loaded target data/PDB/CUUCGG/4wqr-pdb-bundle2_1691_C_A_0_00000.pdb 
# Loading data/PDB/CUUCGG/4wqr-pdb-bundle2_1691_C_A_0_00000.pdb 
# Loaded reference data/PDB/CUUCGG/2KOC_5_C_A_0_00001.pdb 
# Loaded target data/PDB/CUUCGG/2GIO_13_C_A_0_00008.pdb 
# Loading data/PDB/CUUCGG/2GIO_13_C_A_0_00008.pdb 
# Loaded reference data/PDB/CUUCGG/2KOC_5_C_A_0_00001.pdb 
# Loaded target data/PDB/CUUCGG/5h5u-pdb-bundle2_1449_C_h_0_00003.pdb 
# Loading data/PDB/CUUCGG/5h5u-pdb-bundle2_1449_C_h_0_00003.pdb 
# Loaded reference data/PDB/CUUCGG/2KOC_5_C_A_0_00001.pdb 
# Loaded target data/PDB/CUUCGG/4PMI_53_C_A_0_00000.pdb 
# Loading data/PDB/CUUCGG/4PMI_53_C_A_0_00000.pdb 
# Loaded reference data/PDB/CUUCGG/2KOC_5_C_A_0_00001.pdb 
# Loaded target data/PDB/CUUCGG/1A3M_11_C_A_0_00016.pdb 
# Loading data/PDB/CUUCGG/1A3M_11_C_A_0_00016.pdb 
# Loaded reference data/PDB/CUUCGG/2KOC_5_C_A_0_00001.pdb 
# Loaded target data/PDB/CUUCGG/1A3M_11_C_A_0_00002.pdb 
# Loading data/PDB/CUUCGG/1A3M_11_C_A_0_00002.pdb 
# Loaded reference data/PDB/CUUCGG/2KOC_5_C_A_0_00001.pdb 
# Loaded target data/PDB/CUUCGG/4lsk-pdb-bundle4_1691_C_A_0_00000.pdb 
# Loading data/PDB/CUUCGG/4lsk-pdb-bundle4_1691_C_A_0_00000.pdb 
# Loaded reference data/PDB/CUUCGG/2KOC_5_C_A_0_00001.pdb 
# Loaded target data/PDB/CUUCGG/2LQZ_13_C_A_0_00000.pdb 
# Loading data/PDB/CUUCGG/2LQZ_13_C_A_0_00000.pdb 
# Loaded reference data/PDB/CUUCGG/2KOC_5_C_A_0_00001.pdb 
# Loaded target data/PDB/CUUCGG/1BYJ_11_C_A_0_00019.pdb 
# Loading data/PDB/CUUCGG/1BYJ_11_C_A_0_00019.pdb 
# Loaded reference data/PDB/CUUCGG/2KOC_5_C_A_0_00001.pdb 
# Loaded target data/PDB/CUUCGG/6buw-pdb-bundle2_1691_C_A_0_00000.pdb 
# Loading data/PDB/CUUCGG/6buw-pdb-bundle2_1691_C_A_0_00000.pdb 
# Loaded reference data/PDB/CUUCGG/2KOC_5_C_A_0_00001.pdb 
# Loaded target data/PDB/CUUCGG/2HUA_17_C_A_0_00001.pdb 
# Loading data/PDB/CUUCGG/2HUA_17_C_A_0_00001.pdb 
# Loaded reference data/PDB/CUUCGG/2KOC_5_C_A_0_00001.pdb 
# Loaded target data/PDB/CUUCGG/1B36_16_C_A_0_00004.pdb 
# Loading data/PDB/CUUCGG/1B36_16_C_A_0_00004.pdb 
# Loaded reference data/PDB/CUUCGG/2KOC_5_C_A_0_00001.pdb 
# Loaded target data/PDB/CUUCGG/2LHP_16_C_A_0_00004.pdb 
# Loading data/PDB/CUUCGG/2LHP_16_C_A_0_00004.pdb 
# Loaded reference data/PDB/CUUCGG/2KOC_5_C_A_0_00001.pdb 
# Loaded target data/PDB/CUUCGG/4v6c-pdb-bundle1_207_C_U_0_00000.pdb 
# Loading data/PDB/CUUCGG/4v6c-pdb-bundle1_207_C_U_0_00000.pdb 
# Loaded reference data/PDB/CUUCGG/2KOC_5_C_A_0_00001.pdb 
# Loaded target data/PDB/CUUCGG/2M58_12_C_A_0_00001.pdb 
# Loading data/PDB/CUUCGG/2M58_12_C_A_0_00001.pdb 
# Loaded reference data/PDB/CUUCGG/2KOC_5_C_A_0_00001.pdb 
# Loaded target data/PDB/CUUCGG/1BYJ_11_C_A_0_00025.pdb 
# Loading data/PDB/CUUCGG/1BYJ_11_C_A_0_00025.pdb 
# Loaded reference data/PDB/CUUCGG/2KOC_5_C_A_0_00001.pdb 
# Loaded target data/PDB/CUUCGG/4v9p-pdb-bundle6_1028_C_A_0_00002.pdb 
# Loading data/PDB/CUUCGG/4v9p-pdb-bundle6_1028_C_A_0_00002.pdb 
# Loaded reference data/PDB/CUUCGG/2KOC_5_C_A_0_00001.pdb 
# Loaded target data/PDB/CUUCGG/2N3Q_37_C_A_0_00015.pdb 
# Loading data/PDB/CUUCGG/2N3Q_37_C_A_0_00015.pdb 
# Loaded reference data/PDB/CUUCGG/2KOC_5_C_A_0_00001.pdb 
# Loaded target data/PDB/CUUCGG/1FMN_16_C_A_0_00003.pdb 
# Loading data/PDB/CUUCGG/1FMN_16_C_A_0_00003.pdb 
# Loaded reference data/PDB/CUUCGG/2KOC_5_C_A_0_00001.pdb 
# Loaded target data/PDB/CUUCGG/4v4h-pdb-bundle1_419_C_A_0_00001.pdb 
# Loading data/PDB/CUUCGG/4v4h-pdb-bundle1_419_C_A_0_00001.pdb 
# Loaded reference data/PDB/CUUCGG/2KOC_5_C_A_0_00001.pdb 
# Loaded target data/PDB/CUUCGG/2N3Q_37_C_A_0_00001.pdb 
# Loading data/PDB/CUUCGG/2N3Q_37_C_A_0_00001.pdb 
# Loaded reference data/PDB/CUUCGG/2KOC_5_C_A_0_00001.pdb 
# Loaded target data/PDB/CUUCGG/1BYJ_11_C_A_0_00031.pdb 
# Loading data/PDB/CUUCGG/1BYJ_11_C_A_0_00031.pdb 
# Loaded reference data/PDB/CUUCGG/2KOC_5_C_A_0_00001.pdb 
# Loaded target data/PDB/CUUCGG/2NCI_14_C_A_0_00006.pdb 
# Loading data/PDB/CUUCGG/2NCI_14_C_A_0_00006.pdb 
# Loaded reference data/PDB/CUUCGG/2KOC_5_C_A_0_00001.pdb 
# Loaded target data/PDB/CUUCGG/1D6K_283_C_B_0_00015.pdb 
# Loading data/PDB/CUUCGG/1D6K_283_C_B_0_00015.pdb 
# Loaded reference data/PDB/CUUCGG/2KOC_5_C_A_0_00001.pdb 
# Loaded target data/PDB/CUUCGG/5zep-pdb-bundle1_419_C_a_0_00000.pdb 
# Loading data/PDB/CUUCGG/5zep-pdb-bundle1_419_C_a_0_00000.pdb 
# Loaded reference data/PDB/CUUCGG/2KOC_5_C_A_0_00001.pdb 
# Loaded target data/PDB/CUUCGG/1D6K_283_C_B_0_00001.pdb 
# Loading data/PDB/CUUCGG/1D6K_283_C_B_0_00001.pdb 
# Loaded reference data/PDB/CUUCGG/2KOC_5_C_A_0_00001.pdb 
# Loaded target data/PDB/CUUCGG/4y4p-pdb-bundle1_1738_C_A_0_00000.pdb 
# Loading data/PDB/CUUCGG/4y4p-pdb-bundle1_1738_C_A_0_00000.pdb 
# Loaded reference data/PDB/CUUCGG/2KOC_5_C_A_0_00001.pdb 
# Loaded target data/PDB/CUUCGG/5ib7-pdb-bundle2_1691_C_A_0_00000.pdb 
# Loading data/PDB/CUUCGG/5ib7-pdb-bundle2_1691_C_A_0_00000.pdb 
# Loaded reference data/PDB/CUUCGG/2KOC_5_C_A_0_00001.pdb 
# Loaded target data/PDB/CUUCGG/2LWK_15_C_A_0_00003.pdb 
# Loading data/PDB/CUUCGG/2LWK_15_C_A_0_00003.pdb 
# Loaded reference data/PDB/CUUCGG/2KOC_5_C_A_0_00001.pdb 
# Loaded target data/PDB/CUUCGG/4u26-pdb-bundle1_1449_C_A_0_00003.pdb 
# Loading data/PDB/CUUCGG/4u26-pdb-bundle1_1449_C_A_0_00003.pdb 
# Loaded reference data/PDB/CUUCGG/2KOC_5_C_A_0_00001.pdb 
# Loaded target data/PDB/CUUCGG/5f8k-pdb-bundle4_419_C_A_0_00000.pdb 
# Loading data/PDB/CUUCGG/5f8k-pdb-bundle4_419_C_A_0_00000.pdb 
# Loaded reference data/PDB/CUUCGG/2KOC_5_C_A_0_00001.pdb 
# Loaded target data/PDB/CUUCGG/4v50-pdb-bundle1_207_C_A_0_00000.pdb 
# Loading data/PDB/CUUCGG/4v50-pdb-bundle1_207_C_A_0_00000.pdb 
# Loaded reference data/PDB/CUUCGG/2KOC_5_C_A_0_00001.pdb 
# Loaded target data/PDB/CUUCGG/2LUB_16_C_A_0_00001.pdb 
# Loading data/PDB/CUUCGG/2LUB_16_C_A_0_00001.pdb 
# Loaded reference data/PDB/CUUCGG/2KOC_5_C_A_0_00001.pdb 
# Loaded target data/PDB/CUUCGG/4v8e-pdb-bundle1_1691_C_A_0_00000.pdb 
# Loading data/PDB/CUUCGG/4v8e-pdb-bundle1_1691_C_A_0_00000.pdb 
# Loaded reference data/PDB/CUUCGG/2KOC_5_C_A_0_00001.pdb 
# Loaded target data/PDB/CUUCGG/4ypb-pdb-bundle3_419_C_A_0_00000.pdb 
# Loading data/PDB/CUUCGG/4ypb-pdb-bundle3_419_C_A_0_00000.pdb 
# Loaded reference data/PDB/CUUCGG/2KOC_5_C_A_0_00001.pdb 
# Loaded target data/PDB/CUUCGG/2N6S_16_C_A_0_00000.pdb 
# Loading data/PDB/CUUCGG/2N6S_16_C_A_0_00000.pdb 
# Loaded reference data/PDB/CUUCGG/2KOC_5_C_A_0_00001.pdb 
# Loaded target data/PDB/CUUCGG/4z3s-pdb-bundle4_419_C_A_0_00000.pdb 
# Loading data/PDB/CUUCGG/4z3s-pdb-bundle4_419_C_A_0_00000.pdb 
# Loaded reference data/PDB/CUUCGG/2KOC_5_C_A_0_00001.pdb 
# Loaded target data/PDB/CUUCGG/4u50-pdb-bundle2_1923_C_A_0_00002.pdb 
# Loading data/PDB/CUUCGG/4u50-pdb-bundle2_1923_C_A_0_00002.pdb 
# Loaded reference data/PDB/CUUCGG/2KOC_5_C_A_0_00001.pdb 
# Loaded target data/PDB/CUUCGG/4v9b-pdb-bundle1_419_C_A_0_00000.pdb 
# Loading data/PDB/CUUCGG/4v9b-pdb-bundle1_419_C_A_0_00000.pdb 
# Loaded reference data/PDB/CUUCGG/2KOC_5_C_A_0_00001.pdb 
# Loaded target data/PDB/CUUCGG/5hcq-pdb-bundle3_1737_C_A_0_00000.pdb 
# Loading data/PDB/CUUCGG/5hcq-pdb-bundle3_1737_C_A_0_00000.pdb 
# Loaded reference data/PDB/CUUCGG/2KOC_5_C_A_0_00001.pdb 
# Loaded target data/PDB/CUUCGG/5we6-pdb-bundle2_1449_C_a_0_00003.pdb 
# Loading data/PDB/CUUCGG/5we6-pdb-bundle2_1449_C_a_0_00003.pdb 
# Loaded reference data/PDB/CUUCGG/2KOC_5_C_A_0_00001.pdb 
# Loaded target data/PDB/CUUCGG/4v9o-pdb-bundle6_419_C_A_0_00001.pdb 
# Loading data/PDB/CUUCGG/4v9o-pdb-bundle6_419_C_A_0_00001.pdb 
# Loaded reference data/PDB/CUUCGG/2KOC_5_C_A_0_00001.pdb 
# Loaded target data/PDB/CUUCGG/6fkr-pdb-bundle3_1691_C_A_0_00000.pdb 
# Loading data/PDB/CUUCGG/6fkr-pdb-bundle3_1691_C_A_0_00000.pdb 
# Loaded reference data/PDB/CUUCGG/2KOC_5_C_A_0_00001.pdb 
# Loaded target data/PDB/CUUCGG/2N6S_16_C_A_0_00001.pdb 
# Loading data/PDB/CUUCGG/2N6S_16_C_A_0_00001.pdb 
# Loaded reference data/PDB/CUUCGG/2KOC_5_C_A_0_00001.pdb 
# Loaded target data/PDB/CUUCGG/4v4h-pdb-bundle1_1449_C_A_0_00003.pdb 
# Loading data/PDB/CUUCGG/4v4h-pdb-bundle1_1449_C_A_0_00003.pdb 
# Loaded reference data/PDB/CUUCGG/2KOC_5_C_A_0_00001.pdb 
# Loaded target data/PDB/CUUCGG/1FFK_1769_C_0_0_00000.pdb 
# Loading data/PDB/CUUCGG/1FFK_1769_C_0_0_00000.pdb 
# Loaded reference data/PDB/CUUCGG/2KOC_5_C_A_0_00001.pdb 
# Loaded target data/PDB/CUUCGG/4lnt-pdb-bundle1_419_C_A_0_00000.pdb 
# Loading data/PDB/CUUCGG/4lnt-pdb-bundle1_419_C_A_0_00000.pdb 
# Loaded reference data/PDB/CUUCGG/2KOC_5_C_A_0_00001.pdb 
# Loaded target data/PDB/CUUCGG/4v83-pdb-bundle4_1691_C_A_0_00000.pdb 
# Loading data/PDB/CUUCGG/4v83-pdb-bundle4_1691_C_A_0_00000.pdb 
# Loaded reference data/PDB/CUUCGG/2KOC_5_C_A_0_00001.pdb 
# Loaded target data/PDB/CUUCGG/4u1v-pdb-bundle4_1691_C_A_0_00000.pdb 
# Loading data/PDB/CUUCGG/4u1v-pdb-bundle4_1691_C_A_0_00000.pdb 
# Loaded reference data/PDB/CUUCGG/2KOC_5_C_A_0_00001.pdb 
# Loaded target data/PDB/CUUCGG/2LWK_15_C_A_0_00002.pdb 
# Loading data/PDB/CUUCGG/2LWK_15_C_A_0_00002.pdb 
# Loaded reference data/PDB/CUUCGG/2KOC_5_C_A_0_00001.pdb 
# Loaded target data/PDB/CUUCGG/3CD6_1769_C_0_0_00000.pdb 
# Loading data/PDB/CUUCGG/3CD6_1769_C_0_0_00000.pdb 
# Loaded reference data/PDB/CUUCGG/2KOC_5_C_A_0_00001.pdb 
# Loaded target data/PDB/CUUCGG/1F79_12_C_A_0_00000.pdb 
# Loading data/PDB/CUUCGG/1F79_12_C_A_0_00000.pdb 
# Loaded reference data/PDB/CUUCGG/2KOC_5_C_A_0_00001.pdb 
# Loaded target data/PDB/CUUCGG/4v4q-pdb-bundle1_1028_C_A_0_00002.pdb 
# Loading data/PDB/CUUCGG/4v4q-pdb-bundle1_1028_C_A_0_00002.pdb 
# Loaded reference data/PDB/CUUCGG/2KOC_5_C_A_0_00001.pdb 
# Loaded target data/PDB/CUUCGG/1D6K_283_C_B_0_00000.pdb 
# Loading data/PDB/CUUCGG/1D6K_283_C_B_0_00000.pdb 
# Loaded reference data/PDB/CUUCGG/2KOC_5_C_A_0_00001.pdb 
# Loaded target data/PDB/CUUCGG/5wfs-pdb-bundle2_419_C_a_0_00001.pdb 
# Loading data/PDB/CUUCGG/5wfs-pdb-bundle2_419_C_a_0_00001.pdb 
# Loaded reference data/PDB/CUUCGG/2KOC_5_C_A_0_00001.pdb 
# Loaded target data/PDB/CUUCGG/4v9q-pdb-bundle4_419_C_A_0_00000.pdb 
# Loading data/PDB/CUUCGG/4v9q-pdb-bundle4_419_C_A_0_00000.pdb 
# Loaded reference data/PDB/CUUCGG/2KOC_5_C_A_0_00001.pdb 
# Loaded target data/PDB/CUUCGG/4v5q-pdb-bundle1_419_C_A_0_00000.pdb 
# Loading data/PDB/CUUCGG/4v5q-pdb-bundle1_419_C_A_0_00000.pdb 
# Loaded reference data/PDB/CUUCGG/2KOC_5_C_A_0_00001.pdb 
# Loaded target data/PDB/CUUCGG/2NCI_14_C_A_0_00007.pdb 
# Loading data/PDB/CUUCGG/2NCI_14_C_A_0_00007.pdb 
# Loaded reference data/PDB/CUUCGG/2KOC_5_C_A_0_00001.pdb 
# Loaded target data/PDB/CUUCGG/5u4i-pdb-bundle2_1449_C_a_0_00003.pdb 
# Loading data/PDB/CUUCGG/5u4i-pdb-bundle2_1449_C_a_0_00003.pdb 
# Loaded reference data/PDB/CUUCGG/2KOC_5_C_A_0_00001.pdb 
# Loaded target data/PDB/CUUCGG/1D6K_283_C_B_0_00014.pdb 
# Loading data/PDB/CUUCGG/1D6K_283_C_B_0_00014.pdb 
# Loaded reference data/PDB/CUUCGG/2KOC_5_C_A_0_00001.pdb 
# Loaded target data/PDB/CUUCGG/5jc9-pdb-bundle1_1449_C_b_0_00007.pdb 
# Loading data/PDB/CUUCGG/5jc9-pdb-bundle1_1449_C_b_0_00007.pdb 
# Loaded reference data/PDB/CUUCGG/2KOC_5_C_A_0_00001.pdb 
# Loaded target data/PDB/CUUCGG/4wqy-pdb-bundle3_1691_C_A_0_00000.pdb 
# Loading data/PDB/CUUCGG/4wqy-pdb-bundle3_1691_C_A_0_00000.pdb 
# Loaded reference data/PDB/CUUCGG/2KOC_5_C_A_0_00001.pdb 
# Loaded target data/PDB/CUUCGG/1VQ7_1769_C_0_0_00000.pdb 
# Loading data/PDB/CUUCGG/1VQ7_1769_C_0_0_00000.pdb 
# Loaded reference data/PDB/CUUCGG/2KOC_5_C_A_0_00001.pdb 
# Loaded target data/PDB/CUUCGG/4v7v-pdb-bundle1_1028_C_A_0_00002.pdb 
# Loading data/PDB/CUUCGG/4v7v-pdb-bundle1_1028_C_A_0_00002.pdb 
# Loaded reference data/PDB/CUUCGG/2KOC_5_C_A_0_00001.pdb 
# Loaded target data/PDB/CUUCGG/4w2i-pdb-bundle2_1738_C_A_0_00000.pdb 
# Loading data/PDB/CUUCGG/4w2i-pdb-bundle2_1738_C_A_0_00000.pdb 
# Loaded reference data/PDB/CUUCGG/2KOC_5_C_A_0_00001.pdb 
# Loaded target data/PDB/CUUCGG/1BYJ_11_C_A_0_00030.pdb 
# Loading data/PDB/CUUCGG/1BYJ_11_C_A_0_00030.pdb 
# Loaded reference data/PDB/CUUCGG/2KOC_5_C_A_0_00001.pdb 
# Loaded target data/PDB/CUUCGG/5mgp-pdb-bundle1_1691_C_A_0_00000.pdb 
# Loading data/PDB/CUUCGG/5mgp-pdb-bundle1_1691_C_A_0_00000.pdb 
# Loaded reference data/PDB/CUUCGG/2KOC_5_C_A_0_00001.pdb 
# Loaded target data/PDB/CUUCGG/2N3Q_37_C_A_0_00000.pdb 
# Loading data/PDB/CUUCGG/2N3Q_37_C_A_0_00000.pdb 
# Loaded reference data/PDB/CUUCGG/2KOC_5_C_A_0_00001.pdb 
# Loaded target data/PDB/CUUCGG/2M58_12_C_A_0_00000.pdb 
# Loading data/PDB/CUUCGG/2M58_12_C_A_0_00000.pdb 
# Loaded reference data/PDB/CUUCGG/2KOC_5_C_A_0_00001.pdb 
# Loaded target data/PDB/CUUCGG/2N3Q_37_C_A_0_00014.pdb 
# Loading data/PDB/CUUCGG/2N3Q_37_C_A_0_00014.pdb 
# Loaded reference data/PDB/CUUCGG/2KOC_5_C_A_0_00001.pdb 
# Loaded target data/PDB/CUUCGG/1FMN_16_C_A_0_00002.pdb 
# Loading data/PDB/CUUCGG/1FMN_16_C_A_0_00002.pdb 
# Loaded reference data/PDB/CUUCGG/2KOC_5_C_A_0_00001.pdb 
# Loaded target data/PDB/CUUCGG/1BYJ_11_C_A_0_00024.pdb 
# Loading data/PDB/CUUCGG/1BYJ_11_C_A_0_00024.pdb 
# Loaded reference data/PDB/CUUCGG/2KOC_5_C_A_0_00001.pdb 
# Loaded target data/PDB/CUUCGG/2LHP_16_C_A_0_00005.pdb 
# Loading data/PDB/CUUCGG/2LHP_16_C_A_0_00005.pdb 
# Loaded reference data/PDB/CUUCGG/2KOC_5_C_A_0_00001.pdb 
# Loaded target data/PDB/CUUCGG/1B36_16_C_A_0_00005.pdb 
# Loading data/PDB/CUUCGG/1B36_16_C_A_0_00005.pdb 
# Loaded reference data/PDB/CUUCGG/2KOC_5_C_A_0_00001.pdb 
# Loaded target data/PDB/CUUCGG/2E5L_419_C_A_0_00000.pdb 
# Loading data/PDB/CUUCGG/2E5L_419_C_A_0_00000.pdb 
# Loaded reference data/PDB/CUUCGG/2KOC_5_C_A_0_00001.pdb 
# Loaded target data/PDB/CUUCGG/2HUA_17_C_A_0_00000.pdb 
# Loading data/PDB/CUUCGG/2HUA_17_C_A_0_00000.pdb 
# Loaded reference data/PDB/CUUCGG/2KOC_5_C_A_0_00001.pdb 
# Loaded target data/PDB/CUUCGG/6FF4_8_C_6_0_00000.pdb 
# Loading data/PDB/CUUCGG/6FF4_8_C_6_0_00000.pdb 
# Loaded reference data/PDB/CUUCGG/2KOC_5_C_A_0_00001.pdb 
# Loaded target data/PDB/CUUCGG/4wsm-pdb-bundle2_1691_C_A_0_00000.pdb 
# Loading data/PDB/CUUCGG/4wsm-pdb-bundle2_1691_C_A_0_00000.pdb 
# Loaded reference data/PDB/CUUCGG/2KOC_5_C_A_0_00001.pdb 
# Loaded target data/PDB/CUUCGG/6hrm-pdb-bundle1_419_C_1_0_00001.pdb 
# Loading data/PDB/CUUCGG/6hrm-pdb-bundle1_419_C_1_0_00001.pdb 
# Loaded reference data/PDB/CUUCGG/2KOC_5_C_A_0_00001.pdb 
# Loaded target data/PDB/CUUCGG/1BYJ_11_C_A_0_00018.pdb 
# Loading data/PDB/CUUCGG/1BYJ_11_C_A_0_00018.pdb 
# Loaded reference data/PDB/CUUCGG/2KOC_5_C_A_0_00001.pdb 
# Loaded target data/PDB/CUUCGG/2LQZ_13_C_A_0_00001.pdb 
# Loading data/PDB/CUUCGG/2LQZ_13_C_A_0_00001.pdb 
# Loaded reference data/PDB/CUUCGG/2KOC_5_C_A_0_00001.pdb 
# Loaded target data/PDB/CUUCGG/1A3M_11_C_A_0_00003.pdb 
# Loading data/PDB/CUUCGG/1A3M_11_C_A_0_00003.pdb 
# Loaded reference data/PDB/CUUCGG/2KOC_5_C_A_0_00001.pdb 
# Loaded target data/PDB/CUUCGG/4yzv-pdb-bundle2_1691_C_A_0_00000.pdb 
# Loading data/PDB/CUUCGG/4yzv-pdb-bundle2_1691_C_A_0_00000.pdb 
# Loaded reference data/PDB/CUUCGG/2KOC_5_C_A_0_00001.pdb 
# Loaded target data/PDB/CUUCGG/1A3M_11_C_A_0_00017.pdb 
# Loading data/PDB/CUUCGG/1A3M_11_C_A_0_00017.pdb 
# Loaded reference data/PDB/CUUCGG/2KOC_5_C_A_0_00001.pdb 
# Loaded target data/PDB/CUUCGG/2GIO_13_C_A_0_00009.pdb 
# Loading data/PDB/CUUCGG/2GIO_13_C_A_0_00009.pdb 
# Loaded reference data/PDB/CUUCGG/2KOC_5_C_A_0_00001.pdb 
# Loaded target data/PDB/CUUCGG/5e81-pdb-bundle3_419_C_A_0_00000.pdb 
# Loading data/PDB/CUUCGG/5e81-pdb-bundle3_419_C_A_0_00000.pdb 
# Loaded reference data/PDB/CUUCGG/2KOC_5_C_A_0_00001.pdb 
# Loaded target data/PDB/CUUCGG/5MMI_1727_C_A_0_00001.pdb 
# Loading data/PDB/CUUCGG/5MMI_1727_C_A_0_00001.pdb 
# Loaded reference data/PDB/CUUCGG/2KOC_5_C_A_0_00001.pdb 
# Loaded target data/PDB/CUUCGG/4u52-pdb-bundle3_230_C_d_0_00000.pdb 
# Loading data/PDB/CUUCGG/4u52-pdb-bundle3_230_C_d_0_00000.pdb 
# Loaded reference data/PDB/CUUCGG/2KOC_5_C_A_0_00001.pdb 
# Loaded target data/PDB/CUUCGG/1Q81_1769_C_A_0_00000.pdb 
# Loading data/PDB/CUUCGG/1Q81_1769_C_A_0_00000.pdb 
# Loaded reference data/PDB/CUUCGG/2KOC_5_C_A_0_00001.pdb 
# Loaded target data/PDB/CUUCGG/4tue-pdb-bundle1_419_C_A_0_00000.pdb 
# Loading data/PDB/CUUCGG/4tue-pdb-bundle1_419_C_A_0_00000.pdb 
# Loaded reference data/PDB/CUUCGG/2KOC_5_C_A_0_00001.pdb 
# Loaded target data/PDB/CUUCGG/3PIO_1708_C_X_0_00000.pdb 
# Loading data/PDB/CUUCGG/3PIO_1708_C_X_0_00000.pdb 
# Loaded reference data/PDB/CUUCGG/2KOC_5_C_A_0_00001.pdb 
# Loaded target data/PDB/CUUCGG/4wzo-pdb-bundle2_1691_C_A_0_00000.pdb 
# Loading data/PDB/CUUCGG/4wzo-pdb-bundle2_1691_C_A_0_00000.pdb 
# Loaded reference data/PDB/CUUCGG/2KOC_5_C_A_0_00001.pdb 
# Loaded target data/PDB/CUUCGG/4u4u-pdb-bundle4_1923_C_I_0_00002.pdb 
# Loading data/PDB/CUUCGG/4u4u-pdb-bundle4_1923_C_I_0_00002.pdb 
# Loaded reference data/PDB/CUUCGG/2KOC_5_C_A_0_00001.pdb 
# Loaded target data/PDB/CUUCGG/1ZHO_15_C_F_0_00002.pdb 
# Loading data/PDB/CUUCGG/1ZHO_15_C_F_0_00002.pdb 
# Loaded reference data/PDB/CUUCGG/2KOC_5_C_A_0_00001.pdb 
# Loaded target data/PDB/CUUCGG/5zeb-pdb-bundle2_1909_C_A_0_00000.pdb 
# Loading data/PDB/CUUCGG/5zeb-pdb-bundle2_1909_C_A_0_00000.pdb 
# Loaded reference data/PDB/CUUCGG/2KOC_5_C_A_0_00001.pdb 
# Loaded target data/PDB/CUUCGG/4v85-pdb-bundle1_1449_C_A_0_00003.pdb 
# Loading data/PDB/CUUCGG/4v85-pdb-bundle1_1449_C_A_0_00003.pdb 
# Loaded reference data/PDB/CUUCGG/2KOC_5_C_A_0_00001.pdb 
# Loaded target data/PDB/CUUCGG/3jah-pdb-bundle2_2872_C_5_0_00000.pdb 
# Loading data/PDB/CUUCGG/3jah-pdb-bundle2_2872_C_5_0_00000.pdb 
# Loaded reference data/PDB/CUUCGG/2KOC_5_C_A_0_00001.pdb 
# Loaded target data/PDB/CUUCGG/4u1u-pdb-bundle3_1449_C_A_0_00003.pdb 
# Loading data/PDB/CUUCGG/4u1u-pdb-bundle3_1449_C_A_0_00003.pdb 
# Loaded reference data/PDB/CUUCGG/2KOC_5_C_A_0_00001.pdb 
# Loaded target data/PDB/CUUCGG/2N3R_37_C_A_0_00004.pdb 
# Loading data/PDB/CUUCGG/2N3R_37_C_A_0_00004.pdb 
# Loaded reference data/PDB/CUUCGG/2KOC_5_C_A_0_00001.pdb 
# Loaded target data/PDB/CUUCGG/3CCJ_1769_C_0_0_00000.pdb 
# Loading data/PDB/CUUCGG/3CCJ_1769_C_0_0_00000.pdb 
# Loaded reference data/PDB/CUUCGG/2KOC_5_C_A_0_00001.pdb 
# Loaded target data/PDB/CUUCGG/2L3E_18_C_A_0_00003.pdb 
# Loading data/PDB/CUUCGG/2L3E_18_C_A_0_00003.pdb 
# Loaded reference data/PDB/CUUCGG/2KOC_5_C_A_0_00001.pdb 
# Loaded target data/PDB/CUUCGG/2L3E_18_C_A_0_00017.pdb 
# Loading data/PDB/CUUCGG/2L3E_18_C_A_0_00017.pdb 
# Loaded reference data/PDB/CUUCGG/2KOC_5_C_A_0_00001.pdb 
# Loaded target data/PDB/CUUCGG/2N3R_37_C_A_0_00010.pdb 
# Loading data/PDB/CUUCGG/2N3R_37_C_A_0_00010.pdb 
# Loaded reference data/PDB/CUUCGG/2KOC_5_C_A_0_00001.pdb 
# Loaded target data/PDB/CUUCGG/4WFB_1735_C_X_0_00000.pdb 
# Loading data/PDB/CUUCGG/4WFB_1735_C_X_0_00000.pdb 
# Loaded reference data/PDB/CUUCGG/2KOC_5_C_A_0_00001.pdb 
# Loaded target data/PDB/CUUCGG/5hd1-pdb-bundle3_1737_C_A_0_00000.pdb 
# Loading data/PDB/CUUCGG/5hd1-pdb-bundle3_1737_C_A_0_00000.pdb 
# Loaded reference data/PDB/CUUCGG/2KOC_5_C_A_0_00001.pdb 
# Loaded target data/PDB/CUUCGG/5DM6_1691_C_X_0_00000.pdb 
# Loading data/PDB/CUUCGG/5DM6_1691_C_X_0_00000.pdb 
# Loaded reference data/PDB/CUUCGG/2KOC_5_C_A_0_00001.pdb 
# Loaded target data/PDB/CUUCGG/5afi-pdb-bundle1_1449_C_a_0_00003.pdb 
# Loading data/PDB/CUUCGG/5afi-pdb-bundle1_1449_C_a_0_00003.pdb 
# Loaded reference data/PDB/CUUCGG/2KOC_5_C_A_0_00001.pdb 
# Loaded target data/PDB/CUUCGG/6BY5_5_C_A_0_00009.pdb 
# Loading data/PDB/CUUCGG/6BY5_5_C_A_0_00009.pdb 
# Loaded reference data/PDB/CUUCGG/2KOC_5_C_A_0_00001.pdb 
# Loaded target data/PDB/CUUCGG/2QA4_1769_C_0_0_00000.pdb 
# Loading data/PDB/CUUCGG/2QA4_1769_C_0_0_00000.pdb 
# Loaded reference data/PDB/CUUCGG/2KOC_5_C_A_0_00001.pdb 
# Loaded target data/PDB/CUUCGG/6c5l-pdb-bundle2_1691_C_A_0_00000.pdb 
# Loading data/PDB/CUUCGG/6c5l-pdb-bundle2_1691_C_A_0_00000.pdb 
# Loaded reference data/PDB/CUUCGG/2KOC_5_C_A_0_00001.pdb 
# Loaded target data/PDB/CUUCGG/4v57-pdb-bundle4_1691_C_A_0_00000.pdb 
# Loading data/PDB/CUUCGG/4v57-pdb-bundle4_1691_C_A_0_00000.pdb 
# Loaded reference data/PDB/CUUCGG/2KOC_5_C_A_0_00001.pdb 
# Loaded target data/PDB/CUUCGG/4v7u-pdb-bundle3_419_C_A_0_00001.pdb 
# Loading data/PDB/CUUCGG/4v7u-pdb-bundle3_419_C_A_0_00001.pdb 
# Loaded reference data/PDB/CUUCGG/2KOC_5_C_A_0_00001.pdb 
# Loaded target data/PDB/CUUCGG/2LU0_28_C_A_0_00018.pdb 
# Loading data/PDB/CUUCGG/2LU0_28_C_A_0_00018.pdb 
# Loaded reference data/PDB/CUUCGG/2KOC_5_C_A_0_00001.pdb 
# Loaded target data/PDB/CUUCGG/4v50-pdb-bundle3_207_C_A_0_00000.pdb 
# Loading data/PDB/CUUCGG/4v50-pdb-bundle3_207_C_A_0_00000.pdb 
# Loaded reference data/PDB/CUUCGG/2KOC_5_C_A_0_00001.pdb 
# Loaded target data/PDB/CUUCGG/4u4q-pdb-bundle4_1923_C_I_0_00002.pdb 
# Loading data/PDB/CUUCGG/4u4q-pdb-bundle4_1923_C_I_0_00002.pdb 
# Loaded reference data/PDB/CUUCGG/2KOC_5_C_A_0_00001.pdb 
# Loaded target data/PDB/CUUCGG/4v64-pdb-bundle4_1691_C_A_0_00000.pdb 
# Loading data/PDB/CUUCGG/4v64-pdb-bundle4_1691_C_A_0_00000.pdb 
# Loaded reference data/PDB/CUUCGG/2KOC_5_C_A_0_00001.pdb 
# Loaded target data/PDB/CUUCGG/1VQM_1769_C_0_0_00000.pdb 
# Loading data/PDB/CUUCGG/1VQM_1769_C_0_0_00000.pdb 
# Loaded reference data/PDB/CUUCGG/2KOC_5_C_A_0_00001.pdb 
# Loaded target data/PDB/CUUCGG/2L3E_18_C_A_0_00001.pdb 
# Loading data/PDB/CUUCGG/2L3E_18_C_A_0_00001.pdb 
# Loaded reference data/PDB/CUUCGG/2KOC_5_C_A_0_00001.pdb 
# Loaded target data/PDB/CUUCGG/2N3R_37_C_A_0_00006.pdb 
# Loading data/PDB/CUUCGG/2N3R_37_C_A_0_00006.pdb 
# Loaded reference data/PDB/CUUCGG/2KOC_5_C_A_0_00001.pdb 
# Loaded target data/PDB/CUUCGG/5ndj-pdb-bundle3_1741_C_A_0_00000.pdb 
# Loading data/PDB/CUUCGG/5ndj-pdb-bundle3_1741_C_A_0_00000.pdb 
# Loaded reference data/PDB/CUUCGG/2KOC_5_C_A_0_00001.pdb 
# Loaded target data/PDB/CUUCGG/3G6E_1769_C_0_0_00000.pdb 
# Loading data/PDB/CUUCGG/3G6E_1769_C_0_0_00000.pdb 
# Loaded reference data/PDB/CUUCGG/2KOC_5_C_A_0_00001.pdb 
# Loaded target data/PDB/CUUCGG/2N3R_37_C_A_0_00012.pdb 
# Loading data/PDB/CUUCGG/2N3R_37_C_A_0_00012.pdb 
# Loaded reference data/PDB/CUUCGG/2KOC_5_C_A_0_00001.pdb 
# Loaded target data/PDB/CUUCGG/2L3E_18_C_A_0_00015.pdb 
# Loading data/PDB/CUUCGG/2L3E_18_C_A_0_00015.pdb 
# Loaded reference data/PDB/CUUCGG/2KOC_5_C_A_0_00001.pdb 
# Loaded target data/PDB/CUUCGG/5fdv-pdb-bundle3_1691_C_A_0_00000.pdb 
# Loading data/PDB/CUUCGG/5fdv-pdb-bundle3_1691_C_A_0_00000.pdb 
# Loaded reference data/PDB/CUUCGG/2KOC_5_C_A_0_00001.pdb 
# Loaded target data/PDB/CUUCGG/6mtc-pdb-bundle1_2872_C_5_0_00000.pdb 
# Loading data/PDB/CUUCGG/6mtc-pdb-bundle1_2872_C_5_0_00000.pdb 
# Loaded reference data/PDB/CUUCGG/2KOC_5_C_A_0_00001.pdb 
# Loaded target data/PDB/CUUCGG/4ypb-pdb-bundle1_419_C_A_0_00000.pdb 
# Loading data/PDB/CUUCGG/4ypb-pdb-bundle1_419_C_A_0_00000.pdb 
# Loaded reference data/PDB/CUUCGG/2KOC_5_C_A_0_00001.pdb 
# Loaded target data/PDB/CUUCGG/4v7m-pdb-bundle4_1691_C_A_0_00000.pdb 
# Loading data/PDB/CUUCGG/4v7m-pdb-bundle4_1691_C_A_0_00000.pdb 
# Loaded reference data/PDB/CUUCGG/2KOC_5_C_A_0_00001.pdb 
# Loaded target data/PDB/CUUCGG/5jc9-pdb-bundle2_1691_C_T_0_00000.pdb 
# Loading data/PDB/CUUCGG/5jc9-pdb-bundle2_1691_C_T_0_00000.pdb 
# Loaded reference data/PDB/CUUCGG/2KOC_5_C_A_0_00001.pdb 
# Loaded target data/PDB/CUUCGG/4v4h-pdb-bundle3_419_C_A_0_00001.pdb 
# Loading data/PDB/CUUCGG/4v4h-pdb-bundle3_419_C_A_0_00001.pdb 
# Loaded reference data/PDB/CUUCGG/2KOC_5_C_A_0_00001.pdb 
# Loaded target data/PDB/CUUCGG/2GIP_12_C_A_0_00018.pdb 
# Loading data/PDB/CUUCGG/2GIP_12_C_A_0_00018.pdb 
# Loaded reference data/PDB/CUUCGG/2KOC_5_C_A_0_00001.pdb 
# Loaded target data/PDB/CUUCGG/5mdy-pdb-bundle1_419_C_2_0_00002.pdb 
# Loading data/PDB/CUUCGG/5mdy-pdb-bundle1_419_C_2_0_00002.pdb 
# Loaded reference data/PDB/CUUCGG/2KOC_5_C_A_0_00001.pdb 
# Loaded target data/PDB/CUUCGG/3CCE_1769_C_0_0_00000.pdb 
# Loading data/PDB/CUUCGG/3CCE_1769_C_0_0_00000.pdb 
# Loaded reference data/PDB/CUUCGG/2KOC_5_C_A_0_00001.pdb 
# Loaded target data/PDB/CUUCGG/1MFY_14_C_A_0_00008.pdb 
# Loading data/PDB/CUUCGG/1MFY_14_C_A_0_00008.pdb 
# Loaded reference data/PDB/CUUCGG/2KOC_5_C_A_0_00001.pdb 
# Loaded target data/PDB/CUUCGG/5KH8_26_C_A_0_00018.pdb 
# Loading data/PDB/CUUCGG/5KH8_26_C_A_0_00018.pdb 
# Loaded reference data/PDB/CUUCGG/2KOC_5_C_A_0_00001.pdb 
# Loaded target data/PDB/CUUCGG/1A3M_11_C_A_0_00001.pdb 
# Loading data/PDB/CUUCGG/1A3M_11_C_A_0_00001.pdb 
# Loaded reference data/PDB/CUUCGG/2KOC_5_C_A_0_00001.pdb 
# Loaded target data/PDB/CUUCGG/1A3M_11_C_A_0_00015.pdb 
# Loading data/PDB/CUUCGG/1A3M_11_C_A_0_00015.pdb 
# Loaded reference data/PDB/CUUCGG/2KOC_5_C_A_0_00001.pdb 
# Loaded target data/PDB/CUUCGG/5dfe-pdb-bundle3_1691_C_I_0_00000.pdb 
# Loading data/PDB/CUUCGG/5dfe-pdb-bundle3_1691_C_I_0_00000.pdb 
# Loaded reference data/PDB/CUUCGG/2KOC_5_C_A_0_00001.pdb 
# Loaded target data/PDB/CUUCGG/1vy6-pdb-bundle2_1691_C_A_0_00000.pdb 
# Loading data/PDB/CUUCGG/1vy6-pdb-bundle2_1691_C_A_0_00000.pdb 
# Loaded reference data/PDB/CUUCGG/2KOC_5_C_A_0_00001.pdb 
# Loaded target data/PDB/CUUCGG/2HUA_17_C_A_0_00002.pdb 
# Loading data/PDB/CUUCGG/2HUA_17_C_A_0_00002.pdb 
# Loaded reference data/PDB/CUUCGG/2KOC_5_C_A_0_00001.pdb 
# Loaded target data/PDB/CUUCGG/3jbv-pdb-bundle1_419_C_A_0_00001.pdb 
# Loading data/PDB/CUUCGG/3jbv-pdb-bundle1_419_C_A_0_00001.pdb 
# Loaded reference data/PDB/CUUCGG/2KOC_5_C_A_0_00001.pdb 
# Loaded target data/PDB/CUUCGG/1B36_16_C_A_0_00007.pdb 
# Loading data/PDB/CUUCGG/1B36_16_C_A_0_00007.pdb 
# Loaded reference data/PDB/CUUCGG/2KOC_5_C_A_0_00001.pdb 
# Loaded target data/PDB/CUUCGG/4v9f-pdb-bundle1_1769_C_0_0_00000.pdb 
# Loading data/PDB/CUUCGG/4v9f-pdb-bundle1_1769_C_0_0_00000.pdb 
# Loaded reference data/PDB/CUUCGG/2KOC_5_C_A_0_00001.pdb 
# Loaded target data/PDB/CUUCGG/2LHP_16_C_A_0_00007.pdb 
# Loading data/PDB/CUUCGG/2LHP_16_C_A_0_00007.pdb 
# Loaded reference data/PDB/CUUCGG/2KOC_5_C_A_0_00001.pdb 
# Loaded target data/PDB/CUUCGG/5dc3-pdb-bundle4_1923_C_C_0_00002.pdb 
# Loading data/PDB/CUUCGG/5dc3-pdb-bundle4_1923_C_C_0_00002.pdb 
# Loaded reference data/PDB/CUUCGG/2KOC_5_C_A_0_00001.pdb 
# Loaded target data/PDB/CUUCGG/4v9n-pdb-bundle4_1691_C_A_0_00000.pdb 
# Loading data/PDB/CUUCGG/4v9n-pdb-bundle4_1691_C_A_0_00000.pdb 
# Loaded reference data/PDB/CUUCGG/2KOC_5_C_A_0_00001.pdb 
# Loaded target data/PDB/CUUCGG/2LQZ_13_C_A_0_00003.pdb 
# Loading data/PDB/CUUCGG/2LQZ_13_C_A_0_00003.pdb 
# Loaded reference data/PDB/CUUCGG/2KOC_5_C_A_0_00001.pdb 
# Loaded target data/PDB/CUUCGG/4u4z-pdb-bundle2_1923_C_A_0_00002.pdb 
# Loading data/PDB/CUUCGG/4u4z-pdb-bundle2_1923_C_A_0_00002.pdb 
# Loaded reference data/PDB/CUUCGG/2KOC_5_C_A_0_00001.pdb 
# Loaded target data/PDB/CUUCGG/2N3Q_37_C_A_0_00002.pdb 
# Loading data/PDB/CUUCGG/2N3Q_37_C_A_0_00002.pdb 
# Loaded reference data/PDB/CUUCGG/2KOC_5_C_A_0_00001.pdb 
# Loaded target data/PDB/CUUCGG/1BYJ_11_C_A_0_00032.pdb 
# Loading data/PDB/CUUCGG/1BYJ_11_C_A_0_00032.pdb 
# Loaded reference data/PDB/CUUCGG/2KOC_5_C_A_0_00001.pdb 
# Loaded target data/PDB/CUUCGG/1BYJ_11_C_A_0_00026.pdb 
# Loading data/PDB/CUUCGG/1BYJ_11_C_A_0_00026.pdb 
# Loaded reference data/PDB/CUUCGG/2KOC_5_C_A_0_00001.pdb 
# Loaded target data/PDB/CUUCGG/1FMN_16_C_A_0_00000.pdb 
# Loading data/PDB/CUUCGG/1FMN_16_C_A_0_00000.pdb 
# Loaded reference data/PDB/CUUCGG/2KOC_5_C_A_0_00001.pdb 
# Loaded target data/PDB/CUUCGG/2N3Q_37_C_A_0_00016.pdb 
# Loading data/PDB/CUUCGG/2N3Q_37_C_A_0_00016.pdb 
# Loaded reference data/PDB/CUUCGG/2KOC_5_C_A_0_00001.pdb 
# Loaded target data/PDB/CUUCGG/2M58_12_C_A_0_00002.pdb 
# Loading data/PDB/CUUCGG/2M58_12_C_A_0_00002.pdb 
# Loaded reference data/PDB/CUUCGG/2KOC_5_C_A_0_00001.pdb 
# Loaded target data/PDB/CUUCGG/4wu1-pdb-bundle2_1691_C_A_0_00000.pdb 
# Loading data/PDB/CUUCGG/4wu1-pdb-bundle2_1691_C_A_0_00000.pdb 
# Loaded reference data/PDB/CUUCGG/2KOC_5_C_A_0_00001.pdb 
# Loaded target data/PDB/CUUCGG/4v50-pdb-bundle3_1449_C_A_0_00003.pdb 
# Loading data/PDB/CUUCGG/4v50-pdb-bundle3_1449_C_A_0_00003.pdb 
# Loaded reference data/PDB/CUUCGG/2KOC_5_C_A_0_00001.pdb 
# Loaded target data/PDB/CUUCGG/1D6K_283_C_B_0_00002.pdb 
# Loading data/PDB/CUUCGG/1D6K_283_C_B_0_00002.pdb 
# Loaded reference data/PDB/CUUCGG/2KOC_5_C_A_0_00001.pdb 
# Loaded target data/PDB/CUUCGG/6HMA_1735_C_A_0_00000.pdb 
# Loading data/PDB/CUUCGG/6HMA_1735_C_A_0_00000.pdb 
# Loaded reference data/PDB/CUUCGG/2KOC_5_C_A_0_00001.pdb 
# Loaded target data/PDB/CUUCGG/1D6K_283_C_B_0_00016.pdb 
# Loading data/PDB/CUUCGG/1D6K_283_C_B_0_00016.pdb 
# Loaded reference data/PDB/CUUCGG/2KOC_5_C_A_0_00001.pdb 
# Loaded target data/PDB/CUUCGG/4v5k-pdb-bundle2_1691_C_A_0_00000.pdb 
# Loading data/PDB/CUUCGG/4v5k-pdb-bundle2_1691_C_A_0_00000.pdb 
# Loaded reference data/PDB/CUUCGG/2KOC_5_C_A_0_00001.pdb 
# Loaded target data/PDB/CUUCGG/2NCI_14_C_A_0_00005.pdb 
# Loading data/PDB/CUUCGG/2NCI_14_C_A_0_00005.pdb 
# Loaded reference data/PDB/CUUCGG/2KOC_5_C_A_0_00001.pdb 
# Loaded target data/PDB/CUUCGG/4B3S_419_C_A_0_00000.pdb 
# Loading data/PDB/CUUCGG/4B3S_419_C_A_0_00000.pdb 
# Loaded reference data/PDB/CUUCGG/2KOC_5_C_A_0_00001.pdb 
# Loaded target data/PDB/CUUCGG/2LWK_15_C_A_0_00000.pdb 
# Loading data/PDB/CUUCGG/2LWK_15_C_A_0_00000.pdb 
# Loaded reference data/PDB/CUUCGG/2KOC_5_C_A_0_00001.pdb 
# Loaded target data/PDB/CUUCGG/1TLR_10_C_A_0_00019.pdb 
# Loading data/PDB/CUUCGG/1TLR_10_C_A_0_00019.pdb 
# Loaded reference data/PDB/CUUCGG/2KOC_5_C_A_0_00001.pdb 
# Loaded target data/PDB/CUUCGG/5wfk-pdb-bundle1_1691_C_A_0_00000.pdb 
# Loading data/PDB/CUUCGG/5wfk-pdb-bundle1_1691_C_A_0_00000.pdb 
# Loaded reference data/PDB/CUUCGG/2KOC_5_C_A_0_00001.pdb 
# Loaded target data/PDB/CUUCGG/2LWK_15_C_A_0_00014.pdb 
# Loading data/PDB/CUUCGG/2LWK_15_C_A_0_00014.pdb 
# Loaded reference data/PDB/CUUCGG/2KOC_5_C_A_0_00001.pdb 
# Loaded target data/PDB/CUUCGG/5lzz-pdb-bundle2_2872_C_5_0_00000.pdb 
# Loading data/PDB/CUUCGG/5lzz-pdb-bundle2_2872_C_5_0_00000.pdb 
# Loaded reference data/PDB/CUUCGG/2KOC_5_C_A_0_00001.pdb 
# Loaded target data/PDB/CUUCGG/6cfk-pdb-bundle3_1691_C_A_0_00000.pdb 
# Loading data/PDB/CUUCGG/6cfk-pdb-bundle3_1691_C_A_0_00000.pdb 
# Loaded reference data/PDB/CUUCGG/2KOC_5_C_A_0_00001.pdb 
# Loaded target data/PDB/CUUCGG/1VQ8_1769_C_0_0_00000.pdb 
# Loading data/PDB/CUUCGG/1VQ8_1769_C_0_0_00000.pdb 
# Loaded reference data/PDB/CUUCGG/2KOC_5_C_A_0_00001.pdb 
# Loaded target data/PDB/CUUCGG/5i4l-pdb-bundle4_1923_C_E_0_00002.pdb 
# Loading data/PDB/CUUCGG/5i4l-pdb-bundle4_1923_C_E_0_00002.pdb 
# Loaded reference data/PDB/CUUCGG/2KOC_5_C_A_0_00001.pdb 
# Loaded target data/PDB/CUUCGG/1FYO_11_C_A_0_00019.pdb 
# Loading data/PDB/CUUCGG/1FYO_11_C_A_0_00019.pdb 
# Loaded reference data/PDB/CUUCGG/2KOC_5_C_A_0_00001.pdb 
# Loaded target data/PDB/CUUCGG/4v5e-pdb-bundle4_1691_C_A_0_00000.pdb 
# Loading data/PDB/CUUCGG/4v5e-pdb-bundle4_1691_C_A_0_00000.pdb 
# Loaded reference data/PDB/CUUCGG/2KOC_5_C_A_0_00001.pdb 
# Loaded target data/PDB/CUUCGG/5KQE_19_C_A_0_00019.pdb 
# Loading data/PDB/CUUCGG/5KQE_19_C_A_0_00019.pdb 
# Loaded reference data/PDB/CUUCGG/2KOC_5_C_A_0_00001.pdb 
# Loaded target data/PDB/CUUCGG/2N6S_16_C_A_0_00003.pdb 
# Loading data/PDB/CUUCGG/2N6S_16_C_A_0_00003.pdb 
# Loaded reference data/PDB/CUUCGG/2KOC_5_C_A_0_00001.pdb 
# Loaded target data/PDB/CUUCGG/2N6S_16_C_A_0_00002.pdb 
# Loading data/PDB/CUUCGG/2N6S_16_C_A_0_00002.pdb 
# Loaded reference data/PDB/CUUCGG/2KOC_5_C_A_0_00001.pdb 
# Loaded target data/PDB/CUUCGG/5e7k-pdb-bundle1_419_C_A_0_00000.pdb 
# Loading data/PDB/CUUCGG/5e7k-pdb-bundle1_419_C_A_0_00000.pdb 
# Loaded reference data/PDB/CUUCGG/2KOC_5_C_A_0_00001.pdb 
# Loaded target data/PDB/CUUCGG/1FYO_11_C_A_0_00018.pdb 
# Loading data/PDB/CUUCGG/1FYO_11_C_A_0_00018.pdb 
# Loaded reference data/PDB/CUUCGG/2KOC_5_C_A_0_00001.pdb 
# Loaded target data/PDB/CUUCGG/4lt8-pdb-bundle4_1691_C_A_0_00000.pdb 
# Loading data/PDB/CUUCGG/4lt8-pdb-bundle4_1691_C_A_0_00000.pdb 
# Loaded reference data/PDB/CUUCGG/2KOC_5_C_A_0_00001.pdb 
# Loaded target data/PDB/CUUCGG/5KQE_19_C_A_0_00018.pdb 
# Loading data/PDB/CUUCGG/5KQE_19_C_A_0_00018.pdb 
# Loaded reference data/PDB/CUUCGG/2KOC_5_C_A_0_00001.pdb 
# Loaded target data/PDB/CUUCGG/4wt8-pdb-bundle2_1737_C_o_0_00000.pdb 
# Loading data/PDB/CUUCGG/4wt8-pdb-bundle2_1737_C_o_0_00000.pdb 
# Loaded reference data/PDB/CUUCGG/2KOC_5_C_A_0_00001.pdb 
# Loaded target data/PDB/CUUCGG/5lyb-pdb-bundle2_1923_C_A_0_00002.pdb 
# Loading data/PDB/CUUCGG/5lyb-pdb-bundle2_1923_C_A_0_00002.pdb 
# Loaded reference data/PDB/CUUCGG/2KOC_5_C_A_0_00001.pdb 
# Loaded target data/PDB/CUUCGG/1FYO_11_C_A_0_00024.pdb 
# Loading data/PDB/CUUCGG/1FYO_11_C_A_0_00024.pdb 
# Loaded reference data/PDB/CUUCGG/2KOC_5_C_A_0_00001.pdb 
# Loaded target data/PDB/CUUCGG/5u9g-pdb-bundle1_1449_C_A_0_00003.pdb 
# Loading data/PDB/CUUCGG/5u9g-pdb-bundle1_1449_C_A_0_00003.pdb 
# Loaded reference data/PDB/CUUCGG/2KOC_5_C_A_0_00001.pdb 
# Loaded target data/PDB/CUUCGG/6gsj-pdb-bundle4_1691_C_A_0_00000.pdb 
# Loading data/PDB/CUUCGG/6gsj-pdb-bundle4_1691_C_A_0_00000.pdb 
# Loaded reference data/PDB/CUUCGG/2KOC_5_C_A_0_00001.pdb 
# Loaded target data/PDB/CUUCGG/4v5r-pdb-bundle4_1691_C_A_0_00000.pdb 
# Loading data/PDB/CUUCGG/4v5r-pdb-bundle4_1691_C_A_0_00000.pdb 
# Loaded reference data/PDB/CUUCGG/2KOC_5_C_A_0_00001.pdb 
# Loaded target data/PDB/CUUCGG/2LWK_15_C_A_0_00015.pdb 
# Loading data/PDB/CUUCGG/2LWK_15_C_A_0_00015.pdb 
# Loaded reference data/PDB/CUUCGG/2KOC_5_C_A_0_00001.pdb 
# Loaded target data/PDB/CUUCGG/1TLR_10_C_A_0_00018.pdb 
# Loading data/PDB/CUUCGG/1TLR_10_C_A_0_00018.pdb 
# Loaded reference data/PDB/CUUCGG/2KOC_5_C_A_0_00001.pdb 
# Loaded target data/PDB/CUUCGG/5j88-pdb-bundle1_1449_C_b_0_00007.pdb 
# Loading data/PDB/CUUCGG/5j88-pdb-bundle1_1449_C_b_0_00007.pdb 
# Loaded reference data/PDB/CUUCGG/2KOC_5_C_A_0_00001.pdb 
# Loaded target data/PDB/CUUCGG/4v6f-pdb-bundle2_419_C_Z_0_00001.pdb 
# Loading data/PDB/CUUCGG/4v6f-pdb-bundle2_419_C_Z_0_00001.pdb 
# Loaded reference data/PDB/CUUCGG/2KOC_5_C_A_0_00001.pdb 
# Loaded target data/PDB/CUUCGG/2LWK_15_C_A_0_00001.pdb 
# Loading data/PDB/CUUCGG/2LWK_15_C_A_0_00001.pdb 
# Loaded reference data/PDB/CUUCGG/2KOC_5_C_A_0_00001.pdb 
# Loaded target data/PDB/CUUCGG/1D6K_283_C_B_0_00017.pdb 
# Loading data/PDB/CUUCGG/1D6K_283_C_B_0_00017.pdb 
# Loaded reference data/PDB/CUUCGG/2KOC_5_C_A_0_00001.pdb 
# Loaded target data/PDB/CUUCGG/2NCI_14_C_A_0_00004.pdb 
# Loading data/PDB/CUUCGG/2NCI_14_C_A_0_00004.pdb 
# Loaded reference data/PDB/CUUCGG/2KOC_5_C_A_0_00001.pdb 
# Loaded target data/PDB/CUUCGG/5e81-pdb-bundle1_419_C_A_0_00000.pdb 
# Loading data/PDB/CUUCGG/5e81-pdb-bundle1_419_C_A_0_00000.pdb 
# Loaded reference data/PDB/CUUCGG/2KOC_5_C_A_0_00001.pdb 
# Loaded target data/PDB/CUUCGG/1D6K_283_C_B_0_00003.pdb 
# Loading data/PDB/CUUCGG/1D6K_283_C_B_0_00003.pdb 
# Loaded reference data/PDB/CUUCGG/2KOC_5_C_A_0_00001.pdb 
# Loaded target data/PDB/CUUCGG/5j3c-pdb-bundle2_419_C_A_0_00000.pdb 
# Loading data/PDB/CUUCGG/5j3c-pdb-bundle2_419_C_A_0_00000.pdb 
# Loaded reference data/PDB/CUUCGG/2KOC_5_C_A_0_00001.pdb 
# Loaded target data/PDB/CUUCGG/5hcp-pdb-bundle1_1737_C_A_0_00000.pdb 
# Loading data/PDB/CUUCGG/5hcp-pdb-bundle1_1737_C_A_0_00000.pdb 
# Loaded reference data/PDB/CUUCGG/2KOC_5_C_A_0_00001.pdb 
# Loaded target data/PDB/CUUCGG/6CAO_419_C_A_0_00000.pdb 
# Loading data/PDB/CUUCGG/6CAO_419_C_A_0_00000.pdb 
# Loaded reference data/PDB/CUUCGG/2KOC_5_C_A_0_00001.pdb 
# Loaded target data/PDB/CUUCGG/5D5L_60_C_A_0_00000.pdb 
# Loading data/PDB/CUUCGG/5D5L_60_C_A_0_00000.pdb 
# Loaded reference data/PDB/CUUCGG/2KOC_5_C_A_0_00001.pdb 
# Loaded target data/PDB/CUUCGG/4v8a-pdb-bundle1_1691_C_A_0_00000.pdb 
# Loading data/PDB/CUUCGG/4v8a-pdb-bundle1_1691_C_A_0_00000.pdb 
# Loaded reference data/PDB/CUUCGG/2KOC_5_C_A_0_00001.pdb 
# Loaded target data/PDB/CUUCGG/1FMN_16_C_A_0_00001.pdb 
# Loading data/PDB/CUUCGG/1FMN_16_C_A_0_00001.pdb 
# Loaded reference data/PDB/CUUCGG/2KOC_5_C_A_0_00001.pdb 
# Loaded target data/PDB/CUUCGG/2N3Q_37_C_A_0_00017.pdb 
# Loading data/PDB/CUUCGG/2N3Q_37_C_A_0_00017.pdb 
# Loaded reference data/PDB/CUUCGG/2KOC_5_C_A_0_00001.pdb 
# Loaded target data/PDB/CUUCGG/1BYJ_11_C_A_0_00027.pdb 
# Loading data/PDB/CUUCGG/1BYJ_11_C_A_0_00027.pdb 
# Loaded reference data/PDB/CUUCGG/2KOC_5_C_A_0_00001.pdb 
# Loaded target data/PDB/CUUCGG/6b4v-pdb-bundle1_1691_C_B_0_00001.pdb 
# Loading data/PDB/CUUCGG/6b4v-pdb-bundle1_1691_C_B_0_00001.pdb 
# Loaded reference data/PDB/CUUCGG/2KOC_5_C_A_0_00001.pdb 
# Loaded target data/PDB/CUUCGG/5mdv-pdb-bundle1_1691_C_1_0_00000.pdb 
# Loading data/PDB/CUUCGG/5mdv-pdb-bundle1_1691_C_1_0_00000.pdb 
# Loaded reference data/PDB/CUUCGG/2KOC_5_C_A_0_00001.pdb 
# Loaded target data/PDB/CUUCGG/2M58_12_C_A_0_00003.pdb 
# Loading data/PDB/CUUCGG/2M58_12_C_A_0_00003.pdb 
# Loaded reference data/PDB/CUUCGG/2KOC_5_C_A_0_00001.pdb 
# Loaded target data/PDB/CUUCGG/1BYJ_11_C_A_0_00033.pdb 
# Loading data/PDB/CUUCGG/1BYJ_11_C_A_0_00033.pdb 
# Loaded reference data/PDB/CUUCGG/2KOC_5_C_A_0_00001.pdb 
# Loaded target data/PDB/CUUCGG/2N3Q_37_C_A_0_00003.pdb 
# Loading data/PDB/CUUCGG/2N3Q_37_C_A_0_00003.pdb 
# Loaded reference data/PDB/CUUCGG/2KOC_5_C_A_0_00001.pdb 
# Loaded target data/PDB/CUUCGG/6dzi-pdb-bundle1_419_C_h_0_00000.pdb 
# Loading data/PDB/CUUCGG/6dzi-pdb-bundle1_419_C_h_0_00000.pdb 
# Loaded reference data/PDB/CUUCGG/2KOC_5_C_A_0_00001.pdb 
# Loaded target data/PDB/CUUCGG/2LQZ_13_C_A_0_00002.pdb 
# Loading data/PDB/CUUCGG/2LQZ_13_C_A_0_00002.pdb 
# Loaded reference data/PDB/CUUCGG/2KOC_5_C_A_0_00001.pdb 
# Loaded target data/PDB/CUUCGG/4tue-pdb-bundle3_419_C_A_0_00000.pdb 
# Loading data/PDB/CUUCGG/4tue-pdb-bundle3_419_C_A_0_00000.pdb 
# Loaded reference data/PDB/CUUCGG/2KOC_5_C_A_0_00001.pdb 
# Loaded target data/PDB/CUUCGG/5ZET_1909_C_A_0_00000.pdb 
# Loading data/PDB/CUUCGG/5ZET_1909_C_A_0_00000.pdb 
# Loaded reference data/PDB/CUUCGG/2KOC_5_C_A_0_00001.pdb 
# Loaded target data/PDB/CUUCGG/2LHP_16_C_A_0_00006.pdb 
# Loading data/PDB/CUUCGG/2LHP_16_C_A_0_00006.pdb 
# Loaded reference data/PDB/CUUCGG/2KOC_5_C_A_0_00001.pdb 
# Loaded target data/PDB/CUUCGG/1B36_16_C_A_0_00006.pdb 
# Loading data/PDB/CUUCGG/1B36_16_C_A_0_00006.pdb 
# Loaded reference data/PDB/CUUCGG/2KOC_5_C_A_0_00001.pdb 
# Loaded target data/PDB/CUUCGG/2HUA_17_C_A_0_00003.pdb 
# Loading data/PDB/CUUCGG/2HUA_17_C_A_0_00003.pdb 
# Loaded reference data/PDB/CUUCGG/2KOC_5_C_A_0_00001.pdb 
# Loaded target data/PDB/CUUCGG/1A3M_11_C_A_0_00014.pdb 
# Loading data/PDB/CUUCGG/1A3M_11_C_A_0_00014.pdb 
# Loaded reference data/PDB/CUUCGG/2KOC_5_C_A_0_00001.pdb 
# Loaded target data/PDB/CUUCGG/4u24-pdb-bundle4_1691_C_A_0_00000.pdb 
# Loading data/PDB/CUUCGG/4u24-pdb-bundle4_1691_C_A_0_00000.pdb 
# Loaded reference data/PDB/CUUCGG/2KOC_5_C_A_0_00001.pdb 
# Loaded target data/PDB/CUUCGG/1JO7_14_C_A_0_00028.pdb 
# Loading data/PDB/CUUCGG/1JO7_14_C_A_0_00028.pdb 
# Loaded reference data/PDB/CUUCGG/2KOC_5_C_A_0_00001.pdb 
# Loaded target data/PDB/CUUCGG/1A3M_11_C_A_0_00000.pdb 
# Loading data/PDB/CUUCGG/1A3M_11_C_A_0_00000.pdb 
# Loaded reference data/PDB/CUUCGG/2KOC_5_C_A_0_00001.pdb 
# Loaded target data/PDB/CUUCGG/5KH8_26_C_A_0_00019.pdb 
# Loading data/PDB/CUUCGG/5KH8_26_C_A_0_00019.pdb 
# Loaded reference data/PDB/CUUCGG/2KOC_5_C_A_0_00001.pdb 
# Loaded target data/PDB/CUUCGG/5afi-pdb-bundle1_419_C_a_0_00001.pdb 
# Loading data/PDB/CUUCGG/5afi-pdb-bundle1_419_C_a_0_00001.pdb 
# Loaded reference data/PDB/CUUCGG/2KOC_5_C_A_0_00001.pdb 
# Loaded target data/PDB/CUUCGG/4v5q-pdb-bundle3_419_C_A_0_00000.pdb 
# Loading data/PDB/CUUCGG/4v5q-pdb-bundle3_419_C_A_0_00000.pdb 
# Loaded reference data/PDB/CUUCGG/2KOC_5_C_A_0_00001.pdb 
# Loaded target data/PDB/CUUCGG/1MFY_14_C_A_0_00009.pdb 
# Loading data/PDB/CUUCGG/1MFY_14_C_A_0_00009.pdb 
# Loaded reference data/PDB/CUUCGG/2KOC_5_C_A_0_00001.pdb 
# Loaded target data/PDB/CUUCGG/3OW2_1769_C_0_0_00000.pdb 
# Loading data/PDB/CUUCGG/3OW2_1769_C_0_0_00000.pdb 
# Loaded reference data/PDB/CUUCGG/2KOC_5_C_A_0_00001.pdb 
# Loaded target data/PDB/CUUCGG/1JO7_14_C_A_0_00014.pdb 
# Loading data/PDB/CUUCGG/1JO7_14_C_A_0_00014.pdb 
# Loaded reference data/PDB/CUUCGG/2KOC_5_C_A_0_00001.pdb 
# Loaded target data/PDB/CUUCGG/1YJW_1769_C_0_0_00000.pdb 
# Loading data/PDB/CUUCGG/1YJW_1769_C_0_0_00000.pdb 
# Loaded reference data/PDB/CUUCGG/2KOC_5_C_A_0_00001.pdb 
# Loaded target data/PDB/CUUCGG/5j5b-pdb-bundle1_419_C_A_0_00001.pdb 
# Loading data/PDB/CUUCGG/5j5b-pdb-bundle1_419_C_A_0_00001.pdb 
# Loaded reference data/PDB/CUUCGG/2KOC_5_C_A_0_00001.pdb 
# Loaded target data/PDB/CUUCGG/2GIP_12_C_A_0_00019.pdb 
# Loading data/PDB/CUUCGG/2GIP_12_C_A_0_00019.pdb 
# Loaded reference data/PDB/CUUCGG/2KOC_5_C_A_0_00001.pdb 
# Loaded target data/PDB/CUUCGG/4v7t-pdb-bundle2_1691_C_A_0_00000.pdb 
# Loading data/PDB/CUUCGG/4v7t-pdb-bundle2_1691_C_A_0_00000.pdb 
# Loaded reference data/PDB/CUUCGG/2KOC_5_C_A_0_00001.pdb 
# Loaded target data/PDB/CUUCGG/5el7-pdb-bundle2_1691_C_A_0_00000.pdb 
# Loading data/PDB/CUUCGG/5el7-pdb-bundle2_1691_C_A_0_00000.pdb 
# Loaded reference data/PDB/CUUCGG/2KOC_5_C_A_0_00001.pdb 
# Loaded target data/PDB/CUUCGG/5j8a-pdb-bundle1_1449_C_A_0_00003.pdb 
# Loading data/PDB/CUUCGG/5j8a-pdb-bundle1_1449_C_A_0_00003.pdb 
# Loaded reference data/PDB/CUUCGG/2KOC_5_C_A_0_00001.pdb 
# Loaded target data/PDB/CUUCGG/4v8x-pdb-bundle2_1691_C_A_0_00000.pdb 
# Loading data/PDB/CUUCGG/4v8x-pdb-bundle2_1691_C_A_0_00000.pdb 
# Loaded reference data/PDB/CUUCGG/2KOC_5_C_A_0_00001.pdb 
# Loaded target data/PDB/CUUCGG/5O5J_419_C_A_0_00000.pdb 
# Loading data/PDB/CUUCGG/5O5J_419_C_A_0_00000.pdb 
# Loaded reference data/PDB/CUUCGG/2KOC_5_C_A_0_00001.pdb 
# Loaded target data/PDB/CUUCGG/2L3E_18_C_A_0_00014.pdb 
# Loading data/PDB/CUUCGG/2L3E_18_C_A_0_00014.pdb 
# Loaded reference data/PDB/CUUCGG/2KOC_5_C_A_0_00001.pdb 
# Loaded target data/PDB/CUUCGG/2N3R_37_C_A_0_00013.pdb 
# Loading data/PDB/CUUCGG/2N3R_37_C_A_0_00013.pdb 
# Loaded reference data/PDB/CUUCGG/2KOC_5_C_A_0_00001.pdb 
# Loaded target data/PDB/CUUCGG/6c4i-pdb-bundle1_1691_C_A_0_00000.pdb 
# Loading data/PDB/CUUCGG/6c4i-pdb-bundle1_1691_C_A_0_00000.pdb 
# Loaded reference data/PDB/CUUCGG/2KOC_5_C_A_0_00001.pdb 
# Loaded target data/PDB/CUUCGG/2N3R_37_C_A_0_00007.pdb 
# Loading data/PDB/CUUCGG/2N3R_37_C_A_0_00007.pdb 
# Loaded reference data/PDB/CUUCGG/2KOC_5_C_A_0_00001.pdb 
# Loaded target data/PDB/CUUCGG/4zer-pdb-bundle1_1738_C_A_0_00000.pdb 
# Loading data/PDB/CUUCGG/4zer-pdb-bundle1_1738_C_A_0_00000.pdb 
# Loaded reference data/PDB/CUUCGG/2KOC_5_C_A_0_00001.pdb 
# Loaded target data/PDB/CUUCGG/2L3E_18_C_A_0_00000.pdb 
# Loading data/PDB/CUUCGG/2L3E_18_C_A_0_00000.pdb 
# Loaded reference data/PDB/CUUCGG/2KOC_5_C_A_0_00001.pdb 
# Loaded target data/PDB/CUUCGG/5wdt-pdb-bundle1_1691_C_A_0_00000.pdb 
# Loading data/PDB/CUUCGG/5wdt-pdb-bundle1_1691_C_A_0_00000.pdb 
# Loaded reference data/PDB/CUUCGG/2KOC_5_C_A_0_00001.pdb 
# Loaded target data/PDB/CUUCGG/4v7z-pdb-bundle4_1691_C_A_0_00000.pdb 
# Loading data/PDB/CUUCGG/4v7z-pdb-bundle4_1691_C_A_0_00000.pdb 
# Loaded reference data/PDB/CUUCGG/2KOC_5_C_A_0_00001.pdb 
# Loaded target data/PDB/CUUCGG/4v87-pdb-bundle1_1691_C_A_0_00000.pdb 
# Loading data/PDB/CUUCGG/4v87-pdb-bundle1_1691_C_A_0_00000.pdb 
# Loaded reference data/PDB/CUUCGG/2KOC_5_C_A_0_00001.pdb 
# Loaded target data/PDB/CUUCGG/2LU0_28_C_A_0_00019.pdb 
# Loading data/PDB/CUUCGG/2LU0_28_C_A_0_00019.pdb 
# Loaded reference data/PDB/CUUCGG/2KOC_5_C_A_0_00001.pdb 
# Loaded target data/PDB/CUUCGG/4lnt-pdb-bundle3_419_C_A_0_00000.pdb 
# Loading data/PDB/CUUCGG/4lnt-pdb-bundle3_419_C_A_0_00000.pdb 
# Loaded reference data/PDB/CUUCGG/2KOC_5_C_A_0_00001.pdb 
# Loaded target data/PDB/CUUCGG/6c4i-pdb-bundle2_1449_C_a_0_00003.pdb 
# Loading data/PDB/CUUCGG/6c4i-pdb-bundle2_1449_C_a_0_00003.pdb 
# Loaded reference data/PDB/CUUCGG/2KOC_5_C_A_0_00001.pdb 
# Loaded target data/PDB/CUUCGG/5dc3-pdb-bundle2_1923_C_A_0_00002.pdb 
# Loading data/PDB/CUUCGG/5dc3-pdb-bundle2_1923_C_A_0_00002.pdb 
# Loaded reference data/PDB/CUUCGG/2KOC_5_C_A_0_00001.pdb 
# Loaded target data/PDB/CUUCGG/5wdt-pdb-bundle2_1449_C_a_0_00003.pdb 
# Loading data/PDB/CUUCGG/5wdt-pdb-bundle2_1449_C_a_0_00003.pdb 
# Loaded reference data/PDB/CUUCGG/2KOC_5_C_A_0_00001.pdb 
# Loaded target data/PDB/CUUCGG/4ybb-pdb-bundle1_419_C_V_0_00005.pdb 
# Loading data/PDB/CUUCGG/4ybb-pdb-bundle1_419_C_V_0_00005.pdb 
# Loaded reference data/PDB/CUUCGG/2KOC_5_C_A_0_00001.pdb 
# Loaded target data/PDB/CUUCGG/2LU0_28_C_A_0_00009.pdb 
# Loading data/PDB/CUUCGG/2LU0_28_C_A_0_00009.pdb 
# Loaded reference data/PDB/CUUCGG/2KOC_5_C_A_0_00001.pdb 
# Loaded target data/PDB/CUUCGG/4v7v-pdb-bundle3_207_C_A_0_00000.pdb 
# Loading data/PDB/CUUCGG/4v7v-pdb-bundle3_207_C_A_0_00000.pdb 
# Loaded reference data/PDB/CUUCGG/2KOC_5_C_A_0_00001.pdb 
# Loaded target data/PDB/CUUCGG/4v9k-pdb-bundle1_419_C_U_0_00000.pdb 
# Loading data/PDB/CUUCGG/4v9k-pdb-bundle1_419_C_U_0_00000.pdb 
# Loaded reference data/PDB/CUUCGG/2KOC_5_C_A_0_00001.pdb 
# Loaded target data/PDB/CUUCGG/2L3E_18_C_A_0_00004.pdb 
# Loading data/PDB/CUUCGG/2L3E_18_C_A_0_00004.pdb 
# Loaded reference data/PDB/CUUCGG/2KOC_5_C_A_0_00001.pdb 
# Loaded target data/PDB/CUUCGG/5j5b-pdb-bundle1_1449_C_b_0_00007.pdb 
# Loading data/PDB/CUUCGG/5j5b-pdb-bundle1_1449_C_b_0_00007.pdb 
# Loaded reference data/PDB/CUUCGG/2KOC_5_C_A_0_00001.pdb 
# Loaded target data/PDB/CUUCGG/2N3R_37_C_A_0_00003.pdb 
# Loading data/PDB/CUUCGG/2N3R_37_C_A_0_00003.pdb 
# Loaded reference data/PDB/CUUCGG/2KOC_5_C_A_0_00001.pdb 
# Loaded target data/PDB/CUUCGG/6gsl-pdb-bundle2_1691_C_A_0_00000.pdb 
# Loading data/PDB/CUUCGG/6gsl-pdb-bundle2_1691_C_A_0_00000.pdb 
# Loaded reference data/PDB/CUUCGG/2KOC_5_C_A_0_00001.pdb 
# Loaded target data/PDB/CUUCGG/2N3R_37_C_A_0_00017.pdb 
# Loading data/PDB/CUUCGG/2N3R_37_C_A_0_00017.pdb 
# Loaded reference data/PDB/CUUCGG/2KOC_5_C_A_0_00001.pdb 
# Loaded target data/PDB/CUUCGG/2L3E_18_C_A_0_00010.pdb 
# Loading data/PDB/CUUCGG/2L3E_18_C_A_0_00010.pdb 
# Loaded reference data/PDB/CUUCGG/2KOC_5_C_A_0_00001.pdb 
# Loaded target data/PDB/CUUCGG/4v64-pdb-bundle1_1028_C_A_0_00002.pdb 
# Loading data/PDB/CUUCGG/4v64-pdb-bundle1_1028_C_A_0_00002.pdb 
# Loaded reference data/PDB/CUUCGG/2KOC_5_C_A_0_00001.pdb 
# Loaded target data/PDB/CUUCGG/2GIP_12_C_A_0_00009.pdb 
# Loading data/PDB/CUUCGG/2GIP_12_C_A_0_00009.pdb 
# Loaded reference data/PDB/CUUCGG/2KOC_5_C_A_0_00001.pdb 
# Loaded target data/PDB/CUUCGG/5mei-pdb-bundle3_1923_C_A_0_00002.pdb 
# Loading data/PDB/CUUCGG/5mei-pdb-bundle3_1923_C_A_0_00002.pdb 
# Loaded reference data/PDB/CUUCGG/2KOC_5_C_A_0_00001.pdb 
# Loaded target data/PDB/CUUCGG/5h4p-pdb-bundle1_1923_C_1_0_00002.pdb 
# Loading data/PDB/CUUCGG/5h4p-pdb-bundle1_1923_C_1_0_00002.pdb 
# Loaded reference data/PDB/CUUCGG/2KOC_5_C_A_0_00001.pdb 
# Loaded target data/PDB/CUUCGG/1A3M_11_C_A_0_00004.pdb 
# Loading data/PDB/CUUCGG/1A3M_11_C_A_0_00004.pdb 
# Loaded reference data/PDB/CUUCGG/2KOC_5_C_A_0_00001.pdb 
# Loaded target data/PDB/CUUCGG/4v7t-pdb-bundle1_1449_C_A_0_00003.pdb 
# Loading data/PDB/CUUCGG/4v7t-pdb-bundle1_1449_C_A_0_00003.pdb 
# Loaded reference data/PDB/CUUCGG/2KOC_5_C_A_0_00001.pdb 
# Loaded target data/PDB/CUUCGG/5mmm-pdb-bundle1_1727_C_A_0_00001.pdb 
# Loading data/PDB/CUUCGG/5mmm-pdb-bundle1_1727_C_A_0_00001.pdb 
# Loaded reference data/PDB/CUUCGG/2KOC_5_C_A_0_00001.pdb 
# Loaded target data/PDB/CUUCGG/4v7s-pdb-bundle1_207_C_A_0_00000.pdb 
# Loading data/PDB/CUUCGG/4v7s-pdb-bundle1_207_C_A_0_00000.pdb 
# Loaded reference data/PDB/CUUCGG/2KOC_5_C_A_0_00001.pdb 
# Loaded target data/PDB/CUUCGG/1A3M_11_C_A_0_00010.pdb 
# Loading data/PDB/CUUCGG/1A3M_11_C_A_0_00010.pdb 
# Loaded reference data/PDB/CUUCGG/2KOC_5_C_A_0_00001.pdb 
# Loaded target data/PDB/CUUCGG/5KH8_26_C_A_0_00009.pdb 
# Loading data/PDB/CUUCGG/5KH8_26_C_A_0_00009.pdb 
# Loaded reference data/PDB/CUUCGG/2KOC_5_C_A_0_00001.pdb 
# Loaded target data/PDB/CUUCGG/3CMA_1769_C_0_0_00000.pdb 
# Loading data/PDB/CUUCGG/3CMA_1769_C_0_0_00000.pdb 
# Loaded reference data/PDB/CUUCGG/2KOC_5_C_A_0_00001.pdb 
# Loaded target data/PDB/CUUCGG/1vvj-pdb-bundle1_419_C_A_0_00000.pdb 
# Loading data/PDB/CUUCGG/1vvj-pdb-bundle1_419_C_A_0_00000.pdb 
# Loaded reference data/PDB/CUUCGG/2KOC_5_C_A_0_00001.pdb 
# Loaded target data/PDB/CUUCGG/4u6f-pdb-bundle4_1923_C_I_0_00002.pdb 
# Loading data/PDB/CUUCGG/4u6f-pdb-bundle4_1923_C_I_0_00002.pdb 
# Loaded reference data/PDB/CUUCGG/2KOC_5_C_A_0_00001.pdb 
# Loaded target data/PDB/CUUCGG/3j79-pdb-bundle1_2191_C_A_0_00000.pdb 
# Loading data/PDB/CUUCGG/3j79-pdb-bundle1_2191_C_A_0_00000.pdb 
# Loaded reference data/PDB/CUUCGG/2KOC_5_C_A_0_00001.pdb 
# Loaded target data/PDB/CUUCGG/2N3Q_37_C_A_0_00007.pdb 
# Loading data/PDB/CUUCGG/2N3Q_37_C_A_0_00007.pdb 
# Loaded reference data/PDB/CUUCGG/2KOC_5_C_A_0_00001.pdb 
# Loaded target data/PDB/CUUCGG/1BYJ_11_C_A_0_00037.pdb 
# Loading data/PDB/CUUCGG/1BYJ_11_C_A_0_00037.pdb 
# Loaded reference data/PDB/CUUCGG/2KOC_5_C_A_0_00001.pdb 
# Loaded target data/PDB/CUUCGG/2M58_12_C_A_0_00007.pdb 
# Loading data/PDB/CUUCGG/2M58_12_C_A_0_00007.pdb 
# Loaded reference data/PDB/CUUCGG/2KOC_5_C_A_0_00001.pdb 
# Loaded target data/PDB/CUUCGG/1BYJ_11_C_A_0_00023.pdb 
# Loading data/PDB/CUUCGG/1BYJ_11_C_A_0_00023.pdb 
# Loaded reference data/PDB/CUUCGG/2KOC_5_C_A_0_00001.pdb 
# Loaded target data/PDB/CUUCGG/2N3Q_37_C_A_0_00013.pdb 
# Loading data/PDB/CUUCGG/2N3Q_37_C_A_0_00013.pdb 
# Loaded reference data/PDB/CUUCGG/2KOC_5_C_A_0_00001.pdb 
# Loaded target data/PDB/CUUCGG/2HUA_17_C_A_0_00007.pdb 
# Loading data/PDB/CUUCGG/2HUA_17_C_A_0_00007.pdb 
# Loaded reference data/PDB/CUUCGG/2KOC_5_C_A_0_00001.pdb 
# Loaded target data/PDB/CUUCGG/1B36_16_C_A_0_00002.pdb 
# Loading data/PDB/CUUCGG/1B36_16_C_A_0_00002.pdb 
# Loaded reference data/PDB/CUUCGG/2KOC_5_C_A_0_00001.pdb 
# Loaded target data/PDB/CUUCGG/2LHP_16_C_A_0_00002.pdb 
# Loading data/PDB/CUUCGG/2LHP_16_C_A_0_00002.pdb 
# Loaded reference data/PDB/CUUCGG/2KOC_5_C_A_0_00001.pdb 
# Loaded target data/PDB/CUUCGG/2LQZ_13_C_A_0_00006.pdb 
# Loading data/PDB/CUUCGG/2LQZ_13_C_A_0_00006.pdb 
# Loaded reference data/PDB/CUUCGG/2KOC_5_C_A_0_00001.pdb 
# Loaded target data/PDB/CUUCGG/1D6K_283_C_B_0_00007.pdb 
# Loading data/PDB/CUUCGG/1D6K_283_C_B_0_00007.pdb 
# Loaded reference data/PDB/CUUCGG/2KOC_5_C_A_0_00001.pdb 
# Loaded target data/PDB/CUUCGG/5Y87_28_C_B_0_00000.pdb 
# Loading data/PDB/CUUCGG/5Y87_28_C_B_0_00000.pdb 
# Loaded reference data/PDB/CUUCGG/2KOC_5_C_A_0_00001.pdb 
# Loaded target data/PDB/CUUCGG/2NCI_14_C_A_0_00000.pdb 
# Loading data/PDB/CUUCGG/2NCI_14_C_A_0_00000.pdb 
# Loaded reference data/PDB/CUUCGG/2KOC_5_C_A_0_00001.pdb 
# Loaded target data/PDB/CUUCGG/1D6K_283_C_B_0_00013.pdb 
# Loading data/PDB/CUUCGG/1D6K_283_C_B_0_00013.pdb 
# Loaded reference data/PDB/CUUCGG/2KOC_5_C_A_0_00001.pdb 
# Loaded target data/PDB/CUUCGG/4v9o-pdb-bundle2_419_C_A_0_00001.pdb 
# Loading data/PDB/CUUCGG/4v9o-pdb-bundle2_419_C_A_0_00001.pdb 
# Loaded reference data/PDB/CUUCGG/2KOC_5_C_A_0_00001.pdb 
# Loaded target data/PDB/CUUCGG/4u4n-pdb-bundle4_1923_C_I_0_00002.pdb 
# Loading data/PDB/CUUCGG/4u4n-pdb-bundle4_1923_C_I_0_00002.pdb 
# Loaded reference data/PDB/CUUCGG/2KOC_5_C_A_0_00001.pdb 
# Loaded target data/PDB/CUUCGG/4PDB_17_C_I_0_00000.pdb 
# Loading data/PDB/CUUCGG/4PDB_17_C_I_0_00000.pdb 
# Loaded reference data/PDB/CUUCGG/2KOC_5_C_A_0_00001.pdb 
# Loaded target data/PDB/CUUCGG/4B3T_419_C_A_0_00000.pdb 
# Loading data/PDB/CUUCGG/4B3T_419_C_A_0_00000.pdb 
# Loaded reference data/PDB/CUUCGG/2KOC_5_C_A_0_00001.pdb 
# Loaded target data/PDB/CUUCGG/2LUB_16_C_A_0_00007.pdb 
# Loading data/PDB/CUUCGG/2LUB_16_C_A_0_00007.pdb 
# Loaded reference data/PDB/CUUCGG/2KOC_5_C_A_0_00001.pdb 
# Loaded target data/PDB/CUUCGG/2LWK_15_C_A_0_00005.pdb 
# Loading data/PDB/CUUCGG/2LWK_15_C_A_0_00005.pdb 
# Loaded reference data/PDB/CUUCGG/2KOC_5_C_A_0_00001.pdb 
# Loaded target data/PDB/CUUCGG/1TLR_10_C_A_0_00008.pdb 
# Loading data/PDB/CUUCGG/1TLR_10_C_A_0_00008.pdb 
# Loaded reference data/PDB/CUUCGG/2KOC_5_C_A_0_00001.pdb 
# Loaded target data/PDB/CUUCGG/1P6V_26_C_D_0_00001.pdb 
# Loading data/PDB/CUUCGG/1P6V_26_C_D_0_00001.pdb 
# Loaded reference data/PDB/CUUCGG/2KOC_5_C_A_0_00001.pdb 
# Loaded target data/PDB/CUUCGG/2LWK_15_C_A_0_00011.pdb 
# Loading data/PDB/CUUCGG/2LWK_15_C_A_0_00011.pdb 
# Loaded reference data/PDB/CUUCGG/2KOC_5_C_A_0_00001.pdb 
# Loaded target data/PDB/CUUCGG/5hau-pdb-bundle3_1737_C_A_0_00000.pdb 
# Loading data/PDB/CUUCGG/5hau-pdb-bundle3_1737_C_A_0_00000.pdb 
# Loaded reference data/PDB/CUUCGG/2KOC_5_C_A_0_00001.pdb 
# Loaded target data/PDB/CUUCGG/1FYO_11_C_A_0_00008.pdb 
# Loading data/PDB/CUUCGG/1FYO_11_C_A_0_00008.pdb 
# Loaded reference data/PDB/CUUCGG/2KOC_5_C_A_0_00001.pdb 
# Loaded target data/PDB/CUUCGG/5KQE_19_C_A_0_00008.pdb 
# Loading data/PDB/CUUCGG/5KQE_19_C_A_0_00008.pdb 
# Loaded reference data/PDB/CUUCGG/2KOC_5_C_A_0_00001.pdb 
# Loaded target data/PDB/CUUCGG/2N6S_16_C_A_0_00006.pdb 
# Loading data/PDB/CUUCGG/2N6S_16_C_A_0_00006.pdb 
# Loaded reference data/PDB/CUUCGG/2KOC_5_C_A_0_00001.pdb 
# Loaded target data/PDB/CUUCGG/1KP7_14_C_A_0_00000.pdb 
# Loading data/PDB/CUUCGG/1KP7_14_C_A_0_00000.pdb 
# Loaded reference data/PDB/CUUCGG/2KOC_5_C_A_0_00001.pdb 
# Loaded target data/PDB/CUUCGG/1FYO_11_C_A_0_00020.pdb 
# Loading data/PDB/CUUCGG/1FYO_11_C_A_0_00020.pdb 
# Loaded reference data/PDB/CUUCGG/2KOC_5_C_A_0_00001.pdb 
# Loaded target data/PDB/CUUCGG/4v8g-pdb-bundle2_1691_C_A_0_00000.pdb 
# Loading data/PDB/CUUCGG/4v8g-pdb-bundle2_1691_C_A_0_00000.pdb 
# Loaded reference data/PDB/CUUCGG/2KOC_5_C_A_0_00001.pdb 
# Loaded target data/PDB/CUUCGG/1vvj-pdb-bundle4_1691_C_A_0_00000.pdb 
# Loading data/PDB/CUUCGG/1vvj-pdb-bundle4_1691_C_A_0_00000.pdb 
# Loaded reference data/PDB/CUUCGG/2KOC_5_C_A_0_00001.pdb 
# Loaded target data/PDB/CUUCGG/5wfk-pdb-bundle2_1449_C_a_0_00003.pdb 
# Loading data/PDB/CUUCGG/5wfk-pdb-bundle2_1449_C_a_0_00003.pdb 
# Loaded reference data/PDB/CUUCGG/2KOC_5_C_A_0_00001.pdb 
# Loaded target data/PDB/CUUCGG/1FYO_11_C_A_0_00021.pdb 
# Loading data/PDB/CUUCGG/1FYO_11_C_A_0_00021.pdb 
# Loaded reference data/PDB/CUUCGG/2KOC_5_C_A_0_00001.pdb 
# Loaded target data/PDB/CUUCGG/1FYO_11_C_A_0_00009.pdb 
# Loading data/PDB/CUUCGG/1FYO_11_C_A_0_00009.pdb 
# Loaded reference data/PDB/CUUCGG/2KOC_5_C_A_0_00001.pdb 
# Loaded target data/PDB/CUUCGG/2N6S_16_C_A_0_00007.pdb 
# Loading data/PDB/CUUCGG/2N6S_16_C_A_0_00007.pdb 
# Loaded reference data/PDB/CUUCGG/2KOC_5_C_A_0_00001.pdb 
# Loaded target data/PDB/CUUCGG/5KQE_19_C_A_0_00009.pdb 
# Loading data/PDB/CUUCGG/5KQE_19_C_A_0_00009.pdb 
# Loaded reference data/PDB/CUUCGG/2KOC_5_C_A_0_00001.pdb 
# Loaded target data/PDB/CUUCGG/6bz6-pdb-bundle2_1691_C_A_0_00000.pdb 
# Loading data/PDB/CUUCGG/6bz6-pdb-bundle2_1691_C_A_0_00000.pdb 
# Loaded reference data/PDB/CUUCGG/2KOC_5_C_A_0_00001.pdb 
# Loaded target data/PDB/CUUCGG/2LWK_15_C_A_0_00010.pdb 
# Loading data/PDB/CUUCGG/2LWK_15_C_A_0_00010.pdb 
# Loaded reference data/PDB/CUUCGG/2KOC_5_C_A_0_00001.pdb 
# Loaded target data/PDB/CUUCGG/4u26-pdb-bundle3_419_C_A_0_00001.pdb 
# Loading data/PDB/CUUCGG/4u26-pdb-bundle3_419_C_A_0_00001.pdb 
# Loaded reference data/PDB/CUUCGG/2KOC_5_C_A_0_00001.pdb 
# Loaded target data/PDB/CUUCGG/1TLR_10_C_A_0_00009.pdb 
# Loading data/PDB/CUUCGG/1TLR_10_C_A_0_00009.pdb 
# Loaded reference data/PDB/CUUCGG/2KOC_5_C_A_0_00001.pdb 
# Loaded target data/PDB/CUUCGG/2LWK_15_C_A_0_00004.pdb 
# Loading data/PDB/CUUCGG/2LWK_15_C_A_0_00004.pdb 
# Loaded reference data/PDB/CUUCGG/2KOC_5_C_A_0_00001.pdb 
# Loaded target data/PDB/CUUCGG/1YJ9_1769_C_0_0_00000.pdb 
# Loading data/PDB/CUUCGG/1YJ9_1769_C_0_0_00000.pdb 
# Loaded reference data/PDB/CUUCGG/2KOC_5_C_A_0_00001.pdb 
# Loaded target data/PDB/CUUCGG/5X8T_1727_C_A_0_00001.pdb 
# Loading data/PDB/CUUCGG/5X8T_1727_C_A_0_00001.pdb 
# Loaded reference data/PDB/CUUCGG/2KOC_5_C_A_0_00001.pdb 
# Loaded target data/PDB/CUUCGG/4woi-pdb-bundle4_419_C_A_0_00001.pdb 
# Loading data/PDB/CUUCGG/4woi-pdb-bundle4_419_C_A_0_00001.pdb 
# Loaded reference data/PDB/CUUCGG/2KOC_5_C_A_0_00001.pdb 
# Loaded target data/PDB/CUUCGG/4v9q-pdb-bundle1_1691_C_A_0_00000.pdb 
# Loading data/PDB/CUUCGG/4v9q-pdb-bundle1_1691_C_A_0_00000.pdb 
# Loaded reference data/PDB/CUUCGG/2KOC_5_C_A_0_00001.pdb 
# Loaded target data/PDB/CUUCGG/4v8i-pdb-bundle4_1691_C_A_0_00000.pdb 
# Loading data/PDB/CUUCGG/4v8i-pdb-bundle4_1691_C_A_0_00000.pdb 
# Loaded reference data/PDB/CUUCGG/2KOC_5_C_A_0_00001.pdb 
# Loaded target data/PDB/CUUCGG/2NCI_14_C_A_0_00001.pdb 
# Loading data/PDB/CUUCGG/2NCI_14_C_A_0_00001.pdb 
# Loaded reference data/PDB/CUUCGG/2KOC_5_C_A_0_00001.pdb 
# Loaded target data/PDB/CUUCGG/1D6K_283_C_B_0_00012.pdb 
# Loading data/PDB/CUUCGG/1D6K_283_C_B_0_00012.pdb 
# Loaded reference data/PDB/CUUCGG/2KOC_5_C_A_0_00001.pdb 
# Loaded target data/PDB/CUUCGG/1D6K_283_C_B_0_00006.pdb 
# Loading data/PDB/CUUCGG/1D6K_283_C_B_0_00006.pdb 
# Loaded reference data/PDB/CUUCGG/2KOC_5_C_A_0_00001.pdb 
# Loaded target data/PDB/CUUCGG/5lzd-pdb-bundle1_207_C_a_0_00000.pdb 
# Loading data/PDB/CUUCGG/5lzd-pdb-bundle1_207_C_a_0_00000.pdb 
# Loaded reference data/PDB/CUUCGG/2KOC_5_C_A_0_00001.pdb 
# Loaded target data/PDB/CUUCGG/4u4y-pdb-bundle4_1923_C_I_0_00002.pdb 
# Loading data/PDB/CUUCGG/4u4y-pdb-bundle4_1923_C_I_0_00002.pdb 
# Loaded reference data/PDB/CUUCGG/2KOC_5_C_A_0_00001.pdb 
# Loaded target data/PDB/CUUCGG/2LQZ_13_C_A_0_00007.pdb 
# Loading data/PDB/CUUCGG/2LQZ_13_C_A_0_00007.pdb 
# Loaded reference data/PDB/CUUCGG/2KOC_5_C_A_0_00001.pdb 
# Loaded target data/PDB/CUUCGG/2LHP_16_C_A_0_00003.pdb 
# Loading data/PDB/CUUCGG/2LHP_16_C_A_0_00003.pdb 
# Loaded reference data/PDB/CUUCGG/2KOC_5_C_A_0_00001.pdb 
# Loaded target data/PDB/CUUCGG/2HUA_17_C_A_0_00006.pdb 
# Loading data/PDB/CUUCGG/2HUA_17_C_A_0_00006.pdb 
# Loaded reference data/PDB/CUUCGG/2KOC_5_C_A_0_00001.pdb 
# Loaded target data/PDB/CUUCGG/2M58_12_C_A_0_00006.pdb 
# Loading data/PDB/CUUCGG/2M58_12_C_A_0_00006.pdb 
# Loaded reference data/PDB/CUUCGG/2KOC_5_C_A_0_00001.pdb 
# Loaded target data/PDB/CUUCGG/2N3Q_37_C_A_0_00012.pdb 
# Loading data/PDB/CUUCGG/2N3Q_37_C_A_0_00012.pdb 
# Loaded reference data/PDB/CUUCGG/2KOC_5_C_A_0_00001.pdb 
# Loaded target data/PDB/CUUCGG/1FMN_16_C_A_0_00004.pdb 
# Loading data/PDB/CUUCGG/1FMN_16_C_A_0_00004.pdb 
# Loaded reference data/PDB/CUUCGG/2KOC_5_C_A_0_00001.pdb 
# Loaded target data/PDB/CUUCGG/2LKR_63_C_A_0_00009.pdb 
# Loading data/PDB/CUUCGG/2LKR_63_C_A_0_00009.pdb 
# Loaded reference data/PDB/CUUCGG/2KOC_5_C_A_0_00001.pdb 
# Loaded target data/PDB/CUUCGG/1BYJ_11_C_A_0_00022.pdb 
# Loading data/PDB/CUUCGG/1BYJ_11_C_A_0_00022.pdb 
# Loaded reference data/PDB/CUUCGG/2KOC_5_C_A_0_00001.pdb 
# Loaded target data/PDB/CUUCGG/1BYJ_11_C_A_0_00036.pdb 
# Loading data/PDB/CUUCGG/1BYJ_11_C_A_0_00036.pdb 
# Loaded reference data/PDB/CUUCGG/2KOC_5_C_A_0_00001.pdb 
# Loaded target data/PDB/CUUCGG/2N3Q_37_C_A_0_00006.pdb 
# Loading data/PDB/CUUCGG/2N3Q_37_C_A_0_00006.pdb 
# Loaded reference data/PDB/CUUCGG/2KOC_5_C_A_0_00001.pdb 
# Loaded target data/PDB/CUUCGG/5it8-pdb-bundle1_419_C_A_0_00001.pdb 
# Loading data/PDB/CUUCGG/5it8-pdb-bundle1_419_C_A_0_00001.pdb 
# Loaded reference data/PDB/CUUCGG/2KOC_5_C_A_0_00001.pdb 
# Loaded target data/PDB/CUUCGG/4wqu-pdb-bundle3_1691_C_A_0_00000.pdb 
# Loading data/PDB/CUUCGG/4wqu-pdb-bundle3_1691_C_A_0_00000.pdb 
# Loaded reference data/PDB/CUUCGG/2KOC_5_C_A_0_00001.pdb 
# Loaded target data/PDB/CUUCGG/5KH8_26_C_A_0_00008.pdb 
# Loading data/PDB/CUUCGG/5KH8_26_C_A_0_00008.pdb 
# Loaded reference data/PDB/CUUCGG/2KOC_5_C_A_0_00001.pdb 
# Loaded target data/PDB/CUUCGG/1JO7_14_C_A_0_00011.pdb 
# Loading data/PDB/CUUCGG/1JO7_14_C_A_0_00011.pdb 
# Loaded reference data/PDB/CUUCGG/2KOC_5_C_A_0_00001.pdb 
# Loaded target data/PDB/CUUCGG/5mei-pdb-bundle1_1923_C_1_0_00002.pdb 
# Loading data/PDB/CUUCGG/5mei-pdb-bundle1_1923_C_1_0_00002.pdb 
# Loaded reference data/PDB/CUUCGG/2KOC_5_C_A_0_00001.pdb 
# Loaded target data/PDB/CUUCGG/1A3M_11_C_A_0_00011.pdb 
# Loading data/PDB/CUUCGG/1A3M_11_C_A_0_00011.pdb 
# Loaded reference data/PDB/CUUCGG/2KOC_5_C_A_0_00001.pdb 
# Loaded target data/PDB/CUUCGG/4z3s-pdb-bundle1_1691_C_A_0_00000.pdb 
# Loading data/PDB/CUUCGG/4z3s-pdb-bundle1_1691_C_A_0_00000.pdb 
# Loaded reference data/PDB/CUUCGG/2KOC_5_C_A_0_00001.pdb 
# Loaded target data/PDB/CUUCGG/1A3M_11_C_A_0_00005.pdb 
# Loading data/PDB/CUUCGG/1A3M_11_C_A_0_00005.pdb 
# Loaded reference data/PDB/CUUCGG/2KOC_5_C_A_0_00001.pdb 
# Loaded target data/PDB/CUUCGG/4wq1-pdb-bundle4_1691_C_A_0_00000.pdb 
# Loading data/PDB/CUUCGG/4wq1-pdb-bundle4_1691_C_A_0_00000.pdb 
# Loaded reference data/PDB/CUUCGG/2KOC_5_C_A_0_00001.pdb 
# Loaded target data/PDB/CUUCGG/4wf1-pdb-bundle3_1028_C_A_0_00002.pdb 
# Loading data/PDB/CUUCGG/4wf1-pdb-bundle3_1028_C_A_0_00002.pdb 
# Loaded reference data/PDB/CUUCGG/2KOC_5_C_A_0_00001.pdb 
# Loaded target data/PDB/CUUCGG/4v9h-pdb-bundle2_1691_C_A_0_00000.pdb 
# Loading data/PDB/CUUCGG/4v9h-pdb-bundle2_1691_C_A_0_00000.pdb 
# Loaded reference data/PDB/CUUCGG/2KOC_5_C_A_0_00001.pdb 
# Loaded target data/PDB/CUUCGG/5dgv-pdb-bundle3_230_C_b_0_00000.pdb 
# Loading data/PDB/CUUCGG/5dgv-pdb-bundle3_230_C_b_0_00000.pdb 
# Loaded reference data/PDB/CUUCGG/2KOC_5_C_A_0_00001.pdb 
# Loaded target data/PDB/CUUCGG/2GIP_12_C_A_0_00008.pdb 
# Loading data/PDB/CUUCGG/2GIP_12_C_A_0_00008.pdb 
# Loaded reference data/PDB/CUUCGG/2KOC_5_C_A_0_00001.pdb 
# Loaded target data/PDB/CUUCGG/2L3E_18_C_A_0_00011.pdb 
# Loading data/PDB/CUUCGG/2L3E_18_C_A_0_00011.pdb 
# Loaded reference data/PDB/CUUCGG/2KOC_5_C_A_0_00001.pdb 
# Loaded target data/PDB/CUUCGG/4u4r-pdb-bundle3_230_C_d_0_00000.pdb 
# Loading data/PDB/CUUCGG/4u4r-pdb-bundle3_230_C_d_0_00000.pdb 
# Loaded reference data/PDB/CUUCGG/2KOC_5_C_A_0_00001.pdb 
# Loaded target data/PDB/CUUCGG/2N3R_37_C_A_0_00016.pdb 
# Loading data/PDB/CUUCGG/2N3R_37_C_A_0_00016.pdb 
# Loaded reference data/PDB/CUUCGG/2KOC_5_C_A_0_00001.pdb 
# Loaded target data/PDB/CUUCGG/6gz5-pdb-bundle1_2851_C_A_0_00000.pdb 
# Loading data/PDB/CUUCGG/6gz5-pdb-bundle1_2851_C_A_0_00000.pdb 
# Loaded reference data/PDB/CUUCGG/2KOC_5_C_A_0_00001.pdb 
# Loaded target data/PDB/CUUCGG/2N3R_37_C_A_0_00002.pdb 
# Loading data/PDB/CUUCGG/2N3R_37_C_A_0_00002.pdb 
# Loaded reference data/PDB/CUUCGG/2KOC_5_C_A_0_00001.pdb 
# Loaded target data/PDB/CUUCGG/2L3E_18_C_A_0_00005.pdb 
# Loading data/PDB/CUUCGG/2L3E_18_C_A_0_00005.pdb 
# Loaded reference data/PDB/CUUCGG/2KOC_5_C_A_0_00001.pdb 
# Loaded target data/PDB/CUUCGG/5u9f-pdb-bundle1_1691_C_B_0_00004.pdb 
# Loading data/PDB/CUUCGG/5u9f-pdb-bundle1_1691_C_B_0_00004.pdb 
# Loaded reference data/PDB/CUUCGG/2KOC_5_C_A_0_00001.pdb 
# Loaded target data/PDB/CUUCGG/5h5u-pdb-bundle1_1691_C_A_0_00000.pdb 
# Loading data/PDB/CUUCGG/5h5u-pdb-bundle1_1691_C_A_0_00000.pdb 
# Loaded reference data/PDB/CUUCGG/2KOC_5_C_A_0_00001.pdb 
# Loaded target data/PDB/CUUCGG/4v5c-pdb-bundle2_1691_C_A_0_00000.pdb 
# Loading data/PDB/CUUCGG/4v5c-pdb-bundle2_1691_C_A_0_00000.pdb 
# Loaded reference data/PDB/CUUCGG/2KOC_5_C_A_0_00001.pdb 
# Loaded target data/PDB/CUUCGG/4u4r-pdb-bundle2_1923_C_A_0_00002.pdb 
# Loading data/PDB/CUUCGG/4u4r-pdb-bundle2_1923_C_A_0_00002.pdb 
# Loaded reference data/PDB/CUUCGG/2KOC_5_C_A_0_00001.pdb 
# Loaded target data/PDB/CUUCGG/2LU0_28_C_A_0_00008.pdb 
# Loading data/PDB/CUUCGG/2LU0_28_C_A_0_00008.pdb 
# Loaded reference data/PDB/CUUCGG/2KOC_5_C_A_0_00001.pdb 
# Loaded target data/PDB/CUUCGG/4wr6-pdb-bundle4_1691_C_A_0_00000.pdb 
# Loading data/PDB/CUUCGG/4wr6-pdb-bundle4_1691_C_A_0_00000.pdb 
# Loaded reference data/PDB/CUUCGG/2KOC_5_C_A_0_00001.pdb 
# Loaded target data/PDB/CUUCGG/5nwy-pdb-bundle1_1691_C_N_0_00000.pdb 
# Loading data/PDB/CUUCGG/5nwy-pdb-bundle1_1691_C_N_0_00000.pdb 
# Loaded reference data/PDB/CUUCGG/2KOC_5_C_A_0_00001.pdb 
# Loaded target data/PDB/CUUCGG/4y4o-pdb-bundle1_1738_C_A_0_00000.pdb 
# Loading data/PDB/CUUCGG/4y4o-pdb-bundle1_1738_C_A_0_00000.pdb 
# Loaded reference data/PDB/CUUCGG/2KOC_5_C_A_0_00001.pdb 
# Loaded target data/PDB/CUUCGG/4v85-pdb-bundle2_1691_C_A_0_00000.pdb 
# Loading data/PDB/CUUCGG/4v85-pdb-bundle2_1691_C_A_0_00000.pdb 
# Loaded reference data/PDB/CUUCGG/2KOC_5_C_A_0_00001.pdb 
# Loaded target data/PDB/CUUCGG/5lzv-pdb-bundle2_2872_C_5_0_00000.pdb 
# Loading data/PDB/CUUCGG/5lzv-pdb-bundle2_2872_C_5_0_00000.pdb 
# Loaded reference data/PDB/CUUCGG/2KOC_5_C_A_0_00001.pdb 
# Loaded target data/PDB/CUUCGG/4woi-pdb-bundle3_1691_C_A_0_00000.pdb 
# Loading data/PDB/CUUCGG/4woi-pdb-bundle3_1691_C_A_0_00000.pdb 
# Loaded reference data/PDB/CUUCGG/2KOC_5_C_A_0_00001.pdb 
# Loaded target data/PDB/CUUCGG/1vy4-pdb-bundle4_1691_C_A_0_00000.pdb 
# Loading data/PDB/CUUCGG/1vy4-pdb-bundle4_1691_C_A_0_00000.pdb 
# Loaded reference data/PDB/CUUCGG/2KOC_5_C_A_0_00001.pdb 
# Loaded target data/PDB/CUUCGG/2ZJQ_1708_C_X_0_00000.pdb 
# Loading data/PDB/CUUCGG/2ZJQ_1708_C_X_0_00000.pdb 
# Loaded reference data/PDB/CUUCGG/2KOC_5_C_A_0_00001.pdb 
# Loaded target data/PDB/CUUCGG/4v9l-pdb-bundle2_1691_C_A_0_00000.pdb 
# Loading data/PDB/CUUCGG/4v9l-pdb-bundle2_1691_C_A_0_00000.pdb 
# Loaded reference data/PDB/CUUCGG/2KOC_5_C_A_0_00001.pdb 
# Loaded target data/PDB/CUUCGG/4v9q-pdb-bundle2_419_C_A_0_00000.pdb 
# Loading data/PDB/CUUCGG/4v9q-pdb-bundle2_419_C_A_0_00000.pdb 
# Loaded reference data/PDB/CUUCGG/2KOC_5_C_A_0_00001.pdb 
# Loaded target data/PDB/CUUCGG/4v6a-pdb-bundle3_419_C_A_0_00000.pdb 
# Loading data/PDB/CUUCGG/4v6a-pdb-bundle3_419_C_A_0_00000.pdb 
# Loaded reference data/PDB/CUUCGG/2KOC_5_C_A_0_00001.pdb 
# Loaded target data/PDB/CUUCGG/4v9o-pdb-bundle6_1028_C_A_0_00002.pdb 
# Loading data/PDB/CUUCGG/4v9o-pdb-bundle6_1028_C_A_0_00002.pdb 
# Loaded reference data/PDB/CUUCGG/2KOC_5_C_A_0_00001.pdb 
# Loaded target data/PDB/CUUCGG/2N3R_37_C_A_0_00014.pdb 
# Loading data/PDB/CUUCGG/2N3R_37_C_A_0_00014.pdb 
# Loaded reference data/PDB/CUUCGG/2KOC_5_C_A_0_00001.pdb 
# Loaded target data/PDB/CUUCGG/6cfj-pdb-bundle1_1738_C_A_0_00000.pdb 
# Loading data/PDB/CUUCGG/6cfj-pdb-bundle1_1738_C_A_0_00000.pdb 
# Loaded reference data/PDB/CUUCGG/2KOC_5_C_A_0_00001.pdb 
# Loaded target data/PDB/CUUCGG/2L3E_18_C_A_0_00013.pdb 
# Loading data/PDB/CUUCGG/2L3E_18_C_A_0_00013.pdb 
# Loaded reference data/PDB/CUUCGG/2KOC_5_C_A_0_00001.pdb 
# Loaded target data/PDB/CUUCGG/2L3E_18_C_A_0_00007.pdb 
# Loading data/PDB/CUUCGG/2L3E_18_C_A_0_00007.pdb 
# Loaded reference data/PDB/CUUCGG/2KOC_5_C_A_0_00001.pdb 
# Loaded target data/PDB/CUUCGG/2N3R_37_C_A_0_00000.pdb 
# Loading data/PDB/CUUCGG/2N3R_37_C_A_0_00000.pdb 
# Loaded reference data/PDB/CUUCGG/2KOC_5_C_A_0_00001.pdb 
# Loaded target data/PDB/CUUCGG/5afi-pdb-bundle2_1691_C_A_0_00000.pdb 
# Loading data/PDB/CUUCGG/5afi-pdb-bundle2_1691_C_A_0_00000.pdb 
# Loaded reference data/PDB/CUUCGG/2KOC_5_C_A_0_00001.pdb 
# Loaded target data/PDB/CUUCGG/1Q7Y_1769_C_A_0_00000.pdb 
# Loading data/PDB/CUUCGG/1Q7Y_1769_C_A_0_00000.pdb 
# Loaded reference data/PDB/CUUCGG/2KOC_5_C_A_0_00001.pdb 
# Loaded target data/PDB/CUUCGG/2KU0_19_C_A_0_00009.pdb 
# Loading data/PDB/CUUCGG/2KU0_19_C_A_0_00009.pdb 
# Loaded reference data/PDB/CUUCGG/2KOC_5_C_A_0_00001.pdb 
# Loaded target data/PDB/CUUCGG/4v9b-pdb-bundle4_1691_C_A_0_00000.pdb 
# Loading data/PDB/CUUCGG/4v9b-pdb-bundle4_1691_C_A_0_00000.pdb 
# Loaded reference data/PDB/CUUCGG/2KOC_5_C_A_0_00001.pdb 
# Loaded target data/PDB/CUUCGG/4u20-pdb-bundle1_1028_C_A_0_00002.pdb 
# Loading data/PDB/CUUCGG/4u20-pdb-bundle1_1028_C_A_0_00002.pdb 
# Loaded reference data/PDB/CUUCGG/2KOC_5_C_A_0_00001.pdb 
# Loaded target data/PDB/CUUCGG/1A3M_11_C_A_0_00013.pdb 
# Loading data/PDB/CUUCGG/1A3M_11_C_A_0_00013.pdb 
# Loaded reference data/PDB/CUUCGG/2KOC_5_C_A_0_00001.pdb 
# Loaded target data/PDB/CUUCGG/1A3M_11_C_A_0_00007.pdb 
# Loading data/PDB/CUUCGG/1A3M_11_C_A_0_00007.pdb 
# Loaded reference data/PDB/CUUCGG/2KOC_5_C_A_0_00001.pdb 
# Loaded target data/PDB/CUUCGG/1KUQ_8_C_B_0_00000.pdb 
# Loading data/PDB/CUUCGG/1KUQ_8_C_B_0_00000.pdb 
# Loaded reference data/PDB/CUUCGG/2KOC_5_C_A_0_00001.pdb 
# Loaded target data/PDB/CUUCGG/1YKV_220_C_D_0_00001.pdb 
# Loading data/PDB/CUUCGG/1YKV_220_C_D_0_00001.pdb 
# Loaded reference data/PDB/CUUCGG/2KOC_5_C_A_0_00001.pdb 
# Loaded target data/PDB/CUUCGG/4w2g-pdb-bundle4_1691_C_A_0_00000.pdb 
# Loading data/PDB/CUUCGG/4w2g-pdb-bundle4_1691_C_A_0_00000.pdb 
# Loaded reference data/PDB/CUUCGG/2KOC_5_C_A_0_00001.pdb 
# Loaded target data/PDB/CUUCGG/2GIO_13_C_A_0_00019.pdb 
# Loading data/PDB/CUUCGG/2GIO_13_C_A_0_00019.pdb 
# Loaded reference data/PDB/CUUCGG/2KOC_5_C_A_0_00001.pdb 
# Loaded target data/PDB/CUUCGG/4v9l-pdb-bundle1_419_C_T_0_00000.pdb 
# Loading data/PDB/CUUCGG/4v9l-pdb-bundle1_419_C_T_0_00000.pdb 
# Loaded reference data/PDB/CUUCGG/2KOC_5_C_A_0_00001.pdb 
# Loaded target data/PDB/CUUCGG/1BYJ_11_C_A_0_00020.pdb 
# Loading data/PDB/CUUCGG/1BYJ_11_C_A_0_00020.pdb 
# Loaded reference data/PDB/CUUCGG/2KOC_5_C_A_0_00001.pdb 
# Loaded target data/PDB/CUUCGG/2N3Q_37_C_A_0_00010.pdb 
# Loading data/PDB/CUUCGG/2N3Q_37_C_A_0_00010.pdb 
# Loaded reference data/PDB/CUUCGG/2KOC_5_C_A_0_00001.pdb 
# Loaded target data/PDB/CUUCGG/2M58_12_C_A_0_00004.pdb 
# Loading data/PDB/CUUCGG/2M58_12_C_A_0_00004.pdb 
# Loaded reference data/PDB/CUUCGG/2KOC_5_C_A_0_00001.pdb 
# Loaded target data/PDB/CUUCGG/3j7o-pdb-bundle1_2872_C_5_0_00000.pdb 
# Loading data/PDB/CUUCGG/3j7o-pdb-bundle1_2872_C_5_0_00000.pdb 
# Loaded reference data/PDB/CUUCGG/2KOC_5_C_A_0_00001.pdb 
# Loaded target data/PDB/CUUCGG/5u4i-pdb-bundle1_1691_C_A_0_00000.pdb 
# Loading data/PDB/CUUCGG/5u4i-pdb-bundle1_1691_C_A_0_00000.pdb 
# Loaded reference data/PDB/CUUCGG/2KOC_5_C_A_0_00001.pdb 
# Loaded target data/PDB/CUUCGG/2N3Q_37_C_A_0_00004.pdb 
# Loading data/PDB/CUUCGG/2N3Q_37_C_A_0_00004.pdb 
# Loaded reference data/PDB/CUUCGG/2KOC_5_C_A_0_00001.pdb 
# Loaded target data/PDB/CUUCGG/1BYJ_11_C_A_0_00034.pdb 
# Loading data/PDB/CUUCGG/1BYJ_11_C_A_0_00034.pdb 
# Loaded reference data/PDB/CUUCGG/2KOC_5_C_A_0_00001.pdb 
# Loaded target data/PDB/CUUCGG/2HUA_17_C_A_0_00010.pdb 
# Loading data/PDB/CUUCGG/2HUA_17_C_A_0_00010.pdb 
# Loaded reference data/PDB/CUUCGG/2KOC_5_C_A_0_00001.pdb 
# Loaded target data/PDB/CUUCGG/2LQZ_13_C_A_0_00005.pdb 
# Loading data/PDB/CUUCGG/2LQZ_13_C_A_0_00005.pdb 
# Loaded reference data/PDB/CUUCGG/2KOC_5_C_A_0_00001.pdb 
# Loaded target data/PDB/CUUCGG/1BYJ_11_C_A_0_00008.pdb 
# Loading data/PDB/CUUCGG/1BYJ_11_C_A_0_00008.pdb 
# Loaded reference data/PDB/CUUCGG/2KOC_5_C_A_0_00001.pdb 
# Loaded target data/PDB/CUUCGG/2HUA_17_C_A_0_00004.pdb 
# Loading data/PDB/CUUCGG/2HUA_17_C_A_0_00004.pdb 
# Loaded reference data/PDB/CUUCGG/2KOC_5_C_A_0_00001.pdb 
# Loaded target data/PDB/CUUCGG/1B36_16_C_A_0_00001.pdb 
# Loading data/PDB/CUUCGG/1B36_16_C_A_0_00001.pdb 
# Loaded reference data/PDB/CUUCGG/2KOC_5_C_A_0_00001.pdb 
# Loaded target data/PDB/CUUCGG/2LHP_16_C_A_0_00001.pdb 
# Loading data/PDB/CUUCGG/2LHP_16_C_A_0_00001.pdb 
# Loaded reference data/PDB/CUUCGG/2KOC_5_C_A_0_00001.pdb 
# Loaded target data/PDB/CUUCGG/1D6K_283_C_B_0_00010.pdb 
# Loading data/PDB/CUUCGG/1D6K_283_C_B_0_00010.pdb 
# Loaded reference data/PDB/CUUCGG/2KOC_5_C_A_0_00001.pdb 
# Loaded target data/PDB/CUUCGG/5mgp-pdb-bundle2_1449_C_a_0_00003.pdb 
# Loading data/PDB/CUUCGG/5mgp-pdb-bundle2_1449_C_a_0_00003.pdb 
# Loaded reference data/PDB/CUUCGG/2KOC_5_C_A_0_00001.pdb 
# Loaded target data/PDB/CUUCGG/1F6Z_12_C_A_0_00000.pdb 
# Loading data/PDB/CUUCGG/1F6Z_12_C_A_0_00000.pdb 
# Loaded reference data/PDB/CUUCGG/2KOC_5_C_A_0_00001.pdb 
# Loaded target data/PDB/CUUCGG/2NCI_14_C_A_0_00003.pdb 
# Loading data/PDB/CUUCGG/2NCI_14_C_A_0_00003.pdb 
# Loaded reference data/PDB/CUUCGG/2KOC_5_C_A_0_00001.pdb 
# Loaded target data/PDB/CUUCGG/1vvj-pdb-bundle3_419_C_A_0_00000.pdb 
# Loading data/PDB/CUUCGG/1vvj-pdb-bundle3_419_C_A_0_00000.pdb 
# Loaded reference data/PDB/CUUCGG/2KOC_5_C_A_0_00001.pdb 
# Loaded target data/PDB/CUUCGG/1D6K_283_C_B_0_00004.pdb 
# Loading data/PDB/CUUCGG/1D6K_283_C_B_0_00004.pdb 
# Loaded reference data/PDB/CUUCGG/2KOC_5_C_A_0_00001.pdb 
# Loaded target data/PDB/CUUCGG/1XSU_12_C_A_0_00000.pdb 
# Loading data/PDB/CUUCGG/1XSU_12_C_A_0_00000.pdb 
# Loaded reference data/PDB/CUUCGG/2KOC_5_C_A_0_00001.pdb 
# Loaded target data/PDB/CUUCGG/4v7s-pdb-bundle3_207_C_A_0_00000.pdb 
# Loading data/PDB/CUUCGG/4v7s-pdb-bundle3_207_C_A_0_00000.pdb 
# Loaded reference data/PDB/CUUCGG/2KOC_5_C_A_0_00001.pdb 
# Loaded target data/PDB/CUUCGG/4v8c-pdb-bundle2_1691_C_A_0_00000.pdb 
# Loading data/PDB/CUUCGG/4v8c-pdb-bundle2_1691_C_A_0_00000.pdb 
# Loaded reference data/PDB/CUUCGG/2KOC_5_C_A_0_00001.pdb 
# Loaded target data/PDB/CUUCGG/4v9d-pdb-bundle1_1449_C_A_0_00003.pdb 
# Loading data/PDB/CUUCGG/4v9d-pdb-bundle1_1449_C_A_0_00003.pdb 
# Loaded reference data/PDB/CUUCGG/2KOC_5_C_A_0_00001.pdb 
# Loaded target data/PDB/CUUCGG/5we6-pdb-bundle1_1691_C_A_0_00000.pdb 
# Loading data/PDB/CUUCGG/5we6-pdb-bundle1_1691_C_A_0_00000.pdb 
# Loaded reference data/PDB/CUUCGG/2KOC_5_C_A_0_00001.pdb 
# Loaded target data/PDB/CUUCGG/4v4h-pdb-bundle2_1691_C_A_0_00000.pdb 
# Loading data/PDB/CUUCGG/4v4h-pdb-bundle2_1691_C_A_0_00000.pdb 
# Loaded reference data/PDB/CUUCGG/2KOC_5_C_A_0_00001.pdb 
# Loaded target data/PDB/CUUCGG/2LWK_15_C_A_0_00012.pdb 
# Loading data/PDB/CUUCGG/2LWK_15_C_A_0_00012.pdb 
# Loaded reference data/PDB/CUUCGG/2KOC_5_C_A_0_00001.pdb 
# Loaded target data/PDB/CUUCGG/2LWK_15_C_A_0_00006.pdb 
# Loading data/PDB/CUUCGG/2LWK_15_C_A_0_00006.pdb 
# Loaded reference data/PDB/CUUCGG/2KOC_5_C_A_0_00001.pdb 
# Loaded target data/PDB/CUUCGG/4v51-pdb-bundle2_1691_C_A_0_00000.pdb 
# Loading data/PDB/CUUCGG/4v51-pdb-bundle2_1691_C_A_0_00000.pdb 
# Loaded reference data/PDB/CUUCGG/2KOC_5_C_A_0_00001.pdb 
# Loaded target data/PDB/CUUCGG/2N6S_16_C_A_0_00005.pdb 
# Loading data/PDB/CUUCGG/2N6S_16_C_A_0_00005.pdb 
# Loaded reference data/PDB/CUUCGG/2KOC_5_C_A_0_00001.pdb 
# Loaded target data/PDB/CUUCGG/4w4g-pdb-bundle2_1691_C_A_0_00000.pdb 
# Loading data/PDB/CUUCGG/4w4g-pdb-bundle2_1691_C_A_0_00000.pdb 
# Loaded reference data/PDB/CUUCGG/2KOC_5_C_A_0_00001.pdb 
# Loaded target data/PDB/CUUCGG/1KD1_1769_C_A_0_00000.pdb 
# Loading data/PDB/CUUCGG/1KD1_1769_C_A_0_00000.pdb 
# Loaded reference data/PDB/CUUCGG/2KOC_5_C_A_0_00001.pdb 
# Loaded target data/PDB/CUUCGG/1KC8_1769_C_A_0_00000.pdb 
# Loading data/PDB/CUUCGG/1KC8_1769_C_A_0_00000.pdb 
# Loaded reference data/PDB/CUUCGG/2KOC_5_C_A_0_00001.pdb 
# Loaded target data/PDB/CUUCGG/2N6X_20_C_A_0_00008.pdb 
# Loading data/PDB/CUUCGG/2N6X_20_C_A_0_00008.pdb 
# Loaded reference data/PDB/CUUCGG/2KOC_5_C_A_0_00001.pdb 
# Loaded target data/PDB/CUUCGG/4v9p-pdb-bundle3_1691_C_A_0_00000.pdb 
# Loading data/PDB/CUUCGG/4v9p-pdb-bundle3_1691_C_A_0_00000.pdb 
# Loaded reference data/PDB/CUUCGG/2KOC_5_C_A_0_00001.pdb 
# Loaded target data/PDB/CUUCGG/1FYO_11_C_A_0_00023.pdb 
# Loading data/PDB/CUUCGG/1FYO_11_C_A_0_00023.pdb 
# Loaded reference data/PDB/CUUCGG/2KOC_5_C_A_0_00001.pdb 
# Loaded target data/PDB/CUUCGG/1FYO_11_C_A_0_00022.pdb 
# Loading data/PDB/CUUCGG/1FYO_11_C_A_0_00022.pdb 
# Loaded reference data/PDB/CUUCGG/2KOC_5_C_A_0_00001.pdb 
# Loaded target data/PDB/CUUCGG/6ha8-pdb-bundle2_1038_C_a_0_00001.pdb 
# Loading data/PDB/CUUCGG/6ha8-pdb-bundle2_1038_C_a_0_00001.pdb 
# Loaded reference data/PDB/CUUCGG/2KOC_5_C_A_0_00001.pdb 
# Loaded target data/PDB/CUUCGG/2N6X_20_C_A_0_00009.pdb 
# Loading data/PDB/CUUCGG/2N6X_20_C_A_0_00009.pdb 
# Loaded reference data/PDB/CUUCGG/2KOC_5_C_A_0_00001.pdb 
# Loaded target data/PDB/CUUCGG/4u26-pdb-bundle2_1691_C_A_0_00000.pdb 
# Loading data/PDB/CUUCGG/4u26-pdb-bundle2_1691_C_A_0_00000.pdb 
# Loaded reference data/PDB/CUUCGG/2KOC_5_C_A_0_00001.pdb 
# Loaded target data/PDB/CUUCGG/4u53-pdb-bundle4_1923_C_I_0_00002.pdb 
# Loading data/PDB/CUUCGG/4u53-pdb-bundle4_1923_C_I_0_00002.pdb 
# Loaded reference data/PDB/CUUCGG/2KOC_5_C_A_0_00001.pdb 
# Loaded target data/PDB/CUUCGG/4z8c-pdb-bundle3_1737_C_A_0_00000.pdb 
# Loading data/PDB/CUUCGG/4z8c-pdb-bundle3_1737_C_A_0_00000.pdb 
# Loaded reference data/PDB/CUUCGG/2KOC_5_C_A_0_00001.pdb 
# Loaded target data/PDB/CUUCGG/2N6S_16_C_A_0_00004.pdb 
# Loading data/PDB/CUUCGG/2N6S_16_C_A_0_00004.pdb 
# Loaded reference data/PDB/CUUCGG/2KOC_5_C_A_0_00001.pdb 
# Loaded target data/PDB/CUUCGG/5el5-pdb-bundle4_1691_C_A_0_00000.pdb 
# Loading data/PDB/CUUCGG/5el5-pdb-bundle4_1691_C_A_0_00000.pdb 
# Loaded reference data/PDB/CUUCGG/2KOC_5_C_A_0_00001.pdb 
# Loaded target data/PDB/CUUCGG/2LWK_15_C_A_0_00007.pdb 
# Loading data/PDB/CUUCGG/2LWK_15_C_A_0_00007.pdb 
# Loaded reference data/PDB/CUUCGG/2KOC_5_C_A_0_00001.pdb 
# Loaded target data/PDB/CUUCGG/2LWK_15_C_A_0_00013.pdb 
# Loading data/PDB/CUUCGG/2LWK_15_C_A_0_00013.pdb 
# Loaded reference data/PDB/CUUCGG/2KOC_5_C_A_0_00001.pdb 
# Loaded target data/PDB/CUUCGG/5ndv-pdb-bundle1_1923_C_A_0_00002.pdb 
# Loading data/PDB/CUUCGG/5ndv-pdb-bundle1_1923_C_A_0_00002.pdb 
# Loaded reference data/PDB/CUUCGG/2KOC_5_C_A_0_00001.pdb 
# Loaded target data/PDB/CUUCGG/4v7v-pdb-bundle4_1691_C_A_0_00000.pdb 
# Loading data/PDB/CUUCGG/4v7v-pdb-bundle4_1691_C_A_0_00000.pdb 
# Loaded reference data/PDB/CUUCGG/2KOC_5_C_A_0_00001.pdb 
# Loaded target data/PDB/CUUCGG/4v4q-pdb-bundle4_1691_C_A_0_00000.pdb 
# Loading data/PDB/CUUCGG/4v4q-pdb-bundle4_1691_C_A_0_00000.pdb 
# Loaded reference data/PDB/CUUCGG/2KOC_5_C_A_0_00001.pdb 
# Loaded target data/PDB/CUUCGG/1D6K_283_C_B_0_00005.pdb 
# Loading data/PDB/CUUCGG/1D6K_283_C_B_0_00005.pdb 
# Loaded reference data/PDB/CUUCGG/2KOC_5_C_A_0_00001.pdb 
# Loaded target data/PDB/CUUCGG/4l47-pdb-bundle4_1691_C_A_0_00000.pdb 
# Loading data/PDB/CUUCGG/4l47-pdb-bundle4_1691_C_A_0_00000.pdb 
# Loaded reference data/PDB/CUUCGG/2KOC_5_C_A_0_00001.pdb 
# Loaded target data/PDB/CUUCGG/1D6K_283_C_B_0_00011.pdb 
# Loading data/PDB/CUUCGG/1D6K_283_C_B_0_00011.pdb 
# Loaded reference data/PDB/CUUCGG/2KOC_5_C_A_0_00001.pdb 
# Loaded target data/PDB/CUUCGG/4v6f-pdb-bundle3_1691_C_S_0_00000.pdb 
# Loading data/PDB/CUUCGG/4v6f-pdb-bundle3_1691_C_S_0_00000.pdb 
# Loaded reference data/PDB/CUUCGG/2KOC_5_C_A_0_00001.pdb 
# Loaded target data/PDB/CUUCGG/2NCI_14_C_A_0_00002.pdb 
# Loading data/PDB/CUUCGG/2NCI_14_C_A_0_00002.pdb 
# Loaded reference data/PDB/CUUCGG/2KOC_5_C_A_0_00001.pdb 
# Loaded target data/PDB/CUUCGG/4wt8-pdb-bundle1_415_C_V_0_00000.pdb 
# Loading data/PDB/CUUCGG/4wt8-pdb-bundle1_415_C_V_0_00000.pdb 
# Loaded reference data/PDB/CUUCGG/2KOC_5_C_A_0_00001.pdb 
# Loaded target data/PDB/CUUCGG/4u1v-pdb-bundle1_1028_C_A_0_00002.pdb 
# Loading data/PDB/CUUCGG/4u1v-pdb-bundle1_1028_C_A_0_00002.pdb 
# Loaded reference data/PDB/CUUCGG/2KOC_5_C_A_0_00001.pdb 
# Loaded target data/PDB/CUUCGG/2LHP_16_C_A_0_00000.pdb 
# Loading data/PDB/CUUCGG/2LHP_16_C_A_0_00000.pdb 
# Loaded reference data/PDB/CUUCGG/2KOC_5_C_A_0_00001.pdb 
# Loaded target data/PDB/CUUCGG/1B36_16_C_A_0_00000.pdb 
# Loading data/PDB/CUUCGG/1B36_16_C_A_0_00000.pdb 
# Loaded reference data/PDB/CUUCGG/2KOC_5_C_A_0_00001.pdb 
# Loaded target data/PDB/CUUCGG/2HUA_17_C_A_0_00005.pdb 
# Loading data/PDB/CUUCGG/2HUA_17_C_A_0_00005.pdb 
# Loaded reference data/PDB/CUUCGG/2KOC_5_C_A_0_00001.pdb 
# Loaded target data/PDB/CUUCGG/1BYJ_11_C_A_0_00009.pdb 
# Loading data/PDB/CUUCGG/1BYJ_11_C_A_0_00009.pdb 
# Loaded reference data/PDB/CUUCGG/2KOC_5_C_A_0_00001.pdb 
# Loaded target data/PDB/CUUCGG/2LQZ_13_C_A_0_00004.pdb 
# Loading data/PDB/CUUCGG/2LQZ_13_C_A_0_00004.pdb 
# Loaded reference data/PDB/CUUCGG/2KOC_5_C_A_0_00001.pdb 
# Loaded target data/PDB/CUUCGG/4v7x-pdb-bundle2_1691_C_A_0_00000.pdb 
# Loading data/PDB/CUUCGG/4v7x-pdb-bundle2_1691_C_A_0_00000.pdb 
# Loaded reference data/PDB/CUUCGG/2KOC_5_C_A_0_00001.pdb 
# Loaded target data/PDB/CUUCGG/1BYJ_11_C_A_0_00035.pdb 
# Loading data/PDB/CUUCGG/1BYJ_11_C_A_0_00035.pdb 
# Loaded reference data/PDB/CUUCGG/2KOC_5_C_A_0_00001.pdb 
# Loaded target data/PDB/CUUCGG/2N3Q_37_C_A_0_00005.pdb 
# Loading data/PDB/CUUCGG/2N3Q_37_C_A_0_00005.pdb 
# Loaded reference data/PDB/CUUCGG/2KOC_5_C_A_0_00001.pdb 
# Loaded target data/PDB/CUUCGG/2N3Q_37_C_A_0_00011.pdb 
# Loading data/PDB/CUUCGG/2N3Q_37_C_A_0_00011.pdb 
# Loaded reference data/PDB/CUUCGG/2KOC_5_C_A_0_00001.pdb 
# Loaded target data/PDB/CUUCGG/1BYJ_11_C_A_0_00021.pdb 
# Loading data/PDB/CUUCGG/1BYJ_11_C_A_0_00021.pdb 
# Loaded reference data/PDB/CUUCGG/2KOC_5_C_A_0_00001.pdb 
# Loaded target data/PDB/CUUCGG/5doy-pdb-bundle3_1691_C_A_0_00000.pdb 
# Loading data/PDB/CUUCGG/5doy-pdb-bundle3_1691_C_A_0_00000.pdb 
# Loaded reference data/PDB/CUUCGG/2KOC_5_C_A_0_00001.pdb 
# Loaded target data/PDB/CUUCGG/2M58_12_C_A_0_00005.pdb 
# Loading data/PDB/CUUCGG/2M58_12_C_A_0_00005.pdb 
# Loaded reference data/PDB/CUUCGG/2KOC_5_C_A_0_00001.pdb 
# Loaded target data/PDB/CUUCGG/4AY2_8_C_C_0_00000.pdb 
# Loading data/PDB/CUUCGG/4AY2_8_C_C_0_00000.pdb 
# Loaded reference data/PDB/CUUCGG/2KOC_5_C_A_0_00001.pdb 
# Loaded target data/PDB/CUUCGG/1JO7_14_C_A_0_00012.pdb 
# Loading data/PDB/CUUCGG/1JO7_14_C_A_0_00012.pdb 
# Loaded reference data/PDB/CUUCGG/2KOC_5_C_A_0_00001.pdb 
# Loaded target data/PDB/CUUCGG/2GIO_13_C_A_0_00018.pdb 
# Loading data/PDB/CUUCGG/2GIO_13_C_A_0_00018.pdb 
# Loaded reference data/PDB/CUUCGG/2KOC_5_C_A_0_00001.pdb 
# Loaded target data/PDB/CUUCGG/4v57-pdb-bundle1_1028_C_A_0_00002.pdb 
# Loading data/PDB/CUUCGG/4v57-pdb-bundle1_1028_C_A_0_00002.pdb 
# Loaded reference data/PDB/CUUCGG/2KOC_5_C_A_0_00001.pdb 
# Loaded target data/PDB/CUUCGG/4v52-pdb-bundle3_1028_C_A_0_00002.pdb 
# Loading data/PDB/CUUCGG/4v52-pdb-bundle3_1028_C_A_0_00002.pdb 
# Loaded reference data/PDB/CUUCGG/2KOC_5_C_A_0_00001.pdb 
# Loaded target data/PDB/CUUCGG/1A3M_11_C_A_0_00006.pdb 
# Loading data/PDB/CUUCGG/1A3M_11_C_A_0_00006.pdb 
# Loaded reference data/PDB/CUUCGG/2KOC_5_C_A_0_00001.pdb 
# Loaded target data/PDB/CUUCGG/1A3M_11_C_A_0_00012.pdb 
# Loading data/PDB/CUUCGG/1A3M_11_C_A_0_00012.pdb 
# Loaded reference data/PDB/CUUCGG/2KOC_5_C_A_0_00001.pdb 
# Loaded target data/PDB/CUUCGG/4z3s-pdb-bundle2_419_C_A_0_00000.pdb 
# Loading data/PDB/CUUCGG/4z3s-pdb-bundle2_419_C_A_0_00000.pdb 
# Loaded reference data/PDB/CUUCGG/2KOC_5_C_A_0_00001.pdb 
# Loaded target data/PDB/CUUCGG/5mdw-pdb-bundle1_1449_C_2_0_00004.pdb 
# Loading data/PDB/CUUCGG/5mdw-pdb-bundle1_1449_C_2_0_00004.pdb 
# Loaded reference data/PDB/CUUCGG/2KOC_5_C_A_0_00001.pdb 
# Loaded target data/PDB/CUUCGG/6boh-pdb-bundle1_1691_C_B_0_00001.pdb 
# Loading data/PDB/CUUCGG/6boh-pdb-bundle1_1691_C_B_0_00001.pdb 
# Loaded reference data/PDB/CUUCGG/2KOC_5_C_A_0_00001.pdb 
# Loaded target data/PDB/CUUCGG/2KU0_19_C_A_0_00008.pdb 
# Loading data/PDB/CUUCGG/2KU0_19_C_A_0_00008.pdb 
# Loaded reference data/PDB/CUUCGG/2KOC_5_C_A_0_00001.pdb 
# Loaded target data/PDB/CUUCGG/5j5b-pdb-bundle3_1691_C_d_0_00000.pdb 
# Loading data/PDB/CUUCGG/5j5b-pdb-bundle3_1691_C_d_0_00000.pdb 
# Loaded reference data/PDB/CUUCGG/2KOC_5_C_A_0_00001.pdb 
# Loaded target data/PDB/CUUCGG/5D5L_60_C_D_0_00003.pdb 
# Loading data/PDB/CUUCGG/5D5L_60_C_D_0_00003.pdb 
# Loaded reference data/PDB/CUUCGG/2KOC_5_C_A_0_00001.pdb 
# Loaded target data/PDB/CUUCGG/5f8k-pdb-bundle2_419_C_A_0_00000.pdb 
# Loading data/PDB/CUUCGG/5f8k-pdb-bundle2_419_C_A_0_00000.pdb 
# Loaded reference data/PDB/CUUCGG/2KOC_5_C_A_0_00001.pdb 
# Loaded target data/PDB/CUUCGG/3CCQ_1769_C_0_0_00000.pdb 
# Loading data/PDB/CUUCGG/3CCQ_1769_C_0_0_00000.pdb 
# Loaded reference data/PDB/CUUCGG/2KOC_5_C_A_0_00001.pdb 
# Loaded target data/PDB/CUUCGG/4v5p-pdb-bundle2_1691_C_A_0_00000.pdb 
# Loading data/PDB/CUUCGG/4v5p-pdb-bundle2_1691_C_A_0_00000.pdb 
# Loaded reference data/PDB/CUUCGG/2KOC_5_C_A_0_00001.pdb 
# Loaded target data/PDB/CUUCGG/5mdz-pdb-bundle1_1691_C_1_0_00000.pdb 
# Loading data/PDB/CUUCGG/5mdz-pdb-bundle1_1691_C_1_0_00000.pdb 
# Loaded reference data/PDB/CUUCGG/2KOC_5_C_A_0_00001.pdb 
# Loaded target data/PDB/CUUCGG/2N3R_37_C_A_0_00001.pdb 
# Loading data/PDB/CUUCGG/2N3R_37_C_A_0_00001.pdb 
# Loaded reference data/PDB/CUUCGG/2KOC_5_C_A_0_00001.pdb 
# Loaded target data/PDB/CUUCGG/2L3E_18_C_A_0_00006.pdb 
# Loading data/PDB/CUUCGG/2L3E_18_C_A_0_00006.pdb 
# Loaded reference data/PDB/CUUCGG/2KOC_5_C_A_0_00001.pdb 
# Loaded target data/PDB/CUUCGG/4wqu-pdb-bundle2_419_C_A_0_00000.pdb 
# Loading data/PDB/CUUCGG/4wqu-pdb-bundle2_419_C_A_0_00000.pdb 
# Loaded reference data/PDB/CUUCGG/2KOC_5_C_A_0_00001.pdb 
# Loaded target data/PDB/CUUCGG/2L3E_18_C_A_0_00012.pdb 
# Loading data/PDB/CUUCGG/2L3E_18_C_A_0_00012.pdb 
# Loaded reference data/PDB/CUUCGG/2KOC_5_C_A_0_00001.pdb 
# Loaded target data/PDB/CUUCGG/2N3R_37_C_A_0_00015.pdb 
# Loading data/PDB/CUUCGG/2N3R_37_C_A_0_00015.pdb 
# Loaded reference data/PDB/CUUCGG/2KOC_5_C_A_0_00001.pdb 
# Loaded target data/PDB/CUUCGG/4ypb-pdb-bundle4_1691_C_A_0_00000.pdb 
# Loading data/PDB/CUUCGG/4ypb-pdb-bundle4_1691_C_A_0_00000.pdb 
# Loaded reference data/PDB/CUUCGG/2KOC_5_C_A_0_00001.pdb 
# Loaded target data/PDB/CUUCGG/1M90_1769_C_A_0_00000.pdb 
# Loading data/PDB/CUUCGG/1M90_1769_C_A_0_00000.pdb 
# Loaded reference data/PDB/CUUCGG/2KOC_5_C_A_0_00001.pdb 
# Loaded target data/PDB/CUUCGG/5wit-pdb-bundle1_1738_C_A_0_00000.pdb 
# Loading data/PDB/CUUCGG/5wit-pdb-bundle1_1738_C_A_0_00000.pdb 
# Loaded reference data/PDB/CUUCGG/2KOC_5_C_A_0_00001.pdb 
# Loaded target data/PDB/CUUCGG/5ngm-pdb-bundle2_1735_C_A_0_00000.pdb 
# Loading data/PDB/CUUCGG/5ngm-pdb-bundle2_1735_C_A_0_00000.pdb 
# Loaded reference data/PDB/CUUCGG/2KOC_5_C_A_0_00001.pdb 
# Loaded target data/PDB/CUUCGG/4wqf-pdb-bundle3_1691_C_A_0_00000.pdb 
# Loading data/PDB/CUUCGG/4wqf-pdb-bundle3_1691_C_A_0_00000.pdb 
# Loaded reference data/PDB/CUUCGG/2KOC_5_C_A_0_00001.pdb 
# Loaded target data/PDB/CUUCGG/5lyb-pdb-bundle4_1923_C_J_0_00002.pdb 
# Loading data/PDB/CUUCGG/5lyb-pdb-bundle4_1923_C_J_0_00002.pdb 
# Loaded reference data/PDB/CUUCGG/2KOC_5_C_A_0_00001.pdb 
# Loaded target data/PDB/CUUCGG/4v9s-pdb-bundle2_1738_C_A_0_00000.pdb 
# Loading data/PDB/CUUCGG/4v9s-pdb-bundle2_1738_C_A_0_00000.pdb 
# Loaded reference data/PDB/CUUCGG/2KOC_5_C_A_0_00001.pdb 
# Loaded target data/PDB/CUUCGG/3V7E_65_C_C_0_00000.pdb 
# Loading data/PDB/CUUCGG/3V7E_65_C_C_0_00000.pdb 
# Loaded reference data/PDB/CUUCGG/2KOC_5_C_A_0_00001.pdb 
# Loaded target data/PDB/CUUCGG/4u26-pdb-bundle1_419_C_A_0_00001.pdb 
# Loading data/PDB/CUUCGG/4u26-pdb-bundle1_419_C_A_0_00001.pdb 
# Loaded reference data/PDB/CUUCGG/2KOC_5_C_A_0_00001.pdb 
# Loaded target data/PDB/CUUCGG/4v9b-pdb-bundle2_1691_C_A_0_00000.pdb 
# Loading data/PDB/CUUCGG/4v9b-pdb-bundle2_1691_C_A_0_00000.pdb 
# Loaded reference data/PDB/CUUCGG/2KOC_5_C_A_0_00001.pdb 
# Loaded target data/PDB/CUUCGG/4w4g-pdb-bundle1_419_C_A_0_00000.pdb 
# Loading data/PDB/CUUCGG/4w4g-pdb-bundle1_419_C_A_0_00000.pdb 
# Loaded reference data/PDB/CUUCGG/2KOC_5_C_A_0_00001.pdb 
# Loaded target data/PDB/CUUCGG/2LU0_28_C_A_0_00012.pdb 
# Loading data/PDB/CUUCGG/2LU0_28_C_A_0_00012.pdb 
# Loaded reference data/PDB/CUUCGG/2KOC_5_C_A_0_00001.pdb 
# Loaded target data/PDB/CUUCGG/2LU0_28_C_A_0_00006.pdb 
# Loading data/PDB/CUUCGG/2LU0_28_C_A_0_00006.pdb 
# Loaded reference data/PDB/CUUCGG/2KOC_5_C_A_0_00001.pdb 
# Loaded target data/PDB/CUUCGG/6BY5_5_C_A_0_00001.pdb 
# Loading data/PDB/CUUCGG/6BY5_5_C_A_0_00001.pdb 
# Loaded reference data/PDB/CUUCGG/2KOC_5_C_A_0_00001.pdb 
# Loaded target data/PDB/CUUCGG/4v7u-pdb-bundle3_1028_C_A_0_00002.pdb 
# Loading data/PDB/CUUCGG/4v7u-pdb-bundle3_1028_C_A_0_00002.pdb 
# Loaded reference data/PDB/CUUCGG/2KOC_5_C_A_0_00001.pdb 
# Loaded target data/PDB/CUUCGG/4v8a-pdb-bundle3_419_C_A_0_00000.pdb 
# Loading data/PDB/CUUCGG/4v8a-pdb-bundle3_419_C_A_0_00000.pdb 
# Loaded reference data/PDB/CUUCGG/2KOC_5_C_A_0_00001.pdb 
# Loaded target data/PDB/CUUCGG/3jce-pdb-bundle1_207_C_a_0_00000.pdb 
# Loading data/PDB/CUUCGG/3jce-pdb-bundle1_207_C_a_0_00000.pdb 
# Loaded reference data/PDB/CUUCGG/2KOC_5_C_A_0_00001.pdb 
# Loaded target data/PDB/CUUCGG/2N3R_37_C_A_0_00018.pdb 
# Loading data/PDB/CUUCGG/2N3R_37_C_A_0_00018.pdb 
# Loaded reference data/PDB/CUUCGG/2KOC_5_C_A_0_00001.pdb 
# Loaded target data/PDB/CUUCGG/4v5p-pdb-bundle3_419_C_A_0_00000.pdb 
# Loading data/PDB/CUUCGG/4v5p-pdb-bundle3_419_C_A_0_00000.pdb 
# Loaded reference data/PDB/CUUCGG/2KOC_5_C_A_0_00001.pdb 
# Loaded target data/PDB/CUUCGG/1vy4-pdb-bundle2_1691_C_A_0_00000.pdb 
# Loading data/PDB/CUUCGG/1vy4-pdb-bundle2_1691_C_A_0_00000.pdb 
# Loaded reference data/PDB/CUUCGG/2KOC_5_C_A_0_00001.pdb 
# Loaded target data/PDB/CUUCGG/2GIP_12_C_A_0_00012.pdb 
# Loading data/PDB/CUUCGG/2GIP_12_C_A_0_00012.pdb 
# Loaded reference data/PDB/CUUCGG/2KOC_5_C_A_0_00001.pdb 
# Loaded target data/PDB/CUUCGG/2GIP_12_C_A_0_00006.pdb 
# Loading data/PDB/CUUCGG/2GIP_12_C_A_0_00006.pdb 
# Loaded reference data/PDB/CUUCGG/2KOC_5_C_A_0_00001.pdb 
# Loaded target data/PDB/CUUCGG/4v9l-pdb-bundle4_1691_C_A_0_00000.pdb 
# Loading data/PDB/CUUCGG/4v9l-pdb-bundle4_1691_C_A_0_00000.pdb 
# Loaded reference data/PDB/CUUCGG/2KOC_5_C_A_0_00001.pdb 
# Loaded target data/PDB/CUUCGG/5j91-pdb-bundle1_419_C_b_0_00005.pdb 
# Loading data/PDB/CUUCGG/5j91-pdb-bundle1_419_C_b_0_00005.pdb 
# Loaded reference data/PDB/CUUCGG/2KOC_5_C_A_0_00001.pdb 
# Loaded target data/PDB/CUUCGG/6cae-pdb-bundle1_1738_C_A_0_00000.pdb 
# Loading data/PDB/CUUCGG/6cae-pdb-bundle1_1738_C_A_0_00000.pdb 
# Loaded reference data/PDB/CUUCGG/2KOC_5_C_A_0_00001.pdb 
# Loaded target data/PDB/CUUCGG/2KU0_19_C_A_0_00005.pdb 
# Loading data/PDB/CUUCGG/2KU0_19_C_A_0_00005.pdb 
# Loaded reference data/PDB/CUUCGG/2KOC_5_C_A_0_00001.pdb 
# Loaded target data/PDB/CUUCGG/5D5L_60_C_C_0_00002.pdb 
# Loading data/PDB/CUUCGG/5D5L_60_C_C_0_00002.pdb 
# Loaded reference data/PDB/CUUCGG/2KOC_5_C_A_0_00001.pdb 
# Loaded target data/PDB/CUUCGG/4w2f-pdb-bundle3_419_C_A_0_00000.pdb 
# Loading data/PDB/CUUCGG/4w2f-pdb-bundle3_419_C_A_0_00000.pdb 
# Loaded reference data/PDB/CUUCGG/2KOC_5_C_A_0_00001.pdb 
# Loaded target data/PDB/CUUCGG/5czp-pdb-bundle1_1691_C_A_0_00000.pdb 
# Loading data/PDB/CUUCGG/5czp-pdb-bundle1_1691_C_A_0_00000.pdb 
# Loaded reference data/PDB/CUUCGG/2KOC_5_C_A_0_00001.pdb 
# Loaded target data/PDB/CUUCGG/4v57-pdb-bundle1_1449_C_A_0_00003.pdb 
# Loading data/PDB/CUUCGG/4v57-pdb-bundle1_1449_C_A_0_00003.pdb 
# Loaded reference data/PDB/CUUCGG/2KOC_5_C_A_0_00001.pdb 
# Loaded target data/PDB/CUUCGG/4w2i-pdb-bundle4_1691_C_A_0_00000.pdb 
# Loading data/PDB/CUUCGG/4w2i-pdb-bundle4_1691_C_A_0_00000.pdb 
# Loaded reference data/PDB/CUUCGG/2KOC_5_C_A_0_00001.pdb 
# Loaded target data/PDB/CUUCGG/4wt8-pdb-bundle3_1737_C_k_0_00000.pdb 
# Loading data/PDB/CUUCGG/4wt8-pdb-bundle3_1737_C_k_0_00000.pdb 
# Loaded reference data/PDB/CUUCGG/2KOC_5_C_A_0_00001.pdb 
# Loaded target data/PDB/CUUCGG/4v8d-pdb-bundle1_419_C_A_0_00000.pdb 
# Loading data/PDB/CUUCGG/4v8d-pdb-bundle1_419_C_A_0_00000.pdb 
# Loaded reference data/PDB/CUUCGG/2KOC_5_C_A_0_00001.pdb 
# Loaded target data/PDB/CUUCGG/4v52-pdb-bundle3_1449_C_A_0_00003.pdb 
# Loading data/PDB/CUUCGG/4v52-pdb-bundle3_1449_C_A_0_00003.pdb 
# Loaded reference data/PDB/CUUCGG/2KOC_5_C_A_0_00001.pdb 
# Loaded target data/PDB/CUUCGG/4JI1_419_C_A_0_00000.pdb 
# Loading data/PDB/CUUCGG/4JI1_419_C_A_0_00000.pdb 
# Loaded reference data/PDB/CUUCGG/2KOC_5_C_A_0_00001.pdb 
# Loaded target data/PDB/CUUCGG/2GIO_13_C_A_0_00001.pdb 
# Loading data/PDB/CUUCGG/2GIO_13_C_A_0_00001.pdb 
# Loaded reference data/PDB/CUUCGG/2KOC_5_C_A_0_00001.pdb 
# Loaded target data/PDB/CUUCGG/5KH8_26_C_A_0_00006.pdb 
# Loading data/PDB/CUUCGG/5KH8_26_C_A_0_00006.pdb 
# Loaded reference data/PDB/CUUCGG/2KOC_5_C_A_0_00001.pdb 
# Loaded target data/PDB/CUUCGG/1MFY_14_C_A_0_00002.pdb 
# Loading data/PDB/CUUCGG/1MFY_14_C_A_0_00002.pdb 
# Loaded reference data/PDB/CUUCGG/2KOC_5_C_A_0_00001.pdb 
# Loaded target data/PDB/CUUCGG/5KH8_26_C_A_0_00012.pdb 
# Loading data/PDB/CUUCGG/5KH8_26_C_A_0_00012.pdb 
# Loaded reference data/PDB/CUUCGG/2KOC_5_C_A_0_00001.pdb 
# Loaded target data/PDB/CUUCGG/2GIO_13_C_A_0_00015.pdb 
# Loading data/PDB/CUUCGG/2GIO_13_C_A_0_00015.pdb 
# Loaded reference data/PDB/CUUCGG/2KOC_5_C_A_0_00001.pdb 
# Loaded target data/PDB/CUUCGG/2N3Q_37_C_A_0_00008.pdb 
# Loading data/PDB/CUUCGG/2N3Q_37_C_A_0_00008.pdb 
# Loaded reference data/PDB/CUUCGG/2KOC_5_C_A_0_00001.pdb 
# Loaded target data/PDB/CUUCGG/4v51-pdb-bundle4_1691_C_A_0_00000.pdb 
# Loading data/PDB/CUUCGG/4v51-pdb-bundle4_1691_C_A_0_00000.pdb 
# Loaded reference data/PDB/CUUCGG/2KOC_5_C_A_0_00001.pdb 
# Loaded target data/PDB/CUUCGG/2M58_12_C_A_0_00008.pdb 
# Loading data/PDB/CUUCGG/2M58_12_C_A_0_00008.pdb 
# Loaded reference data/PDB/CUUCGG/2KOC_5_C_A_0_00001.pdb 
# Loaded target data/PDB/CUUCGG/2HUA_17_C_A_0_00008.pdb 
# Loading data/PDB/CUUCGG/2HUA_17_C_A_0_00008.pdb 
# Loaded reference data/PDB/CUUCGG/2KOC_5_C_A_0_00001.pdb 
# Loaded target data/PDB/CUUCGG/1BYJ_11_C_A_0_00004.pdb 
# Loading data/PDB/CUUCGG/1BYJ_11_C_A_0_00004.pdb 
# Loaded reference data/PDB/CUUCGG/2KOC_5_C_A_0_00001.pdb 
# Loaded target data/PDB/CUUCGG/1BYJ_11_C_A_0_00010.pdb 
# Loading data/PDB/CUUCGG/1BYJ_11_C_A_0_00010.pdb 
# Loaded reference data/PDB/CUUCGG/2KOC_5_C_A_0_00001.pdb 
# Loaded target data/PDB/CUUCGG/2N3Q_37_C_A_0_00020.pdb 
# Loading data/PDB/CUUCGG/2N3Q_37_C_A_0_00020.pdb 
# Loaded reference data/PDB/CUUCGG/2KOC_5_C_A_0_00001.pdb 
# Loaded target data/PDB/CUUCGG/4v4h-pdb-bundle4_1691_C_A_0_00000.pdb 
# Loading data/PDB/CUUCGG/4v4h-pdb-bundle4_1691_C_A_0_00000.pdb 
# Loaded reference data/PDB/CUUCGG/2KOC_5_C_A_0_00001.pdb 
# Loaded target data/PDB/CUUCGG/1D6K_283_C_B_0_00008.pdb 
# Loading data/PDB/CUUCGG/1D6K_283_C_B_0_00008.pdb 
# Loaded reference data/PDB/CUUCGG/2KOC_5_C_A_0_00001.pdb 
# Loaded target data/PDB/CUUCGG/4u1v-pdb-bundle1_1449_C_A_0_00003.pdb 
# Loading data/PDB/CUUCGG/4u1v-pdb-bundle1_1449_C_A_0_00003.pdb 
# Loaded reference data/PDB/CUUCGG/2KOC_5_C_A_0_00001.pdb 
# Loaded target data/PDB/CUUCGG/4v9p-pdb-bundle5_1691_C_A_0_00000.pdb 
# Loading data/PDB/CUUCGG/4v9p-pdb-bundle5_1691_C_A_0_00000.pdb 
# Loaded reference data/PDB/CUUCGG/2KOC_5_C_A_0_00001.pdb 
# Loaded target data/PDB/CUUCGG/5jc9-pdb-bundle1_419_C_b_0_00005.pdb 
# Loading data/PDB/CUUCGG/5jc9-pdb-bundle1_419_C_b_0_00005.pdb 
# Loaded reference data/PDB/CUUCGG/2KOC_5_C_A_0_00001.pdb 
# Loaded target data/PDB/CUUCGG/4w4g-pdb-bundle4_1691_C_A_0_00000.pdb 
# Loading data/PDB/CUUCGG/4w4g-pdb-bundle4_1691_C_A_0_00000.pdb 
# Loaded reference data/PDB/CUUCGG/2KOC_5_C_A_0_00001.pdb 
# Loaded target data/PDB/CUUCGG/1JJ2_1769_C_0_0_00000.pdb 
# Loading data/PDB/CUUCGG/1JJ2_1769_C_0_0_00000.pdb 
# Loaded reference data/PDB/CUUCGG/2KOC_5_C_A_0_00001.pdb 
# Loaded target data/PDB/CUUCGG/4v7j-pdb-bundle3_419_C_A_0_00000.pdb 
# Loading data/PDB/CUUCGG/4v7j-pdb-bundle3_419_C_A_0_00000.pdb 
# Loaded reference data/PDB/CUUCGG/2KOC_5_C_A_0_00001.pdb 
# Loaded target data/PDB/CUUCGG/2QEX_1769_C_0_0_00000.pdb 
# Loading data/PDB/CUUCGG/2QEX_1769_C_0_0_00000.pdb 
# Loaded reference data/PDB/CUUCGG/2KOC_5_C_A_0_00001.pdb 
# Loaded target data/PDB/CUUCGG/1YHQ_1769_C_0_0_00000.pdb 
# Loading data/PDB/CUUCGG/1YHQ_1769_C_0_0_00000.pdb 
# Loaded reference data/PDB/CUUCGG/2KOC_5_C_A_0_00001.pdb 
# Loaded target data/PDB/CUUCGG/1TLR_10_C_A_0_00007.pdb 
# Loading data/PDB/CUUCGG/1TLR_10_C_A_0_00007.pdb 
# Loaded reference data/PDB/CUUCGG/2KOC_5_C_A_0_00001.pdb 
# Loaded target data/PDB/CUUCGG/5on6-pdb-bundle3_1923_C_A_0_00002.pdb 
# Loading data/PDB/CUUCGG/5on6-pdb-bundle3_1923_C_A_0_00002.pdb 
# Loaded reference data/PDB/CUUCGG/2KOC_5_C_A_0_00001.pdb 
# Loaded target data/PDB/CUUCGG/5tga-pdb-bundle2_1923_C_A_0_00002.pdb 
# Loading data/PDB/CUUCGG/5tga-pdb-bundle2_1923_C_A_0_00002.pdb 
# Loaded reference data/PDB/CUUCGG/2KOC_5_C_A_0_00001.pdb 
# Loaded target data/PDB/CUUCGG/1K73_1769_C_A_0_00000.pdb 
# Loading data/PDB/CUUCGG/1K73_1769_C_A_0_00000.pdb 
# Loaded reference data/PDB/CUUCGG/2KOC_5_C_A_0_00001.pdb 
# Loaded target data/PDB/CUUCGG/1TLR_10_C_A_0_00013.pdb 
# Loading data/PDB/CUUCGG/1TLR_10_C_A_0_00013.pdb 
# Loaded reference data/PDB/CUUCGG/2KOC_5_C_A_0_00001.pdb 
# Loaded target data/PDB/CUUCGG/1FYO_11_C_A_0_00013.pdb 
# Loading data/PDB/CUUCGG/1FYO_11_C_A_0_00013.pdb 
# Loaded reference data/PDB/CUUCGG/2KOC_5_C_A_0_00001.pdb 
# Loaded target data/PDB/CUUCGG/6b4v-pdb-bundle3_1691_C_A_0_00000.pdb 
# Loading data/PDB/CUUCGG/6b4v-pdb-bundle3_1691_C_A_0_00000.pdb 
# Loaded reference data/PDB/CUUCGG/2KOC_5_C_A_0_00001.pdb 
# Loaded target data/PDB/CUUCGG/5KQE_19_C_A_0_00013.pdb 
# Loading data/PDB/CUUCGG/5KQE_19_C_A_0_00013.pdb 
# Loaded reference data/PDB/CUUCGG/2KOC_5_C_A_0_00001.pdb 
# Loaded target data/PDB/CUUCGG/5KQE_19_C_A_0_00007.pdb 
# Loading data/PDB/CUUCGG/5KQE_19_C_A_0_00007.pdb 
# Loaded reference data/PDB/CUUCGG/2KOC_5_C_A_0_00001.pdb 
# Loaded target data/PDB/CUUCGG/2N6S_16_C_A_0_00009.pdb 
# Loading data/PDB/CUUCGG/2N6S_16_C_A_0_00009.pdb 
# Loaded reference data/PDB/CUUCGG/2KOC_5_C_A_0_00001.pdb 
# Loaded target data/PDB/CUUCGG/1FYO_11_C_A_0_00007.pdb 
# Loading data/PDB/CUUCGG/1FYO_11_C_A_0_00007.pdb 
# Loaded reference data/PDB/CUUCGG/2KOC_5_C_A_0_00001.pdb 
# Loaded target data/PDB/CUUCGG/2HEM_10_C_A_0_00002.pdb 
# Loading data/PDB/CUUCGG/2HEM_10_C_A_0_00002.pdb 
# Loaded reference data/PDB/CUUCGG/2KOC_5_C_A_0_00001.pdb 
# Loaded target data/PDB/CUUCGG/4v7t-pdb-bundle1_419_C_A_0_00001.pdb 
# Loading data/PDB/CUUCGG/4v7t-pdb-bundle1_419_C_A_0_00001.pdb 
# Loaded reference data/PDB/CUUCGG/2KOC_5_C_A_0_00001.pdb 
# Loaded target data/PDB/CUUCGG/2N6X_20_C_A_0_00004.pdb 
# Loading data/PDB/CUUCGG/2N6X_20_C_A_0_00004.pdb 
# Loaded reference data/PDB/CUUCGG/2KOC_5_C_A_0_00001.pdb 
# Loaded target data/PDB/CUUCGG/4v4q-pdb-bundle2_1691_C_A_0_00000.pdb 
# Loading data/PDB/CUUCGG/4v4q-pdb-bundle2_1691_C_A_0_00000.pdb 
# Loaded reference data/PDB/CUUCGG/2KOC_5_C_A_0_00001.pdb 
# Loaded target data/PDB/CUUCGG/5w4k-pdb-bundle3_1691_C_A_0_00000.pdb 
# Loading data/PDB/CUUCGG/5w4k-pdb-bundle3_1691_C_A_0_00000.pdb 
# Loaded reference data/PDB/CUUCGG/2KOC_5_C_A_0_00001.pdb 
# Loaded target data/PDB/CUUCGG/2N6X_20_C_A_0_00005.pdb 
# Loading data/PDB/CUUCGG/2N6X_20_C_A_0_00005.pdb 
# Loaded reference data/PDB/CUUCGG/2KOC_5_C_A_0_00001.pdb 
# Loaded target data/PDB/CUUCGG/5on6-pdb-bundle1_1923_C_1_0_00002.pdb 
# Loading data/PDB/CUUCGG/5on6-pdb-bundle1_1923_C_1_0_00002.pdb 
# Loaded reference data/PDB/CUUCGG/2KOC_5_C_A_0_00001.pdb 
# Loaded target data/PDB/CUUCGG/4l47-pdb-bundle2_1691_C_A_0_00000.pdb 
# Loading data/PDB/CUUCGG/4l47-pdb-bundle2_1691_C_A_0_00000.pdb 
# Loaded reference data/PDB/CUUCGG/2KOC_5_C_A_0_00001.pdb 
# Loaded target data/PDB/CUUCGG/2N6S_16_C_A_0_00008.pdb 
# Loading data/PDB/CUUCGG/2N6S_16_C_A_0_00008.pdb 
# Loaded reference data/PDB/CUUCGG/2KOC_5_C_A_0_00001.pdb 
# Loaded target data/PDB/CUUCGG/5KQE_19_C_A_0_00006.pdb 
# Loading data/PDB/CUUCGG/5KQE_19_C_A_0_00006.pdb 
# Loaded reference data/PDB/CUUCGG/2KOC_5_C_A_0_00001.pdb 
# Loaded target data/PDB/CUUCGG/1FYO_11_C_A_0_00006.pdb 
# Loading data/PDB/CUUCGG/1FYO_11_C_A_0_00006.pdb 
# Loaded reference data/PDB/CUUCGG/2KOC_5_C_A_0_00001.pdb 
# Loaded target data/PDB/CUUCGG/5it8-pdb-bundle3_1691_C_A_0_00000.pdb 
# Loading data/PDB/CUUCGG/5it8-pdb-bundle3_1691_C_A_0_00000.pdb 
# Loaded reference data/PDB/CUUCGG/2KOC_5_C_A_0_00001.pdb 
# Loaded target data/PDB/CUUCGG/1FYO_11_C_A_0_00012.pdb 
# Loading data/PDB/CUUCGG/1FYO_11_C_A_0_00012.pdb 
# Loaded reference data/PDB/CUUCGG/2KOC_5_C_A_0_00001.pdb 
# Loaded target data/PDB/CUUCGG/5KQE_19_C_A_0_00012.pdb 
# Loading data/PDB/CUUCGG/5KQE_19_C_A_0_00012.pdb 
# Loaded reference data/PDB/CUUCGG/2KOC_5_C_A_0_00001.pdb 
# Loaded target data/PDB/CUUCGG/1TLR_10_C_A_0_00012.pdb 
# Loading data/PDB/CUUCGG/1TLR_10_C_A_0_00012.pdb 
# Loaded reference data/PDB/CUUCGG/2KOC_5_C_A_0_00001.pdb 
# Loaded target data/PDB/CUUCGG/5hcr-pdb-bundle2_415_C_A_0_00000.pdb 
# Loading data/PDB/CUUCGG/5hcr-pdb-bundle2_415_C_A_0_00000.pdb 
# Loaded reference data/PDB/CUUCGG/2KOC_5_C_A_0_00001.pdb 
# Loaded target data/PDB/CUUCGG/4v7x-pdb-bundle4_1691_C_A_0_00000.pdb 
# Loading data/PDB/CUUCGG/4v7x-pdb-bundle4_1691_C_A_0_00000.pdb 
# Loaded reference data/PDB/CUUCGG/2KOC_5_C_A_0_00001.pdb 
# Loaded target data/PDB/CUUCGG/5j88-pdb-bundle1_419_C_A_0_00001.pdb 
# Loading data/PDB/CUUCGG/5j88-pdb-bundle1_419_C_A_0_00001.pdb 
# Loaded reference data/PDB/CUUCGG/2KOC_5_C_A_0_00001.pdb 
# Loaded target data/PDB/CUUCGG/1TLR_10_C_A_0_00006.pdb 
# Loading data/PDB/CUUCGG/1TLR_10_C_A_0_00006.pdb 
# Loaded reference data/PDB/CUUCGG/2KOC_5_C_A_0_00001.pdb 
# Loaded target data/PDB/CUUCGG/5dgv-pdb-bundle4_1923_C_C_0_00002.pdb 
# Loading data/PDB/CUUCGG/5dgv-pdb-bundle4_1923_C_C_0_00002.pdb 
# Loaded reference data/PDB/CUUCGG/2KOC_5_C_A_0_00001.pdb 
# Loaded target data/PDB/CUUCGG/2ZJR_1708_C_X_0_00000.pdb 
# Loading data/PDB/CUUCGG/2ZJR_1708_C_X_0_00000.pdb 
# Loaded reference data/PDB/CUUCGG/2KOC_5_C_A_0_00001.pdb 
# Loaded target data/PDB/CUUCGG/4u26-pdb-bundle4_1691_C_A_0_00000.pdb 
# Loading data/PDB/CUUCGG/4u26-pdb-bundle4_1691_C_A_0_00000.pdb 
# Loaded reference data/PDB/CUUCGG/2KOC_5_C_A_0_00001.pdb 
# Loaded target data/PDB/CUUCGG/5BR8_419_C_A_0_00000.pdb 
# Loading data/PDB/CUUCGG/5BR8_419_C_A_0_00000.pdb 
# Loaded reference data/PDB/CUUCGG/2KOC_5_C_A_0_00001.pdb 
# Loaded target data/PDB/CUUCGG/4u1v-pdb-bundle1_419_C_A_0_00001.pdb 
# Loading data/PDB/CUUCGG/4u1v-pdb-bundle1_419_C_A_0_00001.pdb 
# Loaded reference data/PDB/CUUCGG/2KOC_5_C_A_0_00001.pdb 
# Loaded target data/PDB/CUUCGG/1D6K_283_C_B_0_00009.pdb 
# Loading data/PDB/CUUCGG/1D6K_283_C_B_0_00009.pdb 
# Loaded reference data/PDB/CUUCGG/2KOC_5_C_A_0_00001.pdb 
# Loaded target data/PDB/CUUCGG/1BYJ_11_C_A_0_00011.pdb 
# Loading data/PDB/CUUCGG/1BYJ_11_C_A_0_00011.pdb 
# Loaded reference data/PDB/CUUCGG/2KOC_5_C_A_0_00001.pdb 
# Loaded target data/PDB/CUUCGG/5el4-pdb-bundle3_419_C_A_0_00000.pdb 
# Loading data/PDB/CUUCGG/5el4-pdb-bundle3_419_C_A_0_00000.pdb 
# Loaded reference data/PDB/CUUCGG/2KOC_5_C_A_0_00001.pdb 
# Loaded target data/PDB/CUUCGG/2LQZ_13_C_A_0_00008.pdb 
# Loading data/PDB/CUUCGG/2LQZ_13_C_A_0_00008.pdb 
# Loaded reference data/PDB/CUUCGG/2KOC_5_C_A_0_00001.pdb 
# Loaded target data/PDB/CUUCGG/4v7v-pdb-bundle2_1691_C_A_0_00000.pdb 
# Loading data/PDB/CUUCGG/4v7v-pdb-bundle2_1691_C_A_0_00000.pdb 
# Loaded reference data/PDB/CUUCGG/2KOC_5_C_A_0_00001.pdb 
# Loaded target data/PDB/CUUCGG/1BYJ_11_C_A_0_00005.pdb 
# Loading data/PDB/CUUCGG/1BYJ_11_C_A_0_00005.pdb 
# Loaded reference data/PDB/CUUCGG/2KOC_5_C_A_0_00001.pdb 
# Loaded target data/PDB/CUUCGG/2HUA_17_C_A_0_00009.pdb 
# Loading data/PDB/CUUCGG/2HUA_17_C_A_0_00009.pdb 
# Loaded reference data/PDB/CUUCGG/2KOC_5_C_A_0_00001.pdb 
# Loaded target data/PDB/CUUCGG/2LKR_63_C_A_0_00006.pdb 
# Loading data/PDB/CUUCGG/2LKR_63_C_A_0_00006.pdb 
# Loaded reference data/PDB/CUUCGG/2KOC_5_C_A_0_00001.pdb 
# Loaded target data/PDB/CUUCGG/5el5-pdb-bundle2_1691_C_A_0_00000.pdb 
# Loading data/PDB/CUUCGG/5el5-pdb-bundle2_1691_C_A_0_00000.pdb 
# Loaded reference data/PDB/CUUCGG/2KOC_5_C_A_0_00001.pdb 
# Loaded target data/PDB/CUUCGG/2M58_12_C_A_0_00009.pdb 
# Loading data/PDB/CUUCGG/2M58_12_C_A_0_00009.pdb 
# Loaded reference data/PDB/CUUCGG/2KOC_5_C_A_0_00001.pdb 
# Loaded target data/PDB/CUUCGG/5j91-pdb-bundle3_1691_C_d_0_00000.pdb 
# Loading data/PDB/CUUCGG/5j91-pdb-bundle3_1691_C_d_0_00000.pdb 
# Loaded reference data/PDB/CUUCGG/2KOC_5_C_A_0_00001.pdb 
# Loaded target data/PDB/CUUCGG/5GAE_1691_C_A_0_00000.pdb 
# Loading data/PDB/CUUCGG/5GAE_1691_C_A_0_00000.pdb 
# Loaded reference data/PDB/CUUCGG/2KOC_5_C_A_0_00001.pdb 
# Loaded target data/PDB/CUUCGG/2N3Q_37_C_A_0_00009.pdb 
# Loading data/PDB/CUUCGG/2N3Q_37_C_A_0_00009.pdb 
# Loaded reference data/PDB/CUUCGG/2KOC_5_C_A_0_00001.pdb 
# Loaded target data/PDB/CUUCGG/5KH8_26_C_A_0_00013.pdb 
# Loading data/PDB/CUUCGG/5KH8_26_C_A_0_00013.pdb 
# Loaded reference data/PDB/CUUCGG/2KOC_5_C_A_0_00001.pdb 
# Loaded target data/PDB/CUUCGG/4v54-pdb-bundle1_207_C_A_0_00000.pdb 
# Loading data/PDB/CUUCGG/4v54-pdb-bundle1_207_C_A_0_00000.pdb 
# Loaded reference data/PDB/CUUCGG/2KOC_5_C_A_0_00001.pdb 
# Loaded target data/PDB/CUUCGG/4ypb-pdb-bundle2_1691_C_A_0_00000.pdb 
# Loading data/PDB/CUUCGG/4ypb-pdb-bundle2_1691_C_A_0_00000.pdb 
# Loaded reference data/PDB/CUUCGG/2KOC_5_C_A_0_00001.pdb 
# Loaded target data/PDB/CUUCGG/2GIO_13_C_A_0_00014.pdb 
# Loading data/PDB/CUUCGG/2GIO_13_C_A_0_00014.pdb 
# Loaded reference data/PDB/CUUCGG/2KOC_5_C_A_0_00001.pdb 
# Loaded target data/PDB/CUUCGG/2GIO_13_C_A_0_00000.pdb 
# Loading data/PDB/CUUCGG/2GIO_13_C_A_0_00000.pdb 
# Loaded reference data/PDB/CUUCGG/2KOC_5_C_A_0_00001.pdb 
# Loaded target data/PDB/CUUCGG/4u56-pdb-bundle2_1923_C_A_0_00002.pdb 
# Loading data/PDB/CUUCGG/4u56-pdb-bundle2_1923_C_A_0_00002.pdb 
# Loaded reference data/PDB/CUUCGG/2KOC_5_C_A_0_00001.pdb 
# Loaded target data/PDB/CUUCGG/1MFY_14_C_A_0_00003.pdb 
# Loading data/PDB/CUUCGG/1MFY_14_C_A_0_00003.pdb 
# Loaded reference data/PDB/CUUCGG/2KOC_5_C_A_0_00001.pdb 
# Loaded target data/PDB/CUUCGG/5KH8_26_C_A_0_00007.pdb 
# Loading data/PDB/CUUCGG/5KH8_26_C_A_0_00007.pdb 
# Loaded reference data/PDB/CUUCGG/2KOC_5_C_A_0_00001.pdb 
# Loaded target data/PDB/CUUCGG/4v8c-pdb-bundle1_1691_C_A_0_00000.pdb 
# Loading data/PDB/CUUCGG/4v8c-pdb-bundle1_1691_C_A_0_00000.pdb 
# Loaded reference data/PDB/CUUCGG/2KOC_5_C_A_0_00001.pdb 
# Loaded target data/PDB/CUUCGG/1JO7_14_C_A_0_00022.pdb 
# Loading data/PDB/CUUCGG/1JO7_14_C_A_0_00022.pdb 
# Loaded reference data/PDB/CUUCGG/2KOC_5_C_A_0_00001.pdb 
# Loaded target data/PDB/CUUCGG/4wro-pdb-bundle3_1691_C_A_0_00000.pdb 
# Loading data/PDB/CUUCGG/4wro-pdb-bundle3_1691_C_A_0_00000.pdb 
# Loaded reference data/PDB/CUUCGG/2KOC_5_C_A_0_00001.pdb 
# Loaded target data/PDB/CUUCGG/2KU0_19_C_A_0_00004.pdb 
# Loading data/PDB/CUUCGG/2KU0_19_C_A_0_00004.pdb 
# Loaded reference data/PDB/CUUCGG/2KOC_5_C_A_0_00001.pdb 
# Loaded target data/PDB/CUUCGG/4v5k-pdb-bundle3_419_C_A_0_00000.pdb 
# Loading data/PDB/CUUCGG/4v5k-pdb-bundle3_419_C_A_0_00000.pdb 
# Loaded reference data/PDB/CUUCGG/2KOC_5_C_A_0_00001.pdb 
# Loaded target data/PDB/CUUCGG/4u20-pdb-bundle1_1449_C_A_0_00003.pdb 
# Loading data/PDB/CUUCGG/4u20-pdb-bundle1_1449_C_A_0_00003.pdb 
# Loaded reference data/PDB/CUUCGG/2KOC_5_C_A_0_00001.pdb 
# Loaded target data/PDB/CUUCGG/1KUQ_32_C_B_0_00001.pdb 
# Loading data/PDB/CUUCGG/1KUQ_32_C_B_0_00001.pdb 
# Loaded reference data/PDB/CUUCGG/2KOC_5_C_A_0_00001.pdb 
# Loaded target data/PDB/CUUCGG/1N32_419_C_A_0_00000.pdb 
# Loading data/PDB/CUUCGG/1N32_419_C_A_0_00000.pdb 
# Loaded reference data/PDB/CUUCGG/2KOC_5_C_A_0_00001.pdb 
# Loaded target data/PDB/CUUCGG/2GIP_12_C_A_0_00007.pdb 
# Loading data/PDB/CUUCGG/2GIP_12_C_A_0_00007.pdb 
# Loaded reference data/PDB/CUUCGG/2KOC_5_C_A_0_00001.pdb 
# Loaded target data/PDB/CUUCGG/2GIP_12_C_A_0_00013.pdb 
# Loading data/PDB/CUUCGG/2GIP_12_C_A_0_00013.pdb 
# Loaded reference data/PDB/CUUCGG/2KOC_5_C_A_0_00001.pdb 
# Loaded target data/PDB/CUUCGG/4v9p-pdb-bundle6_419_C_A_0_00001.pdb 
# Loading data/PDB/CUUCGG/4v9p-pdb-bundle6_419_C_A_0_00001.pdb 
# Loaded reference data/PDB/CUUCGG/2KOC_5_C_A_0_00001.pdb 
# Loaded target data/PDB/CUUCGG/5mdy-pdb-bundle1_1449_C_2_0_00004.pdb 
# Loading data/PDB/CUUCGG/5mdy-pdb-bundle1_1449_C_2_0_00004.pdb 
# Loaded reference data/PDB/CUUCGG/2KOC_5_C_A_0_00001.pdb 
# Loaded target data/PDB/CUUCGG/5fdu-pdb-bundle2_419_C_A_0_00000.pdb 
# Loading data/PDB/CUUCGG/5fdu-pdb-bundle2_419_C_A_0_00000.pdb 
# Loaded reference data/PDB/CUUCGG/2KOC_5_C_A_0_00001.pdb 
# Loaded target data/PDB/CUUCGG/2N3R_37_C_A_0_00019.pdb 
# Loading data/PDB/CUUCGG/2N3R_37_C_A_0_00019.pdb 
# Loaded reference data/PDB/CUUCGG/2KOC_5_C_A_0_00001.pdb 
# Loaded target data/PDB/CUUCGG/5x8p-pdb-bundle1_1727_C_A_0_00001.pdb 
# Loading data/PDB/CUUCGG/5x8p-pdb-bundle1_1727_C_A_0_00001.pdb 
# Loaded reference data/PDB/CUUCGG/2KOC_5_C_A_0_00001.pdb 
# Loaded target data/PDB/CUUCGG/4IOA_1708_C_X_0_00000.pdb 
# Loading data/PDB/CUUCGG/4IOA_1708_C_X_0_00000.pdb 
# Loaded reference data/PDB/CUUCGG/2KOC_5_C_A_0_00001.pdb 
# Loaded target data/PDB/CUUCGG/4u3n-pdb-bundle2_1923_C_A_0_00002.pdb 
# Loading data/PDB/CUUCGG/4u3n-pdb-bundle2_1923_C_A_0_00002.pdb 
# Loaded reference data/PDB/CUUCGG/2KOC_5_C_A_0_00001.pdb 
# Loaded target data/PDB/CUUCGG/2LU0_28_C_A_0_00007.pdb 
# Loading data/PDB/CUUCGG/2LU0_28_C_A_0_00007.pdb 
# Loaded reference data/PDB/CUUCGG/2KOC_5_C_A_0_00001.pdb 
# Loaded target data/PDB/CUUCGG/6BY5_5_C_A_0_00000.pdb 
# Loading data/PDB/CUUCGG/6BY5_5_C_A_0_00000.pdb 
# Loaded reference data/PDB/CUUCGG/2KOC_5_C_A_0_00001.pdb 
# Loaded target data/PDB/CUUCGG/4v5p-pdb-bundle4_1691_C_A_0_00000.pdb 
# Loading data/PDB/CUUCGG/4v5p-pdb-bundle4_1691_C_A_0_00000.pdb 
# Loaded reference data/PDB/CUUCGG/2KOC_5_C_A_0_00001.pdb 
# Loaded target data/PDB/CUUCGG/2LU0_28_C_A_0_00013.pdb 
# Loading data/PDB/CUUCGG/2LU0_28_C_A_0_00013.pdb 
# Loaded reference data/PDB/CUUCGG/2KOC_5_C_A_0_00001.pdb 
# Loaded target data/PDB/CUUCGG/4l47-pdb-bundle3_419_C_A_0_00000.pdb 
# Loading data/PDB/CUUCGG/4l47-pdb-bundle3_419_C_A_0_00000.pdb 
# Loaded reference data/PDB/CUUCGG/2KOC_5_C_A_0_00001.pdb 
# Loaded target data/PDB/CUUCGG/5h5u-pdb-bundle2_419_C_h_0_00001.pdb 
# Loading data/PDB/CUUCGG/5h5u-pdb-bundle2_419_C_h_0_00001.pdb 
# Loaded reference data/PDB/CUUCGG/2KOC_5_C_A_0_00001.pdb 
# Loaded target data/PDB/CUUCGG/4v9d-pdb-bundle3_1691_C_E_0_00000.pdb 
# Loading data/PDB/CUUCGG/4v9d-pdb-bundle3_1691_C_E_0_00000.pdb 
# Loaded reference data/PDB/CUUCGG/2KOC_5_C_A_0_00001.pdb 
# Loaded target data/PDB/CUUCGG/2AU4_20_C_A_0_00000.pdb 
# Loading data/PDB/CUUCGG/2AU4_20_C_A_0_00000.pdb 
# Loaded reference data/PDB/CUUCGG/2KOC_5_C_A_0_00001.pdb 
# Loaded target data/PDB/CUUCGG/1VQP_1769_C_0_0_00000.pdb 
# Loading data/PDB/CUUCGG/1VQP_1769_C_0_0_00000.pdb 
# Loaded reference data/PDB/CUUCGG/2KOC_5_C_A_0_00001.pdb 
# Loaded target data/PDB/CUUCGG/4u24-pdb-bundle1_1449_C_A_0_00003.pdb 
# Loading data/PDB/CUUCGG/4u24-pdb-bundle1_1449_C_A_0_00003.pdb 
# Loaded reference data/PDB/CUUCGG/2KOC_5_C_A_0_00001.pdb 
# Loaded target data/PDB/CUUCGG/1IBL_419_C_A_0_00000.pdb 
# Loading data/PDB/CUUCGG/1IBL_419_C_A_0_00000.pdb 
# Loaded reference data/PDB/CUUCGG/2KOC_5_C_A_0_00001.pdb 
# Loaded target data/PDB/CUUCGG/2LU0_28_C_A_0_00005.pdb 
# Loading data/PDB/CUUCGG/2LU0_28_C_A_0_00005.pdb 
# Loaded reference data/PDB/CUUCGG/2KOC_5_C_A_0_00001.pdb 
# Loaded target data/PDB/CUUCGG/2LU0_28_C_A_0_00011.pdb 
# Loading data/PDB/CUUCGG/2LU0_28_C_A_0_00011.pdb 
# Loaded reference data/PDB/CUUCGG/2KOC_5_C_A_0_00001.pdb 
# Loaded target data/PDB/CUUCGG/5j4d-pdb-bundle2_419_C_0_0_00000.pdb 
# Loading data/PDB/CUUCGG/5j4d-pdb-bundle2_419_C_0_0_00000.pdb 
# Loaded reference data/PDB/CUUCGG/2KOC_5_C_A_0_00001.pdb 
# Loaded target data/PDB/CUUCGG/4wqy-pdb-bundle2_419_C_A_0_00000.pdb 
# Loading data/PDB/CUUCGG/4wqy-pdb-bundle2_419_C_A_0_00000.pdb 
# Loaded reference data/PDB/CUUCGG/2KOC_5_C_A_0_00001.pdb 
# Loaded target data/PDB/CUUCGG/4v7t-pdb-bundle3_419_C_A_0_00001.pdb 
# Loading data/PDB/CUUCGG/4v7t-pdb-bundle3_419_C_A_0_00001.pdb 
# Loaded reference data/PDB/CUUCGG/2KOC_5_C_A_0_00001.pdb 
# Loaded target data/PDB/CUUCGG/6i7v-pdb-bundle1_1449_C_A_0_00003.pdb 
# Loading data/PDB/CUUCGG/6i7v-pdb-bundle1_1449_C_A_0_00003.pdb 
# Loaded reference data/PDB/CUUCGG/2KOC_5_C_A_0_00001.pdb 
# Loaded target data/PDB/CUUCGG/5czp-pdb-bundle2_419_C_A_0_00000.pdb 
# Loading data/PDB/CUUCGG/5czp-pdb-bundle2_419_C_A_0_00000.pdb 
# Loaded reference data/PDB/CUUCGG/2KOC_5_C_A_0_00001.pdb 
# Loaded target data/PDB/CUUCGG/5XYM_1912_C_A_0_00000.pdb 
# Loading data/PDB/CUUCGG/5XYM_1912_C_A_0_00000.pdb 
# Loaded reference data/PDB/CUUCGG/2KOC_5_C_A_0_00001.pdb 
# Loaded target data/PDB/CUUCGG/4u52-pdb-bundle2_1923_C_A_0_00002.pdb 
# Loading data/PDB/CUUCGG/4u52-pdb-bundle2_1923_C_A_0_00002.pdb 
# Loaded reference data/PDB/CUUCGG/2KOC_5_C_A_0_00001.pdb 
# Loaded target data/PDB/CUUCGG/2L3E_18_C_A_0_00008.pdb 
# Loading data/PDB/CUUCGG/2L3E_18_C_A_0_00008.pdb 
# Loaded reference data/PDB/CUUCGG/2KOC_5_C_A_0_00001.pdb 
# Loaded target data/PDB/CUUCGG/2GIP_12_C_A_0_00005.pdb 
# Loading data/PDB/CUUCGG/2GIP_12_C_A_0_00005.pdb 
# Loaded reference data/PDB/CUUCGG/2KOC_5_C_A_0_00001.pdb 
# Loaded target data/PDB/CUUCGG/3T1Y_414_C_A_0_00000.pdb 
# Loading data/PDB/CUUCGG/3T1Y_414_C_A_0_00000.pdb 
# Loaded reference data/PDB/CUUCGG/2KOC_5_C_A_0_00001.pdb 
# Loaded target data/PDB/CUUCGG/2GIP_12_C_A_0_00011.pdb 
# Loading data/PDB/CUUCGG/2GIP_12_C_A_0_00011.pdb 
# Loaded reference data/PDB/CUUCGG/2KOC_5_C_A_0_00001.pdb 
# Loaded target data/PDB/CUUCGG/2KU0_19_C_A_0_00006.pdb 
# Loading data/PDB/CUUCGG/2KU0_19_C_A_0_00006.pdb 
# Loaded reference data/PDB/CUUCGG/2KOC_5_C_A_0_00001.pdb 
# Loaded target data/PDB/CUUCGG/5lzd-pdb-bundle1_1449_C_a_0_00003.pdb 
# Loading data/PDB/CUUCGG/5lzd-pdb-bundle1_1449_C_a_0_00003.pdb 
# Loaded reference data/PDB/CUUCGG/2KOC_5_C_A_0_00001.pdb 
# Loaded target data/PDB/CUUCGG/1JO7_14_C_A_0_00020.pdb 
# Loading data/PDB/CUUCGG/1JO7_14_C_A_0_00020.pdb 
# Loaded reference data/PDB/CUUCGG/2KOC_5_C_A_0_00001.pdb 
# Loaded target data/PDB/CUUCGG/1A3M_11_C_A_0_00008.pdb 
# Loading data/PDB/CUUCGG/1A3M_11_C_A_0_00008.pdb 
# Loaded reference data/PDB/CUUCGG/2KOC_5_C_A_0_00001.pdb 
# Loaded target data/PDB/CUUCGG/6gsl-pdb-bundle4_1691_C_A_0_00000.pdb 
# Loading data/PDB/CUUCGG/6gsl-pdb-bundle4_1691_C_A_0_00000.pdb 
# Loaded reference data/PDB/CUUCGG/2KOC_5_C_A_0_00001.pdb 
# Loaded target data/PDB/CUUCGG/2GIO_13_C_A_0_00016.pdb 
# Loading data/PDB/CUUCGG/2GIO_13_C_A_0_00016.pdb 
# Loaded reference data/PDB/CUUCGG/2KOC_5_C_A_0_00001.pdb 
# Loaded target data/PDB/CUUCGG/5KH8_26_C_A_0_00011.pdb 
# Loading data/PDB/CUUCGG/5KH8_26_C_A_0_00011.pdb 
# Loaded reference data/PDB/CUUCGG/2KOC_5_C_A_0_00001.pdb 
# Loaded target data/PDB/CUUCGG/1MFY_14_C_A_0_00015.pdb 
# Loading data/PDB/CUUCGG/1MFY_14_C_A_0_00015.pdb 
# Loaded reference data/PDB/CUUCGG/2KOC_5_C_A_0_00001.pdb 
# Loaded target data/PDB/CUUCGG/5KH8_26_C_A_0_00005.pdb 
# Loading data/PDB/CUUCGG/5KH8_26_C_A_0_00005.pdb 
# Loaded reference data/PDB/CUUCGG/2KOC_5_C_A_0_00001.pdb 
# Loaded target data/PDB/CUUCGG/1MFY_14_C_A_0_00001.pdb 
# Loading data/PDB/CUUCGG/1MFY_14_C_A_0_00001.pdb 
# Loaded reference data/PDB/CUUCGG/2KOC_5_C_A_0_00001.pdb 
# Loaded target data/PDB/CUUCGG/2GIO_13_C_A_0_00002.pdb 
# Loading data/PDB/CUUCGG/2GIO_13_C_A_0_00002.pdb 
# Loaded reference data/PDB/CUUCGG/2KOC_5_C_A_0_00001.pdb 
# Loaded target data/PDB/CUUCGG/5wis-pdb-bundle3_1691_C_A_0_00000.pdb 
# Loading data/PDB/CUUCGG/5wis-pdb-bundle3_1691_C_A_0_00000.pdb 
# Loaded reference data/PDB/CUUCGG/2KOC_5_C_A_0_00001.pdb 
# Loaded target data/PDB/CUUCGG/6eri-pdb-bundle1_1727_C_A_0_00001.pdb 
# Loading data/PDB/CUUCGG/6eri-pdb-bundle1_1727_C_A_0_00001.pdb 
# Loaded reference data/PDB/CUUCGG/2KOC_5_C_A_0_00001.pdb 
# Loaded target data/PDB/CUUCGG/5wit-pdb-bundle4_419_C_A_0_00000.pdb 
# Loading data/PDB/CUUCGG/5wit-pdb-bundle4_419_C_A_0_00000.pdb 
# Loaded reference data/PDB/CUUCGG/2KOC_5_C_A_0_00001.pdb 
# Loaded target data/PDB/CUUCGG/1YI2_1769_C_0_0_00000.pdb 
# Loading data/PDB/CUUCGG/1YI2_1769_C_0_0_00000.pdb 
# Loaded reference data/PDB/CUUCGG/2KOC_5_C_A_0_00001.pdb 
# Loaded target data/PDB/CUUCGG/1BYJ_11_C_A_0_00013.pdb 
# Loading data/PDB/CUUCGG/1BYJ_11_C_A_0_00013.pdb 
# Loaded reference data/PDB/CUUCGG/2KOC_5_C_A_0_00001.pdb 
# Loaded target data/PDB/CUUCGG/1BYJ_11_C_A_0_00007.pdb 
# Loading data/PDB/CUUCGG/1BYJ_11_C_A_0_00007.pdb 
# Loaded reference data/PDB/CUUCGG/2KOC_5_C_A_0_00001.pdb 
# Loaded target data/PDB/CUUCGG/5LYV_48_C_B_0_00001.pdb 
# Loading data/PDB/CUUCGG/5LYV_48_C_B_0_00001.pdb 
# Loaded reference data/PDB/CUUCGG/2KOC_5_C_A_0_00001.pdb 
# Loaded target data/PDB/CUUCGG/4v5p-pdb-bundle1_419_C_A_0_00000.pdb 
# Loading data/PDB/CUUCGG/4v5p-pdb-bundle1_419_C_A_0_00000.pdb 
# Loaded reference data/PDB/CUUCGG/2KOC_5_C_A_0_00001.pdb 
# Loaded target data/PDB/CUUCGG/3CXC_1769_C_0_0_00000.pdb 
# Loading data/PDB/CUUCGG/3CXC_1769_C_0_0_00000.pdb 
# Loaded reference data/PDB/CUUCGG/2KOC_5_C_A_0_00001.pdb 
# Loaded target data/PDB/CUUCGG/3G71_1769_C_0_0_00000.pdb 
# Loading data/PDB/CUUCGG/3G71_1769_C_0_0_00000.pdb 
# Loaded reference data/PDB/CUUCGG/2KOC_5_C_A_0_00001.pdb 
# Loaded target data/PDB/CUUCGG/1K2G_5_C_A_0_00000.pdb 
# Loading data/PDB/CUUCGG/1K2G_5_C_A_0_00000.pdb 
# Loaded reference data/PDB/CUUCGG/2KOC_5_C_A_0_00001.pdb 
# Loaded target data/PDB/CUUCGG/4JI3_419_C_A_0_00000.pdb 
# Loading data/PDB/CUUCGG/4JI3_419_C_A_0_00000.pdb 
# Loaded reference data/PDB/CUUCGG/2KOC_5_C_A_0_00001.pdb 
# Loaded target data/PDB/CUUCGG/4v8d-pdb-bundle3_419_C_A_0_00000.pdb 
# Loading data/PDB/CUUCGG/4v8d-pdb-bundle3_419_C_A_0_00000.pdb 
# Loaded reference data/PDB/CUUCGG/2KOC_5_C_A_0_00001.pdb 
# Loaded target data/PDB/CUUCGG/5j4d-pdb-bundle1_419_C_A_0_00000.pdb 
# Loading data/PDB/CUUCGG/5j4d-pdb-bundle1_419_C_A_0_00000.pdb 
# Loaded reference data/PDB/CUUCGG/2KOC_5_C_A_0_00001.pdb 
# Loaded target data/PDB/CUUCGG/4w4g-pdb-bundle3_419_C_A_0_00000.pdb 
# Loading data/PDB/CUUCGG/4w4g-pdb-bundle3_419_C_A_0_00000.pdb 
# Loaded reference data/PDB/CUUCGG/2KOC_5_C_A_0_00001.pdb 
# Loaded target data/PDB/CUUCGG/1TLR_10_C_A_0_00010.pdb 
# Loading data/PDB/CUUCGG/1TLR_10_C_A_0_00010.pdb 
# Loaded reference data/PDB/CUUCGG/2KOC_5_C_A_0_00001.pdb 
# Loaded target data/PDB/CUUCGG/1TLR_10_C_A_0_00004.pdb 
# Loading data/PDB/CUUCGG/1TLR_10_C_A_0_00004.pdb 
# Loaded reference data/PDB/CUUCGG/2KOC_5_C_A_0_00001.pdb 
# Loaded target data/PDB/CUUCGG/2LWK_15_C_A_0_00009.pdb 
# Loading data/PDB/CUUCGG/2LWK_15_C_A_0_00009.pdb 
# Loaded reference data/PDB/CUUCGG/2KOC_5_C_A_0_00001.pdb 
# Loaded target data/PDB/CUUCGG/1FJG_419_C_A_0_00000.pdb 
# Loading data/PDB/CUUCGG/1FJG_419_C_A_0_00000.pdb 
# Loaded reference data/PDB/CUUCGG/2KOC_5_C_A_0_00001.pdb 
# Loaded target data/PDB/CUUCGG/1FYO_11_C_A_0_00004.pdb 
# Loading data/PDB/CUUCGG/1FYO_11_C_A_0_00004.pdb 
# Loaded reference data/PDB/CUUCGG/2KOC_5_C_A_0_00001.pdb 
# Loaded target data/PDB/CUUCGG/5KQE_19_C_A_0_00004.pdb 
# Loading data/PDB/CUUCGG/5KQE_19_C_A_0_00004.pdb 
# Loaded reference data/PDB/CUUCGG/2KOC_5_C_A_0_00001.pdb 
# Loaded target data/PDB/CUUCGG/5KQE_19_C_A_0_00010.pdb 
# Loading data/PDB/CUUCGG/5KQE_19_C_A_0_00010.pdb 
# Loaded reference data/PDB/CUUCGG/2KOC_5_C_A_0_00001.pdb 
# Loaded target data/PDB/CUUCGG/5j91-pdb-bundle1_1449_C_b_0_00007.pdb 
# Loading data/PDB/CUUCGG/5j91-pdb-bundle1_1449_C_b_0_00007.pdb 
# Loaded reference data/PDB/CUUCGG/2KOC_5_C_A_0_00001.pdb 
# Loaded target data/PDB/CUUCGG/1FYO_11_C_A_0_00010.pdb 
# Loading data/PDB/CUUCGG/1FYO_11_C_A_0_00010.pdb 
# Loaded reference data/PDB/CUUCGG/2KOC_5_C_A_0_00001.pdb 
# Loaded target data/PDB/CUUCGG/4wt8-pdb-bundle1_415_C_X_0_00001.pdb 
# Loading data/PDB/CUUCGG/4wt8-pdb-bundle1_415_C_X_0_00001.pdb 
# Loaded reference data/PDB/CUUCGG/2KOC_5_C_A_0_00001.pdb 
# Loaded target data/PDB/CUUCGG/2N6X_20_C_A_0_00007.pdb 
# Loading data/PDB/CUUCGG/2N6X_20_C_A_0_00007.pdb 
# Loaded reference data/PDB/CUUCGG/2KOC_5_C_A_0_00001.pdb 
# Loaded target data/PDB/CUUCGG/2HEM_10_C_A_0_00001.pdb 
# Loading data/PDB/CUUCGG/2HEM_10_C_A_0_00001.pdb 
# Loaded reference data/PDB/CUUCGG/2KOC_5_C_A_0_00001.pdb 
# Loaded target data/PDB/CUUCGG/4wt1-pdb-bundle1_419_C_A_0_00000.pdb 
# Loading data/PDB/CUUCGG/4wt1-pdb-bundle1_419_C_A_0_00000.pdb 
# Loaded reference data/PDB/CUUCGG/2KOC_5_C_A_0_00001.pdb 
# Loaded target data/PDB/CUUCGG/2HEM_10_C_A_0_00000.pdb 
# Loading data/PDB/CUUCGG/2HEM_10_C_A_0_00000.pdb 
# Loaded reference data/PDB/CUUCGG/2KOC_5_C_A_0_00001.pdb 
# Loaded target data/PDB/CUUCGG/2N6X_20_C_A_0_00006.pdb 
# Loading data/PDB/CUUCGG/2N6X_20_C_A_0_00006.pdb 
# Loaded reference data/PDB/CUUCGG/2KOC_5_C_A_0_00001.pdb 
# Loaded target data/PDB/CUUCGG/4v90-pdb-bundle2_1691_C_A_0_00000.pdb 
# Loading data/PDB/CUUCGG/4v90-pdb-bundle2_1691_C_A_0_00000.pdb 
# Loaded reference data/PDB/CUUCGG/2KOC_5_C_A_0_00001.pdb 
# Loaded target data/PDB/CUUCGG/1ZHO_15_C_D_0_00001.pdb 
# Loading data/PDB/CUUCGG/1ZHO_15_C_D_0_00001.pdb 
# Loaded reference data/PDB/CUUCGG/2KOC_5_C_A_0_00001.pdb 
# Loaded target data/PDB/CUUCGG/5KQE_19_C_A_0_00011.pdb 
# Loading data/PDB/CUUCGG/5KQE_19_C_A_0_00011.pdb 
# Loaded reference data/PDB/CUUCGG/2KOC_5_C_A_0_00001.pdb 
# Loaded target data/PDB/CUUCGG/5j4b-pdb-bundle1_1738_C_A_0_00000.pdb 
# Loading data/PDB/CUUCGG/5j4b-pdb-bundle1_1738_C_A_0_00000.pdb 
# Loaded reference data/PDB/CUUCGG/2KOC_5_C_A_0_00001.pdb 
# Loaded target data/PDB/CUUCGG/1FYO_11_C_A_0_00011.pdb 
# Loading data/PDB/CUUCGG/1FYO_11_C_A_0_00011.pdb 
# Loaded reference data/PDB/CUUCGG/2KOC_5_C_A_0_00001.pdb 
# Loaded target data/PDB/CUUCGG/1FYO_11_C_A_0_00005.pdb 
# Loading data/PDB/CUUCGG/1FYO_11_C_A_0_00005.pdb 
# Loaded reference data/PDB/CUUCGG/2KOC_5_C_A_0_00001.pdb 
# Loaded target data/PDB/CUUCGG/5KQE_19_C_A_0_00005.pdb 
# Loading data/PDB/CUUCGG/5KQE_19_C_A_0_00005.pdb 
# Loaded reference data/PDB/CUUCGG/2KOC_5_C_A_0_00001.pdb 
# Loaded target data/PDB/CUUCGG/4v8i-pdb-bundle2_1691_C_A_0_00000.pdb 
# Loading data/PDB/CUUCGG/4v8i-pdb-bundle2_1691_C_A_0_00000.pdb 
# Loaded reference data/PDB/CUUCGG/2KOC_5_C_A_0_00001.pdb 
# Loaded target data/PDB/CUUCGG/2LWK_15_C_A_0_00008.pdb 
# Loading data/PDB/CUUCGG/2LWK_15_C_A_0_00008.pdb 
# Loaded reference data/PDB/CUUCGG/2KOC_5_C_A_0_00001.pdb 
# Loaded target data/PDB/CUUCGG/1TLR_10_C_A_0_00005.pdb 
# Loading data/PDB/CUUCGG/1TLR_10_C_A_0_00005.pdb 
# Loaded reference data/PDB/CUUCGG/2KOC_5_C_A_0_00001.pdb 
# Loaded target data/PDB/CUUCGG/1TLR_10_C_A_0_00011.pdb 
# Loading data/PDB/CUUCGG/1TLR_10_C_A_0_00011.pdb 
# Loaded reference data/PDB/CUUCGG/2KOC_5_C_A_0_00001.pdb 
# Loaded target data/PDB/CUUCGG/4www-pdb-bundle2_419_C_V_0_00005.pdb 
# Loading data/PDB/CUUCGG/4www-pdb-bundle2_419_C_V_0_00005.pdb 
# Loaded reference data/PDB/CUUCGG/2KOC_5_C_A_0_00001.pdb 
# Loaded target data/PDB/CUUCGG/5mmm-pdb-bundle2_977_C_a_0_00000.pdb 
# Loading data/PDB/CUUCGG/5mmm-pdb-bundle2_977_C_a_0_00000.pdb 
# Loaded reference data/PDB/CUUCGG/2KOC_5_C_A_0_00001.pdb 
# Loaded target data/PDB/CUUCGG/4l47-pdb-bundle1_419_C_A_0_00000.pdb 
# Loading data/PDB/CUUCGG/4l47-pdb-bundle1_419_C_A_0_00000.pdb 
# Loaded reference data/PDB/CUUCGG/2KOC_5_C_A_0_00001.pdb 
# Loaded target data/PDB/CUUCGG/6fkr-pdb-bundle4_419_C_A_0_00000.pdb 
# Loading data/PDB/CUUCGG/6fkr-pdb-bundle4_419_C_A_0_00000.pdb 
# Loaded reference data/PDB/CUUCGG/2KOC_5_C_A_0_00001.pdb 
# Loaded target data/PDB/CUUCGG/6mte-pdb-bundle1_2872_C_5_0_00000.pdb 
# Loading data/PDB/CUUCGG/6mte-pdb-bundle1_2872_C_5_0_00000.pdb 
# Loaded reference data/PDB/CUUCGG/2KOC_5_C_A_0_00001.pdb 
# Loaded target data/PDB/CUUCGG/4v6c-pdb-bundle3_419_C_N_0_00001.pdb 
# Loading data/PDB/CUUCGG/4v6c-pdb-bundle3_419_C_N_0_00001.pdb 
# Loaded reference data/PDB/CUUCGG/2KOC_5_C_A_0_00001.pdb 
# Loaded target data/PDB/CUUCGG/4v8g-pdb-bundle4_1691_C_A_0_00000.pdb 
# Loading data/PDB/CUUCGG/4v8g-pdb-bundle4_1691_C_A_0_00000.pdb 
# Loaded reference data/PDB/CUUCGG/2KOC_5_C_A_0_00001.pdb 
# Loaded target data/PDB/CUUCGG/2AAR_1708_C_0_0_00000.pdb 
# Loading data/PDB/CUUCGG/2AAR_1708_C_0_0_00000.pdb 
# Loaded reference data/PDB/CUUCGG/2KOC_5_C_A_0_00001.pdb 
# Loaded target data/PDB/CUUCGG/1vvj-pdb-bundle2_1691_C_A_0_00000.pdb 
# Loading data/PDB/CUUCGG/1vvj-pdb-bundle2_1691_C_A_0_00000.pdb 
# Loaded reference data/PDB/CUUCGG/2KOC_5_C_A_0_00001.pdb 
# Loaded target data/PDB/CUUCGG/5j7l-pdb-bundle1_419_C_b_0_00005.pdb 
# Loading data/PDB/CUUCGG/5j7l-pdb-bundle1_419_C_b_0_00005.pdb 
# Loaded reference data/PDB/CUUCGG/2KOC_5_C_A_0_00001.pdb 
# Loaded target data/PDB/CUUCGG/6bu8-pdb-bundle1_1449_C_0_0_00003.pdb 
# Loading data/PDB/CUUCGG/6bu8-pdb-bundle1_1449_C_0_0_00003.pdb 
# Loaded reference data/PDB/CUUCGG/2KOC_5_C_A_0_00001.pdb 
# Loaded target data/PDB/CUUCGG/1BYJ_11_C_A_0_00006.pdb 
# Loading data/PDB/CUUCGG/1BYJ_11_C_A_0_00006.pdb 
# Loaded reference data/PDB/CUUCGG/2KOC_5_C_A_0_00001.pdb 
# Loaded target data/PDB/CUUCGG/6gzq-pdb-bundle1_1740_C_c_0_00000.pdb 
# Loading data/PDB/CUUCGG/6gzq-pdb-bundle1_1740_C_c_0_00000.pdb 
# Loaded reference data/PDB/CUUCGG/2KOC_5_C_A_0_00001.pdb 
# Loaded target data/PDB/CUUCGG/4v9p-pdb-bundle4_419_C_A_0_00001.pdb 
# Loading data/PDB/CUUCGG/4v9p-pdb-bundle4_419_C_A_0_00001.pdb 
# Loaded reference data/PDB/CUUCGG/2KOC_5_C_A_0_00001.pdb 
# Loaded target data/PDB/CUUCGG/1BYJ_11_C_A_0_00012.pdb 
# Loading data/PDB/CUUCGG/1BYJ_11_C_A_0_00012.pdb 
# Loaded reference data/PDB/CUUCGG/2KOC_5_C_A_0_00001.pdb 
# Loaded target data/PDB/CUUCGG/6bz6-pdb-bundle4_1691_C_A_0_00000.pdb 
# Loading data/PDB/CUUCGG/6bz6-pdb-bundle4_1691_C_A_0_00000.pdb 
# Loaded reference data/PDB/CUUCGG/2KOC_5_C_A_0_00001.pdb 
# Loaded target data/PDB/CUUCGG/4v5k-pdb-bundle1_419_C_A_0_00000.pdb 
# Loading data/PDB/CUUCGG/4v5k-pdb-bundle1_419_C_A_0_00000.pdb 
# Loaded reference data/PDB/CUUCGG/2KOC_5_C_A_0_00001.pdb 
# Loaded target data/PDB/CUUCGG/2LKR_63_C_A_0_00005.pdb 
# Loading data/PDB/CUUCGG/2LKR_63_C_A_0_00005.pdb 
# Loaded reference data/PDB/CUUCGG/2KOC_5_C_A_0_00001.pdb 
# Loaded target data/PDB/CUUCGG/4u1v-pdb-bundle3_419_C_A_0_00001.pdb 
# Loading data/PDB/CUUCGG/4u1v-pdb-bundle3_419_C_A_0_00001.pdb 
# Loaded reference data/PDB/CUUCGG/2KOC_5_C_A_0_00001.pdb 
# Loaded target data/PDB/CUUCGG/1MFY_14_C_A_0_00000.pdb 
# Loading data/PDB/CUUCGG/1MFY_14_C_A_0_00000.pdb 
# Loaded reference data/PDB/CUUCGG/2KOC_5_C_A_0_00001.pdb 
# Loaded target data/PDB/CUUCGG/5KH8_26_C_A_0_00004.pdb 
# Loading data/PDB/CUUCGG/5KH8_26_C_A_0_00004.pdb 
# Loaded reference data/PDB/CUUCGG/2KOC_5_C_A_0_00001.pdb 
# Loaded target data/PDB/CUUCGG/4v5c-pdb-bundle4_1691_C_A_0_00000.pdb 
# Loading data/PDB/CUUCGG/4v5c-pdb-bundle4_1691_C_A_0_00000.pdb 
# Loaded reference data/PDB/CUUCGG/2KOC_5_C_A_0_00001.pdb 
# Loaded target data/PDB/CUUCGG/1XST_12_C_A_0_00000.pdb 
# Loading data/PDB/CUUCGG/1XST_12_C_A_0_00000.pdb 
# Loaded reference data/PDB/CUUCGG/2KOC_5_C_A_0_00001.pdb 
# Loaded target data/PDB/CUUCGG/2GIO_13_C_A_0_00003.pdb 
# Loading data/PDB/CUUCGG/2GIO_13_C_A_0_00003.pdb 
# Loaded reference data/PDB/CUUCGG/2KOC_5_C_A_0_00001.pdb 
# Loaded target data/PDB/CUUCGG/2GIO_13_C_A_0_00017.pdb 
# Loading data/PDB/CUUCGG/2GIO_13_C_A_0_00017.pdb 
# Loaded reference data/PDB/CUUCGG/2KOC_5_C_A_0_00001.pdb 
# Loaded target data/PDB/CUUCGG/1MFY_14_C_A_0_00014.pdb 
# Loading data/PDB/CUUCGG/1MFY_14_C_A_0_00014.pdb 
# Loaded reference data/PDB/CUUCGG/2KOC_5_C_A_0_00001.pdb 
# Loaded target data/PDB/CUUCGG/5KH8_26_C_A_0_00010.pdb 
# Loading data/PDB/CUUCGG/5KH8_26_C_A_0_00010.pdb 
# Loaded reference data/PDB/CUUCGG/2KOC_5_C_A_0_00001.pdb 
# Loaded target data/PDB/CUUCGG/4v7t-pdb-bundle1_1028_C_A_0_00002.pdb 
# Loading data/PDB/CUUCGG/4v7t-pdb-bundle1_1028_C_A_0_00002.pdb 
# Loaded reference data/PDB/CUUCGG/2KOC_5_C_A_0_00001.pdb 
# Loaded target data/PDB/CUUCGG/1A3M_11_C_A_0_00009.pdb 
# Loading data/PDB/CUUCGG/1A3M_11_C_A_0_00009.pdb 
# Loaded reference data/PDB/CUUCGG/2KOC_5_C_A_0_00001.pdb 
# Loaded target data/PDB/CUUCGG/5NRG_1735_C_X_0_00000.pdb 
# Loading data/PDB/CUUCGG/5NRG_1735_C_X_0_00000.pdb 
# Loaded reference data/PDB/CUUCGG/2KOC_5_C_A_0_00001.pdb 
# Loaded target data/PDB/CUUCGG/5dat-pdb-bundle2_1923_C_A_0_00002.pdb 
# Loading data/PDB/CUUCGG/5dat-pdb-bundle2_1923_C_A_0_00002.pdb 
# Loaded reference data/PDB/CUUCGG/2KOC_5_C_A_0_00001.pdb 
# Loaded target data/PDB/CUUCGG/4wr6-pdb-bundle2_1691_C_A_0_00000.pdb 
# Loading data/PDB/CUUCGG/4wr6-pdb-bundle2_1691_C_A_0_00000.pdb 
# Loaded reference data/PDB/CUUCGG/2KOC_5_C_A_0_00001.pdb 
# Loaded target data/PDB/CUUCGG/5fci-pdb-bundle4_1923_C_I_0_00002.pdb 
# Loading data/PDB/CUUCGG/5fci-pdb-bundle4_1923_C_I_0_00002.pdb 
# Loaded reference data/PDB/CUUCGG/2KOC_5_C_A_0_00001.pdb 
# Loaded target data/PDB/CUUCGG/2KU0_19_C_A_0_00007.pdb 
# Loading data/PDB/CUUCGG/2KU0_19_C_A_0_00007.pdb 
# Loaded reference data/PDB/CUUCGG/2KOC_5_C_A_0_00001.pdb 
# Loaded target data/PDB/CUUCGG/2GIP_12_C_A_0_00010.pdb 
# Loading data/PDB/CUUCGG/2GIP_12_C_A_0_00010.pdb 
# Loaded reference data/PDB/CUUCGG/2KOC_5_C_A_0_00001.pdb 
# Loaded target data/PDB/CUUCGG/4v64-pdb-bundle1_1449_C_A_0_00003.pdb 
# Loading data/PDB/CUUCGG/4v64-pdb-bundle1_1449_C_A_0_00003.pdb 
# Loaded reference data/PDB/CUUCGG/2KOC_5_C_A_0_00001.pdb 
# Loaded target data/PDB/CUUCGG/5obm-pdb-bundle4_230_C_f_0_00000.pdb 
# Loading data/PDB/CUUCGG/5obm-pdb-bundle4_230_C_f_0_00000.pdb 
# Loaded reference data/PDB/CUUCGG/2KOC_5_C_A_0_00001.pdb 
# Loaded target data/PDB/CUUCGG/6nd6-pdb-bundle1_1738_C_A_0_00000.pdb 
# Loading data/PDB/CUUCGG/6nd6-pdb-bundle1_1738_C_A_0_00000.pdb 
# Loaded reference data/PDB/CUUCGG/2KOC_5_C_A_0_00001.pdb 
# Loaded target data/PDB/CUUCGG/2GIP_12_C_A_0_00004.pdb 
# Loading data/PDB/CUUCGG/2GIP_12_C_A_0_00004.pdb 
# Loaded reference data/PDB/CUUCGG/2KOC_5_C_A_0_00001.pdb 
# Loaded target data/PDB/CUUCGG/3T1H_414_C_A_0_00000.pdb 
# Loading data/PDB/CUUCGG/3T1H_414_C_A_0_00000.pdb 
# Loaded reference data/PDB/CUUCGG/2KOC_5_C_A_0_00001.pdb 
# Loaded target data/PDB/CUUCGG/5we4-pdb-bundle2_1449_C_a_0_00003.pdb 
# Loading data/PDB/CUUCGG/5we4-pdb-bundle2_1449_C_a_0_00003.pdb 
# Loaded reference data/PDB/CUUCGG/2KOC_5_C_A_0_00001.pdb 
# Loaded target data/PDB/CUUCGG/2L3E_18_C_A_0_00009.pdb 
# Loading data/PDB/CUUCGG/2L3E_18_C_A_0_00009.pdb 
# Loaded reference data/PDB/CUUCGG/2KOC_5_C_A_0_00001.pdb 
# Loaded target data/PDB/CUUCGG/4wq1-pdb-bundle2_1691_C_A_0_00000.pdb 
# Loading data/PDB/CUUCGG/4wq1-pdb-bundle2_1691_C_A_0_00000.pdb 
# Loaded reference data/PDB/CUUCGG/2KOC_5_C_A_0_00001.pdb 
# Loaded target data/PDB/CUUCGG/2LU0_28_C_A_0_00010.pdb 
# Loading data/PDB/CUUCGG/2LU0_28_C_A_0_00010.pdb 
# Loaded reference data/PDB/CUUCGG/2KOC_5_C_A_0_00001.pdb 
# Loaded target data/PDB/CUUCGG/2LU0_28_C_A_0_00004.pdb 
# Loading data/PDB/CUUCGG/2LU0_28_C_A_0_00004.pdb 
# Loaded reference data/PDB/CUUCGG/2KOC_5_C_A_0_00001.pdb 
# Loaded target data/PDB/CUUCGG/5ibb-pdb-bundle1_419_C_A_0_00000.pdb 
# Loading data/PDB/CUUCGG/5ibb-pdb-bundle1_419_C_A_0_00000.pdb 
# Loaded reference data/PDB/CUUCGG/2KOC_5_C_A_0_00001.pdb 
# Loaded target data/PDB/CUUCGG/2LU0_28_C_A_0_00000.pdb 
# Loading data/PDB/CUUCGG/2LU0_28_C_A_0_00000.pdb 
# Loaded reference data/PDB/CUUCGG/2KOC_5_C_A_0_00001.pdb 
# Loaded target data/PDB/CUUCGG/5we4-pdb-bundle1_1691_C_A_0_00000.pdb 
# Loading data/PDB/CUUCGG/5we4-pdb-bundle1_1691_C_A_0_00000.pdb 
# Loaded reference data/PDB/CUUCGG/2KOC_5_C_A_0_00001.pdb 
# Loaded target data/PDB/CUUCGG/5tgm-pdb-bundle2_1923_C_A_0_00002.pdb 
# Loading data/PDB/CUUCGG/5tgm-pdb-bundle2_1923_C_A_0_00002.pdb 
# Loaded reference data/PDB/CUUCGG/2KOC_5_C_A_0_00001.pdb 
# Loaded target data/PDB/CUUCGG/6BY5_5_C_A_0_00007.pdb 
# Loading data/PDB/CUUCGG/6BY5_5_C_A_0_00007.pdb 
# Loaded reference data/PDB/CUUCGG/2KOC_5_C_A_0_00001.pdb 
# Loaded target data/PDB/CUUCGG/4v9p-pdb-bundle2_419_C_A_0_00001.pdb 
# Loading data/PDB/CUUCGG/4v9p-pdb-bundle2_419_C_A_0_00001.pdb 
# Loaded reference data/PDB/CUUCGG/2KOC_5_C_A_0_00001.pdb 
# Loaded target data/PDB/CUUCGG/2LU0_28_C_A_0_00014.pdb 
# Loading data/PDB/CUUCGG/2LU0_28_C_A_0_00014.pdb 
# Loaded reference data/PDB/CUUCGG/2KOC_5_C_A_0_00001.pdb 
# Loaded target data/PDB/CUUCGG/4v9i-pdb-bundle3_415_C_A_0_00000.pdb 
# Loading data/PDB/CUUCGG/4v9i-pdb-bundle3_415_C_A_0_00000.pdb 
# Loaded reference data/PDB/CUUCGG/2KOC_5_C_A_0_00001.pdb 
# Loaded target data/PDB/CUUCGG/4v52-pdb-bundle3_419_C_A_0_00001.pdb 
# Loading data/PDB/CUUCGG/4v52-pdb-bundle3_419_C_A_0_00001.pdb 
# Loaded reference data/PDB/CUUCGG/2KOC_5_C_A_0_00001.pdb 
# Loaded target data/PDB/CUUCGG/1IBK_419_C_A_0_00000.pdb 
# Loading data/PDB/CUUCGG/1IBK_419_C_A_0_00000.pdb 
# Loaded reference data/PDB/CUUCGG/2KOC_5_C_A_0_00001.pdb 
# Loaded target data/PDB/CUUCGG/1HNW_419_C_A_0_00000.pdb 
# Loading data/PDB/CUUCGG/1HNW_419_C_A_0_00000.pdb 
# Loaded reference data/PDB/CUUCGG/2KOC_5_C_A_0_00001.pdb 
# Loaded target data/PDB/CUUCGG/5fdu-pdb-bundle1_1738_C_A_0_00000.pdb 
# Loading data/PDB/CUUCGG/5fdu-pdb-bundle1_1738_C_A_0_00000.pdb 
# Loaded reference data/PDB/CUUCGG/2KOC_5_C_A_0_00001.pdb 
# Loaded target data/PDB/CUUCGG/2L3E_18_C_A_0_00019.pdb 
# Loading data/PDB/CUUCGG/2L3E_18_C_A_0_00019.pdb 
# Loaded reference data/PDB/CUUCGG/2KOC_5_C_A_0_00001.pdb 
# Loaded target data/PDB/CUUCGG/6boh-pdb-bundle3_1691_C_A_0_00000.pdb 
# Loading data/PDB/CUUCGG/6boh-pdb-bundle3_1691_C_A_0_00000.pdb 
# Loaded reference data/PDB/CUUCGG/2KOC_5_C_A_0_00001.pdb 
# Loaded target data/PDB/CUUCGG/4GKJ_414_C_A_0_00000.pdb 
# Loading data/PDB/CUUCGG/4GKJ_414_C_A_0_00000.pdb 
# Loaded reference data/PDB/CUUCGG/2KOC_5_C_A_0_00001.pdb 
# Loaded target data/PDB/CUUCGG/5fcj-pdb-bundle4_1923_C_F_0_00002.pdb 
# Loading data/PDB/CUUCGG/5fcj-pdb-bundle4_1923_C_F_0_00002.pdb 
# Loaded reference data/PDB/CUUCGG/2KOC_5_C_A_0_00001.pdb 
# Loaded target data/PDB/CUUCGG/4v88-pdb-bundle2_1923_C_A_0_00002.pdb 
# Loading data/PDB/CUUCGG/4v88-pdb-bundle2_1923_C_A_0_00002.pdb 
# Loaded reference data/PDB/CUUCGG/2KOC_5_C_A_0_00001.pdb 
# Loaded target data/PDB/CUUCGG/5j3c-pdb-bundle1_1691_C_A_0_00000.pdb 
# Loading data/PDB/CUUCGG/5j3c-pdb-bundle1_1691_C_A_0_00000.pdb 
# Loaded reference data/PDB/CUUCGG/2KOC_5_C_A_0_00001.pdb 
# Loaded target data/PDB/CUUCGG/5uq8-pdb-bundle2_419_C_a_0_00000.pdb 
# Loading data/PDB/CUUCGG/5uq8-pdb-bundle2_419_C_a_0_00000.pdb 
# Loaded reference data/PDB/CUUCGG/2KOC_5_C_A_0_00001.pdb 
# Loaded target data/PDB/CUUCGG/5tga-pdb-bundle4_1923_C_J_0_00002.pdb 
# Loading data/PDB/CUUCGG/5tga-pdb-bundle4_1923_C_J_0_00002.pdb 
# Loaded reference data/PDB/CUUCGG/2KOC_5_C_A_0_00001.pdb 
# Loaded target data/PDB/CUUCGG/6cfl-pdb-bundle2_419_C_A_0_00000.pdb 
# Loading data/PDB/CUUCGG/6cfl-pdb-bundle2_419_C_A_0_00000.pdb 
# Loaded reference data/PDB/CUUCGG/2KOC_5_C_A_0_00001.pdb 
# Loaded target data/PDB/CUUCGG/2KU0_19_C_A_0_00003.pdb 
# Loading data/PDB/CUUCGG/2KU0_19_C_A_0_00003.pdb 
# Loaded reference data/PDB/CUUCGG/2KOC_5_C_A_0_00001.pdb 
# Loaded target data/PDB/CUUCGG/5fcj-pdb-bundle2_1923_C_A_0_00002.pdb 
# Loading data/PDB/CUUCGG/5fcj-pdb-bundle2_1923_C_A_0_00002.pdb 
# Loaded reference data/PDB/CUUCGG/2KOC_5_C_A_0_00001.pdb 
# Loaded target data/PDB/CUUCGG/5f8k-pdb-bundle3_1691_C_A_0_00000.pdb 
# Loading data/PDB/CUUCGG/5f8k-pdb-bundle3_1691_C_A_0_00000.pdb 
# Loaded reference data/PDB/CUUCGG/2KOC_5_C_A_0_00001.pdb 
# Loaded target data/PDB/CUUCGG/6c4i-pdb-bundle2_419_C_a_0_00001.pdb 
# Loading data/PDB/CUUCGG/6c4i-pdb-bundle2_419_C_a_0_00001.pdb 
# Loaded reference data/PDB/CUUCGG/2KOC_5_C_A_0_00001.pdb 
# Loaded target data/PDB/CUUCGG/4v57-pdb-bundle1_419_C_A_0_00001.pdb 
# Loading data/PDB/CUUCGG/4v57-pdb-bundle1_419_C_A_0_00001.pdb 
# Loaded reference data/PDB/CUUCGG/2KOC_5_C_A_0_00001.pdb 
# Loaded target data/PDB/CUUCGG/2GIP_12_C_A_0_00000.pdb 
# Loading data/PDB/CUUCGG/2GIP_12_C_A_0_00000.pdb 
# Loaded reference data/PDB/CUUCGG/2KOC_5_C_A_0_00001.pdb 
# Loaded target data/PDB/CUUCGG/6fkr-pdb-bundle2_419_C_A_0_00000.pdb 
# Loading data/PDB/CUUCGG/6fkr-pdb-bundle2_419_C_A_0_00000.pdb 
# Loaded reference data/PDB/CUUCGG/2KOC_5_C_A_0_00001.pdb 
# Loaded target data/PDB/CUUCGG/2GIP_12_C_A_0_00014.pdb 
# Loading data/PDB/CUUCGG/2GIP_12_C_A_0_00014.pdb 
# Loaded reference data/PDB/CUUCGG/2KOC_5_C_A_0_00001.pdb 
# Loaded target data/PDB/CUUCGG/5KH8_26_C_A_0_00014.pdb 
# Loading data/PDB/CUUCGG/5KH8_26_C_A_0_00014.pdb 
# Loaded reference data/PDB/CUUCGG/2KOC_5_C_A_0_00001.pdb 
# Loaded target data/PDB/CUUCGG/1MFY_14_C_A_0_00010.pdb 
# Loading data/PDB/CUUCGG/1MFY_14_C_A_0_00010.pdb 
# Loaded reference data/PDB/CUUCGG/2KOC_5_C_A_0_00001.pdb 
# Loaded target data/PDB/CUUCGG/4v7m-pdb-bundle2_1691_C_A_0_00000.pdb 
# Loading data/PDB/CUUCGG/4v7m-pdb-bundle2_1691_C_A_0_00000.pdb 
# Loaded reference data/PDB/CUUCGG/2KOC_5_C_A_0_00001.pdb 
# Loaded target data/PDB/CUUCGG/2GIO_13_C_A_0_00013.pdb 
# Loading data/PDB/CUUCGG/2GIO_13_C_A_0_00013.pdb 
# Loaded reference data/PDB/CUUCGG/2KOC_5_C_A_0_00001.pdb 
# Loaded target data/PDB/CUUCGG/1ZHO_15_C_H_0_00003.pdb 
# Loading data/PDB/CUUCGG/1ZHO_15_C_H_0_00003.pdb 
# Loaded reference data/PDB/CUUCGG/2KOC_5_C_A_0_00001.pdb 
# Loaded target data/PDB/CUUCGG/2GIO_13_C_A_0_00007.pdb 
# Loading data/PDB/CUUCGG/2GIO_13_C_A_0_00007.pdb 
# Loaded reference data/PDB/CUUCGG/2KOC_5_C_A_0_00001.pdb 
# Loaded target data/PDB/CUUCGG/5KH8_26_C_A_0_00000.pdb 
# Loading data/PDB/CUUCGG/5KH8_26_C_A_0_00000.pdb 
# Loaded reference data/PDB/CUUCGG/2KOC_5_C_A_0_00001.pdb 
# Loaded target data/PDB/CUUCGG/5it8-pdb-bundle1_1449_C_b_0_00007.pdb 
# Loading data/PDB/CUUCGG/5it8-pdb-bundle1_1449_C_b_0_00007.pdb 
# Loaded reference data/PDB/CUUCGG/2KOC_5_C_A_0_00001.pdb 
# Loaded target data/PDB/CUUCGG/1MFY_14_C_A_0_00004.pdb 
# Loading data/PDB/CUUCGG/1MFY_14_C_A_0_00004.pdb 
# Loaded reference data/PDB/CUUCGG/2KOC_5_C_A_0_00001.pdb 
# Loaded target data/PDB/CUUCGG/1JO7_14_C_A_0_00019.pdb 
# Loading data/PDB/CUUCGG/1JO7_14_C_A_0_00019.pdb 
# Loaded reference data/PDB/CUUCGG/2KOC_5_C_A_0_00001.pdb 
# Loaded target data/PDB/CUUCGG/1A3M_11_C_A_0_00019.pdb 
# Loading data/PDB/CUUCGG/1A3M_11_C_A_0_00019.pdb 
# Loaded reference data/PDB/CUUCGG/2KOC_5_C_A_0_00001.pdb 
# Loaded target data/PDB/CUUCGG/4ybb-pdb-bundle1_1449_C_A_0_00003.pdb 
# Loading data/PDB/CUUCGG/4ybb-pdb-bundle1_1449_C_A_0_00003.pdb 
# Loaded reference data/PDB/CUUCGG/2KOC_5_C_A_0_00001.pdb 
# Loaded target data/PDB/CUUCGG/1VQK_1769_C_0_0_00000.pdb 
# Loading data/PDB/CUUCGG/1VQK_1769_C_0_0_00000.pdb 
# Loaded reference data/PDB/CUUCGG/2KOC_5_C_A_0_00001.pdb 
# Loaded target data/PDB/CUUCGG/4v8a-pdb-bundle2_1691_C_A_0_00000.pdb 
# Loading data/PDB/CUUCGG/4v8a-pdb-bundle2_1691_C_A_0_00000.pdb 
# Loaded reference data/PDB/CUUCGG/2KOC_5_C_A_0_00001.pdb 
# Loaded target data/PDB/CUUCGG/4v64-pdb-bundle2_1691_C_A_0_00000.pdb 
# Loading data/PDB/CUUCGG/4v64-pdb-bundle2_1691_C_A_0_00000.pdb 
# Loaded reference data/PDB/CUUCGG/2KOC_5_C_A_0_00001.pdb 
# Loaded target data/PDB/CUUCGG/4v9c-pdb-bundle3_1449_C_A_0_00003.pdb 
# Loading data/PDB/CUUCGG/4v9c-pdb-bundle3_1449_C_A_0_00003.pdb 
# Loaded reference data/PDB/CUUCGG/2KOC_5_C_A_0_00001.pdb 
# Loaded target data/PDB/CUUCGG/1BYJ_11_C_A_0_00016.pdb 
# Loading data/PDB/CUUCGG/1BYJ_11_C_A_0_00016.pdb 
# Loaded reference data/PDB/CUUCGG/2KOC_5_C_A_0_00001.pdb 
# Loaded target data/PDB/CUUCGG/3jce-pdb-bundle1_1449_C_a_0_00003.pdb 
# Loading data/PDB/CUUCGG/3jce-pdb-bundle1_1449_C_a_0_00003.pdb 
# Loaded reference data/PDB/CUUCGG/2KOC_5_C_A_0_00001.pdb 
# Loaded target data/PDB/CUUCGG/1BYJ_11_C_A_0_00002.pdb 
# Loading data/PDB/CUUCGG/1BYJ_11_C_A_0_00002.pdb 
# Loaded reference data/PDB/CUUCGG/2KOC_5_C_A_0_00001.pdb 
# Loaded target data/PDB/CUUCGG/5j30-pdb-bundle3_1691_C_A_0_00000.pdb 
# Loading data/PDB/CUUCGG/5j30-pdb-bundle3_1691_C_A_0_00000.pdb 
# Loaded reference data/PDB/CUUCGG/2KOC_5_C_A_0_00001.pdb 
# Loaded target data/PDB/CUUCGG/2LKR_63_C_A_0_00001.pdb 
# Loading data/PDB/CUUCGG/2LKR_63_C_A_0_00001.pdb 
# Loaded reference data/PDB/CUUCGG/2KOC_5_C_A_0_00001.pdb 
# Loaded target data/PDB/CUUCGG/4v64-pdb-bundle1_419_C_A_0_00001.pdb 
# Loading data/PDB/CUUCGG/4v64-pdb-bundle1_419_C_A_0_00001.pdb 
# Loaded reference data/PDB/CUUCGG/2KOC_5_C_A_0_00001.pdb 
# Loaded target data/PDB/CUUCGG/4v5e-pdb-bundle2_1691_C_A_0_00000.pdb 
# Loading data/PDB/CUUCGG/4v5e-pdb-bundle2_1691_C_A_0_00000.pdb 
# Loaded reference data/PDB/CUUCGG/2KOC_5_C_A_0_00001.pdb 
# Loaded target data/PDB/CUUCGG/4v8i-pdb-bundle3_419_C_A_0_00000.pdb 
# Loading data/PDB/CUUCGG/4v8i-pdb-bundle3_419_C_A_0_00000.pdb 
# Loaded reference data/PDB/CUUCGG/2KOC_5_C_A_0_00001.pdb 
# Loaded target data/PDB/CUUCGG/2NCI_14_C_A_0_00009.pdb 
# Loading data/PDB/CUUCGG/2NCI_14_C_A_0_00009.pdb 
# Loaded reference data/PDB/CUUCGG/2KOC_5_C_A_0_00001.pdb 
# Loaded target data/PDB/CUUCGG/1TLR_10_C_A_0_00015.pdb 
# Loading data/PDB/CUUCGG/1TLR_10_C_A_0_00015.pdb 
# Loaded reference data/PDB/CUUCGG/2KOC_5_C_A_0_00001.pdb 
# Loaded target data/PDB/CUUCGG/5uq7-pdb-bundle1_1691_C_A_0_00000.pdb 
# Loading data/PDB/CUUCGG/5uq7-pdb-bundle1_1691_C_A_0_00000.pdb 
# Loaded reference data/PDB/CUUCGG/2KOC_5_C_A_0_00001.pdb 
# Loaded target data/PDB/CUUCGG/1TLR_10_C_A_0_00001.pdb 
# Loading data/PDB/CUUCGG/1TLR_10_C_A_0_00001.pdb 
# Loaded reference data/PDB/CUUCGG/2KOC_5_C_A_0_00001.pdb 
# Loaded target data/PDB/CUUCGG/4v9n-pdb-bundle2_1691_C_A_0_00000.pdb 
# Loading data/PDB/CUUCGG/4v9n-pdb-bundle2_1691_C_A_0_00000.pdb 
# Loaded reference data/PDB/CUUCGG/2KOC_5_C_A_0_00001.pdb 
# Loaded target data/PDB/CUUCGG/2N6X_20_C_A_0_00002.pdb 
# Loading data/PDB/CUUCGG/2N6X_20_C_A_0_00002.pdb 
# Loaded reference data/PDB/CUUCGG/2KOC_5_C_A_0_00001.pdb 
# Loaded target data/PDB/CUUCGG/4v5k-pdb-bundle4_1691_C_A_0_00000.pdb 
# Loading data/PDB/CUUCGG/4v5k-pdb-bundle4_1691_C_A_0_00000.pdb 
# Loaded reference data/PDB/CUUCGG/2KOC_5_C_A_0_00001.pdb 
# Loaded target data/PDB/CUUCGG/4u3u-pdb-bundle2_1923_C_A_0_00002.pdb 
# Loading data/PDB/CUUCGG/4u3u-pdb-bundle2_1923_C_A_0_00002.pdb 
# Loaded reference data/PDB/CUUCGG/2KOC_5_C_A_0_00001.pdb 
# Loaded target data/PDB/CUUCGG/5KQE_19_C_A_0_00001.pdb 
# Loading data/PDB/CUUCGG/5KQE_19_C_A_0_00001.pdb 
# Loaded reference data/PDB/CUUCGG/2KOC_5_C_A_0_00001.pdb 
# Loaded target data/PDB/CUUCGG/5iqr-pdb-bundle2_1691_C_1_0_00000.pdb 
# Loading data/PDB/CUUCGG/5iqr-pdb-bundle2_1691_C_1_0_00000.pdb 
# Loaded reference data/PDB/CUUCGG/2KOC_5_C_A_0_00001.pdb 
# Loaded target data/PDB/CUUCGG/1FYO_11_C_A_0_00001.pdb 
# Loading data/PDB/CUUCGG/1FYO_11_C_A_0_00001.pdb 
# Loaded reference data/PDB/CUUCGG/2KOC_5_C_A_0_00001.pdb 
# Loaded target data/PDB/CUUCGG/1FYO_11_C_A_0_00015.pdb 
# Loading data/PDB/CUUCGG/1FYO_11_C_A_0_00015.pdb 
# Loaded reference data/PDB/CUUCGG/2KOC_5_C_A_0_00001.pdb 
# Loaded target data/PDB/CUUCGG/4wu1-pdb-bundle4_1691_C_A_0_00000.pdb 
# Loading data/PDB/CUUCGG/4wu1-pdb-bundle4_1691_C_A_0_00000.pdb 
# Loaded reference data/PDB/CUUCGG/2KOC_5_C_A_0_00001.pdb 
# Loaded target data/PDB/CUUCGG/5jc9-pdb-bundle3_1691_C_c_0_00000.pdb 
# Loading data/PDB/CUUCGG/5jc9-pdb-bundle3_1691_C_c_0_00000.pdb 
# Loaded reference data/PDB/CUUCGG/2KOC_5_C_A_0_00001.pdb 
# Loaded target data/PDB/CUUCGG/5KQE_19_C_A_0_00015.pdb 
# Loading data/PDB/CUUCGG/5KQE_19_C_A_0_00015.pdb 
# Loaded reference data/PDB/CUUCGG/2KOC_5_C_A_0_00001.pdb 
# Loaded target data/PDB/CUUCGG/4zer-pdb-bundle2_419_C_A_0_00000.pdb 
# Loading data/PDB/CUUCGG/4zer-pdb-bundle2_419_C_A_0_00000.pdb 
# Loaded reference data/PDB/CUUCGG/2KOC_5_C_A_0_00001.pdb 
# Loaded target data/PDB/CUUCGG/6DMD_57_C_B_0_00001.pdb 
# Loading data/PDB/CUUCGG/6DMD_57_C_B_0_00001.pdb 
# Loaded reference data/PDB/CUUCGG/2KOC_5_C_A_0_00001.pdb 
# Loaded target data/PDB/CUUCGG/1FYO_11_C_A_0_00014.pdb 
# Loading data/PDB/CUUCGG/1FYO_11_C_A_0_00014.pdb 
# Loaded reference data/PDB/CUUCGG/2KOC_5_C_A_0_00001.pdb 
# Loaded target data/PDB/CUUCGG/5KQE_19_C_A_0_00014.pdb 
# Loading data/PDB/CUUCGG/5KQE_19_C_A_0_00014.pdb 
# Loaded reference data/PDB/CUUCGG/2KOC_5_C_A_0_00001.pdb 
# Loaded target data/PDB/CUUCGG/1VQN_1769_C_0_0_00000.pdb 
# Loading data/PDB/CUUCGG/1VQN_1769_C_0_0_00000.pdb 
# Loaded reference data/PDB/CUUCGG/2KOC_5_C_A_0_00001.pdb 
# Loaded target data/PDB/CUUCGG/5KQE_19_C_A_0_00000.pdb 
# Loading data/PDB/CUUCGG/5KQE_19_C_A_0_00000.pdb 
# Loaded reference data/PDB/CUUCGG/2KOC_5_C_A_0_00001.pdb 
# Loaded target data/PDB/CUUCGG/1FYO_11_C_A_0_00000.pdb 
# Loading data/PDB/CUUCGG/1FYO_11_C_A_0_00000.pdb 
# Loaded reference data/PDB/CUUCGG/2KOC_5_C_A_0_00001.pdb 
# Loaded target data/PDB/CUUCGG/6DZP_1909_C_A_0_00000.pdb 
# Loading data/PDB/CUUCGG/6DZP_1909_C_A_0_00000.pdb 
# Loaded reference data/PDB/CUUCGG/2KOC_5_C_A_0_00001.pdb 
# Loaded target data/PDB/CUUCGG/2N6X_20_C_A_0_00003.pdb 
# Loading data/PDB/CUUCGG/2N6X_20_C_A_0_00003.pdb 
# Loaded reference data/PDB/CUUCGG/2KOC_5_C_A_0_00001.pdb 
# Loaded target data/PDB/CUUCGG/4v5c-pdb-bundle3_419_C_A_0_00000.pdb 
# Loading data/PDB/CUUCGG/4v5c-pdb-bundle3_419_C_A_0_00000.pdb 
# Loaded reference data/PDB/CUUCGG/2KOC_5_C_A_0_00001.pdb 
# Loaded target data/PDB/CUUCGG/4v8f-pdb-bundle2_419_C_Z_0_00001.pdb 
# Loading data/PDB/CUUCGG/4v8f-pdb-bundle2_419_C_Z_0_00001.pdb 
# Loaded reference data/PDB/CUUCGG/2KOC_5_C_A_0_00001.pdb 
# Loaded target data/PDB/CUUCGG/1TLR_10_C_A_0_00000.pdb 
# Loading data/PDB/CUUCGG/1TLR_10_C_A_0_00000.pdb 
# Loaded reference data/PDB/CUUCGG/2KOC_5_C_A_0_00001.pdb 
# Loaded target data/PDB/CUUCGG/1TLR_10_C_A_0_00014.pdb 
# Loading data/PDB/CUUCGG/1TLR_10_C_A_0_00014.pdb 
# Loaded reference data/PDB/CUUCGG/2KOC_5_C_A_0_00001.pdb 
# Loaded target data/PDB/CUUCGG/5LYS_48_C_A_0_00000.pdb 
# Loading data/PDB/CUUCGG/5LYS_48_C_A_0_00000.pdb 
# Loaded reference data/PDB/CUUCGG/2KOC_5_C_A_0_00001.pdb 
# Loaded target data/PDB/CUUCGG/4GKK_414_C_A_0_00000.pdb 
# Loading data/PDB/CUUCGG/4GKK_414_C_A_0_00000.pdb 
# Loaded reference data/PDB/CUUCGG/2KOC_5_C_A_0_00001.pdb 
# Loaded target data/PDB/CUUCGG/6gsj-pdb-bundle2_1691_C_A_0_00000.pdb 
# Loading data/PDB/CUUCGG/6gsj-pdb-bundle2_1691_C_A_0_00000.pdb 
# Loaded reference data/PDB/CUUCGG/2KOC_5_C_A_0_00001.pdb 
# Loaded target data/PDB/CUUCGG/2NCI_14_C_A_0_00008.pdb 
# Loading data/PDB/CUUCGG/2NCI_14_C_A_0_00008.pdb 
# Loaded reference data/PDB/CUUCGG/2KOC_5_C_A_0_00001.pdb 
# Loaded target data/PDB/CUUCGG/4v50-pdb-bundle3_1028_C_A_0_00002.pdb 
# Loading data/PDB/CUUCGG/4v50-pdb-bundle3_1028_C_A_0_00002.pdb 
# Loaded reference data/PDB/CUUCGG/2KOC_5_C_A_0_00001.pdb 
# Loaded target data/PDB/CUUCGG/5ndv-pdb-bundle3_1923_C_B_0_00002.pdb 
# Loading data/PDB/CUUCGG/5ndv-pdb-bundle3_1923_C_B_0_00002.pdb 
# Loaded reference data/PDB/CUUCGG/2KOC_5_C_A_0_00001.pdb 
# Loaded target data/PDB/CUUCGG/4lt8-pdb-bundle2_1691_C_A_0_00000.pdb 
# Loading data/PDB/CUUCGG/4lt8-pdb-bundle2_1691_C_A_0_00000.pdb 
# Loaded reference data/PDB/CUUCGG/2KOC_5_C_A_0_00001.pdb 
# Loaded target data/PDB/CUUCGG/5we6-pdb-bundle2_419_C_a_0_00001.pdb 
# Loading data/PDB/CUUCGG/5we6-pdb-bundle2_419_C_a_0_00001.pdb 
# Loaded reference data/PDB/CUUCGG/2KOC_5_C_A_0_00001.pdb 
# Loaded target data/PDB/CUUCGG/5j8a-pdb-bundle1_419_C_A_0_00001.pdb 
# Loading data/PDB/CUUCGG/5j8a-pdb-bundle1_419_C_A_0_00001.pdb 
# Loaded reference data/PDB/CUUCGG/2KOC_5_C_A_0_00001.pdb 
# Loaded target data/PDB/CUUCGG/1vy4-pdb-bundle1_419_C_A_0_00000.pdb 
# Loading data/PDB/CUUCGG/1vy4-pdb-bundle1_419_C_A_0_00000.pdb 
# Loaded reference data/PDB/CUUCGG/2KOC_5_C_A_0_00001.pdb 
# Loaded target data/PDB/CUUCGG/2LKR_63_C_A_0_00000.pdb 
# Loading data/PDB/CUUCGG/2LKR_63_C_A_0_00000.pdb 
# Loaded reference data/PDB/CUUCGG/2KOC_5_C_A_0_00001.pdb 
# Loaded target data/PDB/CUUCGG/1F7Y_32_C_B_0_00001.pdb 
# Loading data/PDB/CUUCGG/1F7Y_32_C_B_0_00001.pdb 
# Loaded reference data/PDB/CUUCGG/2KOC_5_C_A_0_00001.pdb 
# Loaded target data/PDB/CUUCGG/4v5r-pdb-bundle2_1691_C_A_0_00000.pdb 
# Loading data/PDB/CUUCGG/4v5r-pdb-bundle2_1691_C_A_0_00000.pdb 
# Loaded reference data/PDB/CUUCGG/2KOC_5_C_A_0_00001.pdb 
# Loaded target data/PDB/CUUCGG/1BYJ_11_C_A_0_00003.pdb 
# Loading data/PDB/CUUCGG/1BYJ_11_C_A_0_00003.pdb 
# Loaded reference data/PDB/CUUCGG/2KOC_5_C_A_0_00001.pdb 
# Loaded target data/PDB/CUUCGG/4wzd-pdb-bundle1_419_C_A_0_00000.pdb 
# Loading data/PDB/CUUCGG/4wzd-pdb-bundle1_419_C_A_0_00000.pdb 
# Loaded reference data/PDB/CUUCGG/2KOC_5_C_A_0_00001.pdb 
# Loaded target data/PDB/CUUCGG/1JZX_1708_C_A_0_00000.pdb 
# Loading data/PDB/CUUCGG/1JZX_1708_C_A_0_00000.pdb 
# Loaded reference data/PDB/CUUCGG/2KOC_5_C_A_0_00001.pdb 
# Loaded target data/PDB/CUUCGG/4v9k-pdb-bundle3_419_C_A_0_00000.pdb 
# Loading data/PDB/CUUCGG/4v9k-pdb-bundle3_419_C_A_0_00000.pdb 
# Loaded reference data/PDB/CUUCGG/2KOC_5_C_A_0_00001.pdb 
# Loaded target data/PDB/CUUCGG/1BYJ_11_C_A_0_00017.pdb 
# Loading data/PDB/CUUCGG/1BYJ_11_C_A_0_00017.pdb 
# Loaded reference data/PDB/CUUCGG/2KOC_5_C_A_0_00001.pdb 
# Loaded target data/PDB/CUUCGG/1K9M_1769_C_A_0_00000.pdb 
# Loading data/PDB/CUUCGG/1K9M_1769_C_A_0_00000.pdb 
# Loaded reference data/PDB/CUUCGG/2KOC_5_C_A_0_00001.pdb 
# Loaded target data/PDB/CUUCGG/5j7l-pdb-bundle1_1449_C_b_0_00007.pdb 
# Loading data/PDB/CUUCGG/5j7l-pdb-bundle1_1449_C_b_0_00007.pdb 
# Loaded reference data/PDB/CUUCGG/2KOC_5_C_A_0_00001.pdb 
# Loaded target data/PDB/CUUCGG/1A3M_11_C_A_0_00018.pdb 
# Loading data/PDB/CUUCGG/1A3M_11_C_A_0_00018.pdb 
# Loaded reference data/PDB/CUUCGG/2KOC_5_C_A_0_00001.pdb 
# Loaded target data/PDB/CUUCGG/4lt8-pdb-bundle1_419_C_A_0_00000.pdb 
# Loading data/PDB/CUUCGG/4lt8-pdb-bundle1_419_C_A_0_00000.pdb 
# Loaded reference data/PDB/CUUCGG/2KOC_5_C_A_0_00001.pdb 
# Loaded target data/PDB/CUUCGG/2GIO_13_C_A_0_00006.pdb 
# Loading data/PDB/CUUCGG/2GIO_13_C_A_0_00006.pdb 
# Loaded reference data/PDB/CUUCGG/2KOC_5_C_A_0_00001.pdb 
# Loaded target data/PDB/CUUCGG/1JO7_14_C_A_0_00018.pdb 
# Loading data/PDB/CUUCGG/1JO7_14_C_A_0_00018.pdb 
# Loaded reference data/PDB/CUUCGG/2KOC_5_C_A_0_00001.pdb 
# Loaded target data/PDB/CUUCGG/1MFY_14_C_A_0_00005.pdb 
# Loading data/PDB/CUUCGG/1MFY_14_C_A_0_00005.pdb 
# Loaded reference data/PDB/CUUCGG/2KOC_5_C_A_0_00001.pdb 
# Loaded target data/PDB/CUUCGG/5KH8_26_C_A_0_00001.pdb 
# Loading data/PDB/CUUCGG/5KH8_26_C_A_0_00001.pdb 
# Loaded reference data/PDB/CUUCGG/2KOC_5_C_A_0_00001.pdb 
# Loaded target data/PDB/CUUCGG/1MFY_14_C_A_0_00011.pdb 
# Loading data/PDB/CUUCGG/1MFY_14_C_A_0_00011.pdb 
# Loaded reference data/PDB/CUUCGG/2KOC_5_C_A_0_00001.pdb 
# Loaded target data/PDB/CUUCGG/5KH8_26_C_A_0_00015.pdb 
# Loading data/PDB/CUUCGG/5KH8_26_C_A_0_00015.pdb 
# Loaded reference data/PDB/CUUCGG/2KOC_5_C_A_0_00001.pdb 
# Loaded target data/PDB/CUUCGG/5lzd-pdb-bundle2_1691_C_A_0_00000.pdb 
# Loading data/PDB/CUUCGG/5lzd-pdb-bundle2_1691_C_A_0_00000.pdb 
# Loaded reference data/PDB/CUUCGG/2KOC_5_C_A_0_00001.pdb 
# Loaded target data/PDB/CUUCGG/2GIO_13_C_A_0_00012.pdb 
# Loading data/PDB/CUUCGG/2GIO_13_C_A_0_00012.pdb 
# Loaded reference data/PDB/CUUCGG/2KOC_5_C_A_0_00001.pdb 
# Loaded target data/PDB/CUUCGG/2GIP_12_C_A_0_00015.pdb 
# Loading data/PDB/CUUCGG/2GIP_12_C_A_0_00015.pdb 
# Loaded reference data/PDB/CUUCGG/2KOC_5_C_A_0_00001.pdb 
# Loaded target data/PDB/CUUCGG/3CPW_1769_C_0_0_00000.pdb 
# Loading data/PDB/CUUCGG/3CPW_1769_C_0_0_00000.pdb 
# Loaded reference data/PDB/CUUCGG/2KOC_5_C_A_0_00001.pdb 
# Loaded target data/PDB/CUUCGG/6DLT_27_C_A_0_00000.pdb 
# Loading data/PDB/CUUCGG/6DLT_27_C_A_0_00000.pdb 
# Loaded reference data/PDB/CUUCGG/2KOC_5_C_A_0_00001.pdb 
# Loaded target data/PDB/CUUCGG/2GIP_12_C_A_0_00001.pdb 
# Loading data/PDB/CUUCGG/2GIP_12_C_A_0_00001.pdb 
# Loaded reference data/PDB/CUUCGG/2KOC_5_C_A_0_00001.pdb 
# Loaded target data/PDB/CUUCGG/4v7z-pdb-bundle2_1691_C_A_0_00000.pdb 
# Loading data/PDB/CUUCGG/4v7z-pdb-bundle2_1691_C_A_0_00000.pdb 
# Loaded reference data/PDB/CUUCGG/2KOC_5_C_A_0_00001.pdb 
# Loaded target data/PDB/CUUCGG/6ha8-pdb-bundle1_1736_C_A_0_00001.pdb 
# Loading data/PDB/CUUCGG/6ha8-pdb-bundle1_1736_C_A_0_00001.pdb 
# Loaded reference data/PDB/CUUCGG/2KOC_5_C_A_0_00001.pdb 
# Loaded target data/PDB/CUUCGG/2KU0_19_C_A_0_00002.pdb 
# Loading data/PDB/CUUCGG/2KU0_19_C_A_0_00002.pdb 
# Loaded reference data/PDB/CUUCGG/2KOC_5_C_A_0_00001.pdb 
# Loaded target data/PDB/CUUCGG/4v4q-pdb-bundle3_207_C_A_0_00000.pdb 
# Loading data/PDB/CUUCGG/4v4q-pdb-bundle3_207_C_A_0_00000.pdb 
# Loaded reference data/PDB/CUUCGG/2KOC_5_C_A_0_00001.pdb 
# Loaded target data/PDB/CUUCGG/5LYU_48_C_B_0_00001.pdb 
# Loading data/PDB/CUUCGG/5LYU_48_C_B_0_00001.pdb 
# Loaded reference data/PDB/CUUCGG/2KOC_5_C_A_0_00001.pdb 
# Loaded target data/PDB/CUUCGG/5dge-pdb-bundle2_1923_C_A_0_00002.pdb 
# Loading data/PDB/CUUCGG/5dge-pdb-bundle2_1923_C_A_0_00002.pdb 
# Loaded reference data/PDB/CUUCGG/2KOC_5_C_A_0_00001.pdb 
# Loaded target data/PDB/CUUCGG/4w2e-pdb-bundle1_1691_C_A_0_00000.pdb 
# Loading data/PDB/CUUCGG/4w2e-pdb-bundle1_1691_C_A_0_00000.pdb 
# Loaded reference data/PDB/CUUCGG/2KOC_5_C_A_0_00001.pdb 
# Loaded target data/PDB/CUUCGG/4u51-pdb-bundle4_1923_C_I_0_00002.pdb 
# Loading data/PDB/CUUCGG/4u51-pdb-bundle4_1923_C_I_0_00002.pdb 
# Loaded reference data/PDB/CUUCGG/2KOC_5_C_A_0_00001.pdb 
# Loaded target data/PDB/CUUCGG/4u24-pdb-bundle2_1691_C_A_0_00000.pdb 
# Loading data/PDB/CUUCGG/4u24-pdb-bundle2_1691_C_A_0_00000.pdb 
# Loaded reference data/PDB/CUUCGG/2KOC_5_C_A_0_00001.pdb 
# Loaded target data/PDB/CUUCGG/2L3E_18_C_A_0_00018.pdb 
# Loading data/PDB/CUUCGG/2L3E_18_C_A_0_00018.pdb 
# Loaded reference data/PDB/CUUCGG/2KOC_5_C_A_0_00001.pdb 
# Loaded target data/PDB/CUUCGG/5ngm-pdb-bundle1_427_C_A_0_00001.pdb 
# Loading data/PDB/CUUCGG/5ngm-pdb-bundle1_427_C_A_0_00001.pdb 
# Loaded reference data/PDB/CUUCGG/2KOC_5_C_A_0_00001.pdb 
# Loaded target data/PDB/CUUCGG/4v87-pdb-bundle2_419_C_Z_0_00001.pdb 
# Loading data/PDB/CUUCGG/4v87-pdb-bundle2_419_C_Z_0_00001.pdb 
# Loaded reference data/PDB/CUUCGG/2KOC_5_C_A_0_00001.pdb 
# Loaded target data/PDB/CUUCGG/5el7-pdb-bundle4_1691_C_A_0_00000.pdb 
# Loading data/PDB/CUUCGG/5el7-pdb-bundle4_1691_C_A_0_00000.pdb 
# Loaded reference data/PDB/CUUCGG/2KOC_5_C_A_0_00001.pdb 
# Loaded target data/PDB/CUUCGG/5wit-pdb-bundle2_419_C_A_0_00000.pdb 
# Loading data/PDB/CUUCGG/5wit-pdb-bundle2_419_C_A_0_00000.pdb 
# Loaded reference data/PDB/CUUCGG/2KOC_5_C_A_0_00001.pdb 
# Loaded target data/PDB/CUUCGG/4v8x-pdb-bundle4_1691_C_A_0_00000.pdb 
# Loading data/PDB/CUUCGG/4v8x-pdb-bundle4_1691_C_A_0_00000.pdb 
# Loaded reference data/PDB/CUUCGG/2KOC_5_C_A_0_00001.pdb 
# Loaded target data/PDB/CUUCGG/6i7v-pdb-bundle2_1691_C_A_0_00000.pdb 
# Loading data/PDB/CUUCGG/6i7v-pdb-bundle2_1691_C_A_0_00000.pdb 
# Loaded reference data/PDB/CUUCGG/2KOC_5_C_A_0_00001.pdb 
# Loaded target data/PDB/CUUCGG/2OGM_1708_C_0_0_00000.pdb 
# Loading data/PDB/CUUCGG/2OGM_1708_C_0_0_00000.pdb 
# Loaded reference data/PDB/CUUCGG/2KOC_5_C_A_0_00001.pdb 
# Loaded target data/PDB/CUUCGG/2LU0_28_C_A_0_00015.pdb 
# Loading data/PDB/CUUCGG/2LU0_28_C_A_0_00015.pdb 
# Loaded reference data/PDB/CUUCGG/2KOC_5_C_A_0_00001.pdb 
# Loaded target data/PDB/CUUCGG/2LU0_28_C_A_0_00001.pdb 
# Loading data/PDB/CUUCGG/2LU0_28_C_A_0_00001.pdb 
# Loaded reference data/PDB/CUUCGG/2KOC_5_C_A_0_00001.pdb 
# Loaded target data/PDB/CUUCGG/4v7t-pdb-bundle4_1691_C_A_0_00000.pdb 
# Loading data/PDB/CUUCGG/4v7t-pdb-bundle4_1691_C_A_0_00000.pdb 
# Loaded reference data/PDB/CUUCGG/2KOC_5_C_A_0_00001.pdb 
# Loaded target data/PDB/CUUCGG/5zeb-pdb-bundle1_419_C_a_0_00000.pdb 
# Loading data/PDB/CUUCGG/5zeb-pdb-bundle1_419_C_a_0_00000.pdb 
# Loaded reference data/PDB/CUUCGG/2KOC_5_C_A_0_00001.pdb 
# Loaded target data/PDB/CUUCGG/2LU0_28_C_A_0_00017.pdb 
# Loading data/PDB/CUUCGG/2LU0_28_C_A_0_00017.pdb 
# Loaded reference data/PDB/CUUCGG/2KOC_5_C_A_0_00001.pdb 
# Loaded target data/PDB/CUUCGG/2LU0_28_C_A_0_00003.pdb 
# Loading data/PDB/CUUCGG/2LU0_28_C_A_0_00003.pdb 
# Loaded reference data/PDB/CUUCGG/2KOC_5_C_A_0_00001.pdb 
# Loaded target data/PDB/CUUCGG/4u3m-pdb-bundle4_1923_C_I_0_00002.pdb 
# Loading data/PDB/CUUCGG/4u3m-pdb-bundle4_1923_C_I_0_00002.pdb 
# Loaded reference data/PDB/CUUCGG/2KOC_5_C_A_0_00001.pdb 
# Loaded target data/PDB/CUUCGG/4v9o-pdb-bundle5_1691_C_A_0_00000.pdb 
# Loading data/PDB/CUUCGG/4v9o-pdb-bundle5_1691_C_A_0_00000.pdb 
# Loaded reference data/PDB/CUUCGG/2KOC_5_C_A_0_00001.pdb 
# Loaded target data/PDB/CUUCGG/1NJP_1708_C_0_0_00000.pdb 
# Loading data/PDB/CUUCGG/1NJP_1708_C_0_0_00000.pdb 
# Loaded reference data/PDB/CUUCGG/2KOC_5_C_A_0_00001.pdb 
# Loaded target data/PDB/CUUCGG/5j7l-pdb-bundle3_1691_C_d_0_00000.pdb 
# Loading data/PDB/CUUCGG/5j7l-pdb-bundle3_1691_C_d_0_00000.pdb 
# Loaded reference data/PDB/CUUCGG/2KOC_5_C_A_0_00001.pdb 
# Loaded target data/PDB/CUUCGG/4u3n-pdb-bundle3_230_C_d_0_00000.pdb 
# Loading data/PDB/CUUCGG/4u3n-pdb-bundle3_230_C_d_0_00000.pdb 
# Loaded reference data/PDB/CUUCGG/2KOC_5_C_A_0_00001.pdb 
# Loaded target data/PDB/CUUCGG/4DV6_419_C_A_0_00000.pdb 
# Loading data/PDB/CUUCGG/4DV6_419_C_A_0_00000.pdb 
# Loaded reference data/PDB/CUUCGG/2KOC_5_C_A_0_00001.pdb 
# Loaded target data/PDB/CUUCGG/2N3R_37_C_A_0_00009.pdb 
# Loading data/PDB/CUUCGG/2N3R_37_C_A_0_00009.pdb 
# Loaded reference data/PDB/CUUCGG/2KOC_5_C_A_0_00001.pdb 
# Loaded target data/PDB/CUUCGG/4lsk-pdb-bundle2_1691_C_A_0_00000.pdb 
# Loading data/PDB/CUUCGG/4lsk-pdb-bundle2_1691_C_A_0_00000.pdb 
# Loaded reference data/PDB/CUUCGG/2KOC_5_C_A_0_00001.pdb 
# Loaded target data/PDB/CUUCGG/4wpo-pdb-bundle3_1691_C_A_0_00000.pdb 
# Loading data/PDB/CUUCGG/4wpo-pdb-bundle3_1691_C_A_0_00000.pdb 
# Loaded reference data/PDB/CUUCGG/2KOC_5_C_A_0_00001.pdb 
# Loaded target data/PDB/CUUCGG/4wqr-pdb-bundle4_1691_C_A_0_00000.pdb 
# Loading data/PDB/CUUCGG/4wqr-pdb-bundle4_1691_C_A_0_00000.pdb 
# Loaded reference data/PDB/CUUCGG/2KOC_5_C_A_0_00001.pdb 
# Loaded target data/PDB/CUUCGG/5jup-pdb-bundle1_230_C_A_0_00000.pdb 
# Loading data/PDB/CUUCGG/5jup-pdb-bundle1_230_C_A_0_00000.pdb 
# Loaded reference data/PDB/CUUCGG/2KOC_5_C_A_0_00001.pdb 
# Loaded target data/PDB/CUUCGG/1J5A_1708_C_A_0_00000.pdb 
# Loading data/PDB/CUUCGG/1J5A_1708_C_A_0_00000.pdb 
# Loaded reference data/PDB/CUUCGG/2KOC_5_C_A_0_00001.pdb 
# Loaded target data/PDB/CUUCGG/4tue-pdb-bundle4_1691_C_A_0_00000.pdb 
# Loading data/PDB/CUUCGG/4tue-pdb-bundle4_1691_C_A_0_00000.pdb 
# Loaded reference data/PDB/CUUCGG/2KOC_5_C_A_0_00001.pdb 
# Loaded target data/PDB/CUUCGG/4v64-pdb-bundle3_419_C_A_0_00001.pdb 
# Loading data/PDB/CUUCGG/4v64-pdb-bundle3_419_C_A_0_00001.pdb 
# Loaded reference data/PDB/CUUCGG/2KOC_5_C_A_0_00001.pdb 
# Loaded target data/PDB/CUUCGG/2KU0_19_C_A_0_00000.pdb 
# Loading data/PDB/CUUCGG/2KU0_19_C_A_0_00000.pdb 
# Loaded reference data/PDB/CUUCGG/2KOC_5_C_A_0_00001.pdb 
# Loaded target data/PDB/CUUCGG/3jbv-pdb-bundle2_1691_C_b_0_00000.pdb 
# Loading data/PDB/CUUCGG/3jbv-pdb-bundle2_1691_C_b_0_00000.pdb 
# Loaded reference data/PDB/CUUCGG/2KOC_5_C_A_0_00001.pdb 
# Loaded target data/PDB/CUUCGG/2GIP_12_C_A_0_00017.pdb 
# Loading data/PDB/CUUCGG/2GIP_12_C_A_0_00017.pdb 
# Loaded reference data/PDB/CUUCGG/2KOC_5_C_A_0_00001.pdb 
# Loaded target data/PDB/CUUCGG/4v9d-pdb-bundle1_1449_C_Y_0_00007.pdb 
# Loading data/PDB/CUUCGG/4v9d-pdb-bundle1_1449_C_Y_0_00007.pdb 
# Loaded reference data/PDB/CUUCGG/2KOC_5_C_A_0_00001.pdb 
# Loaded target data/PDB/CUUCGG/2GIP_12_C_A_0_00003.pdb 
# Loading data/PDB/CUUCGG/2GIP_12_C_A_0_00003.pdb 
# Loaded reference data/PDB/CUUCGG/2KOC_5_C_A_0_00001.pdb 
# Loaded target data/PDB/CUUCGG/5vp2-pdb-bundle1_1738_C_A_0_00000.pdb 
# Loading data/PDB/CUUCGG/5vp2-pdb-bundle1_1738_C_A_0_00000.pdb 
# Loaded reference data/PDB/CUUCGG/2KOC_5_C_A_0_00001.pdb 
# Loaded target data/PDB/CUUCGG/5KH8_26_C_A_0_00003.pdb 
# Loading data/PDB/CUUCGG/5KH8_26_C_A_0_00003.pdb 
# Loaded reference data/PDB/CUUCGG/2KOC_5_C_A_0_00001.pdb 
# Loaded target data/PDB/CUUCGG/1MFY_14_C_A_0_00007.pdb 
# Loading data/PDB/CUUCGG/1MFY_14_C_A_0_00007.pdb 
# Loaded reference data/PDB/CUUCGG/2KOC_5_C_A_0_00001.pdb 
# Loaded target data/PDB/CUUCGG/4u20-pdb-bundle2_1691_C_A_0_00000.pdb 
# Loading data/PDB/CUUCGG/4u20-pdb-bundle2_1691_C_A_0_00000.pdb 
# Loaded reference data/PDB/CUUCGG/2KOC_5_C_A_0_00001.pdb 
# Loaded target data/PDB/CUUCGG/2GIO_13_C_A_0_00004.pdb 
# Loading data/PDB/CUUCGG/2GIO_13_C_A_0_00004.pdb 
# Loaded reference data/PDB/CUUCGG/2KOC_5_C_A_0_00001.pdb 
# Loaded target data/PDB/CUUCGG/4u55-pdb-bundle4_1923_C_I_0_00002.pdb 
# Loading data/PDB/CUUCGG/4u55-pdb-bundle4_1923_C_I_0_00002.pdb 
# Loaded reference data/PDB/CUUCGG/2KOC_5_C_A_0_00001.pdb 
# Loaded target data/PDB/CUUCGG/2GIO_13_C_A_0_00010.pdb 
# Loading data/PDB/CUUCGG/2GIO_13_C_A_0_00010.pdb 
# Loaded reference data/PDB/CUUCGG/2KOC_5_C_A_0_00001.pdb 
# Loaded target data/PDB/CUUCGG/4lnt-pdb-bundle2_1691_C_A_0_00000.pdb 
# Loading data/PDB/CUUCGG/4lnt-pdb-bundle2_1691_C_A_0_00000.pdb 
# Loaded reference data/PDB/CUUCGG/2KOC_5_C_A_0_00001.pdb 
# Loaded target data/PDB/CUUCGG/5KH8_26_C_A_0_00017.pdb 
# Loading data/PDB/CUUCGG/5KH8_26_C_A_0_00017.pdb 
# Loaded reference data/PDB/CUUCGG/2KOC_5_C_A_0_00001.pdb 
# Loaded target data/PDB/CUUCGG/4v8i-pdb-bundle1_419_C_A_0_00000.pdb 
# Loading data/PDB/CUUCGG/4v8i-pdb-bundle1_419_C_A_0_00000.pdb 
# Loaded reference data/PDB/CUUCGG/2KOC_5_C_A_0_00001.pdb 
# Loaded target data/PDB/CUUCGG/1MFY_14_C_A_0_00013.pdb 
# Loading data/PDB/CUUCGG/1MFY_14_C_A_0_00013.pdb 
# Loaded reference data/PDB/CUUCGG/2KOC_5_C_A_0_00001.pdb 
# Loaded target data/PDB/CUUCGG/1JO7_14_C_A_0_00026.pdb 
# Loading data/PDB/CUUCGG/1JO7_14_C_A_0_00026.pdb 
# Loaded reference data/PDB/CUUCGG/2KOC_5_C_A_0_00001.pdb 
# Loaded target data/PDB/CUUCGG/1VQ4_1769_C_0_0_00000.pdb 
# Loading data/PDB/CUUCGG/1VQ4_1769_C_0_0_00000.pdb 
# Loaded reference data/PDB/CUUCGG/2KOC_5_C_A_0_00001.pdb 
# Loaded target data/PDB/CUUCGG/5IWA_419_C_A_0_00000.pdb 
# Loading data/PDB/CUUCGG/5IWA_419_C_A_0_00000.pdb 
# Loaded reference data/PDB/CUUCGG/2KOC_5_C_A_0_00001.pdb 
# Loaded target data/PDB/CUUCGG/2LHP_16_C_A_0_00008.pdb 
# Loading data/PDB/CUUCGG/2LHP_16_C_A_0_00008.pdb 
# Loaded reference data/PDB/CUUCGG/2KOC_5_C_A_0_00001.pdb 
# Loaded target data/PDB/CUUCGG/1BYJ_11_C_A_0_00001.pdb 
# Loading data/PDB/CUUCGG/1BYJ_11_C_A_0_00001.pdb 
# Loaded reference data/PDB/CUUCGG/2KOC_5_C_A_0_00001.pdb 
# Loaded target data/PDB/CUUCGG/1B36_16_C_A_0_00008.pdb 
# Loading data/PDB/CUUCGG/1B36_16_C_A_0_00008.pdb 
# Loaded reference data/PDB/CUUCGG/2KOC_5_C_A_0_00001.pdb 
# Loaded target data/PDB/CUUCGG/5j88-pdb-bundle3_1691_C_c_0_00000.pdb 
# Loading data/PDB/CUUCGG/5j88-pdb-bundle3_1691_C_c_0_00000.pdb 
# Loaded reference data/PDB/CUUCGG/2KOC_5_C_A_0_00001.pdb 
# Loaded target data/PDB/CUUCGG/1BYJ_11_C_A_0_00015.pdb 
# Loading data/PDB/CUUCGG/1BYJ_11_C_A_0_00015.pdb 
# Loaded reference data/PDB/CUUCGG/2KOC_5_C_A_0_00001.pdb 
# Loaded target data/PDB/CUUCGG/1Q82_1769_C_A_0_00000.pdb 
# Loading data/PDB/CUUCGG/1Q82_1769_C_A_0_00000.pdb 
# Loaded reference data/PDB/CUUCGG/2KOC_5_C_A_0_00001.pdb 
# Loaded target data/PDB/CUUCGG/2CD3_12_C_A_0_00000.pdb 
# Loading data/PDB/CUUCGG/2CD3_12_C_A_0_00000.pdb 
# Loaded reference data/PDB/CUUCGG/2KOC_5_C_A_0_00001.pdb 
# Loaded target data/PDB/CUUCGG/4v57-pdb-bundle3_419_C_A_0_00001.pdb 
# Loading data/PDB/CUUCGG/4v57-pdb-bundle3_419_C_A_0_00001.pdb 
# Loaded reference data/PDB/CUUCGG/2KOC_5_C_A_0_00001.pdb 
# Loaded target data/PDB/CUUCGG/5ib7-pdb-bundle4_1691_C_A_0_00000.pdb 
# Loading data/PDB/CUUCGG/5ib7-pdb-bundle4_1691_C_A_0_00000.pdb 
# Loaded reference data/PDB/CUUCGG/2KOC_5_C_A_0_00001.pdb 
# Loaded target data/PDB/CUUCGG/6i7v-pdb-bundle1_419_C_A_0_00001.pdb 
# Loading data/PDB/CUUCGG/6i7v-pdb-bundle1_419_C_A_0_00001.pdb 
# Loaded reference data/PDB/CUUCGG/2KOC_5_C_A_0_00001.pdb 
# Loaded target data/PDB/CUUCGG/2N3Q_37_C_A_0_00019.pdb 
# Loading data/PDB/CUUCGG/2N3Q_37_C_A_0_00019.pdb 
# Loaded reference data/PDB/CUUCGG/2KOC_5_C_A_0_00001.pdb 
# Loaded target data/PDB/CUUCGG/1BYJ_11_C_A_0_00029.pdb 
# Loading data/PDB/CUUCGG/1BYJ_11_C_A_0_00029.pdb 
# Loaded reference data/PDB/CUUCGG/2KOC_5_C_A_0_00001.pdb 
# Loaded target data/PDB/CUUCGG/5lyb-pdb-bundle3_230_C_d_0_00000.pdb 
# Loading data/PDB/CUUCGG/5lyb-pdb-bundle3_230_C_d_0_00000.pdb 
# Loaded reference data/PDB/CUUCGG/2KOC_5_C_A_0_00001.pdb 
# Loaded target data/PDB/CUUCGG/4v7s-pdb-bundle3_1449_C_A_0_00003.pdb 
# Loading data/PDB/CUUCGG/4v7s-pdb-bundle3_1449_C_A_0_00003.pdb 
# Loaded reference data/PDB/CUUCGG/2KOC_5_C_A_0_00001.pdb 
# Loaded target data/PDB/CUUCGG/5ndk-pdb-bundle2_1741_C_A_0_00000.pdb 
# Loading data/PDB/CUUCGG/5ndk-pdb-bundle2_1741_C_A_0_00000.pdb 
# Loaded reference data/PDB/CUUCGG/2KOC_5_C_A_0_00001.pdb 
# Loaded target data/PDB/CUUCGG/4LF8_419_C_A_0_00000.pdb 
# Loading data/PDB/CUUCGG/4LF8_419_C_A_0_00000.pdb 
# Loaded reference data/PDB/CUUCGG/2KOC_5_C_A_0_00001.pdb 
# Loaded target data/PDB/CUUCGG/1D6K_283_C_B_0_00019.pdb 
# Loading data/PDB/CUUCGG/1D6K_283_C_B_0_00019.pdb 
# Loaded reference data/PDB/CUUCGG/2KOC_5_C_A_0_00001.pdb 
# Loaded target data/PDB/CUUCGG/1TLR_10_C_A_0_00002.pdb 
# Loading data/PDB/CUUCGG/1TLR_10_C_A_0_00002.pdb 
# Loaded reference data/PDB/CUUCGG/2KOC_5_C_A_0_00001.pdb 
# Loaded target data/PDB/CUUCGG/4v8c-pdb-bundle3_419_C_Z_0_00001.pdb 
# Loading data/PDB/CUUCGG/4v8c-pdb-bundle3_419_C_Z_0_00001.pdb 
# Loaded reference data/PDB/CUUCGG/2KOC_5_C_A_0_00001.pdb 
# Loaded target data/PDB/CUUCGG/4WFA_1735_C_X_0_00000.pdb 
# Loading data/PDB/CUUCGG/4WFA_1735_C_X_0_00000.pdb 
# Loaded reference data/PDB/CUUCGG/2KOC_5_C_A_0_00001.pdb 
# Loaded target data/PDB/CUUCGG/6h4n-pdb-bundle1_207_C_a_0_00000.pdb 
# Loading data/PDB/CUUCGG/6h4n-pdb-bundle1_207_C_a_0_00000.pdb 
# Loaded reference data/PDB/CUUCGG/2KOC_5_C_A_0_00001.pdb 
# Loaded target data/PDB/CUUCGG/4v4q-pdb-bundle1_1449_C_A_0_00003.pdb 
# Loading data/PDB/CUUCGG/4v4q-pdb-bundle1_1449_C_A_0_00003.pdb 
# Loaded reference data/PDB/CUUCGG/2KOC_5_C_A_0_00001.pdb 
# Loaded target data/PDB/CUUCGG/4v52-pdb-bundle1_419_C_A_0_00001.pdb 
# Loading data/PDB/CUUCGG/4v52-pdb-bundle1_419_C_A_0_00001.pdb 
# Loaded reference data/PDB/CUUCGG/2KOC_5_C_A_0_00001.pdb 
# Loaded target data/PDB/CUUCGG/5Y85_28_C_B_0_00000.pdb 
# Loading data/PDB/CUUCGG/5Y85_28_C_B_0_00000.pdb 
# Loaded reference data/PDB/CUUCGG/2KOC_5_C_A_0_00001.pdb 
# Loaded target data/PDB/CUUCGG/1TLR_10_C_A_0_00016.pdb 
# Loading data/PDB/CUUCGG/1TLR_10_C_A_0_00016.pdb 
# Loaded reference data/PDB/CUUCGG/2KOC_5_C_A_0_00001.pdb 
# Loaded target data/PDB/CUUCGG/6buw-pdb-bundle4_1691_C_A_0_00000.pdb 
# Loading data/PDB/CUUCGG/6buw-pdb-bundle4_1691_C_A_0_00000.pdb 
# Loaded reference data/PDB/CUUCGG/2KOC_5_C_A_0_00001.pdb 
# Loaded target data/PDB/CUUCGG/2N6X_20_C_A_0_00001.pdb 
# Loading data/PDB/CUUCGG/2N6X_20_C_A_0_00001.pdb 
# Loaded reference data/PDB/CUUCGG/2KOC_5_C_A_0_00001.pdb 
# Loaded target data/PDB/CUUCGG/4v8e-pdb-bundle3_1739_C_A_0_00000.pdb 
# Loading data/PDB/CUUCGG/4v8e-pdb-bundle3_1739_C_A_0_00000.pdb 
# Loaded reference data/PDB/CUUCGG/2KOC_5_C_A_0_00001.pdb 
# Loaded target data/PDB/CUUCGG/5KQE_19_C_A_0_00016.pdb 
# Loading data/PDB/CUUCGG/5KQE_19_C_A_0_00016.pdb 
# Loaded reference data/PDB/CUUCGG/2KOC_5_C_A_0_00001.pdb 
# Loaded target data/PDB/CUUCGG/4v9s-pdb-bundle4_1691_C_A_0_00000.pdb 
# Loading data/PDB/CUUCGG/4v9s-pdb-bundle4_1691_C_A_0_00000.pdb 
# Loaded reference data/PDB/CUUCGG/2KOC_5_C_A_0_00001.pdb 
# Loaded target data/PDB/CUUCGG/1FYO_11_C_A_0_00016.pdb 
# Loading data/PDB/CUUCGG/1FYO_11_C_A_0_00016.pdb 
# Loaded reference data/PDB/CUUCGG/2KOC_5_C_A_0_00001.pdb 
# Loaded target data/PDB/CUUCGG/4v4h-pdb-bundle1_1028_C_A_0_00002.pdb 
# Loading data/PDB/CUUCGG/4v4h-pdb-bundle1_1028_C_A_0_00002.pdb 
# Loaded reference data/PDB/CUUCGG/2KOC_5_C_A_0_00001.pdb 
# Loaded target data/PDB/CUUCGG/1FYO_11_C_A_0_00002.pdb 
# Loading data/PDB/CUUCGG/1FYO_11_C_A_0_00002.pdb 
# Loaded reference data/PDB/CUUCGG/2KOC_5_C_A_0_00001.pdb 
# Loaded target data/PDB/CUUCGG/5i4l-pdb-bundle3_230_C_c_0_00000.pdb 
# Loading data/PDB/CUUCGG/5i4l-pdb-bundle3_230_C_c_0_00000.pdb 
# Loaded reference data/PDB/CUUCGG/2KOC_5_C_A_0_00001.pdb 
# Loaded target data/PDB/CUUCGG/5KQE_19_C_A_0_00002.pdb 
# Loading data/PDB/CUUCGG/5KQE_19_C_A_0_00002.pdb 
# Loaded reference data/PDB/CUUCGG/2KOC_5_C_A_0_00001.pdb 
# Loaded target data/PDB/CUUCGG/4u27-pdb-bundle3_419_C_A_0_00001.pdb 
# Loading data/PDB/CUUCGG/4u27-pdb-bundle3_419_C_A_0_00001.pdb 
# Loaded reference data/PDB/CUUCGG/2KOC_5_C_A_0_00001.pdb 
# Loaded target data/PDB/CUUCGG/4JI7_419_C_A_0_00000.pdb 
# Loading data/PDB/CUUCGG/4JI7_419_C_A_0_00000.pdb 
# Loaded reference data/PDB/CUUCGG/2KOC_5_C_A_0_00001.pdb 
# Loaded target data/PDB/CUUCGG/1FYO_11_C_A_0_00003.pdb 
# Loading data/PDB/CUUCGG/1FYO_11_C_A_0_00003.pdb 
# Loaded reference data/PDB/CUUCGG/2KOC_5_C_A_0_00001.pdb 
# Loaded target data/PDB/CUUCGG/5KQE_19_C_A_0_00003.pdb 
# Loading data/PDB/CUUCGG/5KQE_19_C_A_0_00003.pdb 
# Loaded reference data/PDB/CUUCGG/2KOC_5_C_A_0_00001.pdb 
# Loaded target data/PDB/CUUCGG/5KQE_19_C_A_0_00017.pdb 
# Loading data/PDB/CUUCGG/5KQE_19_C_A_0_00017.pdb 
# Loaded reference data/PDB/CUUCGG/2KOC_5_C_A_0_00001.pdb 
# Loaded target data/PDB/CUUCGG/1FYO_11_C_A_0_00017.pdb 
# Loading data/PDB/CUUCGG/1FYO_11_C_A_0_00017.pdb 
# Loaded reference data/PDB/CUUCGG/2KOC_5_C_A_0_00001.pdb 
# Loaded target data/PDB/CUUCGG/6boh-pdb-bundle1_419_C_A_0_00000.pdb 
# Loading data/PDB/CUUCGG/6boh-pdb-bundle1_419_C_A_0_00000.pdb 
# Loaded reference data/PDB/CUUCGG/2KOC_5_C_A_0_00001.pdb 
# Loaded target data/PDB/CUUCGG/2N6X_20_C_A_0_00000.pdb 
# Loading data/PDB/CUUCGG/2N6X_20_C_A_0_00000.pdb 
# Loaded reference data/PDB/CUUCGG/2KOC_5_C_A_0_00001.pdb 
# Loaded target data/PDB/CUUCGG/1QVG_1769_C_0_0_00000.pdb 
# Loading data/PDB/CUUCGG/1QVG_1769_C_0_0_00000.pdb 
# Loaded reference data/PDB/CUUCGG/2KOC_5_C_A_0_00001.pdb 
# Loaded target data/PDB/CUUCGG/5it8-pdb-bundle2_1691_C_T_0_00000.pdb 
# Loading data/PDB/CUUCGG/5it8-pdb-bundle2_1691_C_T_0_00000.pdb 
# Loaded reference data/PDB/CUUCGG/2KOC_5_C_A_0_00001.pdb 
# Loaded target data/PDB/CUUCGG/6gsl-pdb-bundle3_419_C_A_0_00000.pdb 
# Loading data/PDB/CUUCGG/6gsl-pdb-bundle3_419_C_A_0_00000.pdb 
# Loaded reference data/PDB/CUUCGG/2KOC_5_C_A_0_00001.pdb 
# Loaded target data/PDB/CUUCGG/4wsm-pdb-bundle4_1691_C_A_0_00000.pdb 
# Loading data/PDB/CUUCGG/4wsm-pdb-bundle4_1691_C_A_0_00000.pdb 
# Loaded reference data/PDB/CUUCGG/2KOC_5_C_A_0_00001.pdb 
# Loaded target data/PDB/CUUCGG/4v54-pdb-bundle3_419_C_K_0_00001.pdb 
# Loading data/PDB/CUUCGG/4v54-pdb-bundle3_419_C_K_0_00001.pdb 
# Loaded reference data/PDB/CUUCGG/2KOC_5_C_A_0_00001.pdb 
# Loaded target data/PDB/CUUCGG/1TLR_10_C_A_0_00017.pdb 
# Loading data/PDB/CUUCGG/1TLR_10_C_A_0_00017.pdb 
# Loaded reference data/PDB/CUUCGG/2KOC_5_C_A_0_00001.pdb 
# Loaded target data/PDB/CUUCGG/6G7Z_21_C_A_0_00000.pdb 
# Loading data/PDB/CUUCGG/6G7Z_21_C_A_0_00000.pdb 
# Loaded reference data/PDB/CUUCGG/2KOC_5_C_A_0_00001.pdb 
# Loaded target data/PDB/CUUCGG/1TLR_10_C_A_0_00003.pdb 
# Loading data/PDB/CUUCGG/1TLR_10_C_A_0_00003.pdb 
# Loaded reference data/PDB/CUUCGG/2KOC_5_C_A_0_00001.pdb 
# Loaded target data/PDB/CUUCGG/1D6K_283_C_B_0_00018.pdb 
# Loading data/PDB/CUUCGG/1D6K_283_C_B_0_00018.pdb 
# Loaded reference data/PDB/CUUCGG/2KOC_5_C_A_0_00001.pdb 
# Loaded target data/PDB/CUUCGG/6cfl-pdb-bundle1_1738_C_A_0_00000.pdb 
# Loading data/PDB/CUUCGG/6cfl-pdb-bundle1_1738_C_A_0_00000.pdb 
# Loaded reference data/PDB/CUUCGG/2KOC_5_C_A_0_00001.pdb 
# Loaded target data/PDB/CUUCGG/4DV7_419_C_A_0_00000.pdb 
# Loading data/PDB/CUUCGG/4DV7_419_C_A_0_00000.pdb 
# Loaded reference data/PDB/CUUCGG/2KOC_5_C_A_0_00001.pdb 
# Loaded target data/PDB/CUUCGG/4u1v-pdb-bundle2_1691_C_A_0_00000.pdb 
# Loading data/PDB/CUUCGG/4u1v-pdb-bundle2_1691_C_A_0_00000.pdb 
# Loaded reference data/PDB/CUUCGG/2KOC_5_C_A_0_00001.pdb 
# Loaded target data/PDB/CUUCGG/3jct-pdb-bundle2_1923_C_1_0_00002.pdb 
# Loading data/PDB/CUUCGG/3jct-pdb-bundle2_1923_C_1_0_00002.pdb 
# Loaded reference data/PDB/CUUCGG/2KOC_5_C_A_0_00001.pdb 
# Loaded target data/PDB/CUUCGG/4v4q-pdb-bundle1_207_C_A_0_00000.pdb 
# Loading data/PDB/CUUCGG/4v4q-pdb-bundle1_207_C_A_0_00000.pdb 
# Loaded reference data/PDB/CUUCGG/2KOC_5_C_A_0_00001.pdb 
# Loaded target data/PDB/CUUCGG/1BYJ_11_C_A_0_00028.pdb 
# Loading data/PDB/CUUCGG/1BYJ_11_C_A_0_00028.pdb 
# Loaded reference data/PDB/CUUCGG/2KOC_5_C_A_0_00001.pdb 
# Loaded target data/PDB/CUUCGG/2N3Q_37_C_A_0_00018.pdb 
# Loading data/PDB/CUUCGG/2N3Q_37_C_A_0_00018.pdb 
# Loaded reference data/PDB/CUUCGG/2KOC_5_C_A_0_00001.pdb 
# Loaded target data/PDB/CUUCGG/6hrm-pdb-bundle1_3197_C_1_0_00003.pdb 
# Loading data/PDB/CUUCGG/6hrm-pdb-bundle1_3197_C_1_0_00003.pdb 
# Loaded reference data/PDB/CUUCGG/2KOC_5_C_A_0_00001.pdb 
# Loaded target data/PDB/CUUCGG/4v9p-pdb-bundle6_1449_C_A_0_00003.pdb 
# Loading data/PDB/CUUCGG/4v9p-pdb-bundle6_1449_C_A_0_00003.pdb 
# Loaded reference data/PDB/CUUCGG/2KOC_5_C_A_0_00001.pdb 
# Loaded target data/PDB/CUUCGG/1BYJ_11_C_A_0_00014.pdb 
# Loading data/PDB/CUUCGG/1BYJ_11_C_A_0_00014.pdb 
# Loaded reference data/PDB/CUUCGG/2KOC_5_C_A_0_00001.pdb 
# Loaded target data/PDB/CUUCGG/1B36_16_C_A_0_00009.pdb 
# Loading data/PDB/CUUCGG/1B36_16_C_A_0_00009.pdb 
# Loaded reference data/PDB/CUUCGG/2KOC_5_C_A_0_00001.pdb 
# Loaded target data/PDB/CUUCGG/1BYJ_11_C_A_0_00000.pdb 
# Loading data/PDB/CUUCGG/1BYJ_11_C_A_0_00000.pdb 
# Loaded reference data/PDB/CUUCGG/2KOC_5_C_A_0_00001.pdb 
# Loaded target data/PDB/CUUCGG/2LHP_16_C_A_0_00009.pdb 
# Loading data/PDB/CUUCGG/2LHP_16_C_A_0_00009.pdb 
# Loaded reference data/PDB/CUUCGG/2KOC_5_C_A_0_00001.pdb 
# Loaded target data/PDB/CUUCGG/4v83-pdb-bundle2_1691_C_A_0_00000.pdb 
# Loading data/PDB/CUUCGG/4v83-pdb-bundle2_1691_C_A_0_00000.pdb 
# Loaded reference data/PDB/CUUCGG/2KOC_5_C_A_0_00001.pdb 
# Loaded target data/PDB/CUUCGG/4w2g-pdb-bundle2_1738_C_A_0_00000.pdb 
# Loading data/PDB/CUUCGG/4w2g-pdb-bundle2_1738_C_A_0_00000.pdb 
# Loaded reference data/PDB/CUUCGG/2KOC_5_C_A_0_00001.pdb 
# Loaded target data/PDB/CUUCGG/5nwy-pdb-bundle2_207_C_0_0_00000.pdb 
# Loading data/PDB/CUUCGG/5nwy-pdb-bundle2_207_C_0_0_00000.pdb 
# Loaded reference data/PDB/CUUCGG/2KOC_5_C_A_0_00001.pdb 
# Loaded target data/PDB/CUUCGG/3CCL_1769_C_0_0_00000.pdb 
# Loading data/PDB/CUUCGG/3CCL_1769_C_0_0_00000.pdb 
# Loaded reference data/PDB/CUUCGG/2KOC_5_C_A_0_00001.pdb 
# Loaded target data/PDB/CUUCGG/2GIO_13_C_A_0_00011.pdb 
# Loading data/PDB/CUUCGG/2GIO_13_C_A_0_00011.pdb 
# Loaded reference data/PDB/CUUCGG/2KOC_5_C_A_0_00001.pdb 
# Loaded target data/PDB/CUUCGG/1MFY_14_C_A_0_00012.pdb 
# Loading data/PDB/CUUCGG/1MFY_14_C_A_0_00012.pdb 
# Loaded reference data/PDB/CUUCGG/2KOC_5_C_A_0_00001.pdb 
# Loaded target data/PDB/CUUCGG/5KH8_26_C_A_0_00016.pdb 
# Loading data/PDB/CUUCGG/5KH8_26_C_A_0_00016.pdb 
# Loaded reference data/PDB/CUUCGG/2KOC_5_C_A_0_00001.pdb 
# Loaded target data/PDB/CUUCGG/1MFY_14_C_A_0_00006.pdb 
# Loading data/PDB/CUUCGG/1MFY_14_C_A_0_00006.pdb 
# Loaded reference data/PDB/CUUCGG/2KOC_5_C_A_0_00001.pdb 
# Loaded target data/PDB/CUUCGG/5KH8_26_C_A_0_00002.pdb 
# Loading data/PDB/CUUCGG/5KH8_26_C_A_0_00002.pdb 
# Loaded reference data/PDB/CUUCGG/2KOC_5_C_A_0_00001.pdb 
# Loaded target data/PDB/CUUCGG/2GIO_13_C_A_0_00005.pdb 
# Loading data/PDB/CUUCGG/2GIO_13_C_A_0_00005.pdb 
# Loaded reference data/PDB/CUUCGG/2KOC_5_C_A_0_00001.pdb 
# Loaded target data/PDB/CUUCGG/2GIP_12_C_A_0_00002.pdb 
# Loading data/PDB/CUUCGG/2GIP_12_C_A_0_00002.pdb 
# Loaded reference data/PDB/CUUCGG/2KOC_5_C_A_0_00001.pdb 
# Loaded target data/PDB/CUUCGG/2GIP_12_C_A_0_00016.pdb 
# Loading data/PDB/CUUCGG/2GIP_12_C_A_0_00016.pdb 
# Loaded reference data/PDB/CUUCGG/2KOC_5_C_A_0_00001.pdb 
# Loaded target data/PDB/CUUCGG/5dgv-pdb-bundle2_1923_C_A_0_00002.pdb 
# Loading data/PDB/CUUCGG/5dgv-pdb-bundle2_1923_C_A_0_00002.pdb 
# Loaded reference data/PDB/CUUCGG/2KOC_5_C_A_0_00001.pdb 
# Loaded target data/PDB/CUUCGG/6c5l-pdb-bundle4_1691_C_A_0_00000.pdb 
# Loading data/PDB/CUUCGG/6c5l-pdb-bundle4_1691_C_A_0_00000.pdb 
# Loaded reference data/PDB/CUUCGG/2KOC_5_C_A_0_00001.pdb 
# Loaded target data/PDB/CUUCGG/4v57-pdb-bundle2_1691_C_A_0_00000.pdb 
# Loading data/PDB/CUUCGG/4v57-pdb-bundle2_1691_C_A_0_00000.pdb 
# Loaded reference data/PDB/CUUCGG/2KOC_5_C_A_0_00001.pdb 
# Loaded target data/PDB/CUUCGG/1vy4-pdb-bundle3_419_C_A_0_00000.pdb 
# Loading data/PDB/CUUCGG/1vy4-pdb-bundle3_419_C_A_0_00000.pdb 
# Loaded reference data/PDB/CUUCGG/2KOC_5_C_A_0_00001.pdb 
# Loaded target data/PDB/CUUCGG/2KU0_19_C_A_0_00001.pdb 
# Loading data/PDB/CUUCGG/2KU0_19_C_A_0_00001.pdb 
# Loaded reference data/PDB/CUUCGG/2KOC_5_C_A_0_00001.pdb 
# Loaded target data/PDB/CUUCGG/4www-pdb-bundle2_1028_C_V_0_00006.pdb 
# Loading data/PDB/CUUCGG/4www-pdb-bundle2_1028_C_V_0_00006.pdb 
# Loaded reference data/PDB/CUUCGG/2KOC_5_C_A_0_00001.pdb 
# Loaded target data/PDB/CUUCGG/2N3R_37_C_A_0_00020.pdb 
# Loading data/PDB/CUUCGG/2N3R_37_C_A_0_00020.pdb 
# Loaded reference data/PDB/CUUCGG/2KOC_5_C_A_0_00001.pdb 
# Loaded target data/PDB/CUUCGG/2HHH_415_C_A_0_00000.pdb 
# Loading data/PDB/CUUCGG/2HHH_415_C_A_0_00000.pdb 
# Loaded reference data/PDB/CUUCGG/2KOC_5_C_A_0_00001.pdb 
# Loaded target data/PDB/CUUCGG/4LF9_419_C_A_0_00000.pdb 
# Loading data/PDB/CUUCGG/4LF9_419_C_A_0_00000.pdb 
# Loaded reference data/PDB/CUUCGG/2KOC_5_C_A_0_00001.pdb 
# Loaded target data/PDB/CUUCGG/4yzv-pdb-bundle4_1691_C_A_0_00000.pdb 
# Loading data/PDB/CUUCGG/4yzv-pdb-bundle4_1691_C_A_0_00000.pdb 
# Loaded reference data/PDB/CUUCGG/2KOC_5_C_A_0_00001.pdb 
# Loaded target data/PDB/CUUCGG/5u4i-pdb-bundle2_419_C_a_0_00001.pdb 
# Loading data/PDB/CUUCGG/5u4i-pdb-bundle2_419_C_a_0_00001.pdb 
# Loaded reference data/PDB/CUUCGG/2KOC_5_C_A_0_00001.pdb 
# Loaded target data/PDB/CUUCGG/2N3R_37_C_A_0_00008.pdb 
# Loading data/PDB/CUUCGG/2N3R_37_C_A_0_00008.pdb 
# Loaded reference data/PDB/CUUCGG/2KOC_5_C_A_0_00001.pdb 
# Loaded target data/PDB/CUUCGG/5j8a-pdb-bundle2_1691_C_U_0_00000.pdb 
# Loading data/PDB/CUUCGG/5j8a-pdb-bundle2_1691_C_U_0_00000.pdb 
# Loaded reference data/PDB/CUUCGG/2KOC_5_C_A_0_00001.pdb 
# Loaded target data/PDB/CUUCGG/6bu8-pdb-bundle2_1691_C_A_0_00000.pdb 
# Loading data/PDB/CUUCGG/6bu8-pdb-bundle2_1691_C_A_0_00000.pdb 
# Loaded reference data/PDB/CUUCGG/2KOC_5_C_A_0_00001.pdb 
# Loaded target data/PDB/CUUCGG/2LU0_28_C_A_0_00002.pdb 
# Loading data/PDB/CUUCGG/2LU0_28_C_A_0_00002.pdb 
# Loaded reference data/PDB/CUUCGG/2KOC_5_C_A_0_00001.pdb 
# Loaded target data/PDB/CUUCGG/2LU0_28_C_A_0_00016.pdb 
# Loading data/PDB/CUUCGG/2LU0_28_C_A_0_00016.pdb 
# Loaded reference data/PDB/CUUCGG/2KOC_5_C_A_0_00001.pdb 
# Loaded target data/PDB/CUUCGG/5j4c-pdb-bundle3_1691_C_A_0_00000.pdb 
# Loading data/PDB/CUUCGG/5j4c-pdb-bundle3_1691_C_A_0_00000.pdb 
# Loaded reference data/PDB/CUUCGG/2KOC_5_C_A_0_00001.pdb 
# Loaded target data/PDB/CUUCGG/4v5c-pdb-bundle1_419_C_A_0_00000.pdb 
# Loading data/PDB/CUUCGG/4v5c-pdb-bundle1_419_C_A_0_00000.pdb 
# Loaded reference data/PDB/CUUCGG/2KOC_5_C_A_0_00001.pdb 
# Loaded target data/PDB/CUUCGG/3I55_1769_C_0_0_00000.pdb 
# Loading data/PDB/CUUCGG/3I55_1769_C_0_0_00000.pdb 
# Loaded reference data/PDB/CUUCGG/2KOC_5_C_A_0_00001.pdb 
# Loaded target data/PDB/CUUCGG/2L2J_19_C_A_0_00006.pdb 
# Loading data/PDB/CUUCGG/2L2J_19_C_A_0_00006.pdb 
# Loaded reference data/PDB/CUUCGG/2KOC_5_C_A_0_00001.pdb 
# Loaded target data/PDB/CUUCGG/1EKZ_13_C_B_0_00027.pdb 
# Loading data/PDB/CUUCGG/1EKZ_13_C_B_0_00027.pdb 
# Loaded reference data/PDB/CUUCGG/2KOC_5_C_A_0_00001.pdb 
# Loaded target data/PDB/CUUCGG/1YJN_1769_C_0_0_00000.pdb 
# Loading data/PDB/CUUCGG/1YJN_1769_C_0_0_00000.pdb 
# Loaded reference data/PDB/CUUCGG/2KOC_5_C_A_0_00001.pdb 
# Loaded target data/PDB/CUUCGG/2L2J_19_C_A_0_00012.pdb 
# Loading data/PDB/CUUCGG/2L2J_19_C_A_0_00012.pdb 
# Loaded reference data/PDB/CUUCGG/2KOC_5_C_A_0_00001.pdb 
# Loaded target data/PDB/CUUCGG/4DR2_419_C_A_0_00000.pdb 
# Loading data/PDB/CUUCGG/4DR2_419_C_A_0_00000.pdb 
# Loaded reference data/PDB/CUUCGG/2KOC_5_C_A_0_00001.pdb 
# Loaded target data/PDB/CUUCGG/1EKZ_13_C_B_0_00033.pdb 
# Loading data/PDB/CUUCGG/1EKZ_13_C_B_0_00033.pdb 
# Loaded reference data/PDB/CUUCGG/2KOC_5_C_A_0_00001.pdb 
# Loaded target data/PDB/CUUCGG/5j91-pdb-bundle1_419_C_A_0_00001.pdb 
# Loading data/PDB/CUUCGG/5j91-pdb-bundle1_419_C_A_0_00001.pdb 
# Loaded reference data/PDB/CUUCGG/2KOC_5_C_A_0_00001.pdb 
# Loaded target data/PDB/CUUCGG/4w2f-pdb-bundle4_1691_C_A_0_00000.pdb 
# Loading data/PDB/CUUCGG/4w2f-pdb-bundle4_1691_C_A_0_00000.pdb 
# Loaded reference data/PDB/CUUCGG/2KOC_5_C_A_0_00001.pdb 
# Loaded target data/PDB/CUUCGG/1P5M_28_C_A_0_00003.pdb 
# Loading data/PDB/CUUCGG/1P5M_28_C_A_0_00003.pdb 
# Loaded reference data/PDB/CUUCGG/2KOC_5_C_A_0_00001.pdb 
# Loaded target data/PDB/CUUCGG/1F7I_12_C_A_0_00002.pdb 
# Loading data/PDB/CUUCGG/1F7I_12_C_A_0_00002.pdb 
# Loaded reference data/PDB/CUUCGG/2KOC_5_C_A_0_00001.pdb 
# Loaded target data/PDB/CUUCGG/1ULL_14_C_A_0_00001.pdb 
# Loading data/PDB/CUUCGG/1ULL_14_C_A_0_00001.pdb 
# Loaded reference data/PDB/CUUCGG/2KOC_5_C_A_0_00001.pdb 
# Loaded target data/PDB/CUUCGG/2N6T_19_C_A_0_00000.pdb 
# Loading data/PDB/CUUCGG/2N6T_19_C_A_0_00000.pdb 
# Loaded reference data/PDB/CUUCGG/2KOC_5_C_A_0_00001.pdb 
# Loaded target data/PDB/CUUCGG/1P5M_28_C_A_0_00017.pdb 
# Loading data/PDB/CUUCGG/1P5M_28_C_A_0_00017.pdb 
# Loaded reference data/PDB/CUUCGG/2KOC_5_C_A_0_00001.pdb 
# Loaded target data/PDB/CUUCGG/4v54-pdb-bundle1_419_C_A_0_00001.pdb 
# Loading data/PDB/CUUCGG/4v54-pdb-bundle1_419_C_A_0_00001.pdb 
# Loaded reference data/PDB/CUUCGG/2KOC_5_C_A_0_00001.pdb 
# Loaded target data/PDB/CUUCGG/4v9c-pdb-bundle4_1691_C_A_0_00000.pdb 
# Loading data/PDB/CUUCGG/4v9c-pdb-bundle4_1691_C_A_0_00000.pdb 
# Loaded reference data/PDB/CUUCGG/2KOC_5_C_A_0_00001.pdb 
# Loaded target data/PDB/CUUCGG/2N6W_32_C_A_0_00007.pdb 
# Loading data/PDB/CUUCGG/2N6W_32_C_A_0_00007.pdb 
# Loaded reference data/PDB/CUUCGG/2KOC_5_C_A_0_00001.pdb 
# Loaded target data/PDB/CUUCGG/1RAW_21_C_A_0_00002.pdb 
# Loading data/PDB/CUUCGG/1RAW_21_C_A_0_00002.pdb 
# Loaded reference data/PDB/CUUCGG/2KOC_5_C_A_0_00001.pdb 
# Loaded target data/PDB/CUUCGG/5j5b-pdb-bundle2_1691_C_U_0_00000.pdb 
# Loading data/PDB/CUUCGG/5j5b-pdb-bundle2_1691_C_U_0_00000.pdb 
# Loaded reference data/PDB/CUUCGG/2KOC_5_C_A_0_00001.pdb 
# Loaded target data/PDB/CUUCGG/1AUD_28_C_B_0_00006.pdb 
# Loading data/PDB/CUUCGG/1AUD_28_C_B_0_00006.pdb 
# Loaded reference data/PDB/CUUCGG/2KOC_5_C_A_0_00001.pdb 
# Loaded target data/PDB/CUUCGG/6cfk-pdb-bundle1_1738_C_A_0_00000.pdb 
# Loading data/PDB/CUUCGG/6cfk-pdb-bundle1_1738_C_A_0_00000.pdb 
# Loaded reference data/PDB/CUUCGG/2KOC_5_C_A_0_00001.pdb 
# Loaded target data/PDB/CUUCGG/1EBS_12_C_A_0_00001.pdb 
# Loading data/PDB/CUUCGG/1EBS_12_C_A_0_00001.pdb 
# Loaded reference data/PDB/CUUCGG/2KOC_5_C_A_0_00001.pdb 
# Loaded target data/PDB/CUUCGG/4u24-pdb-bundle1_419_C_A_0_00001.pdb 
# Loading data/PDB/CUUCGG/4u24-pdb-bundle1_419_C_A_0_00001.pdb 
# Loaded reference data/PDB/CUUCGG/2KOC_5_C_A_0_00001.pdb 
# Loaded target data/PDB/CUUCGG/1FYP_11_C_A_0_00012.pdb 
# Loading data/PDB/CUUCGG/1FYP_11_C_A_0_00012.pdb 
# Loaded reference data/PDB/CUUCGG/2KOC_5_C_A_0_00001.pdb 
# Loaded target data/PDB/CUUCGG/4z3s-pdb-bundle3_1691_C_A_0_00000.pdb 
# Loading data/PDB/CUUCGG/4z3s-pdb-bundle3_1691_C_A_0_00000.pdb 
# Loaded reference data/PDB/CUUCGG/2KOC_5_C_A_0_00001.pdb 
# Loaded target data/PDB/CUUCGG/1FYP_11_C_A_0_00006.pdb 
# Loading data/PDB/CUUCGG/1FYP_11_C_A_0_00006.pdb 
# Loaded reference data/PDB/CUUCGG/2KOC_5_C_A_0_00001.pdb 
# Loaded target data/PDB/CUUCGG/4LF4_419_C_A_0_00000.pdb 
# Loading data/PDB/CUUCGG/4LF4_419_C_A_0_00000.pdb 
# Loaded reference data/PDB/CUUCGG/2KOC_5_C_A_0_00001.pdb 
# Loaded target data/PDB/CUUCGG/4wsd-pdb-bundle2_1691_C_A_0_00000.pdb 
# Loading data/PDB/CUUCGG/4wsd-pdb-bundle2_1691_C_A_0_00000.pdb 
# Loaded reference data/PDB/CUUCGG/2KOC_5_C_A_0_00001.pdb 
# Loaded target data/PDB/CUUCGG/1vy5-pdb-bundle4_1691_C_A_0_00000.pdb 
# Loading data/PDB/CUUCGG/1vy5-pdb-bundle4_1691_C_A_0_00000.pdb 
# Loaded reference data/PDB/CUUCGG/2KOC_5_C_A_0_00001.pdb 
# Loaded target data/PDB/CUUCGG/4wf1-pdb-bundle1_1028_C_A_0_00002.pdb 
# Loading data/PDB/CUUCGG/4wf1-pdb-bundle1_1028_C_A_0_00002.pdb 
# Loaded reference data/PDB/CUUCGG/2KOC_5_C_A_0_00001.pdb 
# Loaded target data/PDB/CUUCGG/1M82_10_C_A_0_00002.pdb 
# Loading data/PDB/CUUCGG/1M82_10_C_A_0_00002.pdb 
# Loaded reference data/PDB/CUUCGG/2KOC_5_C_A_0_00001.pdb 
# Loaded target data/PDB/CUUCGG/4v6a-pdb-bundle2_1691_C_A_0_00000.pdb 
# Loading data/PDB/CUUCGG/4v6a-pdb-bundle2_1691_C_A_0_00000.pdb 
# Loaded reference data/PDB/CUUCGG/2KOC_5_C_A_0_00001.pdb 
# Loaded target data/PDB/CUUCGG/4v84-pdb-bundle2_1691_C_A_0_00000.pdb 
# Loading data/PDB/CUUCGG/4v84-pdb-bundle2_1691_C_A_0_00000.pdb 
# Loaded reference data/PDB/CUUCGG/2KOC_5_C_A_0_00001.pdb 
# Loaded target data/PDB/CUUCGG/5vpo-pdb-bundle1_419_C_C_0_00000.pdb 
# Loading data/PDB/CUUCGG/5vpo-pdb-bundle1_419_C_C_0_00000.pdb 
# Loaded reference data/PDB/CUUCGG/2KOC_5_C_A_0_00001.pdb 
# Loaded target data/PDB/CUUCGG/1M82_10_C_A_0_00016.pdb 
# Loading data/PDB/CUUCGG/1M82_10_C_A_0_00016.pdb 
# Loaded reference data/PDB/CUUCGG/2KOC_5_C_A_0_00001.pdb 
# Loaded target data/PDB/CUUCGG/4v9q-pdb-bundle3_1691_C_A_0_00000.pdb 
# Loading data/PDB/CUUCGG/4v9q-pdb-bundle3_1691_C_A_0_00000.pdb 
# Loaded reference data/PDB/CUUCGG/2KOC_5_C_A_0_00001.pdb 
# Loaded target data/PDB/CUUCGG/5IEM_25_C_A_0_00009.pdb 
# Loading data/PDB/CUUCGG/5IEM_25_C_A_0_00009.pdb 
# Loaded reference data/PDB/CUUCGG/2KOC_5_C_A_0_00001.pdb 
# Loaded target data/PDB/CUUCGG/4v95-pdb-bundle4_1691_C_A_0_00000.pdb 
# Loading data/PDB/CUUCGG/4v95-pdb-bundle4_1691_C_A_0_00000.pdb 
# Loaded reference data/PDB/CUUCGG/2KOC_5_C_A_0_00001.pdb 
# Loaded target data/PDB/CUUCGG/1DK1_8_C_B_0_00000.pdb 
# Loading data/PDB/CUUCGG/1DK1_8_C_B_0_00000.pdb 
# Loaded reference data/PDB/CUUCGG/2KOC_5_C_A_0_00001.pdb 
# Loaded target data/PDB/CUUCGG/4v50-pdb-bundle2_1691_C_A_0_00000.pdb 
# Loading data/PDB/CUUCGG/4v50-pdb-bundle2_1691_C_A_0_00000.pdb 
# Loaded reference data/PDB/CUUCGG/2KOC_5_C_A_0_00001.pdb 
# Loaded target data/PDB/CUUCGG/5fdv-pdb-bundle1_1738_C_A_0_00000.pdb 
# Loading data/PDB/CUUCGG/5fdv-pdb-bundle1_1738_C_A_0_00000.pdb 
# Loaded reference data/PDB/CUUCGG/2KOC_5_C_A_0_00001.pdb 
# Loaded target data/PDB/CUUCGG/1G70_54_C_A_0_00002.pdb 
# Loading data/PDB/CUUCGG/1G70_54_C_A_0_00002.pdb 
# Loaded reference data/PDB/CUUCGG/2KOC_5_C_A_0_00001.pdb 
# Loaded target data/PDB/CUUCGG/2KOC_5_C_A_0_00001.pdb 
# Loading data/PDB/CUUCGG/2KOC_5_C_A_0_00001.pdb 
# Loaded reference data/PDB/CUUCGG/2KOC_5_C_A_0_00001.pdb 
# Loaded target data/PDB/CUUCGG/2KOC_5_C_A_0_00015.pdb 
# Loading data/PDB/CUUCGG/2KOC_5_C_A_0_00015.pdb 
# Loaded reference data/PDB/CUUCGG/2KOC_5_C_A_0_00001.pdb 
# Loaded target data/PDB/CUUCGG/4v9o-pdb-bundle8_1449_C_A_0_00003.pdb 
# Loading data/PDB/CUUCGG/4v9o-pdb-bundle8_1449_C_A_0_00003.pdb 
# Loaded reference data/PDB/CUUCGG/2KOC_5_C_A_0_00001.pdb 
# Loaded target data/PDB/CUUCGG/2L2K_19_C_A_0_00008.pdb 
# Loading data/PDB/CUUCGG/2L2K_19_C_A_0_00008.pdb 
# Loaded reference data/PDB/CUUCGG/2KOC_5_C_A_0_00001.pdb 
# Loaded target data/PDB/CUUCGG/4v8b-pdb-bundle2_1691_C_A_0_00000.pdb 
# Loading data/PDB/CUUCGG/4v8b-pdb-bundle2_1691_C_A_0_00000.pdb 
# Loaded reference data/PDB/CUUCGG/2KOC_5_C_A_0_00001.pdb 
# Loaded target data/PDB/CUUCGG/4v67-pdb-bundle2_1691_C_A_0_00000.pdb 
# Loading data/PDB/CUUCGG/4v67-pdb-bundle2_1691_C_A_0_00000.pdb 
# Loaded reference data/PDB/CUUCGG/2KOC_5_C_A_0_00001.pdb 
# Loaded target data/PDB/CUUCGG/1IKD_7_C_A_0_00023.pdb 
# Loading data/PDB/CUUCGG/1IKD_7_C_A_0_00023.pdb 
# Loaded reference data/PDB/CUUCGG/2KOC_5_C_A_0_00001.pdb 
# Loaded target data/PDB/CUUCGG/5WNR_419_C_A_0_00000.pdb 
# Loading data/PDB/CUUCGG/5WNR_419_C_A_0_00000.pdb 
# Loaded reference data/PDB/CUUCGG/2KOC_5_C_A_0_00001.pdb 
# Loaded target data/PDB/CUUCGG/4w2h-pdb-bundle1_419_C_A_0_00000.pdb 
# Loading data/PDB/CUUCGG/4w2h-pdb-bundle1_419_C_A_0_00000.pdb 
# Loaded reference data/PDB/CUUCGG/2KOC_5_C_A_0_00001.pdb 
# Loaded target data/PDB/CUUCGG/1Z31_270_C_A_0_00001.pdb 
# Loading data/PDB/CUUCGG/1Z31_270_C_A_0_00001.pdb 
# Loaded reference data/PDB/CUUCGG/2KOC_5_C_A_0_00001.pdb 
# Loaded target data/PDB/CUUCGG/1C0O_5_C_A_0_00001.pdb 
# Loading data/PDB/CUUCGG/1C0O_5_C_A_0_00001.pdb 
# Loaded reference data/PDB/CUUCGG/2KOC_5_C_A_0_00001.pdb 
# Loaded target data/PDB/CUUCGG/1C0O_5_C_A_0_00015.pdb 
# Loading data/PDB/CUUCGG/1C0O_5_C_A_0_00015.pdb 
# Loaded reference data/PDB/CUUCGG/2KOC_5_C_A_0_00001.pdb 
# Loaded target data/PDB/CUUCGG/5jc9-pdb-bundle1_419_C_A_0_00001.pdb 
# Loading data/PDB/CUUCGG/5jc9-pdb-bundle1_419_C_A_0_00001.pdb 
# Loaded reference data/PDB/CUUCGG/2KOC_5_C_A_0_00001.pdb 
# Loaded target data/PDB/CUUCGG/5fci-pdb-bundle2_1923_C_A_0_00002.pdb 
# Loading data/PDB/CUUCGG/5fci-pdb-bundle2_1923_C_A_0_00002.pdb 
# Loaded reference data/PDB/CUUCGG/2KOC_5_C_A_0_00001.pdb 
# Loaded target data/PDB/CUUCGG/2HGH_32_C_B_0_00007.pdb 
# Loading data/PDB/CUUCGG/2HGH_32_C_B_0_00007.pdb 
# Loaded reference data/PDB/CUUCGG/2KOC_5_C_A_0_00001.pdb 
# Loaded target data/PDB/CUUCGG/5j4d-pdb-bundle3_1691_C_A_0_00000.pdb 
# Loading data/PDB/CUUCGG/5j4d-pdb-bundle3_1691_C_A_0_00000.pdb 
# Loaded reference data/PDB/CUUCGG/2KOC_5_C_A_0_00001.pdb 
# Loaded target data/PDB/CUUCGG/5dat-pdb-bundle4_1923_C_I_0_00002.pdb 
# Loading data/PDB/CUUCGG/5dat-pdb-bundle4_1923_C_I_0_00002.pdb 
# Loaded reference data/PDB/CUUCGG/2KOC_5_C_A_0_00001.pdb 
# Loaded target data/PDB/CUUCGG/1F7G_12_C_A_0_00006.pdb 
# Loading data/PDB/CUUCGG/1F7G_12_C_A_0_00006.pdb 
# Loaded reference data/PDB/CUUCGG/2KOC_5_C_A_0_00001.pdb 
# Loaded target data/PDB/CUUCGG/4u4u-pdb-bundle3_230_C_d_0_00000.pdb 
# Loading data/PDB/CUUCGG/4u4u-pdb-bundle3_230_C_d_0_00000.pdb 
# Loaded reference data/PDB/CUUCGG/2KOC_5_C_A_0_00001.pdb 
# Loaded target data/PDB/CUUCGG/1F7G_12_C_A_0_00012.pdb 
# Loading data/PDB/CUUCGG/1F7G_12_C_A_0_00012.pdb 
# Loaded reference data/PDB/CUUCGG/2KOC_5_C_A_0_00001.pdb 
# Loaded target data/PDB/CUUCGG/2HGH_32_C_B_0_00013.pdb 
# Loading data/PDB/CUUCGG/2HGH_32_C_B_0_00013.pdb 
# Loaded reference data/PDB/CUUCGG/2KOC_5_C_A_0_00001.pdb 
# Loaded target data/PDB/CUUCGG/1F7G_12_C_A_0_00013.pdb 
# Loading data/PDB/CUUCGG/1F7G_12_C_A_0_00013.pdb 
# Loaded reference data/PDB/CUUCGG/2KOC_5_C_A_0_00001.pdb 
# Loaded target data/PDB/CUUCGG/1vy7-pdb-bundle1_419_C_A_0_00000.pdb 
# Loading data/PDB/CUUCGG/1vy7-pdb-bundle1_419_C_A_0_00000.pdb 
# Loaded reference data/PDB/CUUCGG/2KOC_5_C_A_0_00001.pdb 
# Loaded target data/PDB/CUUCGG/5dox-pdb-bundle3_1691_C_A_0_00000.pdb 
# Loading data/PDB/CUUCGG/5dox-pdb-bundle3_1691_C_A_0_00000.pdb 
# Loaded reference data/PDB/CUUCGG/2KOC_5_C_A_0_00001.pdb 
# Loaded target data/PDB/CUUCGG/2HGH_32_C_B_0_00012.pdb 
# Loading data/PDB/CUUCGG/2HGH_32_C_B_0_00012.pdb 
# Loaded reference data/PDB/CUUCGG/2KOC_5_C_A_0_00001.pdb 
# Loaded target data/PDB/CUUCGG/2HGH_32_C_B_0_00006.pdb 
# Loading data/PDB/CUUCGG/2HGH_32_C_B_0_00006.pdb 
# Loaded reference data/PDB/CUUCGG/2KOC_5_C_A_0_00001.pdb 
# Loaded target data/PDB/CUUCGG/4v7y-pdb-bundle2_1691_C_A_0_00000.pdb 
# Loading data/PDB/CUUCGG/4v7y-pdb-bundle2_1691_C_A_0_00000.pdb 
# Loaded reference data/PDB/CUUCGG/2KOC_5_C_A_0_00001.pdb 
# Loaded target data/PDB/CUUCGG/4v67-pdb-bundle1_419_C_A_0_00000.pdb 
# Loading data/PDB/CUUCGG/4v67-pdb-bundle1_419_C_A_0_00000.pdb 
# Loaded reference data/PDB/CUUCGG/2KOC_5_C_A_0_00001.pdb 
# Loaded target data/PDB/CUUCGG/5ib8-pdb-bundle3_419_C_A_0_00000.pdb 
# Loading data/PDB/CUUCGG/5ib8-pdb-bundle3_419_C_A_0_00000.pdb 
# Loaded reference data/PDB/CUUCGG/2KOC_5_C_A_0_00001.pdb 
# Loaded target data/PDB/CUUCGG/1F7G_12_C_A_0_00007.pdb 
# Loading data/PDB/CUUCGG/1F7G_12_C_A_0_00007.pdb 
# Loaded reference data/PDB/CUUCGG/2KOC_5_C_A_0_00001.pdb 
# Loaded target data/PDB/CUUCGG/4v5e-pdb-bundle1_419_C_A_0_00000.pdb 
# Loading data/PDB/CUUCGG/4v5e-pdb-bundle1_419_C_A_0_00000.pdb 
# Loaded reference data/PDB/CUUCGG/2KOC_5_C_A_0_00001.pdb 
# Loaded target data/PDB/CUUCGG/1C0O_5_C_A_0_00014.pdb 
# Loading data/PDB/CUUCGG/1C0O_5_C_A_0_00014.pdb 
# Loaded reference data/PDB/CUUCGG/2KOC_5_C_A_0_00001.pdb 
# Loaded target data/PDB/CUUCGG/4u4n-pdb-bundle3_230_C_d_0_00000.pdb 
# Loading data/PDB/CUUCGG/4u4n-pdb-bundle3_230_C_d_0_00000.pdb 
# Loaded reference data/PDB/CUUCGG/2KOC_5_C_A_0_00001.pdb 
# Loaded target data/PDB/CUUCGG/1C0O_5_C_A_0_00000.pdb 
# Loading data/PDB/CUUCGG/1C0O_5_C_A_0_00000.pdb 
# Loaded reference data/PDB/CUUCGG/2KOC_5_C_A_0_00001.pdb 
# Loaded target data/PDB/CUUCGG/1HR0_419_C_A_0_00000.pdb 
# Loading data/PDB/CUUCGG/1HR0_419_C_A_0_00000.pdb 
# Loaded reference data/PDB/CUUCGG/2KOC_5_C_A_0_00001.pdb 
# Loaded target data/PDB/CUUCGG/5hau-pdb-bundle1_1737_C_A_0_00000.pdb 
# Loading data/PDB/CUUCGG/5hau-pdb-bundle1_1737_C_A_0_00000.pdb 
# Loaded reference data/PDB/CUUCGG/2KOC_5_C_A_0_00001.pdb 
# Loaded target data/PDB/CUUCGG/1Z31_270_C_A_0_00000.pdb 
# Loading data/PDB/CUUCGG/1Z31_270_C_A_0_00000.pdb 
# Loaded reference data/PDB/CUUCGG/2KOC_5_C_A_0_00001.pdb 
# Loaded target data/PDB/CUUCGG/1IKD_7_C_A_0_00022.pdb 
# Loading data/PDB/CUUCGG/1IKD_7_C_A_0_00022.pdb 
# Loaded reference data/PDB/CUUCGG/2KOC_5_C_A_0_00001.pdb 
# Loaded target data/PDB/CUUCGG/2L2K_19_C_A_0_00009.pdb 
# Loading data/PDB/CUUCGG/2L2K_19_C_A_0_00009.pdb 
# Loaded reference data/PDB/CUUCGG/2KOC_5_C_A_0_00001.pdb 
# Loaded target data/PDB/CUUCGG/2KOC_5_C_A_0_00014.pdb 
# Loading data/PDB/CUUCGG/2KOC_5_C_A_0_00014.pdb 
# Loaded reference data/PDB/CUUCGG/2KOC_5_C_A_0_00001.pdb 
# Loaded target data/PDB/CUUCGG/4v7w-pdb-bundle4_1691_C_A_0_00000.pdb 
# Loading data/PDB/CUUCGG/4v7w-pdb-bundle4_1691_C_A_0_00000.pdb 
# Loaded reference data/PDB/CUUCGG/2KOC_5_C_A_0_00001.pdb 
# Loaded target data/PDB/CUUCGG/4wqr-pdb-bundle1_419_C_A_0_00000.pdb 
# Loading data/PDB/CUUCGG/4wqr-pdb-bundle1_419_C_A_0_00000.pdb 
# Loaded reference data/PDB/CUUCGG/2KOC_5_C_A_0_00001.pdb 
# Loaded target data/PDB/CUUCGG/4zer-pdb-bundle3_1691_C_A_0_00000.pdb 
# Loading data/PDB/CUUCGG/4zer-pdb-bundle3_1691_C_A_0_00000.pdb 
# Loaded reference data/PDB/CUUCGG/2KOC_5_C_A_0_00001.pdb 
# Loaded target data/PDB/CUUCGG/2KOC_5_C_A_0_00000.pdb 
# Loading data/PDB/CUUCGG/2KOC_5_C_A_0_00000.pdb 
# Loaded reference data/PDB/CUUCGG/2KOC_5_C_A_0_00001.pdb 
# Loaded target data/PDB/CUUCGG/4ybb-pdb-bundle3_1691_C_A_0_00000.pdb 
# Loading data/PDB/CUUCGG/4ybb-pdb-bundle3_1691_C_A_0_00000.pdb 
# Loaded reference data/PDB/CUUCGG/2KOC_5_C_A_0_00001.pdb 
# Loaded target data/PDB/CUUCGG/1G70_54_C_A_0_00003.pdb 
# Loading data/PDB/CUUCGG/1G70_54_C_A_0_00003.pdb 
# Loaded reference data/PDB/CUUCGG/2KOC_5_C_A_0_00001.pdb 
# Loaded target data/PDB/CUUCGG/4v7t-pdb-bundle1_207_C_A_0_00000.pdb 
# Loading data/PDB/CUUCGG/4v7t-pdb-bundle1_207_C_A_0_00000.pdb 
# Loaded reference data/PDB/CUUCGG/2KOC_5_C_A_0_00001.pdb 
# Loaded target data/PDB/CUUCGG/5el4-pdb-bundle4_1691_C_A_0_00000.pdb 
# Loading data/PDB/CUUCGG/5el4-pdb-bundle4_1691_C_A_0_00000.pdb 
# Loaded reference data/PDB/CUUCGG/2KOC_5_C_A_0_00001.pdb 
# Loaded target data/PDB/CUUCGG/4DR3_419_C_A_0_00000.pdb 
# Loading data/PDB/CUUCGG/4DR3_419_C_A_0_00000.pdb 
# Loaded reference data/PDB/CUUCGG/2KOC_5_C_A_0_00001.pdb 
# Loaded target data/PDB/CUUCGG/4u52-pdb-bundle4_1923_C_I_0_00002.pdb 
# Loading data/PDB/CUUCGG/4u52-pdb-bundle4_1923_C_I_0_00002.pdb 
# Loaded reference data/PDB/CUUCGG/2KOC_5_C_A_0_00001.pdb 
# Loaded target data/PDB/CUUCGG/2UUB_419_C_A_0_00000.pdb 
# Loading data/PDB/CUUCGG/2UUB_419_C_A_0_00000.pdb 
# Loaded reference data/PDB/CUUCGG/2KOC_5_C_A_0_00001.pdb 
# Loaded target data/PDB/CUUCGG/4u27-pdb-bundle2_1691_C_A_0_00000.pdb 
# Loading data/PDB/CUUCGG/4u27-pdb-bundle2_1691_C_A_0_00000.pdb 
# Loaded reference data/PDB/CUUCGG/2KOC_5_C_A_0_00001.pdb 
# Loaded target data/PDB/CUUCGG/2D3O_1708_C_0_0_00000.pdb 
# Loading data/PDB/CUUCGG/2D3O_1708_C_0_0_00000.pdb 
# Loaded reference data/PDB/CUUCGG/2KOC_5_C_A_0_00001.pdb 
# Loaded target data/PDB/CUUCGG/5dgf-pdb-bundle2_1923_C_A_0_00002.pdb 
# Loading data/PDB/CUUCGG/5dgf-pdb-bundle2_1923_C_A_0_00002.pdb 
# Loaded reference data/PDB/CUUCGG/2KOC_5_C_A_0_00001.pdb 
# Loaded target data/PDB/CUUCGG/5vp2-pdb-bundle4_419_C_A_0_00000.pdb 
# Loading data/PDB/CUUCGG/5vp2-pdb-bundle4_419_C_A_0_00000.pdb 
# Loaded reference data/PDB/CUUCGG/2KOC_5_C_A_0_00001.pdb 
# Loaded target data/PDB/CUUCGG/1HNX_419_C_A_0_00000.pdb 
# Loading data/PDB/CUUCGG/1HNX_419_C_A_0_00000.pdb 
# Loaded reference data/PDB/CUUCGG/2KOC_5_C_A_0_00001.pdb 
# Loaded target data/PDB/CUUCGG/5j88-pdb-bundle1_419_C_b_0_00005.pdb 
# Loading data/PDB/CUUCGG/5j88-pdb-bundle1_419_C_b_0_00005.pdb 
# Loaded reference data/PDB/CUUCGG/2KOC_5_C_A_0_00001.pdb 
# Loaded target data/PDB/CUUCGG/5IEM_25_C_A_0_00008.pdb 
# Loading data/PDB/CUUCGG/5IEM_25_C_A_0_00008.pdb 
# Loaded reference data/PDB/CUUCGG/2KOC_5_C_A_0_00001.pdb 
# Loaded target data/PDB/CUUCGG/4v64-pdb-bundle3_1028_C_A_0_00002.pdb 
# Loading data/PDB/CUUCGG/4v64-pdb-bundle3_1028_C_A_0_00002.pdb 
# Loaded reference data/PDB/CUUCGG/2KOC_5_C_A_0_00001.pdb 
# Loaded target data/PDB/CUUCGG/6enu-pdb-bundle1_419_C_a_0_00001.pdb 
# Loading data/PDB/CUUCGG/6enu-pdb-bundle1_419_C_a_0_00001.pdb 
# Loaded reference data/PDB/CUUCGG/2KOC_5_C_A_0_00001.pdb 
# Loaded target data/PDB/CUUCGG/4v51-pdb-bundle3_419_C_A_0_00000.pdb 
# Loading data/PDB/CUUCGG/4v51-pdb-bundle3_419_C_A_0_00000.pdb 
# Loaded reference data/PDB/CUUCGG/2KOC_5_C_A_0_00001.pdb 
# Loaded target data/PDB/CUUCGG/4v9r-pdb-bundle2_1738_C_A_0_00000.pdb 
# Loading data/PDB/CUUCGG/4v9r-pdb-bundle2_1738_C_A_0_00000.pdb 
# Loaded reference data/PDB/CUUCGG/2KOC_5_C_A_0_00001.pdb 
# Loaded target data/PDB/CUUCGG/1M82_10_C_A_0_00017.pdb 
# Loading data/PDB/CUUCGG/1M82_10_C_A_0_00017.pdb 
# Loaded reference data/PDB/CUUCGG/2KOC_5_C_A_0_00001.pdb 
# Loaded target data/PDB/CUUCGG/1M82_10_C_A_0_00003.pdb 
# Loading data/PDB/CUUCGG/1M82_10_C_A_0_00003.pdb 
# Loaded reference data/PDB/CUUCGG/2KOC_5_C_A_0_00001.pdb 
# Loaded target data/PDB/CUUCGG/6HMA_2819_C_A_0_00001.pdb 
# Loading data/PDB/CUUCGG/6HMA_2819_C_A_0_00001.pdb 
# Loaded reference data/PDB/CUUCGG/2KOC_5_C_A_0_00001.pdb 
# Loaded target data/PDB/CUUCGG/1FYP_11_C_A_0_00007.pdb 
# Loading data/PDB/CUUCGG/1FYP_11_C_A_0_00007.pdb 
# Loaded reference data/PDB/CUUCGG/2KOC_5_C_A_0_00001.pdb 
# Loaded target data/PDB/CUUCGG/6hd7-pdb-bundle1_1923_C_1_0_00002.pdb 
# Loading data/PDB/CUUCGG/6hd7-pdb-bundle1_1923_C_1_0_00002.pdb 
# Loaded reference data/PDB/CUUCGG/2KOC_5_C_A_0_00001.pdb 
# Loaded target data/PDB/CUUCGG/1FYP_11_C_A_0_00013.pdb 
# Loading data/PDB/CUUCGG/1FYP_11_C_A_0_00013.pdb 
# Loaded reference data/PDB/CUUCGG/2KOC_5_C_A_0_00001.pdb 
# Loaded target data/PDB/CUUCGG/1AUD_28_C_B_0_00013.pdb 
# Loading data/PDB/CUUCGG/1AUD_28_C_B_0_00013.pdb 
# Loaded reference data/PDB/CUUCGG/2KOC_5_C_A_0_00001.pdb 
# Loaded target data/PDB/CUUCGG/1EBS_12_C_A_0_00000.pdb 
# Loading data/PDB/CUUCGG/1EBS_12_C_A_0_00000.pdb 
# Loaded reference data/PDB/CUUCGG/2KOC_5_C_A_0_00001.pdb 
# Loaded target data/PDB/CUUCGG/5WNS_419_C_A_0_00000.pdb 
# Loading data/PDB/CUUCGG/5WNS_419_C_A_0_00000.pdb 
# Loaded reference data/PDB/CUUCGG/2KOC_5_C_A_0_00001.pdb 
# Loaded target data/PDB/CUUCGG/1AUD_28_C_B_0_00007.pdb 
# Loading data/PDB/CUUCGG/1AUD_28_C_B_0_00007.pdb 
# Loaded reference data/PDB/CUUCGG/2KOC_5_C_A_0_00001.pdb 
# Loaded target data/PDB/CUUCGG/6gsj-pdb-bundle3_419_C_A_0_00000.pdb 
# Loading data/PDB/CUUCGG/6gsj-pdb-bundle3_419_C_A_0_00000.pdb 
# Loaded reference data/PDB/CUUCGG/2KOC_5_C_A_0_00001.pdb 
# Loaded target data/PDB/CUUCGG/4v5q-pdb-bundle2_1691_C_A_0_00000.pdb 
# Loading data/PDB/CUUCGG/4v5q-pdb-bundle2_1691_C_A_0_00000.pdb 
# Loaded reference data/PDB/CUUCGG/2KOC_5_C_A_0_00001.pdb 
# Loaded target data/PDB/CUUCGG/1RAW_21_C_A_0_00003.pdb 
# Loading data/PDB/CUUCGG/1RAW_21_C_A_0_00003.pdb 
# Loaded reference data/PDB/CUUCGG/2KOC_5_C_A_0_00001.pdb 
# Loaded target data/PDB/CUUCGG/2N6W_32_C_A_0_00006.pdb 
# Loading data/PDB/CUUCGG/2N6W_32_C_A_0_00006.pdb 
# Loaded reference data/PDB/CUUCGG/2KOC_5_C_A_0_00001.pdb 
# Loaded target data/PDB/CUUCGG/6DNR_27_C_A_0_00000.pdb 
# Loading data/PDB/CUUCGG/6DNR_27_C_A_0_00000.pdb 
# Loaded reference data/PDB/CUUCGG/2KOC_5_C_A_0_00001.pdb 
# Loaded target data/PDB/CUUCGG/5mdv-pdb-bundle1_1449_C_2_0_00004.pdb 
# Loading data/PDB/CUUCGG/5mdv-pdb-bundle1_1449_C_2_0_00004.pdb 
# Loaded reference data/PDB/CUUCGG/2KOC_5_C_A_0_00001.pdb 
# Loaded target data/PDB/CUUCGG/2N6T_19_C_A_0_00001.pdb 
# Loading data/PDB/CUUCGG/2N6T_19_C_A_0_00001.pdb 
# Loaded reference data/PDB/CUUCGG/2KOC_5_C_A_0_00001.pdb 
# Loaded target data/PDB/CUUCGG/1ULL_14_C_A_0_00000.pdb 
# Loading data/PDB/CUUCGG/1ULL_14_C_A_0_00000.pdb 
# Loaded reference data/PDB/CUUCGG/2KOC_5_C_A_0_00001.pdb 
# Loaded target data/PDB/CUUCGG/1F7I_12_C_A_0_00003.pdb 
# Loading data/PDB/CUUCGG/1F7I_12_C_A_0_00003.pdb 
# Loaded reference data/PDB/CUUCGG/2KOC_5_C_A_0_00001.pdb 
# Loaded target data/PDB/CUUCGG/1P5M_28_C_A_0_00002.pdb 
# Loading data/PDB/CUUCGG/1P5M_28_C_A_0_00002.pdb 
# Loaded reference data/PDB/CUUCGG/2KOC_5_C_A_0_00001.pdb 
# Loaded target data/PDB/CUUCGG/5uym-pdb-bundle1_1449_C_0_0_00003.pdb 
# Loading data/PDB/CUUCGG/5uym-pdb-bundle1_1449_C_0_0_00003.pdb 
# Loaded reference data/PDB/CUUCGG/2KOC_5_C_A_0_00001.pdb 
# Loaded target data/PDB/CUUCGG/3DLL_1708_C_X_0_00000.pdb 
# Loading data/PDB/CUUCGG/3DLL_1708_C_X_0_00000.pdb 
# Loaded reference data/PDB/CUUCGG/2KOC_5_C_A_0_00001.pdb 
# Loaded target data/PDB/CUUCGG/1EKZ_13_C_B_0_00032.pdb 
# Loading data/PDB/CUUCGG/1EKZ_13_C_B_0_00032.pdb 
# Loaded reference data/PDB/CUUCGG/2KOC_5_C_A_0_00001.pdb 
# Loaded target data/PDB/CUUCGG/2L2J_19_C_A_0_00013.pdb 
# Loading data/PDB/CUUCGG/2L2J_19_C_A_0_00013.pdb 
# Loaded reference data/PDB/CUUCGG/2KOC_5_C_A_0_00001.pdb 
# Loaded target data/PDB/CUUCGG/1EKZ_13_C_B_0_00026.pdb 
# Loading data/PDB/CUUCGG/1EKZ_13_C_B_0_00026.pdb 
# Loaded reference data/PDB/CUUCGG/2KOC_5_C_A_0_00001.pdb 
# Loaded target data/PDB/CUUCGG/2L2J_19_C_A_0_00007.pdb 
# Loading data/PDB/CUUCGG/2L2J_19_C_A_0_00007.pdb 
# Loaded reference data/PDB/CUUCGG/2KOC_5_C_A_0_00001.pdb 
# Loaded target data/PDB/CUUCGG/1NBS_152_C_B_0_00000.pdb 
# Loading data/PDB/CUUCGG/1NBS_152_C_B_0_00000.pdb 
# Loaded reference data/PDB/CUUCGG/2KOC_5_C_A_0_00001.pdb 
# Loaded target data/PDB/CUUCGG/2L2J_19_C_A_0_00011.pdb 
# Loading data/PDB/CUUCGG/2L2J_19_C_A_0_00011.pdb 
# Loaded reference data/PDB/CUUCGG/2KOC_5_C_A_0_00001.pdb 
# Loaded target data/PDB/CUUCGG/1EKZ_13_C_B_0_00030.pdb 
# Loading data/PDB/CUUCGG/1EKZ_13_C_B_0_00030.pdb 
# Loaded reference data/PDB/CUUCGG/2KOC_5_C_A_0_00001.pdb 
# Loaded target data/PDB/CUUCGG/5WNP_419_C_A_0_00000.pdb 
# Loading data/PDB/CUUCGG/5WNP_419_C_A_0_00000.pdb 
# Loaded reference data/PDB/CUUCGG/2KOC_5_C_A_0_00001.pdb 
# Loaded target data/PDB/CUUCGG/2L2J_19_C_A_0_00005.pdb 
# Loading data/PDB/CUUCGG/2L2J_19_C_A_0_00005.pdb 
# Loaded reference data/PDB/CUUCGG/2KOC_5_C_A_0_00001.pdb 
# Loaded target data/PDB/CUUCGG/1EKZ_13_C_B_0_00024.pdb 
# Loading data/PDB/CUUCGG/1EKZ_13_C_B_0_00024.pdb 
# Loaded reference data/PDB/CUUCGG/2KOC_5_C_A_0_00001.pdb 
# Loaded target data/PDB/CUUCGG/1ULL_14_C_A_0_00002.pdb 
# Loading data/PDB/CUUCGG/1ULL_14_C_A_0_00002.pdb 
# Loaded reference data/PDB/CUUCGG/2KOC_5_C_A_0_00001.pdb 
# Loaded target data/PDB/CUUCGG/2N6T_19_C_A_0_00003.pdb 
# Loading data/PDB/CUUCGG/2N6T_19_C_A_0_00003.pdb 
# Loaded reference data/PDB/CUUCGG/2KOC_5_C_A_0_00001.pdb 
# Loaded target data/PDB/CUUCGG/4v7u-pdb-bundle1_1449_C_A_0_00003.pdb 
# Loading data/PDB/CUUCGG/4v7u-pdb-bundle1_1449_C_A_0_00003.pdb 
# Loaded reference data/PDB/CUUCGG/2KOC_5_C_A_0_00001.pdb 
# Loaded target data/PDB/CUUCGG/1EKZ_13_C_B_0_00018.pdb 
# Loading data/PDB/CUUCGG/1EKZ_13_C_B_0_00018.pdb 
# Loaded reference data/PDB/CUUCGG/2KOC_5_C_A_0_00001.pdb 
# Loaded target data/PDB/CUUCGG/1F7I_12_C_A_0_00001.pdb 
# Loading data/PDB/CUUCGG/1F7I_12_C_A_0_00001.pdb 
# Loaded reference data/PDB/CUUCGG/2KOC_5_C_A_0_00001.pdb 
# Loaded target data/PDB/CUUCGG/4y4p-pdb-bundle3_1691_C_A_0_00000.pdb 
# Loading data/PDB/CUUCGG/4y4p-pdb-bundle3_1691_C_A_0_00000.pdb 
# Loaded reference data/PDB/CUUCGG/2KOC_5_C_A_0_00001.pdb 
# Loaded target data/PDB/CUUCGG/4v9s-pdb-bundle1_419_C_A_0_00000.pdb 
# Loading data/PDB/CUUCGG/4v9s-pdb-bundle1_419_C_A_0_00000.pdb 
# Loaded reference data/PDB/CUUCGG/2KOC_5_C_A_0_00001.pdb 
# Loaded target data/PDB/CUUCGG/1RAW_21_C_A_0_00001.pdb 
# Loading data/PDB/CUUCGG/1RAW_21_C_A_0_00001.pdb 
# Loaded reference data/PDB/CUUCGG/2KOC_5_C_A_0_00001.pdb 
# Loaded target data/PDB/CUUCGG/2N6W_32_C_A_0_00004.pdb 
# Loading data/PDB/CUUCGG/2N6W_32_C_A_0_00004.pdb 
# Loaded reference data/PDB/CUUCGG/2KOC_5_C_A_0_00001.pdb 
# Loaded target data/PDB/CUUCGG/1EBS_12_C_A_0_00002.pdb 
# Loading data/PDB/CUUCGG/1EBS_12_C_A_0_00002.pdb 
# Loaded reference data/PDB/CUUCGG/2KOC_5_C_A_0_00001.pdb 
# Loaded target data/PDB/CUUCGG/4v9o-pdb-bundle2_1449_C_A_0_00003.pdb 
# Loading data/PDB/CUUCGG/4v9o-pdb-bundle2_1449_C_A_0_00003.pdb 
# Loaded reference data/PDB/CUUCGG/2KOC_5_C_A_0_00001.pdb 
# Loaded target data/PDB/CUUCGG/1FYP_11_C_A_0_00005.pdb 
# Loading data/PDB/CUUCGG/1FYP_11_C_A_0_00005.pdb 
# Loaded reference data/PDB/CUUCGG/2KOC_5_C_A_0_00001.pdb 
# Loaded target data/PDB/CUUCGG/4v52-pdb-bundle1_1028_C_A_0_00002.pdb 
# Loading data/PDB/CUUCGG/4v52-pdb-bundle1_1028_C_A_0_00002.pdb 
# Loaded reference data/PDB/CUUCGG/2KOC_5_C_A_0_00001.pdb 
# Loaded target data/PDB/CUUCGG/1FYP_11_C_A_0_00011.pdb 
# Loading data/PDB/CUUCGG/1FYP_11_C_A_0_00011.pdb 
# Loaded reference data/PDB/CUUCGG/2KOC_5_C_A_0_00001.pdb 
# Loaded target data/PDB/CUUCGG/4v57-pdb-bundle3_1028_C_A_0_00002.pdb 
# Loading data/PDB/CUUCGG/4v57-pdb-bundle3_1028_C_A_0_00002.pdb 
# Loaded reference data/PDB/CUUCGG/2KOC_5_C_A_0_00001.pdb 
# Loaded target data/PDB/CUUCGG/1M82_10_C_A_0_00029.pdb 
# Loading data/PDB/CUUCGG/1M82_10_C_A_0_00029.pdb 
# Loaded reference data/PDB/CUUCGG/2KOC_5_C_A_0_00001.pdb 
# Loaded target data/PDB/CUUCGG/1M82_10_C_A_0_00015.pdb 
# Loading data/PDB/CUUCGG/1M82_10_C_A_0_00015.pdb 
# Loaded reference data/PDB/CUUCGG/2KOC_5_C_A_0_00001.pdb 
# Loaded target data/PDB/CUUCGG/4v6f-pdb-bundle2_419_C_A_0_00000.pdb 
# Loading data/PDB/CUUCGG/4v6f-pdb-bundle2_419_C_A_0_00000.pdb 
# Loaded reference data/PDB/CUUCGG/2KOC_5_C_A_0_00001.pdb 
# Loaded target data/PDB/CUUCGG/3CCS_1769_C_0_0_00000.pdb 
# Loading data/PDB/CUUCGG/3CCS_1769_C_0_0_00000.pdb 
# Loaded reference data/PDB/CUUCGG/2KOC_5_C_A_0_00001.pdb 
# Loaded target data/PDB/CUUCGG/1M82_10_C_A_0_00001.pdb 
# Loading data/PDB/CUUCGG/1M82_10_C_A_0_00001.pdb 
# Loaded reference data/PDB/CUUCGG/2KOC_5_C_A_0_00001.pdb 
# Loaded target data/PDB/CUUCGG/1J5E_419_C_A_0_00000.pdb 
# Loading data/PDB/CUUCGG/1J5E_419_C_A_0_00000.pdb 
# Loaded reference data/PDB/CUUCGG/2KOC_5_C_A_0_00001.pdb 
# Loaded target data/PDB/CUUCGG/5ibb-pdb-bundle4_1691_C_A_0_00000.pdb 
# Loading data/PDB/CUUCGG/5ibb-pdb-bundle4_1691_C_A_0_00000.pdb 
# Loaded reference data/PDB/CUUCGG/2KOC_5_C_A_0_00001.pdb 
# Loaded target data/PDB/CUUCGG/4u24-pdb-bundle3_419_C_A_0_00001.pdb 
# Loading data/PDB/CUUCGG/4u24-pdb-bundle3_419_C_A_0_00001.pdb 
# Loaded reference data/PDB/CUUCGG/2KOC_5_C_A_0_00001.pdb 
# Loaded target data/PDB/CUUCGG/1G70_54_C_A_0_00001.pdb 
# Loading data/PDB/CUUCGG/1G70_54_C_A_0_00001.pdb 
# Loaded reference data/PDB/CUUCGG/2KOC_5_C_A_0_00001.pdb 
# Loaded target data/PDB/CUUCGG/6FAI_230_C_2_0_00000.pdb 
# Loading data/PDB/CUUCGG/6FAI_230_C_2_0_00000.pdb 
# Loaded reference data/PDB/CUUCGG/2KOC_5_C_A_0_00001.pdb 
# Loaded target data/PDB/CUUCGG/5j91-pdb-bundle1_1449_C_A_0_00003.pdb 
# Loading data/PDB/CUUCGG/5j91-pdb-bundle1_1449_C_A_0_00003.pdb 
# Loaded reference data/PDB/CUUCGG/2KOC_5_C_A_0_00001.pdb 
# Loaded target data/PDB/CUUCGG/5doy-pdb-bundle1_1691_C_A_0_00000.pdb 
# Loading data/PDB/CUUCGG/5doy-pdb-bundle1_1691_C_A_0_00000.pdb 
# Loaded reference data/PDB/CUUCGG/2KOC_5_C_A_0_00001.pdb 
# Loaded target data/PDB/CUUCGG/2KOC_5_C_A_0_00016.pdb 
# Loading data/PDB/CUUCGG/2KOC_5_C_A_0_00016.pdb 
# Loaded reference data/PDB/CUUCGG/2KOC_5_C_A_0_00001.pdb 
# Loaded target data/PDB/CUUCGG/4u3n-pdb-bundle4_1923_C_I_0_00002.pdb 
# Loading data/PDB/CUUCGG/4u3n-pdb-bundle4_1923_C_I_0_00002.pdb 
# Loaded reference data/PDB/CUUCGG/2KOC_5_C_A_0_00001.pdb 
# Loaded target data/PDB/CUUCGG/2KOC_5_C_A_0_00002.pdb 
# Loading data/PDB/CUUCGG/2KOC_5_C_A_0_00002.pdb 
# Loaded reference data/PDB/CUUCGG/2KOC_5_C_A_0_00001.pdb 
# Loaded target data/PDB/CUUCGG/5H1S_1727_C_A_0_00001.pdb 
# Loading data/PDB/CUUCGG/5H1S_1727_C_A_0_00001.pdb 
# Loaded reference data/PDB/CUUCGG/2KOC_5_C_A_0_00001.pdb 
# Loaded target data/PDB/CUUCGG/4LF6_419_C_A_0_00000.pdb 
# Loading data/PDB/CUUCGG/4LF6_419_C_A_0_00000.pdb 
# Loaded reference data/PDB/CUUCGG/2KOC_5_C_A_0_00001.pdb 
# Loaded target data/PDB/CUUCGG/5m1j-pdb-bundle2_1923_C_A_0_00002.pdb 
# Loading data/PDB/CUUCGG/5m1j-pdb-bundle2_1923_C_A_0_00002.pdb 
# Loaded reference data/PDB/CUUCGG/2KOC_5_C_A_0_00001.pdb 
# Loaded target data/PDB/CUUCGG/5fdv-pdb-bundle4_419_C_A_0_00000.pdb 
# Loading data/PDB/CUUCGG/5fdv-pdb-bundle4_419_C_A_0_00000.pdb 
# Loaded reference data/PDB/CUUCGG/2KOC_5_C_A_0_00001.pdb 
# Loaded target data/PDB/CUUCGG/6cfj-pdb-bundle2_419_C_A_0_00000.pdb 
# Loading data/PDB/CUUCGG/6cfj-pdb-bundle2_419_C_A_0_00000.pdb 
# Loaded reference data/PDB/CUUCGG/2KOC_5_C_A_0_00001.pdb 
# Loaded target data/PDB/CUUCGG/1IKD_7_C_A_0_00020.pdb 
# Loading data/PDB/CUUCGG/1IKD_7_C_A_0_00020.pdb 
# Loaded reference data/PDB/CUUCGG/2KOC_5_C_A_0_00001.pdb 
# Loaded target data/PDB/CUUCGG/1Z31_270_C_A_0_00002.pdb 
# Loading data/PDB/CUUCGG/1Z31_270_C_A_0_00002.pdb 
# Loaded reference data/PDB/CUUCGG/2KOC_5_C_A_0_00001.pdb 
# Loaded target data/PDB/CUUCGG/4z8c-pdb-bundle1_1737_C_A_0_00000.pdb 
# Loading data/PDB/CUUCGG/4z8c-pdb-bundle1_1737_C_A_0_00000.pdb 
# Loaded reference data/PDB/CUUCGG/2KOC_5_C_A_0_00001.pdb 
# Loaded target data/PDB/CUUCGG/4u56-pdb-bundle4_1923_C_I_0_00002.pdb 
# Loading data/PDB/CUUCGG/4u56-pdb-bundle4_1923_C_I_0_00002.pdb 
# Loaded reference data/PDB/CUUCGG/2KOC_5_C_A_0_00001.pdb 
# Loaded target data/PDB/CUUCGG/1C0O_5_C_A_0_00016.pdb 
# Loading data/PDB/CUUCGG/1C0O_5_C_A_0_00016.pdb 
# Loaded reference data/PDB/CUUCGG/2KOC_5_C_A_0_00001.pdb 
# Loaded target data/PDB/CUUCGG/1C0O_5_C_A_0_00002.pdb 
# Loading data/PDB/CUUCGG/1C0O_5_C_A_0_00002.pdb 
# Loaded reference data/PDB/CUUCGG/2KOC_5_C_A_0_00001.pdb 
# Loaded target data/PDB/CUUCGG/4v6c-pdb-bundle3_207_C_N_0_00000.pdb 
# Loading data/PDB/CUUCGG/4v6c-pdb-bundle3_207_C_N_0_00000.pdb 
# Loaded reference data/PDB/CUUCGG/2KOC_5_C_A_0_00001.pdb 
# Loaded target data/PDB/CUUCGG/2HGH_32_C_B_0_00010.pdb 
# Loading data/PDB/CUUCGG/2HGH_32_C_B_0_00010.pdb 
# Loaded reference data/PDB/CUUCGG/2KOC_5_C_A_0_00001.pdb 
# Loaded target data/PDB/CUUCGG/4v7s-pdb-bundle4_1691_C_A_0_00000.pdb 
# Loading data/PDB/CUUCGG/4v7s-pdb-bundle4_1691_C_A_0_00000.pdb 
# Loaded reference data/PDB/CUUCGG/2KOC_5_C_A_0_00001.pdb 
# Loaded target data/PDB/CUUCGG/4wro-pdb-bundle1_419_C_A_0_00000.pdb 
# Loading data/PDB/CUUCGG/4wro-pdb-bundle1_419_C_A_0_00000.pdb 
# Loaded reference data/PDB/CUUCGG/2KOC_5_C_A_0_00001.pdb 
# Loaded target data/PDB/CUUCGG/5v8i-pdb-bundle4_419_C_A_0_00000.pdb 
# Loading data/PDB/CUUCGG/5v8i-pdb-bundle4_419_C_A_0_00000.pdb 
# Loaded reference data/PDB/CUUCGG/2KOC_5_C_A_0_00001.pdb 
# Loaded target data/PDB/CUUCGG/1F7G_12_C_A_0_00011.pdb 
# Loading data/PDB/CUUCGG/1F7G_12_C_A_0_00011.pdb 
# Loaded reference data/PDB/CUUCGG/2KOC_5_C_A_0_00001.pdb 
# Loaded target data/PDB/CUUCGG/1F7G_12_C_A_0_00005.pdb 
# Loading data/PDB/CUUCGG/1F7G_12_C_A_0_00005.pdb 
# Loaded reference data/PDB/CUUCGG/2KOC_5_C_A_0_00001.pdb 
# Loaded target data/PDB/CUUCGG/2HGH_32_C_B_0_00004.pdb 
# Loading data/PDB/CUUCGG/2HGH_32_C_B_0_00004.pdb 
# Loaded reference data/PDB/CUUCGG/2KOC_5_C_A_0_00001.pdb 
# Loaded target data/PDB/CUUCGG/3CCV_1769_C_0_0_00000.pdb 
# Loading data/PDB/CUUCGG/3CCV_1769_C_0_0_00000.pdb 
# Loaded reference data/PDB/CUUCGG/2KOC_5_C_A_0_00001.pdb 
# Loaded target data/PDB/CUUCGG/1F7G_12_C_A_0_00004.pdb 
# Loading data/PDB/CUUCGG/1F7G_12_C_A_0_00004.pdb 
# Loaded reference data/PDB/CUUCGG/2KOC_5_C_A_0_00001.pdb 
# Loaded target data/PDB/CUUCGG/2HGH_32_C_B_0_00005.pdb 
# Loading data/PDB/CUUCGG/2HGH_32_C_B_0_00005.pdb 
# Loaded reference data/PDB/CUUCGG/2KOC_5_C_A_0_00001.pdb 
# Loaded target data/PDB/CUUCGG/5j7l-pdb-bundle1_419_C_A_0_00001.pdb 
# Loading data/PDB/CUUCGG/5j7l-pdb-bundle1_419_C_A_0_00001.pdb 
# Loaded reference data/PDB/CUUCGG/2KOC_5_C_A_0_00001.pdb 
# Loaded target data/PDB/CUUCGG/2HGH_32_C_B_0_00011.pdb 
# Loading data/PDB/CUUCGG/2HGH_32_C_B_0_00011.pdb 
# Loaded reference data/PDB/CUUCGG/2KOC_5_C_A_0_00001.pdb 
# Loaded target data/PDB/CUUCGG/1F7G_12_C_A_0_00010.pdb 
# Loading data/PDB/CUUCGG/1F7G_12_C_A_0_00010.pdb 
# Loaded reference data/PDB/CUUCGG/2KOC_5_C_A_0_00001.pdb 
# Loaded target data/PDB/CUUCGG/5vpo-pdb-bundle4_1691_C_A_0_00000.pdb 
# Loading data/PDB/CUUCGG/5vpo-pdb-bundle4_1691_C_A_0_00000.pdb 
# Loaded reference data/PDB/CUUCGG/2KOC_5_C_A_0_00001.pdb 
# Loaded target data/PDB/CUUCGG/1C0O_5_C_A_0_00003.pdb 
# Loading data/PDB/CUUCGG/1C0O_5_C_A_0_00003.pdb 
# Loaded reference data/PDB/CUUCGG/2KOC_5_C_A_0_00001.pdb 
# Loaded target data/PDB/CUUCGG/6DLR_27_C_A_0_00000.pdb 
# Loading data/PDB/CUUCGG/6DLR_27_C_A_0_00000.pdb 
# Loaded reference data/PDB/CUUCGG/2KOC_5_C_A_0_00001.pdb 
# Loaded target data/PDB/CUUCGG/4v54-pdb-bundle2_1691_C_A_0_00000.pdb 
# Loading data/PDB/CUUCGG/4v54-pdb-bundle2_1691_C_A_0_00000.pdb 
# Loaded reference data/PDB/CUUCGG/2KOC_5_C_A_0_00001.pdb 
# Loaded target data/PDB/CUUCGG/1C0O_5_C_A_0_00017.pdb 
# Loading data/PDB/CUUCGG/1C0O_5_C_A_0_00017.pdb 
# Loaded reference data/PDB/CUUCGG/2KOC_5_C_A_0_00001.pdb 
# Loaded target data/PDB/CUUCGG/5o2r-pdb-bundle1_1691_C_A_0_00000.pdb 
# Loading data/PDB/CUUCGG/5o2r-pdb-bundle1_1691_C_A_0_00000.pdb 
# Loaded reference data/PDB/CUUCGG/2KOC_5_C_A_0_00001.pdb 
# Loaded target data/PDB/CUUCGG/1Z31_270_C_A_0_00003.pdb 
# Loading data/PDB/CUUCGG/1Z31_270_C_A_0_00003.pdb 
# Loaded reference data/PDB/CUUCGG/2KOC_5_C_A_0_00001.pdb 
# Loaded target data/PDB/CUUCGG/4y4o-pdb-bundle2_419_C_A_0_00000.pdb 
# Loading data/PDB/CUUCGG/4y4o-pdb-bundle2_419_C_A_0_00000.pdb 
# Loaded reference data/PDB/CUUCGG/2KOC_5_C_A_0_00001.pdb 
# Loaded target data/PDB/CUUCGG/4v54-pdb-bundle3_1028_C_K_0_00002.pdb 
# Loading data/PDB/CUUCGG/4v54-pdb-bundle3_1028_C_K_0_00002.pdb 
# Loaded reference data/PDB/CUUCGG/2KOC_5_C_A_0_00001.pdb 
# Loaded target data/PDB/CUUCGG/4v9p-pdb-bundle1_1691_C_A_0_00000.pdb 
# Loading data/PDB/CUUCGG/4v9p-pdb-bundle1_1691_C_A_0_00000.pdb 
# Loaded reference data/PDB/CUUCGG/2KOC_5_C_A_0_00001.pdb 
# Loaded target data/PDB/CUUCGG/4wra-pdb-bundle4_1691_C_A_0_00000.pdb 
# Loading data/PDB/CUUCGG/4wra-pdb-bundle4_1691_C_A_0_00000.pdb 
# Loaded reference data/PDB/CUUCGG/2KOC_5_C_A_0_00001.pdb 
# Loaded target data/PDB/CUUCGG/1IKD_7_C_A_0_00009.pdb 
# Loading data/PDB/CUUCGG/1IKD_7_C_A_0_00009.pdb 
# Loaded reference data/PDB/CUUCGG/2KOC_5_C_A_0_00001.pdb 
# Loaded target data/PDB/CUUCGG/2KOC_5_C_A_0_00003.pdb 
# Loading data/PDB/CUUCGG/2KOC_5_C_A_0_00003.pdb 
# Loaded reference data/PDB/CUUCGG/2KOC_5_C_A_0_00001.pdb 
# Loaded target data/PDB/CUUCGG/2KOC_5_C_A_0_00017.pdb 
# Loading data/PDB/CUUCGG/2KOC_5_C_A_0_00017.pdb 
# Loaded reference data/PDB/CUUCGG/2KOC_5_C_A_0_00001.pdb 
# Loaded target data/PDB/CUUCGG/4v51-pdb-bundle1_419_C_A_0_00000.pdb 
# Loading data/PDB/CUUCGG/4v51-pdb-bundle1_419_C_A_0_00000.pdb 
# Loaded reference data/PDB/CUUCGG/2KOC_5_C_A_0_00001.pdb 
# Loaded target data/PDB/CUUCGG/1G70_54_C_A_0_00000.pdb 
# Loading data/PDB/CUUCGG/1G70_54_C_A_0_00000.pdb 
# Loaded reference data/PDB/CUUCGG/2KOC_5_C_A_0_00001.pdb 
# Loaded target data/PDB/CUUCGG/5j30-pdb-bundle4_419_C_A_0_00000.pdb 
# Loading data/PDB/CUUCGG/5j30-pdb-bundle4_419_C_A_0_00000.pdb 
# Loaded reference data/PDB/CUUCGG/2KOC_5_C_A_0_00001.pdb 
# Loaded target data/PDB/CUUCGG/5WNQ_419_C_A_0_00000.pdb 
# Loading data/PDB/CUUCGG/5WNQ_419_C_A_0_00000.pdb 
# Loaded reference data/PDB/CUUCGG/2KOC_5_C_A_0_00001.pdb 
# Loaded target data/PDB/CUUCGG/5xy3-pdb-bundle1_1588_C_1_0_00000.pdb 
# Loading data/PDB/CUUCGG/5xy3-pdb-bundle1_1588_C_1_0_00000.pdb 
# Loaded reference data/PDB/CUUCGG/2KOC_5_C_A_0_00001.pdb 
# Loaded target data/PDB/CUUCGG/2HW8_15_C_B_0_00000.pdb 
# Loading data/PDB/CUUCGG/2HW8_15_C_B_0_00000.pdb 
# Loaded reference data/PDB/CUUCGG/2KOC_5_C_A_0_00001.pdb 
# Loaded target data/PDB/CUUCGG/4v7j-pdb-bundle2_1691_C_A_0_00000.pdb 
# Loading data/PDB/CUUCGG/4v7j-pdb-bundle2_1691_C_A_0_00000.pdb 
# Loaded reference data/PDB/CUUCGG/2KOC_5_C_A_0_00001.pdb 
# Loaded target data/PDB/CUUCGG/5LYS_48_C_B_0_00001.pdb 
# Loading data/PDB/CUUCGG/5LYS_48_C_B_0_00001.pdb 
# Loaded reference data/PDB/CUUCGG/2KOC_5_C_A_0_00001.pdb 
# Loaded target data/PDB/CUUCGG/4wro-pdb-bundle2_1691_C_A_0_00000.pdb 
# Loading data/PDB/CUUCGG/4wro-pdb-bundle2_1691_C_A_0_00000.pdb 
# Loaded reference data/PDB/CUUCGG/2KOC_5_C_A_0_00001.pdb 
# Loaded target data/PDB/CUUCGG/6h4n-pdb-bundle2_1691_C_A_0_00000.pdb 
# Loading data/PDB/CUUCGG/6h4n-pdb-bundle2_1691_C_A_0_00000.pdb 
# Loaded reference data/PDB/CUUCGG/2KOC_5_C_A_0_00001.pdb 
# Loaded target data/PDB/CUUCGG/4v63-pdb-bundle2_1691_C_A_0_00000.pdb 
# Loading data/PDB/CUUCGG/4v63-pdb-bundle2_1691_C_A_0_00000.pdb 
# Loaded reference data/PDB/CUUCGG/2KOC_5_C_A_0_00001.pdb 
# Loaded target data/PDB/CUUCGG/4v7t-pdb-bundle3_207_C_A_0_00000.pdb 
# Loading data/PDB/CUUCGG/4v7t-pdb-bundle3_207_C_A_0_00000.pdb 
# Loaded reference data/PDB/CUUCGG/2KOC_5_C_A_0_00001.pdb 
# Loaded target data/PDB/CUUCGG/1M82_10_C_A_0_00000.pdb 
# Loading data/PDB/CUUCGG/1M82_10_C_A_0_00000.pdb 
# Loaded reference data/PDB/CUUCGG/2KOC_5_C_A_0_00001.pdb 
# Loaded target data/PDB/CUUCGG/5dfe-pdb-bundle2_419_C_F_0_00000.pdb 
# Loading data/PDB/CUUCGG/5dfe-pdb-bundle2_419_C_F_0_00000.pdb 
# Loaded reference data/PDB/CUUCGG/2KOC_5_C_A_0_00001.pdb 
# Loaded target data/PDB/CUUCGG/1M82_10_C_A_0_00014.pdb 
# Loading data/PDB/CUUCGG/1M82_10_C_A_0_00014.pdb 
# Loaded reference data/PDB/CUUCGG/2KOC_5_C_A_0_00001.pdb 
# Loaded target data/PDB/CUUCGG/5LYU_48_C_A_0_00000.pdb 
# Loading data/PDB/CUUCGG/5LYU_48_C_A_0_00000.pdb 
# Loaded reference data/PDB/CUUCGG/2KOC_5_C_A_0_00001.pdb 
# Loaded target data/PDB/CUUCGG/5czp-pdb-bundle3_1691_C_F_0_00000.pdb 
# Loading data/PDB/CUUCGG/5czp-pdb-bundle3_1691_C_F_0_00000.pdb 
# Loaded reference data/PDB/CUUCGG/2KOC_5_C_A_0_00001.pdb 
# Loaded target data/PDB/CUUCGG/4v6c-pdb-bundle3_1449_C_N_0_00003.pdb 
# Loading data/PDB/CUUCGG/4v6c-pdb-bundle3_1449_C_N_0_00003.pdb 
# Loaded reference data/PDB/CUUCGG/2KOC_5_C_A_0_00001.pdb 
# Loaded target data/PDB/CUUCGG/4u25-pdb-bundle1_1028_C_A_0_00002.pdb 
# Loading data/PDB/CUUCGG/4u25-pdb-bundle1_1028_C_A_0_00002.pdb 
# Loaded reference data/PDB/CUUCGG/2KOC_5_C_A_0_00001.pdb 
# Loaded target data/PDB/CUUCGG/4v87-pdb-bundle3_1691_C_R_0_00000.pdb 
# Loading data/PDB/CUUCGG/4v87-pdb-bundle3_1691_C_R_0_00000.pdb 
# Loaded reference data/PDB/CUUCGG/2KOC_5_C_A_0_00001.pdb 
# Loaded target data/PDB/CUUCGG/1M82_10_C_A_0_00028.pdb 
# Loading data/PDB/CUUCGG/1M82_10_C_A_0_00028.pdb 
# Loaded reference data/PDB/CUUCGG/2KOC_5_C_A_0_00001.pdb 
# Loaded target data/PDB/CUUCGG/1ZHO_15_C_B_0_00000.pdb 
# Loading data/PDB/CUUCGG/1ZHO_15_C_B_0_00000.pdb 
# Loaded reference data/PDB/CUUCGG/2KOC_5_C_A_0_00001.pdb 
# Loaded target data/PDB/CUUCGG/1FYP_11_C_A_0_00010.pdb 
# Loading data/PDB/CUUCGG/1FYP_11_C_A_0_00010.pdb 
# Loaded reference data/PDB/CUUCGG/2KOC_5_C_A_0_00001.pdb 
# Loaded target data/PDB/CUUCGG/1FYP_11_C_A_0_00004.pdb 
# Loading data/PDB/CUUCGG/1FYP_11_C_A_0_00004.pdb 
# Loaded reference data/PDB/CUUCGG/2KOC_5_C_A_0_00001.pdb 
# Loaded target data/PDB/CUUCGG/1HNZ_419_C_A_0_00000.pdb 
# Loading data/PDB/CUUCGG/1HNZ_419_C_A_0_00000.pdb 
# Loaded reference data/PDB/CUUCGG/2KOC_5_C_A_0_00001.pdb 
# Loaded target data/PDB/CUUCGG/5u9g-pdb-bundle1_1691_C_B_0_00004.pdb 
# Loading data/PDB/CUUCGG/5u9g-pdb-bundle1_1691_C_B_0_00004.pdb 
# Loaded reference data/PDB/CUUCGG/2KOC_5_C_A_0_00001.pdb 
# Loaded target data/PDB/CUUCGG/1AUD_28_C_B_0_00004.pdb 
# Loading data/PDB/CUUCGG/1AUD_28_C_B_0_00004.pdb 
# Loaded reference data/PDB/CUUCGG/2KOC_5_C_A_0_00001.pdb 
# Loaded target data/PDB/CUUCGG/1K8A_1769_C_A_0_00000.pdb 
# Loading data/PDB/CUUCGG/1K8A_1769_C_A_0_00000.pdb 
# Loaded reference data/PDB/CUUCGG/2KOC_5_C_A_0_00001.pdb 
# Loaded target data/PDB/CUUCGG/5dox-pdb-bundle2_419_C_A_0_00000.pdb 
# Loading data/PDB/CUUCGG/5dox-pdb-bundle2_419_C_A_0_00000.pdb 
# Loaded reference data/PDB/CUUCGG/2KOC_5_C_A_0_00001.pdb 
# Loaded target data/PDB/CUUCGG/6DCB_11_C_B_0_00000.pdb 
# Loading data/PDB/CUUCGG/6DCB_11_C_B_0_00000.pdb 
# Loaded reference data/PDB/CUUCGG/2KOC_5_C_A_0_00001.pdb 
# Loaded target data/PDB/CUUCGG/1EBS_12_C_A_0_00003.pdb 
# Loading data/PDB/CUUCGG/1EBS_12_C_A_0_00003.pdb 
# Loaded reference data/PDB/CUUCGG/2KOC_5_C_A_0_00001.pdb 
# Loaded target data/PDB/CUUCGG/2N6W_32_C_A_0_00005.pdb 
# Loading data/PDB/CUUCGG/2N6W_32_C_A_0_00005.pdb 
# Loaded reference data/PDB/CUUCGG/2KOC_5_C_A_0_00001.pdb 
# Loaded target data/PDB/CUUCGG/1RAW_21_C_A_0_00000.pdb 
# Loading data/PDB/CUUCGG/1RAW_21_C_A_0_00000.pdb 
# Loaded reference data/PDB/CUUCGG/2KOC_5_C_A_0_00001.pdb 
# Loaded target data/PDB/CUUCGG/5lzs-pdb-bundle2_2872_C_5_0_00000.pdb 
# Loading data/PDB/CUUCGG/5lzs-pdb-bundle2_2872_C_5_0_00000.pdb 
# Loaded reference data/PDB/CUUCGG/2KOC_5_C_A_0_00001.pdb 
# Loaded target data/PDB/CUUCGG/4v9h-pdb-bundle1_419_C_A_0_00000.pdb 
# Loading data/PDB/CUUCGG/4v9h-pdb-bundle1_419_C_A_0_00000.pdb 
# Loaded reference data/PDB/CUUCGG/2KOC_5_C_A_0_00001.pdb 
# Loaded target data/PDB/CUUCGG/4v5e-pdb-bundle3_419_C_A_0_00000.pdb 
# Loading data/PDB/CUUCGG/4v5e-pdb-bundle3_419_C_A_0_00000.pdb 
# Loaded reference data/PDB/CUUCGG/2KOC_5_C_A_0_00001.pdb 
# Loaded target data/PDB/CUUCGG/4LF7_419_C_A_0_00000.pdb 
# Loading data/PDB/CUUCGG/4LF7_419_C_A_0_00000.pdb 
# Loaded reference data/PDB/CUUCGG/2KOC_5_C_A_0_00001.pdb 
# Loaded target data/PDB/CUUCGG/5j8b-pdb-bundle1_1737_C_A_0_00000.pdb 
# Loading data/PDB/CUUCGG/5j8b-pdb-bundle1_1737_C_A_0_00000.pdb 
# Loaded reference data/PDB/CUUCGG/2KOC_5_C_A_0_00001.pdb 
# Loaded target data/PDB/CUUCGG/4v67-pdb-bundle3_419_C_A_0_00000.pdb 
# Loading data/PDB/CUUCGG/4v67-pdb-bundle3_419_C_A_0_00000.pdb 
# Loaded reference data/PDB/CUUCGG/2KOC_5_C_A_0_00001.pdb 
# Loaded target data/PDB/CUUCGG/5ib8-pdb-bundle1_419_C_A_0_00000.pdb 
# Loading data/PDB/CUUCGG/5ib8-pdb-bundle1_419_C_A_0_00000.pdb 
# Loaded reference data/PDB/CUUCGG/2KOC_5_C_A_0_00001.pdb 
# Loaded target data/PDB/CUUCGG/4u1u-pdb-bundle2_1691_C_A_0_00000.pdb 
# Loading data/PDB/CUUCGG/4u1u-pdb-bundle2_1691_C_A_0_00000.pdb 
# Loaded reference data/PDB/CUUCGG/2KOC_5_C_A_0_00001.pdb 
# Loaded target data/PDB/CUUCGG/5lze-pdb-bundle1_1028_C_a_0_00002.pdb 
# Loading data/PDB/CUUCGG/5lze-pdb-bundle1_1028_C_a_0_00002.pdb 
# Loaded reference data/PDB/CUUCGG/2KOC_5_C_A_0_00001.pdb 
# Loaded target data/PDB/CUUCGG/1vy7-pdb-bundle3_419_C_A_0_00000.pdb 
# Loading data/PDB/CUUCGG/1vy7-pdb-bundle3_419_C_A_0_00000.pdb 
# Loaded reference data/PDB/CUUCGG/2KOC_5_C_A_0_00001.pdb 
# Loaded target data/PDB/CUUCGG/1F7I_12_C_A_0_00000.pdb 
# Loading data/PDB/CUUCGG/1F7I_12_C_A_0_00000.pdb 
# Loaded reference data/PDB/CUUCGG/2KOC_5_C_A_0_00001.pdb 
# Loaded target data/PDB/CUUCGG/5t62-pdb-bundle1_1923_C_A_0_00002.pdb 
# Loading data/PDB/CUUCGG/5t62-pdb-bundle1_1923_C_A_0_00002.pdb 
# Loaded reference data/PDB/CUUCGG/2KOC_5_C_A_0_00001.pdb 
# Loaded target data/PDB/CUUCGG/1P5M_28_C_A_0_00001.pdb 
# Loading data/PDB/CUUCGG/1P5M_28_C_A_0_00001.pdb 
# Loaded reference data/PDB/CUUCGG/2KOC_5_C_A_0_00001.pdb 
# Loaded target data/PDB/CUUCGG/1EKZ_13_C_B_0_00019.pdb 
# Loading data/PDB/CUUCGG/1EKZ_13_C_B_0_00019.pdb 
# Loaded reference data/PDB/CUUCGG/2KOC_5_C_A_0_00001.pdb 
# Loaded target data/PDB/CUUCGG/5mdv-pdb-bundle1_419_C_2_0_00002.pdb 
# Loading data/PDB/CUUCGG/5mdv-pdb-bundle1_419_C_2_0_00002.pdb 
# Loaded reference data/PDB/CUUCGG/2KOC_5_C_A_0_00001.pdb 
# Loaded target data/PDB/CUUCGG/1P5M_28_C_A_0_00015.pdb 
# Loading data/PDB/CUUCGG/1P5M_28_C_A_0_00015.pdb 
# Loaded reference data/PDB/CUUCGG/2KOC_5_C_A_0_00001.pdb 
# Loaded target data/PDB/CUUCGG/2N6T_19_C_A_0_00002.pdb 
# Loading data/PDB/CUUCGG/2N6T_19_C_A_0_00002.pdb 
# Loaded reference data/PDB/CUUCGG/2KOC_5_C_A_0_00001.pdb 
# Loaded target data/PDB/CUUCGG/1ULL_14_C_A_0_00003.pdb 
# Loading data/PDB/CUUCGG/1ULL_14_C_A_0_00003.pdb 
# Loaded reference data/PDB/CUUCGG/2KOC_5_C_A_0_00001.pdb 
# Loaded target data/PDB/CUUCGG/1EKZ_13_C_B_0_00025.pdb 
# Loading data/PDB/CUUCGG/1EKZ_13_C_B_0_00025.pdb 
# Loaded reference data/PDB/CUUCGG/2KOC_5_C_A_0_00001.pdb 
# Loaded target data/PDB/CUUCGG/2L2J_19_C_A_0_00004.pdb 
# Loading data/PDB/CUUCGG/2L2J_19_C_A_0_00004.pdb 
# Loaded reference data/PDB/CUUCGG/2KOC_5_C_A_0_00001.pdb 
# Loaded target data/PDB/CUUCGG/1EKZ_13_C_B_0_00031.pdb 
# Loading data/PDB/CUUCGG/1EKZ_13_C_B_0_00031.pdb 
# Loaded reference data/PDB/CUUCGG/2KOC_5_C_A_0_00001.pdb 
# Loaded target data/PDB/CUUCGG/2L2J_19_C_A_0_00010.pdb 
# Loading data/PDB/CUUCGG/2L2J_19_C_A_0_00010.pdb 
# Loaded reference data/PDB/CUUCGG/2KOC_5_C_A_0_00001.pdb 
# Loaded target data/PDB/CUUCGG/1F7I_12_C_A_0_00010.pdb 
# Loading data/PDB/CUUCGG/1F7I_12_C_A_0_00010.pdb 
# Loaded reference data/PDB/CUUCGG/2KOC_5_C_A_0_00001.pdb 
# Loaded target data/PDB/CUUCGG/4v8g-pdb-bundle1_419_C_A_0_00000.pdb 
# Loading data/PDB/CUUCGG/4v8g-pdb-bundle1_419_C_A_0_00000.pdb 
# Loaded reference data/PDB/CUUCGG/2KOC_5_C_A_0_00001.pdb 
# Loaded target data/PDB/CUUCGG/2N6T_19_C_A_0_00006.pdb 
# Loading data/PDB/CUUCGG/2N6T_19_C_A_0_00006.pdb 
# Loaded reference data/PDB/CUUCGG/2KOC_5_C_A_0_00001.pdb 
# Loaded target data/PDB/CUUCGG/1EKZ_13_C_B_0_00009.pdb 
# Loading data/PDB/CUUCGG/1EKZ_13_C_B_0_00009.pdb 
# Loaded reference data/PDB/CUUCGG/2KOC_5_C_A_0_00001.pdb 
# Loaded target data/PDB/CUUCGG/1F7I_12_C_A_0_00004.pdb 
# Loading data/PDB/CUUCGG/1F7I_12_C_A_0_00004.pdb 
# Loaded reference data/PDB/CUUCGG/2KOC_5_C_A_0_00001.pdb 
# Loaded target data/PDB/CUUCGG/4v8n-pdb-bundle2_1691_C_A_0_00000.pdb 
# Loading data/PDB/CUUCGG/4v8n-pdb-bundle2_1691_C_A_0_00000.pdb 
# Loaded reference data/PDB/CUUCGG/2KOC_5_C_A_0_00001.pdb 
# Loaded target data/PDB/CUUCGG/2L2J_19_C_A_0_00014.pdb 
# Loading data/PDB/CUUCGG/2L2J_19_C_A_0_00014.pdb 
# Loaded reference data/PDB/CUUCGG/2KOC_5_C_A_0_00001.pdb 
# Loaded target data/PDB/CUUCGG/1EKZ_13_C_B_0_00035.pdb 
# Loading data/PDB/CUUCGG/1EKZ_13_C_B_0_00035.pdb 
# Loaded reference data/PDB/CUUCGG/2KOC_5_C_A_0_00001.pdb 
# Loaded target data/PDB/CUUCGG/4v4q-pdb-bundle3_419_C_A_0_00001.pdb 
# Loading data/PDB/CUUCGG/4v4q-pdb-bundle3_419_C_A_0_00001.pdb 
# Loaded reference data/PDB/CUUCGG/2KOC_5_C_A_0_00001.pdb 
# Loaded target data/PDB/CUUCGG/2L2J_19_C_A_0_00000.pdb 
# Loading data/PDB/CUUCGG/2L2J_19_C_A_0_00000.pdb 
# Loaded reference data/PDB/CUUCGG/2KOC_5_C_A_0_00001.pdb 
# Loaded target data/PDB/CUUCGG/1EKZ_13_C_B_0_00021.pdb 
# Loading data/PDB/CUUCGG/1EKZ_13_C_B_0_00021.pdb 
# Loaded reference data/PDB/CUUCGG/2KOC_5_C_A_0_00001.pdb 
# Loaded target data/PDB/CUUCGG/5hd1-pdb-bundle1_1737_C_A_0_00000.pdb 
# Loading data/PDB/CUUCGG/5hd1-pdb-bundle1_1737_C_A_0_00000.pdb 
# Loaded reference data/PDB/CUUCGG/2KOC_5_C_A_0_00001.pdb 
# Loaded target data/PDB/CUUCGG/1RAW_21_C_A_0_00004.pdb 
# Loading data/PDB/CUUCGG/1RAW_21_C_A_0_00004.pdb 
# Loaded reference data/PDB/CUUCGG/2KOC_5_C_A_0_00001.pdb 
# Loaded target data/PDB/CUUCGG/2N6W_32_C_A_0_00001.pdb 
# Loading data/PDB/CUUCGG/2N6W_32_C_A_0_00001.pdb 
# Loaded reference data/PDB/CUUCGG/2KOC_5_C_A_0_00001.pdb 
# Loaded target data/PDB/CUUCGG/4wt1-pdb-bundle4_1691_C_A_0_00000.pdb 
# Loading data/PDB/CUUCGG/4wt1-pdb-bundle4_1691_C_A_0_00000.pdb 
# Loaded reference data/PDB/CUUCGG/2KOC_5_C_A_0_00001.pdb 
# Loaded target data/PDB/CUUCGG/4wzd-pdb-bundle4_1691_C_A_0_00000.pdb 
# Loading data/PDB/CUUCGG/4wzd-pdb-bundle4_1691_C_A_0_00000.pdb 
# Loaded reference data/PDB/CUUCGG/2KOC_5_C_A_0_00001.pdb 
# Loaded target data/PDB/CUUCGG/5j30-pdb-bundle2_419_C_A_0_00000.pdb 
# Loading data/PDB/CUUCGG/5j30-pdb-bundle2_419_C_A_0_00000.pdb 
# Loaded reference data/PDB/CUUCGG/2KOC_5_C_A_0_00001.pdb 
# Loaded target data/PDB/CUUCGG/4v52-pdb-bundle4_1691_C_A_0_00000.pdb 
# Loading data/PDB/CUUCGG/4v52-pdb-bundle4_1691_C_A_0_00000.pdb 
# Loaded reference data/PDB/CUUCGG/2KOC_5_C_A_0_00001.pdb 
# Loaded target data/PDB/CUUCGG/5it8-pdb-bundle1_1449_C_A_0_00003.pdb 
# Loading data/PDB/CUUCGG/5it8-pdb-bundle1_1449_C_A_0_00003.pdb 
# Loaded reference data/PDB/CUUCGG/2KOC_5_C_A_0_00001.pdb 
# Loaded target data/PDB/CUUCGG/1AUD_28_C_B_0_00028.pdb 
# Loading data/PDB/CUUCGG/1AUD_28_C_B_0_00028.pdb 
# Loaded reference data/PDB/CUUCGG/2KOC_5_C_A_0_00001.pdb 
# Loaded target data/PDB/CUUCGG/1FYP_11_C_A_0_00000.pdb 
# Loading data/PDB/CUUCGG/1FYP_11_C_A_0_00000.pdb 
# Loaded reference data/PDB/CUUCGG/2KOC_5_C_A_0_00001.pdb 
# Loaded target data/PDB/CUUCGG/1FYP_11_C_A_0_00014.pdb 
# Loading data/PDB/CUUCGG/1FYP_11_C_A_0_00014.pdb 
# Loaded reference data/PDB/CUUCGG/2KOC_5_C_A_0_00001.pdb 
# Loaded target data/PDB/CUUCGG/4v7l-pdb-bundle4_1691_C_A_0_00000.pdb 
# Loading data/PDB/CUUCGG/4v7l-pdb-bundle4_1691_C_A_0_00000.pdb 
# Loaded reference data/PDB/CUUCGG/2KOC_5_C_A_0_00001.pdb 
# Loaded target data/PDB/CUUCGG/4www-pdb-bundle1_1691_C_A_0_00000.pdb 
# Loading data/PDB/CUUCGG/4www-pdb-bundle1_1691_C_A_0_00000.pdb 
# Loaded reference data/PDB/CUUCGG/2KOC_5_C_A_0_00001.pdb 
# Loaded target data/PDB/CUUCGG/4y4o-pdb-bundle4_419_C_A_0_00000.pdb 
# Loading data/PDB/CUUCGG/4y4o-pdb-bundle4_419_C_A_0_00000.pdb 
# Loaded reference data/PDB/CUUCGG/2KOC_5_C_A_0_00001.pdb 
# Loaded target data/PDB/CUUCGG/6DLQ_27_C_A_0_00000.pdb 
# Loading data/PDB/CUUCGG/6DLQ_27_C_A_0_00000.pdb 
# Loaded reference data/PDB/CUUCGG/2KOC_5_C_A_0_00001.pdb 
# Loaded target data/PDB/CUUCGG/5ndk-pdb-bundle3_1741_C_A_0_00000.pdb 
# Loading data/PDB/CUUCGG/5ndk-pdb-bundle3_1741_C_A_0_00000.pdb 
# Loaded reference data/PDB/CUUCGG/2KOC_5_C_A_0_00001.pdb 
# Loaded target data/PDB/CUUCGG/5e81-pdb-bundle4_1691_C_A_0_00000.pdb 
# Loading data/PDB/CUUCGG/5e81-pdb-bundle4_1691_C_A_0_00000.pdb 
# Loaded reference data/PDB/CUUCGG/2KOC_5_C_A_0_00001.pdb 
# Loaded target data/PDB/CUUCGG/1AUD_28_C_B_0_00000.pdb 
# Loading data/PDB/CUUCGG/1AUD_28_C_B_0_00000.pdb 
# Loaded reference data/PDB/CUUCGG/2KOC_5_C_A_0_00001.pdb 
# Loaded target data/PDB/CUUCGG/4v8b-pdb-bundle3_419_C_A_0_00000.pdb 
# Loading data/PDB/CUUCGG/4v8b-pdb-bundle3_419_C_A_0_00000.pdb 
# Loaded reference data/PDB/CUUCGG/2KOC_5_C_A_0_00001.pdb 
# Loaded target data/PDB/CUUCGG/1FYP_11_C_A_0_00028.pdb 
# Loading data/PDB/CUUCGG/1FYP_11_C_A_0_00028.pdb 
# Loaded reference data/PDB/CUUCGG/2KOC_5_C_A_0_00001.pdb 
# Loaded target data/PDB/CUUCGG/1M82_10_C_A_0_00010.pdb 
# Loading data/PDB/CUUCGG/1M82_10_C_A_0_00010.pdb 
# Loaded reference data/PDB/CUUCGG/2KOC_5_C_A_0_00001.pdb 
# Loaded target data/PDB/CUUCGG/6hhq-pdb-bundle1_1923_C_1_0_00002.pdb 
# Loading data/PDB/CUUCGG/6hhq-pdb-bundle1_1923_C_1_0_00002.pdb 
# Loaded reference data/PDB/CUUCGG/2KOC_5_C_A_0_00001.pdb 
# Loaded target data/PDB/CUUCGG/1M82_10_C_A_0_00004.pdb 
# Loading data/PDB/CUUCGG/1M82_10_C_A_0_00004.pdb 
# Loaded reference data/PDB/CUUCGG/2KOC_5_C_A_0_00001.pdb 
# Loaded target data/PDB/CUUCGG/5o61-pdb-bundle1_1909_C_A_0_00000.pdb 
# Loading data/PDB/CUUCGG/5o61-pdb-bundle1_1909_C_A_0_00000.pdb 
# Loaded reference data/PDB/CUUCGG/2KOC_5_C_A_0_00001.pdb 
# Loaded target data/PDB/CUUCGG/2M4Q_11_C_1_0_00000.pdb 
# Loading data/PDB/CUUCGG/2M4Q_11_C_1_0_00000.pdb 
# Loaded reference data/PDB/CUUCGG/2KOC_5_C_A_0_00001.pdb 
# Loaded target data/PDB/CUUCGG/4v5d-pdb-bundle4_1691_C_A_0_00000.pdb 
# Loading data/PDB/CUUCGG/4v5d-pdb-bundle4_1691_C_A_0_00000.pdb 
# Loaded reference data/PDB/CUUCGG/2KOC_5_C_A_0_00001.pdb 
# Loaded target data/PDB/CUUCGG/4v83-pdb-bundle3_419_C_A_0_00000.pdb 
# Loading data/PDB/CUUCGG/4v83-pdb-bundle3_419_C_A_0_00000.pdb 
# Loaded reference data/PDB/CUUCGG/2KOC_5_C_A_0_00001.pdb 
# Loaded target data/PDB/CUUCGG/4y4o-pdb-bundle3_1691_C_A_0_00000.pdb 
# Loading data/PDB/CUUCGG/4y4o-pdb-bundle3_1691_C_A_0_00000.pdb 
# Loaded reference data/PDB/CUUCGG/2KOC_5_C_A_0_00001.pdb 
# Loaded target data/PDB/CUUCGG/4v7v-pdb-bundle3_1028_C_A_0_00002.pdb 
# Loading data/PDB/CUUCGG/4v7v-pdb-bundle3_1028_C_A_0_00002.pdb 
# Loaded reference data/PDB/CUUCGG/2KOC_5_C_A_0_00001.pdb 
# Loaded target data/PDB/CUUCGG/1VQ5_1769_C_0_0_00000.pdb 
# Loading data/PDB/CUUCGG/1VQ5_1769_C_0_0_00000.pdb 
# Loaded reference data/PDB/CUUCGG/2KOC_5_C_A_0_00001.pdb 
# Loaded target data/PDB/CUUCGG/2KOC_5_C_A_0_00013.pdb 
# Loading data/PDB/CUUCGG/2KOC_5_C_A_0_00013.pdb 
# Loaded reference data/PDB/CUUCGG/2KOC_5_C_A_0_00001.pdb 
# Loaded target data/PDB/CUUCGG/5uq8-pdb-bundle1_1691_C_A_0_00000.pdb 
# Loading data/PDB/CUUCGG/5uq8-pdb-bundle1_1691_C_A_0_00000.pdb 
# Loaded reference data/PDB/CUUCGG/2KOC_5_C_A_0_00001.pdb 
# Loaded target data/PDB/CUUCGG/6cfj-pdb-bundle3_1691_C_A_0_00000.pdb 
# Loading data/PDB/CUUCGG/6cfj-pdb-bundle3_1691_C_A_0_00000.pdb 
# Loaded reference data/PDB/CUUCGG/2KOC_5_C_A_0_00001.pdb 
# Loaded target data/PDB/CUUCGG/1KQS_1769_C_0_0_00000.pdb 
# Loading data/PDB/CUUCGG/1KQS_1769_C_0_0_00000.pdb 
# Loaded reference data/PDB/CUUCGG/2KOC_5_C_A_0_00001.pdb 
# Loaded target data/PDB/CUUCGG/5obm-pdb-bundle1_1923_C_A_0_00002.pdb 
# Loading data/PDB/CUUCGG/5obm-pdb-bundle1_1923_C_A_0_00002.pdb 
# Loaded reference data/PDB/CUUCGG/2KOC_5_C_A_0_00001.pdb 
# Loaded target data/PDB/CUUCGG/2KOC_5_C_A_0_00007.pdb 
# Loading data/PDB/CUUCGG/2KOC_5_C_A_0_00007.pdb 
# Loaded reference data/PDB/CUUCGG/2KOC_5_C_A_0_00001.pdb 
# Loaded target data/PDB/CUUCGG/5jup-pdb-bundle2_1923_C_B_0_00002.pdb 
# Loading data/PDB/CUUCGG/5jup-pdb-bundle2_1923_C_B_0_00002.pdb 
# Loaded reference data/PDB/CUUCGG/2KOC_5_C_A_0_00001.pdb 
# Loaded target data/PDB/CUUCGG/1G70_54_C_A_0_00004.pdb 
# Loading data/PDB/CUUCGG/1G70_54_C_A_0_00004.pdb 
# Loaded reference data/PDB/CUUCGG/2KOC_5_C_A_0_00001.pdb 
# Loaded target data/PDB/CUUCGG/4v9a-pdb-bundle2_1691_C_A_0_00000.pdb 
# Loading data/PDB/CUUCGG/4v9a-pdb-bundle2_1691_C_A_0_00000.pdb 
# Loaded reference data/PDB/CUUCGG/2KOC_5_C_A_0_00001.pdb 
# Loaded target data/PDB/CUUCGG/1G70_54_C_A_0_00010.pdb 
# Loading data/PDB/CUUCGG/1G70_54_C_A_0_00010.pdb 
# Loaded reference data/PDB/CUUCGG/2KOC_5_C_A_0_00001.pdb 
# Loaded target data/PDB/CUUCGG/6h4n-pdb-bundle1_1449_C_a_0_00003.pdb 
# Loading data/PDB/CUUCGG/6h4n-pdb-bundle1_1449_C_a_0_00003.pdb 
# Loaded reference data/PDB/CUUCGG/2KOC_5_C_A_0_00001.pdb 
# Loaded target data/PDB/CUUCGG/2KTZ_19_C_A_0_00008.pdb 
# Loading data/PDB/CUUCGG/2KTZ_19_C_A_0_00008.pdb 
# Loaded reference data/PDB/CUUCGG/2KOC_5_C_A_0_00001.pdb 
# Loaded target data/PDB/CUUCGG/4v4h-pdb-bundle3_1449_C_A_0_00003.pdb 
# Loading data/PDB/CUUCGG/4v4h-pdb-bundle3_1449_C_A_0_00003.pdb 
# Loaded reference data/PDB/CUUCGG/2KOC_5_C_A_0_00001.pdb 
# Loaded target data/PDB/CUUCGG/6buw-pdb-bundle1_419_C_A_0_00000.pdb 
# Loading data/PDB/CUUCGG/6buw-pdb-bundle1_419_C_A_0_00000.pdb 
# Loaded reference data/PDB/CUUCGG/2KOC_5_C_A_0_00001.pdb 
# Loaded target data/PDB/CUUCGG/6fkr-pdb-bundle1_1691_C_A_0_00000.pdb 
# Loading data/PDB/CUUCGG/6fkr-pdb-bundle1_1691_C_A_0_00000.pdb 
# Loaded reference data/PDB/CUUCGG/2KOC_5_C_A_0_00001.pdb 
# Loaded target data/PDB/CUUCGG/4v5j-pdb-bundle2_1691_C_A_0_00000.pdb 
# Loading data/PDB/CUUCGG/4v5j-pdb-bundle2_1691_C_A_0_00000.pdb 
# Loaded reference data/PDB/CUUCGG/2KOC_5_C_A_0_00001.pdb 
# Loaded target data/PDB/CUUCGG/1IKD_7_C_A_0_00025.pdb 
# Loading data/PDB/CUUCGG/1IKD_7_C_A_0_00025.pdb 
# Loaded reference data/PDB/CUUCGG/2KOC_5_C_A_0_00001.pdb 
# Loaded target data/PDB/CUUCGG/1Z31_270_C_A_0_00007.pdb 
# Loading data/PDB/CUUCGG/1Z31_270_C_A_0_00007.pdb 
# Loaded reference data/PDB/CUUCGG/2KOC_5_C_A_0_00001.pdb 
# Loaded target data/PDB/CUUCGG/4v54-pdb-bundle1_1449_C_A_0_00003.pdb 
# Loading data/PDB/CUUCGG/4v54-pdb-bundle1_1449_C_A_0_00003.pdb 
# Loaded reference data/PDB/CUUCGG/2KOC_5_C_A_0_00001.pdb 
# Loaded target data/PDB/CUUCGG/5LYV_48_C_A_0_00000.pdb 
# Loading data/PDB/CUUCGG/5LYV_48_C_A_0_00000.pdb 
# Loaded reference data/PDB/CUUCGG/2KOC_5_C_A_0_00001.pdb 
# Loaded target data/PDB/CUUCGG/5O60_1909_C_A_0_00000.pdb 
# Loading data/PDB/CUUCGG/5O60_1909_C_A_0_00000.pdb 
# Loaded reference data/PDB/CUUCGG/2KOC_5_C_A_0_00001.pdb 
# Loaded target data/PDB/CUUCGG/6enf-pdb-bundle1_419_C_a_0_00001.pdb 
# Loading data/PDB/CUUCGG/6enf-pdb-bundle1_419_C_a_0_00001.pdb 
# Loaded reference data/PDB/CUUCGG/2KOC_5_C_A_0_00001.pdb 
# Loaded target data/PDB/CUUCGG/4v6c-pdb-bundle4_1691_C_A_0_00000.pdb 
# Loading data/PDB/CUUCGG/4v6c-pdb-bundle4_1691_C_A_0_00000.pdb 
# Loaded reference data/PDB/CUUCGG/2KOC_5_C_A_0_00001.pdb 
# Loaded target data/PDB/CUUCGG/1F7G_12_C_A_0_00014.pdb 
# Loading data/PDB/CUUCGG/1F7G_12_C_A_0_00014.pdb 
# Loaded reference data/PDB/CUUCGG/2KOC_5_C_A_0_00001.pdb 
# Loaded target data/PDB/CUUCGG/2HGH_32_C_B_0_00015.pdb 
# Loading data/PDB/CUUCGG/2HGH_32_C_B_0_00015.pdb 
# Loaded reference data/PDB/CUUCGG/2KOC_5_C_A_0_00001.pdb 
# Loaded target data/PDB/CUUCGG/5o2r-pdb-bundle2_1449_C_a_0_00003.pdb 
# Loading data/PDB/CUUCGG/5o2r-pdb-bundle2_1449_C_a_0_00003.pdb 
# Loaded reference data/PDB/CUUCGG/2KOC_5_C_A_0_00001.pdb 
# Loaded target data/PDB/CUUCGG/2HGH_32_C_B_0_00001.pdb 
# Loading data/PDB/CUUCGG/2HGH_32_C_B_0_00001.pdb 
# Loaded reference data/PDB/CUUCGG/2KOC_5_C_A_0_00001.pdb 
# Loaded target data/PDB/CUUCGG/1F7G_12_C_A_0_00000.pdb 
# Loading data/PDB/CUUCGG/1F7G_12_C_A_0_00000.pdb 
# Loaded reference data/PDB/CUUCGG/2KOC_5_C_A_0_00001.pdb 
# Loaded target data/PDB/CUUCGG/5e7k-pdb-bundle4_1691_C_A_0_00000.pdb 
# Loading data/PDB/CUUCGG/5e7k-pdb-bundle4_1691_C_A_0_00000.pdb 
# Loaded reference data/PDB/CUUCGG/2KOC_5_C_A_0_00001.pdb 
# Loaded target data/PDB/CUUCGG/4v9p-pdb-bundle2_1449_C_A_0_00003.pdb 
# Loading data/PDB/CUUCGG/4v9p-pdb-bundle2_1449_C_A_0_00003.pdb 
# Loaded reference data/PDB/CUUCGG/2KOC_5_C_A_0_00001.pdb 
# Loaded target data/PDB/CUUCGG/1C0O_5_C_A_0_00013.pdb 
# Loading data/PDB/CUUCGG/1C0O_5_C_A_0_00013.pdb 
# Loaded reference data/PDB/CUUCGG/2KOC_5_C_A_0_00001.pdb 
# Loaded target data/PDB/CUUCGG/4v4q-pdb-bundle3_1028_C_A_0_00002.pdb 
# Loading data/PDB/CUUCGG/4v4q-pdb-bundle3_1028_C_A_0_00002.pdb 
# Loaded reference data/PDB/CUUCGG/2KOC_5_C_A_0_00001.pdb 
# Loaded target data/PDB/CUUCGG/5ngm-pdb-bundle2_2819_C_A_0_00001.pdb 
# Loading data/PDB/CUUCGG/5ngm-pdb-bundle2_2819_C_A_0_00001.pdb 
# Loaded reference data/PDB/CUUCGG/2KOC_5_C_A_0_00001.pdb 
# Loaded target data/PDB/CUUCGG/1vy7-pdb-bundle2_1691_C_A_0_00000.pdb 
# Loading data/PDB/CUUCGG/1vy7-pdb-bundle2_1691_C_A_0_00000.pdb 
# Loaded reference data/PDB/CUUCGG/2KOC_5_C_A_0_00001.pdb 
# Loaded target data/PDB/CUUCGG/1C0O_5_C_A_0_00007.pdb 
# Loading data/PDB/CUUCGG/1C0O_5_C_A_0_00007.pdb 
# Loaded reference data/PDB/CUUCGG/2KOC_5_C_A_0_00001.pdb 
# Loaded target data/PDB/CUUCGG/1C0O_5_C_A_0_00006.pdb 
# Loading data/PDB/CUUCGG/1C0O_5_C_A_0_00006.pdb 
# Loaded reference data/PDB/CUUCGG/2KOC_5_C_A_0_00001.pdb 
# Loaded target data/PDB/CUUCGG/5mdz-pdb-bundle1_1449_C_2_0_00004.pdb 
# Loading data/PDB/CUUCGG/5mdz-pdb-bundle1_1449_C_2_0_00004.pdb 
# Loaded reference data/PDB/CUUCGG/2KOC_5_C_A_0_00001.pdb 
# Loaded target data/PDB/CUUCGG/5el7-pdb-bundle3_419_C_A_0_00000.pdb 
# Loading data/PDB/CUUCGG/5el7-pdb-bundle3_419_C_A_0_00000.pdb 
# Loaded reference data/PDB/CUUCGG/2KOC_5_C_A_0_00001.pdb 
# Loaded target data/PDB/CUUCGG/1C0O_5_C_A_0_00012.pdb 
# Loading data/PDB/CUUCGG/1C0O_5_C_A_0_00012.pdb 
# Loaded reference data/PDB/CUUCGG/2KOC_5_C_A_0_00001.pdb 
# Loaded target data/PDB/CUUCGG/5JVG_1708_C_X_0_00000.pdb 
# Loading data/PDB/CUUCGG/5JVG_1708_C_X_0_00000.pdb 
# Loaded reference data/PDB/CUUCGG/2KOC_5_C_A_0_00001.pdb 
# Loaded target data/PDB/CUUCGG/1Q86_1769_C_A_0_00000.pdb 
# Loading data/PDB/CUUCGG/1Q86_1769_C_A_0_00000.pdb 
# Loaded reference data/PDB/CUUCGG/2KOC_5_C_A_0_00001.pdb 
# Loaded target data/PDB/CUUCGG/2HGH_32_C_B_0_00000.pdb 
# Loading data/PDB/CUUCGG/2HGH_32_C_B_0_00000.pdb 
# Loaded reference data/PDB/CUUCGG/2KOC_5_C_A_0_00001.pdb 
# Loaded target data/PDB/CUUCGG/1F7G_12_C_A_0_00001.pdb 
# Loading data/PDB/CUUCGG/1F7G_12_C_A_0_00001.pdb 
# Loaded reference data/PDB/CUUCGG/2KOC_5_C_A_0_00001.pdb 
# Loaded target data/PDB/CUUCGG/1F7G_12_C_A_0_00015.pdb 
# Loading data/PDB/CUUCGG/1F7G_12_C_A_0_00015.pdb 
# Loaded reference data/PDB/CUUCGG/2KOC_5_C_A_0_00001.pdb 
# Loaded target data/PDB/CUUCGG/5mdw-pdb-bundle1_1691_C_1_0_00000.pdb 
# Loading data/PDB/CUUCGG/5mdw-pdb-bundle1_1691_C_1_0_00000.pdb 
# Loaded reference data/PDB/CUUCGG/2KOC_5_C_A_0_00001.pdb 
# Loaded target data/PDB/CUUCGG/2HGH_32_C_B_0_00014.pdb 
# Loading data/PDB/CUUCGG/2HGH_32_C_B_0_00014.pdb 
# Loaded reference data/PDB/CUUCGG/2KOC_5_C_A_0_00001.pdb 
# Loaded target data/PDB/CUUCGG/1IKD_7_C_A_0_00018.pdb 
# Loading data/PDB/CUUCGG/1IKD_7_C_A_0_00018.pdb 
# Loaded reference data/PDB/CUUCGG/2KOC_5_C_A_0_00001.pdb 
# Loaded target data/PDB/CUUCGG/5hcq-pdb-bundle1_1737_C_A_0_00000.pdb 
# Loading data/PDB/CUUCGG/5hcq-pdb-bundle1_1737_C_A_0_00000.pdb 
# Loaded reference data/PDB/CUUCGG/2KOC_5_C_A_0_00001.pdb 
# Loaded target data/PDB/CUUCGG/4u1u-pdb-bundle1_419_C_A_0_00001.pdb 
# Loading data/PDB/CUUCGG/4u1u-pdb-bundle1_419_C_A_0_00001.pdb 
# Loaded reference data/PDB/CUUCGG/2KOC_5_C_A_0_00001.pdb 
# Loaded target data/PDB/CUUCGG/4u55-pdb-bundle2_1923_C_A_0_00002.pdb 
# Loading data/PDB/CUUCGG/4u55-pdb-bundle2_1923_C_A_0_00002.pdb 
# Loaded reference data/PDB/CUUCGG/2KOC_5_C_A_0_00001.pdb 
# Loaded target data/PDB/CUUCGG/1Z31_270_C_A_0_00006.pdb 
# Loading data/PDB/CUUCGG/1Z31_270_C_A_0_00006.pdb 
# Loaded reference data/PDB/CUUCGG/2KOC_5_C_A_0_00001.pdb 
# Loaded target data/PDB/CUUCGG/1IKD_7_C_A_0_00024.pdb 
# Loading data/PDB/CUUCGG/1IKD_7_C_A_0_00024.pdb 
# Loaded reference data/PDB/CUUCGG/2KOC_5_C_A_0_00001.pdb 
# Loaded target data/PDB/CUUCGG/4v64-pdb-bundle1_207_C_A_0_00000.pdb 
# Loading data/PDB/CUUCGG/4v64-pdb-bundle1_207_C_A_0_00000.pdb 
# Loaded reference data/PDB/CUUCGG/2KOC_5_C_A_0_00001.pdb 
# Loaded target data/PDB/CUUCGG/5j8a-pdb-bundle1_419_C_b_0_00005.pdb 
# Loading data/PDB/CUUCGG/5j8a-pdb-bundle1_419_C_b_0_00005.pdb 
# Loaded reference data/PDB/CUUCGG/2KOC_5_C_A_0_00001.pdb 
# Loaded target data/PDB/CUUCGG/3JAM_229_C_2_0_00000.pdb 
# Loading data/PDB/CUUCGG/3JAM_229_C_2_0_00000.pdb 
# Loaded reference data/PDB/CUUCGG/2KOC_5_C_A_0_00001.pdb 
# Loaded target data/PDB/CUUCGG/2KTZ_19_C_A_0_00009.pdb 
# Loading data/PDB/CUUCGG/2KTZ_19_C_A_0_00009.pdb 
# Loaded reference data/PDB/CUUCGG/2KOC_5_C_A_0_00001.pdb 
# Loaded target data/PDB/CUUCGG/5uym-pdb-bundle2_1691_C_A_0_00000.pdb 
# Loading data/PDB/CUUCGG/5uym-pdb-bundle2_1691_C_A_0_00000.pdb 
# Loaded reference data/PDB/CUUCGG/2KOC_5_C_A_0_00001.pdb 
# Loaded target data/PDB/CUUCGG/1G70_54_C_A_0_00011.pdb 
# Loading data/PDB/CUUCGG/1G70_54_C_A_0_00011.pdb 
# Loaded reference data/PDB/CUUCGG/2KOC_5_C_A_0_00001.pdb 
# Loaded target data/PDB/CUUCGG/2OTJ_1769_C_0_0_00000.pdb 
# Loading data/PDB/CUUCGG/2OTJ_1769_C_0_0_00000.pdb 
# Loaded reference data/PDB/CUUCGG/2KOC_5_C_A_0_00001.pdb 
# Loaded target data/PDB/CUUCGG/1G70_54_C_A_0_00005.pdb 
# Loading data/PDB/CUUCGG/1G70_54_C_A_0_00005.pdb 
# Loaded reference data/PDB/CUUCGG/2KOC_5_C_A_0_00001.pdb 
# Loaded target data/PDB/CUUCGG/5hcq-pdb-bundle2_415_C_A_0_00000.pdb 
# Loading data/PDB/CUUCGG/5hcq-pdb-bundle2_415_C_A_0_00000.pdb 
# Loaded reference data/PDB/CUUCGG/2KOC_5_C_A_0_00001.pdb 
# Loaded target data/PDB/CUUCGG/2KOC_5_C_A_0_00006.pdb 
# Loading data/PDB/CUUCGG/2KOC_5_C_A_0_00006.pdb 
# Loaded reference data/PDB/CUUCGG/2KOC_5_C_A_0_00001.pdb 
# Loaded target data/PDB/CUUCGG/4u3m-pdb-bundle2_1923_C_A_0_00002.pdb 
# Loading data/PDB/CUUCGG/4u3m-pdb-bundle2_1923_C_A_0_00002.pdb 
# Loaded reference data/PDB/CUUCGG/2KOC_5_C_A_0_00001.pdb 
# Loaded target data/PDB/CUUCGG/2KOC_5_C_A_0_00012.pdb 
# Loading data/PDB/CUUCGG/2KOC_5_C_A_0_00012.pdb 
# Loaded reference data/PDB/CUUCGG/2KOC_5_C_A_0_00001.pdb 
# Loaded target data/PDB/CUUCGG/4v5s-pdb-bundle4_1691_C_A_0_00000.pdb 
# Loading data/PDB/CUUCGG/4v5s-pdb-bundle4_1691_C_A_0_00000.pdb 
# Loaded reference data/PDB/CUUCGG/2KOC_5_C_A_0_00001.pdb 
# Loaded target data/PDB/CUUCGG/5tga-pdb-bundle3_230_C_d_0_00000.pdb 
# Loading data/PDB/CUUCGG/5tga-pdb-bundle3_230_C_d_0_00000.pdb 
# Loaded reference data/PDB/CUUCGG/2KOC_5_C_A_0_00001.pdb 
# Loaded target data/PDB/CUUCGG/6bz6-pdb-bundle1_419_C_A_0_00000.pdb 
# Loading data/PDB/CUUCGG/6bz6-pdb-bundle1_419_C_A_0_00000.pdb 
# Loaded reference data/PDB/CUUCGG/2KOC_5_C_A_0_00001.pdb 
# Loaded target data/PDB/CUUCGG/3CCM_1769_C_0_0_00000.pdb 
# Loading data/PDB/CUUCGG/3CCM_1769_C_0_0_00000.pdb 
# Loaded reference data/PDB/CUUCGG/2KOC_5_C_A_0_00001.pdb 
# Loaded target data/PDB/CUUCGG/5WNV_419_C_A_0_00000.pdb 
# Loading data/PDB/CUUCGG/5WNV_419_C_A_0_00000.pdb 
# Loaded reference data/PDB/CUUCGG/2KOC_5_C_A_0_00001.pdb 
# Loaded target data/PDB/CUUCGG/4w2e-pdb-bundle2_419_C_a_0_00000.pdb 
# Loading data/PDB/CUUCGG/4w2e-pdb-bundle2_419_C_a_0_00000.pdb 
# Loaded reference data/PDB/CUUCGG/2KOC_5_C_A_0_00001.pdb 
# Loaded target data/PDB/CUUCGG/5u9f-pdb-bundle1_1449_C_A_0_00003.pdb 
# Loading data/PDB/CUUCGG/5u9f-pdb-bundle1_1449_C_A_0_00003.pdb 
# Loaded reference data/PDB/CUUCGG/2KOC_5_C_A_0_00001.pdb 
# Loaded target data/PDB/CUUCGG/6gsk-pdb-bundle4_1691_C_A_0_00000.pdb 
# Loading data/PDB/CUUCGG/6gsk-pdb-bundle4_1691_C_A_0_00000.pdb 
# Loaded reference data/PDB/CUUCGG/2KOC_5_C_A_0_00001.pdb 
# Loaded target data/PDB/CUUCGG/5wit-pdb-bundle3_1691_C_A_0_00000.pdb 
# Loading data/PDB/CUUCGG/5wit-pdb-bundle3_1691_C_A_0_00000.pdb 
# Loaded reference data/PDB/CUUCGG/2KOC_5_C_A_0_00001.pdb 
# Loaded target data/PDB/CUUCGG/4wqf-pdb-bundle1_1738_C_A_0_00000.pdb 
# Loading data/PDB/CUUCGG/4wqf-pdb-bundle1_1738_C_A_0_00000.pdb 
# Loaded reference data/PDB/CUUCGG/2KOC_5_C_A_0_00001.pdb 
# Loaded target data/PDB/CUUCGG/4DUY_419_C_A_0_00000.pdb 
# Loading data/PDB/CUUCGG/4DUY_419_C_A_0_00000.pdb 
# Loaded reference data/PDB/CUUCGG/2KOC_5_C_A_0_00001.pdb 
# Loaded target data/PDB/CUUCGG/1M82_10_C_A_0_00005.pdb 
# Loading data/PDB/CUUCGG/1M82_10_C_A_0_00005.pdb 
# Loaded reference data/PDB/CUUCGG/2KOC_5_C_A_0_00001.pdb 
# Loaded target data/PDB/CUUCGG/1I6U_16_C_C_0_00000.pdb 
# Loading data/PDB/CUUCGG/1I6U_16_C_C_0_00000.pdb 
# Loaded reference data/PDB/CUUCGG/2KOC_5_C_A_0_00001.pdb 
# Loaded target data/PDB/CUUCGG/5v8i-pdb-bundle2_419_C_A_0_00000.pdb 
# Loading data/PDB/CUUCGG/5v8i-pdb-bundle2_419_C_A_0_00000.pdb 
# Loaded reference data/PDB/CUUCGG/2KOC_5_C_A_0_00001.pdb 
# Loaded target data/PDB/CUUCGG/4wzo-pdb-bundle1_419_C_A_0_00000.pdb 
# Loading data/PDB/CUUCGG/4wzo-pdb-bundle1_419_C_A_0_00000.pdb 
# Loaded reference data/PDB/CUUCGG/2KOC_5_C_A_0_00001.pdb 
# Loaded target data/PDB/CUUCGG/1P9X_1708_C_0_0_00000.pdb 
# Loading data/PDB/CUUCGG/1P9X_1708_C_0_0_00000.pdb 
# Loaded reference data/PDB/CUUCGG/2KOC_5_C_A_0_00001.pdb 
# Loaded target data/PDB/CUUCGG/1M82_10_C_A_0_00011.pdb 
# Loading data/PDB/CUUCGG/1M82_10_C_A_0_00011.pdb 
# Loaded reference data/PDB/CUUCGG/2KOC_5_C_A_0_00001.pdb 
# Loaded target data/PDB/CUUCGG/5xxb-pdb-bundle1_2026_C_1_0_00000.pdb 
# Loading data/PDB/CUUCGG/5xxb-pdb-bundle1_2026_C_1_0_00000.pdb 
# Loaded reference data/PDB/CUUCGG/2KOC_5_C_A_0_00001.pdb 
# Loaded target data/PDB/CUUCGG/5fdv-pdb-bundle2_419_C_A_0_00000.pdb 
# Loading data/PDB/CUUCGG/5fdv-pdb-bundle2_419_C_A_0_00000.pdb 
# Loaded reference data/PDB/CUUCGG/2KOC_5_C_A_0_00001.pdb 
# Loaded target data/PDB/CUUCGG/1AUD_28_C_B_0_00015.pdb 
# Loading data/PDB/CUUCGG/1AUD_28_C_B_0_00015.pdb 
# Loaded reference data/PDB/CUUCGG/2KOC_5_C_A_0_00001.pdb 
# Loaded target data/PDB/CUUCGG/4v9p-pdb-bundle2_207_C_A_0_00000.pdb 
# Loading data/PDB/CUUCGG/4v9p-pdb-bundle2_207_C_A_0_00000.pdb 
# Loaded reference data/PDB/CUUCGG/2KOC_5_C_A_0_00001.pdb 
# Loaded target data/PDB/CUUCGG/1FYP_11_C_A_0_00015.pdb 
# Loading data/PDB/CUUCGG/1FYP_11_C_A_0_00015.pdb 
# Loaded reference data/PDB/CUUCGG/2KOC_5_C_A_0_00001.pdb 
# Loaded target data/PDB/CUUCGG/4v52-pdb-bundle3_207_C_A_0_00000.pdb 
# Loading data/PDB/CUUCGG/4v52-pdb-bundle3_207_C_A_0_00000.pdb 
# Loaded reference data/PDB/CUUCGG/2KOC_5_C_A_0_00001.pdb 
# Loaded target data/PDB/CUUCGG/3AM1_32_C_B_0_00000.pdb 
# Loading data/PDB/CUUCGG/3AM1_32_C_B_0_00000.pdb 
# Loaded reference data/PDB/CUUCGG/2KOC_5_C_A_0_00001.pdb 
# Loaded target data/PDB/CUUCGG/4DR6_419_C_A_0_00000.pdb 
# Loading data/PDB/CUUCGG/4DR6_419_C_A_0_00000.pdb 
# Loaded reference data/PDB/CUUCGG/2KOC_5_C_A_0_00001.pdb 
# Loaded target data/PDB/CUUCGG/4v9p-pdb-bundle8_419_C_A_0_00001.pdb 
# Loading data/PDB/CUUCGG/4v9p-pdb-bundle8_419_C_A_0_00001.pdb 
# Loaded reference data/PDB/CUUCGG/2KOC_5_C_A_0_00001.pdb 
# Loaded target data/PDB/CUUCGG/1FYP_11_C_A_0_00001.pdb 
# Loading data/PDB/CUUCGG/1FYP_11_C_A_0_00001.pdb 
# Loaded reference data/PDB/CUUCGG/2KOC_5_C_A_0_00001.pdb 
# Loaded target data/PDB/CUUCGG/1AUD_28_C_B_0_00029.pdb 
# Loading data/PDB/CUUCGG/1AUD_28_C_B_0_00029.pdb 
# Loaded reference data/PDB/CUUCGG/2KOC_5_C_A_0_00001.pdb 
# Loaded target data/PDB/CUUCGG/5el6-pdb-bundle2_1691_C_A_0_00000.pdb 
# Loading data/PDB/CUUCGG/5el6-pdb-bundle2_1691_C_A_0_00000.pdb 
# Loaded reference data/PDB/CUUCGG/2KOC_5_C_A_0_00001.pdb 
# Loaded target data/PDB/CUUCGG/4v7u-pdb-bundle2_1691_C_A_0_00000.pdb 
# Loading data/PDB/CUUCGG/4v7u-pdb-bundle2_1691_C_A_0_00000.pdb 
# Loaded reference data/PDB/CUUCGG/2KOC_5_C_A_0_00001.pdb 
# Loaded target data/PDB/CUUCGG/2N6W_32_C_A_0_00000.pdb 
# Loading data/PDB/CUUCGG/2N6W_32_C_A_0_00000.pdb 
# Loaded reference data/PDB/CUUCGG/2KOC_5_C_A_0_00001.pdb 
# Loaded target data/PDB/CUUCGG/1RAW_21_C_A_0_00005.pdb 
# Loading data/PDB/CUUCGG/1RAW_21_C_A_0_00005.pdb 
# Loaded reference data/PDB/CUUCGG/2KOC_5_C_A_0_00001.pdb 
# Loaded target data/PDB/CUUCGG/5zep-pdb-bundle2_1909_C_A_0_00000.pdb 
# Loading data/PDB/CUUCGG/5zep-pdb-bundle2_1909_C_A_0_00000.pdb 
# Loaded reference data/PDB/CUUCGG/2KOC_5_C_A_0_00001.pdb 
# Loaded target data/PDB/CUUCGG/5j7l-pdb-bundle1_1449_C_A_0_00003.pdb 
# Loading data/PDB/CUUCGG/5j7l-pdb-bundle1_1449_C_A_0_00003.pdb 
# Loaded reference data/PDB/CUUCGG/2KOC_5_C_A_0_00001.pdb 
# Loaded target data/PDB/CUUCGG/1EKZ_13_C_B_0_00020.pdb 
# Loading data/PDB/CUUCGG/1EKZ_13_C_B_0_00020.pdb 
# Loaded reference data/PDB/CUUCGG/2KOC_5_C_A_0_00001.pdb 
# Loaded target data/PDB/CUUCGG/2L2J_19_C_A_0_00001.pdb 
# Loading data/PDB/CUUCGG/2L2J_19_C_A_0_00001.pdb 
# Loaded reference data/PDB/CUUCGG/2KOC_5_C_A_0_00001.pdb 
# Loaded target data/PDB/CUUCGG/5v8i-pdb-bundle1_1691_C_A_0_00000.pdb 
# Loading data/PDB/CUUCGG/5v8i-pdb-bundle1_1691_C_A_0_00000.pdb 
# Loaded reference data/PDB/CUUCGG/2KOC_5_C_A_0_00001.pdb 
# Loaded target data/PDB/CUUCGG/5tbw-pdb-bundle1_1923_C_1_0_00002.pdb 
# Loading data/PDB/CUUCGG/5tbw-pdb-bundle1_1923_C_1_0_00002.pdb 
# Loaded reference data/PDB/CUUCGG/2KOC_5_C_A_0_00001.pdb 
# Loaded target data/PDB/CUUCGG/4v57-pdb-bundle1_207_C_A_0_00000.pdb 
# Loading data/PDB/CUUCGG/4v57-pdb-bundle1_207_C_A_0_00000.pdb 
# Loaded reference data/PDB/CUUCGG/2KOC_5_C_A_0_00001.pdb 
# Loaded target data/PDB/CUUCGG/1EKZ_13_C_B_0_00034.pdb 
# Loading data/PDB/CUUCGG/1EKZ_13_C_B_0_00034.pdb 
# Loaded reference data/PDB/CUUCGG/2KOC_5_C_A_0_00001.pdb 
# Loaded target data/PDB/CUUCGG/2L2J_19_C_A_0_00015.pdb 
# Loading data/PDB/CUUCGG/2L2J_19_C_A_0_00015.pdb 
# Loaded reference data/PDB/CUUCGG/2KOC_5_C_A_0_00001.pdb 
# Loaded target data/PDB/CUUCGG/1F7I_12_C_A_0_00005.pdb 
# Loading data/PDB/CUUCGG/1F7I_12_C_A_0_00005.pdb 
# Loaded reference data/PDB/CUUCGG/2KOC_5_C_A_0_00001.pdb 
# Loaded target data/PDB/CUUCGG/1P5M_28_C_A_0_00004.pdb 
# Loading data/PDB/CUUCGG/1P5M_28_C_A_0_00004.pdb 
# Loaded reference data/PDB/CUUCGG/2KOC_5_C_A_0_00001.pdb 
# Loaded target data/PDB/CUUCGG/1QVF_1769_C_0_0_00000.pdb 
# Loading data/PDB/CUUCGG/1QVF_1769_C_0_0_00000.pdb 
# Loaded reference data/PDB/CUUCGG/2KOC_5_C_A_0_00001.pdb 
# Loaded target data/PDB/CUUCGG/4u25-pdb-bundle4_1691_C_A_0_00000.pdb 
# Loading data/PDB/CUUCGG/4u25-pdb-bundle4_1691_C_A_0_00000.pdb 
# Loaded reference data/PDB/CUUCGG/2KOC_5_C_A_0_00001.pdb 
# Loaded target data/PDB/CUUCGG/6bu8-pdb-bundle1_419_C_0_0_00001.pdb 
# Loading data/PDB/CUUCGG/6bu8-pdb-bundle1_419_C_0_0_00001.pdb 
# Loaded reference data/PDB/CUUCGG/2KOC_5_C_A_0_00001.pdb 
# Loaded target data/PDB/CUUCGG/1EKZ_13_C_B_0_00008.pdb 
# Loading data/PDB/CUUCGG/1EKZ_13_C_B_0_00008.pdb 
# Loaded reference data/PDB/CUUCGG/2KOC_5_C_A_0_00001.pdb 
# Loaded target data/PDB/CUUCGG/2N6T_19_C_A_0_00007.pdb 
# Loading data/PDB/CUUCGG/2N6T_19_C_A_0_00007.pdb 
# Loaded reference data/PDB/CUUCGG/2KOC_5_C_A_0_00001.pdb 
# Loaded target data/PDB/CUUCGG/1ULL_14_C_A_0_00006.pdb 
# Loading data/PDB/CUUCGG/1ULL_14_C_A_0_00006.pdb 
# Loaded reference data/PDB/CUUCGG/2KOC_5_C_A_0_00001.pdb 
# Loaded target data/PDB/CUUCGG/1F7I_12_C_A_0_00011.pdb 
# Loading data/PDB/CUUCGG/1F7I_12_C_A_0_00011.pdb 
# Loaded reference data/PDB/CUUCGG/2KOC_5_C_A_0_00001.pdb 
# Loaded target data/PDB/CUUCGG/4wf1-pdb-bundle4_1691_C_A_0_00000.pdb 
# Loading data/PDB/CUUCGG/4wf1-pdb-bundle4_1691_C_A_0_00000.pdb 
# Loaded reference data/PDB/CUUCGG/2KOC_5_C_A_0_00001.pdb 
# Loaded target data/PDB/CUUCGG/1M5L_17_C_A_0_00009.pdb 
# Loading data/PDB/CUUCGG/1M5L_17_C_A_0_00009.pdb 
# Loaded reference data/PDB/CUUCGG/2KOC_5_C_A_0_00001.pdb 
# Loaded target data/PDB/CUUCGG/1F7I_12_C_A_0_00007.pdb 
# Loading data/PDB/CUUCGG/1F7I_12_C_A_0_00007.pdb 
# Loaded reference data/PDB/CUUCGG/2KOC_5_C_A_0_00001.pdb 
# Loaded target data/PDB/CUUCGG/4v9p-pdb-bundle8_1449_C_A_0_00003.pdb 
# Loading data/PDB/CUUCGG/4v9p-pdb-bundle8_1449_C_A_0_00003.pdb 
# Loaded reference data/PDB/CUUCGG/2KOC_5_C_A_0_00001.pdb 
# Loaded target data/PDB/CUUCGG/1ULL_14_C_A_0_00004.pdb 
# Loading data/PDB/CUUCGG/1ULL_14_C_A_0_00004.pdb 
# Loaded reference data/PDB/CUUCGG/2KOC_5_C_A_0_00001.pdb 
# Loaded target data/PDB/CUUCGG/2N6T_19_C_A_0_00005.pdb 
# Loading data/PDB/CUUCGG/2N6T_19_C_A_0_00005.pdb 
# Loaded reference data/PDB/CUUCGG/2KOC_5_C_A_0_00001.pdb 
# Loaded target data/PDB/CUUCGG/4u50-pdb-bundle3_230_C_d_0_00000.pdb 
# Loading data/PDB/CUUCGG/4u50-pdb-bundle3_230_C_d_0_00000.pdb 
# Loaded reference data/PDB/CUUCGG/2KOC_5_C_A_0_00001.pdb 
# Loaded target data/PDB/CUUCGG/3CC7_1769_C_0_0_00000.pdb 
# Loading data/PDB/CUUCGG/3CC7_1769_C_0_0_00000.pdb 
# Loaded reference data/PDB/CUUCGG/2KOC_5_C_A_0_00001.pdb 
# Loaded target data/PDB/CUUCGG/2L2J_19_C_A_0_00003.pdb 
# Loading data/PDB/CUUCGG/2L2J_19_C_A_0_00003.pdb 
# Loaded reference data/PDB/CUUCGG/2KOC_5_C_A_0_00001.pdb 
# Loaded target data/PDB/CUUCGG/6ha1-pdb-bundle2_1038_C_a_0_00001.pdb 
# Loading data/PDB/CUUCGG/6ha1-pdb-bundle2_1038_C_a_0_00001.pdb 
# Loaded reference data/PDB/CUUCGG/2KOC_5_C_A_0_00001.pdb 
# Loaded target data/PDB/CUUCGG/1EKZ_13_C_B_0_00022.pdb 
# Loading data/PDB/CUUCGG/1EKZ_13_C_B_0_00022.pdb 
# Loaded reference data/PDB/CUUCGG/2KOC_5_C_A_0_00001.pdb 
# Loaded target data/PDB/CUUCGG/2L2J_19_C_A_0_00017.pdb 
# Loading data/PDB/CUUCGG/2L2J_19_C_A_0_00017.pdb 
# Loaded reference data/PDB/CUUCGG/2KOC_5_C_A_0_00001.pdb 
# Loaded target data/PDB/CUUCGG/5vp2-pdb-bundle2_419_C_A_0_00000.pdb 
# Loading data/PDB/CUUCGG/5vp2-pdb-bundle2_419_C_A_0_00000.pdb 
# Loaded reference data/PDB/CUUCGG/2KOC_5_C_A_0_00001.pdb 
# Loaded target data/PDB/CUUCGG/6buw-pdb-bundle3_419_C_A_0_00000.pdb 
# Loading data/PDB/CUUCGG/6buw-pdb-bundle3_419_C_A_0_00000.pdb 
# Loaded reference data/PDB/CUUCGG/2KOC_5_C_A_0_00001.pdb 
# Loaded target data/PDB/CUUCGG/3jbv-pdb-bundle1_1449_C_A_0_00003.pdb 
# Loading data/PDB/CUUCGG/3jbv-pdb-bundle1_1449_C_A_0_00003.pdb 
# Loaded reference data/PDB/CUUCGG/2KOC_5_C_A_0_00001.pdb 
# Loaded target data/PDB/CUUCGG/5apo-pdb-bundle1_1923_C_5_0_00002.pdb 
# Loading data/PDB/CUUCGG/5apo-pdb-bundle1_1923_C_5_0_00002.pdb 
# Loaded reference data/PDB/CUUCGG/2KOC_5_C_A_0_00001.pdb 
# Loaded target data/PDB/CUUCGG/2N6W_32_C_A_0_00002.pdb 
# Loading data/PDB/CUUCGG/2N6W_32_C_A_0_00002.pdb 
# Loaded reference data/PDB/CUUCGG/2KOC_5_C_A_0_00001.pdb 
# Loaded target data/PDB/CUUCGG/1RAW_21_C_A_0_00007.pdb 
# Loading data/PDB/CUUCGG/1RAW_21_C_A_0_00007.pdb 
# Loaded reference data/PDB/CUUCGG/2KOC_5_C_A_0_00001.pdb 
# Loaded target data/PDB/CUUCGG/5nwy-pdb-bundle2_419_C_0_0_00001.pdb 
# Loading data/PDB/CUUCGG/5nwy-pdb-bundle2_419_C_0_0_00001.pdb 
# Loaded reference data/PDB/CUUCGG/2KOC_5_C_A_0_00001.pdb 
# Loaded target data/PDB/CUUCGG/4wro-pdb-bundle4_419_C_A_0_00000.pdb 
# Loading data/PDB/CUUCGG/4wro-pdb-bundle4_419_C_A_0_00000.pdb 
# Loaded reference data/PDB/CUUCGG/2KOC_5_C_A_0_00001.pdb 
# Loaded target data/PDB/CUUCGG/1PBR_11_C_A_0_00000.pdb 
# Loading data/PDB/CUUCGG/1PBR_11_C_A_0_00000.pdb 
# Loaded reference data/PDB/CUUCGG/2KOC_5_C_A_0_00001.pdb 
# Loaded target data/PDB/CUUCGG/4v9c-pdb-bundle1_1028_C_A_0_00002.pdb 
# Loading data/PDB/CUUCGG/4v9c-pdb-bundle1_1028_C_A_0_00002.pdb 
# Loaded reference data/PDB/CUUCGG/2KOC_5_C_A_0_00001.pdb 
# Loaded target data/PDB/CUUCGG/1FYP_11_C_A_0_00017.pdb 
# Loading data/PDB/CUUCGG/1FYP_11_C_A_0_00017.pdb 
# Loaded reference data/PDB/CUUCGG/2KOC_5_C_A_0_00001.pdb 
# Loaded target data/PDB/CUUCGG/1FYP_11_C_A_0_00003.pdb 
# Loading data/PDB/CUUCGG/1FYP_11_C_A_0_00003.pdb 
# Loaded reference data/PDB/CUUCGG/2KOC_5_C_A_0_00001.pdb 
# Loaded target data/PDB/CUUCGG/3j7q-pdb-bundle1_2872_C_5_0_00000.pdb 
# Loading data/PDB/CUUCGG/3j7q-pdb-bundle1_2872_C_5_0_00000.pdb 
# Loaded reference data/PDB/CUUCGG/2KOC_5_C_A_0_00001.pdb 
# Loaded target data/PDB/CUUCGG/4v83-pdb-bundle1_419_C_A_0_00000.pdb 
# Loading data/PDB/CUUCGG/4v83-pdb-bundle1_419_C_A_0_00000.pdb 
# Loaded reference data/PDB/CUUCGG/2KOC_5_C_A_0_00001.pdb 
# Loaded target data/PDB/CUUCGG/1AUD_28_C_B_0_00003.pdb 
# Loading data/PDB/CUUCGG/1AUD_28_C_B_0_00003.pdb 
# Loaded reference data/PDB/CUUCGG/2KOC_5_C_A_0_00001.pdb 
# Loaded target data/PDB/CUUCGG/1EBS_12_C_A_0_00004.pdb 
# Loading data/PDB/CUUCGG/1EBS_12_C_A_0_00004.pdb 
# Loaded reference data/PDB/CUUCGG/2KOC_5_C_A_0_00001.pdb 
# Loaded target data/PDB/CUUCGG/1AUD_28_C_B_0_00017.pdb 
# Loading data/PDB/CUUCGG/1AUD_28_C_B_0_00017.pdb 
# Loaded reference data/PDB/CUUCGG/2KOC_5_C_A_0_00001.pdb 
# Loaded target data/PDB/CUUCGG/6dzi-pdb-bundle2_1909_C_A_0_00000.pdb 
# Loading data/PDB/CUUCGG/6dzi-pdb-bundle2_1909_C_A_0_00000.pdb 
# Loaded reference data/PDB/CUUCGG/2KOC_5_C_A_0_00001.pdb 
# Loaded target data/PDB/CUUCGG/5Y85_28_C_D_0_00001.pdb 
# Loading data/PDB/CUUCGG/5Y85_28_C_D_0_00001.pdb 
# Loaded reference data/PDB/CUUCGG/2KOC_5_C_A_0_00001.pdb 
# Loaded target data/PDB/CUUCGG/1M82_10_C_A_0_00007.pdb 
# Loading data/PDB/CUUCGG/1M82_10_C_A_0_00007.pdb 
# Loaded reference data/PDB/CUUCGG/2KOC_5_C_A_0_00001.pdb 
# Loaded target data/PDB/CUUCGG/1M82_10_C_A_0_00013.pdb 
# Loading data/PDB/CUUCGG/1M82_10_C_A_0_00013.pdb 
# Loaded reference data/PDB/CUUCGG/2KOC_5_C_A_0_00001.pdb 
# Loaded target data/PDB/CUUCGG/5nwy-pdb-bundle2_1028_C_0_0_00002.pdb 
# Loading data/PDB/CUUCGG/5nwy-pdb-bundle2_1028_C_0_0_00002.pdb 
# Loaded reference data/PDB/CUUCGG/2KOC_5_C_A_0_00001.pdb 
# Loaded target data/PDB/CUUCGG/5hcp-pdb-bundle3_1737_C_A_0_00000.pdb 
# Loading data/PDB/CUUCGG/5hcp-pdb-bundle3_1737_C_A_0_00000.pdb 
# Loaded reference data/PDB/CUUCGG/2KOC_5_C_A_0_00001.pdb 
# Loaded target data/PDB/CUUCGG/4v8b-pdb-bundle1_419_C_A_0_00000.pdb 
# Loading data/PDB/CUUCGG/4v8b-pdb-bundle1_419_C_A_0_00000.pdb 
# Loaded reference data/PDB/CUUCGG/2KOC_5_C_A_0_00001.pdb 
# Loaded target data/PDB/CUUCGG/5dc3-pdb-bundle3_230_C_b_0_00000.pdb 
# Loading data/PDB/CUUCGG/5dc3-pdb-bundle3_230_C_b_0_00000.pdb 
# Loaded reference data/PDB/CUUCGG/2KOC_5_C_A_0_00001.pdb 
# Loaded target data/PDB/CUUCGG/4u51-pdb-bundle2_1923_C_A_0_00002.pdb 
# Loading data/PDB/CUUCGG/4u51-pdb-bundle2_1923_C_A_0_00002.pdb 
# Loaded reference data/PDB/CUUCGG/2KOC_5_C_A_0_00001.pdb 
# Loaded target data/PDB/CUUCGG/2M4Q_11_C_1_0_00003.pdb 
# Loading data/PDB/CUUCGG/2M4Q_11_C_1_0_00003.pdb 
# Loaded reference data/PDB/CUUCGG/2KOC_5_C_A_0_00001.pdb 
# Loaded target data/PDB/CUUCGG/4v54-pdb-bundle3_207_C_K_0_00000.pdb 
# Loading data/PDB/CUUCGG/4v54-pdb-bundle3_207_C_K_0_00000.pdb 
# Loaded reference data/PDB/CUUCGG/2KOC_5_C_A_0_00001.pdb 
# Loaded target data/PDB/CUUCGG/1NJI_1769_C_A_0_00000.pdb 
# Loading data/PDB/CUUCGG/1NJI_1769_C_A_0_00000.pdb 
# Loaded reference data/PDB/CUUCGG/2KOC_5_C_A_0_00001.pdb 
# Loaded target data/PDB/CUUCGG/5dge-pdb-bundle4_1923_C_I_0_00002.pdb 
# Loading data/PDB/CUUCGG/5dge-pdb-bundle4_1923_C_I_0_00002.pdb 
# Loaded reference data/PDB/CUUCGG/2KOC_5_C_A_0_00001.pdb 
# Loaded target data/PDB/CUUCGG/2KOC_5_C_A_0_00004.pdb 
# Loading data/PDB/CUUCGG/2KOC_5_C_A_0_00004.pdb 
# Loaded reference data/PDB/CUUCGG/2KOC_5_C_A_0_00001.pdb 
# Loaded target data/PDB/CUUCGG/4u27-pdb-bundle1_1449_C_A_0_00003.pdb 
# Loading data/PDB/CUUCGG/4u27-pdb-bundle1_1449_C_A_0_00003.pdb 
# Loaded reference data/PDB/CUUCGG/2KOC_5_C_A_0_00001.pdb 
# Loaded target data/PDB/CUUCGG/2KOC_5_C_A_0_00010.pdb 
# Loading data/PDB/CUUCGG/2KOC_5_C_A_0_00010.pdb 
# Loaded reference data/PDB/CUUCGG/2KOC_5_C_A_0_00001.pdb 
# Loaded target data/PDB/CUUCGG/2KMJ_29_C_A_0_00009.pdb 
# Loading data/PDB/CUUCGG/2KMJ_29_C_A_0_00009.pdb 
# Loaded reference data/PDB/CUUCGG/2KOC_5_C_A_0_00001.pdb 
# Loaded target data/PDB/CUUCGG/1G70_54_C_A_0_00013.pdb 
# Loading data/PDB/CUUCGG/1G70_54_C_A_0_00013.pdb 
# Loaded reference data/PDB/CUUCGG/2KOC_5_C_A_0_00001.pdb 
# Loaded target data/PDB/CUUCGG/1G70_54_C_A_0_00007.pdb 
# Loading data/PDB/CUUCGG/1G70_54_C_A_0_00007.pdb 
# Loaded reference data/PDB/CUUCGG/2KOC_5_C_A_0_00001.pdb 
# Loaded target data/PDB/CUUCGG/4wr6-pdb-bundle1_419_C_A_0_00000.pdb 
# Loading data/PDB/CUUCGG/4wr6-pdb-bundle1_419_C_A_0_00000.pdb 
# Loaded reference data/PDB/CUUCGG/2KOC_5_C_A_0_00001.pdb 
# Loaded target data/PDB/CUUCGG/1IKD_7_C_A_0_00026.pdb 
# Loading data/PDB/CUUCGG/1IKD_7_C_A_0_00026.pdb 
# Loaded reference data/PDB/CUUCGG/2KOC_5_C_A_0_00001.pdb 
# Loaded target data/PDB/CUUCGG/1Z31_270_C_A_0_00004.pdb 
# Loading data/PDB/CUUCGG/1Z31_270_C_A_0_00004.pdb 
# Loaded reference data/PDB/CUUCGG/2KOC_5_C_A_0_00001.pdb 
# Loaded target data/PDB/CUUCGG/4v4q-pdb-bundle1_419_C_A_0_00001.pdb 
# Loading data/PDB/CUUCGG/4v4q-pdb-bundle1_419_C_A_0_00001.pdb 
# Loaded reference data/PDB/CUUCGG/2KOC_5_C_A_0_00001.pdb 
# Loaded target data/PDB/CUUCGG/6i7v-pdb-bundle1_419_C_B_0_00005.pdb 
# Loading data/PDB/CUUCGG/6i7v-pdb-bundle1_419_C_B_0_00005.pdb 
# Loaded reference data/PDB/CUUCGG/2KOC_5_C_A_0_00001.pdb 
# Loaded target data/PDB/CUUCGG/1F7H_12_C_A_0_00009.pdb 
# Loading data/PDB/CUUCGG/1F7H_12_C_A_0_00009.pdb 
# Loaded reference data/PDB/CUUCGG/2KOC_5_C_A_0_00001.pdb 
# Loaded target data/PDB/CUUCGG/2L2K_19_C_A_0_00019.pdb 
# Loading data/PDB/CUUCGG/2L2K_19_C_A_0_00019.pdb 
# Loaded reference data/PDB/CUUCGG/2KOC_5_C_A_0_00001.pdb 
# Loaded target data/PDB/CUUCGG/5hau-pdb-bundle2_415_C_A_0_00000.pdb 
# Loading data/PDB/CUUCGG/5hau-pdb-bundle2_415_C_A_0_00000.pdb 
# Loaded reference data/PDB/CUUCGG/2KOC_5_C_A_0_00001.pdb 
# Loaded target data/PDB/CUUCGG/5WNU_419_C_A_0_00000.pdb 
# Loading data/PDB/CUUCGG/5WNU_419_C_A_0_00000.pdb 
# Loaded reference data/PDB/CUUCGG/2KOC_5_C_A_0_00001.pdb 
# Loaded target data/PDB/CUUCGG/4v8g-pdb-bundle3_419_C_A_0_00000.pdb 
# Loading data/PDB/CUUCGG/4v8g-pdb-bundle3_419_C_A_0_00000.pdb 
# Loaded reference data/PDB/CUUCGG/2KOC_5_C_A_0_00001.pdb 
# Loaded target data/PDB/CUUCGG/1F7G_12_C_A_0_00003.pdb 
# Loading data/PDB/CUUCGG/1F7G_12_C_A_0_00003.pdb 
# Loaded reference data/PDB/CUUCGG/2KOC_5_C_A_0_00001.pdb 
# Loaded target data/PDB/CUUCGG/4v6c-pdb-bundle1_1449_C_U_0_00003.pdb 
# Loading data/PDB/CUUCGG/4v6c-pdb-bundle1_1449_C_U_0_00003.pdb 
# Loaded reference data/PDB/CUUCGG/2KOC_5_C_A_0_00001.pdb 
# Loaded target data/PDB/CUUCGG/2HGH_32_C_B_0_00002.pdb 
# Loading data/PDB/CUUCGG/2HGH_32_C_B_0_00002.pdb 
# Loaded reference data/PDB/CUUCGG/2KOC_5_C_A_0_00001.pdb 
# Loaded target data/PDB/CUUCGG/2HGH_32_C_B_0_00016.pdb 
# Loading data/PDB/CUUCGG/2HGH_32_C_B_0_00016.pdb 
# Loaded reference data/PDB/CUUCGG/2KOC_5_C_A_0_00001.pdb 
# Loaded target data/PDB/CUUCGG/1C0O_5_C_A_0_00004.pdb 
# Loading data/PDB/CUUCGG/1C0O_5_C_A_0_00004.pdb 
# Loaded reference data/PDB/CUUCGG/2KOC_5_C_A_0_00001.pdb 
# Loaded target data/PDB/CUUCGG/5ib8-pdb-bundle4_1691_C_A_0_00000.pdb 
# Loading data/PDB/CUUCGG/5ib8-pdb-bundle4_1691_C_A_0_00000.pdb 
# Loaded reference data/PDB/CUUCGG/2KOC_5_C_A_0_00001.pdb 
# Loaded target data/PDB/CUUCGG/1C0O_5_C_A_0_00010.pdb 
# Loading data/PDB/CUUCGG/1C0O_5_C_A_0_00010.pdb 
# Loaded reference data/PDB/CUUCGG/2KOC_5_C_A_0_00001.pdb 
# Loaded target data/PDB/CUUCGG/1C0O_5_C_A_0_00011.pdb 
# Loading data/PDB/CUUCGG/1C0O_5_C_A_0_00011.pdb 
# Loaded reference data/PDB/CUUCGG/2KOC_5_C_A_0_00001.pdb 
# Loaded target data/PDB/CUUCGG/1C0O_5_C_A_0_00005.pdb 
# Loading data/PDB/CUUCGG/1C0O_5_C_A_0_00005.pdb 
# Loaded reference data/PDB/CUUCGG/2KOC_5_C_A_0_00001.pdb 
# Loaded target data/PDB/CUUCGG/4v9d-pdb-bundle1_419_C_Y_0_00005.pdb 
# Loading data/PDB/CUUCGG/4v9d-pdb-bundle1_419_C_Y_0_00005.pdb 
# Loaded reference data/PDB/CUUCGG/2KOC_5_C_A_0_00001.pdb 
# Loaded target data/PDB/CUUCGG/6i7v-pdb-bundle3_1691_C_A_0_00000.pdb 
# Loading data/PDB/CUUCGG/6i7v-pdb-bundle3_1691_C_A_0_00000.pdb 
# Loaded reference data/PDB/CUUCGG/2KOC_5_C_A_0_00001.pdb 
# Loaded target data/PDB/CUUCGG/2HGH_32_C_B_0_00017.pdb 
# Loading data/PDB/CUUCGG/2HGH_32_C_B_0_00017.pdb 
# Loaded reference data/PDB/CUUCGG/2KOC_5_C_A_0_00001.pdb 
# Loaded target data/PDB/CUUCGG/1F7G_12_C_A_0_00016.pdb 
# Loading data/PDB/CUUCGG/1F7G_12_C_A_0_00016.pdb 
# Loaded reference data/PDB/CUUCGG/2KOC_5_C_A_0_00001.pdb 
# Loaded target data/PDB/CUUCGG/4wqu-pdb-bundle1_1738_C_A_0_00000.pdb 
# Loading data/PDB/CUUCGG/4wqu-pdb-bundle1_1738_C_A_0_00000.pdb 
# Loaded reference data/PDB/CUUCGG/2KOC_5_C_A_0_00001.pdb 
# Loaded target data/PDB/CUUCGG/1F7G_12_C_A_0_00002.pdb 
# Loading data/PDB/CUUCGG/1F7G_12_C_A_0_00002.pdb 
# Loaded reference data/PDB/CUUCGG/2KOC_5_C_A_0_00001.pdb 
# Loaded target data/PDB/CUUCGG/2HGH_32_C_B_0_00003.pdb 
# Loading data/PDB/CUUCGG/2HGH_32_C_B_0_00003.pdb 
# Loaded reference data/PDB/CUUCGG/2KOC_5_C_A_0_00001.pdb 
# Loaded target data/PDB/CUUCGG/4w2h-pdb-bundle2_1738_C_A_0_00000.pdb 
# Loading data/PDB/CUUCGG/4w2h-pdb-bundle2_1738_C_A_0_00000.pdb 
# Loaded reference data/PDB/CUUCGG/2KOC_5_C_A_0_00001.pdb 
# Loaded target data/PDB/CUUCGG/5lze-pdb-bundle1_419_C_a_0_00001.pdb 
# Loading data/PDB/CUUCGG/5lze-pdb-bundle1_419_C_a_0_00001.pdb 
# Loaded reference data/PDB/CUUCGG/2KOC_5_C_A_0_00001.pdb 
# Loaded target data/PDB/CUUCGG/2L2K_19_C_A_0_00018.pdb 
# Loading data/PDB/CUUCGG/2L2K_19_C_A_0_00018.pdb 
# Loaded reference data/PDB/CUUCGG/2KOC_5_C_A_0_00001.pdb 
# Loaded target data/PDB/CUUCGG/4v57-pdb-bundle3_207_C_A_0_00000.pdb 
# Loading data/PDB/CUUCGG/4v57-pdb-bundle3_207_C_A_0_00000.pdb 
# Loaded reference data/PDB/CUUCGG/2KOC_5_C_A_0_00001.pdb 
# Loaded target data/PDB/CUUCGG/6d90-pdb-bundle2_2872_C_5_0_00000.pdb 
# Loading data/PDB/CUUCGG/6d90-pdb-bundle2_2872_C_5_0_00000.pdb 
# Loaded reference data/PDB/CUUCGG/2KOC_5_C_A_0_00001.pdb 
# Loaded target data/PDB/CUUCGG/3G4S_1769_C_0_0_00000.pdb 
# Loading data/PDB/CUUCGG/3G4S_1769_C_0_0_00000.pdb 
# Loaded reference data/PDB/CUUCGG/2KOC_5_C_A_0_00001.pdb 
# Loaded target data/PDB/CUUCGG/1Z31_270_C_A_0_00005.pdb 
# Loading data/PDB/CUUCGG/1Z31_270_C_A_0_00005.pdb 
# Loaded reference data/PDB/CUUCGG/2KOC_5_C_A_0_00001.pdb 
# Loaded target data/PDB/CUUCGG/1IKD_7_C_A_0_00027.pdb 
# Loading data/PDB/CUUCGG/1IKD_7_C_A_0_00027.pdb 
# Loaded reference data/PDB/CUUCGG/2KOC_5_C_A_0_00001.pdb 
# Loaded target data/PDB/CUUCGG/4v9k-pdb-bundle4_1691_C_A_0_00000.pdb 
# Loading data/PDB/CUUCGG/4v9k-pdb-bundle4_1691_C_A_0_00000.pdb 
# Loaded reference data/PDB/CUUCGG/2KOC_5_C_A_0_00001.pdb 
# Loaded target data/PDB/CUUCGG/1G70_54_C_A_0_00006.pdb 
# Loading data/PDB/CUUCGG/1G70_54_C_A_0_00006.pdb 
# Loaded reference data/PDB/CUUCGG/2KOC_5_C_A_0_00001.pdb 
# Loaded target data/PDB/CUUCGG/2KMJ_29_C_A_0_00008.pdb 
# Loading data/PDB/CUUCGG/2KMJ_29_C_A_0_00008.pdb 
# Loaded reference data/PDB/CUUCGG/2KOC_5_C_A_0_00001.pdb 
# Loaded target data/PDB/CUUCGG/1G70_54_C_A_0_00012.pdb 
# Loading data/PDB/CUUCGG/1G70_54_C_A_0_00012.pdb 
# Loaded reference data/PDB/CUUCGG/2KOC_5_C_A_0_00001.pdb 
# Loaded target data/PDB/CUUCGG/5tgm-pdb-bundle4_1923_C_I_0_00002.pdb 
# Loading data/PDB/CUUCGG/5tgm-pdb-bundle4_1923_C_I_0_00002.pdb 
# Loaded reference data/PDB/CUUCGG/2KOC_5_C_A_0_00001.pdb 
# Loaded target data/PDB/CUUCGG/4v6g-pdb-bundle4_1691_C_A_0_00000.pdb 
# Loading data/PDB/CUUCGG/4v6g-pdb-bundle4_1691_C_A_0_00000.pdb 
# Loaded reference data/PDB/CUUCGG/2KOC_5_C_A_0_00001.pdb 
# Loaded target data/PDB/CUUCGG/2KOC_5_C_A_0_00011.pdb 
# Loading data/PDB/CUUCGG/2KOC_5_C_A_0_00011.pdb 
# Loaded reference data/PDB/CUUCGG/2KOC_5_C_A_0_00001.pdb 
# Loaded target data/PDB/CUUCGG/2KOC_5_C_A_0_00005.pdb 
# Loading data/PDB/CUUCGG/2KOC_5_C_A_0_00005.pdb 
# Loaded reference data/PDB/CUUCGG/2KOC_5_C_A_0_00001.pdb 
# Loaded target data/PDB/CUUCGG/1SM1_1708_C_0_0_00000.pdb 
# Loading data/PDB/CUUCGG/1SM1_1708_C_0_0_00000.pdb 
# Loaded reference data/PDB/CUUCGG/2KOC_5_C_A_0_00001.pdb 
# Loaded target data/PDB/CUUCGG/6h4n-pdb-bundle1_419_C_a_0_00001.pdb 
# Loading data/PDB/CUUCGG/6h4n-pdb-bundle1_419_C_a_0_00001.pdb 
# Loaded reference data/PDB/CUUCGG/2KOC_5_C_A_0_00001.pdb 
# Loaded target data/PDB/CUUCGG/2CD6_12_C_A_0_00000.pdb 
# Loading data/PDB/CUUCGG/2CD6_12_C_A_0_00000.pdb 
# Loaded reference data/PDB/CUUCGG/2KOC_5_C_A_0_00001.pdb 
# Loaded target data/PDB/CUUCGG/4v50-pdb-bundle1_1449_C_A_0_00003.pdb 
# Loading data/PDB/CUUCGG/4v50-pdb-bundle1_1449_C_A_0_00003.pdb 
# Loaded reference data/PDB/CUUCGG/2KOC_5_C_A_0_00001.pdb 
# Loaded target data/PDB/CUUCGG/1DK1_32_C_B_0_00001.pdb 
# Loading data/PDB/CUUCGG/1DK1_32_C_B_0_00001.pdb 
# Loaded reference data/PDB/CUUCGG/2KOC_5_C_A_0_00001.pdb 
# Loaded target data/PDB/CUUCGG/4v52-pdb-bundle1_207_C_A_0_00000.pdb 
# Loading data/PDB/CUUCGG/4v52-pdb-bundle1_207_C_A_0_00000.pdb 
# Loaded reference data/PDB/CUUCGG/2KOC_5_C_A_0_00001.pdb 
# Loaded target data/PDB/CUUCGG/3CC2_1769_C_0_0_00000.pdb 
# Loading data/PDB/CUUCGG/3CC2_1769_C_0_0_00000.pdb 
# Loaded reference data/PDB/CUUCGG/2KOC_5_C_A_0_00001.pdb 
# Loaded target data/PDB/CUUCGG/6fyx-pdb-bundle1_229_C_2_0_00000.pdb 
# Loading data/PDB/CUUCGG/6fyx-pdb-bundle1_229_C_2_0_00000.pdb 
# Loaded reference data/PDB/CUUCGG/2KOC_5_C_A_0_00001.pdb 
# Loaded target data/PDB/CUUCGG/1M82_10_C_A_0_00012.pdb 
# Loading data/PDB/CUUCGG/1M82_10_C_A_0_00012.pdb 
# Loaded reference data/PDB/CUUCGG/2KOC_5_C_A_0_00001.pdb 
# Loaded target data/PDB/CUUCGG/5j4b-pdb-bundle2_419_C_A_0_00000.pdb 
# Loading data/PDB/CUUCGG/5j4b-pdb-bundle2_419_C_A_0_00000.pdb 
# Loaded reference data/PDB/CUUCGG/2KOC_5_C_A_0_00001.pdb 
# Loaded target data/PDB/CUUCGG/1M82_10_C_A_0_00006.pdb 
# Loading data/PDB/CUUCGG/1M82_10_C_A_0_00006.pdb 
# Loaded reference data/PDB/CUUCGG/2KOC_5_C_A_0_00001.pdb 
# Loaded target data/PDB/CUUCGG/1JZY_1708_C_A_0_00000.pdb 
# Loading data/PDB/CUUCGG/1JZY_1708_C_A_0_00000.pdb 
# Loaded reference data/PDB/CUUCGG/2KOC_5_C_A_0_00001.pdb 
# Loaded target data/PDB/CUUCGG/1AUD_28_C_B_0_00016.pdb 
# Loading data/PDB/CUUCGG/1AUD_28_C_B_0_00016.pdb 
# Loaded reference data/PDB/CUUCGG/2KOC_5_C_A_0_00001.pdb 
# Loaded target data/PDB/CUUCGG/1AUD_28_C_B_0_00002.pdb 
# Loading data/PDB/CUUCGG/1AUD_28_C_B_0_00002.pdb 
# Loaded reference data/PDB/CUUCGG/2KOC_5_C_A_0_00001.pdb 
# Loaded target data/PDB/CUUCGG/1FYP_11_C_A_0_00002.pdb 
# Loading data/PDB/CUUCGG/1FYP_11_C_A_0_00002.pdb 
# Loaded reference data/PDB/CUUCGG/2KOC_5_C_A_0_00001.pdb 
# Loaded target data/PDB/CUUCGG/5o2r-pdb-bundle2_419_C_a_0_00001.pdb 
# Loading data/PDB/CUUCGG/5o2r-pdb-bundle2_419_C_a_0_00001.pdb 
# Loaded reference data/PDB/CUUCGG/2KOC_5_C_A_0_00001.pdb 
# Loaded target data/PDB/CUUCGG/6bz6-pdb-bundle3_419_C_A_0_00000.pdb 
# Loading data/PDB/CUUCGG/6bz6-pdb-bundle3_419_C_A_0_00000.pdb 
# Loaded reference data/PDB/CUUCGG/2KOC_5_C_A_0_00001.pdb 
# Loaded target data/PDB/CUUCGG/5WNT_419_C_A_0_00000.pdb 
# Loading data/PDB/CUUCGG/5WNT_419_C_A_0_00000.pdb 
# Loaded reference data/PDB/CUUCGG/2KOC_5_C_A_0_00001.pdb 
# Loaded target data/PDB/CUUCGG/4u3u-pdb-bundle4_1923_C_I_0_00002.pdb 
# Loading data/PDB/CUUCGG/4u3u-pdb-bundle4_1923_C_I_0_00002.pdb 
# Loaded reference data/PDB/CUUCGG/2KOC_5_C_A_0_00001.pdb 
# Loaded target data/PDB/CUUCGG/1FYP_11_C_A_0_00016.pdb 
# Loading data/PDB/CUUCGG/1FYP_11_C_A_0_00016.pdb 
# Loaded reference data/PDB/CUUCGG/2KOC_5_C_A_0_00001.pdb 
# Loaded target data/PDB/CUUCGG/4JF2_60_C_A_0_00000.pdb 
# Loading data/PDB/CUUCGG/4JF2_60_C_A_0_00000.pdb 
# Loaded reference data/PDB/CUUCGG/2KOC_5_C_A_0_00001.pdb 
# Loaded target data/PDB/CUUCGG/5el7-pdb-bundle1_419_C_A_0_00000.pdb 
# Loading data/PDB/CUUCGG/5el7-pdb-bundle1_419_C_A_0_00000.pdb 
# Loaded reference data/PDB/CUUCGG/2KOC_5_C_A_0_00001.pdb 
# Loaded target data/PDB/CUUCGG/4ybb-pdb-bundle1_419_C_A_0_00001.pdb 
# Loading data/PDB/CUUCGG/4ybb-pdb-bundle1_419_C_A_0_00001.pdb 
# Loaded reference data/PDB/CUUCGG/2KOC_5_C_A_0_00001.pdb 
# Loaded target data/PDB/CUUCGG/1RAW_21_C_A_0_00006.pdb 
# Loading data/PDB/CUUCGG/1RAW_21_C_A_0_00006.pdb 
# Loaded reference data/PDB/CUUCGG/2KOC_5_C_A_0_00001.pdb 
# Loaded target data/PDB/CUUCGG/2N6W_32_C_A_0_00003.pdb 
# Loading data/PDB/CUUCGG/2N6W_32_C_A_0_00003.pdb 
# Loaded reference data/PDB/CUUCGG/2KOC_5_C_A_0_00001.pdb 
# Loaded target data/PDB/CUUCGG/4v64-pdb-bundle3_207_C_A_0_00000.pdb 
# Loading data/PDB/CUUCGG/4v64-pdb-bundle3_207_C_A_0_00000.pdb 
# Loaded reference data/PDB/CUUCGG/2KOC_5_C_A_0_00001.pdb 
# Loaded target data/PDB/CUUCGG/2L2J_19_C_A_0_00016.pdb 
# Loading data/PDB/CUUCGG/2L2J_19_C_A_0_00016.pdb 
# Loaded reference data/PDB/CUUCGG/2KOC_5_C_A_0_00001.pdb 
# Loaded target data/PDB/CUUCGG/4v9i-pdb-bundle2_1737_C_A_0_00000.pdb 
# Loading data/PDB/CUUCGG/4v9i-pdb-bundle2_1737_C_A_0_00000.pdb 
# Loaded reference data/PDB/CUUCGG/2KOC_5_C_A_0_00001.pdb 
# Loaded target data/PDB/CUUCGG/5dgf-pdb-bundle4_1923_C_C_0_00002.pdb 
# Loading data/PDB/CUUCGG/5dgf-pdb-bundle4_1923_C_C_0_00002.pdb 
# Loaded reference data/PDB/CUUCGG/2KOC_5_C_A_0_00001.pdb 
# Loaded target data/PDB/CUUCGG/4v8d-pdb-bundle4_1691_C_A_0_00000.pdb 
# Loading data/PDB/CUUCGG/4v8d-pdb-bundle4_1691_C_A_0_00000.pdb 
# Loaded reference data/PDB/CUUCGG/2KOC_5_C_A_0_00001.pdb 
# Loaded target data/PDB/CUUCGG/2L2J_19_C_A_0_00002.pdb 
# Loading data/PDB/CUUCGG/2L2J_19_C_A_0_00002.pdb 
# Loaded reference data/PDB/CUUCGG/2KOC_5_C_A_0_00001.pdb 
# Loaded target data/PDB/CUUCGG/1VQO_1769_C_0_0_00000.pdb 
# Loading data/PDB/CUUCGG/1VQO_1769_C_0_0_00000.pdb 
# Loaded reference data/PDB/CUUCGG/2KOC_5_C_A_0_00001.pdb 
# Loaded target data/PDB/CUUCGG/1P5M_28_C_A_0_00013.pdb 
# Loading data/PDB/CUUCGG/1P5M_28_C_A_0_00013.pdb 
# Loaded reference data/PDB/CUUCGG/2KOC_5_C_A_0_00001.pdb 
# Loaded target data/PDB/CUUCGG/2N6T_19_C_A_0_00004.pdb 
# Loading data/PDB/CUUCGG/2N6T_19_C_A_0_00004.pdb 
# Loaded reference data/PDB/CUUCGG/2KOC_5_C_A_0_00001.pdb 
# Loaded target data/PDB/CUUCGG/1ULL_14_C_A_0_00005.pdb 
# Loading data/PDB/CUUCGG/1ULL_14_C_A_0_00005.pdb 
# Loaded reference data/PDB/CUUCGG/2KOC_5_C_A_0_00001.pdb 
# Loaded target data/PDB/CUUCGG/1M5L_17_C_A_0_00008.pdb 
# Loading data/PDB/CUUCGG/1M5L_17_C_A_0_00008.pdb 
# Loaded reference data/PDB/CUUCGG/2KOC_5_C_A_0_00001.pdb 
# Loaded target data/PDB/CUUCGG/1F7I_12_C_A_0_00006.pdb 
# Loading data/PDB/CUUCGG/1F7I_12_C_A_0_00006.pdb 
# Loaded reference data/PDB/CUUCGG/2KOC_5_C_A_0_00001.pdb 
# Loaded target data/PDB/CUUCGG/1P5M_28_C_A_0_00007.pdb 
# Loading data/PDB/CUUCGG/1P5M_28_C_A_0_00007.pdb
```

In [17]:

```
cols = ["black","medium blue","pumpkin","brick red","black","white"]
cp = sns.xkcd_palette(cols)

lli = glob.glob("data/PDB/CUUCGG/*.pdb")
ii_2koc = [k for k in range(len(lli)) if ("2KOC" in lli[k]) ] 
ii_6by5 = [k for k in range(len(lli)) if ("6BY5" in lli[k]) ] 
dists = np.array(dists)
dists1 = np.sqrt(np.sum(dists[:,1,4,:]**2,axis=1))

fig,ax = plt.subplots(1,1,figsize=(7.5, 6))
plt.subplots_adjust(left=0.1, bottom=0.1, right=0.95, top=0.98, wspace=0.07, hspace=0.1)

ax.set_xlim(0,1.9)
ax.set_ylim(0,2.2)
CS = ax.contourf(X, Y, Z,levels=levels,cmap=cmap)
CB = fig.colorbar(CS)

ax.scatter(ermsd_pdbs,dists1,s=1.5,c='0.2',label='PDB structures')

ax.scatter(np.array(ermsd_pdbs)[ii_2koc],dists1[ii_2koc],color=cp[2],label="2KOC",marker="s",s=25)
ax.scatter(np.array(ermsd_pdbs)[ii_6by5],dists1[ii_6by5],color=cp[3],label="6BY5",marker="s",s=25)
ax.legend(ncol=1,loc="lower right")
ax.set_xlabel("eRMSD from native (residues 5-10)")
ax.set_ylabel("C6-G9 distance (nm)")
rect = patches.Rectangle((region_A[0][0],region_A[0][1]),region_A[1][0]-region_A[0][0],region_A[1][1]-region_A[0][1],\
                         linewidth=0.5,edgecolor='k',facecolor='none')
ax.add_patch(rect)
rect = patches.Rectangle((region_B[0][0],region_B[0][1]),region_B[1][0]-region_B[0][0],region_B[1][1]-region_B[0][1],\
                         linewidth=0.5,edgecolor='k',facecolor='none')
ax.add_patch(rect)
plt.savefig("figures/2d_projection.png",dpi=600)
plt.savefig("figures/2d_projection.pdf")
plt.show()
plt.close()
```

In [18]:

```
for j in range(len(ermsd_pdbs)):
    if(ermsd_pdbs[j]>1.0 and dists1[j] < 0.75):
        print(j,ermsd_pdbs[j],dists1[j],ll[j])
```

```
7 [1.24982076] 0.6290139006274563 data/PDB/CUUCGG/1F7F_12_C_A_0_00001.pdb
13 [1.04602921] 0.6878930327761845 data/PDB/CUUCGG/1AUD_28_C_B_0_00027.pdb
66 [1.28380599] 0.6079394999035834 data/PDB/CUUCGG/1F7H_12_C_A_0_00010.pdb
91 [1.08355804] 0.6353363899248905 data/PDB/CUUCGG/1EBQ_12_C_A_0_00003.pdb
96 [1.06096606] 0.5912036670630493 data/PDB/CUUCGG/1EKZ_13_C_B_0_00013.pdb
110 [1.22854421] 0.5978147961954856 data/PDB/CUUCGG/1RAW_21_C_A_0_00008.pdb
111 [1.15350321] 0.6370022883056055 data/PDB/CUUCGG/1EBQ_12_C_A_0_00001.pdb
142 [1.24638836] 0.6160342088299764 data/PDB/CUUCGG/1F7H_12_C_A_0_00006.pdb
157 [1.52083807] 0.6671719782529976 data/PDB/CUUCGG/5ngm-pdb-bundle1_1038_C_A_0_00002.pdb
177 [1.17447896] 0.6504740734302774 data/PDB/CUUCGG/1EBR_12_C_A_0_00004.pdb
197 [1.30300945] 0.6209482504333197 data/PDB/CUUCGG/1F7F_12_C_A_0_00003.pdb
198 [1.16994135] 0.6290384416832708 data/PDB/CUUCGG/1EBQ_12_C_A_0_00000.pdb
199 [1.43486036] 0.46994279988645443 data/PDB/CUUCGG/1RAW_21_C_A_0_00009.pdb
211 [1.23473408] 0.6762540137898687 data/PDB/CUUCGG/1F7I_12_C_A_0_00009.pdb
221 [1.3029368] 0.5906035869980171 data/PDB/CUUCGG/1EKZ_13_C_B_0_00000.pdb
225 [1.13040826] 0.6265701899320947 data/PDB/CUUCGG/1EBQ_12_C_A_0_00004.pdb
242 [1.13356836] 0.7344769072422058 data/PDB/CUUCGG/1P6V_26_C_B_0_00000.pdb
266 [1.15353868] 0.6484137823823877 data/PDB/CUUCGG/1F7G_12_C_A_0_00009.pdb
307 [1.26618048] 0.6118273326895443 data/PDB/CUUCGG/1F7F_12_C_A_0_00006.pdb
349 [1.1857004] 0.6309708718375122 data/PDB/CUUCGG/1EBR_12_C_A_0_00003.pdb
380 [1.04907664] 0.6742985512265377 data/PDB/CUUCGG/2KZL_27_C_A_0_00005.pdb
397 [1.22987884] 0.6528579231140934 data/PDB/CUUCGG/1EBR_12_C_A_0_00002.pdb
412 [1.26314972] 0.6162288385348497 data/PDB/CUUCGG/1F7F_12_C_A_0_00005.pdb
413 [1.02764757] 0.7031338438940504 data/PDB/CUUCGG/1EKZ_13_C_B_0_00016.pdb
451 [1.25345605] 0.6554802721563805 data/PDB/CUUCGG/4v6c-pdb-bundle1_207_C_U_0_00000.pdb
461 [1.24991455] 0.6897632818582633 data/PDB/CUUCGG/1D6K_283_C_B_0_00015.pdb
463 [1.51530984] 0.6935553264393707 data/PDB/CUUCGG/1D6K_283_C_B_0_00001.pdb
497 [1.34970802] 0.6916304588175747 data/PDB/CUUCGG/1D6K_283_C_B_0_00014.pdb
600 [1.25319035] 0.557536834662705 data/PDB/CUUCGG/1TLR_10_C_A_0_00019.pdb
623 [1.18410052] 0.542687363711154 data/PDB/CUUCGG/1TLR_10_C_A_0_00018.pdb
627 [1.16534035] 0.6710242884678066 data/PDB/CUUCGG/1D6K_283_C_B_0_00017.pdb
693 [1.4252917] 0.7069600982684604 data/PDB/CUUCGG/4v64-pdb-bundle1_1028_C_A_0_00002.pdb
700 [1.2427608] 0.6600886631094386 data/PDB/CUUCGG/4v7s-pdb-bundle1_207_C_A_0_00000.pdb
719 [1.16636822] 0.6657271748311857 data/PDB/CUUCGG/1D6K_283_C_B_0_00013.pdb
726 [1.05818237] 0.6513678823301239 data/PDB/CUUCGG/1TLR_10_C_A_0_00008.pdb
727 [1.13511031] 0.7345051307434021 data/PDB/CUUCGG/1P6V_26_C_D_0_00001.pdb
745 [1.02731764] 0.6519273924694607 data/PDB/CUUCGG/1TLR_10_C_A_0_00009.pdb
835 [1.11596081] 0.6402134347912577 data/PDB/CUUCGG/1F6Z_12_C_A_0_00000.pdb
838 [1.40806741] 0.6761799137757414 data/PDB/CUUCGG/1D6K_283_C_B_0_00004.pdb
966 [1.20496552] 0.6608572508622078 data/PDB/CUUCGG/1TLR_10_C_A_0_00007.pdb
970 [1.14544239] 0.6542943726425109 data/PDB/CUUCGG/1TLR_10_C_A_0_00013.pdb
991 [1.23535274] 0.6623001009729395 data/PDB/CUUCGG/1TLR_10_C_A_0_00012.pdb
995 [1.30201879] 0.6496680926762899 data/PDB/CUUCGG/1TLR_10_C_A_0_00006.pdb
1025 [1.04994625] 0.6612420482167105 data/PDB/CUUCGG/2KU0_19_C_A_0_00004.pdb
1113 [1.07477894] 0.6599465477963314 data/PDB/CUUCGG/1TLR_10_C_A_0_00005.pdb
1114 [1.05541072] 0.6566257771105066 data/PDB/CUUCGG/1TLR_10_C_A_0_00011.pdb
1151 [1.03732723] 0.6338441447109273 data/PDB/CUUCGG/5obm-pdb-bundle4_230_C_f_0_00000.pdb
1181 [1.05609048] 0.6612899528601833 data/PDB/CUUCGG/2KU0_19_C_A_0_00003.pdb
1209 [1.07140185] 0.6268473466035946 data/PDB/CUUCGG/2LKR_63_C_A_0_00001.pdb
1214 [1.18944084] 0.5399586579433684 data/PDB/CUUCGG/1TLR_10_C_A_0_00015.pdb
1239 [1.10775434] 0.6457453554419958 data/PDB/CUUCGG/1TLR_10_C_A_0_00000.pdb
1240 [1.3505511] 0.6477341216763333 data/PDB/CUUCGG/1TLR_10_C_A_0_00014.pdb
1348 [1.10545963] 0.6596774760899786 data/PDB/CUUCGG/1TLR_10_C_A_0_00002.pdb
1355 [1.39327421] 0.5525485607445217 data/PDB/CUUCGG/1TLR_10_C_A_0_00016.pdb
1379 [1.45248573] 0.645827168988679 data/PDB/CUUCGG/1TLR_10_C_A_0_00017.pdb
1381 [1.24210706] 0.6380430320887976 data/PDB/CUUCGG/1TLR_10_C_A_0_00003.pdb
1382 [1.33012899] 0.6905548185126364 data/PDB/CUUCGG/1D6K_283_C_B_0_00018.pdb
1436 [1.09217244] 0.6360309999772672 data/PDB/CUUCGG/1F7I_12_C_A_0_00002.pdb
1443 [1.67669057] 0.5762059650065263 data/PDB/CUUCGG/1RAW_21_C_A_0_00002.pdb
1445 [1.32995574] 0.6152032407987532 data/PDB/CUUCGG/1AUD_28_C_B_0_00006.pdb
1485 [1.03690751] 0.6389382799574367 data/PDB/CUUCGG/1F7G_12_C_A_0_00006.pdb
1489 [1.12057316] 0.6488820186213259 data/PDB/CUUCGG/1F7G_12_C_A_0_00013.pdb
1497 [1.1608036] 0.625275019204503 data/PDB/CUUCGG/1F7G_12_C_A_0_00007.pdb
1514 [1.25238924] 0.6555211676708157 data/PDB/CUUCGG/4v7t-pdb-bundle1_207_C_A_0_00000.pdb
1526 [1.42342117] 0.7071570901615393 data/PDB/CUUCGG/4v64-pdb-bundle3_1028_C_A_0_00002.pdb
1537 [1.12750455] 0.6325829117106667 data/PDB/CUUCGG/1EBS_12_C_A_0_00000.pdb
1542 [1.14195981] 0.6289445606159191 data/PDB/CUUCGG/1RAW_21_C_A_0_00003.pdb
1543 [1.0601103] 0.6649869735158795 data/PDB/CUUCGG/2N6W_32_C_A_0_00006.pdb
1548 [1.10131288] 0.6517546514051366 data/PDB/CUUCGG/1F7I_12_C_A_0_00003.pdb
1566 [1.11348687] 0.6439498084695249 data/PDB/CUUCGG/1F7I_12_C_A_0_00001.pdb
1569 [1.53885124] 0.4209474840546908 data/PDB/CUUCGG/1RAW_21_C_A_0_00001.pdb
1571 [1.15474486] 0.6273714869949023 data/PDB/CUUCGG/1EBS_12_C_A_0_00002.pdb
1576 [1.26684488] 0.654358084064272 data/PDB/CUUCGG/4v57-pdb-bundle3_1028_C_A_0_00002.pdb
1608 [1.10076983] 0.6349432591956227 data/PDB/CUUCGG/1F7G_12_C_A_0_00011.pdb
1609 [1.1699881] 0.6530922357915064 data/PDB/CUUCGG/1F7G_12_C_A_0_00005.pdb
1612 [1.10507991] 0.6477001900531247 data/PDB/CUUCGG/1F7G_12_C_A_0_00004.pdb
1663 [1.13928165] 0.6284893201956351 data/PDB/CUUCGG/1RAW_21_C_A_0_00000.pdb
1686 [1.15921223] 0.6505547285888429 data/PDB/CUUCGG/1F7I_12_C_A_0_00010.pdb
1690 [1.04093556] 0.6552875114337487 data/PDB/CUUCGG/1F7I_12_C_A_0_00004.pdb
1698 [1.52744369] 0.43260024126171465 data/PDB/CUUCGG/1RAW_21_C_A_0_00004.pdb
1750 [1.21540521] 0.6537041270152071 data/PDB/CUUCGG/1F7G_12_C_A_0_00014.pdb
1770 [1.1062396] 0.6208757225997864 data/PDB/CUUCGG/1F7G_12_C_A_0_00015.pdb
1814 [1.43245993] 0.7082185587589195 data/PDB/CUUCGG/3AM1_32_C_B_0_00000.pdb
1822 [1.62495779] 0.4727774331782183 data/PDB/CUUCGG/1RAW_21_C_A_0_00005.pdb
1830 [1.11261766] 0.6603477331989258 data/PDB/CUUCGG/1EKZ_13_C_B_0_00034.pdb
1832 [1.0893901] 0.6465228110134265 data/PDB/CUUCGG/1F7I_12_C_A_0_00005.pdb
1858 [1.45788262] 0.5074835964310085 data/PDB/CUUCGG/1RAW_21_C_A_0_00007.pdb
1894 [1.05406111] 0.6414196817138739 data/PDB/CUUCGG/1F7H_12_C_A_0_00009.pdb
1913 [1.04582658] 0.6758394397678571 data/PDB/CUUCGG/1F7G_12_C_A_0_00002.pdb
1935 [1.04506323] 0.6209021442676592 data/PDB/CUUCGG/1DK1_32_C_B_0_00001.pdb
1954 [1.15270264] 0.5969868637667803 data/PDB/CUUCGG/1RAW_21_C_A_0_00006.pdb
1955 [1.06005206] 0.6737660078039002 data/PDB/CUUCGG/2N6W_32_C_A_0_00003.pdb
1967 [1.02651767] 0.6669772724705925 data/PDB/CUUCGG/1F7I_12_C_A_0_00006.pdb
```

In [ ]:
