## Supplementary material for "Integrating NMR and Simulations Reveals Motions in the UUCG Tetraloop": Figure 3 (from another version of this preprint: version 2, 2019-09-11)

figure3


### Plot free energy surfaces¶

In this notebook we plot the free energy surfaces onto different collective variables. First, we need to calculate different quantities from trajectories. This is a rather dull excercise, that we did in a separate notebook. Here, we just read the data from pickle files

In [1]:

```
import pickle

# ERMSD - stem only (residues 1-5,10-14)
fname = "data/observables/ermsd_stem_2koc.p"
ermsd_stem_2koc = pickle.load(open(fname,'rb'), encoding='bytes')

fname = "data/observables/ermsd_stem_6by5.p"
ermsd_stem_6by5 = pickle.load(open(fname,'rb'), encoding='bytes')

fname = "data/observables/ermsd_stem.p"
ermsd_stem = pickle.load(open(fname,'rb'), encoding='bytes')

# ERMSD - stem loop (residues 5-10)
fname = "data/observables/ermsd_loop_2koc.p"
ermsd_loop_2koc = pickle.load(open(fname,'rb'), encoding='bytes')

fname = "data/observables/ermsd_loop_6by5.p"
ermsd_loop_6by5 = pickle.load(open(fname,'rb'), encoding='bytes')

fname = "data/observables/ermsd_loop.p"
ermsd_loop = pickle.load(open(fname,'rb'), encoding='bytes')
```

We now need to read the weights of the different ensembles. To complicate things, we have divided our dataset in 4 bins, so that we can estimate errors.

In [2]:

```
import numpy as np
weights_0 = []
weights_A = []
theta=30
for j in range(4):
    dd = np.array([[float(x) for x in line.split()] for line in open("data/weights_set_A2_t_%d_rep_%d.dat" % (theta,j))])
    weights_0.append(dd[:,0])
    weights_A.append(dd[:,1])
  
weights_0 = np.array(weights_0)
weights_A = np.array(weights_A)
```

We define a handy FES class, which essentially takes the weights, makes an histogram and returns the FES with lower and upper error bounds

In [3]:

```
from scipy import stats
import scipy
class FES:
    
    def __init__(self,data,ww,bw=0.1,blocks=4):   
        
        self.data = np.copy(data)
        self.ww = np.copy(ww)
        self.blocks=blocks
        bsize = int(data.shape[0]/blocks)
        self.bins = np.linspace(1.1*np.min(data),1.05*np.max(data),200)

        hists = []
        for j in range(blocks):
            st = j*bsize
            end = st + bsize
            f = bw/self.data[st:end].std()
            kernel = stats.gaussian_kde(data[st:end],weights=ww[j],bw_method=f)
            hh = kernel(self.bins)
            hists.append(hh)
        self.hists = np.array(hists)
        
    def get_fes(self,temp=280):
        
        kbt=0.008314462*temp
        avgs = np.average(self.hists,axis=0)
        stds = np.std(self.hists,axis=0,ddof=1)/np.sqrt(self.blocks)
        fes = -kbt*np.log(avgs)
        fes_up = -kbt*np.log(avgs  + stds)
        fes_low = -kbt*np.log(avgs  - stds)
        mm = np.min(fes)

        return self.bins,fes-mm,fes_low-mm,fes_up-mm
    
    def get_populations(self,r1):
        
        assert(r1[0]<r1[1])
        int1 = np.where((self.bins>r1[0]) & (self.bins<r1[1]))
        p1 = np.sum(self.hists[:,int1[0]],axis=1)/np.sum(self.hists,axis=1)
        return p1
    
    def get_df(self,r1,r2,temp=280):
    
        kbt=0.008314462*temp
        p1 = self.get_populations(r1)
        p2 = self.get_populations(r2)
        return -kbt*(np.log(p1) - np.log(p2))
```

In [4]:

```
fes_loop0 = FES(ermsd_loop,weights_0)
bins_loop0,fes_loop0,fes_loop0_low,fes_loop0_up = fes_loop0.get_fes()

fes_loopA = FES(ermsd_loop,weights_A)
bins_loopA,fes_loopA,fes_loopA_low,fes_loopA_up = fes_loopA.get_fes()

fes_stem0 = FES(ermsd_stem,weights_0)
bins_stem0,fes_stem0,fes_stem0_low,fes_stem0_up = fes_stem0.get_fes()

fes_stemA = FES(ermsd_stem,weights_A)
bins_stemA,fes_stemA,fes_stemA_low,fes_stemA_up = fes_stemA.get_fes()
```

```
/home/sbottaro/Software/anaconda3/lib/python3.7/site-packages/ipykernel_launcher.py:30: RuntimeWarning: divide by zero encountered in log
```

In [10]:

```
import matplotlib.pyplot as plt
import seaborn as sns
cols = ["black","medium blue","pumpkin","brick red","black","white"]
cp = sns.xkcd_palette(cols)


r1 = [0,0.7]
r2 = [0.7,1.4]
r3 = [1.4,2.4]

fig,ax = plt.subplots(2,1,figsize=(3.5,7))
plt.subplots_adjust(left=0.15, bottom=0.07, right=0.95, top=0.91, wspace=0.25, hspace=0.25)


#ax[0].axvline(r1[1],c='k',ls='--',lw=0.5)
#ax[0].axvline(r2[1],c='k',ls='--',lw=0.5)
ax[1].set_ylim(-2.,18)
ax[1].set_xlim(0,2.5)
ax[1].set_ylabel("FES [kJ mol$^{-1}$]")
ax[1].set_yticks([0,4,8,12,16])
ax[1].set_xticks([0,0.5,1.0,1.5,2.0,2.5])
ax[1].text(-0.5,16.5,"b",fontsize=13,fontweight="bold")
ax[1].set_xlabel("eRMSD from native (loop)")
ax[1].plot(bins_loop0,fes_loop0,label="MD",color=cp[0],lw=2)
ax[1].fill_between(bins_loop0,fes_loop0_low,fes_loop0_up,alpha=0.3,color=cp[0])
ax[1].plot(bins_loopA,fes_loopA,label="MD + eNOE",color=cp[1],lw=2)
ax[1].fill_between(bins_loopA,fes_loopA_low,fes_loop0_up,alpha=0.3,color=cp[1])
ax[1].scatter(ermsd_loop_2koc,[-1]*len(ermsd_loop_2koc),color=cp[2],label="2KOC",marker="o",s=10)
ax[1].scatter(ermsd_loop_6by5,[-1]*len(ermsd_loop_6by5),color=cp[3],label="6BY5",marker="o",s=10)

ax[0].text(-0.5,26.5,"a",fontsize=13,fontweight="bold")
ax[0].plot(bins_stem0,fes_stem0,label="MD",color=cp[0],lw=2)
ax[0].fill_between(bins_stem0,fes_stem0_low,fes_stem0_up,alpha=0.3,color=cp[0])
ax[0].plot(bins_stemA,fes_stemA,label="MD + eNOE",color=cp[1],lw=2)
ax[0].fill_between(bins_stemA,fes_stemA_low,fes_stemA_up,alpha=0.3,color=cp[1])
ax[0].scatter(ermsd_stem_2koc,[-0.75]*len(ermsd_stem_2koc),color=cp[2],label="2KOC",marker="o",s=10)
ax[0].scatter(ermsd_stem_6by5,[-0.75]*len(ermsd_stem_6by5),color=cp[3],label="6BY5",marker="o",s=10)
ax[0].set_ylim(-2.0,30)
ax[0].set_yticks([0,5,10,15,20,25])
ax[0].set_xlim(0,2.5)
ax[0].set_ylabel("FES [kJ mol$^{-1}$]")
ax[0].set_xlabel("eRMSD from native (stem)")
ax[0].legend(ncol=2,loc="lower left",bbox_to_anchor=(0.05,1.02),fontsize=10)

plt.savefig("figures/projections_final.pdf")
plt.savefig("figures/projections_final.png",dpi=600)
plt.show()
plt.close()
```

Additionally, we can project the FES on the eRMSD from native, considering the whole system

In [6]:

```
# ERMSD 
fname = "data/observables/ermsd_full.p"
ermsd_full = pickle.load(open(fname,'rb'), encoding='bytes')

fes_full0 = FES(ermsd_full,weights_0)
bins_full0,fes_full0,fes_full0_low,fes_full0_up = fes_full0.get_fes()

fes_fullA = FES(ermsd_full,weights_A)
bins_fullA,fes_fullA,fes_fullA_low,fes_fullA_up = fes_fullA.get_fes()
fig,ax = plt.subplots(1,1,figsize=(7,7))
plt.subplots_adjust(left=0.15, bottom=0.07, right=0.95, top=0.91, wspace=0.25, hspace=0.25)
ax.set_ylim(-2.,18)
ax.set_xlim(0,2.5)
ax.set_ylabel("FES [kJ mol$^{-1}$]")
ax.set_yticks([0,4,8,12,16])
ax.set_xticks([0,0.5,1.0,1.5,2.0,2.5])

ax.set_xlabel("eRMSD from native")
ax.plot(bins_full0,fes_full0,label="MD",color=cp[0],lw=2)
ax.fill_between(bins_full0,fes_full0_low,fes_full0_up,alpha=0.3,color=cp[0])
ax.plot(bins_fullA,fes_fullA,label="MD + eNOE",color=cp[1],lw=2)
ax.fill_between(bins_fullA,fes_fullA_low,fes_fullA_up,alpha=0.3,color=cp[1])

plt.show()
plt.close()
```

And we can do the same for RMSD

In [7]:

```
# RMSD
fname = "data/observables/rmsd_full.p"
rmsd_full = pickle.load(open(fname,'rb'), encoding='bytes')

fes_full0 = FES(rmsd_full,weights_0,bw=0.025)
bins_full0,fes_full0,fes_full0_low,fes_full0_up = fes_full0.get_fes()

fes_fullA = FES(rmsd_full,weights_A,bw=0.025)
bins_fullA,fes_fullA,fes_fullA_low,fes_fullA_up = fes_fullA.get_fes()
```

```
/home/sbottaro/Software/anaconda3/lib/python3.7/site-packages/ipykernel_launcher.py:30: RuntimeWarning: divide by zero encountered in log
/home/sbottaro/Software/anaconda3/lib/python3.7/site-packages/ipykernel_launcher.py:30: RuntimeWarning: invalid value encountered in log
```

In [8]:

```
fig,ax = plt.subplots(1,1,figsize=(7,7))
plt.subplots_adjust(left=0.15, bottom=0.07, right=0.95, top=0.91, wspace=0.25, hspace=0.25)
ax.set_ylim(-2.,25)
ax.set_xlim(0,1.5)
ax.set_ylabel("FES [kJ mol$^{-1}$]")
ax.set_yticks([0,4,8,12,16,20,24])
ax.set_xticks([0,0.5,1.0,1.5])

ax.set_xlabel("rmsd from native [nm]")
ax.plot(bins_full0,fes_full0,label="MD",color=cp[0],lw=2)
ax.fill_between(bins_full0,fes_full0_low,fes_full0_up,alpha=0.3,color=cp[0])
ax.plot(bins_fullA,fes_fullA,label="MD + eNOE",color=cp[1],lw=2)
ax.fill_between(bins_fullA,fes_fullA_low,fes_fullA_up,alpha=0.3,color=cp[1])

plt.show()
plt.close()
```

Note that in the RMSD projection we observe only one minimum: this is because both state A and B are within 0.5nm from native.

In [9]:

```
a = plt.hist(rmsd_full,bins=100)
```

In [ ]:
