## Supplementary material for "Integrating NMR and Simulations Reveals Motions in the UUCG Tetraloop": Figure 2 (from another version of this preprint: version 2, 2019-09-11)

figure2


### Compare simulations with experiments¶

We compare back-calculated and experimental data. Instead of reporting the average difference, we here plot all the data.

You need to execute the following notebooks in order to run successfully the code below:

- back calculate experiment from trajectory
- refine MD simulation

In [6]:

```
import sys
# add BME to the path (should be changed and it depends where the BME script is located)
sys.path.append('/Users/sandrobottaro/Software/BME/')
import bme_reweight as bme

lista1 = [   ["set_A1","eNOE","eNOE_unidir","gn_eNOE"] ]

lista2 = [ ["set_A1","eNOE","eNOE_unidir","gn_eNOE"],\
        ["set_B","NOE","J3","RDC"],\
        ["set_C","RDC1","RDC2"],\
        ["set_D","sPRE"]]
```

In [7]:

```
for i in range(len(lista2)):
    for j in range(1,len(lista2[i])):
        exp = "data/%s/%s.exp.dat" % (lista2[i][0],lista2[i][j])
        calc = "data/%s/%s.calc.dat" % (lista2[i][0],lista2[i][j])
        
        # now for PBD
        rew= bme.Reweight()
        calc = "data/%s/2koc.%s.calc.dat" % (lista2[i][0],lista2[i][j])
        rew.load(exp,calc)
        before,after,neff = rew.optimize(theta=1000)
        chi = rew.weight_exp(exp,calc,"data/comparison/%s/2koc_%s" % (lista2[i][0],lista2[i][j]))
        
        rew= bme.Reweight()
        calc = "data/%s/6by5.%s.calc.dat" % (lista2[i][0],lista2[i][j])
        rew.load(exp,calc)
        before,after,neff = rew.optimize(theta=1000)
        chi = rew.weight_exp(exp,calc,"data/comparison/%s/6by5_%s" % (lista2[i][0],lista2[i][j]))
```

```
# Warning: expt average        C8_H4';G10_H8=2.7098e-05 is larger than maximum value in simulation 1.0950e-05
# Warning: expt average         A4_H2;U11_H3=2.1664e-03 is larger than maximum value in simulation 1.9520e-03
# Warning: expt average         A4_H2;G12_H1=1.4964e-04 is larger than maximum value in simulation 5.0528e-05
# Warning: expt average       U11_H1';U11_H3=1.0555e-04 is smaller than minimum value in simulation 1.1313e-04
# Warning: expt average        U11_H3;U11_H5=2.2328e-04 is larger than maximum value in simulation 1.8584e-04
# Warning: expt average         C5_H5;G10_H1=7.7791e-05 is larger than maximum value in simulation 6.3488e-05
# Warning: expt average         C5_H6;G10_H1=3.4589e-05 is larger than maximum value in simulation 2.9324e-05
# Warning: expt average         G2_H1;C13_H5=8.3836e-05 is larger than maximum value in simulation 7.5302e-05
# Warning: expt average        G2_H1;C14_H1'=7.9748e-05 is larger than maximum value in simulation 7.1334e-05
# Warning: expt average         C3_H41;C3_H5=4.8305e-04 is larger than maximum value in simulation 4.1155e-04
# Warning: expt average       C13_H41;C13_H5=3.4830e-04 is smaller than minimum value in simulation 3.8655e-04
# Warning: expt average         C5_H41;C5_H5=2.6328e-04 is smaller than minimum value in simulation 3.8881e-04
# Warning: expt average        C3_H41;G12_H1=3.3128e-03 is smaller than minimum value in simulation 5.1229e-03
# Warning: expt average        G2_H1;C13_H41=3.2371e-03 is smaller than minimum value in simulation 3.4327e-03
# Warning: expt average        C3_H42;G12_H1=2.1363e-04 is smaller than minimum value in simulation 2.8280e-04
# Warning: expt average        C5_H42;U11_H3=5.6166e-05 is larger than maximum value in simulation 3.1980e-05
# Warning: expt average        C3_H42;U11_H3=4.6153e-05 is larger than maximum value in simulation 1.8533e-05
# Warning: expt average         C5_H42;C5_H5=3.0917e-03 is smaller than minimum value in simulation 4.5465e-03
# Warning: expt average       C13_H42;C13_H5=3.0917e-03 is smaller than minimum value in simulation 4.4892e-03
# Warning: expt average         C3_H42;C3_H5=3.9991e-03 is smaller than minimum value in simulation 4.5956e-03
# Warning: expt average         U6_H2';U7_H6=1.1123e-04 is smaller than minimum value in simulation 1.2521e-04
# Warning: expt average         C5_H1';C5_H6=3.3742e-04 is smaller than minimum value in simulation 5.1395e-04
# Warning: expt average         C8_H4';G9_H8=3.6925e-05 is smaller than minimum value in simulation 5.0185e-05
# Warning: expt average         G9_H2';G9_H8=8.5972e-05 is smaller than minimum value in simulation 2.3119e-04
# Warning: expt average        G10_H8;U11_H5=5.9580e-05 is smaller than minimum value in simulation 1.0024e-04
# Warning: expt average        G10_H8;U11_H6=9.8920e-05 is larger than maximum value in simulation 7.9017e-05
# Warning: expt average          C5_H6;U6_H6=1.6498e-04 is larger than maximum value in simulation 1.2762e-04
# Warning: expt average         G1_H2';G2_H8=3.5527e-03 is smaller than minimum value in simulation 1.0922e-02
# Warning: expt average        G1_H1';G1_H2'=1.6812e-03 is smaller than minimum value in simulation 2.1252e-03
# Warning: expt average       U11_H1';U11_H6=3.6540e-04 is smaller than minimum value in simulation 4.3621e-04
# Warning: expt average         G2_H1';G2_H8=2.7142e-04 is smaller than minimum value in simulation 3.8617e-04
# Warning: expt average         G1_H1';G2_H8=5.5511e-05 is smaller than minimum value in simulation 8.0909e-05
# Warning: expt average         C5_H1';U6_H6=8.2792e-05 is larger than maximum value in simulation 6.6129e-05
# Warning: expt average          G2_H8;C3_H5=2.3344e-04 is larger than maximum value in simulation 1.8452e-04
# Warning: expt average       U11_H1';G12_H8=9.7653e-05 is larger than maximum value in simulation 5.8572e-05
# Warning: expt average         C8_H5";G9_H8=5.2367e-05 is smaller than minimum value in simulation 7.2053e-05
# Warning: expt average         C8_H5';G9_H8=7.5890e-05 is smaller than minimum value in simulation 2.0656e-04
# Warning: expt average         G1_H5";G2_H8=1.5994e-05 is smaller than minimum value in simulation 1.9753e-05
# theta: 1000
# Warning: expt average        U11_H3;U11_H5=2.2328e-04 is larger than maximum value in simulation 1.8558e-04
# Warning: expt average         C3_H41;C3_H5=4.8305e-04 is larger than maximum value in simulation 4.3113e-04
# Warning: expt average       C13_H41;C13_H5=3.4830e-04 is smaller than minimum value in simulation 4.3028e-04
# Warning: expt average         C5_H41;C5_H5=2.6328e-04 is smaller than minimum value in simulation 4.3014e-04
# Warning: expt average         C5_H42;C5_H5=3.0917e-03 is smaller than minimum value in simulation 5.7572e-03
# Warning: expt average       C13_H42;C13_H5=3.0917e-03 is smaller than minimum value in simulation 5.7557e-03
# Warning: expt average         C3_H42;C3_H5=3.9991e-03 is smaller than minimum value in simulation 5.7543e-03
# Warning: expt average         C5_H1';C5_H6=3.3742e-04 is smaller than minimum value in simulation 4.0880e-04
# Warning: expt average         G9_H2';G9_H8=8.5972e-05 is smaller than minimum value in simulation 1.4878e-04
# Warning: expt average        G1_H1';G1_H2'=1.6812e-03 is smaller than minimum value in simulation 2.0841e-03
# Warning: expt average       U11_H1';U11_H6=3.6540e-04 is smaller than minimum value in simulation 4.3557e-04
# Warning: expt average         G2_H1';G2_H8=2.7142e-04 is smaller than minimum value in simulation 3.2636e-04
# Warning: expt average        C8_H4';C8_H5"=3.0917e-03 is smaller than minimum value in simulation 3.7055e-03
# theta: 1000
# Warning: expt average         G2_H1;C13_H6=7.9748e-05 is larger than maximum value in simulation 2.8698e-05
# Warning: expt average         C3_H5;G12_H1=9.7653e-05 is larger than maximum value in simulation 8.9591e-05
# Warning: expt average          U6_H6;G9_H1=3.4966e-05 is larger than maximum value in simulation 2.4703e-05
# Warning: expt average        U6_HO2';C8_H5=2.7142e-04 is larger than maximum value in simulation 2.6324e-04
# Warning: expt average         G9_H1;G10_H8=5.8186e-05 is larger than maximum value in simulation 3.4944e-05
# Warning: expt average        U11_H3;U11_H6=1.7960e-04 is larger than maximum value in simulation 8.5961e-05
# Warning: expt average        C3_H1';G12_H1=1.0418e-04 is larger than maximum value in simulation 5.3480e-05
# Warning: expt average         C5_H41;C5_H6=3.1401e-05 is smaller than minimum value in simulation 4.1193e-05
# Warning: expt average       U6_HO2';U7_H5"=3.5959e-04 is larger than maximum value in simulation 6.9051e-05
# Warning: expt average         C8_H42;C8_H5=7.1185e-03 is larger than maximum value in simulation 5.8576e-03
# Warning: expt average         C8_H5";G9_H1=3.1401e-05 is larger than maximum value in simulation 1.5948e-05
# Warning: expt average       G12_H1;C13_H41=1.6498e-04 is larger than maximum value in simulation 1.6475e-04
# Warning: expt average       G12_H1;C13_H42=6.7978e-05 is larger than maximum value in simulation 6.1231e-05
# Warning: expt average       C14_H42;C14_H5=7.7073e-03 is larger than maximum value in simulation 5.0848e-03
# Warning: expt average         C3_H41;C3_H6=9.3964e-05 is larger than maximum value in simulation 4.2660e-05
# Warning: expt average         G1_H2';G1_H8=4.6713e-04 is larger than maximum value in simulation 3.1946e-04
# Warning: expt average         G1_H4';G2_H8=2.1220e-05 is smaller than minimum value in simulation 3.8759e-05
# Warning: expt average         G2_H4';G2_H8=2.8419e-04 is larger than maximum value in simulation 2.3202e-04
# Warning: expt average         A4_H1';C5_H6=4.4109e-05 is smaller than minimum value in simulation 8.1416e-05
# Warning: expt average        A4_H2;G12_H1'=3.3742e-04 is larger than maximum value in simulation 1.5268e-04
# Warning: expt average         A4_H2;C5_H3'=1.7950e-05 is larger than maximum value in simulation 8.0461e-06
# Warning: expt average         A4_H2';C5_H6=4.1957e-03 is smaller than minimum value in simulation 1.2002e-02
# Warning: expt average          A4_H8;C5_H5=1.5600e-04 is smaller than minimum value in simulation 1.6475e-04
# Warning: expt average        C5_H1';C5_H2'=1.6812e-03 is smaller than minimum value in simulation 1.9487e-03
# Warning: expt average         C5_H3';C5_H6=2.0752e-03 is larger than maximum value in simulation 1.4190e-03
# Warning: expt average         U6_H2';U7_H5=5.8878e-05 is larger than maximum value in simulation 5.0867e-05
# Warning: expt average        U6_H2';U6_H3'=3.9991e-03 is smaller than minimum value in simulation 4.8342e-03
# Warning: expt average         U7_H4';U7_H6=9.5175e-05 is smaller than minimum value in simulation 1.0534e-04
# Warning: expt average         C8_H4';C8_H6=2.7142e-04 is larger than maximum value in simulation 2.1080e-04
# Warning: expt average        G9_H1';G9_H2'=2.7601e-03 is larger than maximum value in simulation 2.0144e-03
# Warning: expt average       G10_H1';G10_H8=3.9613e-04 is larger than maximum value in simulation 3.8207e-04
# Warning: expt average       U11_H3';G12_H8=1.5485e-03 is larger than maximum value in simulation 6.9733e-04
# Warning: expt average      U11_H1';U11_H3'=1.9575e-04 is smaller than minimum value in simulation 2.6420e-04
# Warning: expt average      U11_H1';U11_H4'=8.4859e-04 is larger than maximum value in simulation 8.0735e-04
# Warning: expt average      C13_H1';C13_H2'=1.7524e-03 is smaller than minimum value in simulation 1.7741e-03
# Warning: expt average      C13_H1';C13_H3'=1.4759e-04 is smaller than minimum value in simulation 2.6646e-04
# Warning: expt average       C14_H1';C14_H6=2.7986e-04 is smaller than minimum value in simulation 4.0793e-04
# Warning: expt average      C14_H1';C14_H3'=2.2328e-04 is smaller than minimum value in simulation 2.5335e-04
# Warning: expt average        G1_H2';G2_H4'=2.4783e-04 is larger than maximum value in simulation 2.1142e-04
# Warning: expt average         G1_H5';G2_H8=8.7217e-06 is smaller than minimum value in simulation 1.1025e-05
# Warning: expt average        G1_H4';G1_H5"=2.8875e-03 is smaller than minimum value in simulation 2.9869e-03
# Warning: expt average         C5_H5';C5_H6=1.3781e-04 is smaller than minimum value in simulation 2.2907e-04
# Warning: expt average         C5_H5";C5_H6=4.8305e-04 is larger than maximum value in simulation 3.7150e-04
# Warning: expt average        U7_H2';C8_H5"=5.4865e-05 is larger than maximum value in simulation 3.6800e-05
# Warning: expt average         U7_H5';U7_H6=1.9019e-04 is smaller than minimum value in simulation 2.1762e-04
# Warning: expt average        C8_H4';G9_H5"=9.8547e-04 is smaller than minimum value in simulation 1.5479e-03
# Warning: expt average        C8_H1';C8_H5"=1.1270e-04 is larger than maximum value in simulation 8.1609e-05
# Warning: expt average        C8_H3';C8_H5"=1.4279e-03 is smaller than minimum value in simulation 3.0614e-03
# Warning: expt average        C8_H2';C8_H5"=2.4783e-04 is smaller than minimum value in simulation 2.5979e-04
# Warning: expt average       G10_H5";G10_H8=1.5173e-03 is larger than maximum value in simulation 1.2899e-03
# Warning: expt average       G12_H5";G12_H8=4.9125e-04 is larger than maximum value in simulation 3.6341e-04
# Warning: expt average      C13_H1';C13_H5'=1.2878e-04 is larger than maximum value in simulation 9.7577e-05
# Warning: expt average         G2_H1;C14_H5=1.6042e-04 is larger than maximum value in simulation 4.6006e-05
# Warning: expt average        G2_H1;C13_H1'=7.9748e-05 is larger than maximum value in simulation 5.1500e-05
# Warning: expt average        U6_H1';G10_H1=3.6540e-04 is larger than maximum value in simulation 2.1182e-04
# Warning: expt average       U11_H3;G12_H1'=6.6353e-05 is larger than maximum value in simulation 4.9542e-05
# Warning: expt average       G12_H1;C13_H1'=1.0694e-04 is larger than maximum value in simulation 6.9143e-05
# Warning: expt average       U6_HO2';U7_H2'=7.3339e-04 is smaller than minimum value in simulation 1.8353e-03
# Warning: expt average         U7_H2';G9_H1=1.3595e-04 is larger than maximum value in simulation 1.0919e-04
# Warning: expt average       U11_H2';U11_H3=1.0977e-04 is larger than maximum value in simulation 1.0531e-04
# Warning: expt average        G1_H2';G1_H3'=4.5122e-03 is smaller than minimum value in simulation 4.6250e-03
# Warning: expt average        G1_H2';G1_H4'=5.9304e-04 is larger than maximum value in simulation 3.6764e-04
# Warning: expt average        G2_H1';G2_H2'=2.4694e-03 is larger than maximum value in simulation 2.2267e-03
# Warning: expt average         G2_H2';G2_H8=6.4733e-04 is larger than maximum value in simulation 2.5896e-04
# Warning: expt average         G2_H3';G2_H8=9.3132e-04 is smaller than minimum value in simulation 9.8305e-04
# Warning: expt average        G2_H1';G2_H3'=1.9295e-04 is smaller than minimum value in simulation 2.7146e-04
# Warning: expt average        G2_H1';G2_H4'=1.2181e-03 is larger than maximum value in simulation 6.7412e-04
# Warning: expt average         A4_H1';A4_H2=1.4557e-04 is larger than maximum value in simulation 1.0649e-04
# Warning: expt average         A4_H2';C5_H5=2.3344e-04 is smaller than minimum value in simulation 3.0654e-04
# Warning: expt average        A4_H2';C5_H1'=4.1602e-04 is smaller than minimum value in simulation 7.8883e-04
# Warning: expt average         A4_H2;A4_H2'=8.8172e-05 is larger than maximum value in simulation 5.9003e-05
# Warning: expt average         A4_H3';C5_H6=1.9466e-03 is larger than maximum value in simulation 1.7701e-03
# Warning: expt average          A4_H8;C5_H6=6.8808e-05 is smaller than minimum value in simulation 8.8250e-05
# Warning: expt average        C5_H1';C5_H3'=4.2290e-04 is larger than maximum value in simulation 3.2554e-04
# Warning: expt average         C5_H3';C5_H5=1.7960e-04 is larger than maximum value in simulation 8.9138e-05
# Warning: expt average        C5_H1';C5_H4'=1.6468e-03 is larger than maximum value in simulation 7.1311e-04
# Warning: expt average         C5_H4';C5_H6=5.0814e-04 is larger than maximum value in simulation 2.0064e-04
# Warning: expt average          C5_H6;U6_H5=1.8481e-04 is smaller than minimum value in simulation 1.9590e-04
# Warning: expt average        U6_H1';U6_H2'=1.5805e-03 is smaller than minimum value in simulation 1.8795e-03
# Warning: expt average         U6_H3';U6_H6=3.1634e-03 is larger than maximum value in simulation 3.1533e-03
# Warning: expt average         U7_H1';U7_H6=3.5959e-04 is smaller than minimum value in simulation 3.7240e-04
# Warning: expt average         U7_H3';U7_H6=1.5600e-04 is smaller than minimum value in simulation 1.6031e-04
# Warning: expt average        C8_H3';C8_H4'=2.0312e-03 is smaller than minimum value in simulation 2.2910e-03
# Warning: expt average        C8_H2';C8_H4'=3.8349e-04 is larger than maximum value in simulation 3.4681e-04
# Warning: expt average        G9_H2';G9_H3'=6.0912e-03 is larger than maximum value in simulation 5.3320e-03
# Warning: expt average         G9_H3';G9_H8=5.8186e-05 is larger than maximum value in simulation 4.6395e-05
# Warning: expt average         G9_H4';G9_H8=3.3128e-05 is smaller than minimum value in simulation 3.7377e-05
# Warning: expt average       G10_H3';G10_H8=2.0312e-03 is smaller than minimum value in simulation 3.3491e-03
# Warning: expt average       G10_H3';U11_H5=8.0308e-04 is larger than maximum value in simulation 5.4120e-04
# Warning: expt average       G10_H4';G10_H8=5.0814e-04 is larger than maximum value in simulation 2.5099e-04
# Warning: expt average      U11_H1';U11_H2'=1.5173e-03 is smaller than minimum value in simulation 1.9172e-03
# Warning: expt average       U11_H2';U11_H6=1.9861e-04 is smaller than minimum value in simulation 3.5531e-04
# Warning: expt average       U11_H3';U11_H5=2.0446e-04 is larger than maximum value in simulation 8.6538e-05
# Warning: expt average       C13_H2';C13_H5=1.1572e-04 is larger than maximum value in simulation 5.2568e-05
# Warning: expt average       C13_H3';C13_H5=1.9019e-04 is larger than maximum value in simulation 7.5712e-05
# Warning: expt average         C5_H2';C5_H5=1.3411e-04 is larger than maximum value in simulation 4.1175e-05
# Warning: expt average        C13_H5;C14_H5=5.6302e-04 is larger than maximum value in simulation 2.9017e-04
# Warning: expt average       G10_H1;U11_H1'=1.0555e-04 is larger than maximum value in simulation 5.8090e-05
# Warning: expt average         C3_H42;C3_H6=1.2205e-04 is larger than maximum value in simulation 1.0384e-04
# Warning: expt average         C8_H41;C8_H5=9.3132e-04 is larger than maximum value in simulation 4.4570e-04
# Warning: expt average         C8_H41;C8_H6=1.3781e-04 is larger than maximum value in simulation 4.3822e-05
# Warning: expt average       G12_H1;G12_H22=4.4437e-04 is smaller than minimum value in simulation 8.2151e-03
# Warning: expt average       C14_H41;C14_H5=4.6713e-04 is larger than maximum value in simulation 4.0753e-04
# Warning: expt average       C14_H42;C14_H6=1.4359e-04 is larger than maximum value in simulation 1.0364e-04
# Warning: expt average       U6_H1';G10_H21=3.9991e-03 is larger than maximum value in simulation 2.1556e-03
# Warning: expt average        G1_H1';G1_H5'=1.9575e-04 is larger than maximum value in simulation 1.0883e-04
# Warning: expt average         G1_H5";G1_H8=4.0264e-04 is larger than maximum value in simulation 3.3827e-04
# Warning: expt average        G2_H1';G2_H5"=5.4228e-05 is larger than maximum value in simulation 5.2470e-05
# Warning: expt average         G2_H5";G2_H8=2.9767e-04 is larger than maximum value in simulation 2.9707e-04
# Warning: expt average        C8_H4';G9_H5'=9.1405e-04 is larger than maximum value in simulation 3.4341e-04
# Warning: expt average        C8_H1';C8_H5'=5.4865e-05 is smaller than minimum value in simulation 8.6790e-05
# Warning: expt average        C8_H2';C8_H5'=1.1420e-04 is smaller than minimum value in simulation 1.2620e-04
# Warning: expt average         C8_H5;C8_H5"=2.2311e-05 is larger than maximum value in simulation 1.6263e-05
# Warning: expt average         C8_H5";C8_H6=2.5932e-04 is larger than maximum value in simulation 2.3879e-04
# Warning: expt average        C8_H5";G9_H5'=5.2979e-05 is larger than maximum value in simulation 3.2770e-05
# Warning: expt average      C14_H1';C14_H5'=6.7160e-05 is smaller than minimum value in simulation 7.3121e-05
# Warning: expt average        G1_H2';G1_H5'=1.3781e-04 is larger than maximum value in simulation 4.5876e-05
# Warning: expt average        C5_H1';C5_H5"=1.2205e-04 is larger than maximum value in simulation 5.8131e-05
# Warning: expt average      U11_H1';U11_H5"=7.4042e-05 is larger than maximum value in simulation 5.6697e-05
# Warning: expt average      C13_H1';C13_H5"=9.1596e-05 is larger than maximum value in simulation 6.0334e-05
# theta: 1000
# Warning: expt average         G2_H1;C13_H6=7.9748e-05 is larger than maximum value in simulation 6.8599e-05
# Warning: expt average         C3_H5;G12_H1=9.7653e-05 is larger than maximum value in simulation 7.5908e-05
# Warning: expt average          U6_H6;G9_H1=3.4966e-05 is larger than maximum value in simulation 3.0278e-05
# Warning: expt average         G9_H1;G10_H8=5.8186e-05 is larger than maximum value in simulation 5.2770e-05
# Warning: expt average        U11_H3;U11_H6=1.7960e-04 is larger than maximum value in simulation 8.5314e-05
# Warning: expt average          C8_H5;G9_H1=4.7216e-05 is smaller than minimum value in simulation 4.9765e-05
# Warning: expt average         C5_H41;C5_H6=3.1401e-05 is smaller than minimum value in simulation 4.1907e-05
# Warning: expt average         C8_H42;C8_H5=7.1185e-03 is larger than maximum value in simulation 5.7733e-03
# Warning: expt average         C8_H5";G9_H1=3.1401e-05 is larger than maximum value in simulation 2.6894e-05
# Warning: expt average       G12_H1;C13_H42=6.7978e-05 is larger than maximum value in simulation 5.1608e-05
# Warning: expt average       C14_H42;C14_H5=7.7073e-03 is larger than maximum value in simulation 5.7748e-03
# Warning: expt average         C3_H41;C3_H6=9.3964e-05 is larger than maximum value in simulation 4.1964e-05
# Warning: expt average         G1_H4';G2_H8=2.1220e-05 is smaller than minimum value in simulation 2.1995e-05
# Warning: expt average         G2_H4';G2_H8=2.8419e-04 is larger than maximum value in simulation 1.7235e-04
# Warning: expt average        A4_H2;G12_H1'=3.3742e-04 is larger than maximum value in simulation 3.3228e-04
# Warning: expt average         A4_H2;C5_H3'=1.7950e-05 is larger than maximum value in simulation 1.6203e-05
# Warning: expt average        C5_H1';C5_H2'=1.6812e-03 is smaller than minimum value in simulation 1.7372e-03
# Warning: expt average         C5_H2';C5_H6=2.2999e-04 is smaller than minimum value in simulation 2.4668e-04
# Warning: expt average         U6_H2';U7_H5=5.8878e-05 is larger than maximum value in simulation 3.5614e-05
# Warning: expt average        U6_H2';U6_H3'=3.9991e-03 is smaller than minimum value in simulation 5.6860e-03
# Warning: expt average        C8_H4';G9_H3'=3.4830e-04 is smaller than minimum value in simulation 4.6783e-04
# Warning: expt average         C8_H4';C8_H6=2.7142e-04 is larger than maximum value in simulation 1.7474e-04
# Warning: expt average         G9_H1';G9_H8=4.8569e-03 is larger than maximum value in simulation 3.9620e-03
# Warning: expt average        G9_H1';G9_H2'=2.7601e-03 is larger than maximum value in simulation 1.6959e-03
# Warning: expt average      U11_H1';U11_H3'=1.9575e-04 is smaller than minimum value in simulation 3.2431e-04
# Warning: expt average      U11_H1';U11_H4'=8.4859e-04 is larger than maximum value in simulation 7.8639e-04
# Warning: expt average      C13_H1';C13_H2'=1.7524e-03 is smaller than minimum value in simulation 1.8894e-03
# Warning: expt average      C13_H1';C13_H3'=1.4759e-04 is smaller than minimum value in simulation 3.1652e-04
# Warning: expt average       C14_H1';C14_H6=2.7986e-04 is smaller than minimum value in simulation 3.9820e-04
# Warning: expt average      C14_H1';C14_H3'=2.2328e-04 is smaller than minimum value in simulation 3.2436e-04
# Warning: expt average        G1_H4';G1_H5"=2.8875e-03 is smaller than minimum value in simulation 4.3317e-03
# Warning: expt average         C5_H5';C5_H6=1.3781e-04 is smaller than minimum value in simulation 1.4933e-04
# Warning: expt average         U7_H5';U7_H6=1.9019e-04 is smaller than minimum value in simulation 1.9698e-04
# Warning: expt average        C8_H1';C8_H5"=1.1270e-04 is larger than maximum value in simulation 7.5441e-05
# Warning: expt average        C8_H3';C8_H5"=1.4279e-03 is smaller than minimum value in simulation 2.2398e-03
# Warning: expt average      C13_H1';C13_H5'=1.2878e-04 is larger than maximum value in simulation 9.7628e-05
# Warning: expt average        G2_H1;C13_H1'=7.9748e-05 is larger than maximum value in simulation 6.4215e-05
# Warning: expt average       U11_H3;G12_H1'=6.6353e-05 is larger than maximum value in simulation 6.2582e-05
# Warning: expt average       U6_HO2';U7_H2'=7.3339e-04 is smaller than minimum value in simulation 9.0319e-04
# Warning: expt average        G1_H2';G1_H3'=4.5122e-03 is smaller than minimum value in simulation 5.6702e-03
# Warning: expt average         G1_H3';G2_H8=9.6702e-04 is larger than maximum value in simulation 9.3904e-04
# Warning: expt average        G1_H1';G1_H3'=2.7142e-04 is smaller than minimum value in simulation 3.2223e-04
# Warning: expt average        G1_H2';G1_H4'=5.9304e-04 is larger than maximum value in simulation 3.2314e-04
# Warning: expt average        G1_H1';G1_H4'=6.8269e-04 is larger than maximum value in simulation 6.1313e-04
# Warning: expt average        G2_H1';G2_H2'=2.4694e-03 is larger than maximum value in simulation 2.1800e-03
# Warning: expt average        G2_H1';G2_H3'=1.9295e-04 is smaller than minimum value in simulation 3.2289e-04
# Warning: expt average        G2_H1';G2_H4'=1.2181e-03 is larger than maximum value in simulation 8.4828e-04
# Warning: expt average         A4_H1';A4_H2=1.4557e-04 is larger than maximum value in simulation 1.1148e-04
# Warning: expt average        A4_H2';C5_H1'=4.1602e-04 is larger than maximum value in simulation 3.2848e-04
# Warning: expt average         A4_H3';C5_H6=1.9466e-03 is larger than maximum value in simulation 1.7701e-03
# Warning: expt average        C5_H1';C5_H3'=4.2290e-04 is larger than maximum value in simulation 3.3160e-04
# Warning: expt average         C5_H3';C5_H5=1.7960e-04 is larger than maximum value in simulation 1.1494e-04
# Warning: expt average        C5_H1';C5_H4'=1.6468e-03 is larger than maximum value in simulation 1.3179e-03
# Warning: expt average         C5_H4';C5_H6=5.0814e-04 is larger than maximum value in simulation 2.9809e-04
# Warning: expt average        U6_H1';U6_H2'=1.5805e-03 is smaller than minimum value in simulation 2.0144e-03
# Warning: expt average         U6_H2';C8_H6=3.3212e-04 is larger than maximum value in simulation 2.4410e-04
# Warning: expt average         U7_H1';U7_H6=3.5959e-04 is smaller than minimum value in simulation 4.0028e-04
# Warning: expt average        C8_H3';C8_H4'=2.0312e-03 is smaller than minimum value in simulation 2.2559e-03
# Warning: expt average        C8_H2';C8_H4'=3.8349e-04 is larger than maximum value in simulation 3.1931e-04
# Warning: expt average         G9_H3';G9_H8=5.8186e-05 is larger than maximum value in simulation 3.9877e-05
# Warning: expt average         G9_H4';G9_H8=3.3128e-05 is smaller than minimum value in simulation 4.1609e-05
# Warning: expt average       G10_H2';U11_H5=3.6540e-04 is larger than maximum value in simulation 3.4847e-04
# Warning: expt average       G10_H3';U11_H5=8.0308e-04 is larger than maximum value in simulation 7.7024e-04
# Warning: expt average       G10_H4';G10_H8=5.0814e-04 is larger than maximum value in simulation 2.8811e-04
# Warning: expt average       U11_H2';G12_H8=2.8875e-03 is smaller than minimum value in simulation 4.2150e-03
# Warning: expt average      U11_H1';U11_H2'=1.5173e-03 is smaller than minimum value in simulation 2.0360e-03
# Warning: expt average       U11_H2';U11_H6=1.9861e-04 is smaller than minimum value in simulation 2.0107e-04
# Warning: expt average       U11_H3';U11_H5=2.0446e-04 is larger than maximum value in simulation 1.4950e-04
# Warning: expt average      G12_H2';C13_H1'=3.8975e-04 is larger than maximum value in simulation 3.3476e-04
# Warning: expt average       C13_H1';C13_H6=4.8305e-04 is larger than maximum value in simulation 4.4860e-04
# Warning: expt average       C13_H2';C13_H5=1.1572e-04 is larger than maximum value in simulation 7.1912e-05
# Warning: expt average       C13_H3';C13_H5=1.9019e-04 is larger than maximum value in simulation 1.5321e-04
# Warning: expt average         C5_H2';C5_H5=1.3411e-04 is larger than maximum value in simulation 6.8883e-05
# Warning: expt average       G10_H1;U11_H1'=1.0555e-04 is larger than maximum value in simulation 8.9819e-05
# Warning: expt average         G2_H1;C3_H42=2.0151e-04 is larger than maximum value in simulation 1.8415e-04
# Warning: expt average         C3_H42;C3_H6=1.2205e-04 is larger than maximum value in simulation 1.0421e-04
# Warning: expt average         C8_H41;C8_H5=9.3132e-04 is larger than maximum value in simulation 4.3106e-04
# Warning: expt average         C8_H41;C8_H6=1.3781e-04 is larger than maximum value in simulation 4.1964e-05
# Warning: expt average       G12_H1;G12_H22=4.4437e-04 is smaller than minimum value in simulation 6.7101e-04
# Warning: expt average       C14_H41;C14_H5=4.6713e-04 is larger than maximum value in simulation 4.3099e-04
# Warning: expt average       C14_H42;C14_H6=1.4359e-04 is larger than maximum value in simulation 1.0420e-04
# Warning: expt average        G1_H2';G2_H5"=1.2043e-04 is smaller than minimum value in simulation 1.3225e-04
# Warning: expt average        G1_H1';G1_H5'=1.9575e-04 is larger than maximum value in simulation 1.4894e-04
# Warning: expt average         G1_H5";G1_H8=4.0264e-04 is larger than maximum value in simulation 3.3281e-04
# Warning: expt average        C8_H4';G9_H5'=9.1405e-04 is larger than maximum value in simulation 7.0992e-04
# Warning: expt average        C8_H1';C8_H5'=5.4865e-05 is smaller than minimum value in simulation 8.5572e-05
# Warning: expt average        C8_H2';C8_H5'=1.1420e-04 is smaller than minimum value in simulation 1.4622e-04
# Warning: expt average         C8_H5;C8_H5"=2.2311e-05 is larger than maximum value in simulation 1.5437e-05
# Warning: expt average         C8_H5";C8_H6=2.5932e-04 is larger than maximum value in simulation 2.4203e-04
# Warning: expt average        G1_H2';G1_H5'=1.3781e-04 is larger than maximum value in simulation 1.0241e-04
# Warning: expt average        C5_H1';C5_H5"=1.2205e-04 is larger than maximum value in simulation 7.7464e-05
# Warning: expt average      U11_H1';U11_H5"=7.4042e-05 is larger than maximum value in simulation 6.9059e-05
# theta: 1000
# Warning: expt average        C8_H5;G10_H1'=2.6268e-05 is larger than maximum value in simulation 2.9217e-06
# Warning: expt average         C8_H5;G10_H8=2.4444e-05 is larger than maximum value in simulation 9.8395e-07
# Warning: expt average        C8_H2';G10_H8=8.0755e-06 is larger than maximum value in simulation 1.4181e-06
# Warning: expt average         C8_H3';G9_H8=2.6818e-05 is larger than maximum value in simulation 1.1819e-05
# Warning: expt average         C8_H2';G9_H8=2.3231e-05 is larger than maximum value in simulation 5.6588e-06
# Warning: expt average       C8_H1';G10_H4'=4.6153e-05 is larger than maximum value in simulation 2.3876e-06
# Warning: expt average        C8_H2';G9_H2'=1.0284e-04 is larger than maximum value in simulation 2.9094e-06
# Warning: expt average      G10_H21;U11_H1'=7.3679e-05 is smaller than minimum value in simulation 1.5089e-04
# Warning: expt average         G1_H4';G1_H8=6.9819e-05 is smaller than minimum value in simulation 1.2467e-04
# Warning: expt average        G1_H3';G1_H5'=2.1300e-04 is smaller than minimum value in simulation 3.3545e-04
# Warning: expt average         G2_H2';C3_H6=2.2574e-03 is smaller than minimum value in simulation 3.9188e-03
# Warning: expt average        G2_H3';G2_H5"=4.7026e-04 is smaller than minimum value in simulation 6.9460e-04
# Warning: expt average         C3_H1';C3_H6=2.2001e-04 is smaller than minimum value in simulation 4.5430e-04
# Warning: expt average         C3_H1';A4_H8=2.4620e-05 is smaller than minimum value in simulation 3.9125e-05
# Warning: expt average        C3_H1';C3_H2'=8.5647e-04 is smaller than minimum value in simulation 1.8922e-03
# Warning: expt average         C3_H2';A4_H8=2.0054e-03 is smaller than minimum value in simulation 2.8993e-03
# Warning: expt boundary         C3_H2';C3_H6=1.7857e-03 is larger than maximum value in simulation 3.5816e-04
# Warning: expt average         C3_H2';C3_H6=1.7857e-03 is larger than maximum value in simulation 4.4822e-04
# Warning: expt average        C3_H1';C3_H4'=2.8683e-04 is smaller than minimum value in simulation 7.4164e-04
# Warning: expt average         C3_H4';A4_H8=1.4525e-05 is smaller than minimum value in simulation 2.2458e-05
# Warning: expt boundary          C3_H6;A4_H8=8.9296e-05 is larger than maximum value in simulation 2.2157e-05
# Warning: expt average          C3_H6;A4_H8=8.9296e-05 is larger than maximum value in simulation 3.3422e-05
# Warning: expt average         A4_H1';A4_H8=1.1375e-04 is smaller than minimum value in simulation 3.2355e-04
# Warning: expt average        A4_H1';A4_H2'=6.7424e-04 is smaller than minimum value in simulation 2.2369e-03
# Warning: expt boundary         A4_H2';A4_H8=4.4511e-04 is larger than maximum value in simulation 2.9048e-04
# Warning: expt average         A4_H2';A4_H8=4.4511e-04 is larger than maximum value in simulation 4.0893e-04
# Warning: expt average        A4_H1';A4_H3'=1.5557e-04 is smaller than minimum value in simulation 2.8794e-04
# Warning: expt average         A4_H3';A4_H8=5.8893e-04 is smaller than minimum value in simulation 1.8998e-03
# Warning: expt boundary         A4_H4';A4_H8=2.2361e-04 is larger than maximum value in simulation 1.5661e-04
# Warning: expt average         A4_H4';A4_H8=2.2361e-04 is larger than maximum value in simulation 1.7299e-04
# Warning: expt average        A4_H1';A4_H4'=2.1300e-04 is smaller than minimum value in simulation 5.6839e-04
# Warning: expt boundary         A4_H5';A4_H8=3.7192e-04 is larger than maximum value in simulation 1.8483e-04
# Warning: expt average         A4_H5';A4_H8=3.7192e-04 is larger than maximum value in simulation 2.5904e-04
# Warning: expt average         A4_H5";A4_H8=2.3872e-04 is smaller than minimum value in simulation 2.7679e-04
# Warning: expt average        C5_H2';C5_H3'=2.6749e-03 is smaller than minimum value in simulation 4.8164e-03
# Warning: expt average         C5_H3';U6_H5=2.0626e-04 is smaller than minimum value in simulation 4.9042e-04
# Warning: expt boundary        C5_H3';C5_H4'=2.5470e-03 is larger than maximum value in simulation 1.2979e-03
# Warning: expt average        C5_H3';C5_H4'=2.5470e-03 is larger than maximum value in simulation 1.3695e-03
# Warning: expt average         U6_H3';C8_H5=8.3312e-05 is smaller than minimum value in simulation 2.4473e-04
# Warning: expt boundary         U6_H3';U6_H5=1.2419e-04 is larger than maximum value in simulation 6.0162e-05
# Warning: expt average         U6_H3';U6_H5=1.2419e-04 is larger than maximum value in simulation 1.1939e-04
# Warning: expt average         U6_H4';U7_H5=3.4664e-04 is smaller than minimum value in simulation 6.4210e-04
# Warning: expt average        U6_H1';U6_H4'=5.1596e-04 is smaller than minimum value in simulation 6.8561e-04
# Warning: expt average        U6_H1';U6_H5"=4.7922e-05 is smaller than minimum value in simulation 5.0979e-05
# Warning: expt average        U7_H1';U7_H2'=3.7857e-04 is smaller than minimum value in simulation 1.1639e-03
# Warning: expt average         U7_H2';U7_H5=4.0782e-05 is smaller than minimum value in simulation 1.1572e-04
# Warning: expt average         U7_H2';C8_H6=1.1541e-04 is smaller than minimum value in simulation 1.4210e-04
# Warning: expt average        U7_H1';U7_H4'=3.4064e-04 is smaller than minimum value in simulation 8.3309e-04
# Warning: expt boundary         U7_H4';C8_H6=1.7606e-04 is larger than maximum value in simulation 9.0311e-05
# Warning: expt average         U7_H4';C8_H6=1.7606e-04 is larger than maximum value in simulation 1.4258e-04
# Warning: expt boundary        U7_H3';U7_H4'=4.2364e-03 is larger than maximum value in simulation 2.1367e-03
# Warning: expt average        U7_H3';U7_H4'=4.2364e-03 is larger than maximum value in simulation 2.4391e-03
# Warning: expt average         U7_H5';C8_H5=6.8892e-05 is smaller than minimum value in simulation 2.8397e-04
# Warning: expt average         U7_H5';C8_H6=2.2266e-05 is smaller than minimum value in simulation 4.4124e-05
# Warning: expt average         U7_H5";C8_H5=1.6673e-03 is smaller than minimum value in simulation 3.5711e-03
# Warning: expt boundary         U7_H5";C8_H6=1.8273e-03 is larger than maximum value in simulation 3.6470e-04
# Warning: expt average         U7_H5";C8_H6=1.8273e-03 is larger than maximum value in simulation 1.4476e-03
# Warning: expt boundary        U7_H4';U7_H5"=1.1626e-02 is larger than maximum value in simulation 4.0881e-03
# Warning: expt average        U7_H4';U7_H5"=1.1626e-02 is larger than maximum value in simulation 5.4870e-03
# Warning: expt boundary        U7_H2';U7_H5"=6.0031e-04 is larger than maximum value in simulation 2.0139e-04
# Warning: expt average        U7_H2';U7_H5"=6.0031e-04 is larger than maximum value in simulation 3.1185e-04
# Warning: expt average         C8_H1';C8_H6=2.0298e-04 is smaller than minimum value in simulation 3.9600e-04
# Warning: expt average        C8_H1';C8_H2'=4.5332e-04 is smaller than minimum value in simulation 1.1413e-03
# Warning: expt average         C8_H2';C8_H5=8.8060e-05 is smaller than minimum value in simulation 1.1257e-04
# Warning: expt average        C8_H1';C8_H3'=1.1050e-04 is smaller than minimum value in simulation 2.9142e-04
# Warning: expt boundary        G9_H1';G9_H3'=3.5277e-04 is larger than maximum value in simulation 2.6973e-04
# Warning: expt average        G9_H1';G9_H3'=3.5277e-04 is larger than maximum value in simulation 3.0654e-04
# Warning: expt average        G9_H1';G9_H4'=4.2922e-04 is smaller than minimum value in simulation 1.0001e-03
# Warning: expt boundary        G9_H3';G9_H4'=1.8700e-03 is larger than maximum value in simulation 1.2894e-03
# Warning: expt average        G9_H3';G9_H4'=1.8700e-03 is larger than maximum value in simulation 1.3401e-03
# Warning: expt average        G9_H3';G9_H5"=7.2944e-04 is smaller than minimum value in simulation 3.1424e-03
# Warning: expt average        G9_H1';G9_H5"=7.8860e-05 is smaller than minimum value in simulation 8.3051e-05
# Warning: expt boundary      G10_H3';G10_H5"=6.0446e-03 is larger than maximum value in simulation 4.2827e-03
# Warning: expt average      G10_H3';G10_H5"=6.0446e-03 is larger than maximum value in simulation 5.0108e-03
# Warning: expt average       G12_H1';C13_H6=2.8883e-05 is smaller than minimum value in simulation 3.2634e-05
# Warning: expt average      G12_H1';G12_H2'=9.6885e-04 is smaller than minimum value in simulation 2.0209e-03
# Warning: expt average       G12_H3';C13_H6=2.4268e-04 is smaller than minimum value in simulation 3.9191e-04
# Warning: expt average      G12_H1';G12_H3'=9.5788e-05 is smaller than minimum value in simulation 2.7768e-04
# Warning: expt average      G12_H1';G12_H5'=7.0762e-05 is smaller than minimum value in simulation 7.2239e-05
# Warning: expt average      G12_H1';G12_H5"=3.3631e-05 is smaller than minimum value in simulation 5.2678e-05
# Warning: expt boundary       C13_H5';C13_H6=3.1781e-04 is larger than maximum value in simulation 1.5751e-04
# Warning: expt average       C13_H5';C13_H6=3.1781e-04 is larger than maximum value in simulation 2.4421e-04
# Warning: expt boundary       C13_H5";C13_H6=2.9174e-04 is larger than maximum value in simulation 1.7296e-04
# Warning: expt average       C13_H5";C13_H6=2.9174e-04 is larger than maximum value in simulation 2.8896e-04
# Warning: expt boundary       C14_H2';C14_H6=1.1224e-03 is larger than maximum value in simulation 5.5858e-04
# Warning: expt average       C14_H2';C14_H6=1.1224e-03 is larger than maximum value in simulation 1.0542e-03
# Warning: expt boundary      C14_H2';C14_H3'=6.7728e-03 is larger than maximum value in simulation 5.5975e-03
# Warning: expt average      C14_H2';C14_H3'=6.7728e-03 is larger than maximum value in simulation 5.9480e-03
# theta: 1000
# Constrained optimization: Fall back to L-BFGS_B
# Warning: expt average        C8_H5;G10_H1'=2.6268e-05 is larger than maximum value in simulation 3.0470e-06
# Warning: expt average         C8_H5;G10_H8=2.4444e-05 is larger than maximum value in simulation 2.1545e-06
# Warning: expt average        C8_H2';G10_H8=8.0755e-06 is larger than maximum value in simulation 5.7250e-06
# Warning: expt average         C8_H3';G9_H8=2.6818e-05 is larger than maximum value in simulation 8.0845e-06
# Warning: expt average         C8_H2';G9_H8=2.3231e-05 is larger than maximum value in simulation 3.2440e-06
# Warning: expt average       C8_H1';G10_H4'=4.6153e-05 is larger than maximum value in simulation 4.6880e-06
# Warning: expt average        C8_H2';G9_H2'=1.0284e-04 is larger than maximum value in simulation 3.5052e-06
# Warning: expt average         G1_H3';G1_H8=5.6692e-04 is smaller than minimum value in simulation 1.8508e-03
# Warning: expt average         G1_H4';G1_H8=6.9819e-05 is smaller than minimum value in simulation 1.3809e-04
# Warning: expt average        G1_H3';G1_H5'=2.1300e-04 is smaller than minimum value in simulation 3.8090e-04
# Warning: expt average        G2_H3';G2_H5"=4.7026e-04 is smaller than minimum value in simulation 9.6229e-04
# Warning: expt average         C3_H1';C3_H6=2.2001e-04 is smaller than minimum value in simulation 3.7924e-04
# Warning: expt average        C3_H1';C3_H2'=8.5647e-04 is smaller than minimum value in simulation 1.2190e-03
# Warning: expt average        C3_H1';C3_H4'=2.8683e-04 is smaller than minimum value in simulation 4.5121e-04
# Warning: expt average         A4_H1';A4_H8=1.1375e-04 is smaller than minimum value in simulation 3.1165e-04
# Warning: expt average        A4_H1';A4_H2'=6.7424e-04 is smaller than minimum value in simulation 2.3278e-03
# Warning: expt average        A4_H1';A4_H3'=1.5557e-04 is smaller than minimum value in simulation 2.8927e-04
# Warning: expt average        A4_H1';A4_H4'=2.1300e-04 is smaller than minimum value in simulation 3.0913e-04
# Warning: expt average         A4_H5";A4_H8=2.3872e-04 is smaller than minimum value in simulation 6.8475e-03
# Warning: expt average        C5_H2';C5_H3'=2.6749e-03 is smaller than minimum value in simulation 5.7836e-03
# Warning: expt boundary        C5_H3';C5_H4'=2.5470e-03 is larger than maximum value in simulation 1.2882e-03
# Warning: expt average        C5_H3';C5_H4'=2.5470e-03 is larger than maximum value in simulation 1.3781e-03
# Warning: expt average         U6_H3';C8_H5=8.3312e-05 is smaller than minimum value in simulation 1.3129e-04
# Warning: expt average        U7_H1';U7_H2'=3.7857e-04 is smaller than minimum value in simulation 1.2212e-03
# Warning: expt average         U7_H2';U7_H5=4.0782e-05 is smaller than minimum value in simulation 4.4359e-05
# Warning: expt average         U7_H2';C8_H6=1.1541e-04 is smaller than minimum value in simulation 1.9013e-04
# Warning: expt average        U7_H1';U7_H4'=3.4064e-04 is smaller than minimum value in simulation 3.8837e-04
# Warning: expt boundary        U7_H3';U7_H4'=4.2364e-03 is larger than maximum value in simulation 2.7050e-03
# Warning: expt average        U7_H3';U7_H4'=4.2364e-03 is larger than maximum value in simulation 3.0691e-03
# Warning: expt average         U7_H5';C8_H5=6.8892e-05 is smaller than minimum value in simulation 7.4463e-05
# Warning: expt average         U7_H5';C8_H6=2.2266e-05 is smaller than minimum value in simulation 2.5920e-05
# Warning: expt boundary        U7_H4';U7_H5"=1.1626e-02 is larger than maximum value in simulation 1.7620e-03
# Warning: expt average        U7_H4';U7_H5"=1.1626e-02 is larger than maximum value in simulation 7.5590e-03
# Warning: expt average         C8_H1';C8_H6=2.0298e-04 is smaller than minimum value in simulation 3.8455e-04
# Warning: expt average        C8_H1';C8_H2'=4.5332e-04 is smaller than minimum value in simulation 1.2034e-03
# Warning: expt average        C8_H1';C8_H3'=1.1050e-04 is smaller than minimum value in simulation 3.0784e-04
# Warning: expt boundary        G9_H1';G9_H3'=3.5277e-04 is larger than maximum value in simulation 3.3197e-04
# Warning: expt average        G9_H1';G9_H3'=3.5277e-04 is larger than maximum value in simulation 3.3784e-04
# Warning: expt boundary       G9_H3';G10_H3'=3.1704e-05 is larger than maximum value in simulation 2.4670e-05
# Warning: expt average       G9_H3';G10_H3'=3.1704e-05 is larger than maximum value in simulation 3.1201e-05
# Warning: expt average        G9_H1';G9_H4'=4.2922e-04 is smaller than minimum value in simulation 1.3210e-03
# Warning: expt boundary        G9_H3';G9_H4'=1.8700e-03 is larger than maximum value in simulation 1.2905e-03
# Warning: expt average        G9_H3';G9_H4'=1.8700e-03 is larger than maximum value in simulation 1.3155e-03
# Warning: expt average      G12_H1';G12_H2'=9.6885e-04 is smaller than minimum value in simulation 1.9107e-03
# Warning: expt average       G12_H3';C13_H6=2.4268e-04 is smaller than minimum value in simulation 3.5164e-04
# Warning: expt average      G12_H1';G12_H3'=9.5788e-05 is smaller than minimum value in simulation 3.1891e-04
# Warning: expt average      G12_H1';G12_H5"=3.3631e-05 is smaller than minimum value in simulation 5.0973e-05
# Warning: expt average       C13_H3';C13_H6=1.4250e-03 is smaller than minimum value in simulation 3.5669e-03
# Warning: expt boundary      C14_H2';C14_H3'=6.7728e-03 is larger than maximum value in simulation 5.6602e-03
# Warning: expt average      C14_H2';C14_H3'=6.7728e-03 is larger than maximum value in simulation 5.8427e-03
# theta: 1000
# Constrained optimization: Fall back to L-BFGS_B
# Warning: expt average        G1_H1;C14_H41=8.3836e-05 is smaller than minimum value in simulation 1.9703e-03
# Warning: expt average        G1_H4';G1_H1'=1.1945e-03 is larger than maximum value in simulation 7.2525e-04
# Warning: expt average         G1_H1';G1_H8=6.0346e-04 is larger than maximum value in simulation 4.2110e-04
# Warning: expt average         G1_H8;G1_H2'=6.8269e-04 is larger than maximum value in simulation 3.1946e-04
# Warning: expt average        G1_H8;G1_H5''=1.0236e-03 is larger than maximum value in simulation 3.3827e-04
# Warning: expt average        G2_H1;C13_H41=2.2661e-04 is smaller than minimum value in simulation 3.4327e-03
# Warning: expt average        C14_H42;G2_H1=3.1190e-04 is larger than maximum value in simulation 1.9884e-04
# Warning: expt average         G2_H1;C14_H5=2.6731e-04 is larger than maximum value in simulation 4.6006e-05
# Warning: expt average         G2_H1;C3_H42=4.1957e-03 is larger than maximum value in simulation 4.5634e-04
# Warning: expt average        G2_H1';G2_H2'=6.7551e-03 is larger than maximum value in simulation 2.2267e-03
# Warning: expt average         G1_H1';G2_H8=5.2571e-04 is larger than maximum value in simulation 1.0296e-04
# Warning: expt average         G1_H3';G2_H8=5.2328e-03 is larger than maximum value in simulation 1.6202e-03
# Warning: expt average         G2_H1';G2_H8=6.1409e-04 is larger than maximum value in simulation 4.2689e-04
# Warning: expt average         G2_H8;G2_H3'=4.4037e-03 is larger than maximum value in simulation 1.8466e-03
# Warning: expt average         G2_H8;G2_H5'=1.1332e-02 is larger than maximum value in simulation 1.8962e-03
# Warning: expt average       G2_H2';G2_HO2'=8.5831e-03 is larger than maximum value in simulation 7.8549e-03
# Warning: expt average        C3_H1';C3_H4'=1.9059e-03 is larger than maximum value in simulation 9.6012e-04
# Warning: expt average         G2_H1;C3_H41=1.4759e-04 is smaller than minimum value in simulation 7.7136e-04
# Warning: expt average         C3_H41;C3_H5=6.4132e-03 is larger than maximum value in simulation 4.1155e-04
# Warning: expt average        C3_H41;A4_H61=2.2328e-04 is larger than maximum value in simulation 1.2442e-04
# Warning: expt average         C3_H42;C3_H5=4.9125e-04 is smaller than minimum value in simulation 4.5956e-03
# Warning: expt average        C3_H42;A4_H61=4.4437e-04 is larger than maximum value in simulation 6.5550e-05
# Warning: expt average        C3_H42;A4_H62=3.7132e-04 is larger than maximum value in simulation 8.5529e-05
# Warning: expt average         G2_H1';C3_H6=9.6405e-05 is larger than maximum value in simulation 6.9404e-05
# Warning: expt average         C3_H1';C3_H6=6.9497e-04 is larger than maximum value in simulation 4.7509e-04
# Warning: expt average          C3_H5;C3_H6=5.6430e-03 is larger than maximum value in simulation 4.9491e-03
# Warning: expt average       C3_HO2';C3_H3'=9.4898e-04 is larger than maximum value in simulation 7.8984e-04
# Warning: expt average        A4_H1';C5_H1'=1.2181e-03 is larger than maximum value in simulation 1.4597e-04
# Warning: expt average        A4_H2;G12_H1'=8.1793e-04 is larger than maximum value in simulation 1.5268e-04
# Warning: expt average         A4_H1';A4_H2=1.3411e-04 is larger than maximum value in simulation 1.0649e-04
# Warning: expt average         C5_H1';A4_H2=1.1268e-03 is larger than maximum value in simulation 6.0388e-04
# Warning: expt average        A4_H1';A4_H3'=2.1202e-03 is larger than maximum value in simulation 3.1054e-04
# Warning: expt average        A4_H61;A4_H62=6.4132e-03 is smaller than minimum value in simulation 3.4307e-02
# Warning: expt average        C3_HO2';A4_H8=9.5175e-05 is smaller than minimum value in simulation 1.0970e-04
# Warning: expt average         A4_H1';A4_H8=7.4674e-04 is larger than maximum value in simulation 3.4259e-04
# Warning: expt average         A4_H3';A4_H8=4.9786e-03 is larger than maximum value in simulation 2.7970e-03
# Warning: expt average        A4_H8;A4_H5''=1.4570e-03 is larger than maximum value in simulation 4.8256e-04
# Warning: expt average          A4_H8;C5_H6=7.4042e-05 is smaller than minimum value in simulation 8.8250e-05
# Warning: expt average       A4_H1';A4_HO2'=1.9884e-03 is larger than maximum value in simulation 8.2486e-04
# Warning: expt average       A4_H3';A4_HO2'=3.5527e-03 is larger than maximum value in simulation 7.8453e-04
# Warning: expt average        C5_H1';C5_H2'=1.5485e-03 is smaller than minimum value in simulation 1.9487e-03
# Warning: expt average        C5_H1';C5_H3'=3.4830e-04 is larger than maximum value in simulation 3.2554e-04
# Warning: expt average        C5_H1';C5_H4'=7.7431e-04 is larger than maximum value in simulation 7.1311e-04
# Warning: expt average         C5_H5;C5_H41=5.2328e-03 is larger than maximum value in simulation 4.0494e-04
# Warning: expt average         U6_H5;C5_H41=3.5959e-04 is larger than maximum value in simulation 8.8284e-05
# Warning: expt average        A4_H61;C5_H42=3.7735e-04 is larger than maximum value in simulation 1.5383e-04
# Warning: expt average        C5_H41;C5_H42=2.9401e-02 is smaller than minimum value in simulation 3.4318e-02
# Warning: expt average         C5_H5;C5_H42=5.3476e-04 is smaller than minimum value in simulation 4.5465e-03
# Warning: expt average         A4_H1';C5_H6=6.4773e-05 is smaller than minimum value in simulation 8.1416e-05
# Warning: expt average         A4_H3';C5_H6=8.1315e-03 is larger than maximum value in simulation 1.7701e-03
# Warning: expt average        C5_H6;A4_HO2'=6.9650e-05 is smaller than minimum value in simulation 1.8651e-04
# Warning: expt average         C5_H6;C5_H2'=7.2031e-04 is larger than maximum value in simulation 2.6448e-04
# Warning: expt average         C5_H6;C5_H3'=5.9374e-03 is larger than maximum value in simulation 1.4190e-03
# Warning: expt average          C5_H5;C5_H6=5.9374e-03 is larger than maximum value in simulation 4.8894e-03
# Warning: expt average         C5_H6;C5_H5'=1.1945e-03 is larger than maximum value in simulation 2.6436e-04
# Warning: expt average        C5_H6;C5_H5''=2.0312e-03 is larger than maximum value in simulation 3.7150e-04
# Warning: expt average          C5_H6;U6_H5=5.4399e-04 is larger than maximum value in simulation 5.2036e-04
# Warning: expt average       C5_H3';C5_HO2'=1.0834e-04 is smaller than minimum value in simulation 2.9633e-04
# Warning: expt average        U6_H1';U6_H3'=6.2495e-04 is larger than maximum value in simulation 3.2770e-04
# Warning: expt average        U6_H1';U6_H4'=2.7601e-03 is larger than maximum value in simulation 8.9815e-04
# Warning: expt average        U6_H1';U6_H2'=3.6371e-03 is larger than maximum value in simulation 2.0814e-03
# Warning: expt average         U6_H2';C8_H5=4.5122e-03 is larger than maximum value in simulation 4.0540e-03
# Warning: expt average        U6_H3';U6_H2'=3.2371e-03 is smaller than minimum value in simulation 4.8342e-03
# Warning: expt average         U6_H5;U6_H3'=4.8305e-04 is larger than maximum value in simulation 1.1939e-04
# Warning: expt average         U6_H3';C8_H5=8.1793e-04 is larger than maximum value in simulation 5.2858e-04
# Warning: expt average        U6_H4';U6_H2'=7.3339e-04 is larger than maximum value in simulation 3.7814e-04
# Warning: expt average         C5_H2';U6_H6=6.7551e-03 is larger than maximum value in simulation 5.0333e-03
# Warning: expt average         C5_H3';U6_H6=3.8131e-03 is larger than maximum value in simulation 3.6014e-03
# Warning: expt average          C5_H5;U6_H6=1.1572e-04 is larger than maximum value in simulation 5.5877e-05
# Warning: expt average          C5_H6;U6_H6=2.1052e-04 is larger than maximum value in simulation 1.2762e-04
# Warning: expt average        C5_HO2';U6_H6=1.9295e-04 is larger than maximum value in simulation 1.4903e-04
# Warning: expt average         U6_H2';U6_H6=7.7431e-04 is larger than maximum value in simulation 4.0205e-04
# Warning: expt average          U6_H5;U6_H6=8.1315e-03 is larger than maximum value in simulation 5.3750e-03
# Warning: expt average        U6_H6;U6_HO2'=6.1012e-05 is larger than maximum value in simulation 5.4884e-05
# Warning: expt average       U6_H2';U6_HO2'=1.0708e-02 is larger than maximum value in simulation 9.9815e-03
# Warning: expt average       U6_H3';U6_HO2'=5.6302e-04 is larger than maximum value in simulation 2.5012e-04
# Warning: expt average       U6_H4';U6_HO2'=1.2181e-03 is larger than maximum value in simulation 6.6556e-04
# Warning: expt average       U6_H5';U6_HO2'=1.2205e-04 is larger than maximum value in simulation 3.3693e-05
# Warning: expt average       U6_HO2';U7_H1'=1.9575e-04 is larger than maximum value in simulation 3.6783e-05
# Warning: expt average       U6_HO2';U7_H2'=4.6238e-03 is larger than maximum value in simulation 4.5565e-03
# Warning: expt average        U7_H1';U7_H4'=1.9466e-03 is larger than maximum value in simulation 1.0682e-03
# Warning: expt average        U7_H1';U7_H2'=2.0312e-03 is larger than maximum value in simulation 1.2481e-03
# Warning: expt average         U7_H2';U7_H5=1.7211e-04 is larger than maximum value in simulation 1.4258e-04
# Warning: expt average         U6_H2';U7_H6=4.0926e-04 is larger than maximum value in simulation 2.6192e-04
# Warning: expt average         U7_H1';U7_H6=9.8547e-04 is larger than maximum value in simulation 3.9543e-04
# Warning: expt average         U7_H2';U7_H6=1.0412e-02 is larger than maximum value in simulation 6.3949e-03
# Warning: expt average         U7_H6;U7_H3'=5.2571e-04 is larger than maximum value in simulation 2.3501e-04
# Warning: expt average         U7_H6;U7_H4'=3.2693e-04 is larger than maximum value in simulation 1.3159e-04
# Warning: expt average          U7_H5;U7_H6=7.9161e-03 is larger than maximum value in simulation 5.3494e-03
# Warning: expt average         U7_H6;U7_H5'=7.3339e-04 is larger than maximum value in simulation 4.0620e-04
# Warning: expt average        C8_H1';C8_H4'=1.9884e-03 is larger than maximum value in simulation 1.1780e-03
# Warning: expt average        C8_H1';C8_H2'=2.4156e-03 is larger than maximum value in simulation 1.1961e-03
# Warning: expt average        C8_H4';C8_H2'=3.7735e-04 is larger than maximum value in simulation 3.4681e-04
# Warning: expt average        C8_H2';C8_H5'=1.1270e-04 is smaller than minimum value in simulation 1.2620e-04
# Warning: expt average        C8_H4';C8_H3'=2.8875e-03 is larger than maximum value in simulation 2.6487e-03
# Warning: expt average        C8_H5';C8_H3'=3.0233e-04 is smaller than minimum value in simulation 3.6231e-04
# Warning: expt average       C8_H4';C8_H5''=7.3088e-03 is larger than maximum value in simulation 5.6359e-03
# Warning: expt average        U6_H2';C8_H41=9.3132e-04 is larger than maximum value in simulation 7.1850e-04
# Warning: expt average        U6_H3';C8_H41=1.4868e-03 is larger than maximum value in simulation 8.8078e-04
# Warning: expt average         C8_H5;C8_H41=3.9048e-03 is larger than maximum value in simulation 4.4570e-04
# Warning: expt average        U6_H3';C8_H42=2.1664e-03 is smaller than minimum value in simulation 2.4940e-03
# Warning: expt average       C8_H42;U7_H5''=6.5558e-05 is smaller than minimum value in simulation 1.4343e-04
# Warning: expt average        C8_H41;C8_H42=3.2521e-02 is smaller than minimum value in simulation 3.5031e-02
# Warning: expt average         C8_H5;C8_H42=2.6989e-03 is smaller than minimum value in simulation 5.4691e-03
# Warning: expt average         C8_H5;U7_H3'=3.0707e-04 is smaller than minimum value in simulation 5.6751e-04
# Warning: expt average         C8_H5;U7_H5'=5.9304e-04 is larger than maximum value in simulation 5.6810e-04
# Warning: expt average         C8_H5;C8_H2'=7.3339e-04 is larger than maximum value in simulation 1.2386e-04
# Warning: expt average       C8_H5';C8_H5''=1.3473e-02 is smaller than minimum value in simulation 3.1921e-02
# Warning: expt average         U6_H2';C8_H6=5.5341e-04 is larger than maximum value in simulation 3.3897e-04
# Warning: expt average         C8_H1';C8_H6=1.1945e-03 is larger than maximum value in simulation 4.6085e-04
# Warning: expt average         C8_H2';C8_H6=9.5773e-03 is larger than maximum value in simulation 4.8069e-03
# Warning: expt average          C8_H5;C8_H6=5.5022e-03 is larger than maximum value in simulation 4.8402e-03
# Warning: expt average        C8_H5'';C8_H6=4.2992e-04 is larger than maximum value in simulation 2.3879e-04
# Warning: expt average         G9_H1;G10_H8=1.3231e-04 is larger than maximum value in simulation 3.4944e-05
# Warning: expt average         U6_H1';G9_H1=1.6133e-03 is larger than maximum value in simulation 9.2906e-04
# Warning: expt average         U6_H2';G9_H1=1.0433e-03 is larger than maximum value in simulation 7.3870e-04
# Warning: expt average        U6_HO2';G9_H1=1.9884e-03 is larger than maximum value in simulation 1.4883e-03
# Warning: expt average         U7_H2';G9_H1=2.1052e-04 is larger than maximum value in simulation 1.0919e-04
# Warning: expt average         C8_H1';G9_H1=4.9125e-04 is larger than maximum value in simulation 4.2787e-04
# Warning: expt average         C8_H42;G9_H1=2.6731e-04 is larger than maximum value in simulation 3.8216e-05
# Warning: expt average          C8_H5;G9_H1=2.0747e-04 is larger than maximum value in simulation 1.1520e-04
# Warning: expt average          C8_H6;G9_H1=3.5959e-04 is larger than maximum value in simulation 2.9873e-04
# Warning: expt average        G9_H1';G9_H2'=5.3655e-03 is larger than maximum value in simulation 2.0144e-03
# Warning: expt average        G9_H1';G9_H3'=4.6713e-04 is larger than maximum value in simulation 3.0654e-04
# Warning: expt average        G9_H1';G9_H4'=2.8875e-03 is larger than maximum value in simulation 1.2012e-03
# Warning: expt average        G9_H2';G9_H3'=7.3088e-03 is larger than maximum value in simulation 5.3320e-03
# Warning: expt average        C8_H4';G9_H5'=1.3231e-04 is smaller than minimum value in simulation 1.6035e-04
# Warning: expt average        G9_H1';G9_H5'=1.1123e-04 is larger than maximum value in simulation 6.2269e-05
# Warning: expt average       G9_H3';G9_H5''=7.5050e-03 is larger than maximum value in simulation 5.9389e-03
# Warning: expt average         C8_H4';G9_H8=1.5600e-04 is larger than maximum value in simulation 1.2326e-04
# Warning: expt average        C8_H5'';G9_H8=6.5887e-04 is larger than maximum value in simulation 7.9027e-05
# Warning: expt average         G9_H1';G9_H8=1.1332e-02 is larger than maximum value in simulation 5.8116e-03
# Warning: expt average         G9_H2';G9_H8=4.9125e-04 is larger than maximum value in simulation 3.5401e-04
# Warning: expt average         G9_H3';G9_H8=6.7160e-05 is larger than maximum value in simulation 4.6395e-05
# Warning: expt average         G9_H4';G9_H8=3.2693e-04 is larger than maximum value in simulation 4.0331e-05
# Warning: expt average       G10_H1;G10_H21=8.4859e-04 is larger than maximum value in simulation 5.7144e-04
# Warning: expt average       G10_H1;G10_H22=3.9048e-03 is smaller than minimum value in simulation 7.8172e-03
# Warning: expt average       G10_H1;U11_H1'=1.5173e-04 is larger than maximum value in simulation 5.8090e-05
# Warning: expt average        C5_H2';G10_H1=5.0814e-04 is larger than maximum value in simulation 1.1060e-04
# Warning: expt average        C5_H41;G10_H1=3.1682e-04 is smaller than minimum value in simulation 1.5689e-03
# Warning: expt average        C5_H42;G10_H1=4.2982e-03 is larger than maximum value in simulation 2.7480e-04
# Warning: expt average         C5_H5;G10_H1=4.4437e-04 is larger than maximum value in simulation 6.3488e-05
# Warning: expt average        U6_H1';G10_H1=5.0814e-04 is larger than maximum value in simulation 2.1182e-04
# Warning: expt average         G9_H1;G10_H1=4.1602e-04 is larger than maximum value in simulation 9.3461e-05
# Warning: expt average       U6_H1';G10_H21=4.0264e-04 is smaller than minimum value in simulation 4.6908e-04
# Warning: expt average      G10_H21;G10_H22=9.5773e-03 is smaller than minimum value in simulation 3.4119e-02
# Warning: expt average       C5_H42;G10_H22=6.0291e-05 is larger than maximum value in simulation 3.0140e-05
# Warning: expt average       G10_H8;G10_H4'=1.5485e-03 is larger than maximum value in simulation 2.5099e-04
# Warning: expt average      G10_H8;G10_H5''=7.9161e-03 is larger than maximum value in simulation 1.2899e-03
# Warning: expt average        G10_H8;G9_H3'=1.3446e-03 is larger than maximum value in simulation 6.6994e-04
# Warning: expt average     G10_HO2';G10_H2'=4.7387e-03 is larger than maximum value in simulation 2.8294e-03
# Warning: expt average    G10_H5'';G10_HO2'=9.8547e-04 is larger than maximum value in simulation 1.6922e-04
# Warning: expt average      U11_H1';U11_H2'=4.9786e-03 is larger than maximum value in simulation 2.0304e-03
# Warning: expt average       G10_H21;U11_H3=4.0264e-04 is larger than maximum value in simulation 6.1832e-05
# Warning: expt average       U11_H1';U11_H3=2.2999e-04 is larger than maximum value in simulation 1.2011e-04
# Warning: expt average        U11_H3;U11_H6=2.2999e-04 is larger than maximum value in simulation 8.5961e-05
# Warning: expt average         A4_H2;U11_H3=3.0917e-03 is larger than maximum value in simulation 1.9520e-03
# Warning: expt average        A4_H61;U11_H3=2.6731e-04 is smaller than minimum value in simulation 5.6245e-03
# Warning: expt average        A4_H62;U11_H3=1.3995e-03 is larger than maximum value in simulation 4.4615e-04
# Warning: expt average        C5_H42;U11_H3=1.3595e-04 is larger than maximum value in simulation 3.1980e-05
# Warning: expt average       G10_H2';U11_H6=9.0643e-03 is larger than maximum value in simulation 6.8047e-03
# Warning: expt average       G10_H3';U11_H6=1.7894e-03 is larger than maximum value in simulation 9.2212e-04
# Warning: expt average      G10_HO2';U11_H6=1.3259e-05 is smaller than minimum value in simulation 4.7269e-05
# Warning: expt average       U11_H1';U11_H6=6.2495e-04 is larger than maximum value in simulation 4.6077e-04
# Warning: expt average       U11_H6;U11_H3'=5.6430e-03 is larger than maximum value in simulation 2.0774e-03
# Warning: expt average        U11_H6;U11_H5=3.1634e-03 is smaller than minimum value in simulation 5.2498e-03
# Warning: expt average      U11_H6;U11_H5''=6.9497e-04 is larger than maximum value in simulation 3.8944e-04
# Warning: expt average     U11_H2';U11_HO2'=2.5246e-03 is smaller than minimum value in simulation 2.9598e-03
# Warning: expt average       G12_H1;G12_H21=3.1190e-04 is smaller than minimum value in simulation 5.6147e-04
# Warning: expt average       G12_H1;G12_H22=1.0635e-03 is smaller than minimum value in simulation 8.2151e-03
# Warning: expt average        G12_H1;C3_H41=2.0151e-04 is smaller than minimum value in simulation 5.1229e-03
# Warning: expt average        G12_H1;C3_H42=5.9374e-03 is larger than maximum value in simulation 5.1177e-04
# Warning: expt average      G12_H1';U11_H2'=1.0841e-03 is larger than maximum value in simulation 2.9003e-04
# Warning: expt average      G12_H1';G12_H2'=9.1405e-04 is smaller than minimum value in simulation 2.0209e-03
# Warning: expt average       U11_H2';G12_H8=1.0124e-02 is larger than maximum value in simulation 6.6816e-03
# Warning: expt average      U11_HO2';G12_H8=1.1420e-04 is smaller than minimum value in simulation 1.2840e-04
# Warning: expt average       G12_H1';G12_H8=6.8269e-04 is larger than maximum value in simulation 3.4774e-04
# Warning: expt average      G12_H8;G12_H5''=1.6812e-03 is larger than maximum value in simulation 3.6341e-04
# Warning: expt average     G12_H1';G12_HO2'=2.5246e-03 is larger than maximum value in simulation 1.4823e-03
# Warning: expt average      C13_H1';C13_H5'=4.2992e-04 is larger than maximum value in simulation 9.7577e-05
# Warning: expt average      C13_H1';C13_H2'=3.3906e-03 is larger than maximum value in simulation 1.9151e-03
# Warning: expt average      C13_H1';C13_H3'=6.0346e-04 is larger than maximum value in simulation 2.9328e-04
# Warning: expt average       C13_H41;C13_H5=4.5122e-03 is larger than maximum value in simulation 4.0607e-04
# Warning: expt average       C13_H41;C14_H5=1.8481e-04 is larger than maximum value in simulation 6.6917e-05
# Warning: expt average       C13_H1';C13_H6=1.0043e-03 is larger than maximum value in simulation 4.9257e-04
# Warning: expt average       C13_H41;C13_H6=1.1123e-04 is larger than maximum value in simulation 4.2298e-05
# Warning: expt average        C13_H5;C13_H6=5.3655e-03 is larger than maximum value in simulation 4.8979e-03
# Warning: expt average     C13_H1';C13_HO2'=1.8661e-03 is larger than maximum value in simulation 1.0737e-03
# Warning: expt average     C13_H3';C13_HO2'=2.4783e-04 is smaller than minimum value in simulation 3.4422e-04
# Warning: expt average    C13_HO2';C14_H5''=7.4042e-05 is smaller than minimum value in simulation 9.4531e-05
# Warning: expt average      C14_H2';C14_H1'=3.5527e-03 is larger than maximum value in simulation 1.8334e-03
# Warning: expt average      C14_H1';C14_H3'=5.6302e-04 is larger than maximum value in simulation 2.7747e-04
# Warning: expt average       C14_H41;C13_H5=2.3694e-04 is larger than maximum value in simulation 1.5853e-04
# Warning: expt average       C14_H41;C14_H5=4.7387e-03 is larger than maximum value in simulation 4.0753e-04
# Warning: expt average      C14_H41;C14_H42=2.5769e-02 is smaller than minimum value in simulation 3.3829e-02
# Warning: expt average       C14_H42;C14_H5=4.7501e-04 is smaller than minimum value in simulation 4.6352e-03
# Warning: expt average       C13_H2';C14_H6=8.3538e-03 is larger than maximum value in simulation 5.1924e-03
# Warning: expt average       C14_H1';C14_H6=8.4859e-04 is larger than maximum value in simulation 4.6666e-04
# Warning: expt average       C14_H2';C14_H6=2.5246e-03 is larger than maximum value in simulation 1.0542e-03
# Warning: expt average       C14_H3';C14_H6=5.7880e-03 is larger than maximum value in simulation 2.3308e-03
# Warning: expt average       C14_H41;C14_H6=8.9296e-05 is larger than maximum value in simulation 4.2303e-05
# Warning: expt average        C14_H5;C14_H6=6.2498e-03 is larger than maximum value in simulation 4.8858e-03
# theta: 1000
# Warning: expt average        G1_H1;C14_H41=8.3836e-05 is smaller than minimum value in simulation 9.2617e-05
# Warning: expt average        G1_H1;C14_H42=2.9310e-04 is larger than maximum value in simulation 1.0486e-04
# Warning: expt average        G1_H4';G1_H1'=1.1945e-03 is larger than maximum value in simulation 6.1313e-04
# Warning: expt average         G1_H1';G1_H8=6.0346e-04 is larger than maximum value in simulation 3.8121e-04
# Warning: expt average         G1_H8;G1_H3'=4.4437e-04 is smaller than minimum value in simulation 1.8508e-03
# Warning: expt average        G1_H8;G1_H5''=1.0236e-03 is larger than maximum value in simulation 3.3281e-04
# Warning: expt average        G2_H1;C13_H41=2.2661e-04 is smaller than minimum value in simulation 7.3883e-04
# Warning: expt average        C14_H42;G2_H1=3.1190e-04 is larger than maximum value in simulation 3.0909e-04
# Warning: expt average         G2_H1;C3_H42=4.1957e-03 is larger than maximum value in simulation 1.8415e-04
# Warning: expt average        G2_H1';G2_H2'=6.7551e-03 is larger than maximum value in simulation 2.1800e-03
# Warning: expt average         G1_H1';G2_H8=5.2571e-04 is larger than maximum value in simulation 6.5039e-05
# Warning: expt average         G1_H3';G2_H8=5.2328e-03 is larger than maximum value in simulation 9.3904e-04
# Warning: expt average         G2_H1';G2_H8=6.1409e-04 is larger than maximum value in simulation 3.8176e-04
# Warning: expt average         G2_H8;G2_H3'=4.4037e-03 is larger than maximum value in simulation 3.4787e-03
# Warning: expt average         G2_H8;G2_H5'=1.1332e-02 is larger than maximum value in simulation 6.9373e-04
# Warning: expt average        C3_H1';C3_H4'=1.9059e-03 is larger than maximum value in simulation 1.0429e-03
# Warning: expt average         G2_H1;C3_H41=1.4759e-04 is smaller than minimum value in simulation 1.7331e-04
# Warning: expt average         C3_H41;C3_H5=6.4132e-03 is larger than maximum value in simulation 4.3113e-04
# Warning: expt average         C3_H42;C3_H5=4.9125e-04 is smaller than minimum value in simulation 5.7543e-03
# Warning: expt average         G2_H1';C3_H6=9.6405e-05 is larger than maximum value in simulation 9.2040e-05
# Warning: expt average         C3_H1';C3_H6=6.9497e-04 is larger than maximum value in simulation 4.4622e-04
# Warning: expt average          C3_H5;C3_H6=5.6430e-03 is larger than maximum value in simulation 4.4166e-03
# Warning: expt average       C3_HO2';C3_H3'=9.4898e-04 is larger than maximum value in simulation 4.7684e-04
# Warning: expt average        A4_H1';C5_H1'=1.2181e-03 is larger than maximum value in simulation 3.8023e-05
# Warning: expt average        A4_H2;G12_H1'=8.1793e-04 is larger than maximum value in simulation 3.3228e-04
# Warning: expt average         A4_H1';A4_H2=1.3411e-04 is larger than maximum value in simulation 1.1148e-04
# Warning: expt average         C5_H1';A4_H2=1.1268e-03 is larger than maximum value in simulation 1.0967e-03
# Warning: expt average        A4_H1';A4_H3'=2.1202e-03 is larger than maximum value in simulation 3.1796e-04
# Warning: expt average        A4_H61;A4_H62=6.4132e-03 is smaller than minimum value in simulation 3.4839e-02
# Warning: expt average         A4_H1';A4_H8=7.4674e-04 is larger than maximum value in simulation 5.6653e-04
# Warning: expt average        A4_H8;A4_H5''=1.4570e-03 is smaller than minimum value in simulation 6.8475e-03
# Warning: expt average       A4_H3';A4_HO2'=3.5527e-03 is larger than maximum value in simulation 6.2265e-04
# Warning: expt average        C5_H1';C5_H2'=1.5485e-03 is smaller than minimum value in simulation 1.7372e-03
# Warning: expt average        C5_H1';C5_H3'=3.4830e-04 is larger than maximum value in simulation 3.3160e-04
# Warning: expt average         C5_H5;C5_H41=5.2328e-03 is larger than maximum value in simulation 4.3120e-04
# Warning: expt average         U6_H5;C5_H41=3.5959e-04 is larger than maximum value in simulation 1.7899e-04
# Warning: expt average        C5_H41;C5_H42=2.9401e-02 is smaller than minimum value in simulation 3.4792e-02
# Warning: expt average         C5_H5;C5_H42=5.3476e-04 is smaller than minimum value in simulation 5.7572e-03
# Warning: expt average         A4_H3';C5_H6=8.1315e-03 is larger than maximum value in simulation 1.7701e-03
# Warning: expt average         C5_H1';C5_H6=5.3476e-04 is larger than maximum value in simulation 5.2240e-04
# Warning: expt average         C5_H6;C5_H3'=5.9374e-03 is larger than maximum value in simulation 5.0009e-03
# Warning: expt average          C5_H5;C5_H6=5.9374e-03 is larger than maximum value in simulation 4.4187e-03
# Warning: expt average         C5_H6;C5_H5'=1.1945e-03 is larger than maximum value in simulation 3.6063e-04
# Warning: expt average       C5_H3';C5_HO2'=1.0834e-04 is smaller than minimum value in simulation 2.4986e-04
# Warning: expt average       C5_HO2';U6_H5'=1.3717e-03 is larger than maximum value in simulation 6.7664e-04
# Warning: expt average        U6_H1';U6_H3'=6.2495e-04 is larger than maximum value in simulation 3.3160e-04
# Warning: expt average        U6_H1';U6_H4'=2.7601e-03 is larger than maximum value in simulation 7.4418e-04
# Warning: expt average        U6_H1';U6_H2'=3.6371e-03 is larger than maximum value in simulation 2.3222e-03
# Warning: expt average        U6_H3';U6_H2'=3.2371e-03 is smaller than minimum value in simulation 5.6860e-03
# Warning: expt average         U6_H5;U6_H3'=4.8305e-04 is larger than maximum value in simulation 1.6732e-04
# Warning: expt average         U6_H3';C8_H5=8.1793e-04 is larger than maximum value in simulation 7.4097e-04
# Warning: expt average        U6_H4';U6_H2'=7.3339e-04 is larger than maximum value in simulation 3.3945e-04
# Warning: expt average          C5_H5;U6_H6=1.1572e-04 is larger than maximum value in simulation 9.2641e-05
# Warning: expt average        C5_HO2';U6_H6=1.9295e-04 is larger than maximum value in simulation 1.3760e-04
# Warning: expt average         U6_H2';U6_H6=7.7431e-04 is larger than maximum value in simulation 4.7589e-04
# Warning: expt average          U6_H5;U6_H6=8.1315e-03 is larger than maximum value in simulation 4.6092e-03
# Warning: expt average        U6_H6;U6_HO2'=6.1012e-05 is larger than maximum value in simulation 6.0226e-05
# Warning: expt average       U6_H2';U6_HO2'=1.0708e-02 is larger than maximum value in simulation 7.1939e-03
# Warning: expt average       U6_H3';U6_HO2'=5.6302e-04 is larger than maximum value in simulation 2.5787e-04
# Warning: expt average       U6_H4';U6_HO2'=1.2181e-03 is larger than maximum value in simulation 1.1569e-03
# Warning: expt average       U6_H5';U6_HO2'=1.2205e-04 is larger than maximum value in simulation 4.6048e-05
# Warning: expt average       U6_HO2';U7_H1'=1.9575e-04 is larger than maximum value in simulation 4.4721e-05
# Warning: expt average       U6_HO2';U7_H2'=4.6238e-03 is larger than maximum value in simulation 1.9578e-03
# Warning: expt average        U7_H1';U7_H4'=1.9466e-03 is larger than maximum value in simulation 6.3446e-04
# Warning: expt average        U7_H1';U7_H2'=2.0312e-03 is larger than maximum value in simulation 1.2793e-03
# Warning: expt average         U7_H2';U7_H5=1.7211e-04 is larger than maximum value in simulation 1.5986e-04
# Warning: expt average         U6_H4';U7_H5=2.9538e-03 is larger than maximum value in simulation 2.3091e-03
# Warning: expt average         U6_H2';U7_H6=4.0926e-04 is larger than maximum value in simulation 1.7583e-04
# Warning: expt average         U7_H1';U7_H6=9.8547e-04 is larger than maximum value in simulation 6.6473e-04
# Warning: expt average         U7_H6;U7_H3'=5.2571e-04 is larger than maximum value in simulation 2.1226e-04
# Warning: expt average         U7_H6;U7_H4'=3.2693e-04 is larger than maximum value in simulation 2.4568e-04
# Warning: expt average          U7_H5;U7_H6=7.9161e-03 is larger than maximum value in simulation 4.6069e-03
# Warning: expt average         U7_H6;U7_H5'=7.3339e-04 is larger than maximum value in simulation 5.7921e-04
# Warning: expt average        C8_H1';C8_H4'=1.9884e-03 is larger than maximum value in simulation 1.1069e-03
# Warning: expt average        C8_H1';C8_H2'=2.4156e-03 is larger than maximum value in simulation 1.2488e-03
# Warning: expt average        C8_H4';C8_H2'=3.7735e-04 is larger than maximum value in simulation 3.1931e-04
# Warning: expt average        C8_H2';C8_H5'=1.1270e-04 is smaller than minimum value in simulation 1.4622e-04
# Warning: expt average        C8_H4';C8_H3'=2.8875e-03 is larger than maximum value in simulation 2.8243e-03
# Warning: expt average        C8_H5';C8_H3'=3.0233e-04 is smaller than minimum value in simulation 3.8424e-04
# Warning: expt average       C8_H4';C8_H5''=7.3088e-03 is larger than maximum value in simulation 6.6486e-03
# Warning: expt average        U6_H2';C8_H41=9.3132e-04 is larger than maximum value in simulation 6.7784e-04
# Warning: expt average        U6_H3';C8_H41=1.4868e-03 is larger than maximum value in simulation 9.2662e-04
# Warning: expt average         C8_H5;C8_H41=3.9048e-03 is larger than maximum value in simulation 4.3106e-04
# Warning: expt average        C8_H41;C8_H42=3.2521e-02 is smaller than minimum value in simulation 3.4780e-02
# Warning: expt average         C8_H5;C8_H42=2.6989e-03 is smaller than minimum value in simulation 5.7586e-03
# Warning: expt average         C8_H5;C8_H2'=7.3339e-04 is larger than maximum value in simulation 1.1285e-04
# Warning: expt average       C8_H5';C8_H5''=1.3473e-02 is smaller than minimum value in simulation 3.3201e-02
# Warning: expt average         U6_H2';C8_H6=5.5341e-04 is larger than maximum value in simulation 2.4410e-04
# Warning: expt average         C8_H1';C8_H6=1.1945e-03 is larger than maximum value in simulation 4.1704e-04
# Warning: expt average         C8_H2';C8_H6=9.5773e-03 is larger than maximum value in simulation 3.8122e-03
# Warning: expt average         C8_H41;C8_H6=4.3615e-05 is larger than maximum value in simulation 4.1964e-05
# Warning: expt average          C8_H5;C8_H6=5.5022e-03 is larger than maximum value in simulation 4.4176e-03
# Warning: expt average        C8_H5'';C8_H6=4.2992e-04 is larger than maximum value in simulation 2.4203e-04
# Warning: expt average         G9_H1;G10_H8=1.3231e-04 is larger than maximum value in simulation 5.2770e-05
# Warning: expt average         U6_H1';G9_H1=1.6133e-03 is larger than maximum value in simulation 1.6106e-03
# Warning: expt average         U6_H2';G9_H1=1.0433e-03 is larger than maximum value in simulation 8.1388e-04
# Warning: expt average        U6_HO2';G9_H1=1.9884e-03 is larger than maximum value in simulation 1.6162e-03
# Warning: expt average         C8_H1';G9_H1=4.9125e-04 is larger than maximum value in simulation 3.6699e-04
# Warning: expt average         C8_H42;G9_H1=2.6731e-04 is larger than maximum value in simulation 2.6583e-05
# Warning: expt average          C8_H5;G9_H1=2.0747e-04 is larger than maximum value in simulation 1.0911e-04
# Warning: expt average        G9_H1';G9_H2'=5.3655e-03 is larger than maximum value in simulation 1.6959e-03
# Warning: expt average        G9_H1';G9_H3'=4.6713e-04 is larger than maximum value in simulation 3.3784e-04
# Warning: expt average        G9_H1';G9_H4'=2.8875e-03 is larger than maximum value in simulation 2.6641e-03
# Warning: expt average        G9_H2';G9_H3'=7.3088e-03 is larger than maximum value in simulation 6.5288e-03
# Warning: expt average        C8_H4';G9_H5'=1.3231e-04 is smaller than minimum value in simulation 2.0357e-04
# Warning: expt average        G9_H3';G9_H5'=1.5173e-03 is larger than maximum value in simulation 7.5244e-04
# Warning: expt average       C8_H4';G9_H5''=5.6430e-03 is larger than maximum value in simulation 3.6440e-03
# Warning: expt average       C8_H5';G9_H5''=6.4000e-05 is larger than maximum value in simulation 6.3686e-05
# Warning: expt average         C8_H4';G9_H8=1.5600e-04 is larger than maximum value in simulation 5.1775e-05
# Warning: expt average        C8_H5'';G9_H8=6.5887e-04 is larger than maximum value in simulation 9.2594e-05
# Warning: expt average         G9_H1';G9_H8=1.1332e-02 is larger than maximum value in simulation 3.9620e-03
# Warning: expt average         G9_H2';G9_H8=4.9125e-04 is larger than maximum value in simulation 1.6125e-04
# Warning: expt average         G9_H3';G9_H8=6.7160e-05 is larger than maximum value in simulation 3.9877e-05
# Warning: expt average         G9_H4';G9_H8=3.2693e-04 is larger than maximum value in simulation 5.7585e-05
# Warning: expt average       G10_H1;G10_H21=8.4859e-04 is smaller than minimum value in simulation 1.2979e-02
# Warning: expt average       G10_H1;G10_H22=3.9048e-03 is larger than maximum value in simulation 6.7221e-04
# Warning: expt average       G10_H1;U11_H1'=1.5173e-04 is larger than maximum value in simulation 8.9819e-05
# Warning: expt average        C5_H2';G10_H1=5.0814e-04 is larger than maximum value in simulation 1.7109e-04
# Warning: expt average        C5_H41;G10_H1=3.1682e-04 is smaller than minimum value in simulation 3.5971e-04
# Warning: expt average        C5_H42;G10_H1=4.2982e-03 is larger than maximum value in simulation 4.4027e-04
# Warning: expt average         C5_H5;G10_H1=4.4437e-04 is larger than maximum value in simulation 1.0545e-04
# Warning: expt average         G9_H1;G10_H1=4.1602e-04 is larger than maximum value in simulation 1.8753e-04
# Warning: expt average      G10_H21;U11_H1'=4.4437e-04 is larger than maximum value in simulation 3.7961e-04
# Warning: expt average      G10_H21;G10_H22=9.5773e-03 is smaller than minimum value in simulation 3.4827e-02
# Warning: expt average       C5_H42;G10_H22=6.0291e-05 is larger than maximum value in simulation 1.1141e-05
# Warning: expt average       G10_H8;G10_H4'=1.5485e-03 is larger than maximum value in simulation 2.8811e-04
# Warning: expt average      G10_H8;G10_H5''=7.9161e-03 is larger than maximum value in simulation 5.0433e-03
# Warning: expt average        G10_H8;G9_H2'=2.5932e-04 is larger than maximum value in simulation 2.5775e-04
# Warning: expt average        G10_H8;G9_H3'=1.3446e-03 is larger than maximum value in simulation 4.3213e-04
# Warning: expt average    G10_H5'';G10_HO2'=9.8547e-04 is larger than maximum value in simulation 1.1217e-04
# Warning: expt average      U11_H1';U11_H2'=4.9786e-03 is larger than maximum value in simulation 2.2702e-03
# Warning: expt average       G10_H21;U11_H3=4.0264e-04 is larger than maximum value in simulation 2.4513e-04
# Warning: expt average       U11_H1';U11_H3=2.2999e-04 is larger than maximum value in simulation 1.1912e-04
# Warning: expt average        U11_H3;U11_H6=2.2999e-04 is larger than maximum value in simulation 8.5314e-05
# Warning: expt average        A4_H61;U11_H3=2.6731e-04 is smaller than minimum value in simulation 4.7263e-04
# Warning: expt average        A4_H62;U11_H3=1.3995e-03 is larger than maximum value in simulation 9.0495e-05
# Warning: expt average        C5_H42;U11_H3=1.3595e-04 is larger than maximum value in simulation 9.2570e-05
# Warning: expt average       G10_H3';U11_H6=1.7894e-03 is larger than maximum value in simulation 1.6286e-03
# Warning: expt average      G10_HO2';U11_H6=1.3259e-05 is smaller than minimum value in simulation 1.9806e-05
# Warning: expt average        U11_H6;U11_H5=3.1634e-03 is smaller than minimum value in simulation 4.5956e-03
# Warning: expt average     U11_H1';U11_HO2'=7.3339e-04 is smaller than minimum value in simulation 2.0469e-03
# Warning: expt average       G12_H1;G12_H21=3.1190e-04 is smaller than minimum value in simulation 1.2983e-02
# Warning: expt average       G12_H1;G12_H22=1.0635e-03 is larger than maximum value in simulation 6.7268e-04
# Warning: expt average        G12_H1;C3_H41=2.0151e-04 is smaller than minimum value in simulation 2.5608e-04
# Warning: expt average        G12_H1;C3_H42=5.9374e-03 is larger than maximum value in simulation 3.7729e-04
# Warning: expt average      G12_H1';G12_H2'=9.1405e-04 is smaller than minimum value in simulation 1.9107e-03
# Warning: expt average       U11_H2';G12_H8=1.0124e-02 is larger than maximum value in simulation 9.3655e-03
# Warning: expt average       G12_H1';G12_H8=6.8269e-04 is larger than maximum value in simulation 5.6770e-04
# Warning: expt average      C13_H1';C13_H5'=4.2992e-04 is larger than maximum value in simulation 9.7628e-05
# Warning: expt average      C13_H1';C13_H2'=3.3906e-03 is larger than maximum value in simulation 2.2224e-03
# Warning: expt average      C13_H1';C13_H3'=6.0346e-04 is larger than maximum value in simulation 3.2889e-04
# Warning: expt average       C13_H41;C13_H5=4.5122e-03 is larger than maximum value in simulation 4.3099e-04
# Warning: expt average      G12_HO2';C13_H6=1.3595e-04 is larger than maximum value in simulation 8.2864e-05
# Warning: expt average       C13_H1';C13_H6=1.0043e-03 is larger than maximum value in simulation 4.4860e-04
# Warning: expt average       C13_H41;C13_H6=1.1123e-04 is larger than maximum value in simulation 4.1964e-05
# Warning: expt average        C13_H5;C13_H6=5.3655e-03 is larger than maximum value in simulation 4.4166e-03
# Warning: expt average     C13_H1';C13_HO2'=1.8661e-03 is smaller than minimum value in simulation 1.8732e-03
# Warning: expt average      C14_H2';C14_H1'=3.5527e-03 is larger than maximum value in simulation 2.0592e-03
# Warning: expt average      C14_H1';C14_H3'=5.6302e-04 is larger than maximum value in simulation 3.3470e-04
# Warning: expt average      C14_H1';C14_H4'=1.2181e-03 is larger than maximum value in simulation 7.8567e-04
# Warning: expt average       C14_H41;C14_H5=4.7387e-03 is larger than maximum value in simulation 4.3099e-04
# Warning: expt average      C14_H41;C14_H42=2.5769e-02 is smaller than minimum value in simulation 3.4803e-02
# Warning: expt average       C14_H42;C14_H5=4.7501e-04 is smaller than minimum value in simulation 5.7572e-03
# Warning: expt average       C13_H2';C14_H6=8.3538e-03 is larger than maximum value in simulation 5.8220e-03
# Warning: expt average       C14_H1';C14_H6=8.4859e-04 is larger than maximum value in simulation 5.6137e-04
# Warning: expt average       C14_H41;C14_H6=8.9296e-05 is larger than maximum value in simulation 4.1959e-05
# Warning: expt average        C14_H5;C14_H6=6.2498e-03 is larger than maximum value in simulation 4.4166e-03
# theta: 1000
# Warning: expt average               2-2H5P=2.0000e+00 is larger than maximum value in simulation 1.6438e+00
# Warning: expt average               5-2H5P=3.0000e+00 is larger than maximum value in simulation 1.1122e+00
# Warning: expt average               7-2H5P=5.7000e+00 is larger than maximum value in simulation 3.0909e+00
# Warning: expt average               8-2H5P=1.3000e+00 is smaller than minimum value in simulation 2.3523e+00
# Warning: expt average              11-2H5P=0.0000e+00 is smaller than minimum value in simulation 1.3155e+00
# Warning: expt average               2-1H5P=2.1000e+00 is smaller than minimum value in simulation 4.7079e+00
# Warning: expt average               3-1H5P=0.0000e+00 is smaller than minimum value in simulation 1.0961e+00
# Warning: expt average               5-1H5P=1.8000e+00 is smaller than minimum value in simulation 5.8580e+00
# Warning: expt average               7-1H5P=3.7000e+00 is smaller than minimum value in simulation 5.0607e+00
# Warning: expt average               9-1H5P=9.0000e+00 is larger than maximum value in simulation 3.4087e+00
# Warning: expt average              10-1H5P=2.3000e+00 is smaller than minimum value in simulation 2.4495e+00
# Warning: expt average              13-1H5P=1.7000e+00 is smaller than minimum value in simulation 2.8957e+00
# Warning: expt average               2-C4Pb=7.8000e+00 is smaller than minimum value in simulation 8.7650e+00
# Warning: expt average               7-C4Pb=6.3000e+00 is smaller than minimum value in simulation 7.3774e+00
# Warning: expt average              10-C4Pb=6.1000e+00 is smaller than minimum value in simulation 1.0247e+01
# Warning: expt average              14-C4Pb=7.7000e+00 is smaller than minimum value in simulation 8.0409e+00
# Warning: expt average                4-H3P=3.8000e+00 is smaller than minimum value in simulation 5.5853e+00
# Warning: expt average                7-H3P=3.7000e+00 is smaller than minimum value in simulation 6.3984e+00
# Warning: expt average                8-H3P=7.5000e+00 is smaller than minimum value in simulation 7.5325e+00
# Warning: expt average               1-C4Pe=6.0000e+00 is smaller than minimum value in simulation 7.3177e+00
# Warning: expt average               8-C4Pe=7.1000e+00 is larger than maximum value in simulation 2.1340e+00
# Warning: expt average               9-C4Pe=7.7000e+00 is smaller than minimum value in simulation 7.8565e+00
# Warning: expt average              12-C4Pe=6.5000e+00 is smaller than minimum value in simulation 7.8978e+00
# Warning: expt average             G1-2H5H4=5.2000e+00 is larger than maximum value in simulation 3.6428e+00
# Warning: expt average             C5-2H5H4=1.5000e+00 is larger than maximum value in simulation 1.1144e-01
# Warning: expt average             G9-2H5H4=-4.6000e+00 is smaller than minimum value in simulation 8.9079e+00
# Warning: expt average            U11-2H5H4=4.5000e+00 is larger than maximum value in simulation 1.9227e+00
# Warning: expt average            C13-2H5H4=2.8000e+00 is larger than maximum value in simulation 1.0407e-01
# Warning: expt average             C5-1H5H4=1.0000e+00 is smaller than minimum value in simulation 5.6841e+00
# Warning: expt average             U6-1H5H4=-6.0000e-01 is smaller than minimum value in simulation 4.5576e-01
# Warning: expt average             U7-1H5H4=3.0000e-01 is smaller than minimum value in simulation 1.0820e+00
# Warning: expt average             C8-1H5H4=-3.0000e-01 is smaller than minimum value in simulation 8.9092e-01
# Warning: expt average             G9-1H5H4=-2.8000e+00 is smaller than minimum value in simulation 1.1490e+00
# Warning: expt average            U11-1H5H4=-9.0000e-01 is smaller than minimum value in simulation 1.1603e+00
# Warning: expt average            C13-1H5H4=-1.7000e+00 is smaller than minimum value in simulation 3.8121e+00
# Warning: expt average            C14-1H5H4=-8.0000e-01 is smaller than minimum value in simulation 2.1502e+00
# Warning: expt average              G2-H1H2=7.0000e-01 is larger than maximum value in simulation 6.9371e-01
# Warning: expt average              C3-H1H2=4.0000e-01 is smaller than minimum value in simulation 1.2264e+00
# Warning: expt average              C5-H1H2=4.0000e-01 is smaller than minimum value in simulation 4.3977e-01
# Warning: expt average              U6-H1H2=2.0000e-01 is smaller than minimum value in simulation 4.7003e-01
# Warning: expt average              U7-H1H2=8.0000e+00 is smaller than minimum value in simulation 9.7490e+00
# Warning: expt average              C8-H1H2=9.3000e+00 is smaller than minimum value in simulation 1.0310e+01
# Warning: expt average              G9-H1H2=6.0000e-01 is smaller than minimum value in simulation 1.3688e+00
# Warning: expt average             G10-H1H2=6.0000e-01 is larger than maximum value in simulation 5.1267e-02
# Warning: expt average             U11-H1H2=0.0000e+00 is smaller than minimum value in simulation 9.2752e-01
# Warning: expt average             G12-H1H2=1.2000e+00 is larger than maximum value in simulation 9.3010e-01
# Warning: expt average             C13-H1H2=8.0000e-01 is smaller than minimum value in simulation 1.3732e+00
# Warning: expt average             C14-H1H2=3.0000e-01 is smaller than minimum value in simulation 1.7755e+00
# Warning: expt average              G2-H2H3=6.0000e+00 is larger than maximum value in simulation 5.5278e+00
# Warning: expt average              C5-H2H3=3.3000e+00 is smaller than minimum value in simulation 4.0752e+00
# Warning: expt average              U6-H2H3=5.5000e+00 is larger than maximum value in simulation 5.2475e+00
# Warning: expt average              G9-H2H3=3.4000e+00 is smaller than minimum value in simulation 4.9718e+00
# Warning: expt average             C14-H2H3=5.5000e+00 is smaller than minimum value in simulation 5.7488e+00
# Warning: expt average              C3-H3H4=1.1500e+01 is larger than maximum value in simulation 9.9790e+00
# Warning: expt average              A4-H3H4=1.1200e+01 is larger than maximum value in simulation 9.9682e+00
# Warning: expt average              U6-H3H4=1.1000e+01 is larger than maximum value in simulation 1.0688e+01
# Warning: expt average              G9-H3H4=7.6000e+00 is smaller than minimum value in simulation 9.5958e+00
# Warning: expt average             G10-H3H4=4.9000e+00 is smaller than minimum value in simulation 9.5923e+00
# Warning: expt average             U11-H3H4=1.1300e+01 is larger than maximum value in simulation 9.8068e+00
# Warning: expt average             G12-H3H4=1.1500e+01 is larger than maximum value in simulation 1.0318e+01
# Warning: expt average             C13-H3H4=9.1000e+00 is smaller than minimum value in simulation 9.1016e+00
# Warning: expt average             C14-H3H4=5.3000e+00 is smaller than minimum value in simulation 8.5304e+00
# theta: 1000
# Warning: expt average               3-2H5P=1.7000e+00 is smaller than minimum value in simulation 2.0261e+00
# Warning: expt average               7-2H5P=5.7000e+00 is larger than maximum value in simulation 1.9476e+00
# Warning: expt average               9-2H5P=8.6000e+00 is larger than maximum value in simulation 2.8311e+00
# Warning: expt average              11-2H5P=0.0000e+00 is smaller than minimum value in simulation 1.0097e+00
# Warning: expt average               3-1H5P=0.0000e+00 is smaller than minimum value in simulation 1.0402e+00
# Warning: expt average               5-1H5P=1.8000e+00 is smaller than minimum value in simulation 2.4233e+00
# Warning: expt average               6-1H5P=1.7000e+00 is smaller than minimum value in simulation 3.2091e+00
# Warning: expt average               9-1H5P=9.0000e+00 is larger than maximum value in simulation 6.6739e+00
# Warning: expt average              10-1H5P=2.3000e+00 is smaller than minimum value in simulation 6.6074e+00
# Warning: expt average              11-1H5P=1.7000e+00 is smaller than minimum value in simulation 3.5453e+00
# Warning: expt average               2-C4Pb=7.8000e+00 is smaller than minimum value in simulation 8.6058e+00
# Warning: expt average               7-C4Pb=6.3000e+00 is smaller than minimum value in simulation 8.4191e+00
# Warning: expt average               9-C4Pb=8.0000e+00 is smaller than minimum value in simulation 9.5676e+00
# Warning: expt average                4-H3P=3.8000e+00 is smaller than minimum value in simulation 6.2995e+00
# Warning: expt average                7-H3P=3.7000e+00 is smaller than minimum value in simulation 7.4429e+00
# Warning: expt average                8-H3P=7.5000e+00 is larger than maximum value in simulation 7.3570e+00
# Warning: expt average                9-H3P=7.5000e+00 is smaller than minimum value in simulation 8.0303e+00
# Warning: expt average               1-C4Pe=6.0000e+00 is smaller than minimum value in simulation 7.2816e+00
# Warning: expt average               8-C4Pe=7.1000e+00 is larger than maximum value in simulation 5.6977e-01
# Warning: expt average               9-C4Pe=7.7000e+00 is larger than maximum value in simulation 7.4842e+00
# Warning: expt average             C5-2H5H4=1.5000e+00 is smaller than minimum value in simulation 1.6149e+00
# Warning: expt average             U7-2H5H4=1.2000e+00 is smaller than minimum value in simulation 1.7346e+00
# Warning: expt average             G9-2H5H4=-4.6000e+00 is smaller than minimum value in simulation 3.1503e+00
# Warning: expt average             G1-1H5H4=6.0000e-01 is smaller than minimum value in simulation 2.1439e+00
# Warning: expt average             U6-1H5H4=-6.0000e-01 is smaller than minimum value in simulation 7.9734e-03
# Warning: expt average             C8-1H5H4=-3.0000e-01 is smaller than minimum value in simulation -5.7460e-02
# Warning: expt average             G9-1H5H4=-2.8000e+00 is smaller than minimum value in simulation -8.0299e-02
# Warning: expt average            U11-1H5H4=-9.0000e-01 is smaller than minimum value in simulation 4.8390e-01
# Warning: expt average            C13-1H5H4=-1.7000e+00 is smaller than minimum value in simulation -6.4798e-02
# Warning: expt average            C14-1H5H4=-8.0000e-01 is smaller than minimum value in simulation -2.1497e-02
# Warning: expt average              U6-H1H2=2.0000e-01 is smaller than minimum value in simulation 2.0462e-01
# Warning: expt average              U7-H1H2=8.0000e+00 is smaller than minimum value in simulation 9.2553e+00
# Warning: expt average              C8-H1H2=9.3000e+00 is smaller than minimum value in simulation 9.9207e+00
# Warning: expt average              G9-H1H2=6.0000e-01 is smaller than minimum value in simulation 3.1788e+00
# Warning: expt average             U11-H1H2=0.0000e+00 is smaller than minimum value in simulation 3.0276e-01
# Warning: expt average             C14-H1H2=3.0000e-01 is smaller than minimum value in simulation 9.3117e-01
# Warning: expt average              G2-H2H3=6.0000e+00 is larger than maximum value in simulation 4.6888e+00
# Warning: expt average              C5-H2H3=3.3000e+00 is smaller than minimum value in simulation 4.6369e+00
# Warning: expt average              U6-H2H3=5.5000e+00 is larger than maximum value in simulation 4.7024e+00
# Warning: expt average              U7-H2H3=5.2000e+00 is larger than maximum value in simulation 4.2134e+00
# Warning: expt average              C8-H2H3=4.9000e+00 is larger than maximum value in simulation 4.8406e+00
# Warning: expt average              G9-H2H3=3.4000e+00 is smaller than minimum value in simulation 5.0982e+00
# Warning: expt average             C13-H2H3=5.8000e+00 is larger than maximum value in simulation 4.7976e+00
# Warning: expt average             C14-H2H3=5.5000e+00 is larger than maximum value in simulation 4.7537e+00
# Warning: expt average              C3-H3H4=1.1500e+01 is larger than maximum value in simulation 1.0339e+01
# Warning: expt average              A4-H3H4=1.1200e+01 is larger than maximum value in simulation 8.1825e+00
# Warning: expt average              U6-H3H4=1.1000e+01 is larger than maximum value in simulation 1.0418e+01
# Warning: expt average              U7-H3H4=1.0000e+00 is larger than maximum value in simulation 4.2202e-02
# Warning: expt average              G9-H3H4=7.6000e+00 is smaller than minimum value in simulation 1.0402e+01
# Warning: expt average             G10-H3H4=4.9000e+00 is smaller than minimum value in simulation 7.7457e+00
# Warning: expt average             U11-H3H4=1.1300e+01 is larger than maximum value in simulation 1.0729e+01
# Warning: expt average             G12-H3H4=1.1500e+01 is larger than maximum value in simulation 1.0471e+01
# theta: 1000
# RDC scaling factor 6.2675e-01
# Warning: expt average        3.C.C6-3.C.H6=3.5400e+00 is smaller than minimum value in simulation 5.1008e+00
# Warning: expt average        5.C.C6-5.C.H6=1.0990e+01 is larger than maximum value in simulation 7.9153e+00
# Warning: expt average        6.U.C6-6.U.H6=4.0000e+00 is larger than maximum value in simulation 3.1388e+00
# Warning: expt average        7.U.C6-7.U.H6=3.4700e+00 is smaller than minimum value in simulation 5.3826e+00
# Warning: expt average      11.U.C6-11.U.H6=9.0900e+00 is larger than maximum value in simulation 7.0036e+00
# Warning: expt average      13.C.C6-13.C.H6=2.8300e+00 is smaller than minimum value in simulation 3.8019e+00
# Warning: expt average      14.C.C6-14.C.H6=6.4300e+00 is larger than maximum value in simulation 3.0284e+00
# Warning: expt average        1.G.C8-1.G.H8=7.2900e+00 is larger than maximum value in simulation 2.9296e+00
# Warning: expt average        2.G.C8-2.G.H8=1.6300e+00 is smaller than minimum value in simulation 1.7580e+00
# Warning: expt average        4.A.C8-4.A.H8=6.0200e+00 is smaller than minimum value in simulation 6.2388e+00
# Warning: expt average      10.G.C8-10.G.H8=1.1100e+01 is larger than maximum value in simulation 6.0375e+00
# Warning: expt average      12.G.C8-12.G.H8=5.3600e+00 is smaller than minimum value in simulation 5.8298e+00
# Warning: expt average      1.G.C1'-1.G.H1'=-2.3900e+00 is larger than maximum value in simulation -5.2924e+00
# Warning: expt average      2.G.C1'-2.G.H1'=-3.2000e-01 is larger than maximum value in simulation -3.5194e+00
# Warning: expt average      3.C.C1'-3.C.H1'=6.3600e+00 is larger than maximum value in simulation 9.2760e-01
# Warning: expt average      4.A.C1'-4.A.H1'=6.4400e+00 is larger than maximum value in simulation 5.5545e+00
# Warning: expt average      5.C.C1'-5.C.H1'=4.9500e+00 is larger than maximum value in simulation 2.9690e+00
# Warning: expt average      8.C.C1'-8.C.H1'=-1.4660e+01 is smaller than minimum value in simulation -9.0549e+00
# Warning: expt average      9.G.C1'-9.G.H1'=-5.1400e+00 is smaller than minimum value in simulation -3.8133e+00
# Warning: expt average    11.U.C1'-11.U.H1'=-4.3000e-01 is smaller than minimum value in simulation 2.7640e-01
# Warning: expt average    12.G.C1'-12.G.H1'=6.7700e+00 is larger than maximum value in simulation 6.4605e+00
# Warning: expt average        2.G.N1-2.G.H1=3.0000e-01 is smaller than minimum value in simulation 1.0748e+00
# Warning: expt average        9.G.N1-9.G.H1=-6.3200e+00 is smaller than minimum value in simulation 1.7826e+00
# Warning: expt average      10.G.N1-10.G.H1=-3.8800e+00 is smaller than minimum value in simulation 2.0418e+00
# Warning: expt average      12.G.N1-12.G.H1=-3.7000e-01 is smaller than minimum value in simulation 1.7209e+00
# Warning: expt average      11.U.N3-11.U.H3=-1.7500e+00 is smaller than minimum value in simulation 2.0383e+00
# theta: 1000
# RDC scaling factors 6.2675e-01, 6.2779e-01
# RDC scaling factor 6.1278e-01
# Warning: expt average        5.C.C6-5.C.H6=1.0990e+01 is larger than maximum value in simulation 6.7349e+00
# Warning: expt average        8.C.C6-8.C.H6=-3.6600e+00 is larger than maximum value in simulation -3.7324e+00
# Warning: expt average      11.U.C6-11.U.H6=9.0900e+00 is larger than maximum value in simulation 3.6095e+00
# Warning: expt average      13.C.C6-13.C.H6=2.8300e+00 is smaller than minimum value in simulation 3.6900e+00
# Warning: expt average      14.C.C6-14.C.H6=6.4300e+00 is larger than maximum value in simulation 4.2555e+00
# Warning: expt average        1.G.C8-1.G.H8=7.2900e+00 is larger than maximum value in simulation 1.4610e+00
# Warning: expt average        2.G.C8-2.G.H8=1.6300e+00 is smaller than minimum value in simulation 1.9816e+00
# Warning: expt average        9.G.C8-9.G.H8=3.4000e-01 is larger than maximum value in simulation -2.2605e-01
# Warning: expt average      10.G.C8-10.G.H8=1.1100e+01 is larger than maximum value in simulation 6.9464e+00
# Warning: expt average      1.G.C1'-1.G.H1'=-2.3900e+00 is larger than maximum value in simulation -8.4286e+00
# Warning: expt average      2.G.C1'-2.G.H1'=-3.2000e-01 is larger than maximum value in simulation -1.0067e+00
# Warning: expt average      3.C.C1'-3.C.H1'=6.3600e+00 is larger than maximum value in simulation 2.3631e+00
# Warning: expt average      8.C.C1'-8.C.H1'=-1.4660e+01 is smaller than minimum value in simulation -5.5338e+00
# Warning: expt average      9.G.C1'-9.G.H1'=-5.1400e+00 is smaller than minimum value in simulation -1.1158e+00
# Warning: expt average    12.G.C1'-12.G.H1'=6.7700e+00 is larger than maximum value in simulation 6.3867e+00
# Warning: expt average        9.G.N1-9.G.H1=-6.3200e+00 is smaller than minimum value in simulation 3.3250e-01
# Warning: expt average      10.G.N1-10.G.H1=-3.8800e+00 is smaller than minimum value in simulation 1.5332e+00
# Warning: expt average      11.U.N3-11.U.H3=-1.7500e+00 is smaller than minimum value in simulation -1.0539e+00
# theta: 1000
# RDC scaling factors 6.1278e-01, 6.1275e-01
# RDC scaling factor 1.0834e+00
# Warning: expt average      1.G.C4'-1.G.H4'=-5.7000e+00 is smaller than minimum value in simulation -5.1408e-01
# Warning: expt average        1.G.C8-1.G.H8=1.1770e+01 is larger than maximum value in simulation 5.0641e+00
# Warning: expt average      2.G.C1'-2.G.H1'=-1.0600e+00 is larger than maximum value in simulation -6.0836e+00
# Warning: expt average        3.C.C5-3.C.H5=1.0350e+01 is larger than maximum value in simulation 4.4725e+00
# Warning: expt average      3.C.C2'-3.C.H2'=-1.1820e+01 is smaller than minimum value in simulation -1.1787e+01
# Warning: expt average        4.A.C8-4.A.H8=1.0350e+01 is smaller than minimum value in simulation 1.0785e+01
# Warning: expt average        4.A.C2-4.A.H2=2.5300e+00 is smaller than minimum value in simulation 5.2546e+00
# Warning: expt average      4.A.C2'-4.A.H2'=-5.9700e+00 is larger than maximum value in simulation -6.7002e+00
# Warning: expt average      5.C.C1'-5.C.H1'=8.7800e+00 is larger than maximum value in simulation 5.1322e+00
# Warning: expt average        5.C.C6-5.C.H6=1.7560e+01 is larger than maximum value in simulation 1.3682e+01
# Warning: expt average      5.C.C2'-5.C.H2'=1.0010e+01 is larger than maximum value in simulation 4.0650e-01
# Warning: expt average      6.U.C1'-6.U.H1'=8.3500e+00 is smaller than minimum value in simulation 9.0988e+00
# Warning: expt average        6.U.C6-6.U.H6=6.1300e+00 is larger than maximum value in simulation 5.4257e+00
# Warning: expt average        6.U.C5-6.U.H5=1.5440e+01 is larger than maximum value in simulation 1.4056e+01
# Warning: expt average      7.U.C1'-7.U.H1'=2.6800e+00 is larger than maximum value in simulation -6.0779e-01
# Warning: expt average      7.U.C3'-7.U.H3'=2.7700e+00 is larger than maximum value in simulation 6.5590e-01
# Warning: expt average      8.C.C1'-8.C.H1'=-2.0540e+01 is smaller than minimum value in simulation -1.5652e+01
# Warning: expt average      9.G.C3'-9.G.H3'=1.0200e+01 is larger than maximum value in simulation 6.7008e+00
# Warning: expt average    10.G.C1'-10.G.H1'=-1.9610e+01 is smaller than minimum value in simulation -9.4008e+00
# Warning: expt average    10.G.C2'-10.G.H2'=5.7200e+00 is larger than maximum value in simulation -6.6576e+00
# Warning: expt average      11.U.C6-11.U.H6=1.2430e+01 is larger than maximum value in simulation 1.2106e+01
# Warning: expt average      11.U.C5-11.U.H5=1.0280e+01 is larger than maximum value in simulation 6.0918e+00
# Warning: expt average    12.G.C1'-12.G.H1'=9.2000e-01 is smaller than minimum value in simulation 7.5674e+00
# Warning: expt average      13.C.C6-13.C.H6=3.0700e+00 is smaller than minimum value in simulation 6.5720e+00
# Warning: expt average      13.C.C5-13.C.H5=1.4680e+01 is larger than maximum value in simulation 1.3315e+01
# Warning: expt average    13.C.C2'-13.C.H2'=-1.6320e+01 is smaller than minimum value in simulation -8.2752e+00
# Warning: expt average    14.C.C4'-14.C.H4'=1.4070e+01 is larger than maximum value in simulation 1.4040e+01
# Warning: expt average      14.C.C6-14.C.H6=9.5900e+00 is larger than maximum value in simulation 5.2349e+00
# theta: 1000
# RDC scaling factors 1.0834e+00, 1.0867e+00
# RDC scaling factor 1.0218e+00
# Warning: expt average      1.G.C4'-1.G.H4'=-5.7000e+00 is smaller than minimum value in simulation -4.0096e+00
# Warning: expt average        1.G.C8-1.G.H8=1.1770e+01 is larger than maximum value in simulation 2.4361e+00
# Warning: expt average      2.G.C1'-2.G.H1'=-1.0600e+00 is larger than maximum value in simulation -1.6786e+00
# Warning: expt average        3.C.C5-3.C.H5=1.0350e+01 is larger than maximum value in simulation 8.0674e+00
# Warning: expt average        5.C.C6-5.C.H6=1.7560e+01 is larger than maximum value in simulation 1.1230e+01
# Warning: expt average        5.C.C5-5.C.H5=7.2300e+00 is larger than maximum value in simulation 6.9923e+00
# Warning: expt average      5.C.C2'-5.C.H2'=1.0010e+01 is larger than maximum value in simulation 2.7750e+00
# Warning: expt average        6.U.C5-6.U.H5=1.5440e+01 is larger than maximum value in simulation 1.0845e+01
# Warning: expt average      7.U.C4'-7.U.H4'=-1.1640e+01 is smaller than minimum value in simulation -9.1929e+00
# Warning: expt average      7.U.C1'-7.U.H1'=2.6800e+00 is larger than maximum value in simulation 8.0444e-01
# Warning: expt average      7.U.C3'-7.U.H3'=2.7700e+00 is larger than maximum value in simulation -3.5142e+00
# Warning: expt average      8.C.C4'-8.C.H4'=3.8000e+00 is smaller than minimum value in simulation 8.0474e+00
# Warning: expt average      8.C.C1'-8.C.H1'=-2.0540e+01 is smaller than minimum value in simulation -9.2273e+00
# Warning: expt average        8.C.C6-8.C.H6=-4.8600e+00 is larger than maximum value in simulation -6.2235e+00
# Warning: expt average        8.C.C5-8.C.H5=-1.1690e+01 is smaller than minimum value in simulation -2.4695e+00
# Warning: expt average      9.G.C1'-9.G.H1'=-5.0700e+00 is smaller than minimum value in simulation -1.8604e+00
# Warning: expt average    10.G.C1'-10.G.H1'=-1.9610e+01 is smaller than minimum value in simulation -8.8827e+00
# Warning: expt average    10.G.C2'-10.G.H2'=5.7200e+00 is larger than maximum value in simulation -3.8482e+00
# Warning: expt average      11.U.C6-11.U.H6=1.2430e+01 is larger than maximum value in simulation 6.0187e+00
# Warning: expt average      11.U.C5-11.U.H5=1.0280e+01 is larger than maximum value in simulation 1.0266e+01
# Warning: expt average      12.G.C8-12.G.H8=1.2210e+01 is larger than maximum value in simulation 9.9205e+00
# Warning: expt average      13.C.C6-13.C.H6=3.0700e+00 is smaller than minimum value in simulation 6.1529e+00
# Warning: expt average      13.C.C5-13.C.H5=1.4680e+01 is larger than maximum value in simulation 1.1435e+01
# Warning: expt average    13.C.C2'-13.C.H2'=-1.6320e+01 is smaller than minimum value in simulation -3.8459e+00
# Warning: expt average    14.C.C4'-14.C.H4'=1.4070e+01 is larger than maximum value in simulation 1.0318e+01
# Warning: expt average      14.C.C6-14.C.H6=9.5900e+00 is larger than maximum value in simulation 7.0957e+00
# theta: 1000
# RDC scaling factors 1.0218e+00, 1.0285e+00
# RDC scaling factor 5.9977e-01
# Warning: expt average      1.G.C1'-1.G.H1'=2.9800e+00 is larger than maximum value in simulation -5.0645e+00
# Warning: expt average      2.G.C4'-2.G.H4'=-4.6000e-01 is smaller than minimum value in simulation 4.8048e-01
# Warning: expt average      4.A.C1'-4.A.H1'=7.2200e+00 is larger than maximum value in simulation 5.3154e+00
# Warning: expt average      6.U.C2'-6.U.H2'=1.2820e+01 is larger than maximum value in simulation 3.2051e+00
# Warning: expt average        7.U.C6-7.U.H6=7.3000e+00 is larger than maximum value in simulation 6.8945e+00
# Warning: expt average        7.U.C5-7.U.H5=4.4600e+00 is larger than maximum value in simulation 1.7846e+00
# Warning: expt average      9.G.C2'-9.G.H2'=2.6400e+00 is larger than maximum value in simulation -2.6282e-01
# Warning: expt average      10.G.C8-10.G.H8=1.4760e+01 is larger than maximum value in simulation 5.7776e+00
# Warning: expt average    11.U.C3'-11.U.H3'=-4.7000e+00 is smaller than minimum value in simulation -3.6937e+00
# Warning: expt average      14.C.C5-14.C.H5=1.0600e+00 is smaller than minimum value in simulation 4.7523e+00
# Warning: expt average      6.U.C4'-6.U.H4'=1.5090e+01 is larger than maximum value in simulation 5.4477e+00
# Warning: expt average    10.G.C3'-10.G.H3'=-1.1370e+01 is smaller than minimum value in simulation -6.3218e+00
# Warning: expt average      7.U.C2'-7.U.H2'=-1.2900e+00 is larger than maximum value in simulation -5.2552e+00
# theta: 1000
# RDC scaling factors 5.9977e-01, 6.0018e-01
# RDC scaling factor 4.5234e-01
# Warning: expt average      1.G.C1'-1.G.H1'=2.9800e+00 is larger than maximum value in simulation -6.2218e+00
# Warning: expt average      1.G.C2'-1.G.H2'=-2.6600e+00 is smaller than minimum value in simulation -1.6197e+00
# Warning: expt average      4.A.C1'-4.A.H1'=7.2200e+00 is larger than maximum value in simulation 5.2511e+00
# Warning: expt average      6.U.C2'-6.U.H2'=1.2820e+01 is larger than maximum value in simulation 3.4818e+00
# Warning: expt average        7.U.C6-7.U.H6=7.3000e+00 is larger than maximum value in simulation 4.0218e+00
# Warning: expt average      9.G.C2'-9.G.H2'=2.6400e+00 is larger than maximum value in simulation -1.1593e+00
# Warning: expt average      10.G.C8-10.G.H8=1.4760e+01 is larger than maximum value in simulation 5.1277e+00
# Warning: expt average    11.U.C3'-11.U.H3'=-4.7000e+00 is smaller than minimum value in simulation -4.6038e+00
# Warning: expt average      6.U.C4'-6.U.H4'=1.5090e+01 is larger than maximum value in simulation 3.1917e-01
# Warning: expt average    10.G.C3'-10.G.H3'=-1.1370e+01 is smaller than minimum value in simulation -4.3100e+00
# Warning: expt average      7.U.C2'-7.U.H2'=-1.2900e+00 is larger than maximum value in simulation -2.5750e+00
# theta: 1000
# RDC scaling factors 4.5234e-01, 4.5336e-01
# RDC scaling factor 6.0500e-03
# Warning: expt average             1Gua1H2'=6.5576e-01 is larger than maximum value in simulation 3.3145e-01
# Warning: expt average              1GuaH3'=1.9820e-01 is smaller than minimum value in simulation 3.3574e-01
# Warning: expt average              1GuaH4'=1.2939e+00 is larger than maximum value in simulation 9.3491e-01
# Warning: expt average               1GuaH8=1.7188e+00 is larger than maximum value in simulation 5.9340e-01
# Warning: expt average               2GuaH1=1.4576e-01 is smaller than minimum value in simulation 1.5393e-01
# Warning: expt average              2GuaH4'=5.3169e-01 is smaller than minimum value in simulation 6.4681e-01
# Warning: expt average             2Gua2H5'=9.7922e-01 is larger than maximum value in simulation 9.3054e-01
# Warning: expt average               2GuaH8=5.5815e-01 is larger than maximum value in simulation 2.8459e-01
# Warning: expt average              3CytH41=6.9691e-02 is smaller than minimum value in simulation 1.8705e-01
# Warning: expt average              3CytH42=7.7233e-02 is smaller than minimum value in simulation 3.2283e-01
# Warning: expt average               3CytH5=1.3619e-01 is smaller than minimum value in simulation 2.4462e-01
# Warning: expt average              4AdeH1'=2.4714e-01 is smaller than minimum value in simulation 3.5505e-01
# Warning: expt average               4AdeH2=1.2344e-01 is smaller than minimum value in simulation 1.8934e-01
# Warning: expt average             4Ade1H2'=1.9599e-01 is smaller than minimum value in simulation 2.2153e-01
# Warning: expt average              4AdeH4'=5.0286e-01 is smaller than minimum value in simulation 6.5796e-01
# Warning: expt average              4AdeH61=7.4677e-03 is smaller than minimum value in simulation 1.8234e-01
# Warning: expt average              4AdeH62=-4.8206e-03 is smaller than minimum value in simulation 2.3334e-01
# Warning: expt average               4AdeH8=1.5958e-01 is smaller than minimum value in simulation 2.0369e-01
# Warning: expt average              5CytH1'=2.1959e-01 is smaller than minimum value in simulation 2.7228e-01
# Warning: expt average             5Cyt1H2'=1.9809e-01 is smaller than minimum value in simulation 2.0839e-01
# Warning: expt average              5CytH3'=1.9966e-01 is smaller than minimum value in simulation 2.5809e-01
# Warning: expt average              5CytH41=9.0368e-02 is smaller than minimum value in simulation 2.5432e-01
# Warning: expt average              5CytH42=9.1948e-02 is smaller than minimum value in simulation 4.1964e-01
# Warning: expt average               5CytH5=1.2417e-01 is smaller than minimum value in simulation 2.6912e-01
# Warning: expt average               5CytH6=1.4657e-01 is smaller than minimum value in simulation 2.0803e-01
# Warning: expt average              6UraH1'=1.2072e-01 is smaller than minimum value in simulation 1.4060e-01
# Warning: expt average             6Ura1H2'=1.5312e-01 is smaller than minimum value in simulation 1.7490e-01
# Warning: expt average               6UraH3=2.3618e+00 is larger than maximum value in simulation 3.1610e-01
# Warning: expt average              6UraH3'=2.1742e-01 is smaller than minimum value in simulation 2.5038e-01
# Warning: expt average               6UraH5=1.7169e-01 is smaller than minimum value in simulation 2.8587e-01
# Warning: expt average              7UraH1'=8.2627e-01 is larger than maximum value in simulation 7.1750e-01
# Warning: expt average             7Ura1H2'=3.2446e-01 is larger than maximum value in simulation 2.4390e-01
# Warning: expt average             7Ura1H5'=6.3890e-01 is smaller than minimum value in simulation 6.6991e-01
# Warning: expt average               7UraH6=4.0640e-01 is larger than maximum value in simulation 3.7355e-01
# Warning: expt average             8Cyt1H2'=3.9996e-01 is smaller than minimum value in simulation 4.4855e-01
# Warning: expt average              8CytH3'=4.4405e-01 is smaller than minimum value in simulation 4.5583e-01
# Warning: expt average              8CytH4'=3.0542e-01 is larger than maximum value in simulation 3.0383e-01
# Warning: expt average              8CytH41=4.6025e-01 is smaller than minimum value in simulation 5.2104e-01
# Warning: expt average              8CytH42=6.5893e-01 is larger than maximum value in simulation 6.5526e-01
# Warning: expt average               9GuaH1=4.4689e-01 is larger than maximum value in simulation 1.4877e-01
# Warning: expt average              9GuaH1'=7.9905e-01 is larger than maximum value in simulation 7.5854e-01
# Warning: expt average              9GuaH3'=3.1539e-01 is larger than maximum value in simulation 3.0472e-01
# Warning: expt average             9Gua1H5'=4.9528e-01 is smaller than minimum value in simulation 6.2226e-01
# Warning: expt average             9Gua2H5'=5.3851e-01 is larger than maximum value in simulation 4.2616e-01
# Warning: expt average               9GuaH8=9.6014e-01 is larger than maximum value in simulation 9.4641e-01
# Warning: expt average              10GuaH1=6.7816e-02 is smaller than minimum value in simulation 1.3333e-01
# Warning: expt average             10GuaH1'=1.7740e-01 is smaller than minimum value in simulation 1.8581e-01
# Warning: expt average            10Gua1H2'=1.8282e-01 is smaller than minimum value in simulation 2.0559e-01
# Warning: expt average             10GuaH21=1.0203e-01 is smaller than minimum value in simulation 2.0124e-01
# Warning: expt average             10GuaH22=1.1137e-01 is smaller than minimum value in simulation 1.4818e-01
# Warning: expt average             10GuaH3'=2.2555e-01 is smaller than minimum value in simulation 2.7042e-01
# Warning: expt average            10Gua1H5'=3.6992e-01 is smaller than minimum value in simulation 4.9030e-01
# Warning: expt average            10Gua2H5'=3.5054e-01 is smaller than minimum value in simulation 4.4849e-01
# Warning: expt average              10GuaH8=1.8596e-01 is smaller than minimum value in simulation 2.4095e-01
# Warning: expt average              11UraH3=7.4565e-02 is smaller than minimum value in simulation 1.3338e-01
# Warning: expt average             11UraH3'=2.5973e-01 is smaller than minimum value in simulation 2.8454e-01
# Warning: expt average              11UraH5=1.0542e-01 is smaller than minimum value in simulation 2.9781e-01
# Warning: expt average              11UraH6=1.3466e-01 is smaller than minimum value in simulation 2.0052e-01
# Warning: expt average              12GuaH1=5.8687e-02 is smaller than minimum value in simulation 1.3413e-01
# Warning: expt average             12GuaH1'=2.7069e-01 is smaller than minimum value in simulation 3.2310e-01
# Warning: expt average             12GuaH4'=4.9210e-01 is smaller than minimum value in simulation 5.7974e-01
# Warning: expt average              12GuaH8=1.7404e-01 is smaller than minimum value in simulation 2.1621e-01
# Warning: expt average             13CytH1'=2.5983e-01 is smaller than minimum value in simulation 2.6772e-01
# Warning: expt average            13Cyt1H2'=3.0561e-01 is larger than maximum value in simulation 2.7345e-01
# Warning: expt average             13CytH4'=4.0486e-01 is smaller than minimum value in simulation 5.2940e-01
# Warning: expt average             13CytH41=1.6219e-01 is smaller than minimum value in simulation 2.4028e-01
# Warning: expt average             13CytH42=1.5849e-01 is smaller than minimum value in simulation 4.0345e-01
# Warning: expt average              13CytH5=1.6462e-01 is smaller than minimum value in simulation 3.0119e-01
# Warning: expt average            13Cyt1H5'=5.0428e-01 is smaller than minimum value in simulation 5.9244e-01
# Warning: expt average              13CytH6=1.8168e-01 is smaller than minimum value in simulation 2.2410e-01
# Warning: expt average            14Cyt1H2'=3.4522e-01 is smaller than minimum value in simulation 6.9382e-01
# Warning: expt average             14CytH3'=1.1184e+00 is larger than maximum value in simulation 9.1483e-01
# Warning: expt average             14CytH41=1.2781e+00 is larger than maximum value in simulation 5.8126e-01
# Warning: expt average             14CytH42=1.0972e+00 is larger than maximum value in simulation 9.2257e-01
# Warning: expt average              14CytH6=5.0637e-01 is larger than maximum value in simulation 4.2664e-01
# theta: 1000
# RDC scaling factors 6.0500e-03, 6.1238e-03
# RDC scaling factor 6.0430e-03
# Warning: expt average             1Gua1H2'=6.5576e-01 is larger than maximum value in simulation 3.4403e-01
# Warning: expt average              1GuaH3'=1.9820e-01 is smaller than minimum value in simulation 3.1613e-01
# Warning: expt average              1GuaH4'=1.2939e+00 is larger than maximum value in simulation 1.0671e+00
# Warning: expt average               1GuaH8=1.7188e+00 is larger than maximum value in simulation 5.5994e-01
# Warning: expt average              2GuaH4'=5.3169e-01 is smaller than minimum value in simulation 5.6243e-01
# Warning: expt average             2Gua2H5'=9.7922e-01 is larger than maximum value in simulation 7.9232e-01
# Warning: expt average               2GuaH8=5.5815e-01 is larger than maximum value in simulation 2.9898e-01
# Warning: expt average              3CytH41=6.9691e-02 is smaller than minimum value in simulation 1.3945e-01
# Warning: expt average              3CytH42=7.7233e-02 is smaller than minimum value in simulation 1.6669e-01
# Warning: expt average               3CytH5=1.3619e-01 is smaller than minimum value in simulation 1.6401e-01
# Warning: expt average              4AdeH1'=2.4714e-01 is smaller than minimum value in simulation 2.8615e-01
# Warning: expt average               4AdeH2=1.2344e-01 is smaller than minimum value in simulation 1.3311e-01
# Warning: expt average             4Ade1H2'=1.9599e-01 is smaller than minimum value in simulation 2.2824e-01
# Warning: expt average              4AdeH61=7.4677e-03 is smaller than minimum value in simulation 2.7393e-01
# Warning: expt average              4AdeH62=-4.8206e-03 is smaller than minimum value in simulation 3.5810e-01
# Warning: expt average               4AdeH8=1.5958e-01 is smaller than minimum value in simulation 2.0176e-01
# Warning: expt average              5CytH1'=2.1959e-01 is smaller than minimum value in simulation 3.0253e-01
# Warning: expt average             5Cyt1H2'=1.9809e-01 is smaller than minimum value in simulation 2.1391e-01
# Warning: expt average              5CytH3'=1.9966e-01 is smaller than minimum value in simulation 2.4901e-01
# Warning: expt average              5CytH41=9.0368e-02 is smaller than minimum value in simulation 1.5434e-01
# Warning: expt average              5CytH42=9.1948e-02 is smaller than minimum value in simulation 2.0469e-01
# Warning: expt average               5CytH5=1.2417e-01 is smaller than minimum value in simulation 2.0080e-01
# Warning: expt average               5CytH6=1.4657e-01 is smaller than minimum value in simulation 1.8970e-01
# Warning: expt average              6UraH1'=1.2072e-01 is smaller than minimum value in simulation 1.6426e-01
# Warning: expt average             6Ura1H2'=1.5312e-01 is smaller than minimum value in simulation 1.7574e-01
# Warning: expt average               6UraH3=2.3618e+00 is larger than maximum value in simulation 3.2279e-01
# Warning: expt average              6UraH3'=2.1742e-01 is smaller than minimum value in simulation 2.6018e-01
# Warning: expt average              6UraH4'=2.8013e-01 is smaller than minimum value in simulation 2.9574e-01
# Warning: expt average               6UraH5=1.7169e-01 is smaller than minimum value in simulation 3.2691e-01
# Warning: expt average             7Ura1H2'=3.2446e-01 is larger than maximum value in simulation 2.4977e-01
# Warning: expt average              8CytH4'=3.0542e-01 is larger than maximum value in simulation 2.9704e-01
# Warning: expt average              8CytH41=4.6025e-01 is smaller than minimum value in simulation 5.3794e-01
# Warning: expt average               9GuaH1=4.4689e-01 is larger than maximum value in simulation 1.6039e-01
# Warning: expt average             9Gua1H2'=3.6607e-01 is larger than maximum value in simulation 3.3261e-01
# Warning: expt average              9GuaH3'=3.1539e-01 is larger than maximum value in simulation 2.7841e-01
# Warning: expt average             9Gua1H5'=4.9528e-01 is smaller than minimum value in simulation 5.4926e-01
# Warning: expt average              10GuaH1=6.7816e-02 is smaller than minimum value in simulation 1.3292e-01
# Warning: expt average             10GuaH1'=1.7740e-01 is smaller than minimum value in simulation 2.1406e-01
# Warning: expt average            10Gua1H2'=1.8282e-01 is smaller than minimum value in simulation 2.1923e-01
# Warning: expt average             10GuaH21=1.0203e-01 is smaller than minimum value in simulation 1.5928e-01
# Warning: expt average             10GuaH22=1.1137e-01 is smaller than minimum value in simulation 2.6041e-01
# Warning: expt average             10GuaH3'=2.2555e-01 is smaller than minimum value in simulation 2.8206e-01
# Warning: expt average            10Gua1H5'=3.6992e-01 is smaller than minimum value in simulation 5.1291e-01
# Warning: expt average            10Gua2H5'=3.5054e-01 is smaller than minimum value in simulation 4.7016e-01
# Warning: expt average              10GuaH8=1.8596e-01 is smaller than minimum value in simulation 2.6446e-01
# Warning: expt average              11UraH3=7.4565e-02 is smaller than minimum value in simulation 1.3887e-01
# Warning: expt average              11UraH5=1.0542e-01 is smaller than minimum value in simulation 3.4439e-01
# Warning: expt average              11UraH6=1.3466e-01 is smaller than minimum value in simulation 1.9633e-01
# Warning: expt average              12GuaH1=5.8687e-02 is smaller than minimum value in simulation 1.2876e-01
# Warning: expt average              12GuaH8=1.7404e-01 is smaller than minimum value in simulation 1.8582e-01
# Warning: expt average             13CytH1'=2.5983e-01 is smaller than minimum value in simulation 3.3543e-01
# Warning: expt average             13CytH4'=4.0486e-01 is smaller than minimum value in simulation 5.9638e-01
# Warning: expt average             13CytH41=1.6219e-01 is smaller than minimum value in simulation 2.0557e-01
# Warning: expt average             13CytH42=1.5849e-01 is smaller than minimum value in simulation 3.3451e-01
# Warning: expt average              13CytH5=1.6462e-01 is smaller than minimum value in simulation 2.3517e-01
# Warning: expt average            13Cyt1H5'=5.0428e-01 is smaller than minimum value in simulation 5.5214e-01
# Warning: expt average              13CytH6=1.8168e-01 is smaller than minimum value in simulation 2.0089e-01
# Warning: expt average             14CytH1'=5.7951e-01 is larger than maximum value in simulation 5.7322e-01
# Warning: expt average            14Cyt1H2'=3.4522e-01 is smaller than minimum value in simulation 6.1878e-01
# Warning: expt average             14CytH3'=1.1184e+00 is larger than maximum value in simulation 1.0732e+00
# Warning: expt average             14CytH41=1.2781e+00 is larger than maximum value in simulation 6.6436e-01
# Warning: expt average             14CytH42=1.0972e+00 is larger than maximum value in simulation 9.0540e-01
# theta: 1000
# RDC scaling factors 6.0430e-03, 6.1257e-03
```

Now we collect all the data from the different ensembles

In [8]:

```
import numpy as np
import os
import subprocess
from scipy import stats


def calc_stuff(filename):
    if("RDC" in filename or "sPRE" in filename):
        data_tmp = np.array([[float(x) for x in line.split()[1:]] for line in open(filename) if "#" not in line])
        bb,pb = stats.spearmanr(data_tmp[:,0],data_tmp[:,2])
        aa,pb = stats.spearmanr(data_tmp[:,0],data_tmp[:,3])
    else:
        line = subprocess.check_output(['tail', '-1', ff])
        return float(line.split()[-2]),float(line.split()[-1])
    return bb,aa

ensembles = ["set_A1","2koc","6by5"]
theta=30
data = []
stds = []
for i in range(len(lista2)):
    for j in range(1,len(lista2[i])):
        data_tmp = []
        stds_tmp = []
        for o,ens in enumerate(ensembles):
            if("set" in ens):
                chi2_tmp_before = []
                chi2_tmp_after = []
                for k in range(4):
                    ff = "data/comparison/%s/rw_%s_%s_theta_%d_rep_%d.stats.dat" % (ens,lista2[i][0],lista2[i][j],theta,k)
                    before, after = calc_stuff(ff)
                    chi2_tmp_before.append(before)
                    chi2_tmp_after.append(after)
                avg_b = np.average(chi2_tmp_before)
                avg_a = np.average(chi2_tmp_after)
                std_b = np.std(chi2_tmp_before,ddof=1)
                std_a = np.std(chi2_tmp_after,ddof=1)
                #print(ens,avg_b,avg_a)
                if(ens==ensembles[0]):
                    data_tmp.append(avg_b)
                    stds_tmp.append(std_b)
                data_tmp.append(avg_a)
                stds_tmp.append(std_a)
                
            else:
                ff = "data/comparison/%s/%s_%s.stats.dat" % (lista2[i][0],ens,lista2[i][j])
                before, after = calc_stuff(ff)
                data_tmp.append(before)
        data.append(data_tmp)
        stds.append(stds_tmp)
data = np.array(data)
stds = np.array(stds)
print(data)
```

```
[[1.904725   0.621575   0.9997     0.126     ]
 [0.592925   0.43285    0.5397     0.3019    ]
 [0.6391     0.349125   0.32       0.0613    ]
 [0.4296     0.533125   0.4934     0.6294    ]
 [3.560475   4.09325    4.2772     4.4486    ]
 [0.76087691 0.79921188 0.69318182 0.59420821]
 [0.59757579 0.62394352 0.78216419 0.59345109]
 [0.62747253 0.65384615 0.56043956 0.53846154]
 [0.73784578 0.72282132 0.70901519 0.6944899 ]]
```

And then make a barplot. Most of the commands below are cosmetics

In [9]:

```
import matplotlib.pyplot as plt
import seaborn as sns
import matplotlib as mpl
label_size = 7.5
mpl.rcParams['xtick.labelsize'] = label_size 
mpl.rcParams['ytick.labelsize'] = label_size 
fig, ax = plt.subplots(3,3,figsize=(7.5,7.5))
plt.subplots_adjust(left=0.15, bottom=0.15, right=0.98, top=0.92, wspace=0.3, hspace=0.4)
cols = ["black","medium blue","pumpkin","brick red","black","white"]
cols1 = ["black","medium blue","pumpkin","brick red","leaf","black","white"]

cp = sns.xkcd_palette(cols)
cps = sns.xkcd_palette(cols1)


wi=0.61

suptitle = ["DATASET A","DATASET B","DATASET C","DATASET D"]
titles = [["bidir-eNOE","unidir-eNOE","gn-eNOE"],\
          ["NOE","3J couplings","RDC"],["RDC1","RDC2","sPRE"]]
labs = ["MD","MD+eNOE","2KOC","6BY5"]

mm = [2,1.0,0.6,0.8,5,1,1,1,1]
xx = [0.2,1.2,2.7,3.7]

l=0
for k in range(3):
    for i in range(3):

        for m in range(data.shape[1]):
            if(m<2):
                ax[k,i].bar(xx[m],data[l,m],width=wi,color=cp[m],alpha=0.7,edgecolor=cps[m],yerr=stds[l,m],label=labs[m])
            else:
                ax[k,i].bar(xx[m],data[l,m],width=wi,color=cp[m],alpha=0.7,edgecolor=cps[m],label=labs[m])
        #print(data[l,:])    
        ax[k,i].set_xticks([])
        ax[k,i].set_xlabel(titles[k][i],fontsize=9)
        yy = np.round(np.linspace(0,mm[l] ,3), 2)
        ax[k,i].set_yticks(yy)
        ax[k,i].spines['right'].set_visible(False)
        ax[k,i].spines['top'].set_visible(False)
        if(i==0):
            if(k==2):
                ax[k,i].set_ylabel(r"$\rho $")
            else:
                ax[k,i].set_ylabel("$\chi^2$")
        if(i==2 and k==1):
                ax[k,i].set_ylabel(r"$\rho $")

        if(i==1 and k != 2):
            ax[k,i].set_title(suptitle[k],fontsize=10)
        if(k==1 and i==0):
            ax[k,i].legend(ncol=4,loc="lower left",bbox_to_anchor=(0.5,-1.8),fontsize=10)
        if(k==2):
            if(i==2):
                ax[k,i].set_title(suptitle[k+1],fontsize=10)
            if(i==1):
                ax[k,i].text(-1,1.1,suptitle[k],fontsize=10,ha="center")
        l += 1

plt.savefig("figures/comparison_final.pdf")
plt.savefig("figures/comparison_final.png",dpi=500)
plt.show()
plt.close()
```

In [ ]:
