## Supplementary material for "Integrating NMR and Simulations Reveals Motions in the UUCG Tetraloop": SI4 (from another version of this preprint: version 2, 2019-09-11)

SI4\_compare\_full


### Compare simulations with experiments¶

We here compare back-calculated and experimental data.

You need to execute the following notebooks in order to run successfully the code below:

- create figure 2

We first calculate RMSE, chi2, pearson correlation coefficient and number of violations

In [1]:

```
import sys


lista2 = [ ["set_A1","eNOE","eNOE_unidir","gn_eNOE"],\
        ["set_B","NOE","J3","RDC"],\
        ["set_C","RDC1","RDC2"],\
        ["set_D","sPRE"]]
```

Now we collect all the data from the different ensembles

In [2]:

```
import numpy as np
import os
import subprocess
from scipy import stats


def calc_stuff(filename):
    dd = np.array([[float(x) for x in line.split()[1:]] for line in open(filename) if "#" not in line])
    return dd

ensembles = ["set_A1","2koc","6by5"]
theta=30
for i in range(len(lista2)):
    for j in range(1,len(lista2[i])):
        data_tmp = []
        stds_tmp = []
        #fig, ax = plt.subplots(3,1,figsize=(6,18))
        for o,ens in enumerate(ensembles):
            d = []
            if(ens==ensembles[0]):
                for k in range(4):
                    ff = "data/comparison/%s/rw_%s_%s_theta_%d_rep_%d.stats.dat" % (ens,lista2[i][0],lista2[i][j],theta,k)
                    d.append(calc_stuff(ff))
            else:
                ff = "data/comparison/%s/%s_%s.stats.dat" % (lista2[i][0],ens,lista2[i][j])
                d.append(calc_stuff(ff))
            d = np.array(d)
            print(d.shape)
            if(d.shape[2]==4):
                ee = d[0,:,0]
                ees = d[0,:,1]
                bb = d[:,:,3]
            else:
                ee = d[0,:,4]
                ees = d[0,:,5]
                bb = d[:,:,7]
   
            aa = np.average(bb,axis=0)
            ss = np.std(bb,axis=0,ddof=1)
            #ax[o].errorbar(ee,aa,xerr=ees,yerr=ss,fmt="o")
            #ax[o].set_title("%s %s" % (lista2[i][j],ens)) 
            #mma = np.max([ax[o].get_ylim()[1], ax[o].get_xlim()[1]])
            #mmi = np.min([ax[o].get_ylim()[0], ax[o].get_xlim()[0]])
            #ax[o].set_ylim(mmi,mma)
            #ax[o].set_xlim(mmi,mma)
            if(o!=10):
                diff = (bb-ee)**2
                if(lista2[i][j]=="gn_eNOE"):
                    #print(bb.shape,diff.shape,ee.shape)
                    for w in range(diff.shape[0]):
                        diff[w,np.where((bb[w,:]-ee)<0.0)] = 0.0
                rmse = np.sqrt(np.average(diff,axis=1))
                chi2 = np.average(diff/ees**2,axis=1)
                viol = [sum((np.where(diff/ees**2>1)[0])==y) for y in range(diff.shape[0])]
                rho = [ stats.spearmanr(bb[y,:],ee)[0] for y in range(diff.shape[0])]
                print("%11s %7s" % (lista2[i][j],ens),end=" ")
                print("RMSE: %4.2f+/-%-4.2f " % (np.average(rmse), np.std(rmse,ddof=1)), end=" ")
                print("Chi2: %4.2f+/-%-4.2f " % (np.average(chi2), np.std(chi2,ddof=1)), end=" ")
                print("rho: %4.2f+/-%-4.2f " % (np.average(rho), np.std(rho,ddof=1)), end=" ")
                print("viol: %3d+/-%-3.0f " % (np.average(viol), np.std(viol,ddof=1)))
                
                #print()
        #fig.show()
        #plt.close()
        print("")
```

```
(4, 63, 8)
       eNOE  set_A1 RMSE: 0.33+/-0.04  Chi2: 0.62+/-0.15  rho: 0.94+/-0.00  viol:  11+/-3   
(1, 63, 8)
       eNOE    2koc RMSE: 0.46+/-nan   Chi2: 1.00+/-nan   rho: 0.91+/-nan   viol:  14+/-nan 
(1, 63, 8)
       eNOE    6by5 RMSE: 0.14+/-nan   Chi2: 0.13+/-nan   rho: 0.99+/-nan   viol:   1+/-nan 

(4, 177, 8)
eNOE_unidir  set_A1 RMSE: 0.39+/-0.02  Chi2: 0.43+/-0.04  rho: 0.91+/-0.00  viol:  23+/-4   
(1, 177, 8)
eNOE_unidir    2koc RMSE: 0.44+/-nan   Chi2: 0.54+/-nan   rho: 0.92+/-nan   viol:  30+/-nan 
(1, 177, 8)
eNOE_unidir    6by5 RMSE: 0.34+/-nan   Chi2: 0.30+/-nan   rho: 0.94+/-nan   viol:   3+/-nan 

(4, 84, 8)
    gn_eNOE  set_A1 RMSE: 0.52+/-0.07  Chi2: 1.17+/-0.27  rho: 0.89+/-0.00  viol:  10+/-2   
(1, 84, 8)
    gn_eNOE    2koc RMSE: 0.97+/-nan   Chi2: 3.34+/-nan   rho: 0.90+/-nan   viol:  14+/-nan 
(1, 84, 8)
    gn_eNOE    6by5 RMSE: 0.84+/-nan   Chi2: 2.47+/-nan   rho: 0.84+/-nan   viol:   9+/-nan 

(4, 251, 8)
        NOE  set_A1 RMSE: 0.72+/-0.01  Chi2: 0.53+/-0.01  rho: 0.71+/-0.00  viol:  29+/-1   
(1, 251, 8)
        NOE    2koc RMSE: 0.74+/-nan   Chi2: 0.49+/-nan   rho: 0.74+/-nan   viol:  30+/-nan 
(1, 251, 8)
        NOE    6by5 RMSE: 0.76+/-nan   Chi2: 0.63+/-nan   rho: 0.72+/-nan   viol:  32+/-nan 

(4, 96, 4)
         J3  set_A1 RMSE: 3.03+/-0.07  Chi2: 4.09+/-0.19  rho: 0.73+/-0.01  viol:  44+/-1   
(1, 96, 4)
         J3    2koc RMSE: 3.10+/-nan   Chi2: 4.27+/-nan   rho: 0.65+/-nan   viol:  53+/-nan 
(1, 96, 4)
         J3    6by5 RMSE: 3.15+/-nan   Chi2: 4.42+/-nan   rho: 0.64+/-nan   viol:  56+/-nan 

(4, 32, 4)
        RDC  set_A1 RMSE: 4.03+/-0.21  Chi2: 16.27+/-1.76  rho: 0.80+/-0.02  viol:  25+/-1   
(1, 32, 4)
        RDC    2koc RMSE: 3.75+/-nan   Chi2: 14.07+/-nan   rho: 0.69+/-nan   viol:  29+/-nan 
(1, 32, 4)
        RDC    6by5 RMSE: 4.46+/-nan   Chi2: 19.93+/-nan   rho: 0.60+/-nan   viol:  28+/-nan 

(4, 39, 4)
       RDC1  set_A1 RMSE: 8.12+/-0.35  Chi2: 33.68+/-2.98  rho: 0.62+/-0.03  viol:  34+/-1   
(1, 39, 4)
       RDC1    2koc RMSE: 6.09+/-nan   Chi2: 18.90+/-nan   rho: 0.78+/-nan   viol:  33+/-nan 
(1, 39, 4)
       RDC1    6by5 RMSE: 8.08+/-nan   Chi2: 33.27+/-nan   rho: 0.60+/-nan   viol:  37+/-nan 

(4, 14, 4)
       RDC2  set_A1 RMSE: 7.22+/-0.08  Chi2: 10.77+/-0.24  rho: 0.65+/-0.03  viol:  11+/-1   
(1, 14, 4)
       RDC2    2koc RMSE: 6.65+/-nan   Chi2: 9.15+/-nan   rho: 0.56+/-nan   viol:  12+/-nan 
(1, 14, 4)
       RDC2    6by5 RMSE: 7.47+/-nan   Chi2: 11.53+/-nan   rho: 0.54+/-nan   viol:  13+/-nan 

(4, 91, 4)
       sPRE  set_A1 RMSE: 0.30+/-0.00  Chi2: 190.66+/-2.42  rho: 0.72+/-0.01  viol:  83+/-2   
(1, 91, 4)
       sPRE    2koc RMSE: 0.32+/-nan   Chi2: 174.79+/-nan   rho: 0.73+/-nan   viol:  87+/-nan 
(1, 91, 4)
       sPRE    6by5 RMSE: 0.31+/-nan   Chi2: 144.66+/-nan   rho: 0.72+/-nan   viol:  84+/-nan
```

```
/anaconda3/lib/python3.7/site-packages/numpy/core/_methods.py:140: RuntimeWarning: Degrees of freedom <= 0 for slice
  keepdims=keepdims)
/anaconda3/lib/python3.7/site-packages/numpy/core/_methods.py:130: RuntimeWarning: invalid value encountered in true_divide
  ret, rcount, out=ret, casting='unsafe', subok=False)
/anaconda3/lib/python3.7/site-packages/numpy/core/_methods.py:132: RuntimeWarning: invalid value encountered in double_scalars
  ret = ret.dtype.type(ret / rcount)
```

Now we plot the experimental data against the back-calculated averages. Large violations in MD are listed, sorted by the deviation (relative to the error)

In [4]:

```
import numpy as np
import os
import subprocess
from scipy import stats
import matplotlib.pyplot as plt

def calc_stuff(filename):
    dd = np.array([[float(x) for x in line.split()[1:]] for line in open(filename) if "#" not in line])
    lab = [line.split()[0] for line in open(filename) if "#" not in line]

    return lab,dd

ensembles = ["set_A1","2koc","6by5"]
theta=30
for i in range(len(lista2)):
    for j in range(1,len(lista2[i])):
        data_tmp = []
        stds_tmp = []
        #fig, ax = plt.subplots(1,1,figsize=(6,6))
        fig, ax = plt.subplots(1,1,figsize=(6,6))
        for o,ens in enumerate(ensembles):
            d = []
            if(ens==ensembles[0]):
                for k in range(4):
                    ff = "data/comparison/%s/rw_%s_%s_theta_%d_rep_%d.stats.dat" % (ens,lista2[i][0],lista2[i][j],theta,k)
                    lab, dd = calc_stuff(ff)
                    d.append(dd)
            else:
                ff = "data/comparison/%s/%s_%s.stats.dat" % (lista2[i][0],ens,lista2[i][j])
                lab, dd = calc_stuff(ff)
                d.append(dd)
            d = np.array(d)
            if(d.shape[2]==4):
                ee = d[0,:,0]
                ees = d[0,:,1]
                bb = d[:,:,3]
            else:
                ee = d[0,:,4]
                ees = d[0,:,5]
                bb = d[:,:,7]
   
            # plot only the MD ensemble
            if(o!=0): continue
            aa = np.average(bb,axis=0)
            ss = np.std(bb,axis=0,ddof=1)
            ax.errorbar(ee,aa,xerr=ees,yerr=ss,fmt="o")
            ax.set_title("%s %s" % (lista2[i][j],ens)) 
            mma = np.max([ax.get_ylim()[1], ax.get_xlim()[1]])
            mmi = np.min([ax.get_ylim()[0], ax.get_xlim()[0]])
            ax.set_ylim(mmi,mma)
            ax.set_xlim(mmi,mma)
            ax.set_xlabel("Experimental NOE [$\AA$]")
            ax.set_ylabel("Simulation NOE [$\AA$]")
            ax.plot([0,7],[0,7],c='k',ls="--")
            diff = (aa-ee)**2
            print("List of large violations %s" % lista2[i][j])
            print("%15s %5s %5s %5s " % ("Label","exp","calc","deviation"))
            if(lista2[i][j]=="gn_eNOE"):
                diff[np.where((aa-ee)<0.0)] = 0.0

            red = diff/ees**2    
            viol = (np.argsort(red)[::-1])
            for el in viol:
                if(red[el]>1.5):
                    print("%15s %5.2f %5.2f %5.2f " % (lab[el],ee[el],aa[el],red[el]))
            
        fig.show()
        #plt.savefig("eNOE_nn1.pdf",dpi=600)
        #plt.close()
        print("")
        
plt.show()
```

```
List of large violations eNOE
          Label   exp  calc deviation 
   G2_H1;C3_H41  3.26  3.93  4.20 
  C3_H42;U11_H3  5.28  6.20  3.06 
  G10_H8;U11_H5  5.06  4.31  2.22 
   C8_H5';G9_H8  4.86  4.14  2.21 
   G9_H2';G9_H8  4.76  4.11  1.88 
   C5_H41;C5_H5  3.95  3.44  1.68 

List of large violations eNOE_unidir
          Label   exp  calc deviation 
   U6_H6;C8_H41  3.42  4.21  2.35 
 G10_H3';U11_H5  3.28  4.01  2.20 
  C8_H4';G9_H5"  3.17  2.50  1.99 
 U6_HO2';U7_H2'  3.33  2.63  1.97 
 U6_H1';G10_H21  2.51  3.04  1.96 
 G10_H4';G10_H8  3.54  4.28  1.96 
   U7_H2';G9_H1  4.41  5.30  1.82 
  C8_H4';G9_H5'  3.21  3.84  1.74 
   A4_H3';C5_H6  2.83  3.38  1.71 
   G2_H1;C13_H6  4.82  5.76  1.67 
  G1_H2';G1_H5'  4.40  5.25  1.66 

List of large violations gn_eNOE
          Label   exp  calc deviation 
   C8_H5;G10_H8  5.87  9.00 28.46 
  C8_H2';G9_H2'  4.62  6.89 24.06 
   C3_H2';C3_H6  2.87  3.79 10.22 
 C8_H1';G10_H4'  5.28  6.81  8.36 
   U7_H5";C8_H6  2.86  3.50  4.95 
  U7_H4';U7_H5"  2.10  2.43  2.52 
  U7_H3';U7_H4'  2.49  2.81  1.75 

List of large violations NOE
          Label   exp  calc deviation 
   G2_H8;G2_H5'  2.11  3.87 16.04 
   G2_H1;C3_H42  2.49  4.46 10.08 
  A4_H1';C5_H1'  3.06  5.62  7.44 
  G12_H1;C3_H42  2.35  3.69  5.95 
   A4_H3';C5_H6  2.23  3.38  5.34 
  C5_H42;G10_H1  2.48  3.79  4.47 
   C3_H41;C3_H5  2.32  3.42  4.12 
G10_H5'';G10_HO2'  3.17  5.08  3.56 
   C5_H5;C5_H41  2.40  3.44  3.20 
 A4_H3';A4_HO2'  2.56  3.65  2.71 
 G10_H8;G10_H4'  2.94  4.28  2.44 
  C5_H6;C5_H5''  2.81  4.01  2.31 
 C13_H41;C13_H5  2.46  3.32  2.03 
   G1_H3';G2_H8  2.40  3.19  1.86 
   C5_H3';U6_H6  2.53  3.38  1.76 
  A4_H8;A4_H5''  2.97  4.09  1.62 

List of large violations J3
          Label   exp  calc deviation 
       G9-2H5H4 -4.60 10.44 100.57 
        10-2H5P  1.30 12.18 52.59 
        10-1H5P  2.30  9.26 21.55 
         9-2H5P  8.60  2.52 16.43 
         9-1H5P  9.00  3.00 15.98 
       G10-H3H4  4.90 10.85 15.75 
         8-C4Pe  7.10  1.31 14.90 
       C14-H3H4  5.30 10.59 12.43 
       G9-1H5H4 -2.80  1.82  9.50 
         7-2H5P  5.70  2.00  6.09 
      C13-1H5H4 -1.70  1.97  5.98 
         1-C4Pe  6.00  9.63  5.85 
          6-H3P  7.30  3.80  5.45 
         3-1H5P  0.00  3.45  5.30 
         7-C4Pb  6.30  9.59  4.80 
        10-C4Pb  6.10  2.89  4.57 
          7-H3P  3.70  6.88  4.49 
         9-C4Pe  7.70  4.55  4.41 
        14-C4Pb  7.70 10.79  4.26 
        12-C4Pe  6.50  9.59  4.24 
         13-H3P  9.30  6.31  3.98 
         2-C4Pb  7.80 10.77  3.91 
      U11-2H5H4  4.50  1.62  3.69 
          5-H3P  9.40  6.62  3.44 
      U11-1H5H4 -0.90  1.75  3.13 
         9-C4Pb  8.00 10.64  3.10 
          4-H3P  3.80  6.39  2.99 
        G9-H2H3  3.40  5.98  2.96 
       U6-1H5H4 -0.60  1.97  2.93 
      C14-1H5H4 -0.80  1.64  2.66 
        G2-H2H3  6.00  3.72  2.31 
         11-H3P  8.00  5.87  2.02 
       C13-H2H3  5.80  3.71  1.94 
         7-1H5P  3.70  5.75  1.87 
        U7-H1H2  8.00 10.00  1.78 
       G1-2H5H4  5.20  3.21  1.76 
        13-1H5P  1.70  3.68  1.75 
        11-2H5P  0.00  1.96  1.71 
       C13-H3H4  9.10 11.05  1.69 
         6-1H5P  1.70  3.64  1.67 
        G9-H1H2  0.60  2.53  1.66 
         5-1H5P  1.80  3.71  1.63 

List of large violations RDC
          Label   exp  calc deviation 
8.C.C1'-8.C.H1' -14.66 -3.21 131.14 
  5.C.C6-5.C.H6 10.99  4.40 43.46 
1.G.C1'-1.G.H1' -2.39 -8.82 41.35 
10.G.C8-10.G.H8 11.10  4.78 39.99 
  9.G.N1-9.G.H1 -6.32 -0.02 39.69 
10.G.N1-10.G.H1 -3.88  1.82 32.51 
11.U.C6-11.U.H6  9.09  3.51 31.18 
  1.G.C8-1.G.H8  7.29  2.72 20.90 
3.C.C1'-3.C.H1'  6.36  1.99 19.10 
  9.G.C8-9.G.H8  0.34 -3.95 18.39 
  4.A.C8-4.A.H8  6.02  2.63 11.50 
11.U.N3-11.U.H3 -1.75  1.45 10.24 
14.C.C6-14.C.H6  6.43  3.28  9.93 
11.U.C1'-11.U.H1' -0.43  2.56  8.92 
9.G.C1'-9.G.H1' -5.14 -2.23  8.45 
12.G.C8-12.G.H8  5.36  2.53  7.99 
2.G.C1'-2.G.H1' -0.32 -3.13  7.92 
7.U.C1'-7.U.H1' -1.95 -4.49  6.43 
5.C.C1'-5.C.H1'  4.95  7.40  6.02 
  3.C.C6-3.C.H6  3.54  1.42  4.50 
13.C.C1'-13.C.H1'  6.53  8.12  2.54 
  7.U.C6-7.U.H6  3.47  1.97  2.25 
  2.G.C8-2.G.H8  1.63  3.10  2.16 
  8.C.C6-8.C.H6 -3.66 -5.02  1.84 
6.U.C1'-6.U.H1'  6.89  8.23  1.79 

List of large violations RDC1
          Label   exp  calc deviation 
8.C.C1'-8.C.H1' -20.54 -4.65 128.82 
10.G.C1'-10.G.H1' -19.61 -4.86 111.03 
10.G.C2'-10.G.H2'  5.72 -7.98 95.74 
7.U.C3'-7.U.H3'  2.77 -9.88 81.65 
13.C.C2'-13.C.H2' -16.32 -4.28 73.94 
9.G.C3'-9.G.H3' 10.20 -1.79 73.40 
  6.U.C5-6.U.H5 15.44  3.51 72.58 
13.C.C5-13.C.H5 14.68  2.86 71.27 
5.C.C2'-5.C.H2' 10.01 -1.61 68.85 
  8.C.C5-8.C.H5 -11.69 -0.10 68.53 
  5.C.C6-5.C.H6 17.56  6.38 63.78 
14.C.C4'-14.C.H4' 14.07  3.41 58.00 
7.U.C1'-7.U.H1'  2.68 -6.51 43.05 
12.G.C1'-12.G.H1'  0.92  9.58 38.27 
12.G.C8-12.G.H8 12.21  3.67 37.17 
  1.G.C8-1.G.H8 11.77  3.94 31.28 
11.U.C6-11.U.H6 12.43  5.08 27.53 
  5.C.C5-5.C.H5  7.23  0.23 25.00 
  4.A.C8-4.A.H8 10.35  3.81 21.80 
  3.C.C5-3.C.H5 10.35  4.62 16.73 
14.C.C6-14.C.H6  9.59  4.75 11.94 
8.C.C4'-8.C.H4'  3.80  8.57 11.60 
11.U.C5-11.U.H5 10.28  5.53 11.49 
7.U.C4'-7.U.H4' -11.64 -6.96 11.18 
4.A.C2'-4.A.H2' -5.97 -9.85  7.69 
8.C.C3'-8.C.H3' -14.78 -11.01  7.24 
6.U.C1'-6.U.H1'  8.35 11.93  6.55 
2.G.C1'-2.G.H1' -1.06 -4.55  6.22 
  8.C.C6-8.C.H6 -4.86 -7.27  2.97 
3.C.C2'-3.C.H2' -11.82 -14.18  2.84 
13.C.C1'-13.C.H1'  9.66 11.79  2.31 
5.C.C1'-5.C.H1'  8.78 10.73  1.95 
9.G.C1'-9.G.H1' -5.07 -3.24  1.71 
14.C.C1'-14.C.H1' 12.23 10.51  1.51 

List of large violations RDC2
          Label   exp  calc deviation 
6.U.C4'-6.U.H4' 15.09 -0.08 47.54 
10.G.C8-10.G.H8 14.76  3.95 24.13 
1.G.C1'-1.G.H1'  2.98 -7.31 21.87 
6.U.C2'-6.U.H2' 12.82  2.66 21.33 
10.G.C3'-10.G.H3' -11.37 -5.68  6.68 
  7.U.C5-7.U.H5  4.46 -1.22  6.67 
  7.U.C6-7.U.H6  7.30  1.62  6.66 
9.G.C2'-9.G.H2'  2.64 -2.82  6.16 
7.U.C2'-7.U.H2' -1.29 -5.13  3.04 
11.U.C3'-11.U.H3' -4.70 -1.06  2.74 

List of large violations sPRE
          Label   exp  calc deviation 
        5CytH42  0.09  0.44 3803.58 
        3CytH42  0.08  0.43 2792.39 
        11UraH5  0.11  0.40 1421.19 
        5CytH41  0.09  0.29 1080.53 
         6UraH5  0.17  0.43 1015.88 
       13CytH42  0.16  0.47 665.76 
        13CytH5  0.16  0.34 579.80 
         3CytH5  0.14  0.32 474.20 
      14Cyt1H2'  0.35  0.71 361.32 
        10GuaH8  0.19  0.44 351.49 
        3CytH41  0.07  0.24 327.19 
      10Gua1H5'  0.37  0.64 324.23 
         5CytH5  0.12  0.30 314.62 
        5CytH1'  0.22  0.33 258.96 
       13CytH41  0.16  0.31 250.72 
       14CytH1'  0.58  0.43 223.14 
      10Gua2H5'  0.35  0.61 214.14 
       10GuaH3'  0.23  0.33 179.34 
        2GuaH4'  0.53  0.75 155.78 
        6UraH1'  0.12  0.19 148.39 
        5CytH3'  0.20  0.28 128.15 
        6UraH3'  0.22  0.28 118.35 
       9Gua2H5'  0.54  0.33 103.34 
       12GuaH1'  0.27  0.36 92.75 
        4AdeH1'  0.25  0.33 92.15 
       10GuaH21  0.10  0.22 81.00 
        4AdeH61  0.01  0.29 76.98 
         6UraH3  2.36  0.28 76.67 
        8CytH41  0.46  0.69 72.38 
       6Ura1H2'  0.15  0.20 64.98 
       12GuaH4'  0.49  0.72 64.84 
        6UraH4'  0.28  0.37 63.91 
      13Cyt1H2'  0.31  0.27 60.58 
        11UraH6  0.13  0.22 58.63 
       13CytH1'  0.26  0.32 57.00 
       1Gua1H2'  0.66  0.35 55.82 
      13Cyt1H5'  0.50  0.62 48.32 
        11UraH3  0.07  0.14 47.70 
       5Cyt1H2'  0.20  0.23 46.63 
       4Ade1H2'  0.20  0.24 40.14 
        8CytH3'  0.44  0.52 38.94 
        4AdeH4'  0.50  0.71 37.20 
        14CytH6  0.51  0.33 37.15 
       7Ura1H2'  0.32  0.26 34.72 
       2Gua2H5'  0.98  0.69 34.55 
       10GuaH22  0.11  0.23 33.70 
         4AdeH2  0.12  0.18 32.28 
        1GuaH3'  0.20  0.44 31.63 
      11Ura1H2'  0.20  0.23 31.56 
        10GuaH1  0.07  0.15 30.89 
         4AdeH8  0.16  0.21 30.88 
       14CytH3'  1.12  0.65 29.50 
        4AdeH62 -0.00  0.27 26.97 
         5CytH6  0.15  0.21 26.50 
        1GuaH4'  1.29  0.93 25.18 
        8CytH1'  0.28  0.33 20.75 
        12GuaH1  0.06  0.14 20.38 
         1GuaH8  1.72  0.79 16.60 
      10Gua1H2'  0.18  0.22 16.25 
       10GuaH1'  0.18  0.20 15.39 
        8CytH4'  0.31  0.33 15.27 
       11UraH3'  0.26  0.28 14.68 
         9GuaH1  0.45  0.29 14.66 
       7Ura1H5'  0.64  0.76 14.36 
       13CytH4'  0.40  0.69 13.25 
        2GuaH1'  0.46  0.42 12.26 
         8CytH5  0.29  0.34 11.91 
        12GuaH8  0.17  0.22 11.56 
       8Cyt1H2'  0.40  0.47  9.96 
        7UraH1'  0.83  0.60  9.37 
        13CytH6  0.18  0.22  8.85 
        9GuaH3'  0.32  0.30  6.20 
       14CytH42  1.10  0.68  5.79 
         2GuaH8  0.56  0.27  4.62 
       2Gua1H5'  0.58  0.68  3.37 
       14CytH41  1.28  0.58  2.56 
         7UraH5  0.64  0.72  2.31 
         2GuaH1  0.15  0.17  2.22 
         9GuaH8  0.96  0.83  1.79 
        9GuaH1'  0.80  0.72  1.70 
       14CytH4'  0.87  0.80  1.54
```

In [ ]:
