## Supplementary material for "Integrating NMR and Simulations Reveals Motions in the UUCG Tetraloop": SI3 (from another version of this preprint: version 2, 2019-09-11)

SI3\_BME


### Integrate simulations and experimental data¶

In this notebook we refine the MD simulation to match experimental data using the Bayesian/MaxEnt approach (BME). In brief, what we use as input is:

1. A set of experimental constraints ${F\_1^{exp} ... F\_m^{exp}}$ and associated errors ${\sigma\_1... \sigma\_m}$
2. The experimental observables back-calculated for each of the $n$ frames in the simulation ($F(x\_j)\_{i}^{calc}, i=1...m, j=1...n$)
3. The initial weights $w^0\_1...w^0\_n$. In a plain MD simulation the initial weights are uniform ($w\_i=1/n$), but we here use metadynamics, and the weights are given by $w^0\_j \propto \exp( V(x\_j)/kbt)$, where $V$is the final bias potential. The appendix at the end of this notebook describes how to calculate the bias using Plumed.

The output of the procedure is a new set of weights ${w\_1...w\_n}$ that are such that the new averages $<F\_i^{calc}> = \sum\_j^n w\_j F(x\_j)\_{i}^{calc} \approx F\_i^{exp} $. Clearly, the new weights can be used to calculate any desired quantity or distribution.

These steps are largely automatized by our BME script, that can be found here. In essence, the user has to perform the following actions:

```
# initialize the class, specifying the initial weights (if non-uniform)
rew = bme.Reweight(w0=bias,kbt=kbt)
# load the experimental and back-calculated data files
rew.load(exp,calc)
# Run the optimization. A regularization parameter, theta, has to be specified
chi2_before,chi2_after,effective_frames = rew.optimize(theta=t)
# Calculate new averages with the new weights. Note that the exp and calc can be the original 
data used as input (training set), or new data (cross-validation or test set)
# statistics are written to a file
chi = rew.weight_exp(exp,calc,out_filename)
```

We choose the regularization parameter theta by scanning different theta values using the dataset A1 as training, monitoring at the same time the agreement with datasets A0, B, C, and D. We will then pick the theta by identifying the 'elbow' in the $theta$ versus $\chi^2$ plot, and making sure that the choice performs well on the cross-validation sets. To be fully general, one can also try to use dataset B (or C or D or A1) for training and all the rest as cross-validation.

In [4]:

```
import sys
import numpy as np
import os
# add BME to the path (should be changed and it depends where the BME script is located)
sys.path.append('/Users/sandrobottaro/Software/BME/')
import bme_reweight as bme

# set the intial weights from metadynamics. The bias is passed along with kbt, so 
# that the initial weights are set proportional to exp(bias/kbt)
kbt = 0.008314462*280
bias = [float(line.split()[1]) for line in open("data/bias") if "#" not in line]

# this is a list of available data
lista = [["set_A0","eNOE","eNOE_unidir","gn_eNOE"],\
         ["set_A1","eNOE","eNOE_unidir","gn_eNOE"],\
        ["set_B","NOE","RDC","J3"],\
        ["set_C","RDC1","RDC2"],\
        ["set_D","sPRE"]]
lista1 = [["set_A0","eNOE","eNOE_unidir","gn_eNOE"],\
         ["set_A1","eNOE","eNOE_unidir","gn_eNOE"],\
        ["set_B","NOE","RDC","J3"],\
        ["set_C","RDC1","RDC2"],\
        ["set_D","sPRE"]]

lista1 = [["set_A1","eNOE","eNOE_unidir","gn_eNOE"]]
# Scanning different theta values. Theta is a free parameter that will be set later.
#thetas = [1,3,6,10,30,60,100,300,600,1000,5000,10000]
thetas = [2,10,20,30,50,100,250,500,1000,10000]
#thetas = [2,10,20,50,250,500]
#thetas = [30,60,100,300,600,1000,5000,10000]
%mkdir -p data/comparison
%mkdir -p data/comparison/set_A1
%mkdir -p data/comparison/set_A0
%mkdir -p data/comparison/set_B
%mkdir -p data/comparison/set_C
%mkdir -p data/comparison/set_D
```

In [5]:

```
for t in thetas:
    for i in range(len(lista1)):
        rew= bme.Reweight(w0=bias,kbt=kbt)
        if(0==0):
            for j in range(1,len(lista1[i])):
                # define experimental and calculated datafile
                exp = "data/%s/%s.exp.dat" % (lista1[i][0],lista1[i][j])
                calc = "data/%s/%s.calc.dat" % (lista1[i][0],lista1[i][j])
                # Load data
                rew.load(exp,calc)
                # Optimize
            before,after,neff = rew.optimize(theta=t)
            # now loop over the other experimental data for cross-validation
            for l in range(len(lista)):
                for m in range(1,len(lista[l])):
                    exp2 = "data/%s/%s.exp.dat" % (lista[l][0],lista[l][m])
                    calc2 = "data/%s/%s.calc.dat" % (lista[l][0],lista[l][m])
                    chi = rew.weight_exp(exp2,calc2,"data/comparison/%s/rw_%s_%s_theta_%d" % (lista1[i][0],lista[l][0],lista[l][m],t))
                    
        else:
            
            print("IMPOSSIBLE",lista1[i][0],t)
```

```
# Assuming weights given as minus free energies. w=exp(bias/kbt) kbt=  2.3280 
# Set non-uniform initial weights from file. Sum= 11503.331447473747 100001
# Warning: expt average        U11_H3;U11_H6=1.7960e-04 is larger than maximum value in simulation 1.0536e-04
# Warning: expt average         C5_H41;C5_H6=3.1401e-05 is smaller than minimum value in simulation 3.4403e-05
# Warning: expt average      C13_H1';C13_H3'=1.4759e-04 is smaller than minimum value in simulation 1.7333e-04
# Warning: expt average         C5_H4';C5_H6=5.0814e-04 is larger than maximum value in simulation 4.7732e-04
# Warning: expt average       G10_H4';G10_H8=5.0814e-04 is larger than maximum value in simulation 4.1958e-04
# Warning: expt average         C3_H1';C3_H6=2.2001e-04 is smaller than minimum value in simulation 2.5194e-04
# Warning: expt average        C3_H1';C3_H2'=8.5647e-04 is smaller than minimum value in simulation 8.9282e-04
# Warning: expt average         A4_H1';A4_H8=1.1375e-04 is smaller than minimum value in simulation 1.9556e-04
# Warning: expt average        A4_H1';A4_H2'=6.7424e-04 is smaller than minimum value in simulation 9.5273e-04
# Warning: expt average        A4_H1';A4_H3'=1.5557e-04 is smaller than minimum value in simulation 1.6570e-04
# Warning: expt average        A4_H1';A4_H4'=2.1300e-04 is smaller than minimum value in simulation 2.2308e-04
# Warning: expt average        U7_H1';U7_H2'=3.7857e-04 is smaller than minimum value in simulation 8.1448e-04
# Warning: expt average         C8_H1';C8_H6=2.0298e-04 is smaller than minimum value in simulation 2.4134e-04
# Warning: expt average        C8_H1';C8_H2'=4.5332e-04 is smaller than minimum value in simulation 8.5016e-04
# Warning: expt average        C8_H1';C8_H3'=1.1050e-04 is smaller than minimum value in simulation 1.8231e-04
# Warning: expt average      G12_H1';G12_H3'=9.5788e-05 is smaller than minimum value in simulation 1.8902e-04
# theta: 2
# Constrained optimization: Fall back to L-BFGS_B
```

```
---------------------------------------------------------------------------
KeyboardInterrupt                         Traceback (most recent call last)
<ipython-input-5-b9d8e5dba126> in <module>
     10                 rew.load(exp,calc)
     11                 # Optimize
---> 12             before,after,neff = rew.optimize(theta=t)
     13             # now loop over the other experimental data for cross-validation
     14             for l in range(len(lista)):

~/Software/BME/bme_reweight.py in optimize(self, theta, method)
    454                 #                           bounds=self.bounds)
    455                 print("# Constrained optimization: Fall back to L-BFGS_B")
--> 456                 result = optimize.minimize(func_maxent_gauss,lambdas,options=opt,method=meth,jac=True,bounds=self.bounds)
    457             else:
    458                 if meth == 'trust':

/anaconda3/lib/python3.7/site-packages/scipy/optimize/_minimize.py in minimize(fun, x0, args, method, jac, hess, hessp, bounds, constraints, tol, callback, options)
    599     elif meth == 'l-bfgs-b':
    600         return _minimize_lbfgsb(fun, x0, args, jac, bounds,
--> 601                                 callback=callback, **options)
    602     elif meth == 'tnc':
    603         return _minimize_tnc(fun, x0, args, jac, bounds, callback=callback,

/anaconda3/lib/python3.7/site-packages/scipy/optimize/lbfgsb.py in _minimize_lbfgsb(fun, x0, args, jac, bounds, disp, maxcor, ftol, gtol, eps, maxfun, maxiter, iprint, callback, maxls, **unknown_options)
    333             # until the completion of the current minimization iteration.
    334             # Overwrite f and g:
--> 335             f, g = func_and_grad(x)
    336         elif task_str.startswith(b'NEW_X'):
    337             # new iteration

/anaconda3/lib/python3.7/site-packages/scipy/optimize/lbfgsb.py in func_and_grad(x)
    283     else:
    284         def func_and_grad(x):
--> 285             f = fun(x, *args)
    286             g = jac(x, *args)
    287             return f, g

/anaconda3/lib/python3.7/site-packages/scipy/optimize/optimize.py in function_wrapper(*wrapper_args)
    298     def function_wrapper(*wrapper_args):
    299         ncalls[0] += 1
--> 300         return function(*(wrapper_args + args))
    301 
    302     return ncalls, function_wrapper

/anaconda3/lib/python3.7/site-packages/scipy/optimize/optimize.py in __call__(self, x, *args)
     61     def __call__(self, x, *args):
     62         self.x = numpy.asarray(x).copy()
---> 63         fg = self.fun(x, *args)
     64         self.jac = fg[1]
     65         return fg[0]

~/Software/BME/bme_reweight.py in func_maxent_gauss(lambdas)
    359             ww /= zz
    360             # new averages
--> 361             avg = np.sum((ww[:,np.newaxis]*self.sim_data), axis=0)
    362 
    363             # errors are rescaled by factor theta

KeyboardInterrupt:
```

In [6]:

```
# import plotting libraries
import matplotlib.pyplot as plt
import seaborn as sns 
import subprocess

cp = sns.color_palette("colorblind", 12)

for i in range(len(lista1)):
    
    fig, axs = plt.subplots(1,5,figsize=(15,6))
    p = 0
    for l in range(len(lista)):

        for m in range(1,len(lista[l])):
            data = []
            for t in thetas:
       
                filename =  "data/comparison/%s/rw_%s_%s_theta_%d.stats.dat" %\
                                            (lista1[i][0],lista[l][0],lista[l][m],t)
                #print(filename,os.path.isfile(filename))
                if(os.path.isfile(filename)): 
                    line = subprocess.check_output(['head', '-1', filename])
                    srel = str(line.split()[-1]).split("=")[-1][:-1]
                    neff = np.exp(float(srel))
                    #print(neff)
                    line = subprocess.check_output(['tail', '-1', filename])
                    chi_b = float(line.split()[-2])
                    chi_a = float(line.split()[-1])
                    #print(filename,chi_a)
                    data.append([t,neff,chi_a,chi_b])
                else:
                    data.append([t,0,0,0])
            data = np.array(data)

            if(lista[i][0]==lista[l][0]):
                axs[l].plot(data[:,1],data[:,2],"-+",color=cp[p],label=lista[l][m],lw=1.5,ls="--")
            else:
                    
                axs[l].plot(data[:,1],data[:,2],"-+",color=cp[p],label=lista[l][m],lw=1)
            axs[l].scatter(1,data[-1,3],marker="*",c='k')
            ax2 = axs[l].twiny()

            axs[l].set(xscale="log")
            ax2.set(xscale="log")
            ax2.plot(data[:,1],data[:,2],c='w',lw=0.001)
            ax2.set_xticks(data[:,1])
            ax2.set_xticklabels(["%d"% s for s in thetas],rotation=60)
            p += 1
            if(l==1):
                axs[l].set_title("Training: %s" % lista1[i][0])
            if(l==0):
                axs[l].set_ylabel(r"$\chi^2$")
            axs[l].set_xlabel("Neff")
            ax2.set_xlabel(r"$\theta$")
        axs[l].legend()
        lims = axs[l].get_ylim()
        axs[l].set_ylim(0,lims[1])
    plt.show()
    plt.savefig("figures/theta_scan_train_%s.pdf" % (lista[i][0]))
    plt.close()
```

```
<Figure size 1500x600 with 17 Axes>
```

### $\theta$ selection¶

Let us focus on the first plot in the firs row. As theta is set to small values (top x axis), the $\chi^2$ becomes smaller and smaller (y axis). This is expected because we use these data (set A) for training. Note also that when we choose a small $\theta$, the number of effective frames becomes small: this is because we are essentially 'throwing away' some of the frames by giving them a very small weight. The other plots in the first row show the agreement on the cross-validation sets. By construction, dataset A1 is almost identical to dataset A, and it has the same behavior. The use of eNOE slighty improves the agreement with RDC in dataset B, but worsens the agreement with dataset D (sPRE).
In the second row we show the results by using set A1 for training and A,B,C,D as cross validation.
In the third row we show the results by using set B for training and A,A1,C,D as cross validation.
In the fourth row we show the results by using set C for training and A,A1,B,D as cross validation.
In the last row we show the results by using set D for training and A,A1,B,C as cross validation.

Note that only when using eNOE for training the agreement on the cross-validation datasets do not worsen significantly. We therefore use dataset A for training in our work, and set $\theta=30$. To get an error estimate, we split the simulation into four blocks and reweight the four blocks independently.

In [7]:

```
nblocks = 4
theta=30
bsize = int(len(bias)/nblocks)

exp1 = "data/set_A1/eNOE.exp.dat" 
calc1 = "data/set_A1/eNOE.calc.dat" 
exp2 = "data/set_A1/eNOE_unidir.exp.dat" 
calc2 = "data/set_A1/eNOE_unidir.calc.dat"
exp3 = "data/set_A1/gn_eNOE.exp.dat" 
calc3 = "data/set_A1/gn_eNOE.calc.dat" 

for p in range(nblocks):
 
    start = p*bsize
    stop = start+bsize

    rew= bme.Reweight(w0=bias,kbt=kbt)
    rew.load(exp1,calc1,rows=range(start,stop))
    rew.load(exp2,calc2,rows=range(start,stop))
    rew.load(exp3,calc3,rows=range(start,stop))
    
    # Optimize
    before,after,neff = rew.optimize(theta=theta)
            
    # print weights to file
    w_opt = rew.get_weights()
    w_0 = np.exp(np.array(bias)[range(start,stop)]/kbt)
    w_0 /= np.sum(w_0)
    stri = ""
    for ii in range(len(w_opt)):
        stri += "%10.4e %10.4e \n" % (w_0[ii],w_opt[ii])
    fhw = open("data/weights_%s_t_%d_rep_%d.dat" % ("set_A1",theta,p),"w")
    fhw.write(stri)
    fhw.close()
    
    # now loop over the other experimental data for cross-validation
    for l in range(len(lista)):
        for m in range(1,len(lista[l])):
            exp_c = "data/%s/%s.exp.dat" % (lista[l][0],lista[l][m])
            calc_c = "data/%s/%s.calc.dat" % (lista[l][0],lista[l][m])
            chi = rew.weight_exp(exp_c,calc_c,\
                                 "data/comparison/%s/rw_%s_%s_theta_%d_rep_%d"\
                                  % ("set_A1",lista[l][0],lista[l][m],theta,p),rows=range(start,stop))
                
    
# do it again, to get weights wihtout bins - useful for 2d projections
rew= bme.Reweight(w0=bias,kbt=kbt)
rew.load(exp1,calc1)
rew.load(exp2,calc2)
rew.load(exp3,calc3)
           
# Optimize
before,after,neff = rew.optimize(theta=theta)
w_opt = rew.get_weights()
w_0 = np.exp(np.array(bias)/kbt)
w_0 /= np.sum(w_0)
stri = ""
for ii in range(len(w_opt)):
    stri += "%10.4e %10.4e \n" % (w_0[ii],w_opt[ii])
fhw = open("data/weights_%s_t_%d.dat" % ("set_A1",theta),"w")
fhw.write(stri)
fhw.close()
```

```
# Assuming weights given as minus free energies. w=exp(bias/kbt) kbt=  2.3280 
# Set non-uniform initial weights from file. Sum= 11503.331447473747 100001
# Warning: expt average         G2_H1;C13_H6=7.9748e-05 is larger than maximum value in simulation 4.0578e-05
# Warning: expt average        U11_H3;U11_H6=1.7960e-04 is larger than maximum value in simulation 1.0253e-04
# Warning: expt average         C5_H41;C5_H6=3.1401e-05 is smaller than minimum value in simulation 3.4807e-05
# Warning: expt average      C13_H1';C13_H3'=1.4759e-04 is smaller than minimum value in simulation 2.0084e-04
# Warning: expt average         C5_H4';C5_H6=5.0814e-04 is larger than maximum value in simulation 4.4971e-04
# Warning: expt average       G10_H4';G10_H8=5.0814e-04 is larger than maximum value in simulation 3.7553e-04
# Warning: expt average         C3_H1';C3_H6=2.2001e-04 is smaller than minimum value in simulation 2.5787e-04
# Warning: expt average        C3_H1';C3_H2'=8.5647e-04 is smaller than minimum value in simulation 9.7491e-04
# Warning: expt average        C3_H1';C3_H4'=2.8683e-04 is smaller than minimum value in simulation 3.3040e-04
# Warning: expt average         A4_H1';A4_H8=1.1375e-04 is smaller than minimum value in simulation 1.9556e-04
# Warning: expt average        A4_H1';A4_H2'=6.7424e-04 is smaller than minimum value in simulation 1.0877e-03
# Warning: expt average        A4_H1';A4_H3'=1.5557e-04 is smaller than minimum value in simulation 1.9359e-04
# Warning: expt average        A4_H1';A4_H4'=2.1300e-04 is smaller than minimum value in simulation 2.3085e-04
# Warning: expt average        U7_H1';U7_H2'=3.7857e-04 is smaller than minimum value in simulation 8.9415e-04
# Warning: expt average         C8_H1';C8_H6=2.0298e-04 is smaller than minimum value in simulation 2.4948e-04
# Warning: expt average        C8_H1';C8_H2'=4.5332e-04 is smaller than minimum value in simulation 8.6363e-04
# Warning: expt average        C8_H1';C8_H3'=1.1050e-04 is smaller than minimum value in simulation 1.8317e-04
# Warning: expt average      G12_H1';G12_H2'=9.6885e-04 is smaller than minimum value in simulation 1.0576e-03
# Warning: expt average      G12_H1';G12_H3'=9.5788e-05 is smaller than minimum value in simulation 2.0192e-04
# Warning: expt average      G12_H1';G12_H5"=3.3631e-05 is smaller than minimum value in simulation 3.4526e-05
# theta: 30
# Constrained optimization: Fall back to L-BFGS_B
# RDC scaling factors 7.3452e-01, 7.0709e-01
# RDC scaling factors 1.1145e+00, 1.0236e+00
# RDC scaling factors 6.1904e-01, 5.9723e-01
# RDC scaling factors 6.1926e-03, 6.1850e-03
# Assuming weights given as minus free energies. w=exp(bias/kbt) kbt=  2.3280 
# Set non-uniform initial weights from file. Sum= 11503.331447473747 100001
# Warning: expt average         C5_H41;C5_H5=2.6328e-04 is smaller than minimum value in simulation 2.7802e-04
# Warning: expt average        U11_H3;U11_H6=1.7960e-04 is larger than maximum value in simulation 1.0536e-04
# Warning: expt average         C5_H41;C5_H6=3.1401e-05 is smaller than minimum value in simulation 3.4403e-05
# Warning: expt average      C13_H1';C13_H3'=1.4759e-04 is smaller than minimum value in simulation 1.7333e-04
# Warning: expt average        G2_H1';G2_H3'=1.9295e-04 is smaller than minimum value in simulation 2.0908e-04
# Warning: expt average         C5_H4';C5_H6=5.0814e-04 is larger than maximum value in simulation 4.3686e-04
# Warning: expt average       G10_H4';G10_H8=5.0814e-04 is larger than maximum value in simulation 3.3470e-04
# Warning: expt average         C3_H1';C3_H6=2.2001e-04 is smaller than minimum value in simulation 2.6356e-04
# Warning: expt average        C3_H1';C3_H2'=8.5647e-04 is smaller than minimum value in simulation 9.3465e-04
# Warning: expt average         A4_H1';A4_H8=1.1375e-04 is smaller than minimum value in simulation 2.0075e-04
# Warning: expt average        A4_H1';A4_H2'=6.7424e-04 is smaller than minimum value in simulation 9.5273e-04
# Warning: expt average        A4_H1';A4_H3'=1.5557e-04 is smaller than minimum value in simulation 1.6570e-04
# Warning: expt average        A4_H1';A4_H4'=2.1300e-04 is smaller than minimum value in simulation 2.2308e-04
# Warning: expt average        U7_H1';U7_H2'=3.7857e-04 is smaller than minimum value in simulation 8.1448e-04
# Warning: expt average         C8_H1';C8_H6=2.0298e-04 is smaller than minimum value in simulation 2.4195e-04
# Warning: expt average        C8_H1';C8_H2'=4.5332e-04 is smaller than minimum value in simulation 8.8685e-04
# Warning: expt average        C8_H1';C8_H3'=1.1050e-04 is smaller than minimum value in simulation 1.8286e-04
# Warning: expt average      G12_H1';G12_H3'=9.5788e-05 is smaller than minimum value in simulation 1.8902e-04
# theta: 30
# Constrained optimization: Fall back to L-BFGS_B
# RDC scaling factors 6.9135e-01, 7.2539e-01
# RDC scaling factors 1.0043e+00, 1.0355e+00
# RDC scaling factors 6.1683e-01, 5.8468e-01
# RDC scaling factors 6.1923e-03, 6.1876e-03
# Assuming weights given as minus free energies. w=exp(bias/kbt) kbt=  2.3280 
# Set non-uniform initial weights from file. Sum= 11503.331447473747 100001
# Warning: expt average        U11_H3;U11_H6=1.7960e-04 is larger than maximum value in simulation 1.0162e-04
# Warning: expt average         C5_H41;C5_H6=3.1401e-05 is smaller than minimum value in simulation 3.5711e-05
# Warning: expt average      C13_H1';C13_H3'=1.4759e-04 is smaller than minimum value in simulation 1.8502e-04
# Warning: expt average        G2_H1';G2_H3'=1.9295e-04 is smaller than minimum value in simulation 1.9618e-04
# Warning: expt average         C5_H4';C5_H6=5.0814e-04 is larger than maximum value in simulation 4.4115e-04
# Warning: expt average       G10_H4';G10_H8=5.0814e-04 is larger than maximum value in simulation 3.3656e-04
# Warning: expt average         C3_H1';C3_H6=2.2001e-04 is smaller than minimum value in simulation 2.5194e-04
# Warning: expt average        C3_H1';C3_H2'=8.5647e-04 is smaller than minimum value in simulation 1.0128e-03
# Warning: expt average        C3_H1';C3_H4'=2.8683e-04 is smaller than minimum value in simulation 3.0865e-04
# Warning: expt average         A4_H1';A4_H8=1.1375e-04 is smaller than minimum value in simulation 1.9792e-04
# Warning: expt average        A4_H1';A4_H2'=6.7424e-04 is smaller than minimum value in simulation 9.9353e-04
# Warning: expt average        A4_H1';A4_H3'=1.5557e-04 is smaller than minimum value in simulation 1.9005e-04
# Warning: expt average        A4_H1';A4_H4'=2.1300e-04 is smaller than minimum value in simulation 2.7430e-04
# Warning: expt average        U7_H1';U7_H2'=3.7857e-04 is smaller than minimum value in simulation 8.9849e-04
# Warning: expt average         C8_H1';C8_H6=2.0298e-04 is smaller than minimum value in simulation 2.5141e-04
# Warning: expt average        C8_H1';C8_H2'=4.5332e-04 is smaller than minimum value in simulation 8.8274e-04
# Warning: expt average        C8_H1';C8_H3'=1.1050e-04 is smaller than minimum value in simulation 1.8587e-04
# Warning: expt average      G12_H1';G12_H3'=9.5788e-05 is smaller than minimum value in simulation 1.9118e-04
# Warning: expt average      G12_H1';G12_H5"=3.3631e-05 is smaller than minimum value in simulation 3.4571e-05
# theta: 30
# Constrained optimization: Fall back to L-BFGS_B
# RDC scaling factors 7.0718e-01, 6.8458e-01
# RDC scaling factors 1.0634e+00, 9.9498e-01
# RDC scaling factors 6.6067e-01, 5.8331e-01
# RDC scaling factors 6.2253e-03, 6.1513e-03
# Assuming weights given as minus free energies. w=exp(bias/kbt) kbt=  2.3280 
# Set non-uniform initial weights from file. Sum= 11503.331447473747 100001
# Warning: expt average         C5_H41;C5_H5=2.6328e-04 is smaller than minimum value in simulation 2.6424e-04
# Warning: expt average        U11_H3;U11_H6=1.7960e-04 is larger than maximum value in simulation 1.0155e-04
# Warning: expt average         C5_H41;C5_H6=3.1401e-05 is smaller than minimum value in simulation 3.4777e-05
# Warning: expt average      C13_H1';C13_H3'=1.4759e-04 is smaller than minimum value in simulation 1.9801e-04
# Warning: expt average         C5_H4';C5_H6=5.0814e-04 is larger than maximum value in simulation 4.7732e-04
# Warning: expt average       G10_H4';G10_H8=5.0814e-04 is larger than maximum value in simulation 4.1958e-04
# Warning: expt average        G1_H3';G1_H5'=2.1300e-04 is smaller than minimum value in simulation 2.1332e-04
# Warning: expt average         C3_H1';C3_H6=2.2001e-04 is smaller than minimum value in simulation 2.5693e-04
# Warning: expt average        C3_H1';C3_H2'=8.5647e-04 is smaller than minimum value in simulation 8.9282e-04
# Warning: expt average        C3_H1';C3_H4'=2.8683e-04 is smaller than minimum value in simulation 2.8936e-04
# Warning: expt average         A4_H1';A4_H8=1.1375e-04 is smaller than minimum value in simulation 2.0058e-04
# Warning: expt average        A4_H1';A4_H2'=6.7424e-04 is smaller than minimum value in simulation 9.6466e-04
# Warning: expt average        A4_H1';A4_H3'=1.5557e-04 is smaller than minimum value in simulation 1.8249e-04
# Warning: expt average        A4_H1';A4_H4'=2.1300e-04 is smaller than minimum value in simulation 2.8829e-04
# Warning: expt average        U7_H1';U7_H2'=3.7857e-04 is smaller than minimum value in simulation 8.6651e-04
# Warning: expt average         C8_H1';C8_H6=2.0298e-04 is smaller than minimum value in simulation 2.4134e-04
# Warning: expt average        C8_H1';C8_H2'=4.5332e-04 is smaller than minimum value in simulation 8.5016e-04
# Warning: expt average        C8_H1';C8_H3'=1.1050e-04 is smaller than minimum value in simulation 1.8231e-04
# Warning: expt average      G12_H1';G12_H2'=9.6885e-04 is smaller than minimum value in simulation 1.0130e-03
# Warning: expt average      G12_H1';G12_H3'=9.5788e-05 is smaller than minimum value in simulation 1.9629e-04
# Warning: expt average      G12_H1';G12_H5"=3.3631e-05 is smaller than minimum value in simulation 3.4826e-05
# theta: 30
# Constrained optimization: Fall back to L-BFGS_B
# RDC scaling factors 6.5480e-01, 6.9124e-01
# RDC scaling factors 1.0142e+00, 1.0186e+00
# RDC scaling factors 5.7153e-01, 5.6226e-01
# RDC scaling factors 6.1197e-03, 6.1870e-03
# Assuming weights given as minus free energies. w=exp(bias/kbt) kbt=  2.3280 
# Set non-uniform initial weights from file. Sum= 11503.331447473747 100001
# Warning: expt average        U11_H3;U11_H6=1.7960e-04 is larger than maximum value in simulation 1.0536e-04
# Warning: expt average         C5_H41;C5_H6=3.1401e-05 is smaller than minimum value in simulation 3.4403e-05
# Warning: expt average      C13_H1';C13_H3'=1.4759e-04 is smaller than minimum value in simulation 1.7333e-04
# Warning: expt average         C5_H4';C5_H6=5.0814e-04 is larger than maximum value in simulation 4.7732e-04
# Warning: expt average       G10_H4';G10_H8=5.0814e-04 is larger than maximum value in simulation 4.1958e-04
# Warning: expt average         C3_H1';C3_H6=2.2001e-04 is smaller than minimum value in simulation 2.5194e-04
# Warning: expt average        C3_H1';C3_H2'=8.5647e-04 is smaller than minimum value in simulation 8.9282e-04
# Warning: expt average         A4_H1';A4_H8=1.1375e-04 is smaller than minimum value in simulation 1.9556e-04
# Warning: expt average        A4_H1';A4_H2'=6.7424e-04 is smaller than minimum value in simulation 9.5273e-04
# Warning: expt average        A4_H1';A4_H3'=1.5557e-04 is smaller than minimum value in simulation 1.6570e-04
# Warning: expt average        A4_H1';A4_H4'=2.1300e-04 is smaller than minimum value in simulation 2.2308e-04
# Warning: expt average        U7_H1';U7_H2'=3.7857e-04 is smaller than minimum value in simulation 8.1448e-04
# Warning: expt average         C8_H1';C8_H6=2.0298e-04 is smaller than minimum value in simulation 2.4134e-04
# Warning: expt average        C8_H1';C8_H2'=4.5332e-04 is smaller than minimum value in simulation 8.5016e-04
# Warning: expt average        C8_H1';C8_H3'=1.1050e-04 is smaller than minimum value in simulation 1.8231e-04
# Warning: expt average      G12_H1';G12_H3'=9.5788e-05 is smaller than minimum value in simulation 1.8902e-04
# theta: 30
# Constrained optimization: Fall back to L-BFGS_B
```

### appendix: calculating the final bias potential for each frame¶

A first step we take the average HILLS file

`bash ../script/medie.sh < data/HILLS.0 200000 1000000 > data/HILLS.0.avg`

Then, calculate the bias for each frame using the following `plumed.weights.dat` file.

```
RESTART
MOLINFO STRUCTURE=uucg_gmx.pdb
WHOLEMOLECULES ENTITY0=1-447
ee3: ERMSD ATOMS=@lcs-1,@lcs-2,@lcs-3,@lcs-4,@lcs-5,@lcs-6,@lcs-7,@lcs-8,@lcs-9,@lcs-10,@lcs-11,@lcs-12,@lcs-13,@lcs-14 REFERENCE=2koc_gmx.pdb CUTOFF=3.2
metad: METAD ARG=ee3 PACE=500000001 HEIGHT=0.0 SIGMA=0.1 FILE=HILLS.0.avg TEMP=280 BIASFACTOR=15 GRID_MIN=0.0 GRID_MAX=6.0
bias: REWEIGHT_BIAS TEMP=280
PRINT ARG=metad.bias FILE=bias STRIDE=1 RESTART=NO
```

And run plumed driver

`plumed driver --plumed plumed.weights.dat --mf_xtc traj_temp_f_0.xtc`

This produces a file `bias` that contains the averaged final bias, and the weights can be estimated as $w\_i \propto exp(bias/kt)$

In [ ]:
