## Supplementary material for "Integrating NMR and Simulations Reveals Motions in the UUCG Tetraloop": SI2 (from another version of this preprint: version 2, 2019-09-11)

SI2\_back\_calculate


### Back-calculate experimental data from structure¶

Here we back-calcuate the experimental quantities from structures and trjactories. We first need to define three small helper functions

In [1]:

```
import numpy as np

# helper functions, subsititute strings. This is because the name of hydrogens is a mess
alt = {"H2'":"1H2'","H5''":"2H5'","H5'":"1H5'","HO2'":"2HO'","H5\"":"2H5'"}
def sub(ss):
    at = ss.split("_")[1]
    if(at in alt):
        at = alt[at]
    return ss.split("_")[0] + "-" + at

# read experimental datafile and returns a list of labels and experimental values
def read_exp(f_exp):
    
    labels = []
    vals = []
    fh = open(f_exp)
    
    for line in fh:
        if("#" not in line):
            r1 = line.split()[0].split(";")[0]
            r2 = line.split()[0].split(";")[1]
            v1 = np.sort([r1,r2])
            qq = v1[0] +"/"+ v1[1]
            if(qq in labels):
                print("# DUPLICATE. Skipping data.."),
                print(qq,vals[labels.index(qq)], line),
            else:
                vals.append([float(line.split()[1]),float(line.split()[2])])
                labels.append(qq)
            
    fh.close()
    return labels,vals


# find indeces in topology corresponding to labels in experimental datafile
def get_idxs(labels,top):
    
    atoms = []
    for atom in top.atoms:
        aa = str(atom).split("-")[1]
        if(aa in alt): aa = alt[aa]
        atoms.append("%s-%s" % (str(atom).split("-")[0],aa))
    pairs = []
    for el in labels:
        ss = el.split("/")
        at1= sub(ss[0])
        at2 = sub(ss[1])
        if(at1 in atoms and at2 in atoms):
            pairs.append([atoms.index(at1),atoms.index(at2)])
        else:
            print("# Warning: Either %s or %s are missing" % (at1,at2)) 
            return 0
    print("# Found %d pairs out of %d" % (len(pairs),len(labels)))
    return np.array(pairs)


def write(data,fname):
    fh = open(fname,"w")
    stri = ""
    for j in range(data.shape[0]):
        stri += "%10d "% j 
        stri += " ".join(["%10.4e" % data[j,k] for k in range(data.shape[1])])
        stri += "\n"
    fh.write(stri)
    fh.close()
```

### Dataset A0¶

The first dataset that we consider is the original set of eNOE data published in "Nichols, Parker J., et al. "High-resolution small RNA structures from exact nuclear Overhauser enhancement measurements without additional restraints." Communications biology 1.1 (2018): 61."

#### bidirectional exact noes (eNOE).¶

From the original list of 72 distances, 10 datapoints were removed, corresponding to distances between H41-H42, H21-H22, H5'-H5'' and H5-H6 protons in the same nucleotide. 6 URA H2' 7 URA H6 bi-dir 4.56 and 6 URA H2' 6 URA H6 bi-dir 3.83 are duplicated and were also removed. The experimental error was set to 10% of the eNOE. For all NOE data used here we use the standard $^{-6}$ averaging, i.e. $\text{NOE}\_{\text{CALC}} = (\sum\_i w\_i r\_i^{-6})^{-1/6}$

In [2]:

```
import mdtraj as md

top = "data/PDB/2koc_gmx.pdb"
traj = "data/traj_temp_f_0.xtc"
labels,vals = read_exp("data/set_A0/eNOE.exp.dat")

#######################
md_trj = md.load(traj,top=top)
pairs = get_idxs(labels,md_trj.topology)
# convert to Angstrom
dists_md = 10*md.compute_distances(md_trj,pairs)
# write to file 
write(dists_md,"data/set_A0/eNOE.calc.dat")

pdbs = ["2koc","6by5"]
dists_pdbs = []
for  p in pdbs:
    top = "data/PDB/%s.pdb" % p 
    traj = "data/PDB/%s.pdb" % p 
    md_trj = md.load(traj,top=top)
    pairs = get_idxs(labels,md_trj.topology)
    dists = 10.0*md.compute_distances(md_trj,pairs)
    write(dists,"data/set_A0/%s.eNOE.calc.dat" %p)
```

```
# Found 62 pairs out of 62
```

```
/anaconda3/lib/python3.7/site-packages/mdtraj/formats/pdb/pdbfile.py:198: UserWarning: Unlikely unit cell vectors detected in PDB file likely resulting from a dummy CRYST1 record. Discarding unit cell vectors.
  'cell vectors.', category=UserWarning)
/anaconda3/lib/python3.7/site-packages/mdtraj/core/trajectory.py:419: UserWarning: top= kwarg ignored since file contains topology information
  warnings.warn('top= kwarg ignored since file contains topology information')
```

```
# Found 62 pairs out of 62
# Found 62 pairs out of 62
```

#### unidirectional eNOE¶

From the original list of 189 distances, 6 intra-residue NOEs were removed as described above, and RGUA H1' 3 RCYT H6 uni-dir 5.01 was duplicated. The experimental error was set to 15%. After a preliminary refinement round we found it difficult to refine the simulation so as to match the following eNOE: C13\_H5;C13\_H5'';G12\_H5'';C13\_H5; #
C5\_H5;C5\_H5''; A4\_H62;U11\_H3; A4\_H61;U11\_H3; C5\_H4';C5\_H5; U7\_H5'';G9\_H1. Since these eNOEs have a high value of sigma fit, they were removed from the dataset.

In [3]:

```
labels,vals = read_exp("data/set_A0/eNOE_unidir.exp.dat")


top = "data/PDB/2koc_gmx.pdb"
traj = "data/traj_temp_f_0.xtc"
#######################
md_trj = md.load(traj,top=top)
pairs = get_idxs(labels,md_trj.topology)
# convert to Angstrom
dists_md = 10*md.compute_distances(md_trj,pairs)
# write to file 
write(dists_md,"data/set_A0/eNOE_unidir.calc.dat")

pdbs = ["2koc","6by5"]
dists_pdbs = []
for  p in pdbs:
    top = "data/PDB/%s.pdb" % p 
    traj = "data/PDB/%s.pdb" % p 
    md_trj = md.load(traj,top=top)
    pairs = get_idxs(labels,md_trj.topology)
    dists = 10.0*md.compute_distances(md_trj,pairs)
    write(dists,"data/set_A0/%s.eNOE_unidir.calc.dat" %p)
```

```
# Found 177 pairs out of 177
# Found 177 pairs out of 177
# Found 177 pairs out of 177
```

#### gn-eNOE¶

From 88 datapoints, 9 were removed and 5 RCYT H5" 5 RCYT H5, 13 RCYT H2' 13 RCYT H5 were duplicated, resulting in 77 upper-limits distances. The experimental error was set to 10%

In [4]:

```
labels,vals = read_exp("data/set_A0/gn_eNOE.exp.dat")
top = "data/PDB/2koc_gmx.pdb"
traj = "data/traj_temp_f_0.xtc"
#######################
md_trj = md.load(traj,top=top)
pairs = get_idxs(labels,md_trj.topology)
# convert to Angstrom
dists_md = 10*md.compute_distances(md_trj,pairs)
# write to file 
write(dists_md,"data/set_A0/gn_eNOE.calc.dat")

pdbs = ["2koc","6by5"]
dists_pdbs = []
for  p in pdbs:
    top = "data/PDB/%s.pdb" % p 
    traj = "data/PDB/%s.pdb" % p 
    md_trj = md.load(traj,top=top)
    pairs = get_idxs(labels,md_trj.topology)
    dists = 10.0*md.compute_distances(md_trj,pairs)
    write(dists,"data/set_A0/%s.gn_eNOE.calc.dat" %p)
```

```
# Found 77 pairs out of 77
# Found 77 pairs out of 77
# Found 77 pairs out of 77
```

### Dataset A1¶

Dataset A1 is identical to A, with the following additions. Dataset A1 is the one used in the Manuscript. Dataset A is used for comparative purposes only.

```
NEW bidirectional eNOE
C8_H4';G10_H8  5.77 0.577

New gn-eNOE, used as lower-bound distances 
 C8_H5;G10_H1'  5.8 0.58 
 C8_H5;G10_H8 5.87 0.587  
 C8_H2';G10_H8 7.06 0.706
 C8_H3';G9_H8 5.78 0.578 
 C8_H2';G9_H8 5.92 0.592 
 C8_H1';G10_H4' 5.28 0.528 
 C8_H2';G9_H2' 4.62 0.462
```

In [5]:

```
import mdtraj as md

top = "data/PDB/2koc_gmx.pdb"
traj = "data/traj_temp_f_0.xtc"
labels,vals = read_exp("data/set_A1/eNOE.exp.dat")

#######################
md_trj = md.load(traj,top=top)
pairs = get_idxs(labels,md_trj.topology)
# convert to Angstrom
dists_md = 10*md.compute_distances(md_trj,pairs)
# write to file 
write(dists_md,"data/set_A1/eNOE.calc.dat")

pdbs = ["2koc","6by5"]
dists_pdbs = []
for  p in pdbs:
    top = "data/PDB/%s.pdb" % p 
    traj = "data/PDB/%s.pdb" % p 
    md_trj = md.load(traj,top=top)
    pairs = get_idxs(labels,md_trj.topology)
    dists = 10.0*md.compute_distances(md_trj,pairs)
    write(dists,"data/set_A1/%s.eNOE.calc.dat" %p)
```

```
# Found 63 pairs out of 63
# Found 63 pairs out of 63
# Found 63 pairs out of 63
```

In [6]:

```
labels,vals = read_exp("data/set_A1/eNOE_unidir.exp.dat")


top = "data/PDB/2koc_gmx.pdb"
traj = "data/traj_temp_f_0.xtc"
#######################
md_trj = md.load(traj,top=top)
pairs = get_idxs(labels,md_trj.topology)
# convert to Angstrom
dists_md = 10*md.compute_distances(md_trj,pairs)
# write to file 
write(dists_md,"data/set_A1/eNOE_unidir.calc.dat")

pdbs = ["2koc","6by5"]
dists_pdbs = []
for  p in pdbs:
    top = "data/PDB/%s.pdb" % p 
    traj = "data/PDB/%s.pdb" % p 
    md_trj = md.load(traj,top=top)
    pairs = get_idxs(labels,md_trj.topology)
    dists = 10.0*md.compute_distances(md_trj,pairs)
    write(dists,"data/set_A1/%s.eNOE_unidir.calc.dat" %p)
```

```
# Found 177 pairs out of 177
# Found 177 pairs out of 177
# Found 177 pairs out of 177
```

In [7]:

```
labels,vals = read_exp("data/set_A1/gn_eNOE.exp.dat")
top = "data/PDB/2koc_gmx.pdb"
traj = "data/traj_temp_f_0.xtc"
#######################
md_trj = md.load(traj,top=top)
pairs = get_idxs(labels,md_trj.topology)
# convert to Angstrom
dists_md = 10*md.compute_distances(md_trj,pairs)
# write to file 
write(dists_md,"data/set_A1/gn_eNOE.calc.dat")

pdbs = ["2koc","6by5"]
dists_pdbs = []
for  p in pdbs:
    top = "data/PDB/%s.pdb" % p 
    traj = "data/PDB/%s.pdb" % p 
    md_trj = md.load(traj,top=top)
    pairs = get_idxs(labels,md_trj.topology)
    dists = 10.0*md.compute_distances(md_trj,pairs)
    write(dists,"data/set_A1/%s.gn_eNOE.calc.dat" %p)
```

```
# Found 84 pairs out of 84
# Found 84 pairs out of 84
# Found 84 pairs out of 84
```

### Dataset B¶

Data taken from the study “High-Resolution NMR Structure of a RNA model system: the 14mer cUUCGg tetraloop hairpin RNA“ and corresponding PDB structure 2KOC.

#### NOE¶

We sourced the 251 unambigous NOE data from the .mr restraint file deposited on the PDB. Experimental error was calculated as $\sigma = (rmax-rmin)/2$

In [8]:

```
labels,vals = read_exp("data/set_B/NOE.exp.dat")

#######################
top = "data/PDB/2koc_gmx.pdb"
traj = "data/traj_temp_f_0.xtc"
md_trj = md.load(traj,top=top)
pairs = get_idxs(labels,md_trj.topology)
# convert to Angstrom
dists_md = 10*md.compute_distances(md_trj,pairs)
# write to file 
write(dists_md,"data/set_B/NOE.calc.dat")

pdbs = ["2koc","6by5"]
dists_pdbs = []
for  p in pdbs:
    top = "data/PDB/%s.pdb" % p 
    traj = "data/PDB/%s.pdb" % p 
    md_trj = md.load(traj,top=top)
    pairs = get_idxs(labels,md_trj.topology)
    dists = 10.0*md.compute_distances(md_trj,pairs)
    write(dists,"data/set_B/%s.NOE.calc.dat" %p)
```

```
# Found 251 pairs out of 251
# Found 251 pairs out of 251
# Found 251 pairs out of 251
```

### Scalar couplings¶

Available 3J couplings relative to angles

- 1H5-P, 2H5-P, C4-P (angle $\beta$). Data for residue 1 are omitted because P is not present at the 5' end in simulations.
- H3-P, C4-P(+1) (angle $\epsilon$). Data for C2-P are not used because I could not find valid Karplus relationships.
- 2H5H4, 1H5H4 (angle $\gamma$).Data for C4-1H5/2H5 are not used because I could not find valid Karplus relationships.
- H1'-H2', H2'-H3', H3'-H4' for the sugar.
  The error was taken 1.5HZ for all data. In total, there are 96 scalar couplings. For calculating the scalar coupling from structure we used the Karplus relationship as defined in baRNAba.

In [9]:

```
# read experimental data
import barnaba as bb
from barnaba import definitions
jj = [l for l in (definitions.couplings_idx)]
exp = []
dd = []
for line in open("data/set_B/J3.exp.dat"):
    if("#" not in line):
        a = line.split()[0].split("-")[0]
        try: a = int(a)
        except: a = int(a[1:])
        b = line.split()[0].split("-")[1]
        dd.append("%d-%s" % (a,b))
        
        exp.append([float(x) for x in line.split()[1:]])
exp = np.array(exp)

jjr = []
for  c in range(14):
    for j in jj:
        jjr.append("%d-%s" % (c+1,j))
idxs = [jjr.index(el) for el in dd]
```

In [10]:

```
# calculate scalar couplings for trajectory and write to file
top = "data/PDB/2koc_gmx.pdb"
native = "data/PDB/2koc_gmx.pdb"
traj = "data/traj_temp_f_0.xtc"

couplings,res = bb.jcouplings(traj,topology=top) 
data = couplings[:,:,:].reshape(-1,len(jjr))[:,idxs]
write(data,"data/set_B/J3.calc.dat")

couplings_2koc,res_2koc = bb.jcouplings("data/PDB/2koc.pdb",topology="data/PDB/2koc.pdb")
data_2koc = couplings_2koc[:,:,:].reshape(-1,len(jjr))[:,idxs]
write(data_2koc,"data/set_B/2koc.J3.calc.dat")

couplings_6by5,res_6by5 = bb.jcouplings("data/PDB/6by5.pdb",topology="data/PDB/6by5.pdb")
data_6by5 = couplings_6by5[:,:,:].reshape(-1,len(jjr))[:,idxs]
write(data_6by5,"data/set_B/6by5.J3.calc.dat")
```

```
# Loading data/traj_temp_f_0.xtc 
# Loading data/PDB/2koc.pdb 
# Loading data/PDB/6by5.pdb
```

#### RDC¶

32 RDC were taken from the .mr restraint file deposited on the PDB.
For each samples, RDC are calculated using the software pales

```
`pales -pdb sample.pdb -inD exp/rdc_set_c_bme.dat -outD outfile -H -pf1 -wv 0.022`
```

The ensemble average is then rescaled globally by a factor

$L = \sum\_i \text{exp}\_i \text{avg}\_i/\sum\_i \text{avg}\_i \text{avg}\_i$

Where the sum runs over the experimental data and $\text{avg}\_i = \sum\_j w\_j F(x\_j)$
is the pales prediction $F(x)$ calculated by averaging over the configurations $x$.
The error sigma was set to 1Hz.

### Dataset C¶

RDC taken from "Simultaneous NMR characterisation of multiple minima in the free energy landscape of an RNA UUCG tetraloop" are devided in two sub-datasets, RDC\_1 ( 39 datapoints) and RDC\_2 (14 datapoints). Back-calculation was performed as in set B. The experimental error was set to 1.4Hz (RDC\_1) and 2.2 Hz (RDC\_2) as described in the original paper.

### Dataset D¶

Solvent PRE data were obtained from the authors of the study "RNA structure refinement using NMR solvent accessibility data. Sci Rep. 2017; 7: 5393." The back-calculation of experimental measure from structure was performed using the program from Chun Tang's lab, available here using a probe radius of 0.35 nm.

In [ ]:
