## Supplementary material for "Integrating NMR and Simulations Reveals Motions in the UUCG Tetraloop": SI1 (from another version of this preprint: version 2, 2019-09-11)

SI1\_MD


### Simulation setup¶

We have performed MD simulation of an RNA 14-mer with sequence GGCACUUCGGUGCC.
Molecular dynamics simulations were performed using the GROMACS 2016.4 software package in
combination with PLUMED 2.4. Simulations wer intialized in ideal A-form, fully stacked conformations generated using the Make-NA web server . The system was solvated in a truncated dodecahedral box with 6997 TIP4P-D water molecules and neutralized by adding K+ counterions. RNA was modeled using the DESRES force-field,
available here.
The initial conformations were minimized in vacuum first, followed by a minimization in water and equilibration in NPT ensemble at 280K and 1 bar for 1 ns. Production runs were performed in the canonical ensemble using stochastic velocity rescaling thermostat. All bonds were constrained using the LINCS algorithm,and equations of motion were integrated with a 2fs time-step. To enhance sampling we have used well-tempered Metadynamics (WT-METAD) in combination with solute tempering in its REST2 formulation. 20 geometrically-distributed replicas spanning the range $\lambda$=0.37-1 (corresponding to an effective temperature range 280K-760K) were simulated for 1.0 µs per replica. The eRMSD (with cutoff=3.2) from the native reference structure (PDB 2KOC) was used as a biasing collective variable. A Gaussian bias of width 0.1 was deposited every 500 steps, and the initial height of 0.46 kJ/mol was decreased with a biasfactor 15.

### Convergence¶

We first consider the time series/distribution of the distance from the reference structure (PDB 2KOC, model 1).
RMSD and eRMSD can be calculated as described in the notebook calculate\_observables. We perform the calculation in a different notebook and save the data in a pickle file, since the same quantities will be used several times throughout the analysis. We split the simulation in four blocks of 250ns each, and observe the histograms calculated over these blocks.

In [1]:

```
import pickle
import numpy as np
import matplotlib.pyplot as plt

# plot RMSD from native
fname = "data/observables/rmsd_full.p"
rmsd_full = pickle.load(open(fname,'rb'), encoding='bytes')

plt.figure(figsize=(8,6))
plt.plot(np.arange(rmsd_full.shape[0])/100.,rmsd_full,lw=0.2)
plt.ylabel("RMSD from native [nm]")
plt.xlabel("time [ns]")
plt.show()

plt.figure(figsize=(8,6))
bins = plt.hist(rmsd_full,bins=80,density=True,alpha=0.5,label="Total")
nblocks=4
blocksize=int(rmsd_full.shape[0]/nblocks)
for j in range(nblocks):
    start=j*(blocksize)
    end= start+blocksize
    hh, ee = np.histogram(rmsd_full[start:end],bins=bins[1],density=True)
    plt.plot(0.5*(ee[1:]+ee[:-1]),hh,label="block %d" % (j+1))
plt.legend()
plt.xlabel("RMSD from native [nm]")
```

```
<Figure size 800x600 with 1 Axes>
```

Out[1]:

```
Text(0.5, 0, 'RMSD from native [nm]')
```

In [2]:

```
# plot ermsd from native
fname = "data/observables/ermsd_full.p"
ermsd_full = pickle.load(open(fname,'rb'), encoding='bytes')

plt.figure(figsize=(8,6))
plt.plot(np.arange(ermsd_full.shape[0])/100.,ermsd_full,lw=0.2)
plt.ylabel("ermsd from native [nm]")
plt.xlabel("time [ns]")
plt.show()

plt.figure(figsize=(8,6))
bins = plt.hist(ermsd_full,bins=80,density=True,alpha=0.25,label="Total")
nblocks=4
blocksize=int(ermsd_full.shape[0]/nblocks)
for j in range(nblocks):
    start=j*(blocksize)
    end= start+blocksize
    hh, ee = np.histogram(ermsd_full[start:end],bins=bins[1],density=True)
    plt.plot(0.5*(ee[1:]+ee[:-1]),hh,label="block %d" % (j+1))
plt.legend()
plt.xlabel("ermsd from native [nm]")
```

Out[2]:

```
Text(0.5, 0, 'ermsd from native [nm]')
```

Two items are worth highlighting: i) we deliberately consider the biased distribution to check convergence and ii) since eRMSD is more accurate than RMSD in distinguishing different conformations, we observe that the eRMSD distributions in the four blocks are more different one from the other compared to the RMSD.

### Assess diffusion of replicas in temperature space¶

When performing parallel-tempering schemes it is important to check that replicas diffuse in temperature space. We monitor this by analyzing the log file produced by gromacs using the perl script demux.pl.
The file "replica\_temp.xvg" contains 21 columns, each one indicating the temperature index.

In [3]:

```
#import os
#cmd = "echo 0.002 | data/script/demux.pl data/mdrun.log"
#os.system(cmd)
#os.system("mv *.xvg data")
```

For example, we can check what happens for the replica 0

In [4]:

```
import numpy as np
import matplotlib.pyplot as plt

replica_temps = np.array([[float(x) for x in line.split()] for line in open("data/replica_temp.xvg")])

# replica 1 
plt.title("Replica 0")
plt.plot(replica_temps[:,0]/1000,replica_temps[:,1])
plt.xlabel("time (ns)")
plt.yticks(range(20))
plt.ylabel("replica temperature")
plt.ylim((0,20))
plt.show()
```

More generally, we calculate the number of round trips in temperature space. A complete round trip means that goes from T=T\_0 to T=T\_19 and back to T\_0

In [5]:

```
nrep = replica_temps.shape[1]-1
nframes = replica_temps.shape[0]
states = np.zeros((nframes,nrep))
up = np.zeros(nrep)
down = np.zeros(nrep)
for j in range(nframes):
    for k in range(1,nrep+1):
        if(int(replica_temps[j,k])==0): 
            states[j,k-1] = -1
            if(states[j-1,k-1]==1): down[k-1] += 1 
        elif(int(replica_temps[j,k])==(replica_temps.shape[1]-2)):
            states[j,k-1] = 1
            if(states[j-1,k-1]==-1): up[k-1] += 1 
        else:
            states[j,k-1] = states[j-1,k-1]
```

In [6]:

```
# And do a barplot
plt.bar(np.arange(nrep),down)
plt.xlabel("replica index")
plt.ylabel("# of round trips in temperature space")
plt.xticks(np.arange(nrep))
plt.savefig("figures/round_trips.png",dpi=500)
plt.show()
```

We can see that replica 5 goes to the highest temperature and comes back to T\_0 only once

In [7]:

```
# replica 1 
plt.title("Replica 5")
plt.plot(replica_temps[:,0]/1000,replica_temps[:,6])
plt.xlabel("time (ns)")
plt.yticks(range(20))
plt.ylabel("replica temperature")
plt.ylim((0,20))
plt.show()
```
