## Supplementary material for "Integrating NMR and Simulations Reveals Motions in the UUCG Tetraloop": SI0 (from another version of this preprint: version 2, 2019-09-11)

00\_calculate\_observables


### Calculate quantities from trajectory¶

In this notebook we calculate RMSD/eRMSD from 2KOC, backbone angles and distances between 6-membered rings. These quantities are useful throughout the other notebooks. Therefore, we calculate them only once.

In [1]:

```
import barnaba as bb
import pickle
import os
# native structure
native = "data/PDB/2koc_gmx.pdb"
# topology 
top = "data/PDB/2koc_gmx.pdb"
# trajectory file
trj = "data/traj_temp_f_0.xtc"

# 2KOC and 6BY5 files from PDB database 
top1 = "data/PDB/2koc.pdb"
top2 = "data/PDB/6by5.pdb"

# makedir
%mkdir -p data/observables
```

In [2]:

```
# calculate ermsd from 2KOC, full structure 
fname = "data/observables/ermsd_full_2koc.p"
if os.path.isfile(fname):
    ermsd_full_2koc = pickle.load(open(fname,'rb'), encoding='bytes')
else:
    ermsd_full_2koc = bb.ermsd(native,top1,topology=top1)
    pickle.dump(ermsd_full_2koc,open(fname, 'wb'))

fname = "data/observables/ermsd_full_6by5.p"
if os.path.isfile(fname):
    ermsd_full_6by5 = pickle.load(open(fname,'rb'), encoding='bytes')
else:
    ermsd_full_6by5 = bb.ermsd(native,top2,topology=top2)
    pickle.dump(ermsd_full_6by5,open(fname, 'wb'))
    
fname = "data/observables/ermsd_full.p"
if os.path.isfile(fname):
    ermsd_full = pickle.load(open(fname,'rb'), encoding='bytes')
else:
    ermsd_full = bb.ermsd(native,trj,topology=top)
    pickle.dump(ermsd_full,open(fname, 'wb'))
```

In [3]:

```
# calculate rmsd from 2KOC, full structure 
fname = "data/observables/rmsd_full_2koc.p"
if os.path.isfile(fname):
    rmsd_full_2koc = pickle.load(open(fname,'rb'), encoding='bytes')
else:
    rmsd_full_2koc = bb.rmsd(native,top1,topology=top1,heavy_atom=True)
    pickle.dump(rmsd_full_2koc,open(fname, 'wb'))

fname = "data/observables/rmsd_full_6by5.p"
if os.path.isfile(fname):
    rmsd_full_6by5 = pickle.load(open(fname,'rb'), encoding='bytes')
else:
    rmsd_full_6by5 = bb.rmsd(native,top2,topology=top2,heavy_atom=True)
    pickle.dump(rmsd_full_6by5,open(fname, 'wb'))
    
fname = "data/observables/rmsd_full.p"
if os.path.isfile(fname):
    rmsd_full = pickle.load(open(fname,'rb'), encoding='bytes')
else:
    rmsd_full = bb.rmsd(native,trj,topology=top,heavy_atom=True)
    pickle.dump(rmsd_full,open(fname, 'wb'))
```

In [4]:

```
# calculate ermsd from 2KOC, stem only  
fname = "data/observables/ermsd_stem_2koc.p"
if os.path.isfile(fname):
    ermsd_stem_2koc = pickle.load(open(fname,'rb'), encoding='bytes')
else:
    ermsd_stem_2koc = bb.ermsd(native,top1,topology=top1,residues_ref=[0,1,2,3,4,9,10,11,12,13],residues_target=[0,1,2,3,4,9,10,11,12,13])
    pickle.dump(ermsd_stem_2koc,open(fname, 'wb'))

fname = "data/observables/ermsd_stem_6by5.p"
if os.path.isfile(fname):
    ermsd_stem_6by5 = pickle.load(open(fname,'rb'), encoding='bytes')
else:
    ermsd_stem_6by5 = bb.ermsd(native,top2,topology=top2,residues_ref=[0,1,2,3,4,9,10,11,12,13],residues_target=[0,1,2,3,4,9,10,11,12,13])
    pickle.dump(ermsd_stem_6by5,open(fname, 'wb'))
    
fname = "data/observables/ermsd_stem.p"
if os.path.isfile(fname):
    ermsd_stem = pickle.load(open(fname,'rb'), encoding='bytes')
else:
    ermsd_stem = bb.ermsd(native,trj,topology=top,residues_ref=[0,1,2,3,4,9,10,11,12,13],residues_target=[0,1,2,3,4,9,10,11,12,13])
    pickle.dump(ermsd_stem,open(fname, 'wb'))
```

In [5]:

```
# calculate ermsd from 2KOC, loop only  

fname = "data/observables/ermsd_loop_2koc.p"
if os.path.isfile(fname):
    ermsd_loop_2koc = pickle.load(open(fname,'rb'), encoding='bytes')
else:
    ermsd_loop_2koc = bb.ermsd(native,top1,topology=top1,residues_ref=[4,5,6,7,8,9],residues_target=[4,5,6,7,8,9])
    pickle.dump(ermsd_loop_2koc,open(fname, 'wb'))

fname = "data/observables/ermsd_loop_6by5.p"
if os.path.isfile(fname):
    ermsd_loop_6by5 = pickle.load(open(fname,'rb'), encoding='bytes')
else:
    ermsd_loop_6by5 = bb.ermsd(native,top2,topology=top2,residues_ref=[4,5,6,7,8,9],residues_target=[4,5,6,7,8,9])
    pickle.dump(ermsd_loop_6by5,open(fname, 'wb'))
    
fname = "data/observables/ermsd_loop.p"
if os.path.isfile(fname):
    ermsd_loop = pickle.load(open(fname,'rb'), encoding='bytes')
else:
    ermsd_loop = bb.ermsd(native,trj,topology=top,residues_ref=[4,5,6,7,8,9],residues_target=[4,5,6,7,8,9])
    pickle.dump(ermsd_loop,open(fname, 'wb'))
```

In [6]:

```
# calculate backbone angles  

fname = "data/observables/angles_2koc.p"
if os.path.isfile(fname):
    [angles_2koc,res_2koc] = pickle.load(open(fname,'rb'), encoding='bytes')
else:
    [angles_2koc,res_2koc] = bb.backbone_angles(top1,topology=top1)
    pickle.dump( [angles_2koc,res_2koc] ,open(fname, 'wb'))

fname = "data/observables/angles_6by5.p"
if os.path.isfile(fname):
    [angles_6by5,res_6by5] = pickle.load(open(fname,'rb'), encoding='bytes')
else:
    [angles_6by5,res_6by5] = bb.backbone_angles(top2,topology=top2)
    pickle.dump( [angles_6by5,res_6by5] ,open(fname, 'wb'))

fname = "data/observables/angles.p"
if os.path.isfile(fname):
    [angles,res] = pickle.load(open(fname,'rb'), encoding='bytes')
else:
    [angles,res] = bb.backbone_angles(trj,topology=top)
    pickle.dump( [angles,res] ,open(fname, 'wb'))
```

In [7]:

```
# calculate vectors connecting the centers of the six-mbemberd rings
fname = "data/observables/rvecs_2koc.p"
if os.path.isfile(fname):
    [rvecs_2koc,res_2koc] = pickle.load(open(fname,'rb'), encoding='bytes')
else:
    [rvecs_2koc,res_2koc] = bb.dump_rvec(top1,topology=top1,cutoff=100)
    pickle.dump( [rvecs_2koc,res_2koc] ,open(fname, 'wb'))

fname = "data/observables/rvecs_6by5.p"
if os.path.isfile(fname):
    [rvecs_6by5,res_6by5] = pickle.load(open(fname,'rb'), encoding='bytes')
else:
    [rvecs_6by5,res_6by5] = bb.dump_rvec(top2,topology=top2,cutoff=100)
    pickle.dump( [rvecs_6by5,res_6by5] ,open(fname, 'wb'))

fname = "data/observables/rvecs.p"
if os.path.isfile(fname):
    [rvecs,res] = pickle.load(open(fname,'rb'), encoding='bytes')
else:
    [rvecs,res] = bb.dump_rvec(trj,topology=top,cutoff=100)
    pickle.dump( [rvecs,res] ,open(fname, 'wb'))
```
